# Dietary restriction transforms the protein sulfhydrome in a tissue-specific and cystathionine γ-lyase-dependent manner

**DOI:** 10.1101/869271

**Authors:** Nazmin Bithi, Christopher Link, Rui Wang, Belinda Willard, Christopher Hine

## Abstract

Hydrogen sulfide (H_2_S) is a cytoprotective redox-active metabolite that signals through protein sulfhydration (R-SS_n_H). Despite the known importance of sulfhydration on relatively few identified proteins, tissue-specific sulfhydrome profiles and their associated functions are not well characterized, specifically under conditions known to modulate H_2_S production. We hypothesized that dietary restriction (DR), which increases lifespan and boosts endogenous H_2_S production, expands functional tissue-specific sulfhydromes. Here, we found that 50% DR enriched total sulfhydrated proteins in liver, kidney, muscle, and brain but decreased these in heart of adult male mice. DR promoted sulfhydration in numerous metabolic and aging-related pathways. Mice lacking the H_2_S producing enzyme cystathionine γ-lyase (CGL) had decreased liver and kidney protein sulfhydration and failed to functionally augment their sulfhydrome in response to DR. Overall, we defined tissue- and CGL-dependent sulfhydromes and how diet transforms their makeup, underscoring the breadth for DR and H_2_S to impact biological processes and organismal health.

**One Sentence Summary:** Dietary restriction altered the tissue-specific enrichment of sulfhydrated proteins and their downstream signaling pathways in liver, kidney, skeletal muscle, brain, heart, and plasma that was partly dependent on the hydrogen sulfide producing enzyme cystathionine γ-lyase.

## Introduction

The search for anti-aging interventions predates the modern medical and scientific era. Currently, the best studied and effective interventions and genetic models to extend both lifespan and healthspan across evolutionary boundaries are dietary restriction (DR) (Fontana and Partridge, 2015) and disruption of the growth hormone (GH)/insulin-like growth factor 1 (IGF-1) axis (Junnila et al., 2013). Largely defined as reduced nutrient intake without malnutrition, DR encompasses decreased total daily caloric intake, the removal of specific macronutrients such as amino acids, and/or intermittent fasting. In addition to defending against aging-related diseases (Mattison et al., 2017; Sinclair, 2005; Weindruch and Walford, 1982), DR also provides stress resistance (Mitchell et al., 2010; Peng et al., 2012) and metabolic fitness (Fabbiano et al., 2016; Plaisance et al., 2011). Emerging DR studies in humans indicate an average daily caloric reduction of approximately 12% over two years resulted in favorable improvements in metabolic rate, endocrine activity, and redox status (Redman et al., 2018). Similar to DR, mice lacking adequate GH production (Flurkey et al., 2001) or GH receptor (GHRKO) signaling (Bonkowski et al., 2006) have increased lifespans, resistance to a number of stressors, and metabolic fitness (Brown-Borg, 2015). Interestingly, GHR deficiency in humans reduces pro-aging signaling, cancer, and diabetes (Guevara-Aguirre et al., 2011). It is hypothesized that common molecular pathways enriched or affected by DR and GH/IGF-1 axis disruption are central to improved aging (Li et al., 2014). Identifying these shared pathways and elucidating their mechanism of action will shed light onto the underpinnings of aging and usher more targeted diagnostics and therapies into the clinic.

A common molecular phenomenon amongst several dietary (Hine et al., 2015; Nakano et al., 2015; Yoshida et al., 2018), pharmaceutical (Wiliński et al., 2013), and genetic (Hine et al., 2017; Wei and Kenyon, 2016) models of longevity is the increased production capacity and/or bioavailability of hydrogen sulfide (H_2_S) gas (Hine et al., 2018). H_2_S and its HS^−^ anion and S^2−^ ion, herein referred simply as H_2_S, is historically classified as an environmental and occupational hazard due to its inhibition of mitochondrial respiration (Szabo et al., 2013; Wu et al., 2011) and lethality at elevated levels (Hendrickson et al., 2004). However, in the later part of the 20^th^ and early part of the 21^st^ Centuries, functional and beneficial endogenous production of H_2_S via enzymatic catalysis of sulfur containing amino acids (Stipanuk and Beck, 1982) was first discovered in the brain, aorta, and vascular smooth muscle cells (Abe and Kimura, 1996; Hosoki et al., 1997; Zhao et al., 2001). Exogenously provided H_2_S, either in gas form or via donor molecules, increases lifespan in model organisms (Miller and Roth, 2007), prevents multi-tissue ischemic-reperfusion injury (Elrod et al., 2007; Hine et al., 2015), improves cardiovascular health (Das et al., 2018; Yang et al., 2008), and is recognized as a potential therapeutic agent against aging-associated diseases (Zhang et al., 2013). Conversely, deficiencies in endogenous H_2_S levels or production correlate with and/or are causative of hypertension (Yang et al., 2008), colitis (Flannigan et al., 2014), neurodegeneration (Sbodio et al., 2016), and glioma growth (Takano et al., 2014) while H_2_S levels in humans are reported to decline during aging, particularly those suffering from COPD (Chen et al., 2005; Perridon et al., 2016). Thus, the field of H_2_S in biology, physiology, and medicine has blossomed (Wallace and Wang, 2015; Wang, 2012), with endogenous H_2_S serving as a therapeutic target and the third functional gasotransmitter alongside carbon monoxide and nitric oxide (Paul and Snyder, 2018).

Due to the multiphasic dose response of H_2_S, life evolved with several mechanisms for its controlled production and utilization. In mammals, H_2_S is enzymatically produced during the metabolism of sulfur containing amino acids, primarily cysteine and homocysteine, via the activities of cystathionine β-synthase (CBS), cystathionine γ-lyase (CGL), and 3-mercaptopyruvate sulfurtransferase (3-MST) (Kabil and Banerjee, 2014). CBS and CGL, both central to the transsulfuration pathway, require pyridoxal phosphate (PLP) as co-factor for α,β-elimination or β-replacement of the thiol group on cysteine or homocysteine to produce H_2_S (Singh and Banerjee, 2011). 3-MST produces H_2_S via sulfane sulfur and 3-mercaptopyruvate, which is synthesized by the conversion of cysteine to 3-mercaptopyruvate via cysteine aminotransferase (CAT) (Shibuya et al., 2009). CGL and CBS are predominantly cytoplasmic, while 3-MST is principally mitochondrial (Kimura, 2014). In addition to their differences in subcellular localization, CGL, CBS, and 3-MST are differentially expressed and active in tissue-specific manners, with CGL contributing the majority of enzymatic H_2_S production in the kidney, liver, and endothelium (Kabil et al., 2011b; Longchamp et al., 2018; Yang et al., 2019). Lack of CGL expression and/or activity is attributed to hypertension (Yang et al., 2008), neurodegeneration (Paul et al., 2014), and the inability to properly respond to dietary and endocrine cues (Hine et al., 2015; Hine et al., 2017; Ishii et al., 2010; Kabil et al., 2011a; Longchamp et al., 2018; Nakano et al., 2015). Remarkably, it was recently found that increased CGL expression in liver is a hallmark of numerous dietary, genetic, hormonal, and pharmaceutical mouse models of extended lifespan and may serve as a molecular biomarker associated with longevity (Tyshkovskiy et al., 2019).

The downstream mechanisms as to how enhanced CGL expression and subsequent endogenous H_2_S production are utilized and impart cellular, tissue, and systemic benefits rests on the versatility of sulfur to have oxidation states from −2 to +6 and accept or donate electrons (Filipovic et al., 2018). With the sulfur on H_2_S having an oxidation state of −2, indicative of a reducing agent, it results in H_2_S performing several non-mutually exclusive biochemical reactions. These include mitochondrial electron transfer, alterations of iron-sulfur and heme centers, antioxidant activity, and protein posttranslational modification via sulfhydration; aka persulfidation or s-sulfhydration, of reactive cysteine residues to form persulfide tails of various lengths (R-SS_n_H) (Filipovic, 2015). While the benefits of a readily diffusible antioxidant to counter oxidative damage (Harman, 2009) and improvement in mitochondrial bioenergetics to delay the onset of aging-related disorders (Bratic and Larsson, 2013) are easily identifiable, the act of sulfhydration and its extent to initiate improved fitness, lifespan, and healthspan is less well understood.

Like other protein posttranslational modifications, sulfhydration of readily active cysteine residues potentially alters a protein’s structure, function, stability, and/or macromolecular interactions. (Filipovic et al., 2018; Kimura, 2019; Mustafa et al., 2009). The sulfhydration process generally involves: 1) oxidation of the cysteine residue(s) on the targeted protein, as H_2_S cannot solely react with a reduced thiol group, followed by nucleophilic attack of the oxidized thiol and/or disulfide by H_2_S, or 2) reaction between H_2_S derived polysulfides (H_2_S_n_) with a reduced protein thiol (Gao et al., 2015; Kimura, 2019). It is estimated that 10 to 25% of proteins in the rodent liver, a strong generator of H_2_S in the body (Kabil et al., 2011b), are sulfhydrated (Mustafa et al., 2009) and this modification typically increases the reactivity of the modified cysteine (Paul and Snyder, 2012). Examples of metabolic, stress resistance, and aging-related proteins identified to have functional sulfhydration modifications include, but are not limited to, glyceraldehyde phosphate dehydrogenase (GAPDH) at Cys^150^ (Gao et al., 2015; Mustafa et al., 2009), nuclear factor κB (NF-κB) at Cys^38^ (Sen et al., 2012), the Kir 6.1 subunit of endothelial ATP-sensitive potassium channel at C^43^ (Mustafa et al., 2011), sirtuin 1 (SIRT1) at Cys^371^, Cys^374^, Cys^395^, and Cys^398^ (Du et al., 2019), and the E3 ubiquitin ligase Parkin at Cys95 (Vandiver et al., 2013). The majority of studies identifying and examining the biochemical functions of sulfhydration are primarily constructed on cell culture models and/or with purified proteins *in vitro*. One of the few studies to examine the total number and identity of sulfhydrated proteins, or sulfhydrome, of an organism *in vivo* was completed in the plant *Arabidopsis thaliana*, with the discovery of 2,015 proteins, or approximately 5% of the entire proteome, as sulfhydrated (Aroca et al., 2017). However, the sulfhydrome profiles and their associated biological functions of major organs in a mammal, particularly under dietary and/or genetic conditions known to impact endogenous H_2_S production, have not been described.

Utilizing the relatively new approach for the identification of sulfhydrated proteins developed by Gao, *et al.*, termed the Biotin-Thiol-Assay (BTA) (Gao et al., 2015), we set out to test the hypothesis that interventions known to increase lifespan, improve metabolic and stress resiliency, and boost endogenous H_2_S production would essentially expand and/or alter tissue-specific sulfhydrome profiles *in vivo*. To address this hypothesis, we employed a daily 50% caloric restriction for 1-week versus *ad libitum* (AL) feeding in 6-month male CGL wild type (WT) and total body knock out (KO) mice to examine for diet-, tissue- and genotype-specific changes in H_2_S production (*i*), protein sulfhydration (*ii*), and biological pathways impacted by protein sulfhydration (*iii*) (Figure 1A). This type and duration of dietary intervention was chosen as it was previously shown to induce functional CGL-dependent hepatic H_2_S production in WT mice (Hine et al., 2015). Here, we identified 977, 1086, 459, 431, 884 and 160 total sulfhydrated proteins and their associated biological pathways in the liver, kidney, heart, muscle (quadriceps), brain and plasma, respectively. DR enriched the number of sulfhydrated proteins in liver, kidney, muscle, and brain while it decreased these in the heart and had minimal impact in the plasma. DR in WT mice promoted sulfhydration of proteins involved in numerous pathways, notably metabolic and protein/nitrogen homeostasis. CGL KO mice had an overall decrease in sulfhydrated proteins in the liver and kidney and failed to appropriately augment their sulfhydrome in response to DR. These findings suggest DR plays an important role in H_2_S production and transformation of the protein sulfhydrome to ultimately promote stress resistance, metabolic fitness, neuroprotection, and longevity.

**Figure 1:**
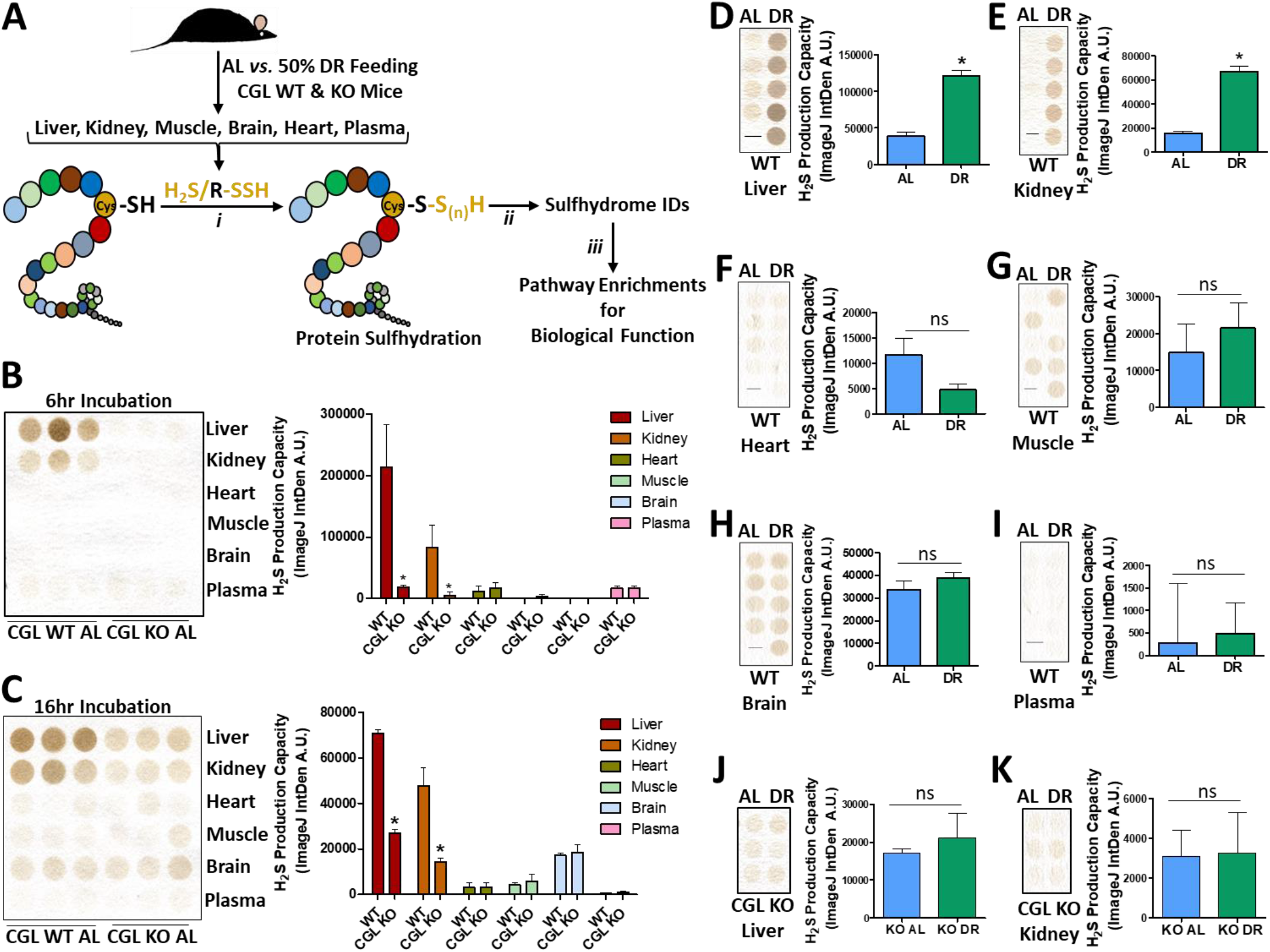
Diet and CGL status impact H_2_S production in a tissue specific manner. (**A**) Graphical presentation of the overarching experimental setup. 6-month old male cystathionine γ-lyase (CGL) WT and total body KO mice were placed on *ad libitum* (AL) or 50% dietary restriction (DR) diets for 1 week prior to tissue harvest. Tissues were analyzed for (*i*) H_2_S production capacity via the lead acetate/lead sulfide method, (*ii*) protein sulfhydration profiles via the biotin thiol (BTA) assay, and (iii) biological pathway enrichment/function of the identified sulfhydrated proteins. (**B-C**) H_2_S production capacity in tissues in AL fed CGL WT (n= 3 mice/group) and CGL KO (n=3 mice/group) mice at short 6 hr (**B**) and long 16 hr (**C**) exposures. The asterisk indicates the significance of the difference in the average tissue-specific H_2_S production capacity between the CGL WT and KO groups; * p < 0.05. (**D-I**) H_2_S production capacity in tissues harvested from CGL WT mice after one week of AL (n=4 mice/group) or 50% DR (n=5 mice/group) feeding. (**J-K**) H_2_S production capacity in liver (**J**) and kidney (**K**) in CGL KO mice after one week of AL (n=3 mice/group) or 50% DR (n=3 mice/group) feeding. Error bars are ± SEM. The asterisks indicate the significance of the difference between the dietary treatment groups; *p < 0.05. See also Supplemental Figure 1.

## Results

### Diet and CGL status impact tissue specific H_2_S production

To establish the impact of 1-week 50% DR on mouse physiology, H_2_S production, and sulfhydrome profiles, the daily food intake was first measured to determine the correct amount of modified AIN-93G rodent diet to restrict to achieve a 50% reduction in calories consumed. *Ad libitum* (AL) mice were then provided 24-hour access to the rodent diet and the 50% DR (DR) mice were fed their calculated allotment near the start of their dark phase at 7pm to avoid disturbances in circadian rhythms and feeding patterns between the two groups (Acosta-Rodriguez et al., 2017; Froy et al., 2009). Average daily food intake monitored over the 1-week intervention showed the AL group consumed 0.2g food/g mouse, or approximately 12 kcal/mouse, per day while the DR group consumed 0.1g food/g mouse, or approximately 6 kcal/mouse, per day (Supplemental Figure 1A, B). DR for 7 days reduced body mass similarly in both CGL WT and KO mice by 5-10% of initial weight (Supplemental Figure 1C, D). After the 1-week dietary intervention liver, kidney, heart, muscle (quadriceps), brain, and plasma were collected for downstream H_2_S production capacity analysis via the lead acetate/lead sulfide endpoint method (Hine and Mitchell, 2017) and for sulfhydration profiling. Similar to previous results obtained in our laboratory regarding the tissue specificity of CGL, CBS, and 3MST protein expression, H_2_S production, and the dependence on CGL for H_2_S production (Yang et al., 2019), and in line with unbiased ENCODE transcriptome data of H_2_S producing and consuming genes obtained from NCBI Gene Database (Yue et al., 2014) (Supplemental Figure 1E-H), we show H_2_S production capacity in AL fed mice is highest in liver and kidney, and this is dependent on CGL (Figure 1B, C). Heart, muscle, brain, and plasma showed low H_2_S production capacities with little to no dependence on CGL (Figure 1B, C). DR increased H_2_S production capacity in liver (Figure 1D) and kidney (Figure 1E), but had little to no impact in heart, muscle, brain, or plasma (Figure 1F-I). As CGL status is important for liver and kidney H_2_S production, we next focused on the impact of DR on H_2_S production in these two organs in CGL KO mice. DR failed to increase detectable H_2_S production capacity in the liver and kidney of CGL KO mice (Figure 1J, K). Thus, we confirm that H_2_S production capacity is highest and malleable via DR in liver and kidney, which are dependent on CGL, while other tissues tested have relatively low H_2_S production capacity independent of CGL status or diet.

### DR enriches the sulfhydrome in liver, kidney, muscle, and brain

To first verify the ability of the modified BTA (Gao et al., 2015) (Supplemental Figure 2A) to detect sulfhydrated proteins from tissue samples, we ran Western blot analysis on the final liver BTA elution −/+ DTT for α-Tubulin and GAPDH, two hepatic proteins previously shown to be sulfhydrated (Mustafa et al., 2009) (Supplemental Figure 2B). α-Tubulin and GAPDH were detected in the sulfhydration-specific +DTT elution (Gao et al., 2015) from WT mice but not CGL KO mice nor in the −DTT elution lanes (Supplemental Figure 2B). Pre-treatment of NaHS on WT livers *ex vivo* prior to performing the BTA assay not only failed to increase detection of these two proteins, but resulted in decreased detection. This is indicative that *ex vivo* addition of NaHS/H_2_S alone cannot sulfhydrate proteins without an oxidant to activate the cysteine residue or without formation of a polysulfide prior to modifying the cysteine residue, as noted in previous reports (Filipovic et al., 2018; Gao et al., 2015), and that *ex vivo* addition of NaHS may act as a reducing equivalent, similar to DTT, and remove the sulfhydration modification prior to the BTA assay (Supplemental Figure 2B). Thus, utilizing this modified BTA technique followed by mass spectrometry-based quantitative proteomics analysis of peptide spectral counts (Supplemental Figure 2A, Supplemental Data File 1), we set out to examine tissue-, dietary-, and CGL-dependent sulfhydrome profiles on major the tissue samples listed in Figure 1A.

**Figure 2:**
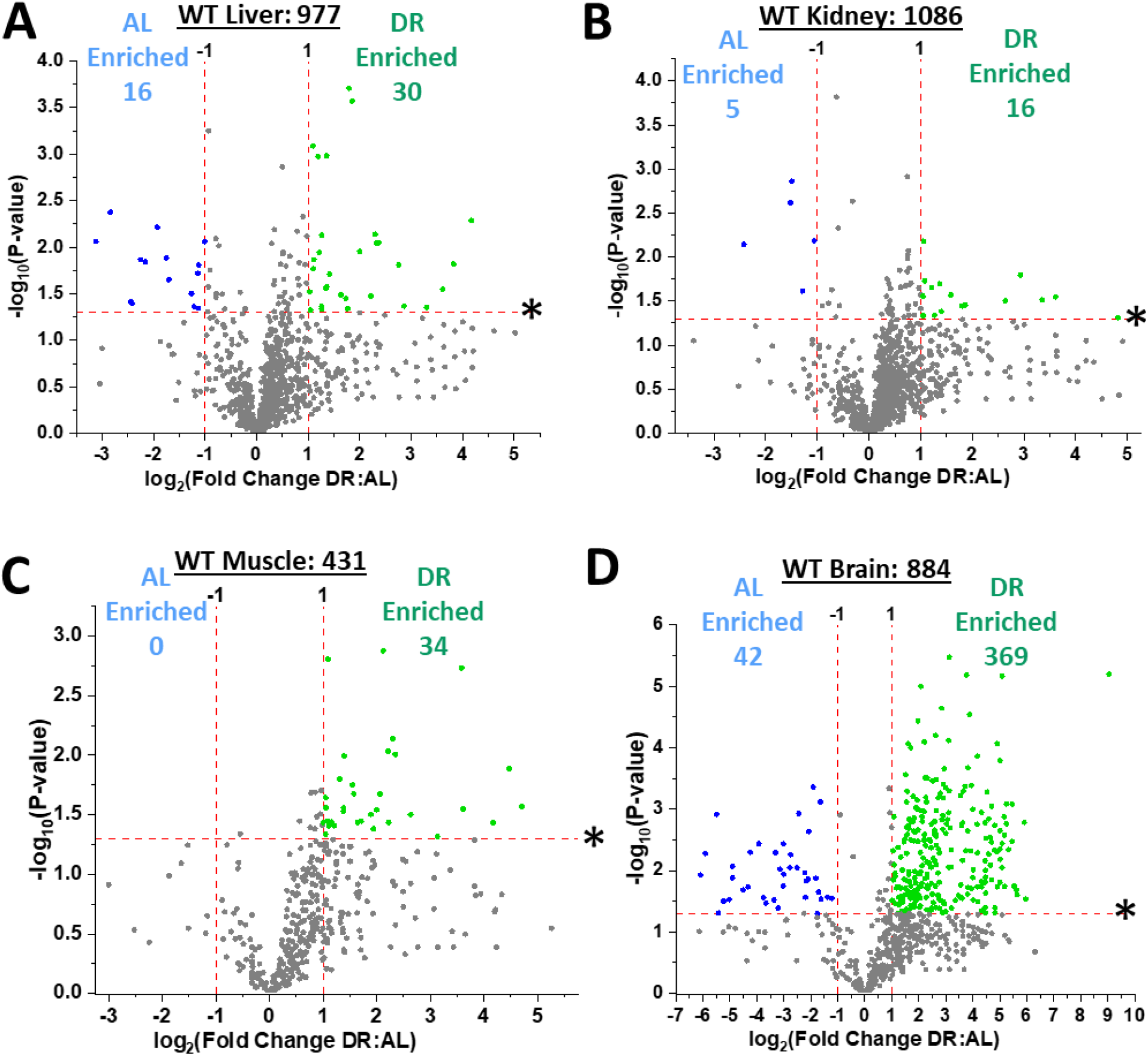
Dietary restriction enriches the sulfhydration profiles in liver, kidney, muscle, and brain. (**A-D**) Volcano plots showing biologically- and statistically-significant differentially abundant sulfhydrated proteins in liver (**A**), kidney (**B**), quadriceps muscle (**C**), and whole brain (**D**) from *ad libitum* (AL; n = 4 mice/group) versus 50% dietary restriction (DR; n = 5 mice/group) fed cystathionine γ-lyase (CGL) WT mice. The log_2_(Fold Change DR:AL) X-axis displays the average fold change in spectral counts for each identified sulfhydrated protein while the −log_10_ Y-axis displays the calculated p-value when comparing the individual spectral count values for each identified sulfhydrated protein in a specific tissue from AL versus DR fed mice. The non-axial red dotted vertical lines highlight the biological significance threshold of +/−2-fold change in spectral counts between DR versus AL, while the non-axial red dotted horizontal line with asterisk highlights our statistical significance threshold of p <0.05. The number of total sulfhydrated proteins identified in each tissue are given next to the tissue name, while blue (AL enriched) and green (DR enriched) colored dots and text indicate sulfhydrated proteins reaching both biological- and statistical- thresholds. Gray color dots indicate sulfhydrated proteins not reaching the criteria for both biological and statistical significance for enrichment under either diet. See also Supplemental Figure 2.

Of the six tissues tested in WT mice via the BTA, four were found to have enriched sulfhydrome profiles in response to DR when compared to AL feeding (Figures 2A-D, Supplementary Figures 2C-F, Supplemental Tables 1-4). These tissues included liver (Figure 2A, Supplementary Figure 2C, Supplemental Table 1), kidney (Figure 2B, Supplementary Figure 2D, Supplemental Table 2), muscle (Figure 2C, Supplementary Figure 2E, Supplemental Table 3), and brain (Figure 2D, Supplementary Figure 2F, Supplemental Table 4). Total sulfhydrated proteins identified in each tissue independent of diet included 977 in liver, 1086 in kidney, 431 in muscle, and 884 in brain. Identified proteins enriched under DR, displayed as green dots, included 30 in liver, 16 in kidney, 34 in muscle, and 369 in brain that met a biological- and statistical-significance threshold of at least a 2-fold increase in DR:AL spectral count ratio with a p-value <0.05, respectively (Figures 2A-D). Likewise, proteins enriched under AL feeding in these four tissues, displayed as blue dots, included 16 in liver, 5 in kidney, 0 in muscle, and 42 in brain that met a biological- and statistical-significance threshold of at least a 2-fold increase in AL:DR spectral count ratio with a p-value <0.05, respectively (Figures 2A-D). In addition, sulfhydrated proteins not meeting these thresholds for biological- and statistical significance, displayed as gray dots, were overall skewed in all four tissues for enrichment under DR feeding (Figures 2A-D). Thus, DR enhances the number of sulfhydrated proteins in liver, kidney, muscle, and brain. However, the concerted biological functions and pathways affected by these sulfhydration events, their commonality across tissue types, and how they are altered as a function of diet are yet to be classified.

### Numerous pathways enriched with sulfhydrated proteins dependent on diet and organ

In the four organs positively enriched for sulfhydrated proteins after 1-week 50% DR, a total of 1,854 individual proteins were identified, with 209, or 11.3%, shared amongst liver, kidney, muscle, and brain (Figure 3A, Supplemental Table 5). Biological function and pathway enrichment via g:Profiler analysis (Raudvere et al., 2019) of the shared 209 sulfhydrated proteins utilizing the KEGG database (Kanehisa et al., 2016) revealed a total of 15 pathways, with carbon metabolism, proteasome, and valine, leucine, and isoleucine degradation as the top three most significantly enriched (Figure 3B). As this previous analysis did not take into account diet-related changes in the sulfhydromes of these four organs, we next compared the proteins and pathways enriched for sulfhydration under AL and DR feeding (Figures 3C-G, Supplementary Tables 6-9). A total of 63 individual proteins were identified to be enriched for sulfhydration under AL feeding with 0 shared (Figure 3C), while a total of 429 proteins were identified to be enriched for sulfhydration under DR feeding with 0 shared (Figure 3D).

**Figure 3:**
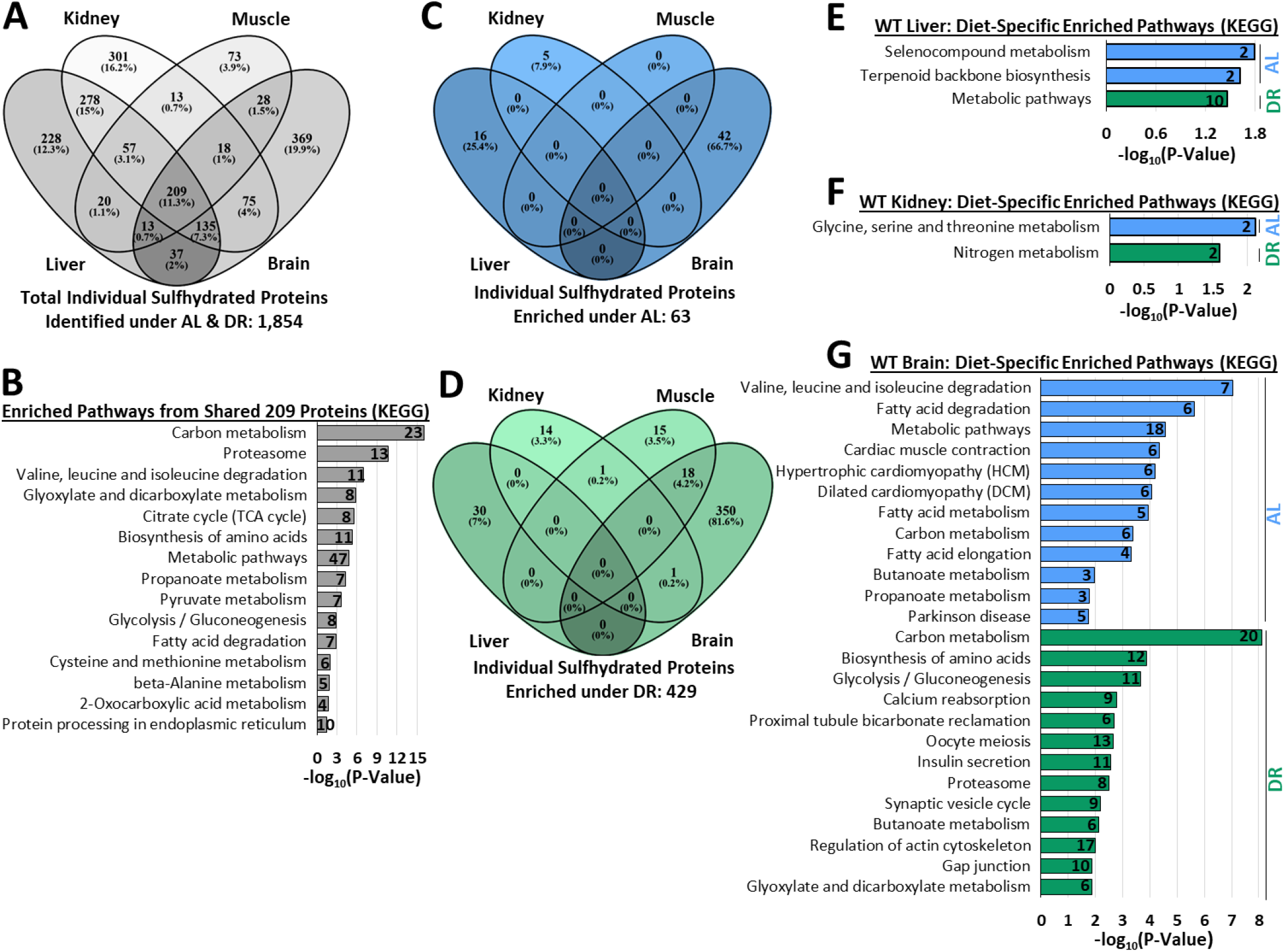
Multiple pathways enriched with sulfhydrated proteins dependent on diet and organ. (**A**) 4-way Venn diagram showing the abundance of shared and non-shared sulfhydrated proteins identified in liver, kidney, muscle, and brain in CGL WT mice. 209 sulfhydrated proteins were shared amongst all 4 organs. n = 9 mice total; 4 AL and 5 DR. (**B**) KEGG biological function and pathway enrichment of the shared 209 proteins found in liver, kidney, muscle, and brain via g:Profiler analysis. The number in the bar indicates the number of individual sulfhydrated proteins identified for that specific pathway. (**C-D**) 4-way Venn diagram showing shared and non-shared sulfhydrated proteins enriched under AL (**C**) and DR (**D**) feeding. (**E-G**) KEGG biological function and pathway enrichment of AL enriched (blue bars) or DR enriched (green bars) sulfhydrated proteins in liver (**E**), kidney (**F**), and brain (**G**). The number inside the bar indicates the quantity of sulfhydrated proteins involved in that specific pathway. Statistical significance for pathway enrichment plotted as −log_10_ (P-Value) and obtained via the g:Profiler g:SCS algorithm for KEGG database analysis. See also Supplemental Figure 3.

As no common sulfhydrated proteins were enriched as a function of diet in all four organs, we next examined tissue-specific functional enrichment analysis of sulfhydrated proteins. Liver contained 2 pathways enriched; selenocompound metabolism and terpenoid backbone biosynthesis, under AL and 1 pathway enriched; metabolic pathways, under DR feeding (Figure 3E, Supplemental Table 6). Kidney contained 1 pathway enriched; glycine, serine, and threonine metabolism, under AL and 1 pathway; nitrogen metabolism, under DR feeding (Figure 3F, Supplemental Table 7). No pathways were enriched from the diet-specific sulfhydrated proteins in muscle. Brain contained 12 pathways enriched under AL and 13 enriched under DR feeding (Figure 3G, Supplemental Table 9). The top three pathways enriched under AL feeding in the brain included valine, leucine, and isoleucine degradation, fatty acid degradation, and metabolic pathways, while the top three pathways enriched under DR feeding in the brain included carbon metabolism, biosynthesis of amino acids, and glycolysis/gluconeogenesis (Figure 3G).

Sulfhydrated proteins not significantly impacted by diet, which primarily compose the majority of the sulfhydromes in liver, kidney, muscle, and brain, still trended for enrichment under DR (Figures 2A-D; gray dots). Function and pathway enrichment of these proteins revealed 34, 41, 15, and 13 pathways in liver, kidney, muscle, and brain, respectively (Supplemental Figure 3A-D). Ten of the pathways identified were shared amongst all four tissues, and include metabolic processes, carbon metabolism, valine, leucine and isoleucine degradation, biosynthesis of amino acids, glyoxylate and dicarboxylate metabolism, cysteine and methionine metabolism, pyruvate metabolism, proteasome, citrate cycle (TCA cycle), and glycolysis/gluconeogenesis. While not shared amongst all tissues, pathways related to aging-related neurodegenerative Parkinson and Huntington diseases were enriched with sulfhydrated proteins in both kidney and muscle (Supplemental Figure 3B, C). Thus, we have identified 4 major metabolic tissues/organs that respond to dietary restriction via enriching their total number of sulfhydrated proteins, with many of the proteins falling into specific biological pathways and functions.

### DR decreases heart and negligibly impacts plasma sulfhydrome profiles

While DR enriched the sulfhydromes of liver, kidney, muscle, and brain, it failed to do so in heart and plasma (Figure 4A, B; Supplemental Figure 4A, B; and Supplemental Tables 10, 11). In heart, a total of 459 sulfhydrated proteins were identified, with 45 enriched under AL feeding and 17 enriched under DR feeding (Figure 4A; Supplemental Figure 4A; and Supplemental Table 10). In plasma, a total of 160 sulfhydrated proteins were identified, with 3 enriched under AL feeding and 0 enriched under DR feeding (Figure 4B; Supplemental Figure 4B; and Supplemental Table 11). Between heart and plasma, in total 78 common sulfhydrated proteins were identified (Figure 4C; Supplemental Table 12), of which 4 biological functions/pathways were enriched, and include complement and coagulation cascades, staphylococcus aureus infection, HIF-1 signaling, and glycolysis/gluconeogenesis (Figure 4D). Of the sulfhydrated proteins enriched under AL and DR feeding in heart and plasma, 0 were common between the two tissues (Figure 4E). As no common sulfhydrated proteins were enriched as a function of diet in heart and plasma, we next examined tissue-specific functional and pathway enrichment analysis of sulfhydrated proteins. Heart contained 12 pathways enriched; with the top three being carbon metabolism, valine, leucine, and isoleucine degradation, and citrate/TCA cycle, under AL and 3 pathways enriched; complement and coagulation cascades, ferroptosis, and staphylococcus aureus infection, under DR feeding (Figure 4F, Supplemental Table 13). Plasma had no diet-specific pathway enrichment of sulfhydrated proteins (Supplemental Table 14).

**Figure 4:**
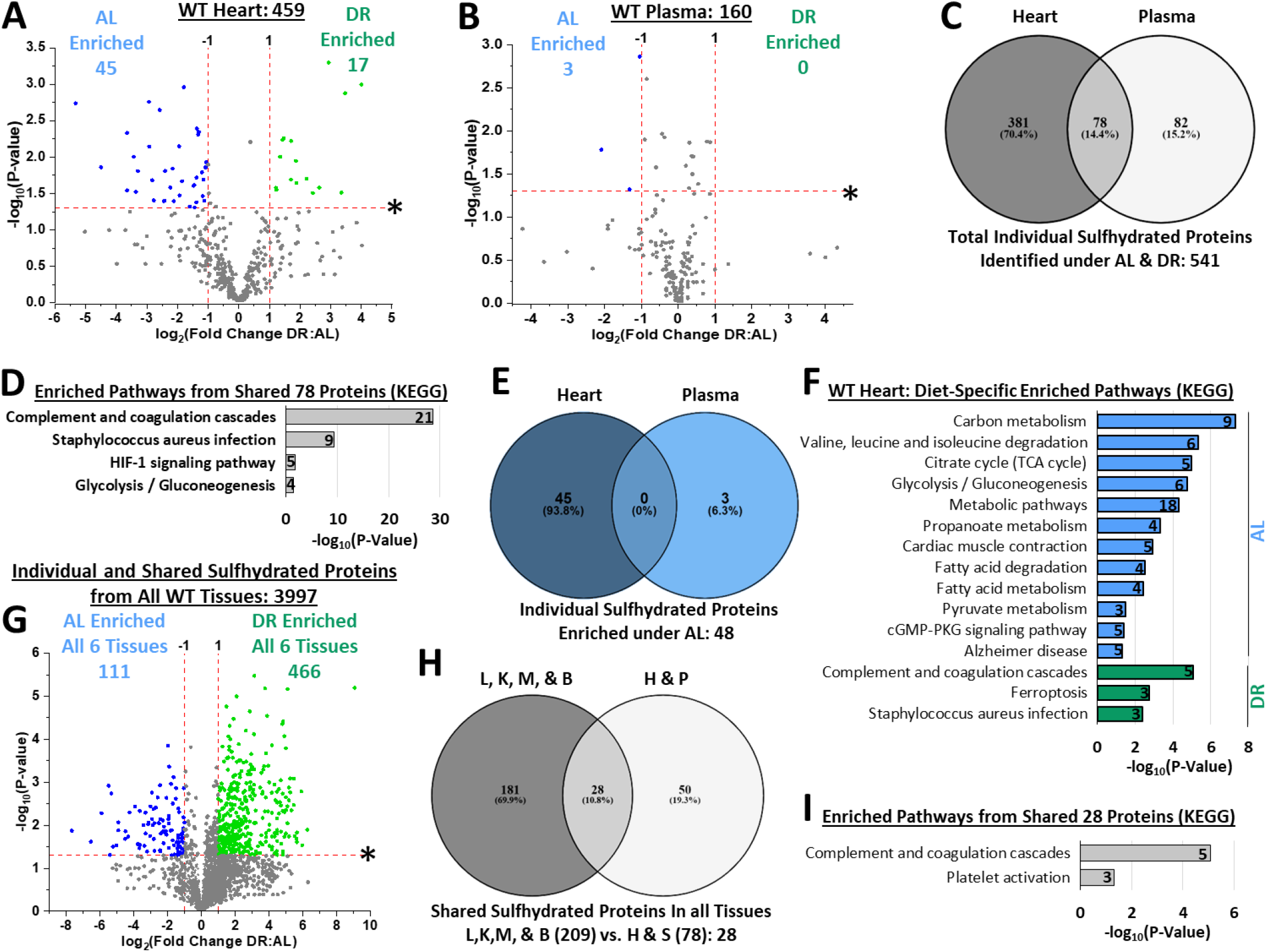
Sulfhydromes of heart and plasma respond differently to diet compared to those of liver, kidney, muscle, and brain. (**A-B**). Volcano plots showing biologically- and statistically-significant differentially abundant sulfhydrated proteins in heart (**A**), and plasma (**B**) from AL (n = 4 mice/group) versus DR (n = 5 mice/group) fed CGL WT mice. The log_2_(Fold Change DR:AL) X-axis displays the average fold change in spectral counts for each identified sulfhydrated protein while the −log_10_ Y-axis displays the calculated p-value when comparing the individual spectral count values for each identified sulfhydrated protein in a specific tissue from AL versus DR fed mice. The number of total sulfhydrated proteins identified in each tissue are given next to the tissue name, while blue (AL enriched) and green (DR enriched) colored dots and text indicate sulfhydrated proteins reaching both biological- and statistical- thresholds. Gray color dots indicate sulfhydrated proteins not reaching the criteria for both biological and statistical significance for enrichment under either diet. (**C**) Venn diagram showing the abundance of shared and non-shared sulfhydrated proteins in heart and plasma in CGL WT mice. 78 sulfhydrated proteins were shared amongst these two tissues. (**D**) KEGG biological function and pathway enrichment of the shared 78 proteins found in heart and plasma via g:Profiler analysis. The number in the bar indicates the number of individual sulfhydrated proteins identified for that specific pathway. (**E**) Overlap of AL enriched sulfhydrated proteins in heart and plasma. (**F**) KEGG biological function and pathway enrichment of AL enriched (blue bars) or DR enriched (green bars) sulfhydrated proteins in heart. (**G)** Comprehensive volcano plot showing diet-related protein sulfhydration enrichment in liver, kidney, muscle, brain, heart, and plasma between AL and DR fed mice. (**H**) Venn diagram to show shared and unshared sulfhydrated proteins, with 28 shared amongst all 6 tissues. (**I**) KEGG biological function and pathway enrichment of the shared 28 proteins found in all 6 tissues via g:Profiler analysis. The number in the bar indicates the number of individual sulfhydrated proteins identified for that specific pathway. Statistical significance for pathway enrichment plotted as −log_10_ (P-Value) and obtained via the g:Profiler g:SCS algorithm for KEGG database analysis. See also Supplemental Figure 4.

Sulfhydrated proteins not significantly impacted by diet did not trend for enrichment under either diet type (Figure 4A, B; gray dots). Function and pathway enrichment of these proteins revealed 27 and 6 pathways enriched in the heart and plasma, respectively (Supplemental Figure 4C, D). In heart, the top three pathways enriched include carbon metabolism, metabolic pathways, and pyruvate metabolism (Supplemental Figure 4C). In plasma, the top three pathways enriched include complement and coagulation cascades, staphylococcus aureus infection, and prion diseases (Supplemental Figure 4D). One shared pathway amongst this population of sulfhydrated proteins in heart and plasma included complement and coagulation cascades. Similar to kidney and muscle, the heart also showed enrichment for sulfhydrated proteins in the aging-related Parkinson and Alzheimer disease pathways (Supplemental Figure 4C).

When combining all six tissues analyzed in CGL WT mice, 3,997 individual and shared sulfhydrated proteins were identified, with 111 and 466 statistically and biologically enriched under AL and DR feeding, respectively (Figure 4G). The remaining proteins not meeting statistical and biological criteria still trended for enrichment under DR feeding (Figure 4G). To examine shared sulfhydrated proteins amongst all six tissues, we compared the 209 proteins identified as shared in liver, kidney, muscle, and brain with the 78 proteins identified as shared in heart and plasma. A total of 28 proteins were identified as common in all six tissues (Figure 4H, Supplemental Table 15). These 28 were involved in 2 blood-centric biological functions/pathways, which included complement and coagulation cascades, and platelet activation (Figure 4I).

Thus, diet impacts the mammalian sulfhydrome in a tissue-specific manner, with the majority of tissues tested having expanded and enhanced sulfhydromes after 1-week of 50% DR. However, increases or decreases in tissue-specific sulfhydromes surprisingly only correlated to H_2_S production capacity in liver and kidney, which were CGL-dependent for their DR-induced increase in H_2_S production. This is indicative of sulfhydration being more than just the presence of H_2_S modifying a protein, and suggests a more complicated systemic regulation of tissue-specific sulfhydrome maintenance either at baseline or when faced with a stressor, such as restricted food access.

### CGL deficiency limits the sulfhydrome under AL and DR feeding

DR-enhanced H_2_S production capacity (Hine et al., 2015; Yoshida et al., 2018) (Figure 1D, E, J, and K), stress resistance (Hine et al., 2015), endocrine response (Hine et al., 2017), angiogenesis (Longchamp et al., 2018), and lifespan (Kabil et al., 2011a) requires CGL activity. Thus, we next tested the CGL requirement for diet-induced changes in the sulfhydrome. As the dependency of H_2_S production in liver and kidney were primarily CGL derived (Figure 1A, B), we chose these same two tissues to examine CGL-dependent and independent sulfhydration at baseline and under DR.

In the liver, a total of 698 sulfhydrated proteins were detected in CGL KO mice; an approximate 30% reduction compared to CGL WT mice (Figure 5A, Supplemental Figure 5A, and Supplemental Table 16). Of these 698 proteins, 4 were enriched under AL feeding while 25 were enriched under DR feeding (Figure 5A, and Supplemental Table 16). In comparing total liver sulfhydrated proteins of WT and CGL KO mice, we discovered a total of 1,021 individual proteins, with 323; or 31.6%, unique to WT mice, 44; or 4.3 %, unique to KO mice, and 654; or 64.1%, shared between WT and CGL KO mice (Figure 5B). Biological function and pathway enrichment via g:Profiler analysis of these shared 654 sulfhydrated proteins utilizing the KEGG database revealed over 25 pathways, with the top three being metabolic pathways, carbon metabolism, and valine, leucine, and isoleucine degradation (Figure 5C). Pathway enrichment for the CGL-dependent 323 sulfhydrated proteins was found to total 8, and included aging and health-related targets such as metabolic pathways, oxidative phosphorylation, Parkinson disease, Huntington disease, non-alcoholic fatty liver disease, lysosome, Alzheimer disease, and thermogenesis (Figure 5D). Interestingly, no functions/pathways were enriched from the 44 unique sulfhydrated proteins found only in the livers of CGL KO mice. In comparing AL diet-enriched sulfhydrated proteins in WT and CGL KO mice, we found a total of 19 individual proteins enriched under AL, with 15; or 78.9%, unique to WT mice, 3, or 15.8%, unique to KO mice, and 1, or 5.3%, shared between WT and CGL KO mice (Figure 5E). The 3 proteins unique to CGL KO mice and the 1 protein shared between WT and CGL KO mice failed to account for any significant pathway enrichment. However the 15 proteins unique to AL fed WT mice livers resulted in the same 2 enriched pathways as detected previously (Figure 3E), and included selenocompound metabolism, and terpenoid backbone biosynthesis (Figure 5F). In comparing DR-enriched sulfhydrated proteins in WT and CGL KO mice, we found a total of 55 individual proteins enriched under DR, with 30; or 54.5%, unique to WT mice, 25; or 45.5%, unique to KO mice, and 0 shared between WT and CGL KO mice (Figure 5G). Interestingly, the 25 sulfhydrated proteins unique to CGL KO mice failed to account for any functional enrichment. The 30 sulfhydrated proteins unique for CGL WT mice under DR provided the same enriched function detected previously (Figure 3E), that being metabolic pathways (Figure 5H). Sulfhydrated proteins not significantly impacted by diet in CGL KO livers did not trend for enrichment under either diet type (Figure 5A; gray dots). Function and pathway enrichment of these proteins revealed 30 pathways (Supplemental Figure 5B, Supplemental Table 17). The top three pathways enriched include metabolic pathways, carbon metabolism, and valine, leucine, and isoleucine degradation (Supplemental Figure 5B).

**Figure 5:**
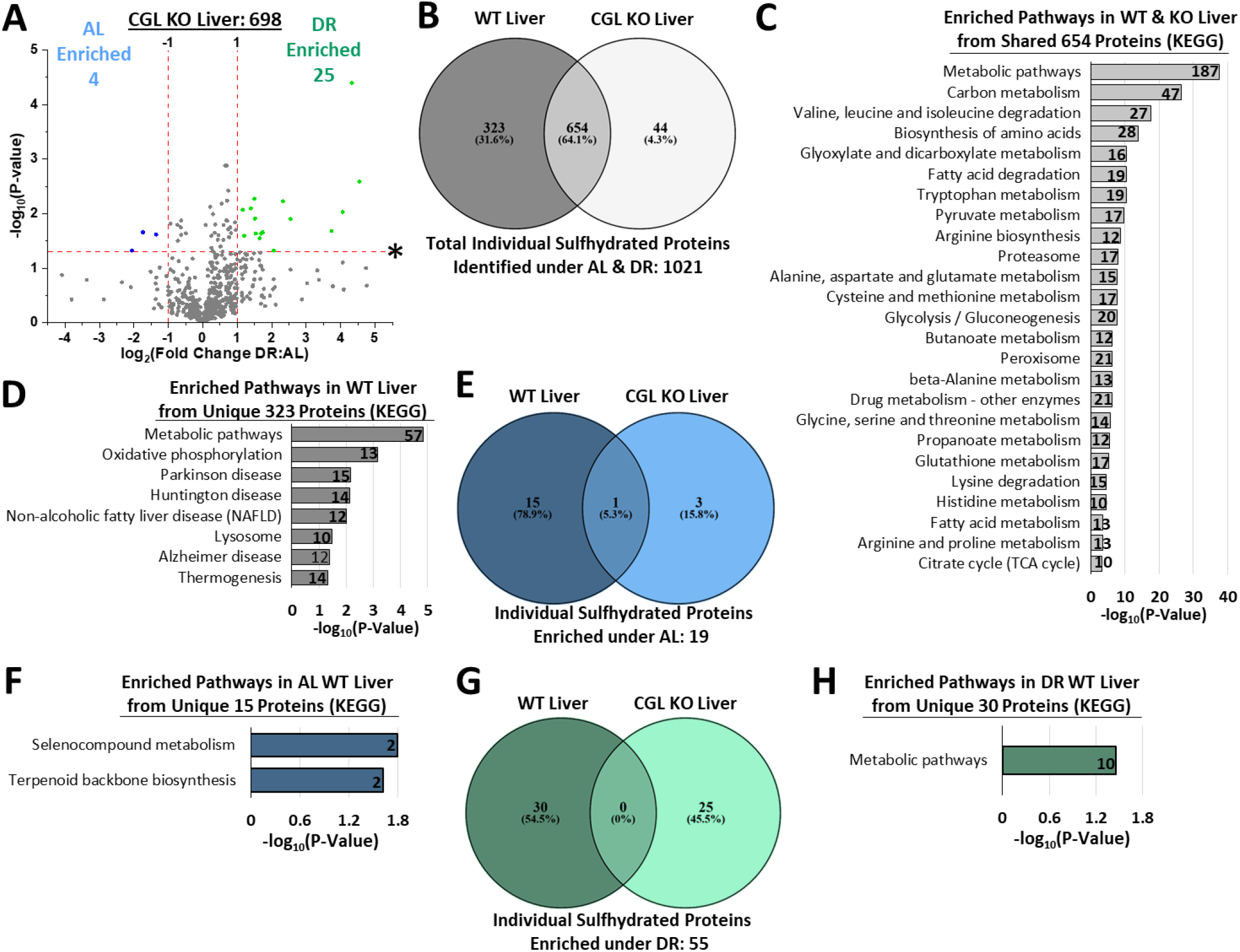
CGL dependent and independent sulfhydration and pathway enrichment in the liver. (**A**) Volcano plot showing biologically- and statistically-significant differentially abundant sulfhydrated proteins in liver from AL (n = 3 mice/group) versus DR (n = 3 mice/group) fed CGL KO mice. The log_2_(Fold Change DR:AL) X-axis displays the average fold change in spectral counts for each identified sulfhydrated protein while the −log_10_ Y-axis displays the calculated p-value when comparing the individual spectral count values for each identified sulfhydrated protein in liver from AL versus DR fed mice. The number of total sulfhydrated proteins identified was 698, while blue (AL enriched) and green (DR enriched) colored dots and text indicate sulfhydrated proteins reaching both biological (*2-fold enrichment*)- and statistical (p < 0.05)- thresholds. Gray color dots indicate sulfhydrated proteins not reaching the criteria for both biological and statistical significance under either diet. (**B**) Venn diagram to show shared and unshared sulfhydrated proteins between CGL WT and KO livers, with 654 common. (**C-D**) Enriched function/pathway analysis of the shared 654 proteins (**C**) or the 323 unique to CGL WT liver (**D**). Numbers inside the bars indicate the number of sulfhydrated proteins involved in that specific pathway. (**E**) Venn diagram to examine the number of shared and unshared AL enriched sulfhydrated proteins in WT and CGL KO livers. (**F**) Enriched function/pathway analysis of the 15 unique sulfhydrated proteins found in AL fed WT livers, similar to Figure 3E. (**G**) Venn diagram to examine the number of shared and unshared DR enriched sulfhydrated proteins in WT and CGL KO livers. (**H**) Enriched function/pathway analysis of the 30 unique sulfhydrated proteins found in DR fed WT livers, similar to Figure 3E. See also Supplemental Figure 5.

In the kidney, a total of 869 sulfhydrated proteins were detected in CGL KO mice; an approximate 20% reduction compared to CGL WT mice (Figure 6A, Supplemental Figure 6A, and Supplemental Table 18). Of these 869 proteins, 10 were enriched under AL feeding while only 5 were enriched under DR feeding (Figure 6A, and Supplemental Table 18). In comparing total kidney sulfhydrated proteins of WT and CGL KO mice, we discovered a total of 1,341 individual proteins, with 472; or 35.2%, unique; to WT mice, 268; or 20 %, unique to KO mice, and 601; or 44.8%, shared between WT and CGL KO mice (Figure 6B). Biological function and pathway enrichment via g:Profiler analysis of these shared 601 sulfhydrated proteins utilizing the KEGG database revealed over of 25 pathways, with the top three being metabolic pathways, carbon metabolism, and valine, leucine, and isoleucine degradation (Figure 6C). Pathway enrichment for the CGL-dependent 472 sulfhydrated proteins was found to total 2, and included metabolic pathways, and valine, leucine, and isoleucine degradation (Figure 6D) with no pathways enriched from the 268 proteins unique to CGL KO kidney. In comparing AL diet-enriched sulfhydrated proteins in WT and CGL KO mice, we found a total of 15 individual proteins enriched under AL, with 5; or 33.3%, unique to WT mice, 10; or 66.7%, unique to KO mice, and 0 shared between WT and CGL KO mice (Figure 6E). The 10 kidney proteins unique to AL fed CGL KO mice failed to account for any pathway enrichment. However the 5 kidney proteins unique to AL fed WT mice resulted in the same enriched pathway as detected previously (Figure 3F), and included glycine, serine, and threonine metabolism (Figure 6F). In comparing DR-enriched kidney sulfhydrated proteins in WT and CGL KO mice, we found a total of 21 individual proteins enriched under DR, with 16; or 76.2%, unique to WT mice, 5; or 23.8%, unique to KO mice, and 0 shared between WT and CGL KO mice (Figure 6G). The 5 sulfhydrated proteins unique to CGL KO mice failed to account for any functional enrichment. The 16 sulfhydrated proteins unique for CGL WT mice under DR provided the same enriched function detected previously (Figure 3F), that being nitrogen metabolism (Figure 6H). Sulfhydrated proteins not significantly impacted by diet in CGL KO kidneys trend for enrichment under AL feeding (Figure 6A; gray dots), which is opposite for what was detected in CGL WT kidneys (Figure 2B). Function and pathway enrichment of these proteins revealed 31 pathways (Supplemental Figure 6B, Supplemental Table 19). The top three pathways enriched include metabolic pathways, carbon metabolism, and citrate (TCA) cycle (Supplemental Figure 6B).

**Figure 6:**
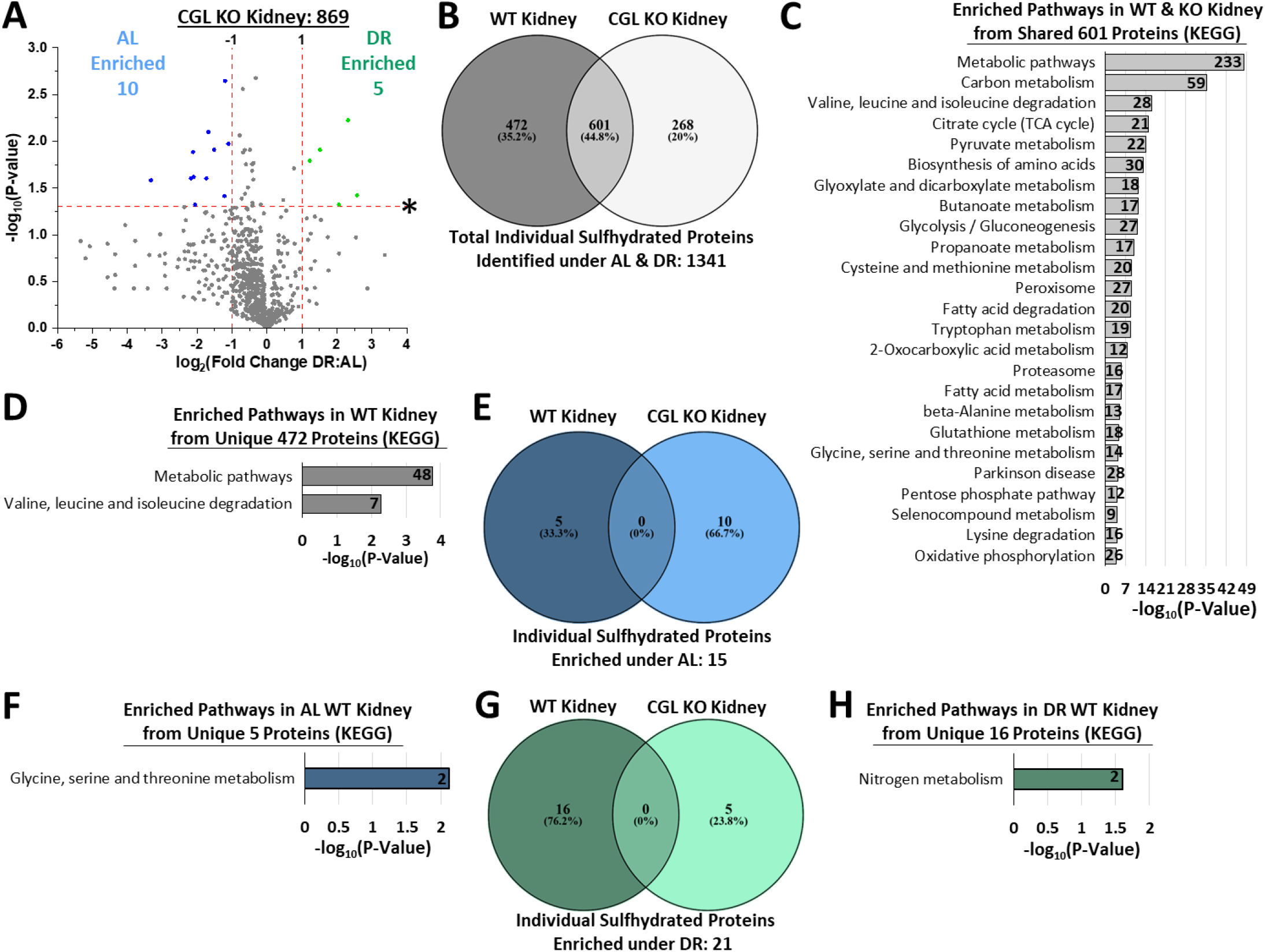
CGL dependent and independent sulfhydration and pathway enrichment in the kidney. (**A**) Volcano plot showing biologically- and statistically-significant differentially abundant sulfhydrated proteins in kidney from AL (n = 3 mice/group) versus DR (n = 3 mice/group) fed CGL KO mice. The log_2_(Fold Change DR:AL) X-axis displays the average fold change in spectral counts for each identified sulfhydrated protein while the −log_10_ Y-axis displays the calculated p-value when comparing the individual spectral count values for each identified sulfhydrated protein in kidney from AL versus DR fed mice. The number of total sulfhydrated proteins identified was 869, while blue (AL enriched) and green (DR enriched) colored dots and text indicate sulfhydrated proteins reaching both biological (*2-fold enrichment*)- and statistical (p < 0.05)- thresholds. Gray color dots indicate sulfhydrated proteins not reaching the criteria for both biological and statistical significance under either diet. (**B**) Venn diagram to show shared and unshared sulfhydrated proteins between CGL WT and KO kidneys, with 601 common. (**C-D**) Enriched function/pathway analysis of the shared 601 proteins (**C**) or the 472 unique to CGL WT kidney (**D**). Numbers inside the bars indicate the number of sulfhydrated proteins involved in that specific pathway. (**E**) Venn diagram to examine the number of shared and unshared AL enriched sulfhydrated proteins in WT and CGL KO kidneys. (**F**) Enriched function/pathway analysis of the 5 unique sulfhydrated proteins found in AL fed WT livers, similar to Figure 3F. (**G**) Venn diagram to examine the number of shared and unshared DR enriched sulfhydrated proteins in WT and CGL KO kidneys. (**H**) Enriched function/pathway analysis of the 16 unique sulfhydrated proteins found in DR fed WT kidneys, similar to Figure 3F. See also Supplemental Figure 6.

In examining CGL-dependent and independent sulfhydration regulation in the liver and kidney under both feeding conditions, we discovered CGL is required for approximately 30% of total proteins sulfhydrated in the liver and 20% in the kidney (Figure 5B and Figure 6B). Additionally, CGL is required for concerted sulfhydration of proteins for functional pathway enrichment under DR (Figure 5G, H and Figure 6G, H). Further examining the extent and requirement for CGL in shaping the sulfhydrome under DR in these two tissues, we re-plotted individual and shared sulfhydrated proteins from WT liver and kidney (Figure 7A) and from CGL KO liver and kidney (Figure 7B). In total, 2,063 proteins were detected in WT liver and kidney and 1,567 proteins in KO liver and kidney, resulting in an approximate 25% decrease in sulfhydrated proteins due to loss of CGL. While the fold enrichment in sulfhydrated proteins meeting both statistical and biological thresholds between DR and AL is similar between WT and KO; 2.2-fold and 2.1-fold, respectively, we have already demonstrated that the sulfhydrated proteins enriched in CGL KO mice, but not WT mice, for both of these tissues fail to fall under specific functional pathway enrichment. This is indicative that CGL may not just produce H_2_S for random sulfhydration or in itself randomly sulfhydrate proteins, but that CGL may process specificity for its protein targets. Also remarkable is when visually examining the plots for the sulfhydrated proteins that failed to meet statistical and biological thresholds of significance for enrichment under AL versus DR in liver and kidney (Figure 7A, B; gray dots). This subset of sulfhydrated proteins is prominently shifted towards enrichment under DR in WT mice, with ~75% with a DR:AL average spectral count ratio above 1 (Figure 7A). Conversely, this same subset of sulfhydrated proteins is shifted towards enrichment under AL in CGL KO mice, with only 38% having a DR:AL average spectral count ratio above 1 (Figure 7B). The overall reductions in sulfhydrated proteins in liver and kidney of CGL KO mice are proportional to their relative H_2_S production capacities, as seen in Figure 7C, where we show a strong, positive correlation between both CGL WT and KO tissue-specific H_2_S production capacities under AL feeding and the total numbers of sulfhydrated proteins identified (N = 8, r = 0.881, p = 0.0072). Removing CGL KO tissues from this analysis still provides an almost identical Spearman r-value and positive correlation, albeit one with a less-significant p-value (Supplemental Figure 7A) (N = 6, r = 0.8857, p = 0.0333), indicative that sulfhydration in CGL KO liver and kidney is appropriate for their levels of H_2_S production capacity. Thus, CGL-independent H_2_S production processes via CBS, 3MST, or non-enzymatic production (Yang et al., 2019) can contribute to sulfhydration in these two tissues, however CGL is required for DR-mediated sulfhydration of specific proteins for functional pathway enrichment as well as an amplified shift of the entire sulfhydrome.

**Figure 7:**
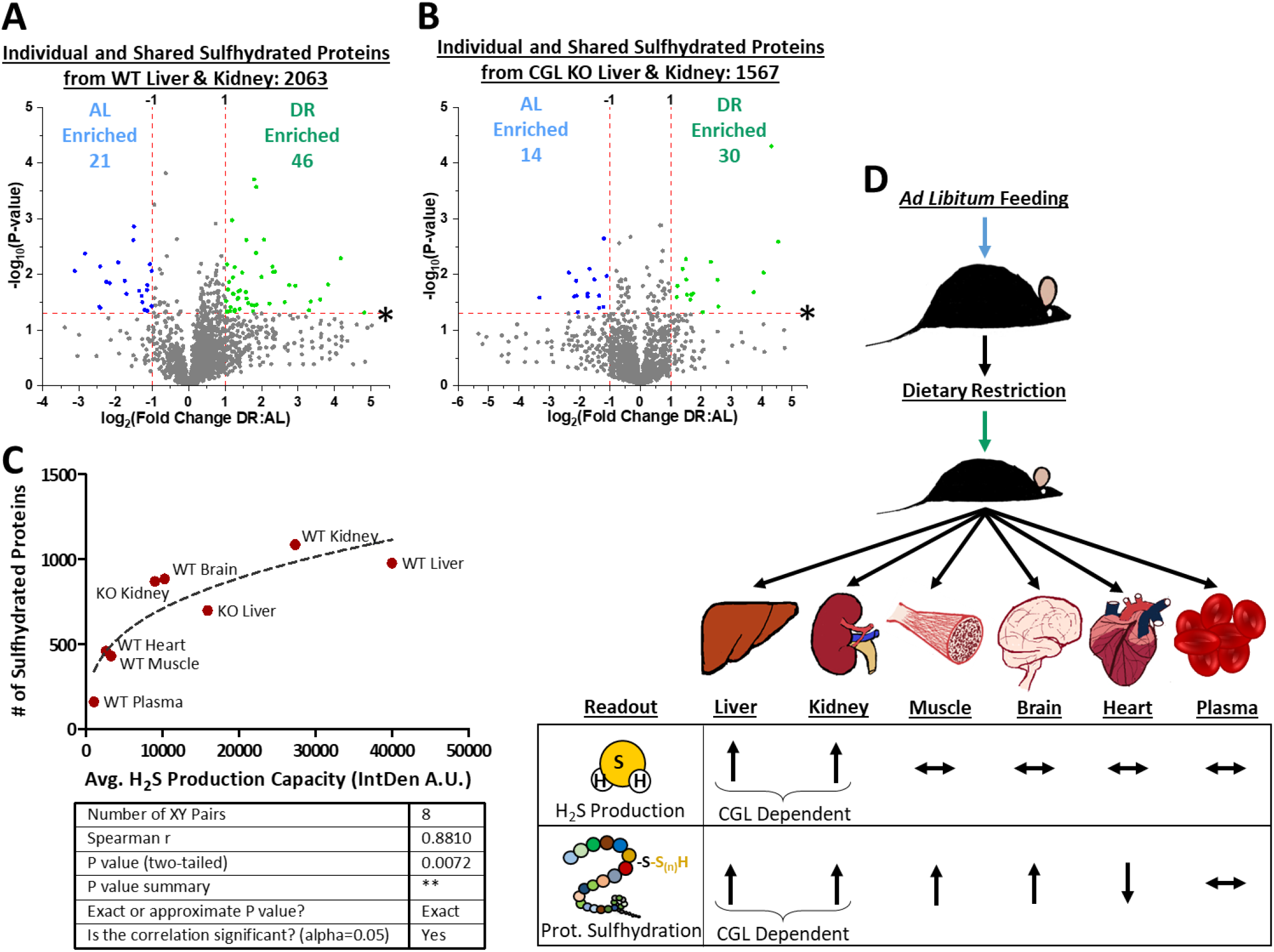
CGL is required for DR-mediated enrichment of the entire liver and kidney sulfhydrome. (**A-B**) Volcano plots re-imaging data from Figures 2, 5, and 6 to show biologically- and statistically-significant differentially abundant sulfhydrated proteins in kidney and liver from CGL WT AL (n = 4 mice/group) versus CGL WT DR (n = 5 mice/group) (**A**) and in kidney and liver from CGL KO AL (n = 3 mice/group) versus CGL KO DR (n = 3 mice/group) (**B**). The log_2_(Fold Change DR:AL) X-axis displays the average fold change in spectral counts for each identified sulfhydrated protein while the −log_10_ Y-axis displays the calculated p-value when comparing the individual spectral count values for each identified sulfhydrated protein in kidney and liver from AL versus DR fed mice. Blue (AL enriched) and green (DR enriched) colored dots and text indicate sulfhydrated proteins reaching both biological (*2-fold enrichment*)- and statistical (p < 0.05)- thresholds. Gray color dots indicate sulfhydrated proteins not reaching the criteria for both biological and statistical significance under either diet. (**C**) Correlation between average H_2_S production capacity and total number of sulfhydrated proteins in CGL WT and KO tissues. Arbitrary average H_2_S production capacity values determined from ImageJ IntDen function using the lead acetate/lead sulfide H_2_S assay as shown in Figure 1C (n = 3 samples/tissue) were plotted on the X-axis while the total number of sulfhydrated proteins stated in Figures 2, 4, 5, and 6 were plotted on the Y-axis (n = 9 samples/tissue for CGL WT and n = 6 samples/tissue for CGL KO). Graphpad Prism was used to fit the trendline and the given statistics calculated via XY analysis nonparametric correlation (Spearman) test with a two-tailed P- value and 95% confidence interval. (**D**) Model figure depicting the impact of DR on endogenous H_2_S production and protein sulfhydration in tissue specific and CGL-dependent manners. Arrows pointing up indicate an increase, arrows pointing down indicate a decrease, and horizontal arrows indicate no change. See also Supplemental Figure 7.

## Discussion

In the current study, we examined the multi-tissue and systemic extent to which dietary restriction (DR), a well-studied anti-aging intervention (Fontana and Partridge, 2015), impacts H_2_S production capacity and the sulfhydromes in liver, kidney, muscle, brain, heart, and plasma. In addition, we hypothesized the H_2_S producing enzyme cystathionine γ-lyase (CGL), which is required for stress resistance and angiogenesis benefits of short-term DR (Hine et al., 2015; Longchamp et al., 2018; Trocha et al., 2019), is partially required for baseline and diet-induced changes in tissue-specific sulfhydromes. We found H_2_S production is strongest in liver and kidney and primarily driven by CGL in these tissues, while other tissues had relatively weak H_2_S production independent of CGL. Short-term 1-week 50% DR increased H_2_S production capacity in a CGL-dependent manner in liver and kidney, while it had minimal effect on production capacities in the other tissues. Interestingly, despite only liver and kidney showing increased H_2_S production after DR, the sulfhydromes of muscle and brain were also expanded and enriched after one week of DR. The concerted augmentation for sulfhydrated proteins in liver and kidney to enrich specific biological functions and pathways was CGL-dependent. Also surprising, the heart sulfhydrome was diminished after DR. These results are summarized in the model Figure 7D. Below, we discuss these findings with regard to their contexts in the fields of aging and sulfide-redox biology while also addressing this study’s limitations and future directions.

Enhanced bioavailability and/or production capacity of endogenous H_2_S, particularly via CGL activity, has only recently been recognized as a common phenomenon and proposed mechanisms of action for dietary, genetic, and pharmacological models of longevity and healthy aging across evolutionary boundaries (Hine et al., 2018; Kabil et al., 2011a; Tyshkovskiy et al., 2019). However, the molecular mechanisms underlying many of the pleiotropic cellular, physiological, and systemic benefits of H_2_S, particularly and in the context of diet and aging, have remained unclear. As aging is a multifaceted and complex decline in numerous physiological and metabolic functions, the role H_2_S plays in combating these declines theoretically should not be limited to one molecular pathway, receptor, system, or mechanism. Most likely, it is through H_2_S performing several non-mutually exclusive functions related to antioxidant and redox homeostasis (Kabil et al., 2014), mitochondrial electron transfer (Luna-Sanchez et al., 2017), and protein modification and signaling via persulfidation/sulfhydration (Filipovic et al., 2018; Gao et al., 2015; Mustafa et al., 2009). Here, we focused on the latter mechanism, as little has been revealed regarding the entirety and scope of sulfhydration in mammals across several organ systems while under an anti-aging intervention. Additionally, uncovering the multi-organ sulfhydrome provides a resource and a primer for numerous downstream mechanistic and biochemical studies related to basic physiological processes and potential clinical issues impacted by, or the lack of, protein sulfhydration modifications.

By what means does protein sulfhydration, and its augmentation during DR and other anti-aging interventions, serve to extend lifespan and healthspan? As sulfhydration offers a protective barrier on protein cysteine residues from irreversible oxidative modifications (Filipovic et al., 2018), it first protects the proteome from unwanted oxidative stress and prevents the need to synthesize new proteins to replace those that have been damaged. It also offers neuroprotection by preventing protein misfolding and amyloid formation linked to a number of neurodegenerative diseases (Rosario-Alomar et al., 2015). Most importantly, it enhances the anti-aging and cytoprotective activities of longevity and metabolism associated proteins such as, but not limited to, glyceraldehyde phosphate dehydrogenase (GAPDH) (Gao et al., 2015; Mustafa et al., 2009), nuclear factor κB (NF-κB) (Sen et al., 2012), endothelial ATP-sensitive potassium channels (Mustafa et al., 2011), sirtuin 1 (SIRT1) (Du et al., 2019), and the E3 ubiquitin ligase parkin (Vandiver et al., 2013). As we have uncovered in this study well over 1,000 sulfhydrated proteins involved in numerous biological pathways across multiple tissues, these results hint at a far reaching and potent H_2_S-mediated change in global proteome stability and function.

As expected, sulfhydrome profiles were expanded under DR in the liver and kidney and this was CGL-dependent. While this is not the first study to recognize relative changes in tissue-specific protein sulfhydration as a function of CGL (Wedmann et al., 2016), it is the first to reveal the individual identities and abundances of these proteins, their functional and pathway involvement, and the dependence of both tissue-type, diet, and CGL to impact their sulfhydration status. These sulfhydration events in liver and kidney imparted on numerous metabolic, cellular homeostasis, and signaling pathways affected by the aging process, notably the proteasome, Parkinson’s disease, and Huntington’s disease. What we did not expect is the increase in global sulfhydration in muscle and brain under DR, as both of these tissues fail to augment H_2_S production capacity under DR. This indicates DR-mediated enhanced sulfhydration in these tissues may not be tissue autonomous, and the sources of H_2_S that enhance sulfhydration in these tissues arise from distal tissues, such as the liver or kidney, or from diet-modifiable H_2_S producing endothelial cells residing within the tissue (Longchamp et al., 2018). While the idea of H_2_S itself being a circulating gasotransmitter in blood is somewhat controversial, mostly due to a number of issues regarding H_2_S detection methodology preferences (Olson, 2009), our data show these weak producing tissues still augment their sulfhydromes under DR. The failure to detect increased H_2_S production capacity or enriched sulfhydration in plasma may suggest DR does not induce systemic increases in sulfide availability and local enrichment of sulfhydration may be tissue autonomous via direct enzymatic sulfhydration of proteins via CGL, CBS, and/or 3-MST. However, only plasma was assayed here and not whole blood including white and red blood cells, of which the latter act as a sink for H_2_S via exchange in the microcirculation of strong H_2_S-producing tissues (Jennings, 2013). Thus, we cannot entirely rule out DR-enhanced H_2_S or persulfides in circulation as the mechanism for augmented sulfhydration in marginally H_2_S producing tissues. Future studies will need to test the requirement and protein-protein interactions for CGL, CBS, and 3-MST in DR induced sulfhydration changes as well as identify sources of H_2_S and/or polysulfides for sulfhydration in brain, muscle, and other weak H_2_S producing tissues.

The absolute number of sulfhydrated proteins in the brain and their expansion under DR was unforeseen and surprising. This was primarily due to the lack of detectable DR-enhanced H_2_S production (Figure 1H) along with the purported sensitivity to H_2_S-induced toxicity in the central nervous system (CNS) owning to its low or non-existent sulfide quinone oxidoreductase (SQR) expression (Supplemental Figure 1H) (Lagoutte et al., 2010; Linden et al., 2012). SQR is a key mitochondrial enzyme catalyzing the initial steps of H_2_S oxidation, detoxification, and removal (Horsman and Miller, 2016; Libiad et al., 2014). However, H_2_S does serve beneficial roles in the brain, as decreased H_2_S or disruption of H_2_S producing enzymes in the brain or neuronal cells are associated with aging-related Parkinson’s, Huntington’s, and Alzheimer’s disease-like phenotypes (Paul and Snyder, 2018), while controlled H_2_S exposure serves to protect against cerebral ischemia-reperfusion injury (Yin et al., 2013) and slow the progression of neurodegeneration (Kamat et al., 2013). This suggests the lack of SQR enables H_2_S that reaches or is produced in the CNS to be utilized primarily for protein sulfhydration, and the benefits of H_2_S in the CNS are primarily derived through sulfhydration. An example of this transformation and biological impact in the brain can be seen when examining pyruvate kinase (PKM), the protein in the brain with the greatest increase; > 500-fold, in sulfhydration enrichment under DR detected in our study. PKM catalyzes the final rate-limiting step of glycolysis to produce pyruvate and ATP, and modification of cysteine residues allosterically regulate PKM’s enzymatic activity (Gao et al., 2015; Mitchell et al., 2018). Importantly, it was previously shown that oxidation or nitrosylation of key cysteine residues prevent and/or destabilize the active tetrameric form of PKM and drive PKM to its inactive monomeric form (Mitchell et al., 2018), while CGL generated H_2_S increases PKM activity without altering PKM protein levels (Gao et al., 2015). Thus, DR may stimulate glycolytic flux in the brain via increased sulfhydration of PKM and maintenance of its active state. The functional impact of sulfhydration on other individual proteins identified here but not in previous studies will be of great interest and focus for numerous future studies.

Higher SQR activity in liver, kidney or heart potentially limits protein sulfhydration via the oxidation of and removal of H_2_S. This may explain the decrease in sulfhydration in heart upon DR, as H_2_S may impart its beneficial cardiac functions (Kondo et al., 2013; Polhemus et al., 2013), particularly under ischemic-reperfusion injury, via SQR-mediated mitochondrial oxidation of H_2_S rather than sulfhydration-centric mechanisms. Future studies need to examine SQR expression or activity in cardiac tissue in response to DR to elucidate its DR-induced loss of sulfhydration. Conversely, the decrease in cardiac sulfhydration upon DR may not be solely due to increased H_2_S oxidation and removal via SQR, but through potential depersulfidation of proteins via the thioredoxin (Trx) system (Doka et al., 2016; Wedmann et al., 2016). The function and potential benefit of removing sulfhydryl groups on proteins in cardiac tissue upon DR is not well understood or described in our study. However, Trx mediated cleavage of cysteine persulfides results in H_2_S release (Wedmann et al., 2016). Thus, a potential hypothesis is that depersulfidation in the heart releases H_2_S and/or polysulfides for potential beneficial use elsewhere, perhaps via SQR for mitochondrial protection and appropriate changes in metabolism and fuel usage during ischemic reperfusion. Further examination into Trx depersulfidase activity during DR, and if it is stunted in tissues with enhanced sulfhydrome profiles and boosted in tissues with decreased sulfhydrome profiles, is warranted. This also offers an alternative explanation for the changes in tissue-specific sulfhydromes as a function of DR that is mutually exclusive of H_2_S production capacities. While SRQ and Trx may impart some regulation into sulfhydrome profiles under different dietary cues, our results in Figure 7C support the hypothesis that in general the overall number of sulfhydrated proteins in a tissue is strongly correlated to its H_2_S production capacity under *ad libitum* feeding conditions.

Reactive oxygen and nitrogen species (RONS), much like H_2_S, display hormetic dose responses and act as secondary cellular messengers. Similarly, transiently increased RONS production during DR through increased mitochondrial fatty acid oxidation is thought to drive many of the benefits of DR and DR-mimetics (Calabrese and Mattson, 2011; Hine and Mitchell, 2012; Ristow and Schmeisser, 2011; Ristow et al., 2009). Thus, in light of the requirement for the activation of cysteine residues via oxidation prior to sulfhydration by H_2_S (Filipovic et al., 2018; Gao et al., 2015), it is appropriate to consider the interdependence of DR-mediated augmentation in RONS and H_2_S for promoting changes in the sulfhydrome during DR and together driving adaptive mechanisms of antioxidant protection, stress resistance, and longevity. Likewise, non-dietary means to extend lifespan in the worm *Caenorhabditis elegans* via germline loss, low levels of paraquat, or mild inhibition of respiration simultaneously triggered the generation of RONS and H_2_S, with the boost of both hormetic factors responsible and necessary for the stress resistance and slowed aging found in these manipulations (Wei and Kenyon, 2016). Thus, this hypothesis can be framed in such a way that the same cues that lead to increased H_2_S production also boost non-harmful RONS production, and these RONS prime cysteine residues for sulfhydration. This ultimately alters the protein’s structure, interactions, and function concurrent with protecting the protein’s cysteine residues from irreversible over-oxidation. Taking into context these biological complexities of the sulfhydration process that may not be fully recapitulated in cell culture systems or *in vitro* with exogenous addition of H_2_S highlights the need for more studies to be done *in vivo* and under various conditions to take snapshots of malleable and dynamic tissue-specific and/or organism-specific sulfhydromes.

Limitations to our study include the lack of sulfhydration site identification on individual proteins and their mechanisms of action. While we identified specific proteins as sulfhydrated, we did not shown what cysteine residue or residues are sulfhydrated. Additionally, the pathway enrichment analysis simply shows the involvement of these sulfhydrated proteins in specific pathways, but the direction of metabolic flux or pathway activation/silencing is not inferred. Our biological experimental setup may also limit or influence the sulfhydromes identified, as we used male mice at 6-months of age, tissues harvested in the morning, and under a a 1-week dietary intervention. Changing the variables of sex, age, harvest timing, and diet could impact the tissue specific sulfhydromes, by either expanding them or condensing them. Insights into how these variables impact H_2_S and sulfide biology include the sex-, diet-, and circadian-dependent differences in growth hormone and thyroid hormone signaling, of which are potent regulators of H_2_S production (Hine et al., 2017). Additionally, diurnal fluctuations in plasma H_2_S exist (Jin et al., 2017), so time of tissue harvest may impact sulfhydrome profiles. Likewise, technical variables in our experimental setup possibly influence the sulfhydromes identified. As we examined whole tissue lysates, we may be missing tissue region-specific sulfhydration enrichments. The ratio of NM-biotin to protein added during the BTA could also affect the results, Gao, et al. previously showed how lower concentrations of NM-biotin result in selective labeling of the more highly reactive sulfhydrated cysteine groups versus the less reactive non-sulfhydrated cysteine thiol (Gao et al., 2015). By increasing the amount of NM-biotin added, it could be possible to diminish the overall number of sulfhydrated proteins identified via labeling both the sulfhydrated cysteine and the non-sulfhydrated cysteines and thus preventing elution with DTT, and/or offering more selective isolation to proteins with multiple sulfhydration modifications. Thus, due to the dynamic rather than static nature of the sulfhydrome, the biological or technical experimental factors listed above, or use of another sulfhydration/persulfidation detecting methodology such as the biotin-switch (Mustafa et al., 2009) or tag-switch methods (Kouroussis et al., 2019), generation of alternative sulfhydrome profiles is likely. Despite these limitations, our work provides a starting point by identifying these proteins and their sulfhydration changes specific to tissue, diet, and CGL status. Future work is needed to identify the residue or residues of each protein sulfhydrated, how these residues are sulfhydrated; i.e. passive or targeted, and the functional changes in protein stability, location, and activity induced by these modifications.

In summary, we reveal the global sulfhydration profile alterations in major metabolic organs under *ad libitum* and calorically restricted feeding. We show the importance of CGL enzyme for protein sulfhydration in liver and kidney. Ultimately, this study establishes a data resource for dietary restriction and H_2_S-related research while prompting the need to decipher downstream effects of these sulfhydrated proteins in metabolism, stress resistance, and longevity for ultimate use in developing sulfhydrated protein-based interventions and diagnostics.

## Supporting information

Supplemental Tabe 4

## Acknowledgments

We thank Drs. Xing-Huang Gao and Maria Hatzoglou of Case Western Reserve University for technical insights and advice regarding the BTA method, Paul Minkler and Ling Li from the Proteomics and Metabolomics Core at the Cleveland Clinic for technical assistance and advice regarding proteomics and LC-MS/MS analysis, and Drs. Yoko Henderson and Jie Yang for constructive comments on experimental design.

## Author contributions

NB and CH conceptualized the project; NB, BW, and CH designed experiments; NB, CL, BW, and CH performed experiments and analyzed data, RW contributed valuable CGL KO mouse models; and NB and CH wrote the paper with all authors editing and approving the manuscript.

## Data and materials availability

The authors declare that the majority of the data supporting the findings of this study are available within the paper and its supplementary information files and tables. Further information and requests for resources and reagents should be directed to and will be fulfilled by the Lead Contact, Christopher Hine.

## Materials and Methods

### Animal Husbandry and Diet Intervention

All experiments were performed under the approval of the Cleveland Clinic Institutional Animal Care and Use Committee (IACUC), protocol number 2016-1778, and followed the National Institutes of Health Guide for the Care and Use of Laboratory Animals. Mice were bred, weaned between 21-23 days of age, and maintained under standard barrier housing conditions in the Cleveland Clinic Lerner Research Institute Biological Resource Unit on a 12-hour light/ 12-hour dark cycle, temperature between 20–23°C, 30%–70% relative humidity, and with initial *ad libitum* access to standard rodent food (Envigo #2918) and drinking water until the dietary intervention. Mice utilized were F2 generation 6-month old cystathionine γ-lyase (CGL) wildtype (WT) and total body knock out (KO) male mice obtained from initial parental CGL Het × Het breeding to generate CGL WT and KO F1s, and then breeding CGL WT × WT or CGL KO × KO F1s to generate the littermate and/or age-matched F2 experimental animals. Mice were group housed during all experimental procedures, with 3-5 mice per cage. The CGL WT and KO mice were originally generated on a mixed 129/C57BL/6 background as described previously (Hine et al., 2015; Yang et al., 2008) and then subsequently rederived into pathogen free C57/BL6 mice at Jackson Laboratories prior to colony establishment at the Cleveland Clinic. At 6-months of age, mice were switched to the experimental AIN-93G-based diet (Research Diets D10012G-2V-Formula 1) and exposed to *ad libitum* access for several days to adapt to the new food and for monitoring intake. The experimental diet consists of 20% of calories from protein, 64% of calories from carbohydrate, and 16% calories from fat, and has a 2x concentrations of mineral mix S10022G, vitamin mix V10037, and choline bitartrate to ensure proper nutrition and avoid malnutrition during the dietary restriction period. The powdered food mix was added in a 1:1 ratio (grams:mL) to a 2% agar (Sigma #A1296) solution in water before solidification to a semi-solid consistency that lessens the potential for hoarding of food in group housing of mice and increases accuracy of measuring food consumption as large chunks can’t be taken at one time and the food doesn’t crumble like with standard rodent chows. *Ad libitum* food intake per cage was measured daily for four days to determine the correct amount to restrict to achieve a 50% reduction in food intake, and thus a 50% reduction in calories consumed. After randomly assigning *ad libitum* (AL) or diet restriction (DR) feeding to the cages, food intake and body mass were measured over the 1-week intervention. AL fed mice were provided 24-hour access to the diet and the DR mice fed their calculated allotment near the start of their dark phase at 7pm to limit disturbances in circadian rhythms and feeding patterns between the two groups (Acosta-Rodriguez et al., 2017; Froy et al., 2009). After the 1-week dietary intervention, mice were euthanized in the late morning (between 9am-12pm) via isoflurane anesthesia overdose followed by cervical dislocation and harvest of liver, kidney, heart, muscle (quadriceps), brain, and plasma. Plasma was collected via retro-orbital bleed and the blood immediately placed into lithium-heparin-coated tubes (Terumo #T-MLH). Tubes were centrifuged to separate RBCs from the serum/plasma. Collected tissues were immediately placed in 1.5 mL centrifuge tubes, flash frozen in liquid nitrogen, and then stored at −80°C until further analysis.

### Lead acetate/lead sulfide assay to detect enzymatic H_2_S production

The endogenous H_2_S production capacity of tissues were measured by following the lead acetate/lead sulfide method previously described (Hine et al., 2015; Hine and Mitchell, 2017). Briefly, ~80mg of tissue is first placed in 1.5 mL microcentrifuge tubes containing 250 μL of 1x passive lysis buffer (Promega) and homogenized, followed by multiple rounds of flash freezing/thawing using liquid nitrogen. After homogenization and lysis, tubes were briefly spun down to remove debris and the protein supernatant was saved. Protein concentration of the supernatant was measured with bicinchoninic acid assay (BCA) kit (Bio-Rad) followed by normalization of proteins via the addition of 1x passive lysis buffer. Next, the lead acetate/lead sulfide assay is setup by initially preparing the reaction mixture of 10mM L-cysteine (Sigma #168149) and 1mM pyridoxal phosphate (PLP) (Sigma #9255) in PBS, with 150 μL placed into each well of a 96-well plate. 100 μg of proteins from each tissue or 20 μL of plasma are added to each respective well, then the plate is overlaid with filter paper embedded with lead acetate (Sigma #316512) and an addition weight on top, and incubated at 37 °C until a desirable amount of lead sulfide is detected for proper quantification using ImageJ densitometry analysis via the IntDen function.

### Biotin thiol assay (BTA)

The isolation and detection of sulfhydrated proteins were performed using an adaptation to the biotin thiol assay (BTA) as previously described by Gao, *et al*. (Gao et al., 2015). To isolate sulfhydrated proteins, tissues were first lysed with RIPA lysis buffer (Thermo Fisher Scientific, # 89900) containing protease inhibitor cocktail (Thermo Fisher Scientific, #78415). Protein concentration was determined via BCA kit (Bio-Rad) and all samples normalized to the same protein concentration. Next, 7 mg of protein were incubated with 343 μM Maleimide-PEG2-biotin (Thermo Fisher Scientific, #21901BID) at room temperature for 30 minutes with agitation by precipitating the alkylated protein with 1 mL of 100% cold acetone by incubating at −20°C for 30 min. Additional washes were performed with 1 mL of 75% cold acetone followed by centrifugation and removal of acetone. Proteins were then resuspended in 0.25 mL of suspension buffer (RIPA + 1% SDS, pH 7.5) followed by adding 0.75 mL of neutralization buffer (30 mM Tris, 1 mM EDTA, 150 mM NaCl, 0.5 % Triton X-100, pH 7.5). Alkylated proteins were incubated in 0.39 mL of streptavidin-agarose resin (Thermo Scientific, #20347) contained in spin columns (Pierce, Catalog No. 69705) and kept rotating overnight at 4°C. Avidin beads were washed six times in 0.85 mL of wash buffer 1 (30 mM Tris, 1 mM EDTA, 150 mM NaCl, 0.5 % Triton X-100, pH 7.5) followed by another six washes with 0.85 mL of wash buffer 2 (30 mM Tris, 1 mM EDTA, 600 mM NaCl, 0.5 % Triton X-100, pH 7.5) and finally another three washes with 0.85 mL of wash buffer 3 (30 mM Tris, 1 mM EDTA, 100 mM NaCl, pH 7.5). Resin with bound proteins was first incubated with 500 μL elution buffer without DTT for 30 min at 25°C which serves as a negative control and then subsequently with 500 μL 20 mM DTT for 30 min at 25°C to elute sulfhydrated proteins. The addition of DTT selectively elutes sulfhydrated proteins, as it breaks the S-S bond but not the S-NM-Biotin bond, as can be seen with the −DTT control gel lanes having a lower detectable protein compared to the +DTT gel lanes (Supplemental Figure 2 B, C, D, E, F; Supplemental Figure 4 A, B; Supplemental Figure 5 A; and Supplemental Figure 6 A), supporting the effectiveness of the BTA method. Eluted proteins were run through Amicon Ultracel 10K (Millipore, #UFC501096) to concentrate in a final volume of 50μL. After obtaining this final protein solution with sulfhydrated proteins, subsequent gel electrophoresis analysis and mass spectrometry-based label-free protein identification is commenced to visualize, identify, and quantify the sulfhydrated proteins.

### Gel Electrophoresis and in-gel digestion

1.5 mm thick hand cast sodium dodecyl sulfate (SDS)-12% polyacrylamide gels were used to separate and visualize purified sulfhydrated proteins via staining or Western Blot relative to −DTT controls and input controls via gel electrophoresis. 0.75 mm thick one-dimensional (SDS)-12% polyacrylamide hand cast gels were used to performed gel electrophoresis to separate sulfhydrated proteins for in-gel digestion and downstream HPLC-MS/MS analysis. For both of these processes, 11 μL of proteins from input, eluted −DTT samples, and +DTT samples, were mixed and denatured by boiling at 100°C with 2μL of 5x Laemmli loading buffer (Fisher Scientific, Catalog No. 39001) and loaded into each lane, with 7 μL of PageRuler Plus Prestained (Thermo Fisher #26619) protein ladder in the first lane. Gels were run at 135 volts for 2 hours for imaging only and Western blot endpoints, and for 13-15 minutes for gels purposed for obtaining sulhfydrated proteins for in-gel digestion and HPLC-MS/MS analysis. Gels for imaging only were then carefully washed with ddH_2_O for 10 minutes, protein bands stained with colloidal blue staining kit (Thermo Fisher, #LC6025) for 3 hours at room temperature, and followed by another 3 hours of washing with ddH_2_O. Clean and visible protein containing gels were then scanned (Epson, #LW8W004863) to obtain images and then stored at 4°C in 5% acetic acid and 10% glycerol solution in a heat-sealable roll stock pouches (Fisher Scientific, #0181226E). Proteins in gels used for Western Blotting were then transferred to polyvinylidene difluoride membranes (Whatman), blotted for GAPDH (Abcam #ab8245) or α-tubulin (Abcam #ab4074) followed by horseradish peroxidase-conjugated secondary antibody (Abcam #97051) and visualized using SuperSignal West Femto Maximym Sensitivity Substrate (Thermo Scientific #34096) on an Amersham Imager 600 (General Electric). Gels used for in-gel digestion and downstream HPLC-MS/MS analysis were then washed for half an hour with ddH_2_O at room temperature. Protein bands were visualized by staining with gel code blue stain reagent (Thermo Fisher scientific, #24590) for one hour followed by washing the gel with ddH_2_O for another one hour at room temperature with gentle shaking. Clean and visible protein containing gels were scanned by using the Epson Scanner. A clean scalpel was first used to excise the entire band, and then the band was further cut into smaller bands of approximate 1 mm^3^. All of the smaller gel pieces from the original band were transferred to 0.2 mL of wash buffer (50% ethanol and 5% acetic acid) in a 1.5 mL tube and washed overnight at room temperature. The following day gel pieces were dehydrated with acetonitrile and dried in a SpeedVac for 6 minutes. Sulfhydrated proteins in the dried bands were reduced with 0.1 mL of 65 mM dithiothreitol (DTT) (Fisher Scientific # BP172-5) at room temperature for 30 minutes, followed by alkylation for another 30 minutes at room temperature with 0.1 mL of 162 mM iodoacetamide (Fisher Scientific, #AC122270050). Gel pieces were dehydrated two times with 0.2 mL acetonitrile and rehydration with 0.2 mL of 100 mM ammonium bicarbonate. Then, 40 μL of 10 ng/μL trypsin (Promega, #V5111) was added into the dried gel pieces with an additional 20 μL of 50 mM ammonium bicarbonate added to completely cover the gel pieces and kept overnight at room temperature for tryptic digestion. 0.08 mL of extraction buffer (50% acetonitrile + 5% formic acid) was added twice for 10 minutes at room temperature each time to extract the digested peptides from the gel pieces followed by micro centrifuging for 10 seconds. Supernatants were combined and transferred into a new 0.5 mL centrifuge tube and dried at room temperature in a Speedvac for 3 hours. 30 μL of 1% acetic acid was added into the dried peptides and transferred into an HPLC vial w/ cap (Sun-Sri, #200 050 & 501 313) for further mass spectrometric analysis.

### HPLC-Tandem mass spectrometric analysis/Orbitrap tribrid fusion lumos analysis

Digested peptides were analyzed on a ThermoFisher Scientific UltiMate 3000 HPLC system (ThermoFisher Scientific, Bremen, Germany) interfaced with a ThermoFisher Scientific Orbitrap Fusion Lumos Tribrid mass spectrometer (Thermo Scientific, Bremen, Germany). Liquid chromatography was performed prior to MS/MS analysis for peptide separation. The HPLC column used is a Dionex 15 cm × 75 μm Acclaim Pepmap C18, 2 μm, 100 Å reversed-phase capillary chromatography column. 5 μL volumes of the peptide extract were injected and peptides eluted from the column by a 90-minute acetonitrile/0.1% formic acid gradient at a flow rate of 0.30 μL/min and introduced to the source of the mass spectrometer on-line. Nano electrospray ion source was operated at 2.3 kV. The digest was analyzed using the data dependent multitask capability of the instrument acquiring full scan mass spectra using a Fourier Transform (FT) orbitrap analyzer to determine peptide molecular weights and collision induced dissociation (CID) MS/MS product ion spectra with an ion-trap analyzer to determine the amino acid sequence in successive instrument scans. The MS method used in this study was a data-dependent acquisition (DIA) with 3 second duty cycle. It includes one full scan at a resolution of 120,000 followed by as many MS/MS scans as possible on the most abundant ions in that full scan. Dynamic exclusion was enabled with a repeat count of 1 and ions within 10 ppm of the fragmented mass were excluded for 60 seconds. Chromatograms obtains from this HPLC-MS/MS method are included in Supplemental Data File 1.

### Bioinformatics for Peptide identification and quantification

In order to do label free quantitative and qualitative proteomics analysis we used Mascot, SEQUEST and Scaffold software packages. These software were used for protein identification by converting raw spectrometric data into protein IDs and for relative quantification of protein abundance via label-free spectral counting. For protein identification, three search engines were used including Mascot, Sequest which is bundled into Proteome Discoverer 1.4, and X!Tandem which is bundled into Scaffold 4.8.7. For the Mascot searches, primary raw MS/MS data were converted into its MGF (Mascot Generic File) format files by using Discoverer Daemon 1.4 software (Licensed under Cleveland Clinic Proteomics Core). The parameters selected for these searches include the following: Database: SwissProt, Taxonomy: Mus Musculus, Enzyme: Trypsin, Fixed modification: Carbamidomethyl(C), Variable modification: Oxidation of Methionine, Precursor Mass Tolerance: 10 ppm, Fragment Mass Tolerance: 0.8 Da, Peptide charge: 2+, 3+ and 4+, with the monoisotopic and decoy database present.

Scaffold, an online database software (version Scaffold_4.8.7, Proteome Software Inc., Portland, OR) (Craig and Beavis, 2004) was used to validate peptide and protein identifications and to create a comprehensive lists of target proteins. Protein and peptide identifications were validated using a decoy database strategy (reverse sequence of each protein for use as a decoy) and the decoy rate was set to zero, which may limit the total number of proteins identified but enables a higher confidence in protein identity for those that do meet all stringent criteria. Other settings for Scaffold included only allowing positive IDs if they could be established at greater than 99.9% probability and contained at least 2 identified peptides by the Peptide Prophet algorithm (Keller et al., 2002; Nesvizhskii et al., 2003) with Scaffold delta-mass correction. Proteins that contained similar peptides and could not be differentiated based on MS/MS analysis alone were grouped to satisfy the principles of parsimony. Additionally, to remove any false positive hits, we individually analyzed all of the proteins identified to ensure each one contained at least one cysteine residue by using the mouse protein amino acid sequences from UniProt Knowledgebase protein database (Proteome_ID/Tax_ID: UP000000589/10090). Those that did not contain at least one cysteine residue, and thus could not theoretically be sulfhydrated (approx. ~ 2-4% of our initial findings), were removed from our database and from further analysis. Peptide identifications were also required to exceed specific database search engine thresholds, including Mascot identifications required at least 40 ion score, Sequest identifications required at least XCorr (+1) is 1.5, XCorr (+2) is 2, XCorr (+3) is 2.25, and XCorr(+4) is 2.5, and X!Tandem identifications required at least 2.

Label-free spectral counting was used to determine relative differences in sulfhydrated protein abundance between AL versus DR fed groups within a specific tissue (Asara et al., 2008). The spectral counts, which are defined as the total number of spectra identified for a specific protein (Lundgren et al., 2010), have integer values ranging from 0 for proteins below the level of detection up to 505. Theoretically, the higher the spectral count for a given protein equates to higher abundance of that protein in a given tissue sample, where the number of spectra matched to peptides from a specific protein are used as a surrogate measure of protein abundance (Choi et al., 2008). The average spectral counts for each protein in each tissue were calculated, and the ratio of average spectral counts amongst AL (n = 4 CGL WT, 3 CGL KO) compared to DR (n = 5 CGL WT, 3 CGL KO) were used to determine fold-change between the two diet types. Spectral count values below the level of detection were originally automatically assigned a value of 0, however, these were changed to 0.1 for purposes of statistical analysis and plotting in log scale in volcano plots.

### Biological Function and Pathway Enrichment

The online-based web server g:Profiler (https://biit.cs.ut.ee/gprofiler/gost) (Raudvere et al., 2019) was used for functional enrichment analysis of identified sulfhydrated proteins. Gene ID’s or accession number of enriched proteins were used in the g:GOSt (Gene Group Functional Profiling) identifier tool to detect biological pathways from the KEGG database (Kanehisa et al., 2016) significantly enriched with sulfhydrated proteins. These significance values are given a threshold of p < 0.05 and were auto-calculated by the software’s proprietary g:SCS algorithm that utilizes multiple testing correction for p-values gained from pathway enrichment analysis. In setting the boundaries for significance determination, it corresponds to an experiment-wide threshold of a=0.05, with the g:SCS threshold pre-calculated for list sizes up to 1000 accession number or gene ID terms and analytically approximates a threshold t corresponding to the 5% upper quantile of randomly generated queries of that size. As per the g:Profiler description, all actual p-values resulting from the query are automatically transformed to corrected p-values by multiplying these to the ratio of the approximate threshold t and the initial experiment-wide threshold a=0.05 with consideration to the underlying gene sets annotated to terms of each organism from the KEGG database, and therefore gives a tighter threshold to significant results (Raudvere et al., 2019).

### Statistical analysis

Statistical significance and data display were generated in Microsoft Excel, GraphPad Prism, Origin, and Venny 2.1.0 software. The significant differences between the two diet groups within the same genotype in regards to H_2_S production or spectral counts for individual sulfhydrated proteins from specific tissues were analyzed by Student’s t-test, with a level of significance set to p < 0.05 and n-values equaling 4 mice for WT AL, 5 mice for WT DR, 3 mice for KO AL, and 3 mice for KO DR for the majority of the experiments and data analysis. Data was arranged in Microsoft Excel to show both relative fold-changes in average spectral counts for an individual protein as well as the t-test p-value, which can be found in Supplemental Tables 1, 2, 3, 4, 10, 11, 16, and 18. Bar-graph data in Figure 1 are displayed as means +/−SEM with n-values between 3 and 5 as indicated in the figure legend. To determine the correlation between average H_2_S production capacity and total number of sulfhydrated proteins in CGL WT and KO tissues, the arbitrary average H_2_S production capacity values determined from ImageJ IntDen function using the lead acetate/lead sulfide H_2_S assay as shown in Figure 1C (n = 3 samples/tissue) were plotted on the X-axis while the total number of sulfhydrated proteins stated in Figures 2, 4, 5, and 6 were plotted on the Y-axis (n = 9 samples/tissue for CGL WT and n = 6 samples/tissue for CGL KO). GraphPad Prism was used to fit a logarithmic trendline and the given statistics calculated via XY analysis correlation function using a nonparametric correlation (Spearman) test with a two-tailed P-value and 95% confidence interval. Origin software (https://www.originlab.com/) was used for volcano plots to display biologically- and statistically-significant differentially abundant sulfhydrated proteins in tissues from AL versus DR fed mice. The log_2_(Fold Change DR:AL) X-axis displays the average fold change in spectral counts for each identified protein while the −log_10_ Y-axis displays the calculated p-value when comparing the individual spectral count values for each identified protein in a specific tissue from AL versus DR fed mice. The non-axial red dotted vertical lines highlight the biological significance threshold of +/−2-fold change in spectral counts between DR versus AL, while the non-axial red dotted horizontal lines with asterisks highlight our statistical significance threshold that was set to p <0.05. Venny 2.1.0 software was used to generate Venn diagrams and determine differences and commonalities in sulfhydrated proteins between tissues, diets, and genotypes. Statistical analysis for pathway enrichment via g:SCS algorithm is described in the above section regarding the g:Profiler online functional enrichment analysis tool, and further specific information regarding this statistical approach can be found at the following webpage: https://biit.cs.ut.ee/gprofiler/page/docs. The calculated p-values and identities of proteins for each enriched pathway are found in Supplemental Tables 6, 7, 8, 9, 13, 14, 17, and 19.

## Supplemental Materials

### Supplemental Data File

Mass Spectrum Chromatograms from each animal and tissue analyzed

### Supplemental Tables 1-19

Data highlighted in Green are enriched under DR, data highlighted in Blue are enriched under AL, and data highlighted in Grey do not meet statistical and/or biological significance thresholds when comparing spectral counts for a specific protein between AL vs. DR. Biological significance set to ≥ 2-fold difference in average spectral counts and statistical significance set to p < 0.05 using a Student’s t-test.

Supplemental Table 1: WT Liver Sulfhydrome

Supplemental Table 2: WT Kidney Sulfhydrome

Supplemental Table 3: WT Muscle Sulfhydrome

Supplemental Table 4: WT Brain Sulfhydrome

Supplemental Table 5: Accession Numbers for the 209 Shared Sulfhydrated Proteins in WT Liver, Kidney, Muscle, and Brain

Supplemental Table 6: Pathway Enrichment in WT Liver for Sulfhydrated Proteins

Supplemental Table 7: Pathway Enrichment in WT Kidney for Sulfhydrated Proteins

Supplemental Table 8: Pathway Enrichment in WT Muscle for Sulfhydrated Proteins

Supplemental Table 9: Pathway Enrichment in WT Brain for Sulfhydrated Proteins

Supplemental Table 10: WT Heart Sulfhydrome

Supplemental Table 11: WT Plasma Sulfhydrome

Supplemental Table 12: Accession Numbers for the 78 Shared Sulfhydrated Proteins in WT Heart and Plasma

Supplemental Table 13: Pathway Enrichment in WT Heart for Sulfhydrated Proteins

Supplemental Table 14: Pathway Enrichment in WT Plasma for Sulfhydrated Proteins

Supplemental Table 15: Accession Numbers for the 28 Shared Sulfhydrated Proteins in all 6 WT Tissues

Supplemental Table 16: CGL KO Liver Sulfhydrome

Supplemental Table 17: Pathway Enrichment in CGL KO Liver for Sulfhydrated Proteins

Supplemental Table 18: CGL KO Kidney Sulfhydrome

Supplemental Table 19: Pathway Enrichment in CGL KO Kidney for Sulfhydrated Proteins

### Supplemental Figures

**Supplemental Figure 1:**
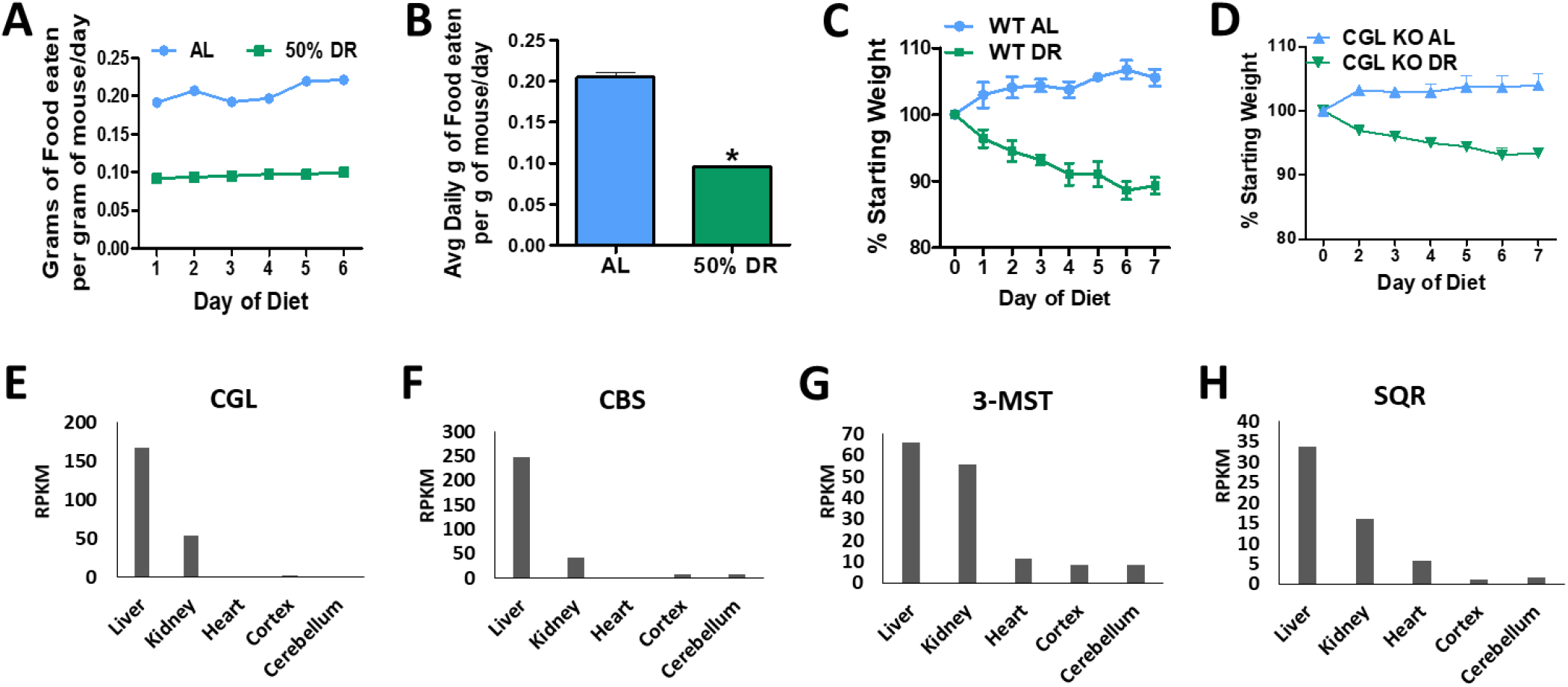
Food intake and changes in body mass as a result of 1 week 50% Dietary Restriction. (**A-B**) Daily (**A**) and average (**B**) food intake of modified AIN-93G rodent diet displayed as grams of food eaten per gram of mouse body mass per day over a 1 week period of *ad libitum* (AL) (n=7 mice/group) or 50% dietary restriction (DR) (n=8 mice/group) feeding. Asterisk indicates the statistical significance between AL versus DR; ∗p < 0.05. Error bars are ± SEM. (**C-D**) Body masses of cystathionine γ-lyase (CGL) wildtype (WT) (**C**) and total body CGL knock out (KO) (**D**) mice over the 7-day dietary intervention normalized to % initial starting weight. WT AL n=4 mice/group, WT DR n=5 mice/group, KO AL n=3 mice/group, and KO DR n=3 mice/group. Error bars are ± SEM. (**E-H**) RNA expression profiling data sets of H_2_S producing (**E-G**) and consuming (**H**) proteins generated by the Mouse ENCODE project (Yue et al., 2014) and extracted from the NCBI Mouse Gene Database. CGL = cystathionine γ-lyase, CBS = cystathionine β-synthase, 3-MST = 3-mercaptopyruvate sulfurtransferase, SQR = sulfide:quinone oxidoreductase, RPKM = reads per kilobase of transcript, per million mapped reads. See also Figure 1.

**Supplemental Figure 2:**
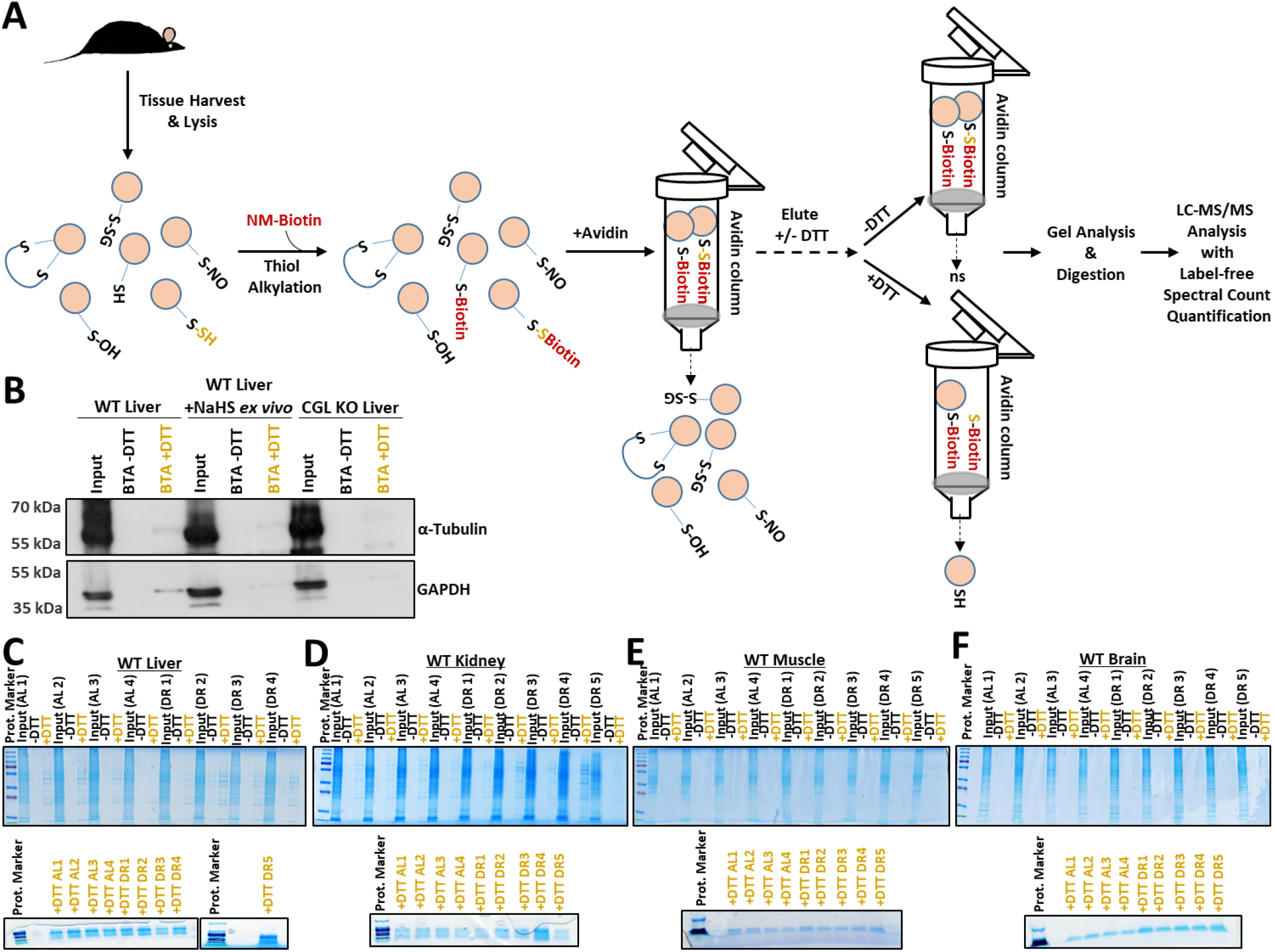
Modified biotin thiol assay (BTA) to isolate and detect differentially sulfhydrated proteins in mouse tissues after one week of dietary restriction. (**A**) Schematic presentation of the biotin-thiol-assay (BTA) adapted from the method developed by Gao, et al (Gao et al., 2015) to isolate, detect, and quantify sulfhydrated proteins in mouse tissue. Tissues are first homogenized and lysed, the protein concentrations equally normalized, and then lysates subjected to thiol- and persulfide-specific (shown in yellow text) alkylation and binding to NM-biotin. Subsequent addition of biotin bound and unbound proteins into an avidin column isolates those bound to biotin, with several buffered wash steps and elution without dithiothreitol (−DTT) to remove non-specific non-sulfhydrated proteins attached to the column. A final buffered elution with DTT (+DTT) reduces and cleaves the disulfide bond between the cysteine-attached sulfur and the persulfide (shown in yellow text) bound to the biotin/avidin column, thus eluting the sulfhydrated proteins for downstream gel-based analysis (**B-F**) as well as mass spectrometry based LC-MS/MS analysis coupled with label-free spectral counting to measure relative sulfhydrated protein abundance between different samples as a function of diet and/or genotype. (**B**) Validation of the BTA via Western blot analysis of liver lysates treated *ex vivo* −/+ sodium hydrosulfide (NaHS) from CGL WT and KO mice for the known sulfhydrated proteins α-tubulin and GAPDH following application of the BTA. (**C-F**) SDS-PAGE gel electrophoresis followed by colloidal staining on liver (**C**), kidney (**D**), muscle (**E**), and brain (**F**) protein lysate input loading controls as well as −DTT and +DTT eluates from the BTA derived from *ad libitum* (AL, n=4) or 50% dietary restriction (DR; n=5) fed cystathionine γ-lyase (CGL) WT mice (top gels). Bottom SDS-PAGE gel images are from the same +DTT eluates shown in the upper gels but run for a short period of time; 13-15 minutes, prior to excising the entire protein lane to ensure all of that tissue’s sulfhydrated proteins across the full spectrum of protein masses are accounted for in the subsequent downstream LC-MS/MS proteomics analysis. Yellow font for +DTT indicates the experimental conditions in which the sulfhydrated proteins were eluted and isolated for future gel and LC-MS/MS proteomics analysis. See also Figure 2.

**Supplemental Figure 3:**
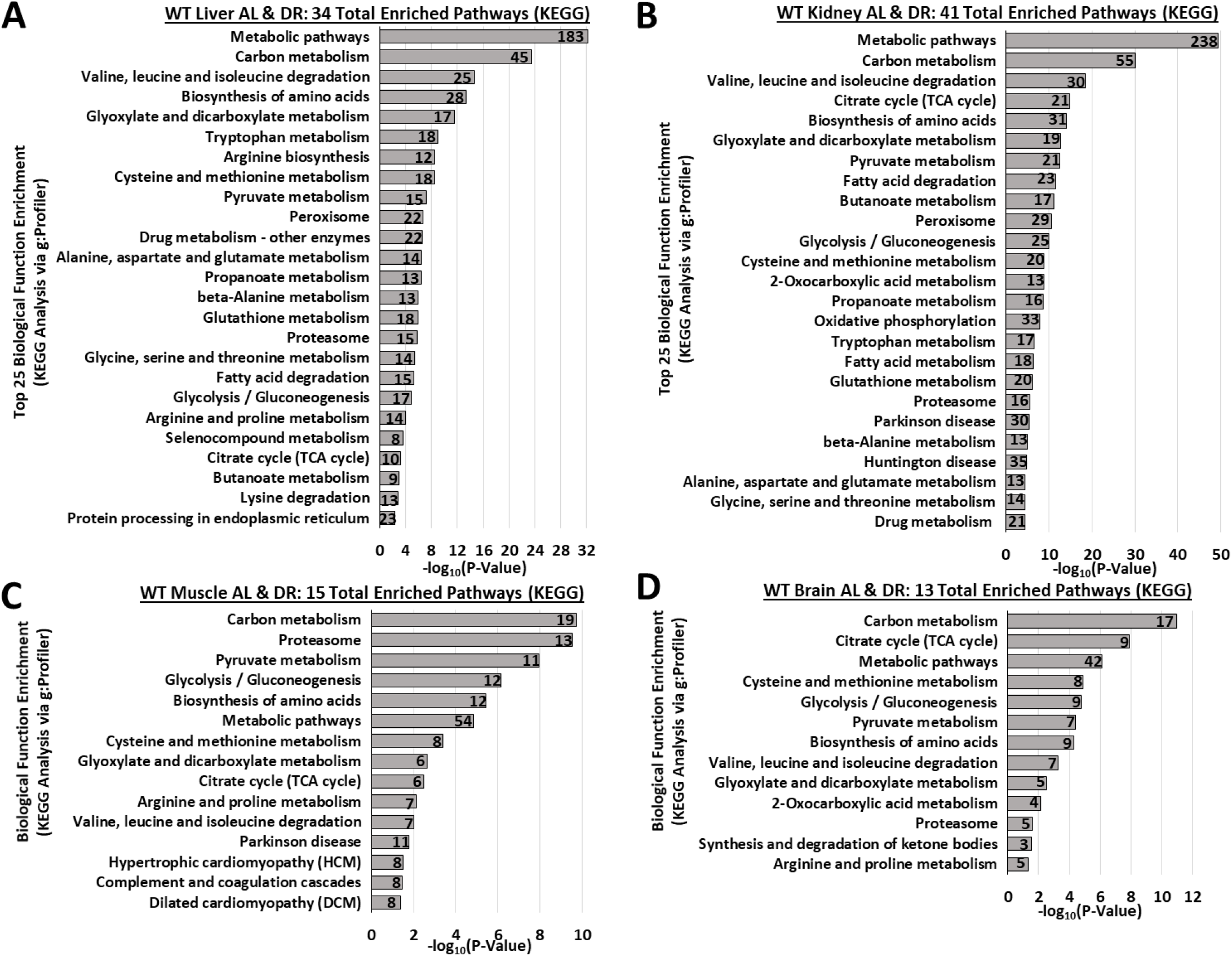
Biological function and pathway enrichment for sulfhydrated proteins not significantly changed by diet. (**A-D**) KEGG biological function and pathway enrichment via g:Profiler analysis of sulfhydrated proteins whose relative abundance was not changed under AL or DR feeding in liver (**A**), kidney (**B**), muscle (**C**), and brain (**D**). The number inside the bar indicates the quantity of sulfhydrated proteins involved in that specific pathway. n = 9 mice in total; 4 in the AL group and 5 in the DR group. Statistical significance for pathway enrichment plotted as −log_10_ (P-Value) and obtained via the g:Profiler g:SCS algorithm for KEGG database analysis. See also Figure 3.

**Supplemental Figure 4:**
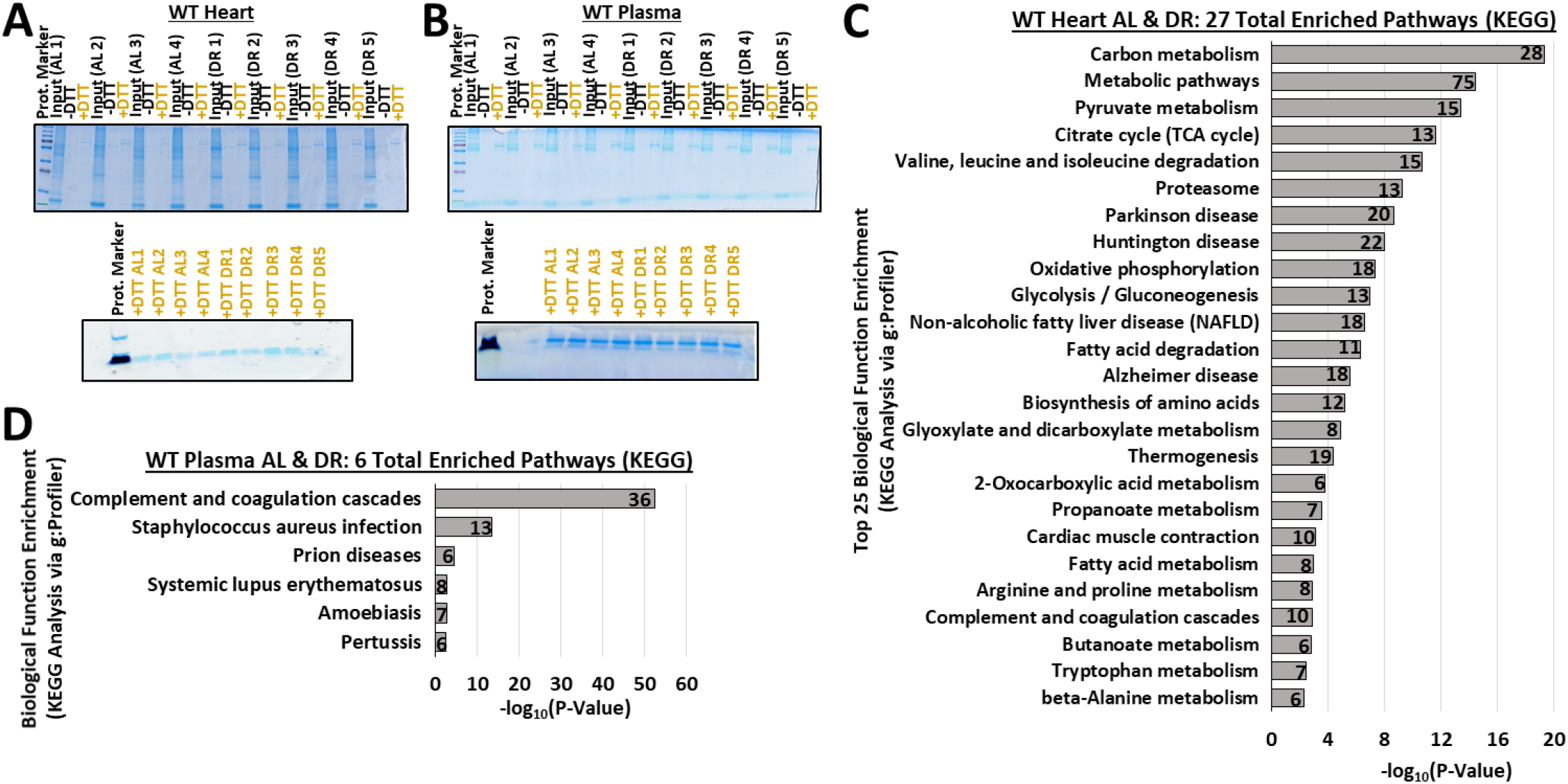
Sulfhydration analysis and pathway enrichment in heart and plasma. (**A-B**). SDS-PAGE gel electrophoresis followed by colloidal staining on heart (**A**), and plasma (**B**) protein lysate input loading controls as well as −DTT and +DTT eluates from the BTA derived from AL (n=4) or DR (n=5) fed cystathionine γ-lyase (CGL) WT mice (top gels). Bottom SDS-PAGE gel images are from the same +DTT eluates shown in the upper gels but run briefly prior to excising the entire protein lane to ensure all of that tissue’s sulfhydrated proteins across the full spectrum of masses are accounted for in the subsequent downstream LC-MS/MS proteomics analysis. Yellow font for +DTT indicates the experimental conditions in which the sulfhydrated proteins were eluted and isolated for future gel and LC-MS/MS proteomics analysis. (**C-D**) KEGG biological function and pathway enrichment via g:Profiler analysis of sulfhydrated proteins whose relative abundance was not changed under AL or DR feeding in heart (**C**), and plasma (**D**). The number inside the bar indicates the number of sulfhydrated proteins involved in that specific pathway. n = 9 mice in total; 4 in the AL group and 5 in the DR group. Statistical significance for pathway enrichment plotted as −log_10_ (P-Value) and obtained via the g:Profiler g:SCS algorithm for KEGG database analysis. See also Figure 4.

**Supplemental Figure 5:**
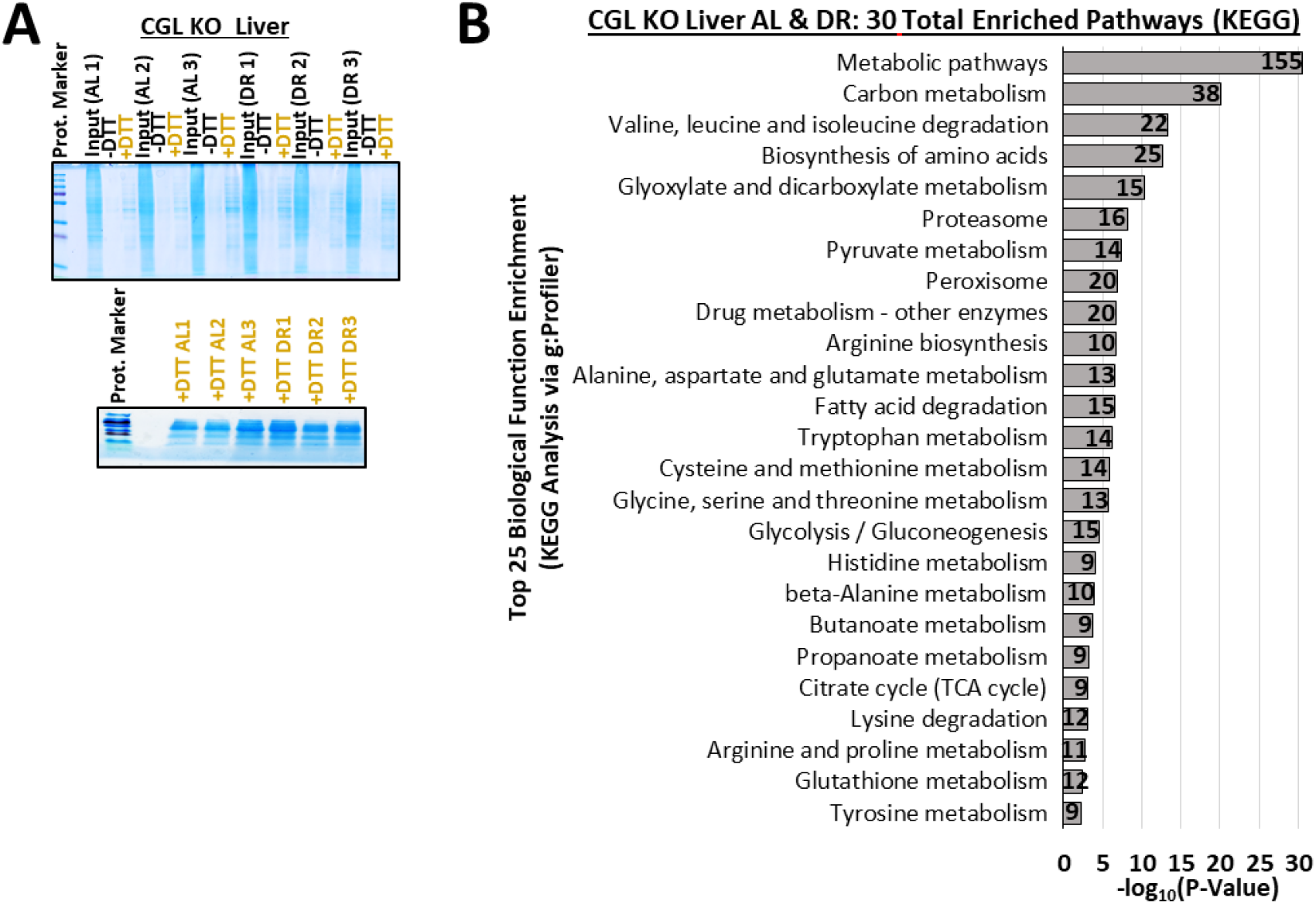
Sulfhydration analysis and pathway enrichment in CGL KO liver. **(A)** SDS-PAGE gel electrophoresis followed by colloidal staining on liver protein lysate input loading controls as well as −DTT and +DTT eluates from the BTA derived from AL (n=3) or DR (n=3) fed cystathionine γ-lyase (CGL) KO mice (top gels). Bottom SDS-PAGE gel images are from the same +DTT eluates shown in the upper gels but run briefly prior to excising the entire protein lane to ensure all of the liver’s sulfhydrated proteins across the full spectrum of masses are accounted for in the subsequent downstream LC-MS/MS proteomics analysis. Yellow font for +DTT indicates the experimental conditions in which the sulfhydrated proteins were eluted and isolated for future gel and LC-MS/MS proteomics analysis. (**B**) KEGG biological function and pathway enrichment via g:Profiler analysis of sulfhydrated proteins whose relative abundance was not changed under AL or DR feeding in liver of CGL KO mice. The number inside the bar indicates the number of sulfhydrated proteins involved in that specific pathway. n = 6 mice in total; 3 in the AL group and 3 in the DR group. Statistical significance for pathway enrichment plotted as −log_10_ (P-Value) and obtained via the g:Profiler g:SCS algorithm for KEGG database analysis. See also Figure 5.

**Supplemental Figure 6:**
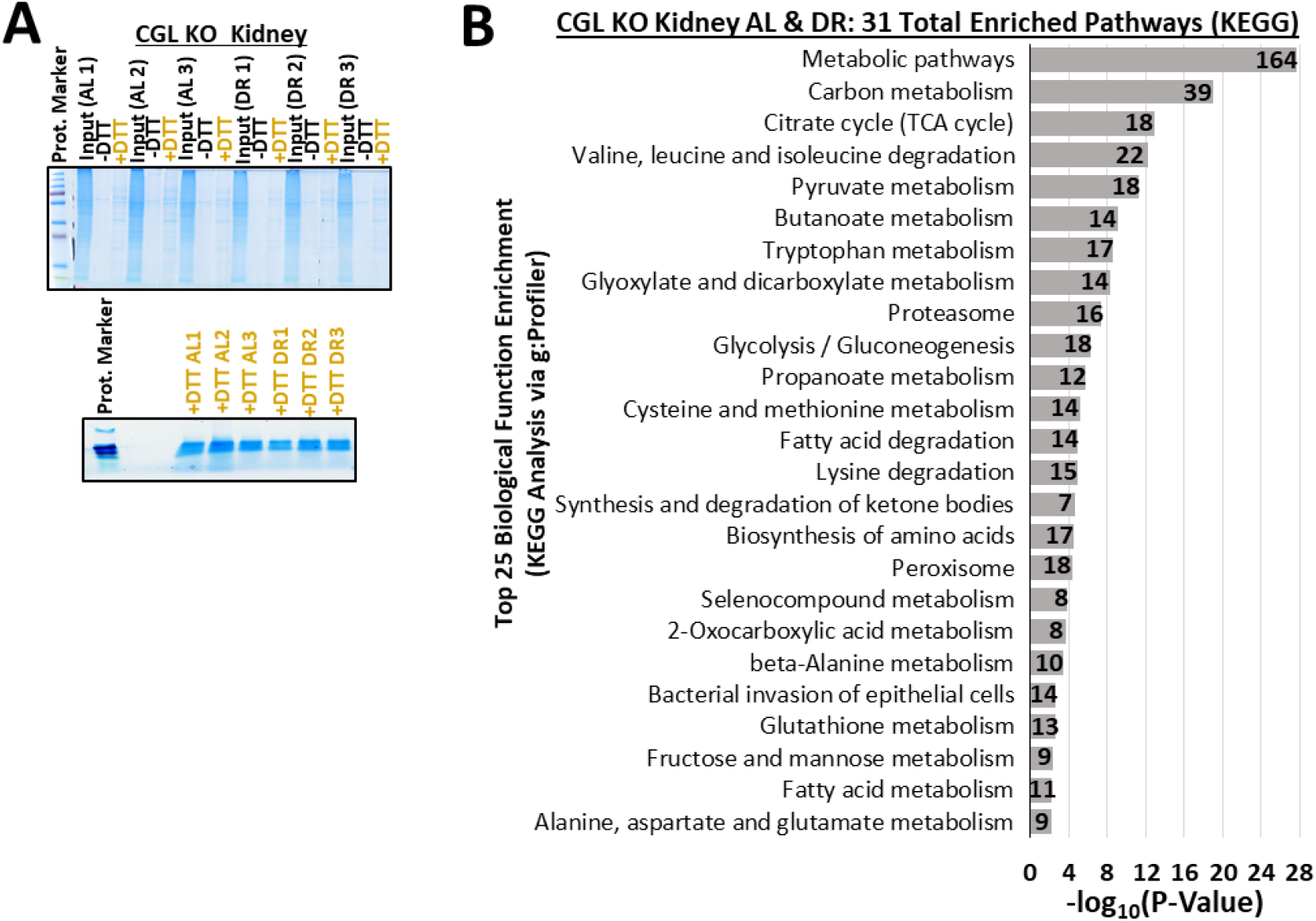
Sulfhydration analysis and pathway enrichment in CGL KO kidney. **(A)** SDS-PAGE gel electrophoresis followed by colloidal staining on kidney protein lysate input loading controls as well as −DTT and +DTT eluates from the BTA derived from AL (n=3) or DR (n=3) fed cystathionine γ-lyase (CGL) KO mice (top gels). Bottom SDS-PAGE gel images are from the same +DTT eluates shown in the upper gels but run briefly prior to excising the entire protein lane to ensure all of the kidney’s sulfhydrated proteins across the full spectrum of masses are accounted for in the subsequent downstream LC-MS/MS proteomics analysis. Yellow font for +DTT indicates the experimental conditions in which the sulfhydrated proteins were eluted and isolated for future gel and LC-MS/MS proteomics analysis. (**B**) KEGG biological function and pathway enrichment via g:Profiler analysis of sulfhydrated proteins whose relative abundance was not changed under AL or DR feeding in kidney of CGL KO mice. The number inside the bar indicates the number of sulfhydrated proteins involved in that specific pathway. n = 6 mice in total; 3 in the AL group and 3 in the DR group. Statistical significance for pathway enrichment plotted as −log_10_ (P-Value) and obtained via the g:Profiler g:SCS algorithm for KEGG database analysis. See also Figure 6.

**Supplemental Figure 7:**
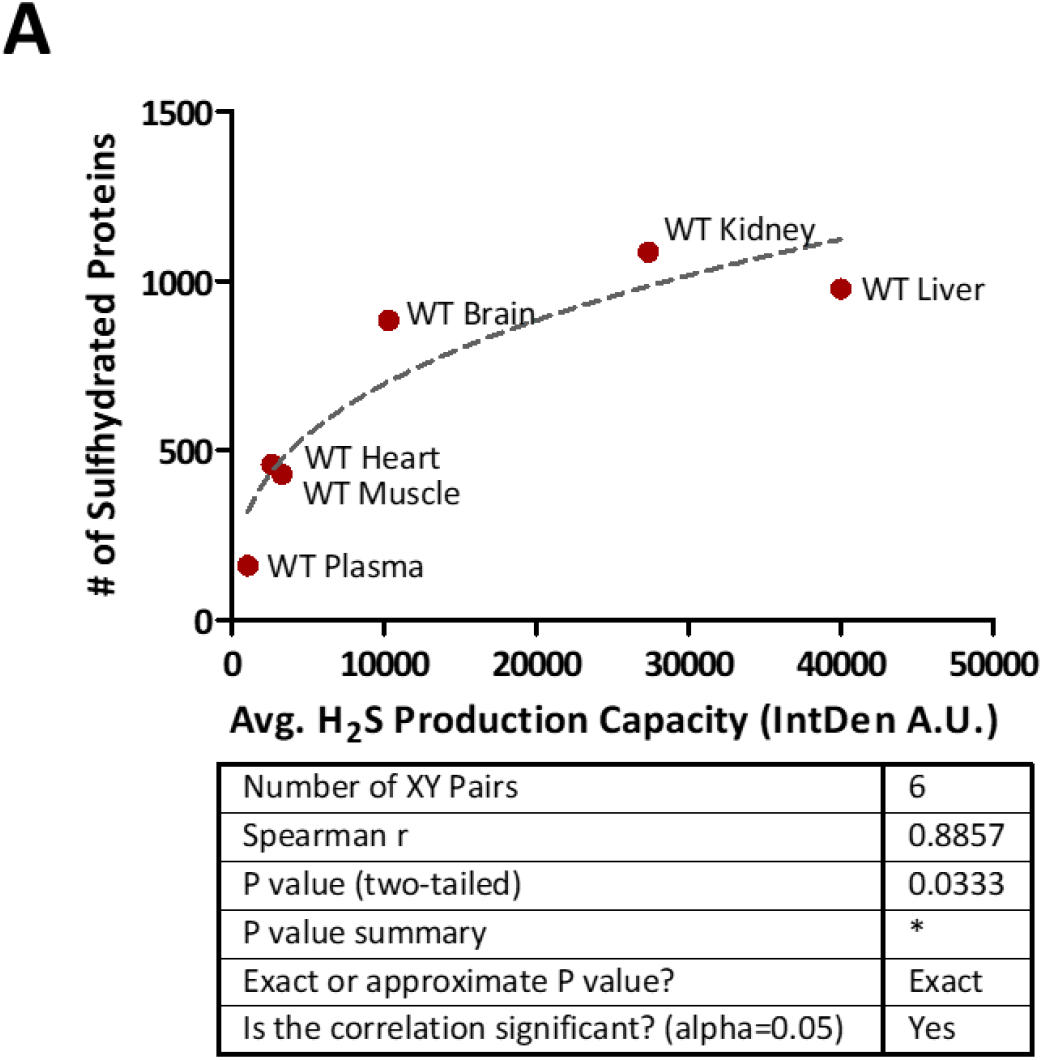
Positive correlation between H_2_S production capacity and number of sulfhydrated proteins identified in CGL WT tissues. Correlation between average H_2_S production capacity and total number of sulfhydrated proteins in CGL WT and KO tissues. Arbitrary average H_2_S production capacity values determined from ImageJ IntDen function using the lead acetate/lead sulfide H_2_S assay as shown in Figure 1C (n = 3 samples/tissue) were plotted on the X-axis while the total number of sulfhydrated proteins stated in Figures 2 and were plotted on the Y-axis (n = 9 samples/tissue). GraphPad Prism was used to fit the trendline and the given statistics calculated via XY analysis nonparametric correlation (Spearman) test with a two-tailed P- value and 95% confidence interval. See also Figure 7.

