## Supplemental Data File 1 for "Dietary restriction transforms the protein sulfhydrome in a tissue-specific and cystathionine γ-lyase-dependent manner"

### WT Liver Chromatograms from Label Free Quantification

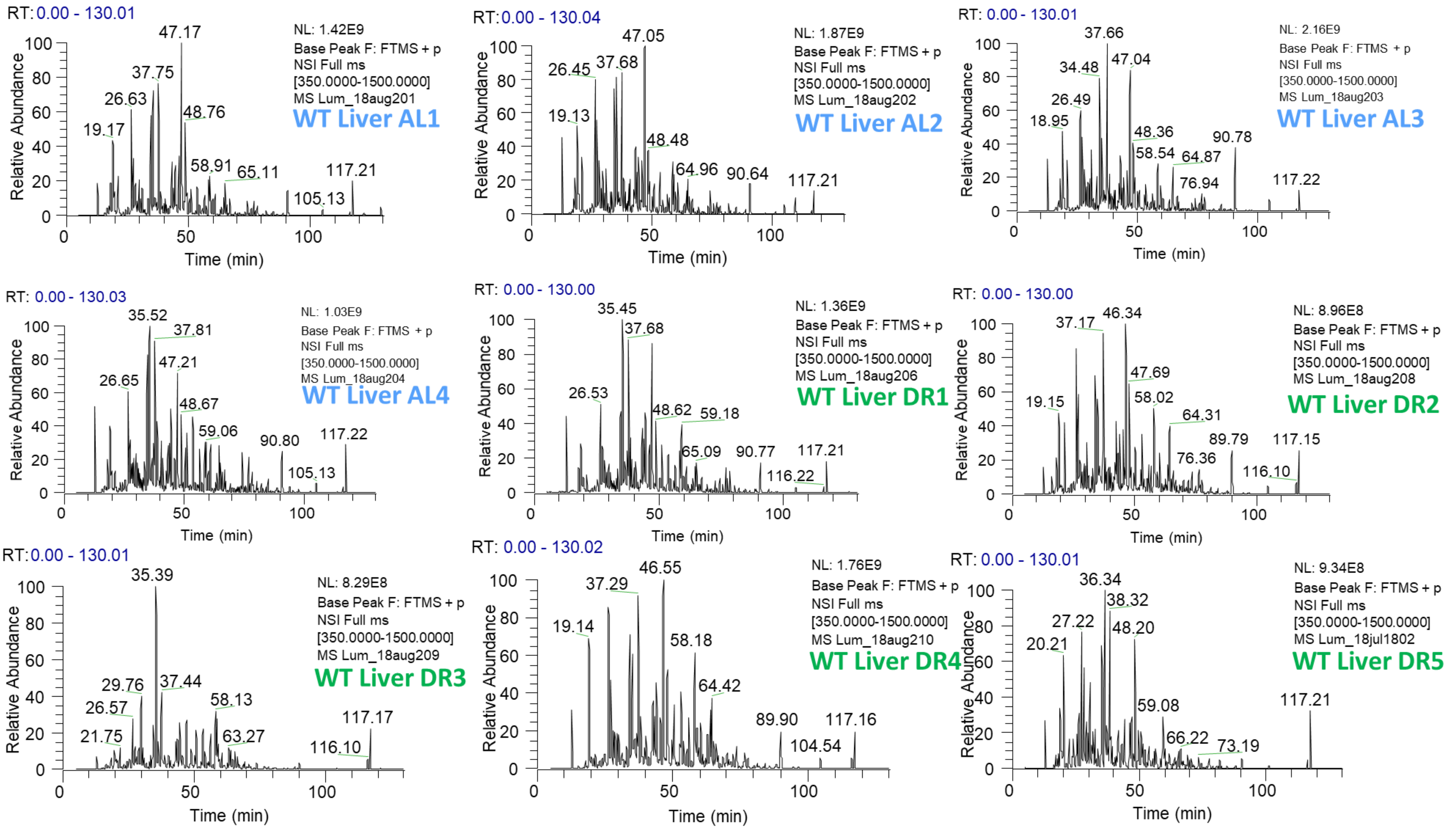

### WT Kidney Chromatograms from Label Free Quantification

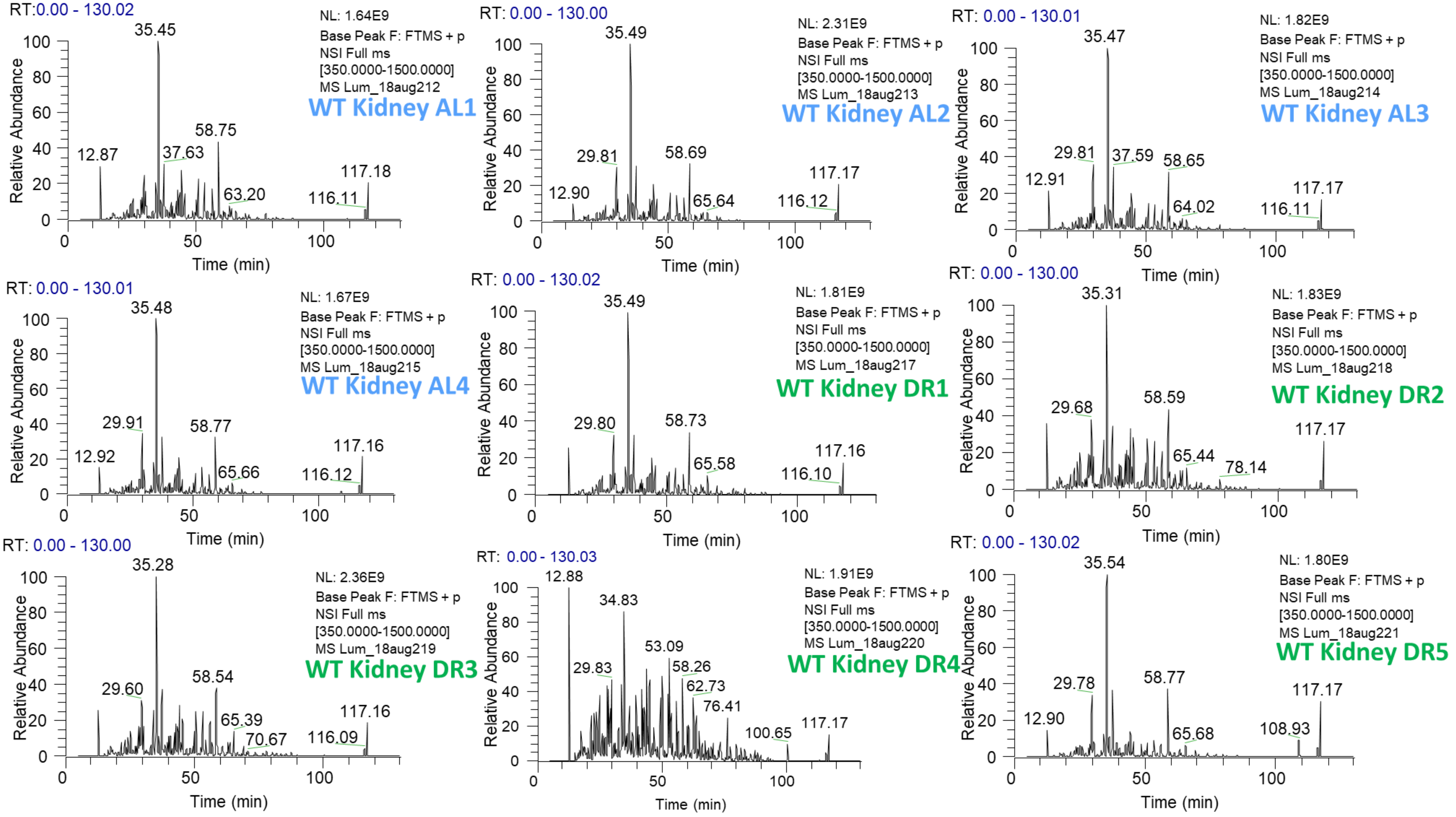

### WT Muscle Chromatograms from Label Free Quantification

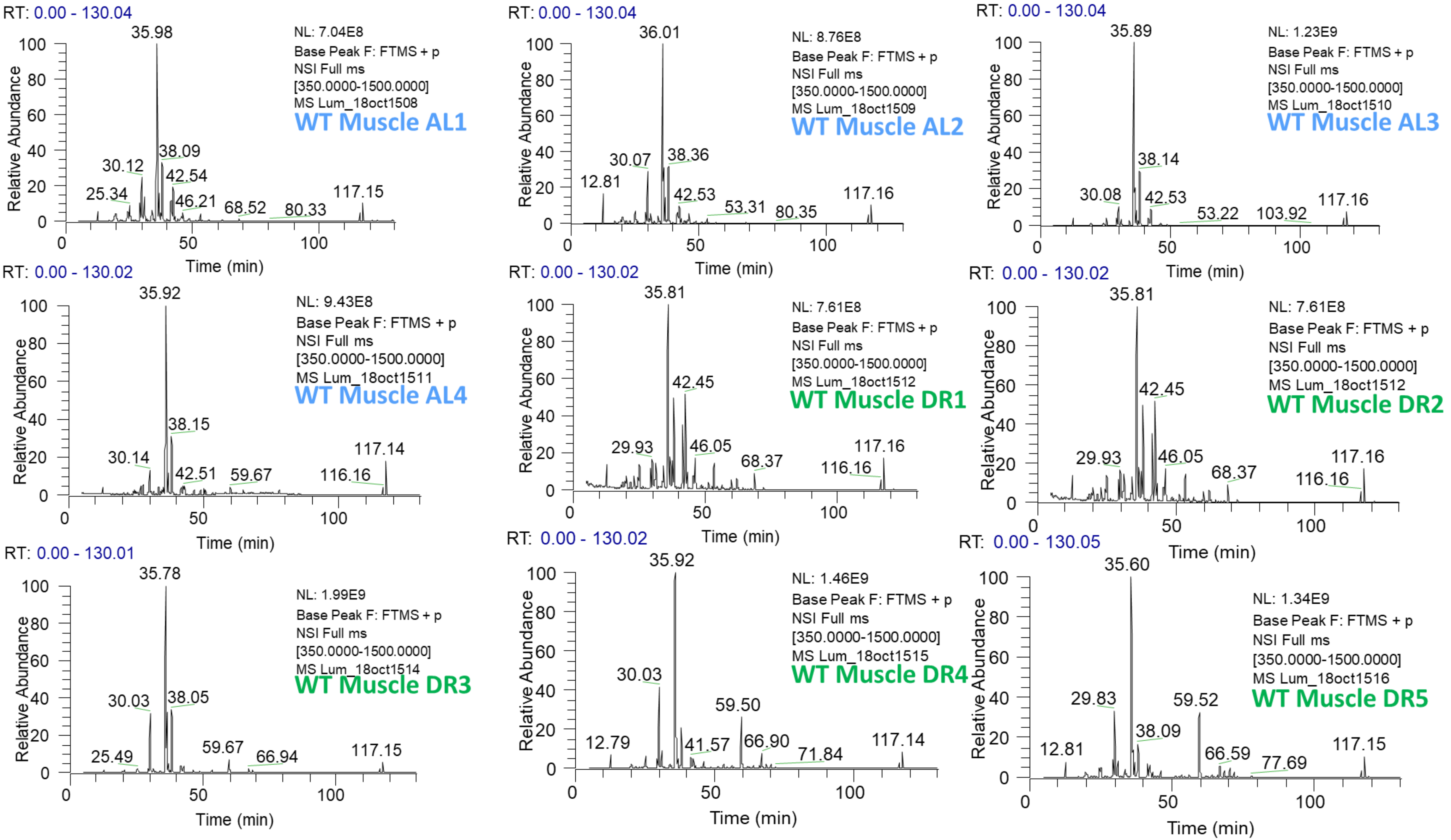

### WT Brain Chromatograms from Label Free Quantification

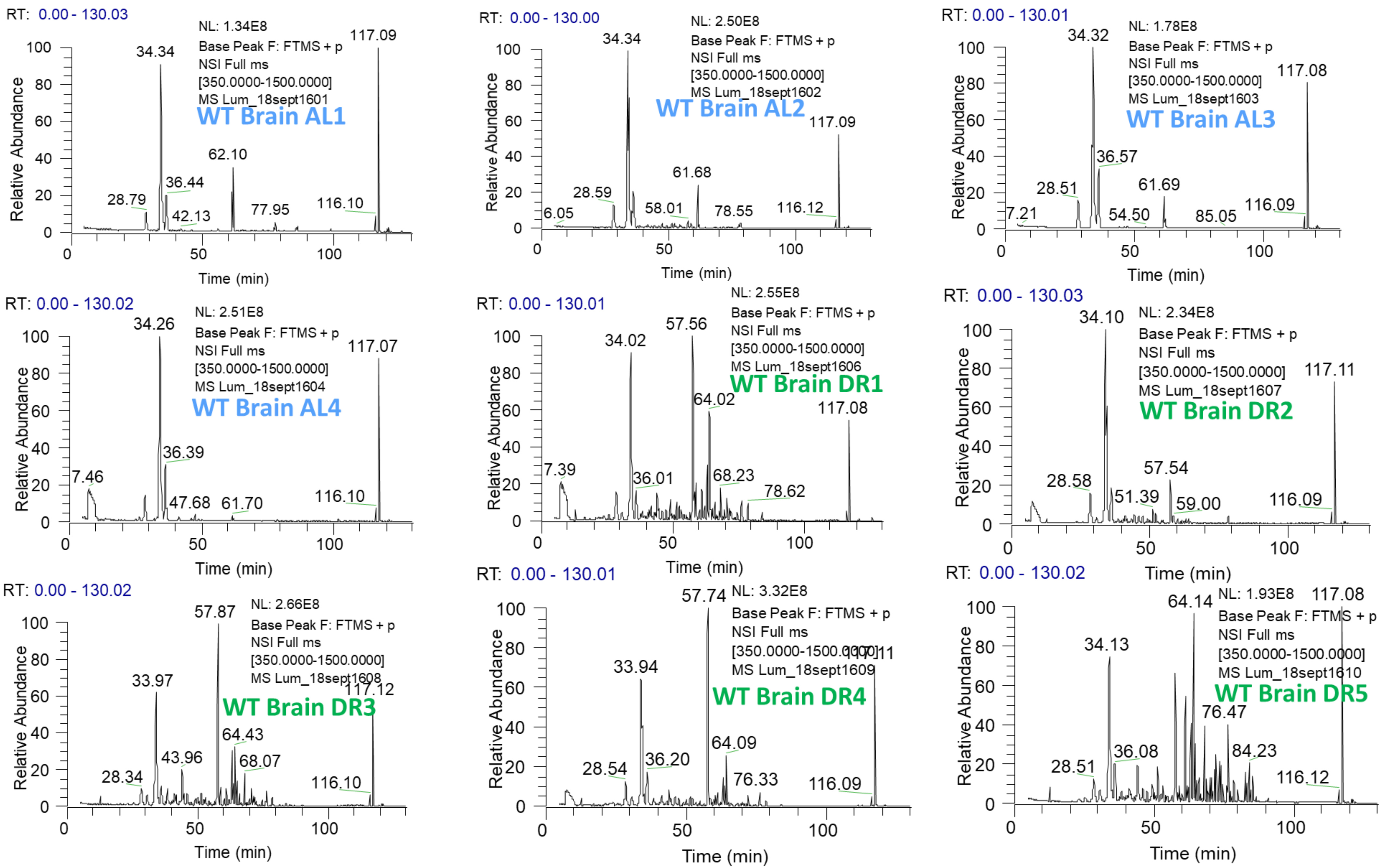

### WT Heart Chromatograms from Label Free Quantification

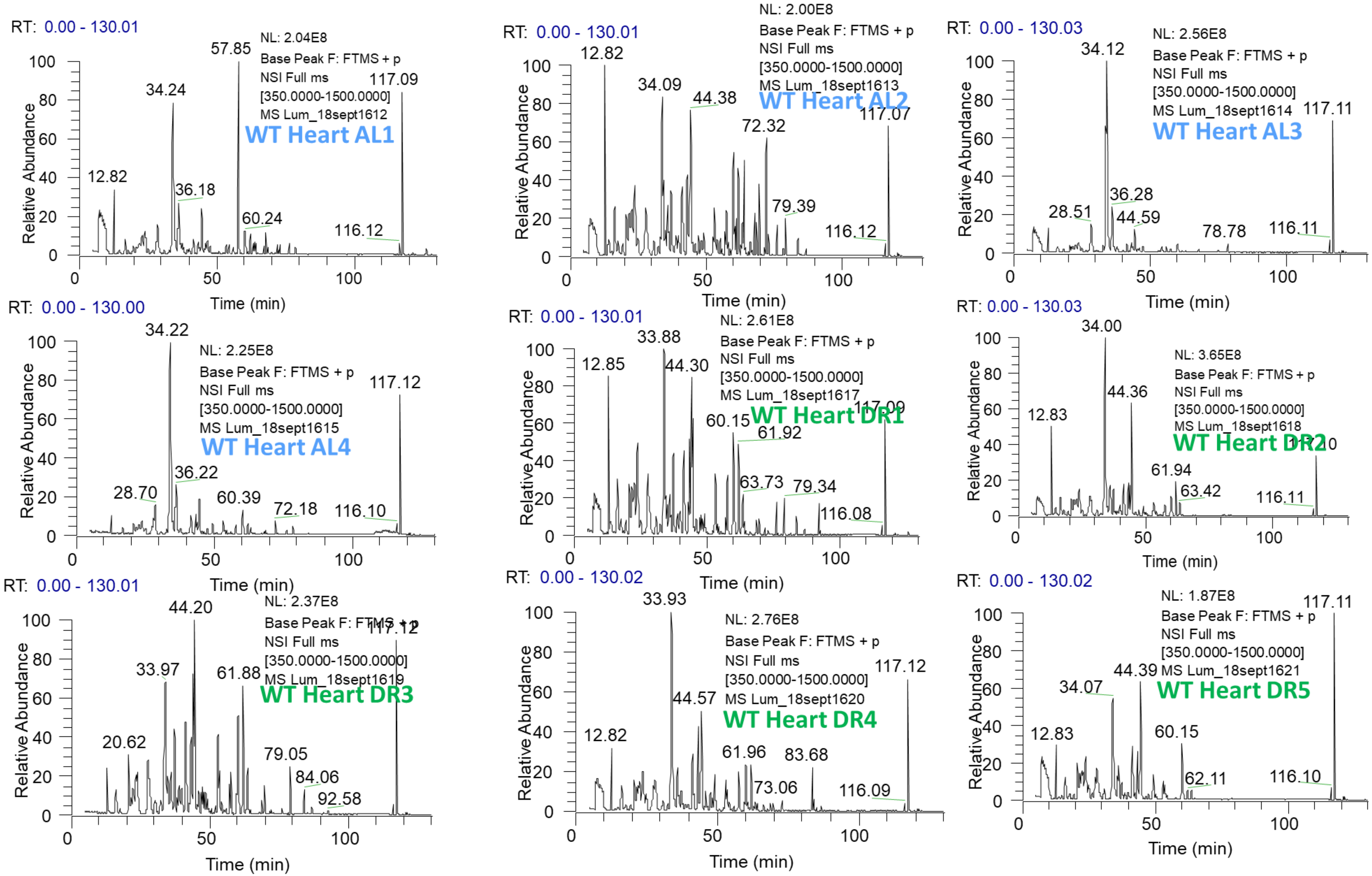

### WT Serum Chromatograms from Label Free Quantification

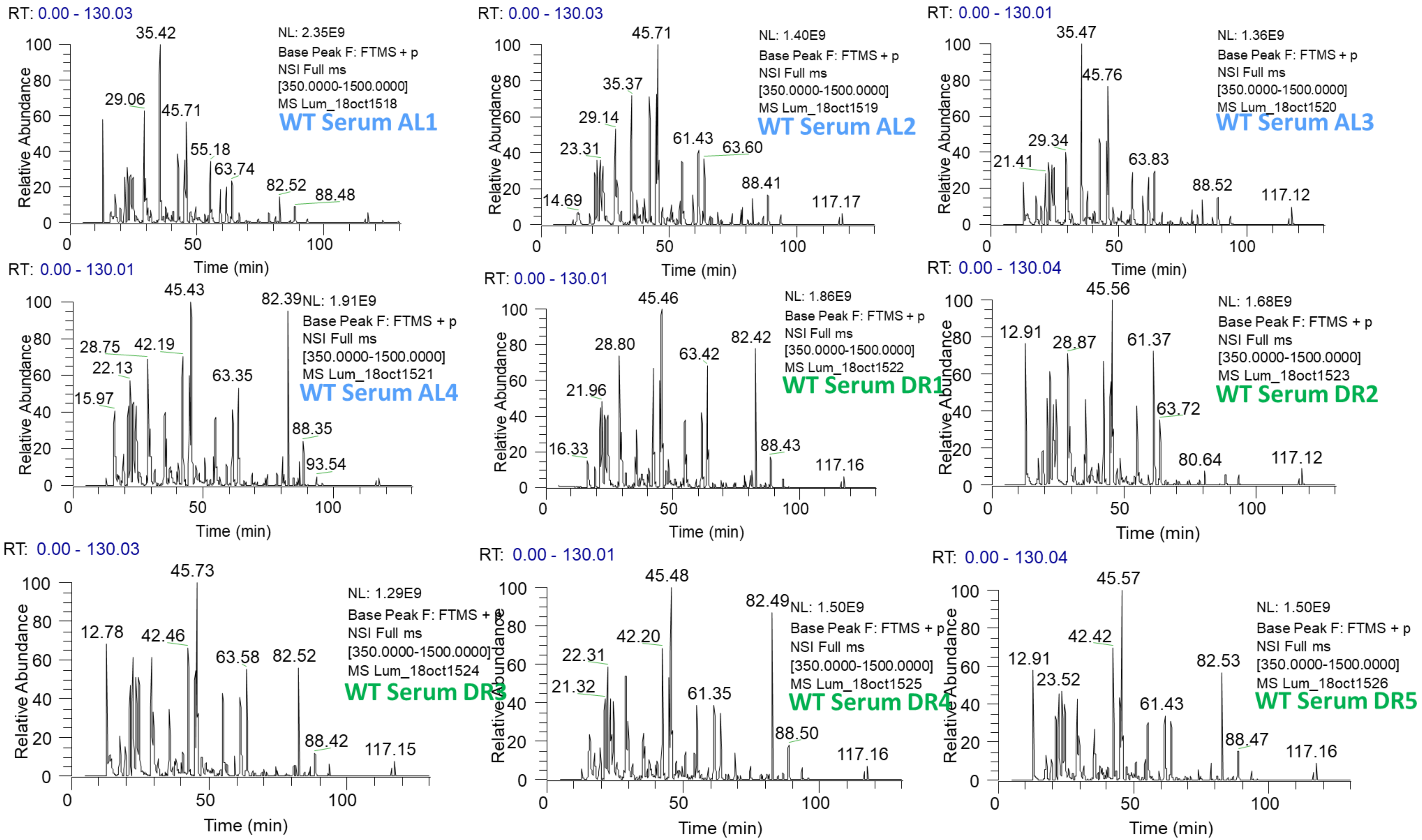

### CGL KO Liver Chromatograms from Label Free Quantification

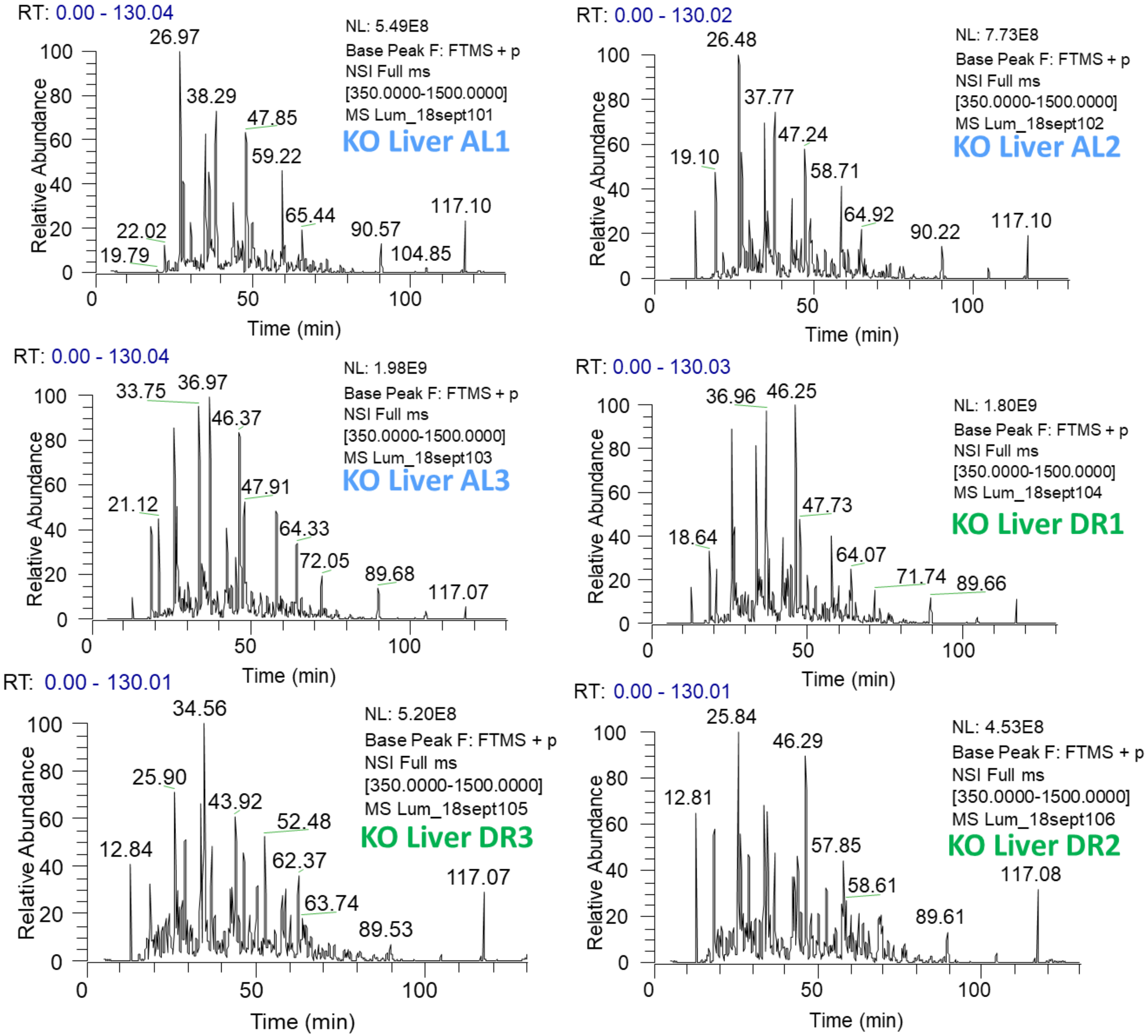

### CGL KO Kidney Chromatograms from Label Free Quantification

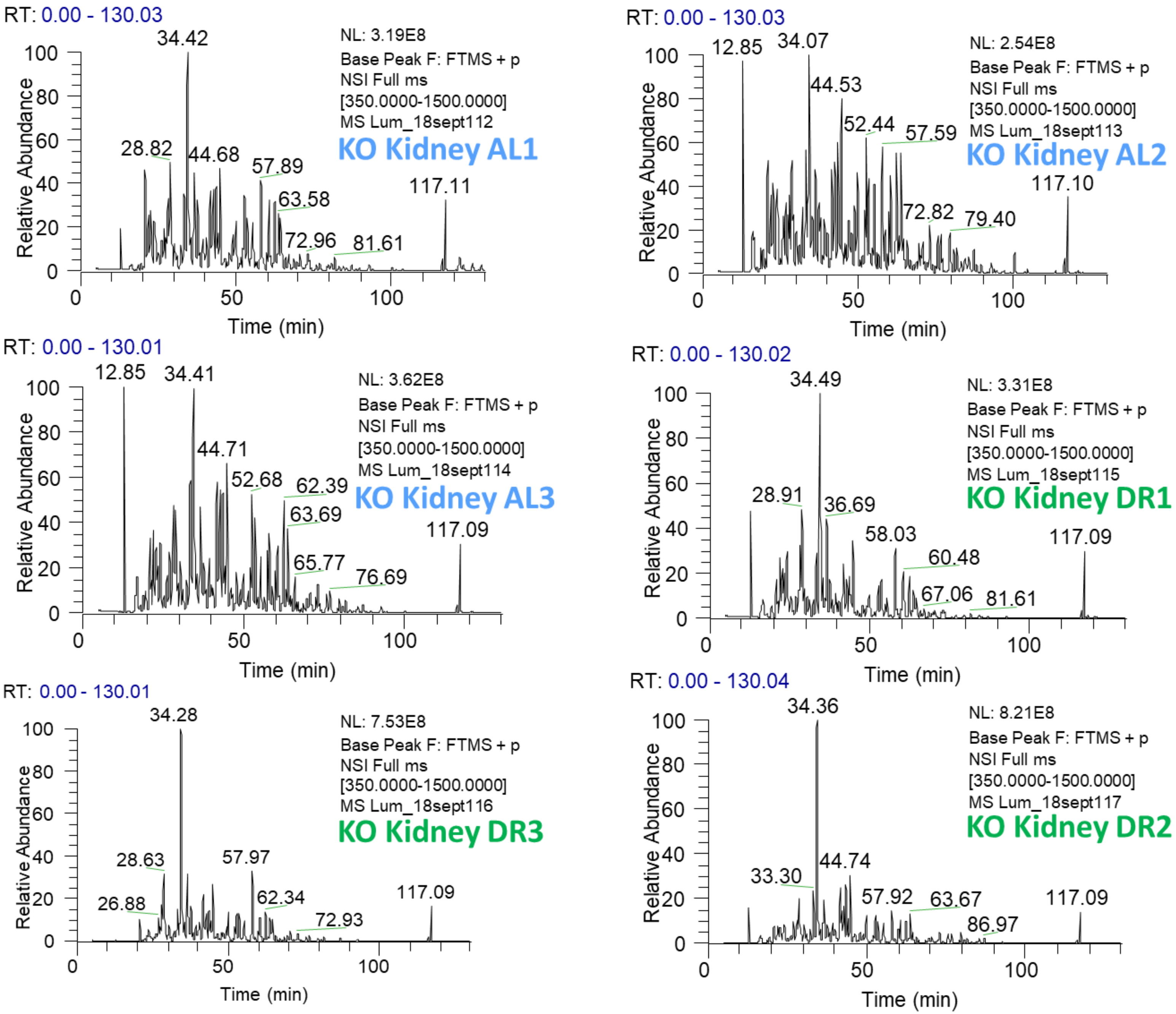
