## Supplemental Table 1 for "Dietary restriction transforms the protein sulfhydrome in a tissue-specific and cystathionine γ-lyase-dependent manner"

| Supplemental Table 1: Dietary Impact on the Liver Sulfhydrome | | | | | | |
|---|---|---|---|---|---|---|
| Protein Name | Accession Number | Alternate ID | Molecular Weight | Cysteine Residues | DR/AL Spectral Count Ratio | P-value |
| O-acetyl-ADP-ribose deacetylase MACROD1 | Q922B1 | Macrod1 | 35 kDa | 10 C | 18.000 | 0.00514 |
| Pyruvate kinase PKLR | P53657 | Pklr | 62 kDa | 6 C | 14.154 | 0.01512 |
| Uroporphyrinogen decarboxylase | P70697 | Urod | 41 kDa | 5 C | 12.200 | 0.02824 |
| 3-hydroxyacyl-CoA dehydrogenase type-2 | O08756 | Hsd17b10 | 27 kDa | 2 C | 9.846 | 0.04406 |
| Prelamin-A/C | P48678 | Lmna | 74 kDa | 5 C | 7.275 | 0.04266 |
| Polyadenylate-binding protein 2 | Q8CCS6 (+1) | Pabpn1 | 32 kDa | 2 C | 6.769 | 0.01549 |
| Acyl-CoA synthetase family member 2, mitochondrial | Q8VCW8 | Acsf2 | 68 kDa | 12 C | 5.161 | 0.00895 |
| Pro-cathepsin H | P49935 | Ctsh | 37 kDa | 10 C | 5.000 | 0.00903 |
| DnaJ homolog subfamily C member 3 | Q91YW3 | Dnajc3 | 57 kDa | 8 C | 4.923 | 0.00726 |
| Pyridoxal phosphate homeostasis protein | Q9Z2Y8 | Prosc | 30 kDa | 3 C | 4.645 | 0.03341 |
| Endoplasmin | P08113 | Hsp90b1 | 92 kDa | 5 C | 4.000 | 0.01106 |
| Polymeric immunoglobulin receptor | O70570 | Pigr | 85 kDa | 23 C | 3.613 | 0.00027 |
| Acyl-coenzyme A synthetase ACSM1, mitochondrial | Q91VA0 | Acsm1 | 65 kDa | 14 C | 3.467 | 0.0002 |
| Prostaglandin E synthase 3 | Q9R0Q7 | Ptges3 | 19 kDa | 5 C | 3.400 | 0.04556 |
| ES1 protein homolog, mitochondrial | Q9D172 | D10Jhu81e | 28 kDa | 6 C | 3.317 | 0.03557 |
| Peptidyl-prolyl cis-trans isomerase FKBP2 | P45878 | Fkbp2 | 15 kDa | 3 C | 3.097 | 0.03249 |
| Phospholipase B-like 1 | Q8VCI0 | Plbd1 | 63 kDa | 7 C | 2.667 | 0.01942 |
| Hydroxymethylglutaryl-CoA synthase, mitochondrial | P54869 | Hmgcs2 | 57 kDa | 9 C | 2.560 | 0.00104 |
| Carboxylesterase 1C | P23953 | Ces1c | 61 kDa | 5 C | 2.560 | 0.02656 |
| NADH dehydrogenase [ubiquinone] 1 alpha subcomplex subunit 8 | Q9DCJ5 | Ndufa8 | 20 kDa | 8 C | 2.537 | 0.02746 |
| NADH dehydrogenase [ubiquinone] iron-sulfur protein 6, mitochondrial | P52503 | Ndufs6 | 13 kDa | 3 C | 2.400 | 0.00743 |
| Membrane-associated progesterone receptor component 1 | O55022 | Pgrmc1 | 22 kDa | 2 C | 2.400 | 0.04315 |
| Complement factor I | Q61129 | Cfi | 67 kDa | 40 C | 2.400 | 0.046 |
| GrpE protein homolog 1, mitochondrial | Q99LP6 | Grpel1 | 24 kDa | 4 C | 2.323 | 0.01133 |
| Enoyl-CoA delta isomerase 1, mitochondrial | P42125 | Eci1 | 32 kDa | 5 C | 2.286 | 0.00106 |
| Sorting nexin-3 | O70492 | Snx3 | 19 kDa | 1 C | 2.160 | 0.01359 |
| Long-chain specific acyl-CoA dehydrogenase, mitochondrial | P51174 | Acadl | 48 kDa | 7 C | 2.142 | 0.01708 |
| Enoyl-CoA hydratase domain-containing protein 2, mitochondrial | Q3TLP5 | Echdc2 | 32 kDa | 6 C | 2.133 | 0.00081 |
| D-dopachrome decarboxylase | O35215 | Ddt | 13 kDa | 2 C | 2.059 | 0.04739 |
| ELAV-like protein 1 | P70372 | Elavl1 | 36 kDa | 3 C | 2.039 | 0.03002 |
| Alcohol dehydrogenase [NADP(+)] | Q9JII6 | Akr1a1 | 37 kDa | 4 C | 32.400 | 0.08295 |
| Long-chain-fatty-acid--CoA ligase 1 | P41216 | Acsl1 | 78 kDa | 18 C | 24.400 | 0.07944 |
| 60S acidic ribosomal protein P0 | P14869 | Rplp0 | 34 kDa | 3 C | 18.400 | 0.13087 |
| Citrate lyase subunit beta-like protein, mitochondrial | Q8R4N0 | Clybl | 38 kDa | 6 C | 18.200 | 0.07401 |
| Nucleoporin SEH1 | Q8R2U0 (+1) | Seh1l | 40 kDa | 10 C | 18.200 | 0.1953 |
| Prohibitin | P67778 | Phb | 30 kDa | 1 C | 16.600 | 0.26887 |
| Acyl-CoA-binding domain-containing protein 5 | Q5XG73 | Acbd5 | 57 kDa | 9 C | 16.400 | 0.13044 |
| Isoform 2 of Peptidyl-prolyl cis-trans isomerase H | Q9D868-2 | Ppih | 19 kDa | 5 C | 16.000 | 0.06328 |
| Tetratricopeptide repeat protein 36 | Q8VBW8 | Ttc36 | 20 kDa | 1 C | 14.600 | 0.33097 |
| 40S ribosomal protein S5 | P97461 | Rps5 | 23 kDa | 3 C | 14.400 | 0.07963 |
| Cytochrome c oxidase subunit 6B1 | P56391 | Cox6b1 | 10 kDa | 4 C | 14.400 | 0.14941 |
| Clusterin | Q06890 | Clu | 52 kDa | 11 C | 12.600 | 0.31777 |
| Myosin light chain 3 | P09542 | Myl3 | 22 kDa | 2 C | 12.600 | 0.31777 |
| Cytochrome c1, heme protein, mitochondrial | Q9D0M3 | Cyc1 | 35 kDa | 14 C | 12.600 | 0.19301 |
| Vesicular integral-membrane protein VIP36 | Q9DBH5 | Lman2 | 40 kDa | 6 C | 12.600 | 0.19301 |
| Isoform 2 of Basigin | P18572-2 | Bsg | 30 kDa | 8 C | 12.600 | 0.31777 |
| 6-phosphogluconolactonase | Q9CQ60 | Pgls | 27 kDa | 4 C | 12.400 | 0.06758 |
| Very long-chain specific acyl-CoA dehydrogenase, mitochondrial | P50544 | Acadvl | 71 kDa | 7 C | 12.400 | 0.11712 |
| Myosin-6 | Q02566 | Myh6 | 224 kDa | 14 C | 10.800 | 0.40708 |
| Acyl-CoA dehydrogenase family member 10 | Q8K370 | Acad10 | 119 kDa | 16 C | 10.600 | 0.20691 |
| GTP-binding protein SAR1b | Q9CQC9 | Sar1b | 22 kDa | 2 C | 10.400 | 0.09207 |
| 60S ribosomal protein L11 | Q9CXW4 | Rpl11 | 20 kDa | 4 C | 10.400 | 0.09207 |
| Ubiquinone biosynthesis protein COQ9, mitochondrial | Q8K1Z0 | Coq9 | 35 kDa | 3 C | 10.400 | 0.09207 |
| Vesicle-associated membrane protein-associated protein B | Q9QY76 | Vapb | 27 kDa | 4 C | 10.400 | 0.16156 |
| Annexin A5 | P48036 | Anxa5 | 36 kDa | 1 C | 9.292 | 0.05787 |
| Glutathione S-transferase kappa 1 | Q9DCM2 | Gstk1 | 26 kDa | 2 C | 8.800 | 0.40708 |
| Inter alpha-trypsin inhibitor, heavy chain 4 | A6X935 (+1) | Itih4 | 105 kDa | 3 C | 8.800 | 0.40708 |
| Polymerase delta-interacting protein 3 | Q8BG81 | Poldip3 | 46 kDa | 4 C | 8.800 | 0.40708 |
| Synaptosomal-associated protein 47 | Q8R570 | Snap47 | 47 kDa | 4 C | 8.800 | 0.40708 |
| Coiled-coil-helix-coiled-coil-helix domain-containing protein 2 | Q9D1L0 | Chchd2 | 16 kDa | 4 C | 8.600 | 0.19301 |
| Carnitine O-palmitoyltransferase 2, mitochondrial | P52825 | Cpt2 | 74 kDa | 10 C | 8.600 | 0.27239 |
| Isoform 2 of Acylamino-acid-releasing enzyme | Q8R146-2 | Apeh | 80 kDa | 16 C | 8.600 | 0.27239 |
| Carbonic anhydrase 1 | P13634 | Ca1 | 28 kDa | 1 C | 8.600 | 0.27239 |
| Ribose-phosphate pyrophosphokinase 1 | Q9D7G0 | Prps1 | 35 kDa | 9 C | 8.400 | 0.10695 |
| Cytochrome P450 2C54 | Q6XVG2 | Cyp2c54 | 56 kDa | 13 C | 8.400 | 0.10695 |
| Eukaryotic translation initiation factor 5 | P59325 | Eif5 | 49 kDa | 8 C | 8.400 | 0.10695 |
| Integrin beta-1 | P09055 | Itgb1 | 88 kDa | 57 C | 8.400 | 0.10695 |
| Ig gamma-3 chain C region | P03987 | | 44 kDa | 10 C | 8.400 | 0.10695 |
| Deoxyribose-phosphate aldolase | Q91YP3 | Dera | 35 kDa | 6 C | 8.400 | 0.10695 |
| Synapse-associated protein 1 | Q9D5V6 | Syap1 | 41 kDa | 1 C | 8.400 | 0.10695 |
| Peroxisomal trans-2-enoyl-CoA reductase | Q99MZ7 | Pecr | 32 kDa | 5 C | 8.400 | 0.10695 |
| DEP domain-containing mTOR-interacting protein | Q570Y9 | Deptor | 46 kDa | 9 C | 7.446 | 0.17095 |
| 60S ribosomal protein L10a | P53026 | Rpl10a | 25 kDa | 1 C | 6.800 | 0.40708 |
| Acyl-coenzyme A synthetase ACSM5, mitochondrial | Q8BGA8 | Acsm5 | 64 kDa | 15 C | 6.800 | 0.40708 |
| CDKN2A-interacting protein | Q8BI72 | Cdkn2aip | 60 kDa | 9 C | 6.800 | 0.40708 |

| | | | | | | |
|---|---|---|---|---|---|---|
| Costars family protein ABRACL | Q4KML4 | Abracl | 9 kDa | 1 C | 6.800 | 0.40708 |
| CD2-associated protein | Q9JLQ0 | Cd2ap | 70 kDa | 3 C | 6.600 | 0.23224 |
| Septin-4 | P28661 (+1) | 43347 | 55 kDa | 8 C | 6.600 | 0.23224 |
| 3-oxoacyl-[acyl-carrier-protein] synthase, mitochondrial | Q9D404 | Oxsm | 49 kDa | 11 C | 6.600 | 0.23224 |
| Peroxisomal targeting signal 2 receptor | P97865 | Pex7 | 36 kDa | 12 C | 6.600 | 0.23224 |
| WD repeat-containing protein 26 | Q8C6G8 | Wdr26 | 71 kDa | 17 C | 6.600 | 0.23224 |
| Platelet glycoprotein 4 | Q08857 | Cd36 | 53 kDa | 10 C | 6.600 | 0.23224 |
| V-type proton ATPase subunit E 1 | P50518 | Atp6v1e1 | 26 kDa | 1 C | 6.600 | 0.23224 |
| Junctional adhesion molecule A | O88792 | F11r | 32 kDa | 5 C | 6.600 | 0.23224 |
| Pyruvate carboxylase, mitochondrial | Q05920 | Pc | 130 kDa | 13 C | 6.600 | 0.23224 |
| Dehydrogenase/reductase SDR family member 4 | Q99LB2 | Dhrs4 | 30 kDa | 4 C | 6.600 | 0.23224 |
| Protein FAM136A | Q9CR98 | Fam136a | 16 kDa | 9 C | 6.600 | 0.23224 |
| 40S ribosomal protein S3 | P62908 | Rps3 | 27 kDa | 3 C | 6.600 | 0.23224 |
| Amine oxidase [flavin-containing] B | Q8BW75 | Maob | 59 kDa | 9 C | 6.600 | 0.23224 |
| Extracellular superoxide dismutase [Cu-Zn] | O09164 | Sod3 | 27 kDa | 6 C | 6.600 | 0.23224 |
| Gamma-aminobutyric acid receptor-associated protein-like 2 | P60521 | Gabarapl2 | 14 kDa | 1 C | 6.600 | 0.23224 |
| L-xylulose reductase | Q91X52 | Dcxr | 26 kDa | 4 C | 6.600 | 0.23224 |
| Lupus La protein homolog | P32067 | Ssb | 48 kDa | 3 C | 6.600 | 0.23224 |
| Uridine diphosphate glucose pyrophosphatase | Q9D142 | Nudt14 | 24 kDa | 4 C | 6.600 | 0.23224 |
| Tubulin alpha-1B chain | P05213 | Tuba1b | 50 kDa | 12 C | 6.277 | 0.20831 |
| ATP synthase subunit alpha, mitochondrial | Q03265 | Atp5a1 | 60 kDa | 2 C | 5.600 | 0.05346 |
| Sepiapterin reductase | Q64105 | Spr | 28 kDa | 10 C | 5.252 | 0.11179 |
| Corticosteroid 11-beta-dehydrogenase isozyme 1 | P50172 | Hsd11b1 | 32 kDa | 3 C | 5.200 | 0.11864 |
| Retinol dehydrogenase 7 | O88451 | Rdh7 | 36 kDa | 8 C | 5.046 | 0.19618 |
| Ribonuclease 4 | Q9JJH1 | Rnase4 | 17 kDa | 8 C | 4.985 | 0.14139 |
| 60S acidic ribosomal protein P1 | P47955 | Rplp1 | 11 kDa | 2 C | 4.985 | 0.06576 |
| Isovaleryl-CoA dehydrogenase, mitochondrial | Q9JHI5 | Ivd | 46 kDa | 7 C | 4.985 | 0.10435 |
| NADH dehydrogenase [ubiquinone] 1 beta subcomplex subunit 7 | Q9CR61 | Ndufb7 | 16 kDa | 4 C | 4.800 | 0.40708 |
| Eukaryotic translation initiation factor 4 gamma 1 | Q6NZJ6 | Eif4g1 | 176 kDa | 15 C | 4.800 | 0.40708 |
| L-gulonolactone oxidase | P58710 | Gulo | 50 kDa | 11 C | 4.800 | 0.40708 |
| NADH dehydrogenase [ubiquinone] 1 alpha subcomplex subunit 5 | Q9CPP6 | Ndufa5 | 13 kDa | 1 C | 4.800 | 0.40708 |
| Ig heavy chain V region 102 | P01750 | | 13 kDa | 3 C | 4.800 | 0.40708 |
| Isoform 4 of Apoptotic chromatin condensation inducer in the nucleus | Q9JIX8-4 | Acin1 | 146 kDa | 7 C | 4.800 | 0.40708 |
| Phenylalanine-4-hydroxylase | P16331 | Pah | 52 kDa | 9 C | 4.800 | 0.40708 |
| Isoform 4 of Heterogeneous nuclear ribonucleoproteins C1/C2 | Q9Z204-4 | Hnrnpc | 32 kDa | 1 C | 4.800 | 0.40708 |
| Molybdopterin synthase sulfur carrier subunit | Q9Z224 | Mocs2 | 10 kDa | 4 C | 4.800 | 0.40708 |
| Phospholysine phosphohistidine inorganic pyrophosphate phosphatase | Q9D7I5 (+1) | Lhpp | 29 kDa | 6 C | 4.800 | 0.40708 |
| Dynactin subunit 2 | Q99KJ8 | Dctn2 | 44 kDa | 2 C | 4.800 | 0.40708 |
| Trafficking protein particle complex subunit 3 | O55013 | Trappc3 | 20 kDa | 3 C | 4.800 | 0.40708 |
| Cytochrome P450 2D26 | Q8CIM7 | Cyp2d26 | 57 kDa | 5 C | 4.800 | 0.40708 |
| Ribonuclease T2-A | C0HKG5 | Rnaset2a | 30 kDa | 10 C | 4.800 | 0.40708 |
| Vesicle-trafficking protein SEC22b | O08547 | Sec22b | 25 kDa | 3 C | 4.800 | 0.40708 |
| Apoptosis-associated speck-like protein containing a CARD | Q9EPB4 | Pycard | 21 kDa | 1 C | 4.800 | 0.40708 |
| Arylacetamide deacetylase | Q99PG0 | Aadac | 45 kDa | 3 C | 4.800 | 0.40708 |
| Gamma-glutamylaminecyclotransferase | Q923B0 | Ggact | 17 kDa | 4 C | 4.800 | 0.40708 |
| Mannose-binding protein C | P41317 | Mbl2 | 26 kDa | 7 C | 4.492 | 0.28537 |
| Formimidoyltransferase-cyclodeaminase | Q91XD4 | Ftcd | 59 kDa | 11 C | 4.388 | 0.13584 |
| Alanine--glyoxylate aminotransferase 2, mitochondrial | Q3UEG6 | Agxt2 | 57 kDa | 14 C | 4.369 | 0.10905 |
| Haloacid dehalogenase-like hydrolase domain-containing protein 3 | Q9CYW4 | Hdhd3 | 28 kDa | 4 C | 4.369 | 0.10905 |
| Small nuclear ribonucleoprotein Sm D2 | P62317 | Snrpd2 | 14 kDa | 2 C | 4.369 | 0.20465 |
| Junction plakoglobin | Q02257 | Jup | 82 kDa | 13 C | 4.117 | 0.37608 |
| Asialoglycoprotein receptor 1 | P34927 | Asgr1 | 33 kDa | 10 C | 4.000 | 0.05041 |
| Acylpyruvase FAHD1, mitochondrial | Q8R0F8 | Fahd1 | 25 kDa | 6 C | 4.000 | 0.09046 |
| Ferritin heavy chain | P09528 | Fth1 | 21 kDa | 3 C | 3.902 | 0.09801 |
| Glyoxalase domain-containing protein 4 | Q9CPV4 | Glod4 | 33 kDa | 5 C | 3.877 | 0.422 |
| OTU domain-containing protein 6B | Q8K2H2 | Otud6b | 34 kDa | 4 C | 3.815 | 0.21303 |
| Alcohol dehydrogenase class-3 | P28474 | Adh5 | 40 kDa | 14 C | 3.815 | 0.21303 |
| ADP/ATP translocase 2 | P51881 | Slc25a5 | 33 kDa | 4 C | 3.815 | 0.21303 |
| UDP-glucuronosyltransferase 1-1 | Q63886 | Ugt1a1 | 60 kDa | 13 C | 3.815 | 0.21303 |
| Transmembrane emp24 domain-containing protein 10 | Q9D1D4 | Tmed10 | 25 kDa | 3 C | 3.815 | 0.21303 |
| Cytochrome c oxidase subunit 4 isoform 1, mitochondrial | P19783 | Cox4i1 | 20 kDa | 1 C | 3.815 | 0.31135 |
| NADH dehydrogenase [ubiquinone] iron-sulfur protein 8, mitochondrial | Q8K3J1 | Ndufs8 | 24 kDa | 8 C | 3.815 | 0.21303 |
| Inhibitor of carbonic anhydrase | Q9DBD0 | Ica | 77 kDa | 35 C | 3.800 | 0.16948 |
| G-rich sequence factor 1 | Q8C5Q4 | Grsf1 | 53 kDa | 9 C | 3.754 | 0.16469 |
| Cytidine deaminase | P56389 | Cda | 16 kDa | 7 C | 3.754 | 0.16469 |
| NADH dehydrogenase [ubiquinone] iron-sulfur protein 3, mitochondrial | Q9DCT2 | Ndufs3 | 30 kDa | 3 C | 3.673 | 0.11552 |
| Cytochrome b5 type B | Q9CQX2 | Cyb5b | 16 kDa | 1 C | 3.639 | 0.10561 |
| Electron transfer flavoprotein-ubiquinone oxidoreductase, mitochondrial | Q921G7 | Etfdh | 68 kDa | 15 C | 3.467 | 0.08748 |
| Fatty aldehyde dehydrogenase | P47740 | Aldh3a2 | 54 kDa | 8 C | 3.360 | 0.08826 |
| Trifunctional enzyme subunit beta, mitochondrial | Q99JY0 | Hadhb | 51 kDa | 5 C | 3.345 | 0.25578 |
| Interferon-activable protein 204 | P0DOV2 | Ifi204 | 69 kDa | 13 C | 3.262 | 0.34006 |
| Vesicle-associated membrane protein-associated protein A | Q9WV55 | Vapa | 28 kDa | 4 C | 3.262 | 0.34006 |
| UDP-glucuronosyltransferase 2B17 | P17717 | Ugt2b17 | 61 kDa | 10 C | 3.200 | 0.21217 |
| Tubulin-specific chaperone A | P48428 | Tbca | 13 kDa | 1 C | 3.200 | 0.2954 |
| 3-oxo-5-beta-steroid 4-dehydrogenase | Q8VCX1 | Akr1d1 | 37 kDa | 6 C | 3.200 | 0.2954 |
| Glutathione S-transferase theta-1 | Q64471 | Gstt1 | 27 kDa | 5 C | 3.200 | 0.21217 |
| SAP domain-containing ribonucleoprotein | Q9D1J3 | Sarnp | 24 kDa | 1 C | 3.200 | 0.2954 |
| Phosphoribosyl pyrophosphate synthase-associated protein 1 | Q9D0M1 | Prpsap1 | 39 kDa | 6 C | 3.200 | 0.2954 |

| | | | | | | |
|---|---|---|---|---|---|---|
| Caspase-6 | O08738 | Casp6 | 32 kDa | 10 C | 3.176 | 0.12963 |
| Insulin-like growth factor-binding protein 2 | P47877 | Igfbp2 | 33 kDa | 3 C | 3.138 | 0.12115 |
| Hemoglobin subunit alpha | P01942 | Hba | 15 kDa | 1 C | 3.100 | 0.22745 |
| Prostaglandin reductase 2 | Q8VDQ1 | Ptgr2 | 38 kDa | 8 C | 3.000 | 0.1315 |
| UPF0568 protein C14orf166 homolog | Q9CQE8 | | 28 kDa | 2 C | 2.982 | 0.20323 |
| Septin-8 | Q8CHH9 (+1 | 43351 | 50 kDa | 6 C | 2.890 | 0.22151 |
| ADP-ribosylation factor GTPase-activating protein 2 | Q99K28 | Arfgap2 | 57 kDa | 6 C | 2.890 | 0.24302 |
| Serine/arginine-rich splicing factor 3 | P84104 (+1) | Srsf3 | 19 kDa | 4 C | 2.751 | 0.25132 |
| Acidic leucine-rich nuclear phosphoprotein 32 family member A | O35381 | Anp32a | 29 kDa | 3 C | 2.720 | 0.08543 |
| Vimentin | P20152 | Vim | 54 kDa | 1 C | 2.708 | 0.55962 |
| DNA-(apurinic or apyrimidinic site) lyase | P28352 | Apex1 | 35 kDa | 7 C | 2.646 | 0.37403 |
| Nicotinate-nucleotide pyrophosphorylase [carboxylating] | Q91X91 | Qprt | 32 kDa | 6 C | 2.646 | 0.44946 |
| N(G),N(G)-dimethylarginine dimethylaminohydrolase 1 | Q9CWS0 | Ddah1 | 31 kDa | 7 C | 2.620 | 0.22167 |
| Receptor expression-enhancing protein 6 | Q9JM62 (+1) | Reep6 | 22 kDa | 3 C | 2.618 | 0.23967 |
| Perilipin-3 | Q9DBG5 | Plin3 | 47 kDa | 2 C | 2.618 | 0.42866 |
| Apolipoprotein E | P08226 | Apoe | 36 kDa | 1 C | 2.595 | 0.09036 |
| Isoform 3 of Septin-9 | Q80UG5-3 | 43352 | 65 kDa | 6 C | 2.585 | 0.28648 |
| Ig gamma-2B chain C region | P01867 (+1) | Igh-3 | 44 kDa | 13 C | 2.585 | 0.28648 |
| Ig kappa chain V-II region 7S34.1 | P01630 | | 12 kDa | 3 C | 2.585 | 0.28648 |
| Succinate--CoA ligase [GDP-forming] subunit beta, mitochondrial | Q9Z2I8 | Suclg2 | 47 kDa | 5 C | 2.585 | 0.28648 |
| BAG family molecular chaperone regulator 1 | Q60739 | Bag1 | 40 kDa | 5 C | 2.585 | 0.28648 |
| Succinate--CoA ligase [ADP/GDP-forming] subunit alpha, mitochondrial | Q9WUM5 | Suclg1 | 36 kDa | 6 C | 2.585 | 0.28648 |
| Cysteine and glycine-rich protein 1 | P97315 | Csrp1 | 21 kDa | 15 C | 2.585 | 0.28648 |
| m7GpppX diphosphatase | Q9DAR7 | Dcps | 39 kDa | 2 C | 2.585 | 0.28648 |
| GDP-mannose 4,6 dehydratase | Q8K0C9 | Gmds | 42 kDa | 6 C | 2.585 | 0.28648 |
| Alanine--tRNA ligase, cytoplasmic | Q8BGQ7 | Aars | 107 kDa | 15 C | 2.585 | 0.28648 |
| Cysteine and glycine-rich protein 3 | P50462 | Csrp3 | 21 kDa | 16 C | 2.582 | 0.30789 |
| Protein NipSnap homolog 1 | O55125 | Nipsnap1 | 33 kDa | 4 C | 2.581 | 0.21579 |
| Iron-sulfur cluster assembly 2 homolog, mitochondrial | Q9DCB8 | Isca2 | 17 kDa | 4 C | 2.560 | 0.05807 |
| Glutamate--cysteine ligase regulatory subunit | O09172 | Gclm | 31 kDa | 6 C | 2.537 | 0.10333 |
| Cytochrome c oxidase subunit 5B, mitochondrial | P19536 | Cox5b | 14 kDa | 5 C | 2.533 | 0.08088 |
| Maleylacetoacetate isomerase | Q9WVL0 | Gstz1 | 24 kDa | 3 C | 2.500 | 0.12458 |
| Cathepsin Z | Q9WUU7 | Ctsz | 34 kDa | 12 C | 2.490 | 0.17874 |
| N-fatty-acyl-amino acid synthase/hydrolase PM20D1 | Q8C165 | Pm20d1 | 56 kDa | 2 C | 2.400 | 0.19113 |
| Desmoplakin | E9Q557 | Dsp | 333 kDa | 43 C | 2.400 | 0.47163 |
| MICOS complex subunit Mic19 | Q9CRB9 | Chchd3 | 26 kDa | 4 C | 2.374 | 0.36373 |
| UPF0598 protein C8orf82 homolog | Q8VE95 | | 24 kDa | 6 C | 2.341 | 0.07969 |
| Short-chain specific acyl-CoA dehydrogenase, mitochondrial | Q07417 | Acads | 45 kDa | 5 C | 2.324 | 0.33353 |
| DCN1-like protein 1 | Q9QZ73 | Dcun1d1 | 30 kDa | 4 C | 2.323 | 0.18103 |
| Cob(I)yrinic acid a,c-diamide adenosyltransferase, mitochondrial | Q9D273 | Mmab | 26 kDa | 5 C | 2.291 | 0.54004 |
| Translationally-controlled tumor protein | P63028 | Tpt1 | 19 kDa | 2 C | 2.267 | 0.15576 |
| Estradiol 17-beta-dehydrogenase 8 | P50171 (+1) | Hsd17b8 | 27 kDa | 4 C | 2.255 | 0.34243 |
| NEDD8-conjugating enzyme Ubc12 | P61082 | Ube2m | 21 kDa | 5 C | 2.240 | 0.16501 |
| Enoyl-CoA hydratase domain-containing protein 3, mitochondrial | Q9D7J9 | Echdc3 | 32 kDa | 5 C | 2.234 | 0.06732 |
| Ubiquitin-conjugating enzyme E2 G1 | P62254 | Ube2g1 | 20 kDa | 2 C | 2.218 | 0.29199 |
| Beta-mannosidase | Q8K2I4 | Manba | 101 kDa | 12 C | 2.196 | 0.21336 |
| Tumor protein D54 | Q9CYZ2 | Tpd52l2 | 24 kDa | 1 C | 2.185 | 0.5549 |
| Peflin | Q8BFY6 | Pef1 | 29 kDa | 5 C | 2.157 | 0.41194 |
| Cytochrome b-c1 complex subunit 2, mitochondrial | Q9DB77 | Uqcrc2 | 48 kDa | 1 C | 2.100 | 0.15297 |
| Lysosomal protective protein | P16675 | Ctsa | 54 kDa | 11 C | 2.092 | 0.62111 |
| Delta-1-pyrroline-5-carboxylate dehydrogenase, mitochondrial | Q8CHT0 | Aldh4a1 | 62 kDa | 8 C | 2.050 | 0.35177 |
| Ig kappa chain V-V region L6 (Fragment) | P01638 | | 13 kDa | 3 C | 2.050 | 0.4724 |
| Methionine aminopeptidase 1 | Q8BP48 | Metap1 | 43 kDa | 17 C | 2.039 | 0.10181 |
| Malectin | Q6ZQI3 | Mlec | 32 kDa | 3 C | 2.031 | 0.50045 |
| Arylsulfatase B | P50429 | Arsb | 60 kDa | 10 C | 2.031 | 0.50045 |
| Peptidyl-prolyl cis-trans isomerase NIMA-interacting 4 | Q9CWW6 | Pin4 | 14 kDa | 1 C | 2.031 | 0.50045 |
| Exosome complex component RRP41 | Q921I9 | Exosc4 | 26 kDa | 5 C | 2.020 | 0.31158 |
| Hydroxyacid-oxoacid transhydrogenase, mitochondrial | Q8R0N6 | Adhfe1 | 50 kDa | 8 C | 2.000 | 0.07298 |
| Alpha-soluble NSF attachment protein | Q9DB05 | Napa | 33 kDa | 8 C | 2.000 | 0.09956 |
| Cathepsin B | P10605 | Ctsb | 37 kDa | 16 C | 2.000 | 0.1405 |
| StAR-related lipid transfer protein 5 | Q9EPQ7 | Stard5 | 24 kDa | 8 C | 1.964 | 0.00759 |
| Hemoglobin subunit beta-1 | P02088 | Hbb-b1 | 16 kDa | 2 C | 1.953 | 0.2518 |
| Trans-1,2-dihydrobenzene-1,2-diol dehydrogenase | Q9DBB8 | Dhdh | 36 kDa | 6 C | 1.951 | 0.15458 |
| Methylglutaconyl-CoA hydratase, mitochondrial | Q9JLZ3 | Auh | 33 kDa | 5 C | 1.951 | 0.02433 |
| Purine nucleoside phosphorylase | P23492 | Pnp | 32 kDa | 5 C | 1.943 | 0.0661 |
| Phosphomannomutase 2 | Q9Z2M7 | Pmm2 | 28 kDa | 6 C | 1.927 | 0.50564 |
| Delta(3,5)-Delta(2,4)-dienoyl-CoA isomerase, mitochondrial | O35459 | Ech1 | 36 kDa | 6 C | 1.920 | 0.0945 |
| GMP reductase 2 | Q99L27 | Gmpr2 | 38 kDa | 9 C | 1.917 | 0.10348 |
| Ras-related protein Rab-11B | P46638 | Rab11b | 24 kDa | 2 C | 1.905 | 0.30786 |
| UDP-glucose 6-dehydrogenase | O70475 | Ugdh | 55 kDa | 12 C | 1.900 | 0.40751 |
| Protein-glucosylgalactosylhydroxylysine glucosidase | Q8BP56 | Pgghg | 76 kDa | 8 C | 1.898 | 0.18087 |
| Isoform 2 of ATP-dependent (S)-NAD(P)H-hydrate dehydratase | Q9CZ42-2 | Naxd | 32 kDa | 8 C | 1.891 | 0.38865 |
| Exosome complex component RRP43 | Q9D753 | Exosc8 | 30 kDa | 10 C | 1.891 | 0.47061 |
| Heterogeneous nuclear ribonucleoprotein M | Q9D0E1 | Hnrnpm | 78 kDa | 6 C | 1.867 | 0.00468 |
| Exosome complex component RRP4 | Q8VBV3 | Exosc2 | 33 kDa | 4 C | 1.855 | 0.28959 |
| Phosphatidylinositol transfer protein beta isoform | P53811 | Pitpnb | 31 kDa | 5 C | 1.840 | 0.26516 |
| Isoform 2 of WD repeat domain phosphoinositide-interacting protein 4 | Q91VM3-2 | Wdr45 | 38 kDa | 14 C | 1.832 | 0.31542 |
| Regulator of microtubule dynamics protein 3 | Q3UJU9 | Rmdn3 | 52 kDa | 6 C | 1.832 | 0.31542 |

| | | | | | | |
|---|---|---|---|---|---|---|
| Splicing factor U2AF 65 kDa subunit | P26369 | U2af2 | 54 kDa | 6 C | 1.829 | 0.1363 |
| Dipeptidyl peptidase 1 | P97821 | Ctsc | 52 kDa | 14 C | 1.829 | 0.1363 |
| Histidine triad nucleotide-binding protein 1 | P70349 | Hint1 | 14 kDa | 2 C | 1.829 | 0.05551 |
| 2,4-dienoyl-CoA reductase, mitochondrial | Q9CQ62 | Decr1 | 36 kDa | 5 C | 1.818 | 0.12381 |
| Interferon-induced 35 kDa protein homolog | Q9D8C4 | Ifi35 | 32 kDa | 4 C | 1.810 | 0.28516 |
| Thiosulfate sulfurtransferase | P52196 | Tst | 33 kDa | 4 C | 1.803 | 0.26181 |
| Medium-chain specific acyl-CoA dehydrogenase, mitochondrial | P45952 | Acadm | 46 kDa | 8 C | 1.800 | 0.15527 |
| ATP synthase subunit O, mitochondrial | Q9DB20 | Atp5o | 23 kDa | 1 C | 1.800 | 0.26556 |
| Lysosomal alpha-mannosidase | O09159 | Man2b1 | 115 kDa | 10 C | 1.800 | 0.43065 |
| Carbonyl reductase [NADPH] 1 | P48758 | Cbr1 | 31 kDa | 5 C | 1.795 | 0.41714 |
| Glucosidase 2 subunit beta | O08795 (+1) | Prkcsh | 59 kDa | 17 C | 1.778 | 0.3 |
| Triokinase/FMN cyclase | Q8VC30 | Tkfc | 60 kDa | 5 C | 1.776 | 0.31128 |
| Omega-amidase NIT2 | Q9JHW2 | Nit2 | 31 kDa | 2 C | 1.774 | 0.03249 |
| Protein arginine N-methyltransferase 5 | Q8CIG8 | Prmt5 | 73 kDa | 12 C | 1.770 | 0.67451 |
| Receptor of activated protein C kinase 1 | P68040 | Rack1 | 35 kDa | 8 C | 1.768 | 0.01459 |
| NADH dehydrogenase [ubiquinone] flavoprotein 2, mitochondrial | Q9D6J6 | Ndufv2 | 27 kDa | 6 C | 1.768 | 0.02555 |
| Isoform Soluble of Catechol O-methyltransferase | O88587-2 | Comt | 25 kDa | 4 C | 1.760 | 0.21492 |
| Extracellular matrix protein 1 | Q61508 | Ecm1 | 63 kDa | 29 C | 1.760 | 0.07907 |
| Heme-binding protein 1 | Q9R257 | Hebp1 | 21 kDa | 2 C | 1.759 | 0.15211 |
| Carboxypeptidase Q | Q9WVJ3 (+1 | Cpq | 52 kDa | 1 C | 1.756 | 0.19746 |
| Methylmalonyl-CoA epimerase, mitochondrial | Q9D1I5 | Mcee | 19 kDa | 2 C | 1.756 | 0.33212 |
| 60S ribosomal protein L22 | P67984 | Rpl22 | 15 kDa | 1 C | 1.756 | 0.33212 |
| Kininogen-1 | O08677 | Kng1 | 73 kDa | 19 C | 1.752 | 0.09986 |
| Fumarate hydratase, mitochondrial | P97807 (+1) | Fh | 54 kDa | 4 C | 1.752 | 0.52504 |
| Pregnancy zone protein | Q61838 | Pzp | 166 kDa | 24 C | 1.745 | 0.09876 |
| Protein phosphatase 1A | P49443 | Ppm1a | 42 kDa | 10 C | 1.745 | 0.02163 |
| ATP synthase subunit d, mitochondrial | Q9DCX2 | Atp5h | 19 kDa | 1 C | 1.733 | 0.17858 |
| Low molecular weight phosphotyrosine protein phosphatase | Q9D358 | Acp1 | 18 kDa | 8 C | 1.725 | 0.40961 |
| Sorbin and SH3 domain-containing protein 1 | Q62417 | Sorbs1 | 143 kDa | 4 C | 1.723 | 0.17615 |
| Trifunctional enzyme subunit alpha, mitochondrial | Q8BMS1 | Hadha | 83 kDa | 12 C | 1.723 | 0.49231 |
| Agmatinase, mitochondrial | A2AS89 | Agmat | 38 kDa | 9 C | 1.723 | 0.00672 |
| Isoform 2 of Ataxin-2 | O70305-2 | Atxn2 | 129 kDa | 15 C | 1.714 | 0.2675 |
| SEC14-like protein 2 | Q99J08 | Sec14l2 | 46 kDa | 8 C | 1.713 | 0.46254 |
| Annexin A6 | P14824 | Anxa6 | 76 kDa | 8 C | 1.712 | 0.59584 |
| Protein canopy homolog 2 | Q9QXT0 | Cnpy2 | 21 kDa | 6 C | 1.700 | 0.19962 |
| Serpin B6 | Q60854 | Serpinb6 | 43 kDa | 6 C | 1.700 | 0.248 |
| Glucose-6-phosphate isomerase | P06745 | Gpi | 63 kDa | 4 C | 1.700 | 0.39539 |
| Ornithine carbamoyltransferase, mitochondrial | P11725 | Otc | 40 kDa | 2 C | 1.656 | 0.00518 |
| Argininosuccinate lyase | Q91YI0 | Asl | 52 kDa | 13 C | 1.653 | 0.41103 |
| Desmoglein-1-beta | Q7TSF1 | Dsg1b | 114 kDa | 18 C | 1.639 | 0.73006 |
| Beta-ureidopropionase | Q8VC97 | Upb1 | 44 kDa | 10 C | 1.621 | 0.092 |
| Isoform 2 of Clathrin interactor 1 | Q99KN9-2 | Clint1 | 51 kDa | 1 C | 1.620 | 0.30527 |
| ELKS/Rab6-interacting/CAST family member 1 | Q99MI1 | Erc1 | 128 kDa | 4 C | 1.607 | 0.30283 |
| Alpha-2-HS-glycoprotein | P29699 | Ahsg | 37 kDa | 14 C | 1.600 | 0.10179 |
| Ubiquitin-fold modifier-conjugating enzyme 1 | Q9CR09 | Ufc1 | 19 kDa | 3 C | 1.600 | 0.1674 |
| Glutathione S-transferase omega-1 | O09131 | Gsto1 | 27 kDa | 4 C | 1.600 | 0.5552 |
| Glycerol-3-phosphate dehydrogenase [NAD(+)], cytoplasmic | P13707 | Gpd1 | 38 kDa | 11 C | 1.600 | 0.03515 |
| Chromobox protein homolog 1 | P83917 | Cbx1 | 21 kDa | 2 C | 1.600 | 0.16352 |
| Sarcosine dehydrogenase, mitochondrial | Q99LB7 | Sardh | 102 kDa | 18 C | 1.600 | 0.18777 |
| Proliferation-associated protein 2G4 | P50580 | Pa2g4 | 44 kDa | 6 C | 1.600 | 0.26249 |
| Ficolin-1 | O70165 | Fcn1 | 36 kDa | 10 C | 1.600 | 0.34334 |
| Ceruloplasmin | Q61147 | Cp | 121 kDa | 14 C | 1.600 | 0.50876 |
| 3-ketoacyl-CoA thiolase, mitochondrial | Q8BWT1 | Acaa2 | 42 kDa | 8 C | 1.577 | 0.15487 |
| Coatomer subunit zeta-1 | P61924 | Copz1 | 20 kDa | 1 C | 1.574 | 0.35748 |
| Acyl-protein thioesterase 1 | P97823 (+1) | Lypla1 | 25 kDa | 6 C | 1.574 | 0.35748 |
| cAMP-dependent protein kinase type I-alpha regulatory subunit | Q9DBC7 | Prkar1a | 43 kDa | 5 C | 1.574 | 0.2594 |
| Aspartate aminotransferase, mitochondrial | P05202 | Got2 | 47 kDa | 7 C | 1.568 | 0.06385 |
| Glutathione S-transferase A3 | P30115 | Gsta3 | 25 kDa | 1 C | 1.564 | 0.01247 |
| Fumarylacetoacetate hydrolase domain-containing protein 2A | Q3TC72 | Fahd2 | 35 kDa | 6 C | 1.561 | 0.34459 |
| Phenazine biosynthesis-like domain-containing protein 2 | Q9CXN7 | Pbld2 | 32 kDa | 4 C | 1.560 | 0.18626 |
| Isoform 2 of F-actin-capping protein subunit beta | P47757-2 | Capzb | 31 kDa | 5 C | 1.558 | 0.05184 |
| Leucine-rich repeat-containing protein 59 | Q922Q8 | Lrrc59 | 35 kDa | 8 C | 1.550 | 0.52681 |
| Legumain | O89017 | Lgmn | 49 kDa | 7 C | 1.550 | 0.57674 |
| Uricase | P25688 | Uox | 35 kDa | 4 C | 1.545 | 0.19053 |
| S-phase kinase-associated protein 1 | Q9WTX5 | Skp1 | 19 kDa | 3 C | 1.538 | 0.05753 |
| Glutamate dehydrogenase 1, mitochondrial | P26443 | Glud1 | 61 kDa | 6 C | 1.538 | 0.31543 |
| Poly(rC)-binding protein 1 | P60335 | Pcbp1 | 37 kDa | 9 C | 1.533 | 0.08449 |
| Gamma-soluble NSF attachment protein | Q9CWZ7 | Napg | 35 kDa | 6 C | 1.528 | 0.31411 |
| Zinc phosphodiesterase ELAC protein 2 | Q80Y81 (+1) | Elac2 | 93 kDa | 23 C | 1.527 | 0.54905 |
| Farnesyl pyrophosphate synthase | Q920E5 | Fdps | 41 kDa | 2 C | 1.527 | 0.54905 |
| Ubiquitin carboxyl-terminal hydrolase isozyme L3 | Q9JKB1 | Uchl3 | 26 kDa | 3 C | 1.527 | 0.54905 |
| Proteasome activator complex subunit 2 | P97372 | Psme2 | 27 kDa | 4 C | 1.527 | 0.54905 |
| Alpha-actinin-4 | P57780 | Actn4 | 105 kDa | 8 C | 1.527 | 0.04095 |
| Phosphoglucomutase-1 | Q9D0F9 | Pgm1 | 61 kDa | 10 C | 1.518 | 0.15861 |
| Glutathione S-transferase A1 | P13745 | Gsta1 | 26 kDa | 2 C | 1.515 | 0.00187 |
| Phytanoyl-CoA dioxygenase, peroxisomal | O35386 | Phyh | 39 kDa | 8 C | 1.511 | 0.15173 |
| Ferritin light chain 1 | P29391 | Ftl1 | 21 kDa | 1 C | 1.509 | 0.08322 |
| Isoform Mt-VDAC1 of Voltage-dependent anion-selective channel protein 1 | Q60932-2 | Vdac1 | 31 kDa | 2 C | 1.500 | 0.30144 |

| | | | | | | |
|---|---|---|---|---|---|---|
| Cysteine sulfinic acid decarboxylase | Q9DBE0 | Csad | 55 kDa | 11 C | 1.500 | 0.31021 |
| Retinol-binding protein 4 | Q00724 | Rbp4 | 23 kDa | 6 C | 1.500 | 0.32686 |
| Ubiquitin-fold modifier 1 | P61961 | Ufm1 | 9 kDa | 1 C | 1.500 | 0.32686 |
| L-lactate dehydrogenase A chain | P06151 | Ldha | 36 kDa | 6 C | 1.500 | 0.5205 |
| Proteasome subunit alpha type-2 | P49722 | Psma2 | 26 kDa | 2 C | 1.500 | 0.04642 |
| Polyribonucleotide nucleotidyltransferase 1, mitochondrial | Q8K1R3 | Pnpt1 | 86 kDa | 16 C | 1.500 | 0.41122 |
| Moesin | P26041 | Msn | 68 kDa | 2 C | 1.497 | 0.43833 |
| Mitochondrial amidoxime reducing component 2 | Q922Q1 | 43161 | 38 kDa | 11 C | 1.497 | 0.50139 |
| Haptoglobin | Q61646 | Hp | 39 kDa | 9 C | 1.491 | 0.10968 |
| Urocanate hydratase | Q8VC12 | Uroc1 | 75 kDa | 12 C | 1.490 | 0.07155 |
| 2-oxo-4-hydroxy-4-carboxy-5-ureidoimidazoline decarboxylase | Q283N4 | Urad | 20 kDa | 3 C | 1.486 | 0.1355 |
| Nucleoside diphosphate kinase 3 | Q9WV85 | Nme3 | 19 kDa | 3 C | 1.486 | 0.36446 |
| Polyubiquitin-C | P0CG50 | Ubc | 83 kDa | 6 C | 1.482 | 0.01595 |
| Transcription elongation factor A protein 3 | P23881 | Tcea3 | 39 kDa | 13 C | 1.477 | 0.75324 |
| Granulins | P28798 | Grn | 63 kDa | 88 C | 1.477 | 0.75324 |
| COP9 signalosome complex subunit 8 | Q8VBV7 | Cops8 | 23 kDa | 1 C | 1.477 | 0.75324 |
| Transthyretin | P07309 | Ttr | 16 kDa | 2 C | 1.467 | 0.07862 |
| Actin-related protein 2/3 complex subunit 2 | Q9CVB6 | Arpc2 | 34 kDa | 2 C | 1.467 | 0.32488 |
| Ketohexokinase | P97328 | Khk | 33 kDa | 10 C | 1.462 | 0.08651 |
| Thioredoxin | P10639 | Txn | 12 kDa | 6 C | 1.462 | 0.16831 |
| Ubiquitin-like-conjugating enzyme ATG3 | Q9CPX6 | Atg3 | 36 kDa | 8 C | 1.461 | 0.66058 |
| Malate dehydrogenase, mitochondrial | P08249 | Mdh2 | 36 kDa | 8 C | 1.461 | 0.0305 |
| Eukaryotic translation initiation factor 2 subunit 1 | Q6ZWX6 | Eif2s1 | 36 kDa | 5 C | 1.456 | 0.42974 |
| CD5 antigen-like | Q9QWK4 | Cd5l | 39 kDa | 26 C | 1.455 | 0.12979 |
| Immunoglobulin J chain | P01592 | Jchain | 18 kDa | 8 C | 1.455 | 0.20985 |
| Major urinary protein 3 | P04939 | Mup3 | 21 kDa | 5 C | 1.455 | 0.2675 |
| 14 kDa phosphohistidine phosphatase | Q9DAK9 | Phpt1 | 14 kDa | 3 C | 1.450 | 0.06758 |
| Isoform Peroxisomal of Serine--pyruvate aminotransferase, mitochondrial | O35423-2 | Agxt | 44 kDa | 7 C | 1.450 | 0.30309 |
| Methyltransferase-like 26 | Q9DCS2 | Mettl26 | 23 kDa | 6 C | 1.446 | 0.03839 |
| Cytochrome c oxidase assembly factor 7 | Q921H9 | Coa7 | 26 kDa | 13 C | 1.440 | 0.02218 |
| Regucalcin | Q64374 | Rgn | 33 kDa | 9 C | 1.430 | 0.18061 |
| Heterogeneous nuclear ribonucleoprotein A3 | Q8BG05 | Hnrnpa3 | 40 kDa | 4 C | 1.429 | 0.05336 |
| Immunoglobulin-binding protein 1 | Q61249 | Igbp1 | 39 kDa | 3 C | 1.427 | 0.56919 |
| Retinal dehydrogenase 1 | P24549 | Aldh1a1 | 54 kDa | 11 C | 1.426 | 0.01131 |
| Beta-2-glycoprotein 1 | Q01339 | Apoh | 39 kDa | 23 C | 1.422 | 0.1715 |
| Chromobox protein homolog 3 | P23198 | Cbx3 | 21 kDa | 3 C | 1.422 | 0.25302 |
| Gamma-butyrobetaine dioxygenase | Q924Y0 | Bbox1 | 45 kDa | 9 C | 1.422 | 0.01178 |
| Polypyrimidine tract-binding protein 1 | P17225 | Ptbp1 | 56 kDa | 3 C | 1.420 | 0.47985 |
| Phenazine biosynthesis-like domain-containing protein 1 | Q9DCG6 | Pbld1 | 32 kDa | 3 C | 1.418 | 0.30703 |
| Acyl-coenzyme A thioesterase 4 | Q8BWN8 | Acot4 | 46 kDa | 6 C | 1.415 | 0.29316 |
| NADH dehydrogenase [ubiquinone] flavoprotein 1, mitochondrial | Q91YT0 | Ndufv1 | 51 kDa | 12 C | 1.413 | 0.00137 |
| 2-iminobutanoate/2-iminopropanoate deaminase | P52760 | Rida | 14 kDa | 1 C | 1.412 | 0.05753 |
| Density-regulated protein | Q9CQJ6 | Denr | 22 kDa | 7 C | 1.410 | 0.67382 |
| Argininosuccinate synthase | P16460 | Ass1 | 47 kDa | 5 C | 1.404 | 0.64667 |
| Isoform 2 of Heterogeneous nuclear ribonucleoprotein A3 | Q8BG05-2 | Hnrnpa3 | 37 kDa | 4 C | 1.402 | 0.06557 |
| Glyoxylate reductase/hydroxypyruvate reductase | Q91Z53 | Grhpr | 35 kDa | 7 C | 1.400 | 0.05095 |
| Isoform 2 of Thioredoxin reductase 2, mitochondrial | Q9JLT4-2 | Txnrd2 | 53 kDa | 11 C | 1.400 | 0.06758 |
| UBX domain-containing protein 1 | Q922Y1 | Ubxn1 | 34 kDa | 2 C | 1.400 | 0.10133 |
| Isopentenyl-diphosphate Delta-isomerase 1 | P58044 | Idi1 | 26 kDa | 8 C | 1.400 | 0.19301 |
| Elongation factor 1-beta | O70251 | Eef1b | 25 kDa | 3 C | 1.400 | 0.30967 |
| UMP-CMP kinase | Q9DBP5 | Cmpk1 | 22 kDa | 6 C | 1.400 | 0.0339 |
| Carboxylesterase 3B | Q8VCU1 | Ces3b | 63 kDa | 7 C | 1.388 | 0.3776 |
| Aminoacylase-1 | Q99JW2 | Acy1 | 46 kDa | 4 C | 1.387 | 0.23691 |
| Rho GDP-dissociation inhibitor 1 | Q99PT1 | Arhgdia | 23 kDa | 1 C | 1.387 | 0.23691 |
| S-formylglutathione hydrolase | Q9R0P3 | Esd | 31 kDa | 10 C | 1.387 | 0.38175 |
| Isoform 2 of Disks large homolog 1 | Q811D0-2 | Dlg1 | 100 kDa | 7 C | 1.385 | 0.56724 |
| Methylmalonate-semialdehyde dehydrogenase [acylating], mitochondrial | Q9EQ20 | Aldh6a1 | 58 kDa | 8 C | 1.384 | 0.01085 |
| NADH dehydrogenase [ubiquinone] 1 beta subcomplex subunit 10 | Q9DCS9 | Ndufb10 | 21 kDa | 5 C | 1.383 | 0.60244 |
| Ribosome-recycling factor, mitochondrial | Q9D6S7 | Mrrf | 29 kDa | 2 C | 1.383 | 0.67898 |
| 1,2-dihydroxy-3-keto-5-methylthiopentene dioxygenase | Q99JT9 | Adi1 | 22 kDa | 1 C | 1.382 | 0.22208 |
| Histidine-rich glycoprotein | Q9ESB3 | Hrg | 59 kDa | 17 C | 1.382 | 0.32853 |
| Oxygen-dependent coproporphyrinogen-III oxidase, mitochondrial | P36552 | Cpox | 50 kDa | 10 C | 1.378 | 0.01519 |
| Ribonuclease inhibitor | Q91VI7 | Rnh1 | 50 kDa | 30 C | 1.378 | 0.06526 |
| Proteasome subunit alpha type-3 | O70435 | Psma3 | 28 kDa | 4 C | 1.378 | 0.15604 |
| Actin-related protein 2/3 complex subunit 4 | P59999 | Arpc4 | 20 kDa | 4 C | 1.371 | 0.10901 |
| Heterogeneous nuclear ribonucleoprotein A1 | P49312 | Hnrnpa1 | 34 kDa | 2 C | 1.371 | 0.16455 |
| UPF0160 protein MYG1, mitochondrial | Q9JK81 | Myg1 | 43 kDa | 7 C | 1.371 | 0.26365 |
| Alpha-enolase | P17182 | Eno1 | 47 kDa | 6 C | 1.370 | 0.2165 |
| Acyl-coenzyme A thioesterase THEM4 | Q3UUI3 | Them4 | 26 kDa | 4 C | 1.365 | 0.11509 |
| Endonuclease G, mitochondrial | O08600 | Endog | 32 kDa | 2 C | 1.365 | 0.43912 |
| Myosin light polypeptide 6 | Q60605 (+1) | Myl6 | 17 kDa | 3 C | 1.365 | 0.46138 |
| Cofilin-2 | P45591 | Cfl2 | 19 kDa | 2 C | 1.360 | 0.14557 |
| Clathrin light chain A | O08585 | Clta | 26 kDa | 1 C | 1.360 | 0.30309 |
| Plasminogen activator inhibitor 1 RNA-binding protein | Q9CY58 | Serbp1 | 45 kDa | 2 C | 1.360 | 0.60793 |
| 39S ribosomal protein L53, mitochondrial | Q9D1H8 | Mrpl53 | 13 kDa | 3 C | 1.352 | 0.63279 |
| Adenosylhomocysteinase | P50247 | Ahcy | 48 kDa | 9 C | 1.352 | 0.03384 |
| Endoplasmic reticulum resident protein 44 | Q9D1Q6 | Erp44 | 47 kDa | 7 C | 1.350 | 0.36983 |
| Phosphoglycerate kinase 1 | P09411 | Pgk1 | 45 kDa | 7 C | 1.350 | 0.17861 |

| | | | | | | |
|---|---|---|---|---|---|---|
| Dihydropteridine reductase | Q8BVI4 | Qdpr | 26 kDa | 4 C | 1.347 | 0.12498 |
| Galectin-1 | P16045 | Lgals1 | 15 kDa | 6 C | 1.345 | 0.10493 |
| 60 kDa heat shock protein, mitochondrial | P63038 | Hspd1 | 61 kDa | 3 C | 1.343 | 0.04026 |
| Na(+)/H(+) exchange regulatory cofactor NHE-RF1 | P70441 | Slc9a3r1 | 39 kDa | 5 C | 1.342 | 0.12604 |
| Fructose-bisphosphate aldolase C | P05063 | Aldoc | 39 kDa | 7 C | 1.341 | 0.66196 |
| Actin, alpha skeletal muscle | P68134 | Acta1 | 42 kDa | 6 C | 1.341 | 0.12168 |
| Electron transfer flavoprotein subunit beta | Q9DCW4 | Etfb | 28 kDa | 4 C | 1.337 | 0.09492 |
| Hydroxyacyl-coenzyme A dehydrogenase, mitochondrial | Q61425 | Hadh | 34 kDa | 5 C | 1.333 | 0.10439 |
| COP9 signalosome complex subunit 5 | O35864 | Cops5 | 38 kDa | 4 C | 1.333 | 0.13952 |
| Aldose 1-epimerase | Q8K157 | Galm | 38 kDa | 4 C | 1.333 | 0.20985 |
| AN1-type zinc finger protein 6 | Q9DCH6 | Zfand6 | 24 kDa | 12 C | 1.333 | 0.29235 |
| Copper homeostasis protein cutC homolog | Q9D8X1 | Cutc | 29 kDa | 7 C | 1.333 | 0.36489 |
| Mesencephalic astrocyte-derived neurotrophic factor | Q9CXI5 | Manf | 20 kDa | 8 C | 1.333 | 0.39693 |
| Secernin-3 | Q3TMH2 | Scrn3 | 48 kDa | 7 C | 1.333 | 0.4652 |
| Actin, cytoplasmic 2 | P63260 | Actg1 | 42 kDa | 6 C | 1.330 | 0.10041 |
| Peptidyl-prolyl cis-trans isomerase FKBP3 | Q62446 | Fkbp3 | 25 kDa | 1 C | 1.325 | 0.75368 |
| Tight junction protein ZO-1 | P39447 | Tjp1 | 195 kDa | 7 C | 1.325 | 0.79102 |
| 14-3-3 protein eta | P68510 | Ywhah | 28 kDa | 3 C | 1.320 | 0.51133 |
| NADH-ubiquinone oxidoreductase 75 kDa subunit, mitochondrial | Q91VD9 | Ndufs1 | 80 kDa | 18 C | 1.319 | 0.12418 |
| Elongation factor 1-delta | P57776 | Eef1d | 31 kDa | 2 C | 1.318 | 0.31264 |
| Isoform 2 of 14-3-3 protein theta | P68254-2 | Ywhaq | 28 kDa | 5 C | 1.318 | 0.4483 |
| Heterogeneous nuclear ribonucleoprotein L | Q8R081 | Hnrnpl | 64 kDa | 11 C | 1.314 | 0.50538 |
| Proteasome subunit alpha type-5 | Q9Z2U1 | Psma5 | 26 kDa | 3 C | 1.312 | 0.10901 |
| Peroxiredoxin-2 | Q61171 | Prdx2 | 22 kDa | 3 C | 1.309 | 0.09191 |
| Alpha-actinin-1 | Q7TPR4 | Actn1 | 103 kDa | 11 C | 1.309 | 0.29664 |
| Serum albumin | P07724 | Alb | 69 kDa | 36 C | 1.305 | 0.30176 |
| Radixin | P26043 | Rdx | 69 kDa | 1 C | 1.304 | 0.54733 |
| Transgelin | P37804 | Tagln | 23 kDa | 1 C | 1.302 | 0.81882 |
| Macrophage migration inhibitory factor | P34884 | Mif | 13 kDa | 3 C | 1.300 | 0.22745 |
| NFU1 iron-sulfur cluster scaffold homolog, mitochondrial | Q9QZ23 | Nfu1 | 29 kDa | 5 C | 1.300 | 0.46084 |
| Ig heavy chain V region MOPC 21 (Fragment) | P01783 | | 15 kDa | 3 C | 1.300 | 0.71228 |
| Transforming protein RhoA | Q9QUI0 | Rhoa | 22 kDa | 6 C | 1.300 | 0.71228 |
| Myeloid-derived growth factor | Q9CPT4 | Mydgf | 18 kDa | 2 C | 1.300 | 0.74876 |
| Protein DDI1 homolog 2 | A2ADY9 | Ddi2 | 45 kDa | 8 C | 1.297 | 0.1652 |
| Dihydrolipoyl dehydrogenase, mitochondrial | O08749 | Dld | 54 kDa | 9 C | 1.292 | 0.19631 |
| Heterogeneous nuclear ribonucleoprotein A/B | Q99020 | Hnrnpab | 31 kDa | 2 C | 1.291 | 0.10133 |
| Dimethylglycine dehydrogenase, mitochondrial | Q9DBT9 | Dmgdh | 97 kDa | 4 C | 1.289 | 0.24139 |
| Ig alpha chain C region | P01878 | | 37 kDa | 13 C | 1.289 | 0.40708 |
| Eukaryotic translation initiation factor 2 subunit 3, X-linked | Q9Z0N1 | Eif2s3x | 51 kDa | 10 C | 1.286 | 0.72181 |
| Propionyl-CoA carboxylase beta chain, mitochondrial | Q99MN9 | Pccb | 58 kDa | 11 C | 1.286 | 0.05688 |
| Isoform 2 of Nitrilase homolog 1 | Q8VDK1-2 | Nit1 | 32 kDa | 13 C | 1.280 | 0.07862 |
| Actin-related protein 2/3 complex subunit 3 | Q9JM76 | Arpc3 | 21 kDa | 4 C | 1.280 | 0.19962 |
| Major urinary protein 1 | P11588 | Mup1 | 21 kDa | 5 C | 1.280 | 0.3493 |
| Histidine triad nucleotide-binding protein 2, mitochondrial | Q9D0S9 | Hint2 | 17 kDa | 1 C | 1.280 | 0.35582 |
| 3'(2'),5'-bisphosphate nucleotidase 1 | Q9Z0S1 | Bpnt1 | 33 kDa | 6 C | 1.280 | 0.41207 |
| Probable D-lactate dehydrogenase, mitochondrial | Q7TNG8 | Ldhd | 52 kDa | 12 C | 1.280 | 0.42169 |
| Ubiquitin-conjugating enzyme E2 N | P61089 | Ube2n | 17 kDa | 1 C | 1.280 | 0.48156 |
| Indolethylamine N-methyltransferase | P40936 | Inmt | 29 kDa | 11 C | 1.280 | 0.48156 |
| Arginase-1 | Q61176 | Arg1 | 35 kDa | 3 C | 1.273 | 0.23303 |
| Aldehyde dehydrogenase, cytosolic 1 | O35945 | Aldh1a7 | 55 kDa | 8 C | 1.273 | 0.02277 |
| Vitronectin | P29788 | Vtn | 55 kDa | 14 C | 1.269 | 0.50958 |
| Fatty acid-binding protein, liver | P12710 | Fabp1 | 14 kDa | 1 C | 1.268 | 0.00646 |
| Na(+)/H(+) exchange regulatory cofactor NHE-RF3 | Q9JIL4 | Pdzk1 | 56 kDa | 7 C | 1.268 | 0.12003 |
| 14-3-3 protein beta/alpha | Q9CQV8 | Ywhab | 28 kDa | 2 C | 1.265 | 0.37367 |
| Liver carboxylesterase 1 | Q8VCC2 | Ces1 | 63 kDa | 7 C | 1.265 | 0.04895 |
| Glyceraldehyde-3-phosphate dehydrogenase | P16858 | Gapdh | 36 kDa | 5 C | 1.261 | 0.18312 |
| NAD kinase 2, mitochondrial | Q8C5H8 | Nadk2 | 51 kDa | 9 C | 1.261 | 0.13994 |
| Peroxiredoxin-1 | P35700 | Prdx1 | 22 kDa | 4 C | 1.260 | 0.02531 |
| FAS-associated death domain protein | Q61160 | Fadd | 23 kDa | 3 C | 1.257 | 0.4758 |
| Protein-glutamine gamma-glutamyltransferase K | Q9JLF6 | Tgm1 | 90 kDa | 16 C | 1.257 | 0.63174 |
| 2-oxoglutarate dehydrogenase, mitochondrial | Q60597 | Ogdh | 116 kDa | 21 C | 1.257 | 0.46637 |
| 28S ribosomal protein S22, mitochondrial | Q9CXW2 | Mrps22 | 41 kDa | 2 C | 1.253 | 0.57978 |
| Pyridoxine-5'-phosphate oxidase | Q91XF0 | Pnpo | 30 kDa | 6 C | 1.253 | 0.60249 |
| Flavin reductase (NADPH) | Q923D2 | Blvrb | 22 kDa | 2 C | 1.253 | 0.13923 |
| Superoxide dismutase [Cu-Zn] | P08228 | Sod1 | 16 kDa | 3 C | 1.252 | 0.27712 |
| Ran-specific GTPase-activating protein | P34022 | Ranbp1 | 24 kDa | 3 C | 1.250 | 0.1514 |
| Heterogeneous nuclear ribonucleoprotein K | P61979 (+1) | Hnrnpk | 51 kDa | 5 C | 1.250 | 0.31212 |
| Proteasome subunit beta type-5 | O55234 | Psmb5 | 29 kDa | 3 C | 1.247 | 0.12218 |
| BolA-like protein 1 | Q9D8S9 | Bola1 | 14 kDa | 3 C | 1.244 | 0.12498 |
| Proteasome subunit alpha type-7 | Q9Z2U0 | Psma7 | 28 kDa | 3 C | 1.244 | 0.27316 |
| Mitotic checkpoint protein BUB3 | Q9WVA3 | Bub3 | 37 kDa | 7 C | 1.244 | 0.28741 |
| Cystathionine gamma-lyase | Q8VCN5 | Cth | 44 kDa | 11 C | 1.244 | 0.39766 |
| Calreticulin | P14211 | Calr | 48 kDa | 6 C | 1.243 | 0.25955 |
| Galectin-9 | O08573 (+2) | Lgals9 | 40 kDa | 7 C | 1.240 | 0.19301 |
| Proteasome subunit alpha type-4 | Q9R1P0 | Psma4 | 29 kDa | 5 C | 1.236 | 0.12552 |
| Carbonic anhydrase 2 | P00920 | Ca2 | 29 kDa | 2 C | 1.236 | 0.17917 |
| Choline dehydrogenase, mitochondrial | Q8BJ64 | Chdh | 66 kDa | 12 C | 1.236 | 0.32819 |
| Regulator of microtubule dynamics protein 1 | Q9DCV4 | Rmdn1 | 35 kDa | 5 C | 1.236 | 0.32819 |

| | | | | | | |
|---|---|---|---|---|---|---|
| Serine--tRNA ligase, cytoplasmic | P26638 | Sars | 58 kDa | 9 C | 1.236 | 0.56421 |
| 4-trimethylaminobutyraldehyde dehydrogenase | Q9JLJ2 | Aldh9a1 | 54 kDa | 17 C | 1.235 | 0.15318 |
| Ubiquitin-conjugating enzyme E2 L3 | P68037 | Ube2l3 | 18 kDa | 3 C | 1.234 | 0.23946 |
| Glutathione peroxidase 1 | P11352 | Gpx1 | 22 kDa | 4 C | 1.233 | 0.1466 |
| Adenylate kinase 2, mitochondrial | Q9WTP6 | Ak2 | 26 kDa | 5 C | 1.231 | 0.11765 |
| Annexin A2 | P07356 | Anxa2 | 39 kDa | 5 C | 1.231 | 0.69041 |
| Fructose-bisphosphate aldolase A | P05064 | Aldoa | 39 kDa | 8 C | 1.229 | 0.07603 |
| Cytochrome b-c1 complex subunit 1, mitochondrial | Q9CZ13 | Uqcrc1 | 53 kDa | 11 C | 1.224 | 0.5261 |
| Proteasome activator complex subunit 1 | P97371 | Psme1 | 29 kDa | 3 C | 1.224 | 0.60187 |
| Carboxylesterase 1D | Q8VCT4 | Ces1d | 62 kDa | 5 C | 1.222 | 0.00913 |
| Peroxiredoxin-6 | O08709 | Prdx6 | 25 kDa | 2 C | 1.221 | 0.18887 |
| Proteasome subunit beta type-1 | O09061 | Psmb1 | 26 kDa | 5 C | 1.219 | 0.11728 |
| Secernin-2 | Q8VCA8 | Scrn2 | 47 kDa | 10 C | 1.219 | 0.46181 |
| Persulfide dioxygenase ETHE1, mitochondrial | Q9DCM0 | Ethe1 | 28 kDa | 9 C | 1.217 | 0.5762 |
| Methylthioribulose-1-phosphate dehydratase | Q9WVQ5 | Apip | 27 kDa | 11 C | 1.213 | 0.08238 |
| Toll-interacting protein | Q9QZ06 | Tollip | 30 kDa | 4 C | 1.213 | 0.60547 |
| Elongation factor 1-alpha 1 | P10126 | Eef1a1 | 50 kDa | 6 C | 1.213 | 0.15604 |
| Nucleoside diphosphate kinase A | P15532 | Nme1 | 17 kDa | 2 C | 1.211 | 0.16334 |
| Fatty acid-binding protein, brain | P51880 | Fabp7 | 15 kDa | 5 C | 1.210 | 0.67899 |
| Aldehyde dehydrogenase family 8 member A1 | Q8BH00 | Aldh8a1 | 54 kDa | 13 C | 1.207 | 0.71216 |
| Stress-70 protein, mitochondrial | P38647 | Hspa9 | 73 kDa | 5 C | 1.206 | 0.37637 |
| Inorganic pyrophosphatase 2, mitochondrial | Q91VM9 | Ppa2 | 38 kDa | 8 C | 1.206 | 0.21515 |
| Protein disulfide-isomerase A3 | P27773 | Pdia3 | 57 kDa | 8 C | 1.203 | 0.12999 |
| Fumarylacetoacetase | P35505 | Fah | 46 kDa | 6 C | 1.203 | 0.07594 |
| Isoform 2 of DAZ-associated protein 1 | Q9JII5-2 | Dazap1 | 43 kDa | 4 C | 1.200 | 0.08762 |
| Serine/threonine-protein phosphatase 5 | Q60676 | Ppp5c | 57 kDa | 11 C | 1.200 | 0.34334 |
| Carboxylesterase 1F | Q91WU0 | Ces1f | 62 kDa | 7 C | 1.200 | 0.06221 |
| Nucleolysin TIAR | P70318 | Tial1 | 43 kDa | 6 C | 1.200 | 0.20584 |
| Sulfite oxidase, mitochondrial | Q8R086 | Suox | 61 kDa | 9 C | 1.200 | 0.20636 |
| 14-3-3 protein epsilon | P62259 | Ywhae | 29 kDa | 3 C | 1.200 | 0.21116 |
| FAD-linked sulfhydryl oxidase ALR | P56213 | Gfer | 23 kDa | 8 C | 1.200 | 0.21648 |
| 40S ribosomal protein SA | P14206 | Rpsa | 33 kDa | 2 C | 1.200 | 0.24904 |
| Hydroxyacylglutathione hydrolase, mitochondrial | Q99KB8 | Hagh | 34 kDa | 8 C | 1.200 | 0.27561 |
| Hsc70-interacting protein | Q99L47 | St13 | 42 kDa | 3 C | 1.200 | 0.32326 |
| Ribosyldihydronicotinamide dehydrogenase [quinone] | Q9JI75 | Nqo2 | 26 kDa | 4 C | 1.200 | 0.32819 |
| Copper chaperone for superoxide dismutase | Q9WU84 | Ccs | 29 kDa | 10 C | 1.200 | 0.35582 |
| NADH dehydrogenase [ubiquinone] 1 alpha subcomplex subunit 2 | Q9CQ75 | Ndufa2 | 11 kDa | 2 C | 1.200 | 0.40708 |
| Cleavage and polyadenylation specificity factor subunit 5 | Q9CQF3 | Nudt21 | 26 kDa | 1 C | 1.200 | 0.48033 |
| Coatomer subunit epsilon | O89079 | Cope | 35 kDa | 4 C | 1.200 | 0.82713 |
| Tubulin beta-5 chain | P99024 | Tubb5 | 50 kDa | 8 C | 1.200 | 0.82713 |
| Dihydropyrimidinase | Q9EQF5 | Dpys | 57 kDa | 9 C | 1.197 | 0.40381 |
| Acetyl-CoA acetyltransferase, mitochondrial | Q8QZT1 | Acat1 | 45 kDa | 6 C | 1.195 | 0.05367 |
| Glutaredoxin-1 | Q9QUH0 | Glrx | 12 kDa | 5 C | 1.193 | 0.8297 |
| Isoform 2 of Tropomyosin alpha-3 chain | P21107-2 | Tpm3 | 29 kDa | 1 C | 1.192 | 0.19614 |
| Isoform 3 of Heterogeneous nuclear ribonucleoprotein D0 | Q60668-3 | Hnrnpd | 33 kDa | 3 C | 1.191 | 0.30573 |
| Cell division control protein 42 homolog | P60766 | Cdc42 | 21 kDa | 7 C | 1.190 | 0.72398 |
| Thioredoxin-dependent peroxide reductase, mitochondrial | P20108 | Prdx3 | 28 kDa | 4 C | 1.190 | 0.23775 |
| Proteasome subunit alpha type-1 | Q9R1P4 | Psma1 | 30 kDa | 5 C | 1.189 | 0.47389 |
| Major urinary protein 2 | P11589 | Mup2 | 21 kDa | 5 C | 1.188 | 0.43477 |
| Histidine ammonia-lyase | P35492 | Hal | 72 kDa | 12 C | 1.187 | 0.26877 |
| Exosome complex exonuclease RRP42 | Q9D0M0 | Exosc7 | 32 kDa | 12 C | 1.185 | 0.66639 |
| 3-ketoacyl-CoA thiolase A, peroxisomal | Q921H8 | Acaa1a | 44 kDa | 8 C | 1.182 | 0.30387 |
| Transketolase | P40142 | Tkt | 68 kDa | 12 C | 1.182 | 0.17747 |
| Serine/threonine-protein phosphatase PP1-alpha catalytic subunit | P62137 | Ppp1ca | 38 kDa | 13 C | 1.181 | 0.13939 |
| 4-hydroxyphenylpyruvate dioxygenase | P49429 | Hpd | 45 kDa | 4 C | 1.180 | 0.19924 |
| Delta-aminolevulinic acid dehydratase | P10518 | Alad | 36 kDa | 8 C | 1.178 | 0.21242 |
| Heterogeneous nuclear ribonucleoproteins A2/B1 | O88569 | Hnrnpa2b1 | 37 kDa | 1 C | 1.178 | 0.18024 |
| 14-3-3 protein gamma | P61982 | Ywhag | 28 kDa | 3 C | 1.173 | 0.23129 |
| Glutaryl-CoA dehydrogenase, mitochondrial | Q60759 | Gcdh | 49 kDa | 9 C | 1.173 | 0.51455 |
| Serotransferrin | Q921I1 | Tf | 77 kDa | 38 C | 1.173 | 0.05541 |
| Alcohol dehydrogenase 1 | P00329 | Adh1 | 40 kDa | 15 C | 1.171 | 0.40708 |
| Superoxide dismutase [Mn], mitochondrial | P09671 | Sod2 | 25 kDa | 4 C | 1.169 | 0.4247 |
| Cytochrome c, somatic | P62897 | Cycs | 12 kDa | 2 C | 1.168 | 0.29784 |
| Proteasome subunit beta type-7 | P70195 | Psmb7 | 30 kDa | 6 C | 1.167 | 0.64702 |
| Phosphotriesterase-related protein | Q60866 | Pter | 39 kDa | 6 C | 1.165 | 0.46555 |
| Pterin-4-alpha-carbinolamine dehydratase | P61458 | Pcbd1 | 12 kDa | 1 C | 1.164 | 0.4758 |
| Antithrombin-III | P32261 | Serpinc1 | 52 kDa | 9 C | 1.164 | 0.53892 |
| Eukaryotic translation initiation factor 6 | O55135 | Eif6 | 27 kDa | 8 C | 1.164 | 0.66626 |
| Eukaryotic translation initiation factor 4H | Q9WUK2 | Eif4h | 27 kDa | 1 C | 1.160 | 0.60931 |
| Carbonic anhydrase 3 | P16015 | Ca3 | 29 kDa | 5 C | 1.158 | 0.08858 |
| Succinate-semialdehyde dehydrogenase, mitochondrial | Q8BWF0 | Aldh5a1 | 56 kDa | 10 C | 1.157 | 0.18837 |
| Cytochrome c oxidase subunit 5A, mitochondrial | P12787 | Cox5a | 16 kDa | 4 C | 1.156 | 0.50972 |
| Sorbitol dehydrogenase | Q64442 | Sord | 38 kDa | 10 C | 1.155 | 0.18133 |
| Phosphatidylethanolamine-binding protein 1 | P70296 | Pebp1 | 21 kDa | 3 C | 1.151 | 0.30663 |
| Dihydropyrimidinase-related protein 3 | Q62188 | Dpysl3 | 62 kDa | 7 C | 1.150 | 0.50235 |
| Heterogeneous nuclear ribonucleoprotein D-like | Q9Z130 | Hnrnpdl | 34 kDa | 3 C | 1.150 | 0.63824 |
| Heterogeneous nuclear ribonucleoprotein F | Q9Z2X1 | Hnrnpf | 46 kDa | 6 C | 1.149 | 0.4897 |
| Drebrin-like protein | Q62418 | Dbnl | 49 kDa | 5 C | 1.148 | 0.41125 |

| | | | | | | |
|---|---|---|---|---|---|---|
| Malate dehydrogenase, cytoplasmic | P14152 | Mdh1 | 37 kDa | 3 C | 1.148 | 0.42896 |
| Fibrinogen beta chain | Q8K0E8 | Fgb | 55 kDa | 12 C | 1.146 | 0.32309 |
| Glutaredoxin-3 | Q9CQM9 | Glrx3 | 38 kDa | 5 C | 1.145 | 0.22208 |
| 3-hydroxyisobutyrate dehydrogenase, mitochondrial | Q99L13 | Hibadh | 35 kDa | 12 C | 1.143 | 0.13059 |
| Oligoribonuclease, mitochondrial | Q9D8S4 | Rexo2 | 27 kDa | 4 C | 1.143 | 0.29235 |
| PDZ and LIM domain protein 1 | O70400 | Pdlim1 | 36 kDa | 8 C | 1.143 | 0.56202 |
| Protein transport protein Sec31A | Q3UPL0 | Sec31a | 134 kDa | 18 C | 1.143 | 0.57083 |
| Exosome complex component MTR3 | Q8BTW3 | Exosc6 | 28 kDa | 6 C | 1.143 | 0.57083 |
| Ig lambda-1 chain C region | P01843 | | 12 kDa | 3 C | 1.143 | 0.57083 |
| Peroxisomal acyl-coenzyme A oxidase 1 | Q9R0H0 | Acox1 | 75 kDa | 7 C | 1.141 | 0.11625 |
| Glutathione reductase, mitochondrial | P47791 | Gsr | 54 kDa | 11 C | 1.139 | 0.31355 |
| Protein ABHD14B | Q8VCR7 | Abhd14b | 22 kDa | 2 C | 1.138 | 0.12687 |
| Carbonic anhydrase 5A, mitochondrial | P23589 | Ca5a | 34 kDa | 7 C | 1.138 | 0.37757 |
| Cystathionine beta-synthase | Q91WT9 | Cbs | 62 kDa | 13 C | 1.138 | 0.59746 |
| Serine hydroxymethyltransferase, cytosolic | P50431 | Shmt1 | 53 kDa | 10 C | 1.138 | 0.36202 |
| Thioredoxin-like protein 1 | Q8CDN6 | Txnl1 | 32 kDa | 7 C | 1.138 | 0.32629 |
| Bifunctional epoxide hydrolase 2 | P34914 | Ephx2 | 63 kDa | 10 C | 1.137 | 0.22591 |
| Eukaryotic peptide chain release factor GTP-binding subunit ERF3A | Q8R050 (+1) | Gspt1 | 69 kDa | 14 C | 1.136 | 0.78655 |
| 78 kDa glucose-regulated protein | P20029 | Hspa5 | 72 kDa | 1 C | 1.134 | 0.19937 |
| 60S ribosomal protein L30 | P62889 | Rpl30 | 13 kDa | 3 C | 1.133 | 0.40708 |
| Isoform 2 of Alpha-aminoadipic semialdehyde dehydrogenase | Q9DBF1-2 | Aldh7a1 | 56 kDa | 9 C | 1.133 | 0.20212 |
| Acetyl-CoA acetyltransferase, cytosolic | Q8CAY6 | Acat2 | 41 kDa | 8 C | 1.133 | 0.20171 |
| Cytosolic non-specific dipeptidase | Q9D1A2 | Cndp2 | 53 kDa | 8 C | 1.129 | 0.27316 |
| Adrenodoxin, mitochondrial | P46656 | Fdx1 | 20 kDa | 8 C | 1.129 | 0.50538 |
| Calcyclin-binding protein | Q9CXW3 | Cacybp | 27 kDa | 2 C | 1.129 | 0.86271 |
| Peptidyl-prolyl cis-trans isomerase NIMA-interacting 1 | Q9QUR7 | Pin1 | 18 kDa | 2 C | 1.129 | 0.87198 |
| 5-hydroxyisourate hydrolase | Q9CRB3 | Urah | 14 kDa | 3 C | 1.126 | 0.32316 |
| Inorganic pyrophosphatase | Q9D819 | Ppa1 | 33 kDa | 8 C | 1.126 | 0.4423 |
| Proteasome subunit beta type-3 | Q9R1P1 | Psmb3 | 23 kDa | 5 C | 1.126 | 0.53151 |
| 4-hydroxy-2-oxoglutarate aldolase, mitochondrial | Q9DCU9 | Hoga1 | 35 kDa | 6 C | 1.125 | 0.48033 |
| Probable imidazolonepropionase | Q9DBA8 | Amdhd1 | 46 kDa | 11 C | 1.123 | 0.19113 |
| Quinone oxidoreductase | P47199 | Cryz | 35 kDa | 5 C | 1.122 | 0.1932 |
| Protein disulfide-isomerase A4 | P08003 | Pdia4 | 72 kDa | 6 C | 1.121 | 0.10616 |
| Transaldolase | Q93092 | Taldo1 | 37 kDa | 3 C | 1.120 | 0.29235 |
| Microsomal glutathione S-transferase 1 | Q91VS7 | Mgst1 | 18 kDa | 1 C | 1.120 | 0.70009 |
| Ethanolamine-phosphate phospho-lyase | Q8BWU8 | Etnppl | 55 kDa | 10 C | 1.120 | 0.76501 |
| NADH dehydrogenase [ubiquinone] 1 alpha subcomplex subunit 7 | Q9Z1P6 | Ndufa7 | 13 kDa | 1 C | 1.120 | 0.68453 |
| 28S ribosomal protein S28, mitochondrial | Q9CY16 | Mrps28 | 21 kDa | 3 C | 1.120 | 0.68453 |
| Prolyl endopeptidase | Q9QUR6 | Prep | 81 kDa | 17 C | 1.120 | 0.86293 |
| Optineurin | Q8K3K8 | Optn | 67 kDa | 12 C | 1.114 | 0.83985 |
| Na(+)/H(+) exchange regulatory cofactor NHE-RF2 | Q9JHL1 | Slc9a3r2 | 37 kDa | 6 C | 1.114 | 0.83985 |
| 60S ribosomal protein L5 | P47962 | Rpl5 | 34 kDa | 4 C | 1.113 | 0.89744 |
| Eukaryotic translation initiation factor 1 | P48024 | Eif1 | 13 kDa | 2 C | 1.110 | 0.77292 |
| Ig gamma-1 chain C region secreted form | P01868 (+1) | Ighg1 | 36 kDa | 12 C | 1.110 | 0.90231 |
| 60S ribosomal protein L12 | P35979 | Rpl12 | 18 kDa | 3 C | 1.110 | 0.36446 |
| Aldo-keto reductase family 1 member C13 | Q8VC28 | Akr1c13 | 37 kDa | 10 C | 1.108 | 0.50972 |
| Adenosine kinase | P55264 | Adk | 40 kDa | 6 C | 1.107 | 0.46011 |
| Heat shock cognate 71 kDa protein | P63017 | Hspa8 | 71 kDa | 4 C | 1.106 | 0.60993 |
| TAR DNA-binding protein 43 | Q921F2 | Tardbp | 45 kDa | 7 C | 1.105 | 0.59936 |
| Polyadenylate-binding protein 1 | P29341 | Pabpc1 | 71 kDa | 4 C | 1.105 | 0.49035 |
| Aconitate hydratase, mitochondrial | Q99KI0 | Aco2 | 85 kDa | 13 C | 1.102 | 0.24941 |
| 40S ribosomal protein S28 | P62858 | Rps28 | 8 kDa | 1 C | 1.100 | 0.65169 |
| Thioredoxin domain-containing protein 12 | Q9CQU0 | Txndc12 | 19 kDa | 3 C | 1.100 | 0.72943 |
| Myotrophin | P62774 | Mtpn | 13 kDa | 3 C | 1.100 | 0.77107 |
| Serine beta-lactamase-like protein LACTB, mitochondrial | Q9EP89 | Lactb | 61 kDa | 4 C | 1.100 | 0.70955 |
| Alpha-taxilin | Q6PAM1 | Txlna | 62 kDa | 8 C | 1.100 | 0.93656 |
| Hydroxymethylglutaryl-CoA lyase, mitochondrial | P38060 | Hmgcl | 34 kDa | 8 C | 1.099 | 0.20741 |
| Cathepsin D | P18242 | Ctsd | 45 kDa | 8 C | 1.098 | 0.80528 |
| LIM and SH3 domain protein 1 | Q61792 | Lasp1 | 30 kDa | 7 C | 1.098 | 0.54104 |
| Heat shock protein HSP 90-beta | P11499 | Hsp90ab1 | 83 kDa | 6 C | 1.096 | 0.86824 |
| Selenocysteine lyase | Q9JLI6 | Scly | 47 kDa | 8 C | 1.095 | 0.70009 |
| Dynein light chain 2, cytoplasmic | Q9D0M5 | Dynll2 | 10 kDa | 2 C | 1.095 | 0.72537 |
| Aldehyde dehydrogenase, mitochondrial | P47738 | Aldh2 | 57 kDa | 9 C | 1.093 | 0.53907 |
| Lactoylglutathione lyase | Q9CPU0 | Glo1 | 21 kDa | 3 C | 1.092 | 0.60122 |
| Kynurenine--oxoglutarate transaminase 3 | Q71RI9 (+1) | Kyat3 | 51 kDa | 10 C | 1.091 | 0.48033 |
| Protein disulfide-isomerase | P09103 | P4hb | 57 kDa | 7 C | 1.090 | 0.41749 |
| Ig kappa chain C region | P01837 | | 12 kDa | 3 C | 1.087 | 0.68873 |
| Kynureninase | Q9CXF0 | Kynu | 52 kDa | 8 C | 1.086 | 0.56303 |
| Protein phosphatase 1B | P36993 | Ppm1b | 43 kDa | 12 C | 1.086 | 0.70955 |
| Crk-like protein | P47941 | Crkl | 34 kDa | 2 C | 1.086 | 0.78724 |
| Actin-related protein 2/3 complex subunit 1A | Q9R0Q6 | Arpc1a | 42 kDa | 10 C | 1.086 | 0.79797 |
| Spermidine synthase | Q64674 | Srm | 34 kDa | 10 C | 1.084 | 0.88833 |
| Ig mu chain C region | P01872 | Ighm | 50 kDa | 19 C | 1.082 | 0.73384 |
| Glutathione S-transferase Mu 1 | P10649 | Gstm1 | 26 kDa | 2 C | 1.080 | 0.63295 |
| Cysteine-rich protein 2 | Q9DCT8 | Crip2 | 23 kDa | 14 C | 1.077 | 0.66466 |
| S-methylmethionine--homocysteine S-methyltransferase BHMT2 | Q91WS4 | Bhmt2 | 40 kDa | 10 C | 1.076 | 0.69619 |
| Diphosphoinositol polyphosphate phosphohydrolase 2 | Q8R2U6 | Nudt4 | 20 kDa | 4 C | 1.075 | 0.93862 |
| Isoform Cytoplasmic+peroxisomal of Peroxiredoxin-5, mitochondrial | P99029-2 | Prdx5 | 17 kDa | 6 C | 1.073 | 0.52624 |

| | | | | | | |
|---|---|---|---|---|---|---|
| Cytosolic purine 5'-nucleotidase | Q3V1L4 | Nt5c2 | 65 kDa | 8 C | 1.073 | 0.85901 |
| Profilin-1 | P62962 | Pfn1 | 15 kDa | 3 C | 1.073 | 0.60931 |
| Methylcrotonoyl-CoA carboxylase beta chain, mitochondrial | Q3ULD5 | Mccc2 | 61 kDa | 10 C | 1.070 | 0.19798 |
| Cytochrome b-c1 complex subunit Rieske, mitochondrial | Q9CR68 | Uqcrfs1 | 29 kDa | 5 C | 1.067 | 0.62055 |
| 3-hydroxyanthranilate 3,4-dioxygenase | Q78JT3 | Haao | 33 kDa | 3 C | 1.067 | 0.64685 |
| Glutathione synthetase | P51855 | Gss | 52 kDa | 4 C | 1.067 | 0.72327 |
| Carboxymethylenebutenolidase homolog | Q8R1G2 | Cmbl | 28 kDa | 6 C | 1.067 | 0.75026 |
| Ig kappa chain V-II region 26-10 | P01631 | | 12 kDa | 2 C | 1.067 | 0.77107 |
| Proteasome subunit beta type-6 | Q60692 | Psmb6 | 25 kDa | 4 C | 1.067 | 0.81532 |
| Serine/threonine-protein phosphatase 2A catalytic subunit alpha isoform | P63330 | Ppp2ca | 36 kDa | 10 C | 1.067 | 0.82494 |
| Platelet-activating factor acetylhydrolase IB subunit beta | Q61206 | Pafah1b2 | 26 kDa | 3 C | 1.067 | 0.84337 |
| Vigilin | Q8VDJ3 | Hdlbp | 142 kDa | 10 C | 1.067 | 0.88641 |
| PDZ and LIM domain protein 5 | Q8CI51 | Pdlim5 | 63 kDa | 22 C | 1.067 | 0.90718 |
| Mitochondrial peptide methionine sulfoxide reductase | Q9D6Y7 | Msra | 26 kDa | 4 C | 1.065 | 0.28067 |
| Fructose-1,6-bisphosphatase 1 | Q9QXD6 | Fbp1 | 37 kDa | 7 C | 1.063 | 0.35045 |
| 3-ketoacyl-CoA thiolase B, peroxisomal | Q8VCH0 | Acaa1b | 44 kDa | 9 C | 1.062 | 0.71839 |
| WD repeat domain phosphoinositide-interacting protein 3 | Q9CR39 | Wdr45b | 38 kDa | 15 C | 1.062 | 0.90712 |
| Calnexin | P35564 | Canx | 67 kDa | 7 C | 1.062 | 0.91547 |
| Selenide, water dikinase 1 | Q8BH69 | Sephs1 | 43 kDa | 9 C | 1.062 | 0.73583 |
| Isocitrate dehydrogenase [NADP] cytoplasmic | O88844 | Idh1 | 47 kDa | 7 C | 1.060 | 0.57928 |
| Succinate dehydrogenase [ubiquinone] flavoprotein subunit, mitochondrial | Q8K2B3 | Sdha | 73 kDa | 19 C | 1.060 | 0.49107 |
| Ester hydrolase C11orf54 homolog | Q91V76 | | 35 kDa | 8 C | 1.059 | 0.70175 |
| 4-aminobutyrate aminotransferase, mitochondrial | P61922 | Abat | 56 kDa | 12 C | 1.058 | 0.79101 |
| Proteasome subunit beta type-2 | Q9R1P3 | Psmb2 | 23 kDa | 3 C | 1.056 | 0.6394 |
| Serine/threonine-protein phosphatase PP1-beta catalytic subunit | P62141 | Ppp1cb | 37 kDa | 14 C | 1.055 | 0.6537 |
| Electron transfer flavoprotein subunit alpha, mitochondrial | Q99LC5 | Etfa | 35 kDa | 6 C | 1.055 | 0.77427 |
| Peptidyl-prolyl cis-trans isomerase A | P17742 | Ppia | 18 kDa | 3 C | 1.052 | 0.54914 |
| Peroxisomal multifunctional enzyme type 2 | P51660 | Hsd17b4 | 79 kDa | 9 C | 1.051 | 0.65463 |
| Non-specific lipid-transfer protein | P32020 | Scp2 | 59 kDa | 11 C | 1.051 | 0.70777 |
| COP9 signalosome complex subunit 3 | O88543 | Cops3 | 48 kDa | 10 C | 1.050 | 0.94546 |
| RNA-binding motif, single-stranded-interacting protein 1 | Q91W59 | Rbms1 | 44 kDa | 4 C | 1.050 | 0.94546 |
| Molybdopterin synthase catalytic subunit | Q9Z223 | Mocs2 | 21 kDa | 4 C | 1.049 | 0.91917 |
| Septin-2 | P42208 | 43345 | 42 kDa | 8 C | 1.048 | 0.86574 |
| Methionine adenosyltransferase 2 subunit beta | Q99LB6 (+1) | Mat2b | 37 kDa | 7 C | 1.046 | 0.7746 |
| Endoplasmic reticulum resident protein 29 | P57759 | Erp29 | 29 kDa | 1 C | 1.046 | 0.80014 |
| 40S ribosomal protein S21 | Q9CQR2 | Rps21 | 9 kDa | 2 C | 1.043 | 0.8348 |
| Fucose mutarotase | Q8R2K1 | Fuom | 17 kDa | 3 C | 1.043 | 0.84211 |
| Isoform 2 of Cytosol aminopeptidase | Q9CPY7-2 | Lap3 | 53 kDa | 7 C | 1.042 | 0.58973 |
| Tetratricopeptide repeat protein 38 | A3KMP2 | Ttc38 | 52 kDa | 9 C | 1.042 | 0.73583 |
| Apoptosis-inducing factor 1, mitochondrial | Q9Z0X1 | Aifm1 | 67 kDa | 4 C | 1.042 | 0.62055 |
| Presequence protease, mitochondrial | Q8K411 (+1) | Pitrm1 | 117 kDa | 20 C | 1.040 | 0.89433 |
| N-myc-interactor | O35309 | Nmi | 35 kDa | 8 C | 1.040 | 0.89433 |
| Calpain small subunit 1 | O88456 | Capns1 | 28 kDa | 2 C | 1.040 | 0.89433 |
| Exosome complex component CSL4 | Q9DAA6 | Exosc1 | 21 kDa | 6 C | 1.040 | 0.93617 |
| Endoribonuclease LACTB2 | Q99KR3 | Lactb2 | 33 kDa | 5 C | 1.037 | 0.70502 |
| Hydroxymethylglutaryl-CoA synthase, cytoplasmic | Q8JZK9 | Hmgcs1 | 58 kDa | 11 C | 1.037 | 0.87189 |
| Vitamin D-binding protein | P21614 | Gc | 54 kDa | 28 C | 1.034 | 0.81652 |
| Neutral alpha-glucosidase AB | Q8BHN3 | Ganab | 107 kDa | 8 C | 1.034 | 0.87539 |
| Pyridoxal kinase | Q8K183 | Pdxk | 35 kDa | 5 C | 1.032 | 0.83879 |
| Selenide, water dikinase 2 | P97364 | Sephs2 | 48 kDa | 7 C | 1.030 | 0.75157 |
| Destrin | Q9R0P5 | Dstn | 19 kDa | 6 C | 1.029 | 0.80723 |
| Hsp90 co-chaperone Cdc37 | Q61081 | Cdc37 | 45 kDa | 9 C | 1.029 | 0.86758 |
| UV excision repair protein RAD23 homolog A | P54726 | Rad23a | 40 kDa | 1 C | 1.029 | 0.91958 |
| Mitochondrial intermembrane space import and assembly protein 40 | Q8VEA4 | Chchd4 | 16 kDa | 7 C | 1.029 | 0.93563 |
| UV excision repair protein RAD23 homolog B | P54728 | Rad23b | 44 kDa | 1 C | 1.029 | 0.87189 |
| Complement component C8 gamma chain | Q8VCG4 | C8g | 23 kDa | 3 C | 1.026 | 0.77236 |
| Protein disulfide-isomerase A6 | Q922R8 | Pdia6 | 48 kDa | 7 C | 1.022 | 0.85285 |
| Molybdenum cofactor biosynthesis protein 1 | Q5RKZ7 | Mocs1 | 70 kDa | 15 C | 1.022 | 0.87862 |
| Proteasome subunit beta type-4 | P99026 | Psmb4 | 29 kDa | 2 C | 1.022 | 0.89433 |
| Catalase | P24270 | Cat | 60 kDa | 5 C | 1.021 | 0.87688 |
| Nucleoside diphosphate kinase B | Q01768 | Nme2 | 17 kDa | 2 C | 1.020 | 0.82909 |
| Cofilin-1 | P18760 | Cfl1 | 19 kDa | 4 C | 1.020 | 0.88565 |
| Toll-like receptor 3 | Q99MB1 | Tlr3 | 104 kDa | 15 C | 1.020 | 0.95486 |
| Glutamate--cysteine ligase catalytic subunit | P97494 | Gclc | 73 kDa | 14 C | 1.018 | 0.98464 |
| Thioredoxin domain-containing protein 17 | Q9CQM5 | Txndc17 | 14 kDa | 6 C | 1.018 | 0.93563 |
| Eukaryotic translation initiation factor 4B | Q8BGD9 | Eif4b | 69 kDa | 3 C | 1.018 | 0.96151 |
| Transitional endoplasmic reticulum ATPase | Q01853 | Vcp | 89 kDa | 12 C | 1.017 | 0.90461 |
| Microtubule-associated protein RP/EB family member 1 | Q61166 | Mapre1 | 30 kDa | 3 C | 1.015 | 0.98328 |
| 40S ribosomal protein S20 | P60867 | Rps20 | 13 kDa | 2 C | 1.013 | 0.91958 |
| GTP cyclohydrolase 1 feedback regulatory protein | P99025 | Gchfr | 10 kDa | 2 C | 1.013 | 0.9509 |
| Far upstream element-binding protein 1 | Q91WJ8 | Fubp1 | 69 kDa | 3 C | 1.013 | 0.96151 |
| Alpha-1-antitrypsin 1-4 | Q00897 | Serpina1d | 46 kDa | 3 C | 1.011 | 0.97108 |
| Cordon-bleu protein-like 1 | Q3UMF0 | Cobll1 | 137 kDa | 10 C | 1.011 | 0.97207 |
| Adenylyl cyclase-associated protein 1 | P40124 | Cap1 | 52 kDa | 6 C | 1.010 | 0.98354 |
| Isoform 2 of Gelsolin | P13020-2 | Gsn | 81 kDa | 7 C | 1.009 | 0.97523 |
| Scaffold attachment factor B1 | D3YXK2 | Safb | 105 kDa | 9 C | 1.007 | 0.98621 |
| Stress-induced-phosphoprotein 1 | Q60864 | Stip1 | 63 kDa | 11 C | 1.002 | 0.97276 |
| Carboxylesterase 3A | Q63880 (+1) | Ces3a | 63 kDa | 7 C | 1.000 | 1 |

| | | | | | | |
|---|---|---|---|---|---|---|
| Polymerase delta-interacting protein 2 | Q91VA6 | Poldip2 | 42 kDa | 4 C | 1.000 | 1 |
| S-adenosylmethionine synthase isoform type-2 | Q3THS6 | Mat2a | 44 kDa | 6 C | 1.000 | 1 |
| Stromal cell-derived factor 2-like protein 1 | Q9ESP1 | Sdf2l1 | 24 kDa | 4 C | 1.000 | 1 |
| Mitochondrial antiviral-signaling protein | Q8VCF0 | Mavs | 53 kDa | 8 C | 1.000 | 1 |
| 26S proteasome non-ATPase regulatory subunit 9 | Q9CR00 | Psmd9 | 25 kDa | 3 C | 1.000 | 1 |
| Vinculin | Q64727 | Vcl | 117 kDa | 10 C | 0.997 | 0.9875 |
| Glutathione S-transferase Mu 2 | P15626 | Gstm2 | 26 kDa | 3 C | 0.995 | 0.98549 |
| N-acyl-aromatic-L-amino acid amidohydrolase (carboxylate-forming) | Q91XE4 | Acy3 | 35 kDa | 8 C | 0.991 | 0.98362 |
| Fibrinogen alpha chain | E9PV24 | Fga | 87 kDa | 13 C | 0.990 | 0.96465 |
| Glycine N-methyltransferase | Q9QXF8 | Gnmt | 33 kDa | 8 C | 0.990 | 0.9239 |
| Selenoprotein F | Q9ERR7 | Selenof | 18 kDa | 7 C | 0.988 | 0.93563 |
| Estradiol 17 beta-dehydrogenase 5 | P70694 | Akr1c6 | 37 kDa | 7 C | 0.987 | 0.93834 |
| Aspartate aminotransferase, cytoplasmic | P05201 | Got1 | 46 kDa | 5 C | 0.985 | 0.9208 |
| Fatty acid-binding protein, adipocyte | P04117 | Fabp4 | 15 kDa | 2 C | 0.985 | 0.94479 |
| Caspase-3 | P70677 | Casp3 | 31 kDa | 8 C | 0.985 | 0.96377 |
| Galactose-1-phosphate uridylyltransferase | Q03249 | Galt | 43 kDa | 7 C | 0.985 | 0.9673 |
| Protein DJ-1 | Q99LX0 | Park7 | 20 kDa | 4 C | 0.983 | 0.87174 |
| Hypoxanthine-guanine phosphoribosyltransferase | P00493 | Hprt1 | 25 kDa | 4 C | 0.982 | 0.86758 |
| Calponin-3 | Q9DAW9 | Cnn3 | 36 kDa | 3 C | 0.982 | 0.95328 |
| Betaine--homocysteine S-methyltransferase 1 | O35490 | Bhmt | 45 kDa | 8 C | 0.981 | 0.84109 |
| Carbamoyl-phosphate synthase [ammonia], mitochondrial | Q8C196 | Cps1 | 165 kDa | 21 C | 0.980 | 0.95808 |
| Enoyl-CoA hydratase, mitochondrial | Q8BH95 | Echs1 | 31 kDa | 7 C | 0.979 | 0.84211 |
| Probable ATP-dependent RNA helicase DDX17 | Q501J6 | Ddx17 | 72 kDa | 11 C | 0.978 | 0.95535 |
| GTP cyclohydrolase 1 | Q05915 | Gch1 | 27 kDa | 3 C | 0.978 | 0.92978 |
| Alpha-1-antitrypsin 1-1 | P07758 (+1) | Serpina1a | 46 kDa | 3 C | 0.977 | 0.95532 |
| Sialic acid synthase | Q99J77 | Nans | 40 kDa | 8 C | 0.976 | 0.95002 |
| Ribokinase | Q8R1Q9 | Rbks | 34 kDa | 9 C | 0.976 | 0.96051 |
| Aspartyl aminopeptidase | Q9Z2W0 | Dnpep | 52 kDa | 10 C | 0.975 | 0.82467 |
| Isoaspartyl peptidase/L-asparaginase | Q8C0M9 | Asrgl1 | 34 kDa | 8 C | 0.975 | 0.9095 |
| Iron-sulfur cluster assembly enzyme ISCU, mitochondrial | Q9D7P6 | Iscu | 18 kDa | 4 C | 0.971 | 0.87862 |
| 6-phosphofructo-2-kinase/fructose-2,6-bisphosphatase 1 | P70266 | Pfkfb1 | 55 kDa | 10 C | 0.970 | 0.87862 |
| Ubiquitin carboxyl-terminal hydrolase 5 | P56399 | Usp5 | 96 kDa | 16 C | 0.967 | 0.88838 |
| Peroxisomal sarcosine oxidase | Q9D826 | Pipox | 44 kDa | 11 C | 0.966 | 0.81652 |
| Isoamyl acetate-hydrolyzing esterase 1 homolog | Q9DB29 | Iah1 | 28 kDa | 8 C | 0.963 | 0.86518 |
| Selenium-binding protein 1 | P17563 | Selenbp1 | 53 kDa | 10 C | 0.962 | 0.71909 |
| Deoxynucleoside triphosphate triphosphohydrolase SAMHD1 | Q60710 | Samhd1 | 73 kDa | 17 C | 0.960 | 0.82798 |
| Growth factor receptor-bound protein 2 | Q60631 | Grb2 | 25 kDa | 2 C | 0.960 | 0.92469 |
| Fructose-bisphosphate aldolase B | Q91Y97 | Aldob | 40 kDa | 8 C | 0.958 | 0.76232 |
| 2-hydroxyacyl-CoA lyase 1 | Q9QXE0 | Hacl1 | 64 kDa | 17 C | 0.958 | 0.73266 |
| Alanine aminotransferase 1 | Q8QZR5 | Gpt | 55 kDa | 14 C | 0.957 | 0.69138 |
| Dihydrolipoyllysine-residue succinyltransferase component of 2-oxoglutarate dehydrogenase complex, mito | Q9D2G2 | Dlst | 49 kDa | 6 C | 0.956 | 0.77879 |
| Plasminogen | P20918 | Plg | 91 kDa | 48 C | 0.955 | 0.8373 |
| Methionine-R-sulfoxide reductase B1 | Q9JLC3 | Msrb1 | 13 kDa | 6 C | 0.953 | 0.92477 |
| DnaJ homolog subfamily B member 11 | Q99KV1 | Dnajb11 | 41 kDa | 5 C | 0.950 | 0.88729 |
| Succinate dehydrogenase [ubiquinone] iron-sulfur subunit, mitochondrial | Q9CQA3 | Sdhb | 32 kDa | 14 C | 0.949 | 0.62756 |
| Isoform 2 of Sorbin and SH3 domain-containing protein 2 | Q3UTJ2-2 | Sorbs2 | 145 kDa | 3 C | 0.947 | 0.87801 |
| Microsomal triglyceride transfer protein large subunit | O08601 | Mttp | 99 kDa | 11 C | 0.945 | 0.40708 |
| Aflatoxin B1 aldehyde reductase member 2 | Q8CG76 | Akr7a2 | 41 kDa | 8 C | 0.945 | 0.7746 |
| Homogentisate 1,2-dioxygenase | O09173 | Hgd | 50 kDa | 14 C | 0.944 | 0.51296 |
| Phosphoglycerate mutase 1 | Q9DBJ1 | Pgam1 | 29 kDa | 2 C | 0.944 | 0.70895 |
| S-adenosylmethionine synthase isoform type-1 | Q91X83 | Mat1a | 44 kDa | 10 C | 0.943 | 0.79935 |
| Proteasome subunit beta type-8 | P28063 | Psmb8 | 30 kDa | 5 C | 0.941 | 0.71556 |
| Complement C3 | P01027 | C3 | 186 kDa | 27 C | 0.941 | 0.79042 |
| Serine/threonine-protein phosphatase 6 catalytic subunit | Q9CQR6 | Ppp6c | 35 kDa | 12 C | 0.939 | 0.78605 |
| Glycine dehydrogenase (decarboxylating), mitochondrial | Q91W43 | Gldc | 113 kDa | 24 C | 0.937 | 0.72076 |
| 40S ribosomal protein S12 | P63323 | Rps12 | 15 kDa | 7 C | 0.936 | 0.44615 |
| Heterogeneous nuclear ribonucleoprotein H | O35737 | Hnrnph1 | 49 kDa | 1 C | 0.933 | 0.68453 |
| Alpha-methylacyl-CoA racemase | O09174 | Amacr | 42 kDa | 6 C | 0.933 | 0.7746 |
| Triosephosphate isomerase | P17751 | Tpi1 | 32 kDa | 9 C | 0.933 | 0.4758 |
| Serine/threonine-protein phosphatase 2A 55 kDa regulatory subunit B alpha isoform | Q6P1F6 | Ppp2r2a | 52 kDa | 9 C | 0.933 | 0.76248 |
| Ig kappa chain V-V region MOPC 149 | P01636 | | 12 kDa | 2 C | 0.933 | 0.79797 |
| Aspartoacylase | Q8R3P0 | Aspa | 35 kDa | 8 C | 0.933 | 0.81719 |
| Sorting nexin-12 | O70493 | Snx12 | 19 kDa | 3 C | 0.926 | 0.85808 |
| Actin-related protein 2/3 complex subunit 1B | Q9WV32 | Arpc1b | 41 kDa | 14 C | 0.923 | 0.74864 |
| 2-aminoethanethiol dioxygenase | Q6PDY2 | Ado | 28 kDa | 7 C | 0.923 | 0.77236 |
| Bleomycin hydrolase | Q8R016 | Blmh | 53 kDa | 4 C | 0.923 | 0.86391 |
| Far upstream element-binding protein 2 | Q3U0V1 | Khsrp | 77 kDa | 8 C | 0.921 | 0.34262 |
| Protein NDRG2 | Q9QYG0 | Ndrg2 | 41 kDa | 6 C | 0.920 | 0.85338 |
| Mannose-binding protein A | P39039 | Mbl1 | 25 kDa | 8 C | 0.920 | 0.65169 |
| Major urinary protein 20 | Q5FW60 | Mup20 | 21 kDa | 4 C | 0.920 | 0.84337 |
| Isoform Rpn10B of 26S proteasome non-ATPase regulatory subunit 4 | O35226-2 | Psmd4 | 41 kDa | 4 C | 0.919 | 0.78676 |
| SH3 domain-binding glutamic acid-rich-like protein | Q9JJU8 | Sh3bgrl | 13 kDa | 2 C | 0.914 | 0.6059 |
| Lysosome-associated membrane glycoprotein 2 | P17047 | Lamp2 | 46 kDa | 9 C | 0.914 | 0.7746 |
| Ubiquitin-conjugating enzyme E2 variant 2 | Q9D2M8 | Ube2v2 | 16 kDa | 1 C | 0.914 | 0.7746 |
| Hemopexin | Q91X72 | Hpx | 51 kDa | 13 C | 0.914 | 0.61024 |
| Copine-1 | Q8C166 | Cpne1 | 59 kDa | 13 C | 0.913 | 0.86082 |
| Glutathione S-transferase Mu 3 | P19639 | Gstm3 | 26 kDa | 4 C | 0.912 | 0.74864 |
| Actin-related protein 3 | Q99JY9 | Actr3 | 47 kDa | 8 C | 0.912 | 0.34262 |

| | | | | | | |
|---|---|---|---|---|---|---|
| Peptidyl-prolyl cis-trans isomerase D | Q9CR16 | Ppid | 41 kDa | 7 C | 0.912 | 0.40082 |
| Glutathione S-transferase P 1 | P19157 | Gstp1 | 24 kDa | 3 C | 0.911 | 0.30664 |
| Spectrin alpha chain, non-erythrocytic 1 | P16546 | Sptan1 | 285 kDa | 14 C | 0.910 | 0.78788 |
| Copine-3 | Q8BT60 | Cpne3 | 60 kDa | 13 C | 0.909 | 0.53197 |
| Protein phosphatase 1 regulatory subunit 7 | Q3UM45 | Ppp1r7 | 41 kDa | 2 C | 0.908 | 0.4423 |
| Glucosamine 6-phosphate N-acetyltransferase | Q9JK38 | Gnpnat1 | 21 kDa | 6 C | 0.907 | 0.73976 |
| Elongin-B | P62869 | Elob | 13 kDa | 1 C | 0.907 | 0.76627 |
| Eukaryotic translation initiation factor 5A-1 | P63242 | Eif5a | 17 kDa | 4 C | 0.906 | 0.20622 |
| Serine protease inhibitor A3K | P07759 | Serpina3k | 47 kDa | 4 C | 0.905 | 0.55146 |
| Dihydropyrimidinase-related protein 2 | O08553 | Dpysl2 | 62 kDa | 7 C | 0.905 | 0.50958 |
| Ig kappa chain V-V region K2 (Fragment) | P01635 | | 13 kDa | 3 C | 0.900 | 0.40708 |
| Coactosin-like protein | Q9CQI6 | Cotl1 | 16 kDa | 2 C | 0.900 | 0.65169 |
| Thioredoxin domain-containing protein 5 | Q91W90 | Txndc5 | 46 kDa | 12 C | 0.896 | 0.37378 |
| Isoform 2 of Cellular nucleic acid-binding protein | P53996-2 (+ | Cnbp | 19 kDa | 22 C | 0.896 | 0.71046 |
| Splicing factor 1 | Q64213 (+2) | Sf1 | 70 kDa | 4 C | 0.896 | 0.7165 |
| Serine-threonine kinase receptor-associated protein | Q9Z1Z2 | Strap | 38 kDa | 6 C | 0.892 | 0.65849 |
| Methylmalonyl-CoA mutase, mitochondrial | P16332 | Mut | 83 kDa | 8 C | 0.889 | 0.57913 |
| Caspase-7 | P97864 | Casp7 | 34 kDa | 11 C | 0.889 | 0.71556 |
| Glutathione peroxidase 3 | P46412 | Gpx3 | 25 kDa | 3 C | 0.889 | 0.77236 |
| Uncharacterized protein C1orf50 homolog | Q5EBG8 | | 22 kDa | 3 C | 0.889 | 0.77236 |
| Phospholipid hydroperoxide glutathione peroxidase, mitochondrial | O70325 (+1) | Gpx4 | 22 kDa | 10 C | 0.888 | 0.22655 |
| Peptidyl-prolyl cis-trans isomerase FKBP4 | P30416 | Fkbp4 | 52 kDa | 7 C | 0.887 | 0.2361 |
| Cold shock domain-containing protein E1 | Q91W50 | Csde1 | 89 kDa | 15 C | 0.886 | 0.57504 |
| Arsenite methyltransferase | Q91WU5 | As3mt | 42 kDa | 12 C | 0.880 | 0.71666 |
| Golgi reassembly-stacking protein 2 | Q99JX3 | Gorasp2 | 47 kDa | 4 C | 0.873 | 0.44799 |
| Exosome complex component RRP45 | Q9JHI7 | Exosc9 | 49 kDa | 11 C | 0.873 | 0.88989 |
| NHP2-like protein 1 | Q9D0T1 | Snu13 | 14 kDa | 4 C | 0.873 | 0.88989 |
| Trimethyllysine dioxygenase, mitochondrial | Q91ZE0 | Tmlhe | 50 kDa | 11 C | 0.870 | 0.69041 |
| Thioredoxin reductase 1, cytoplasmic | Q9JMH6 | Txnrd1 | 67 kDa | 21 C | 0.869 | 0.32534 |
| Src substrate cortactin | Q60598 | Cttn | 61 kDa | 3 C | 0.869 | 0.29347 |
| BAG family molecular chaperone regulator 5 | Q8CI32 | Bag5 | 51 kDa | 10 C | 0.868 | 0.76199 |
| Fibrinogen gamma chain | Q8VCM7 | Fgg | 49 kDa | 12 C | 0.867 | 0.10133 |
| Peroxiredoxin-4 | O08807 | Prdx4 | 31 kDa | 4 C | 0.866 | 0.22831 |
| Inosine-5'-monophosphate dehydrogenase 2 | P24547 | Impdh2 | 56 kDa | 7 C | 0.865 | 0.72753 |
| Aldehyde oxidase 3 | G3X982 | Aox3 | 147 kDa | 38 C | 0.863 | 0.4172 |
| Septin-11 | Q8C1B7 (+2) | 43354 | 50 kDa | 6 C | 0.862 | 0.37757 |
| Plastin-2 | Q61233 | Lcp1 | 70 kDa | 11 C | 0.862 | 0.48052 |
| Malignant T-cell-amplified sequence 1 | Q9DB27 | Mcts1 | 21 kDa | 4 C | 0.862 | 0.51036 |
| Zyxin | Q62523 | Zyx | 61 kDa | 23 C | 0.862 | 0.58373 |
| 14-3-3 protein zeta/delta | P63101 | Ywhaz | 28 kDa | 3 C | 0.858 | 0.04574 |
| Insulin-degrading enzyme | Q9JHR7 | Ide | 118 kDa | 13 C | 0.857 | 0.21684 |
| Macrophage-capping protein | P24452 | Capg | 39 kDa | 5 C | 0.857 | 0.69323 |
| Selenium-binding protein 2 | Q63836 | Selenbp2 | 53 kDa | 10 C | 0.855 | 0.3264 |
| 182 kDa tankyrase-1-binding protein | P58871 | Tnks1bp1 | 182 kDa | 23 C | 0.854 | 0.68449 |
| Elongation factor 2 | P58252 | Eef2 | 95 kDa | 7 C | 0.853 | 0.28741 |
| Protein-glutamine gamma-glutamyltransferase 2 | P21981 | Tgm2 | 77 kDa | 20 C | 0.850 | 0.73997 |
| Epsin-1 | Q80VP1 (+1) | Epn1 | 60 kDa | 2 C | 0.849 | 0.66424 |
| 40S ribosomal protein S17 | P63276 | Rps17 | 16 kDa | 1 C | 0.848 | 0.83599 |
| Tubulin-folding cofactor B | Q9D1E6 | Tbcb | 27 kDa | 5 C | 0.842 | 0.35062 |
| Pyrethroid hydrolase Ces2e | Q8BK48 | Ces2e | 62 kDa | 7 C | 0.842 | 0.43477 |
| Adapter molecule crk | Q64010 | Crk | 34 kDa | 1 C | 0.842 | 0.49326 |
| Transcription elongation factor A protein 1 | P10711 | Tcea1 | 34 kDa | 8 C | 0.842 | 0.49938 |
| 39S ribosomal protein L49, mitochondrial | Q9CQ40 | Mrpl49 | 19 kDa | 1 C | 0.839 | 0.80039 |
| Peroxisomal coenzyme A diphosphatase NUDT7 | Q99P30 (+1) | Nudt7 | 27 kDa | 4 C | 0.836 | 0.30309 |
| Probable aminopeptidase NPEPL1 | Q6NSR8 | Npepl1 | 56 kDa | 16 C | 0.835 | 0.23076 |
| Osteoclast-stimulating factor 1 | Q62422 | Ostf1 | 24 kDa | 4 C | 0.835 | 0.42479 |
| NADP-dependent malic enzyme | P06801 | Me1 | 64 kDa | 11 C | 0.831 | 0.28928 |
| Septin-7 | O55131 | 43350 | 51 kDa | 6 C | 0.828 | 0.44285 |
| Acylcarnitine hydrolase | Q91WG0 | Ces2c | 62 kDa | 5 C | 0.826 | 0.81642 |
| Isochorismatase domain-containing protein 2A | P85094 | Isoc2a | 22 kDa | 6 C | 0.826 | 0.05184 |
| Nucleoporin Nup43 | P59235 | Nup43 | 42 kDa | 10 C | 0.825 | 0.81752 |
| ADP-sugar pyrophosphatase | Q9JKX6 | Nudt5 | 24 kDa | 5 C | 0.825 | 0.81752 |
| Ig kappa chain V-III region PC 7043 | P01665 (+1) | | 12 kDa | 2 C | 0.820 | 0.81691 |
| 5'-3' exoribonuclease 2 | Q9DBR1 (+1 | Xrn2 | 109 kDa | 17 C | 0.820 | 0.73528 |
| Protein NipSnap homolog 3B | Q9CQE1 | Nipsnap3b | 28 kDa | 2 C | 0.819 | 0.09212 |
| Small glutamine-rich tetratricopeptide repeat-containing protein alpha | Q8BJU0 | Sgta | 34 kDa | 4 C | 0.816 | 0.75448 |
| Methylosome protein 50 | Q99J09 | Wdr77 | 37 kDa | 12 C | 0.816 | 0.77922 |
| Small ubiquitin-related modifier 3 | Q9Z172 | Sumo3 | 12 kDa | 3 C | 0.816 | 0.75448 |
| Sulfatase-modifying factor 1 | Q8R0F3 | Sumf1 | 41 kDa | 11 C | 0.813 | 0.57714 |
| Isochorismatase domain-containing protein 1 | Q91V64 | Isoc1 | 32 kDa | 5 C | 0.813 | 0.57714 |
| Cleavage and polyadenylation specificity factor subunit 6 | Q6NVF9 | Cpsf6 | 59 kDa | 3 C | 0.810 | 0.51961 |
| Heat shock 70 kDa protein 4 | Q61316 | Hspa4 | 94 kDa | 14 C | 0.807 | 0.17666 |
| Leukotriene A-4 hydrolase | P24527 | Lta4h | 69 kDa | 11 C | 0.806 | 0.67103 |
| Protein AMBP | Q07456 | Ambp | 39 kDa | 16 C | 0.800 | 0.81985 |
| Inositol monophosphatase 1 | O55023 | Impa1 | 30 kDa | 6 C | 0.800 | 0.08317 |
| Actin-related protein 2 | P61161 | Actr2 | 45 kDa | 5 C | 0.800 | 0.17353 |
| Protein LSM12 homolog | Q9D0R8 | Lsm12 | 22 kDa | 5 C | 0.800 | 0.19301 |
| Kynurenine--oxoglutarate transaminase 1 | Q8BTY1 | Kyat1 | 48 kDa | 7 C | 0.800 | 0.2542 |

| | | | | | | |
|---|---|---|---|---|---|---|
| Ubiquitin recognition factor in ER-associated degradation protein 1 | P70362 | Ufd1 | 34 kDa | 5 C | 0.800 | 0.29235 |
| Eukaryotic translation initiation factor 1A, X-chromosomal | Q8BMJ3 | Eif1ax | 16 kDa | 2 C | 0.800 | 0.40708 |
| Actin-related protein 2/3 complex subunit 5 | Q9CPW4 | Arpc5 | 16 kDa | 1 C | 0.800 | 0.40708 |
| Glutaredoxin-related protein 5, mitochondrial | Q80Y14 | Glrx5 | 16 kDa | 2 C | 0.800 | 0.48033 |
| Ig heavy chain V-III region A4 | P01796 (+3) | | 13 kDa | 2 C | 0.800 | 0.48033 |
| Isoleucine--tRNA ligase, mitochondrial | Q8BIJ6 | Iars2 | 113 kDa | 20 C | 0.800 | 0.55146 |
| Calcium-regulated heat stable protein 1 | Q9CR86 | Carhsp1 | 16 kDa | 4 C | 0.800 | 0.60141 |
| PITH domain-containing protein 1 | Q8BWR2 | Pithd1 | 24 kDa | 4 C | 0.800 | 0.63871 |
| Ig kappa chain V19-17 | P01633 | Igk-V19-17 | 16 kDa | 2 C | 0.800 | 0.74723 |
| Proteasome subunit alpha type-6 | Q9QUM9 | Psma6 | 27 kDa | 8 C | 0.800 | 0.03035 |
| Mitochondrial fission 1 protein | Q9CQ92 | Fis1 | 17 kDa | 1 C | 0.800 | 0.19962 |
| Lysosome-associated membrane glycoprotein 1 | P11438 | Lamp1 | 44 kDa | 8 C | 0.800 | 0.40708 |
| Acyl-coenzyme A amino acid N-acyltransferase 1 | A2AKK5 | Acnat1 | 46 kDa | 7 C | 0.800 | 0.46084 |
| High mobility group protein B1 | P63158 | Hmgb1 | 25 kDa | 3 C | 0.800 | 0.52793 |
| Beta-glucuronidase | P12265 | Gusb | 74 kDa | 8 C | 0.800 | 0.53562 |
| Protein SEC13 homolog | Q9D1M0 | Sec13 | 36 kDa | 9 C | 0.800 | 0.74337 |
| Ran-binding protein 3 | Q9CT10 | Ranbp3 | 53 kDa | 5 C | 0.800 | 0.74337 |
| Galectin-3-binding protein | Q07797 | Lgals3bp | 64 kDa | 16 C | 0.793 | 0.7714 |
| Ornithine aminotransferase, mitochondrial | P29758 | Oat | 48 kDa | 7 C | 0.789 | 0.42319 |
| Hydroxyacid oxidase 1 | Q9WU19 | Hao1 | 41 kDa | 5 C | 0.783 | 0.11163 |
| Transgelin-2 | Q9WVA4 | Tagln2 | 22 kDa | 3 C | 0.781 | 0.33209 |
| Galectin-3 | P16110 | Lgals3 | 28 kDa | 1 C | 0.776 | 0.79463 |
| Isoform 2 of Heterogeneous nuclear ribonucleoprotein Q | Q7TMK9-2 | Syncrip | 63 kDa | 4 C | 0.771 | 0.29221 |
| Ig heavy chain V region AC38 205.12 | P06330 | | 13 kDa | 2 C | 0.769 | 0.43477 |
| Septin-10 | Q8C650 | 43353 | 52 kDa | 14 C | 0.753 | 0.29767 |
| Ig kappa chain V-III region PC 2880/PC 1229 | P01654 | | 12 kDa | 2 C | 0.753 | 0.49778 |
| Xanthine dehydrogenase/oxidase | Q00519 | Xdh | 147 kDa | 37 C | 0.753 | 0.20735 |
| Dynein light chain Tctex-type 3 | P56387 | Dynlt3 | 13 kDa | 6 C | 0.749 | 0.3438 |
| Leucine-rich repeat flightless-interacting protein 1 | Q3UZ39 | Lrrfip1 | 79 kDa | 12 C | 0.748 | 0.34529 |
| Alpha-1-antitrypsin 1-2 | P22599 | Serpina1b | 46 kDa | 3 C | 0.747 | 0.56178 |
| Splicing factor, proline- and glutamine-rich | Q8VIJ6 | Sfpq | 75 kDa | 7 C | 0.736 | 0.2581 |
| Glia maturation factor beta | Q9CQI3 | Gmfb | 17 kDa | 3 C | 0.735 | 0.35247 |
| Dihydropyrimidine dehydrogenase [NADP(+)] | Q8CHR6 | Dpyd | 111 kDa | 35 C | 0.734 | 0.06347 |
| START domain-containing protein 10 | Q9JMD3 | Stard10 | 33 kDa | 7 C | 0.729 | 0.06015 |
| Selenoprotein P | P70274 | Selenop | 43 kDa | 18 C | 0.728 | 0.28828 |
| Heterogeneous nuclear ribonucleoprotein U | Q8VEK3 (+1) | Hnrnpu | 88 kDa | 14 C | 0.728 | 0.454 |
| Hepatoma-derived growth factor | P51859 | Hdgf | 26 kDa | 2 C | 0.724 | 0.26828 |
| Complement component C8 alpha chain | Q8K182 | C8a | 66 kDa | 29 C | 0.720 | 0.49004 |
| Proteasome assembly chaperone 1 | Q9JK23 | Psmg1 | 33 kDa | 15 C | 0.720 | 0.60641 |
| Ribosome-binding protein 1 | Q99PL5 | Rrbp1 | 173 kDa | 8 C | 0.719 | 0.11054 |
| Cytosolic 10-formyltetrahydrofolate dehydrogenase | Q8R0Y6 | Aldh1l1 | 99 kDa | 15 C | 0.715 | 0.09759 |
| Anamorsin | Q8WTY4 | Ciapin1 | 33 kDa | 10 C | 0.712 | 0.46533 |
| Phosphopantothenoylcysteine decarboxylase | Q8BZB2 | Ppcdc | 22 kDa | 6 C | 0.710 | 0.33707 |
| AH receptor-interacting protein | O08915 | Aip | 38 kDa | 8 C | 0.709 | 0.4744 |
| Plastin-3 | Q99K51 | Pls3 | 71 kDa | 9 C | 0.708 | 0.13939 |
| DnaJ homolog subfamily C member 7 | Q9QYI3 | Dnajc7 | 56 kDa | 14 C | 0.699 | 0.57299 |
| RNA-binding protein FUS | P56959 | Fus | 53 kDa | 4 C | 0.691 | 0.31842 |
| Tyrosine--tRNA ligase, cytoplasmic | Q91WQ3 | Yars | 59 kDa | 7 C | 0.688 | 0.51697 |
| Phosphatidylcholine transfer protein | P53808 | Pctp | 25 kDa | 4 C | 0.680 | 0.2958 |
| Multifunctional protein ADE2 | Q9DCL9 | Paics | 47 kDa | 13 C | 0.676 | 0.13004 |
| Isoform 6 of Palladin | Q9ET54-6 | Palld | 151 kDa | 17 C | 0.667 | 0.21818 |
| WW domain-binding protein 2 | P97765 | Wbp2 | 28 kDa | 2 C | 0.655 | 0.08707 |
| Nascent polypeptide-associated complex subunit alpha, muscle-specific form | P70670 | Naca | 220 kDa | 16 C | 0.650 | 0.046 |
| Pyrroline-5-carboxylate reductase 3 | Q9DCC4 | Pycr3 | 29 kDa | 8 C | 0.648 | 0.66366 |
| Dual specificity protein phosphatase 3 | Q9D7X3 | Dusp3 | 20 kDa | 4 C | 0.648 | 0.07271 |
| Ig kappa chain V-V region L7 (Fragment) | P01642 | Gm10881 | 13 kDa | 2 C | 0.644 | 0.5262 |
| Xaa-Pro dipeptidase | Q11136 | Pepd | 55 kDa | 17 C | 0.637 | 0.06781 |
| NIF3-like protein 1 | Q9EQ80 | Nif3l1 | 42 kDa | 8 C | 0.634 | 0.23876 |
| S-methyl-5'-thioadenosine phosphorylase | Q9CQ65 | Mtap | 31 kDa | 10 C | 0.629 | 0.03179 |
| von Willebrand factor A domain-containing protein 5A | Q99KC8 | Vwa5a | 87 kDa | 13 C | 0.619 | 0.16109 |
| DNA-directed RNA polymerases I, II, and III subunit RPABC3 | Q923G2 | Polr2h | 17 kDa | 1 C | 0.610 | 0.19322 |
| Gephyrin | Q8BUV3 | Gphn | 83 kDa | 13 C | 0.610 | 0.2094 |
| Nucleolin | P09405 | Ncl | 77 kDa | 1 C | 0.610 | 0.54851 |
| Creatine kinase B-type | Q04447 | Ckb | 43 kDa | 5 C | 0.610 | 0.54851 |
| Calpastatin | P51125 (+1) | Cast | 85 kDa | 4 C | 0.600 | 0.00963 |
| Guanine deaminase | Q9R111 | Gda | 51 kDa | 9 C | 0.594 | 0.1095 |
| Aldehyde oxidase 1 | O54754 | Aox1 | 147 kDa | 41 C | 0.586 | 0.06277 |
| G protein-regulated inducer of neurite outgrowth 3 | Q8BWS5 | Gprin3 | 80 kDa | 16 C | 0.585 | 0.20381 |
| Glycogen phosphorylase, liver form | Q9ET01 | Pygl | 97 kDa | 8 C | 0.585 | 0.14761 |
| Tubulin polymerization-promoting protein | Q7TQD2 | Tppp | 24 kDa | 3 C | 0.583 | 0.11485 |
| Glutamine synthetase | P15105 | Glul | 42 kDa | 13 C | 0.581 | 0.03133 |
| 5-oxoprolinase | Q8K010 | Oplah | 138 kDa | 25 C | 0.578 | 0.25598 |
| Beta-arrestin-1 | Q8BWG8 | Arrb1 | 47 kDa | 8 C | 0.578 | 0.00811 |
| Stromal cell-derived factor 2 | Q9DCT5 | Sdf2 | 23 kDa | 4 C | 0.578 | 0.03456 |
| Isoform 2 of Lymphocyte-specific protein 1 | P19973-2 | Lsp1 | 37 kDa | 2 C | 0.575 | 0.49263 |
| Early endosome antigen 1 | Q8BL66 | Eea1 | 161 kDa | 20 C | 0.572 | 0.4769 |
| Target of Myb protein 1 | O88746 | Tom1 | 54 kDa | 4 C | 0.560 | 0.05547 |
| Protein argonaute-2 | Q8CJG0 | Ago2 | 97 kDa | 22 C | 0.560 | 0.34303 |

| | | | | | | |
|---|---|---|---|---|---|---|
| Prolow-density lipoprotein receptor-related protein 1 | Q91ZX7 | Lrp1 | 505 kDa | 332 C | 0.551 | 0.34752 |
| ADP-ribosylation factor-binding protein GGA1 | Q8R0H9 | Gga1 | 70 kDa | 6 C | 0.547 | 0.07506 |
| Caprin-1 | Q60865 | Caprin1 | 78 kDa | 3 C | 0.533 | 0.05753 |
| Eukaryotic translation initiation factor 3 subunit I | Q9QZD9 | Eif3i | 36 kDa | 7 C | 0.533 | 0.01765 |
| Nuclear protein localization protein 4 homolog | P60670 (+1) | Nploc4 | 68 kDa | 17 C | 0.528 | 0.25981 |
| BAG family molecular chaperone regulator 3 | Q9JLV1 | Bag3 | 62 kDa | 4 C | 0.523 | 0.00056 |
| Fatty acid-binding protein, epidermal | Q05816 | Fabp5 | 15 kDa | 6 C | 0.523 | 0.02655 |
| Ethylmalonyl-CoA decarboxylase | Q9D9V3 | Echdc1 | 35 kDa | 6 C | 0.520 | 0.17667 |
| RNA-binding protein EWS | Q61545 | Ewsr1 | 68 kDa | 5 C | 0.518 | 0.04895 |
| Alanine aminotransferase 2 | Q8BGT5 | Gpt2 | 58 kDa | 14 C | 0.510 | 0.08809 |
| ERBB receptor feedback inhibitor 1 | Q99JZ7 | Errfi1 | 50 kDa | 14 C | 0.510 | 0.08809 |
| Serpin B8 | O08800 | Serpinb8 | 42 kDa | 11 C | 0.510 | 0.33285 |
| GMP reductase 1 | Q9DCZ1 | Gmpr | 37 kDa | 9 C | 0.500 | 0.1514 |
| Actin-related protein 10 | Q9QZB7 | Actr10 | 46 kDa | 10 C | 0.468 | 0.35426 |
| Complement C4-B | P01029 | C4b | 193 kDa | 29 C | 0.462 | 0.14286 |
| Ras GTPase-activating protein-binding protein 2 | P97379 (+1) | G3bp2 | 54 kDa | 1 C | 0.449 | 0.22315 |
| Growth arrest-specific protein 2 | P11862 | Gas2 | 35 kDa | 11 C | 0.438 | 0.39455 |
| Ethanolamine-phosphate cytidylyltransferase | Q922E4 | Pcyt2 | 45 kDa | 8 C | 0.433 | 0.34907 |
| Shootin-1 | Q8K2Q9 | Shtn1 | 71 kDa | 9 C | 0.375 | 0.06431 |
| Intracellular hyaluronan-binding protein 4 | Q9JKS5 | Habp4 | 46 kDa | 3 C | 0.350 | 0.28281 |
| Inositol-3-phosphate synthase 1 | Q9JHU9 | Isyna1 | 61 kDa | 11 C | 0.339 | 0.4419 |
| Ig kappa chain V-V region HP R16.7 | P01644 (+1) | | 12 kDa | 2 C | 0.326 | 0.13993 |
| NudC domain-containing protein 2 | Q9CQ48 | Nudcd2 | 18 kDa | 3 C | 0.320 | 0.1413 |
| Alpha-mannosidase 2C1 | Q91W89 | Man2c1 | 116 kDa | 24 C | 0.308 | 0.11326 |
| Epidermal growth factor receptor | Q01279 | Egfr | 135 kDa | 60 C | 0.273 | 0.1029 |
| Filamin-B | Q80X90 | Flnb | 278 kDa | 43 C | 0.125 | 0.12176 |
| Protein SGT1 homolog | Q9CX34 | Sugt1 | 38 kDa | 5 C | 0.125 | 0.12176 |
| Cysteine desulfurase, mitochondrial | Q9Z1J3 | Nfs1 | 51 kDa | 7 C | 0.121 | 0.29235 |
| O-phosphoseryl-tRNA(Sec) selenium transferase | Q6P6M7 | Sepsecs | 55 kDa | 13 C | 0.497 | 0.00869 |
| Molybdenum cofactor sulfurase | Q14CH1 | Mocos | 95 kDa | 24 C | 0.457 | 0.01553 |
| Catenin alpha-1 | P26231 | Ctnna1 | 100 kDa | 12 C | 0.455 | 0.04496 |
| Serine/threonine-protein phosphatase CPPED1 | Q8BFS6 | Cpped1 | 35 kDa | 6 C | 0.453 | 0.01903 |
| Complement factor H | P06909 | Cfh | 139 kDa | 82 C | 0.432 | 0.0433 |
| Protein farnesyltransferase/geranylgeranyltransferase type-1 subunit alpha | Q61239 | Fnta | 44 kDa | 3 C | 0.416 | 0.03148 |
| Thioredoxin reductase 3 | Q99MD6 | Txnrd3 | 71 kDa | 18 C | 0.307 | 0.02231 |
| Alpha-1-antitrypsin 1-5 | Q00898 | Serpina1e | 46 kDa | 4 C | 0.297 | 0.01301 |
| Properdin | P11680 | Cfp | 50 kDa | 44 C | 0.263 | 0.00608 |
| Uncharacterized protein C2orf72 homolog | Q9CYS6 (+1) | | 30 kDa | 5 C | 0.224 | 0.01434 |
| Protein farnesyltransferase subunit beta | Q8K2I1 | Fntb | 49 kDa | 17 C | 0.224 | 0.01434 |
| Heat shock 70 kDa protein 4L | P48722 | Hspa4l | 94 kDa | 15 C | 0.210 | 0.0136 |
| 2-amino-3-ketobutyrate coenzyme A ligase, mitochondrial | O88986 | Gcat | 45 kDa | 10 C | 0.187 | 0.03985 |
| Microtubule-associated protein 4 | P27546 (+1) | Map4 | 117 kDa | 10 C | 0.184 | 0.03822 |
| Cytoplasmic protein NCK1 | Q99M51 | Nck1 | 43 kDa | 4 C | 0.140 | 0.0042 |
| 2-amino-3-carboxymuconate-6-semialdehyde decarboxylase | Q8R519 | Acmsd | 38 kDa | 7 C | 0.115 | 0.00864 |
