## Supplemental Table 2 for "Dietary restriction transforms the protein sulfhydrome in a tissue-specific and cystathionine γ-lyase-dependent manner"

| Supplemental Table 1: Dietary Impact on the Kidney Sulfhydrome | | | | | | |
|---|---|---|---|---|---|---|
| Protein Name | Accession Number | Alternate ID | Molecular Weight | Cysteine Residues | DR/AL Spectral Count Ratio | P-value |
| Xanthine dehydrogenase/oxidase | Q00519 | Xdh | 147 kDa | 37 C | 28.200 | 0.04859 |
| Protein-glucosylgalactosylhydroxylysine glucosidase | Q8BP56 | Pgghg | 76 kDa | 8 C | 12.200 | 0.02824 |
| Splicing factor 3B subunit 4 | Q8QZY9 | Sf3b4 | 44 kDa | 1 C | 10.200 | 0.0307 |
| 5-oxoprolinase | Q8K010 | Oplah | 138 kDa | 25 C | 7.608 | 0.016 |
| Phosphoserine phosphatase | Q99LS3 | Psph | 25 kDa | 4 C | 6.215 | 0.03147 |
| Carbonic anhydrase 5B | Q9QZA0 | Ca5b | 37 kDa | 7 C | 3.636 | 0.03504 |
| Serpin B6 | Q60854 | Serpinb6 | 43 kDa | 6 C | 3.478 | 0.03593 |
| Acylphosphatase-2 | P56375 | Acyp2 | 12 kDa | 1 C | 3.000 | 0.02695 |
| Ubiquitin-conjugating enzyme E2 L3 | P68037 | Ube2l3 | 18 kDa | 3 C | 2.640 | 0.04164 |
| Protein arginine N-methyltransferase 1 | Q9JIF0 | Prmt1 | 42 kDa | 11 C | 2.581 | 0.02017 |
| Prolow-density lipoprotein receptor-related protein 1 | Q91ZX7 | Lrp1 | 505 kDa | 332 C | 2.400 | 0.046 |
| F-actin-capping protein subunit alpha-2 | P47754 | Capza2 | 33 kDa | 3 C | 2.320 | 0.02218 |
| Carbonic anhydrase 3 | P16015 | Ca3 | 29 kDa | 5 C | 2.109 | 0.01857 |
| Caspase-6 | O08738 | Casp6 | 32 kDa | 10 C | 2.080 | 0.0066 |
| Isoform 2 of Eukaryotic peptide chain release factor GTP-binding subunit ERF3A | Q8R050-2 | Gspt1 | 69 kDa | 14 C | 2.065 | 0.04608 |
| GMP reductase 2 | Q99L27 | Gmpr2 | 38 kDa | 9 C | 2.057 | 0.02779 |
| Liprin-beta-1 | Q8C8U0 | Ppfibp1 | 109 kDa | 11 C | 30.000 | 0.09061 |
| Golgin subfamily A member 4 | Q91VW5 | Golga4 | 258 kDa | 25 C | 28.600 | 0.37 |
| Clathrin heavy chain 1 | Q68FD5 | Cltc | 192 kDa | 31 C | 22.800 | 0.40708 |
| Proteasome assembly chaperone 1 | Q9JK23 | Psmg1 | 33 kDa | 15 C | 20.400 | 0.15628 |
| Iron-sulfur cluster co-chaperone protein HscB | Q8K3A0 | Hscb | 27 kDa | 7 C | 18.400 | 0.08903 |
| Alanine--glyoxylate aminotransferase 2 | Q3UEG6 | Agxt2 | 57 kDa | 14 C | 18.400 | 0.15069 |
| Iron-sulfur cluster assembly 2 homolog | Q9DCB8 | Isca2 | 17 kDa | 4 C | 16.400 | 0.17932 |
| Ras GTPase-activating-like protein IQGAP1 | Q9JKF1 | Iqgap1 | 189 kDa | 14 C | 16.400 | 0.26009 |
| Matrin-3 | Q8K310 | Matr3 | 95 kDa | 9 C | 14.600 | 0.25098 |
| Myotubularin | Q9Z2C5 | Mtm1 | 70 kDa | 9 C | 12.400 | 0.11712 |
| Golgin subfamily A member 1 | Q9CW79 | Golga1 | 87 kDa | 8 C | 12.400 | 0.11712 |
| Glucosamine 6-phosphate N-acetyltransferase | Q9JK38 | Gnpnat1 | 21 kDa | 6 C | 12.400 | 0.11712 |
| Ribose-phosphate pyrophosphokinase 1 | Q9D7G0 | Prps1 | 35 kDa | 9 C | 12.400 | 0.20459 |
| Ankycorbin | Q9EP71 | Rai14 | 109 kDa | 12 C | 12.369 | 0.13366 |
| Cubilin | Q9JLB4 | Cubn | 399 kDa | 155 C | 12.200 | 0.07758 |
| N-myc-interactor | O35309 | Nmi | 35 kDa | 8 C | 10.600 | 0.20691 |
| Isoform 3 of Programmed cell death 6-interacting protein | Q9WU78-3 | Pdcd6ip | 97 kDa | 10 C | 10.400 | 0.16156 |
| Eukaryotic translation initiation factor 1 | P48024 | Eif1 | 13 kDa | 2 C | 10.400 | 0.16156 |
| GRIP and coiled-coil domain-containing protein 2 | Q8CHG3 | Gcc2 | 194 kDa | 17 C | 8.800 | 0.40708 |
| Prefoldin subunit 3 | P61759 | Vbp1 | 22 kDa | 3 C | 8.800 | 0.40708 |
| Coiled-coil domain-containing protein 22 | Q9JIG7 | Ccdc22 | 71 kDa | 8 C | 8.800 | 0.40708 |
| Glutamyl-tRNA(Gln) amidotransferase subunit A | Q9CZN8 | Qrsl1 | 57 kDa | 11 C | 8.677 | 0.14907 |
| Intersectin-2 | Q9Z0R6 | Itsn2 | 189 kDa | 20 C | 8.600 | 0.19301 |
| Actin-related protein 10 | Q9QZB7 | Actr10 | 46 kDa | 10 C | 8.600 | 0.19301 |
| BAG family molecular chaperone regulator 5 | Q8CI32 | Bag5 | 51 kDa | 10 C | 8.600 | 0.27239 |
| FYVE and coiled-coil domain-containing protein 1 | Q8VDC1 | Fyco1 | 162 kDa | 32 C | 8.600 | 0.27239 |
| DnaJ homolog subfamily C member 9 | Q91WN1 | Dnajc9 | 30 kDa | 3 C | 8.600 | 0.27239 |
| Alpha-aminoadipic semialdehyde synthase | Q99K67 | Aass | 103 kDa | 13 C | 8.600 | 0.27239 |
| Mitogen-activated protein kinase 1 | P63085 | Mapk1 | 41 kDa | 7 C | 8.400 | 0.10695 |
| General vesicular transport factor p115 | Q9Z1Z0 | Uso1 | 107 kDa | 19 C | 8.400 | 0.10695 |
| Tyrosine-protein phosphatase non-receptor type 11 | P35235 | Ptpn11 | 68 kDa | 10 C | 8.400 | 0.10695 |
| Methylmalonyl-CoA epimerase | Q9D1I5 | Mcee | 19 kDa | 2 C | 8.400 | 0.10695 |
| Aminomethyltransferase | Q8CFA2 | Amt | 44 kDa | 8 C | 8.000 | 0.05861 |
| Isoform 4 of Cingulin-like protein 1 | Q6AW69-4 | Cgnl1 | 148 kDa | 13 C | 7.252 | 0.0627 |
| Early endosome antigen 1 | Q8BL66 | Eea1 | 161 kDa | 20 C | 6.913 | 0.05444 |
| U5 small nuclear ribonucleoprotein 200 kDa helicase | Q6P4T2 | Snrnp200 | 245 kDa | 29 C | 6.800 | 0.40708 |
| Isoform 2 of Alpha-ketoglutarate-dependent dioxygenase FTO | Q8BGW1-2 | Fto | 55 kDa | 15 C | 6.800 | 0.40708 |
| Proteasome activator complex subunit 2 | P97372 | Psme2 | 27 kDa | 4 C | 6.800 | 0.40708 |
| Ig heavy chain V regions TEPC 15/S107/HPCM1/HPCM2/HPCM3 | P01787 | | 14 kDa | 2 C | 6.800 | 0.40708 |
| Plectin | Q9QXS1 | Plec | 534 kDa | 34 C | 6.600 | 0.23224 |

| | | | | | | |
|---|---|---|---|---|---|---|
| Isoform 2 of Cysteine--tRNA ligase | Q9ER72-2 | Cars | 86 kDa | 11 C | 6.600 | 0.23224 |
| Eukaryotic translation initiation factor 5 | P59325 | Eif5 | 49 kDa | 8 C | 6.600 | 0.23224 |
| COP9 signalosome complex subunit 8 | Q8VBV7 | Cops8 | 23 kDa | 1 C | 6.600 | 0.23224 |
| Phosphopantothenate--cysteine ligase | Q8VDG5 | Ppcs | 34 kDa | 1 C | 6.600 | 0.23224 |
| EGF-containing fibulin-like extracellular matrix protein 1 | Q8BPB5 | Efemp1 | 55 kDa | 40 C | 6.600 | 0.23224 |
| UPF0568 protein C14orf166 homolog | Q9CQE8 | | 28 kDa | 2 C | 6.582 | 0.1777 |
| Periplakin | Q9R269 | Ppl | 204 kDa | 11 C | 6.215 | 0.19904 |
| Exosome complex component RRP41 | Q921I9 | Exosc4 | 26 kDa | 5 C | 5.164 | 0.14599 |
| Aminopeptidase N | P97449 | Anpep | 110 kDa | 8 C | 5.000 | 0.06231 |
| Nucleoprotein TPR | F6ZDS4 | Tpr | 274 kDa | 7 C | 4.958 | 0.1339 |
| Myosin-9 | Q8VDD5 | Myh9 | 226 kDa | 21 C | 4.904 | 0.09433 |
| Protein argonaute-2 | Q8CJG0 | Ago2 | 97 kDa | 22 C | 4.904 | 0.209 |
| WD repeat-containing protein 92 | Q8BGF3 | Wdr92 | 40 kDa | 7 C | 4.800 | 0.40708 |
| Dimethylaniline monooxygenase [N-oxide-forming] 1 | P50285 | Fmo1 | 60 kDa | 10 C | 4.800 | 0.40708 |
| Heterogeneous nuclear ribonucleoprotein L-like | Q921F4 | Hnrnpll | 64 kDa | 15 C | 4.800 | 0.40708 |
| ER membrane protein complex subunit 8 | O70378 | Emc8 | 23 kDa | 8 C | 4.800 | 0.40708 |
| T-complex protein 1 subunit theta | P42932 | Cct8 | 60 kDa | 10 C | 4.800 | 0.40708 |
| Optineurin | Q8K3K8 | Optn | 67 kDa | 12 C | 4.800 | 0.40708 |
| Transmembrane and immunoglobulin domain-containing protein 1 | Q9D7L8 | Tmigd1 | 29 kDa | 8 C | 4.800 | 0.40708 |
| 39S ribosomal protein L46 | Q9EQI8 | Mrpl46 | 32 kDa | 3 C | 4.800 | 0.40708 |
| Isoform 2 of Myosin phosphatase Rho-interacting protein | P97434-2 | Mprip | 118 kDa | 15 C | 4.565 | 0.11638 |
| E3 SUMO-protein ligase RanBP2 | Q9ERU9 | Ranbp2 | 341 kDa | 67 C | 4.400 | 0.08505 |
| Syntaxin-7 | O70439 | Stx7 | 30 kDa | 3 C | 4.369 | 0.10905 |
| Ferritin heavy chain | P09528 | Fth1 | 21 kDa | 3 C | 4.308 | 0.06634 |
| ELKS/Rab6-interacting/CAST family member 1 | Q99MI1 | Erc1 | 128 kDa | 4 C | 3.896 | 0.21001 |
| Pre-mRNA-splicing factor ATP-dependent RNA helicase DHX15 | O35286 | Dhx15 | 91 kDa | 14 C | 3.754 | 0.16469 |
| Low molecular weight phosphotyrosine protein phosphatase | Q9D358 | Acp1 | 18 kDa | 8 C | 3.754 | 0.08807 |
| Cysteine and glycine-rich protein 2 | P97314 | Csrp2 | 21 kDa | 16 C | 3.673 | 0.11552 |
| Alpha-soluble NSF attachment protein | Q9DB05 | Napa | 33 kDa | 8 C | 3.639 | 0.05924 |
| Isoform HK1 of Hexokinase-1 | P17710-3 | Hk1 | 102 kDa | 21 C | 3.624 | 0.13989 |
| Guanine deaminase | Q9R111 | Gda | 51 kDa | 9 C | 3.500 | 0.05103 |
| Isoform 2 of Pleckstrin homology domain-containing family A member 7 | Q3UIL6-2 | Plekha7 | 144 kDa | 10 C | 3.376 | 0.12329 |
| G-rich sequence factor 1 | Q8C5Q4 | Grsf1 | 53 kDa | 9 C | 3.355 | 0.11315 |
| Cingulin | P59242 | Cgn | 136 kDa | 4 C | 3.345 | 0.25578 |
| Isoform 4 of TRIO and F-actin-binding protein | Q99KW3-4 | Triobp | 218 kDa | 34 C | 3.309 | 0.10533 |
| Isoform 2 of A-kinase anchor protein 9 | Q70FJ1-2 | Akap9 | 434 kDa | 55 C | 3.262 | 0.34006 |
| Complement component 1 Q subcomponent-binding protein | O35658 | C1qbp | 31 kDa | 6 C | 3.262 | 0.34006 |
| Basement membrane-specific heparan sulfate proteoglycan core protein | Q05793 | Hspg2 | 398 kDa | 188 C | 3.262 | 0.34006 |
| E3 ubiquitin-protein ligase NEDD4 | P46935 | Nedd4 | 103 kDa | 9 C | 3.262 | 0.43068 |
| Pyrroline-5-carboxylate reductase 3 | Q9DCC4 | Pycr3 | 29 kDa | 8 C | 3.262 | 0.43068 |
| Uveal autoantigen with coiled-coil domains and ankyrin repeats | Q8CGB3 | Uaca | 161 kDa | 22 C | 3.216 | 0.06919 |
| Sepiapterin reductase | Q64105 | Spr | 28 kDa | 10 C | 3.200 | 0.21217 |
| ATP synthase F(0) complex subunit B1 | Q9CQQ7 | Atp5f1 | 29 kDa | 2 C | 3.200 | 0.21217 |
| Isoform 3 of DEP domain-containing mTOR-interacting protein | Q570Y9-3 | Deptor | 45 kDa | 9 C | 3.200 | 0.21217 |
| Histone-binding protein RBBP7 | Q60973 | Rbbp7 | 48 kDa | 6 C | 3.200 | 0.2954 |
| Isoform 2 of RNA-binding protein 39 | Q8VH51-2 | Rbm39 | 59 kDa | 5 C | 3.200 | 0.21217 |
| Isoform 3 of Septin-9 | Q80UG5-3 | 43352 | 65 kDa | 6 C | 3.200 | 0.21217 |
| Protein angel homolog 2 | Q8K1C0 | Angel2 | 62 kDa | 18 C | 3.200 | 0.21217 |
| Serine--tRNA ligase | Q9JJL8 | Sars2 | 58 kDa | 9 C | 3.165 | 0.18498 |
| Small nuclear ribonucleoprotein-associated protein N | P63163 | Snrpn | 25 kDa | 3 C | 3.138 | 0.12115 |
| Complement factor H | P06909 | Cfh | 139 kDa | 82 C | 3.111 | 0.09853 |
| Protein phosphatase 1B | P36993 | Ppm1b | 43 kDa | 12 C | 3.022 | 0.21297 |
| InaD-like protein | Q63ZW7 | Patj | 199 kDa | 17 C | 3.000 | 0.08187 |
| Alpha-2-HS-glycoprotein | P29699 | Ahsg | 37 kDa | 14 C | 2.945 | 0.21747 |
| Isoform 2 of Nuclear protein localization protein 4 homolog | P60670-2 | Nploc4 | 64 kDa | 17 C | 2.865 | 0.06782 |
| CDGSH iron-sulfur domain-containing protein 1 | Q91WS0 | Cisd1 | 12 kDa | 3 C | 2.852 | 0.25123 |
| CAP-Gly domain-containing linker protein 2 | Q9Z0H8 | Clip2 | 116 kDa | 9 C | 2.852 | 0.25123 |
| Protein phosphatase 1 regulatory subunit 12A | Q9DBR7 | Ppp1r12a | 115 kDa | 8 C | 2.813 | 0.11253 |
| Chromobox protein homolog 1 | P83917 | Cbx1 | 21 kDa | 2 C | 2.791 | 0.07682 |
| Pyruvate carboxylase | Q05920 | Pc | 130 kDa | 13 C | 2.725 | 0.27526 |

| | | | | | | |
|---|---|---|---|---|---|---|
| UDP-glucuronosyltransferase 1-7C | Q6ZQM8 | Ugt1a7c | 60 kDa | 13 C | 2.708 | 0.55962 |
| Nuclear mitotic apparatus protein 1 | E9Q7G0 | Numa1 | 236 kDa | 19 C | 2.648 | 0.13207 |
| OTU domain-containing protein 6B | Q8K2H2 | Otud6b | 34 kDa | 4 C | 2.646 | 0.37403 |
| Alpha-mannosidase 2C1 | Q91W89 | Man2c1 | 116 kDa | 6 C | 2.627 | 0.32165 |
| 39S ribosomal protein L19 | Q9D338 | Mrpl19 | 34 kDa | 6 C | 2.618 | 0.29134 |
| Urocanate hydratase | Q8VC12 | Uroc1 | 75 kDa | 12 C | 2.618 | 0.3343 |
| Insulin-like growth factor-binding protein 7 | Q61581 | Igfbp7 | 29 kDa | 18 C | 2.606 | 0.234 |
| Mitochondrial-processing peptidase subunit alpha | Q9DC61 | Pmpca | 58 kDa | 9 C | 2.600 | 0.07603 |
| Striatin-3 | Q9ERG2 | Strn3 | 87 kDa | 8 C | 2.585 | 0.28648 |
| Profilin-2 | Q9JJV2 | Pfn2 | 15 kDa | 6 C | 2.585 | 0.28648 |
| Translin | Q62348 | Tsn | 26 kDa | 2 C | 2.585 | 0.28648 |
| Isoform 3 of Cytidine and dCMP deaminase domain-containing protein 1 | Q8BMD5-3 | Cdadc1 | 56 kDa | 17 C | 2.585 | 0.28648 |
| Cleavage and polyadenylation specificity factor subunit 6 | Q6NVF9 | Cpsf6 | 59 kDa | 3 C | 2.585 | 0.28648 |
| CAP-Gly domain-containing linker protein 1 | Q922J3 | Clip1 | 156 kDa | 11 C | 2.585 | 0.28648 |
| Vitronectin | P29788 | Vtn | 55 kDa | 14 C | 2.585 | 0.28648 |
| Pyridoxine-5'-phosphate oxidase | Q91XF0 | Pnpo | 30 kDa | 6 C | 2.585 | 0.28648 |
| Stromal cell-derived factor 2-like protein 1 | Q9ESP1 | Sdf2l1 | 24 kDa | 4 C | 2.582 | 0.30789 |
| Murinoglobulin-1 | P28665 | Mug1 | 165 kDa | 25 C | 2.581 | 0.05912 |
| Vigilin | Q8VDJ3 | Hdlbp | 142 kDa | 10 C | 2.549 | 0.11855 |
| Ran-specific GTPase-activating protein | P34022 | Ranbp1 | 24 kDa | 3 C | 2.537 | 0.10333 |
| Sideroflexin-1 | Q99JR1 | Sfxn1 | 36 kDa | 5 C | 2.504 | 0.38169 |
| Septin-6 | Q9R1T4 | 43349 | 50 kDa | 8 C | 2.495 | 0.16894 |
| ADP/ATP translocase 1 | P48962 | Slc25a4 | 33 kDa | 4 C | 2.477 | 0.29331 |
| Molybdenum cofactor biosynthesis protein 1 | Q5RKZ7 | Mocs1 | 70 kDa | 15 C | 2.470 | 0.20313 |
| Isoform A2 of Drebrin | Q9QXS6-2 | Dbn1 | 42 kDa | 13 C | 2.470 | 0.26696 |
| Agmatinase | A2AS89 | Agmat | 38 kDa | 9 C | 2.470 | 0.26696 |
| Galactose-1-phosphate uridylyltransferase | Q03249 | Galt | 43 kDa | 7 C | 2.470 | 0.3611 |
| Transcobalamin-2 | O88968 | Tcn2 | 48 kDa | 8 C | 2.448 | 0.34757 |
| Spectrin alpha chain | P16546 | Sptan1 | 285 kDa | 14 C | 2.422 | 0.06624 |
| Phosphopantothenoylcysteine decarboxylase | Q8BZB2 | Ppcdc | 22 kDa | 6 C | 2.361 | 0.24201 |
| Armadillo repeat-containing protein 1 | Q9D7A8 | Armc1 | 31 kDa | 4 C | 2.348 | 0.16468 |
| Polyadenylate-binding protein-interacting protein 1 | Q8VE62 | Paip1 | 46 kDa | 6 C | 2.323 | 0.06468 |
| Acidic leucine-rich nuclear phosphoprotein 32 family member A | O35381 | Anp32a | 29 kDa | 3 C | 2.323 | 0.18103 |
| Indolethylamine N-methyltransferase | P40936 | Inmt | 29 kDa | 11 C | 2.320 | 0.08994 |
| Thioredoxin reductase 3 | Q99MD6 | Txnrd3 | 71 kDa | 18 C | 2.311 | 0.12838 |
| Aldo-keto reductase family 1 member C21 | Q91WR5 | Akr1c21 | 37 kDa | 9 C | 2.305 | 0.39983 |
| Isoform 2 of Septin-8 | Q8CHH9-2 | 43351 | 50 kDa | 6 C | 2.300 | 0.08999 |
| Annexin A6 | P14824 | Anxa6 | 76 kDa | 8 C | 2.280 | 0.24452 |
| Ceruloplasmin | Q61147 | Cp | 121 kDa | 14 C | 2.275 | 0.36049 |
| Peroxidasin homolog | Q3UQ28 | Pxdn | 165 kDa | 46 C | 2.255 | 0.11596 |
| Ubiquinone biosynthesis protein COQ9 | Q8K1Z0 | Coq9 | 35 kDa | 3 C | 2.255 | 0.34243 |
| Lupus La protein homolog | P32067 | Ssb | 48 kDa | 3 C | 2.255 | 0.34243 |
| Ig gamma-2A chain C region secreted form | P01864 | | 37 kDa | 9 C | 2.250 | 0.06521 |
| Protein farnesyltransferase subunit beta | Q8K2I1 | Fntb | 49 kDa | 17 C | 2.250 | 0.12918 |
| Glycine N-acyltransferase-like protein Keg1 | Q9DCY0 | Keg1 | 34 kDa | 8 C | 2.240 | 0.41392 |
| Zinc phosphodiesterase ELAC protein 2 | Q80Y81 | Elac2 | 93 kDa | 23 C | 2.218 | 0.19366 |
| Complement factor B | P04186 | Cfb | 85 kDa | 20 C | 2.200 | 0.06758 |
| Epidermal growth factor receptor substrate 15-like 1 | Q60902 | Eps15l1 | 99 kDa | 3 C | 2.133 | 0.43093 |
| Sulfotransferase 1 family member D1 | Q3UZZ6 | Sult1d1 | 35 kDa | 3 C | 2.122 | 0.30465 |
| Ubiquitin-like modifier-activating enzyme 1 | Q02053 | Uba1 | 118 kDa | 21 C | 2.092 | 0.62111 |
| Mitochondrial 2-oxoglutarate/malate carrier protein | Q9CR62 | Slc25a11 | 34 kDa | 3 C | 2.092 | 0.62111 |
| Small nuclear ribonucleoprotein Sm D2 | P62317 | Snrpd2 | 14 kDa | 2 C | 2.092 | 0.62111 |
| NIF3-like protein 1 | Q9EQ80 | Nif3l1 | 42 kDa | 8 C | 2.090 | 0.19811 |
| Alanine--tRNA ligase | Q8BGQ7 | Aars | 107 kDa | 15 C | 2.090 | 0.3153 |
| Glutamyl aminopeptidase | P16406 | Enpep | 108 kDa | 9 C | 2.080 | 0.22422 |
| N(G) | Q99LD8 | Ddah2 | 30 kDa | 6 C | 2.065 | 0.13934 |
| StAR-related lipid transfer protein 5 | Q9EPQ7 | Stard5 | 24 kDa | 8 C | 2.057 | 0.07271 |
| Platelet-activating factor acetylhydrolase IB subunit beta | Q61206 | Pafah1b2 | 26 kDa | 3 C | 2.055 | 0.21575 |
| C-1-tetrahydrofolate synthase | Q922D8 | Mthfd1 | 101 kDa | 12 C | 2.031 | 0.50045 |
| Isoform 4 of Ankyrin-2 | Q8C8R3-4 | Ank2 | 433 kDa | 19 C | 2.031 | 0.50045 |

| | | | | | | |
|---|---|---|---|---|---|---|
| Interferon-induced 35 kDa protein homolog | Q9D8C4 | Ifi35 | 32 kDa | 4 C | 2.031 | 0.50045 |
| Protein FAM151A | Q8QZW3 | Fam151a | 67 kDa | 6 C | 2.031 | 0.50045 |
| ATP-dependent RNA helicase DDX3X | Q62167 | Ddx3x | 73 kDa | 7 C | 2.031 | 0.50045 |
| Thiopurine S-methyltransferase | O55060 | Tpmt | 28 kDa | 5 C | 2.031 | 0.50045 |
| Coatomer subunit beta' | O55029 | Copb2 | 102 kDa | 15 C | 2.031 | 0.50045 |
| Dynactin subunit 2 | Q99KJ8 | Dctn2 | 44 kDa | 2 C | 2.031 | 0.50045 |
| BH3-interacting domain death agonist | P70444 | Bid | 22 kDa | 2 C | 2.031 | 0.50045 |
| Inositol oxygenase | Q9QXN5 | Miox | 33 kDa | 6 C | 2.027 | 0.22137 |
| Trifunctional enzyme subunit alpha | Q8BMS1 | Hadha | 83 kDa | 12 C | 2.018 | 0.28203 |
| Isoform 2 of Septin-11 | Q8C1B7-2 | 43354 | 49 kDa | 6 C | 2.000 | 0.05815 |
| Nucleolysin TIAR | P70318 | Tial1 | 43 kDa | 6 C | 2.000 | 0.10533 |
| Malonyl-CoA-acyl carrier protein transacylase | Q8R3F5 | Mcat | 42 kDa | 12 C | 2.000 | 0.1365 |
| Peptidyl-prolyl cis-trans isomerase D | Q9CR16 | Ppid | 41 kDa | 7 C | 2.000 | 0.16579 |
| 28S ribosomal protein S28 | Q9CY16 | Mrps28 | 21 kDa | 3 C | 2.000 | 0.22639 |
| Neutral and basic amino acid transport protein rBAT | Q91WV7 | Slc3a1 | 78 kDa | 8 C | 2.000 | 0.34382 |
| Golgi reassembly-stacking protein 2 | Q99JX3 | Gorasp2 | 47 kDa | 4 C | 2.000 | 0.02695 |
| Ubiquitin-40S ribosomal protein S27a | P62983 | Rps27a | 18 kDa | 6 C | 2.000 | 0.03833 |
| Glutathione S-transferase Mu 5 | P48774 | Gstm5 | 27 kDa | 7 C | 1.990 | 0.33599 |
| Desmin | P31001 | Des | 53 kDa | 1 C | 1.988 | 0.58109 |
| Dipeptidase 1 | P31428 | Dpep1 | 46 kDa | 8 C | 1.980 | 0.20848 |
| Isoform 3 of Rab11 family-interacting protein 3 | Q8CHD8-3 | Rab11fip3 | 124 kDa | 44 C | 1.971 | 0.24169 |
| Isoform 1 of Glycerol kinase | Q64516-1 | Gk | 57 kDa | 15 C | 1.964 | 0.41663 |
| Prolyl endopeptidase | Q9QUR6 | Prep | 81 kDa | 17 C | 1.951 | 0.21116 |
| Presequence protease | Q8K411 | Pitrm1 | 117 kDa | 20 C | 1.947 | 0.02246 |
| Proteasome activator complex subunit 1 | P97371 | Psme1 | 29 kDa | 3 C | 1.943 | 0.14724 |
| GTP:AMP phosphotransferase AK3 | Q9WTP7 | Ak3 | 25 kDa | 1 C | 1.927 | 0.50564 |
| Very long-chain acyl-CoA synthetase | O35488 | Slc27a2 | 70 kDa | 14 C | 1.926 | 0.3427 |
| Ig kappa chain V-V region MOPC 41 | P01639 | Gm5571 | 14 kDa | 3 C | 1.920 | 0.05646 |
| Haloacid dehalogenase-like hydrolase domain-containing protein 3 | Q9CYW4 | Hdhd3 | 28 kDa | 4 C | 1.920 | 0.23088 |
| O-phosphoseryl-tRNA(Sec) selenium transferase | Q6P6M7 | Sepsecs | 55 kDa | 13 C | 1.915 | 0.38387 |
| Cytochrome c oxidase subunit 2 | P00405 | Mtco2 | 26 kDa | 3 C | 1.905 | 0.15768 |
| D-beta-hydroxybutyrate dehydrogenase | Q80XN0 | Bdh1 | 38 kDa | 6 C | 1.898 | 0.38002 |
| Talin-1 | P26039 | Tln1 | 270 kDa | 38 C | 1.892 | 0.15549 |
| Lamin-B1 | P14733 | Lmnb1 | 67 kDa | 4 C | 1.891 | 0.47061 |
| Succinate--CoA ligase [GDP-forming] subunit beta | Q9Z2I8 | Suclg2 | 47 kDa | 5 C | 1.877 | 0.19941 |
| Enoyl-CoA hydratase domain-containing protein 3 | Q9D7J9 | Echdc3 | 32 kDa | 5 C | 1.867 | 0.10901 |
| Prostaglandin reductase 2 | Q8VDQ1 | Ptgr2 | 38 kDa | 8 C | 1.867 | 0.10901 |
| FAS-associated death domain protein | Q61160 | Fadd | 23 kDa | 3 C | 1.867 | 0.10901 |
| Dynein light chain 1 | P63168 | Dynll1 | 10 kDa | 3 C | 1.867 | 0.30847 |
| 4-hydroxyphenylpyruvate dioxygenase | P49429 | Hpd | 45 kDa | 4 C | 1.867 | 0.0339 |
| N-acetyl-D-glucosamine kinase | Q9QZ08 | Nagk | 37 kDa | 6 C | 1.855 | 0.28959 |
| Isoform 2 of Small glutamine-rich tetratricopeptide repeat-containing protein alpha | Q8BJU0-2 | Sgta | 34 kDa | 4 C | 1.843 | 0.56658 |
| Vimentin | P20152 | Vim | 54 kDa | 1 C | 1.840 | 0.29767 |
| Methionine-R-sulfoxide reductase B2 | Q78J03 | Msrb2 | 19 kDa | 9 C | 1.829 | 0.04559 |
| Regulator of nonsense transcripts 1 | Q9EPU0 | Upf1 | 124 kDa | 23 C | 1.820 | 0.30258 |
| Proteasome subunit beta type-4 | P99026 | Psmb4 | 29 kDa | 2 C | 1.818 | 0.11765 |
| Leukotriene A-4 hydrolase | P24527 | Lta4h | 69 kDa | 11 C | 1.818 | 0.27729 |
| Ankyrin repeat and SAM domain-containing protein 4B | Q8K3X6 | Anks4b | 48 kDa | 6 C | 1.809 | 0.48935 |
| Protein arginine N-methyltransferase 5 | Q8CIG8 | Prmt5 | 73 kDa | 12 C | 1.807 | 0.26522 |
| Isoform 2 of Acyl-protein thioesterase 1 | P97823-2 | Lypla1 | 23 kDa | 6 C | 1.806 | 0.10938 |
| Fatty acid-binding protein | P51880 | Fabp7 | 15 kDa | 5 C | 1.806 | 0.10938 |
| Pregnancy zone protein | Q61838 | Pzp | 166 kDa | 24 C | 1.800 | 0.06129 |
| Protein-glutamine gamma-glutamyltransferase 2 | P21981 | Tgm2 | 77 kDa | 20 C | 1.800 | 0.17433 |
| Ig gamma-3 chain C region | P03987 | | 44 kDa | 10 C | 1.800 | 0.19301 |
| Catenin beta-1 | Q02248 | Ctnnb1 | 85 kDa | 11 C | 1.800 | 0.2958 |
| Isoform 2 of Cadherin-related family member 5 | Q8VHF2-2 | Cdhr5 | 73 kDa | 7 C | 1.800 | 0.47363 |
| Gephyrin | Q8BUV3 | Gphn | 83 kDa | 13 C | 1.787 | 0.16411 |
| Proteasome subunit beta type-1 | O09061 | Psmb1 | 26 kDa | 5 C | 1.785 | 0.20402 |
| Ig kappa chain V-II region 26-10 | P01631 | | 12 kDa | 2 C | 1.778 | 0.07331 |
| Serpin B8 | O08800 | Serpinb8 | 42 kDa | 11 C | 1.778 | 0.19113 |

| | | | | | | |
|---|---|---|---|---|---|---|
| Gamma-soluble NSF attachment protein | Q9CWZ7 | Napg | 35 kDa | 6 C | 1.776 | 0.31128 |
| Lipoma-preferred partner homolog | Q8BFW7 | Lpp | 66 kDa | 27 C | 1.776 | 0.36014 |
| Eukaryotic initiation factor 4A-II | P10630 | Eif4a2 | 46 kDa | 4 C | 1.774 | 0.43608 |
| Fatty acid-binding protein | Q05816 | Fabp5 | 15 kDa | 6 C | 1.760 | 0.01967 |
| Heterogeneous nuclear ribonucleoprotein D-like | Q9Z130 | Hnrnpdl | 34 kDa | 3 C | 1.760 | 0.01967 |
| Isoform 2 of Heterogeneous nuclear ribonucleoprotein M | Q9D0E1-2 | Hnrnpm | 74 kDa | 6 C | 1.760 | 0.01967 |
| Dynein light chain Tctex-type 3 | P56387 | Dynlt3 | 13 kDa | 6 C | 1.757 | 0.40486 |
| Calponin-3 | Q9DAW9 | Cnn3 | 36 kDa | 3 C | 1.752 | 0.45748 |
| Isoform 2 of Palladin | Q9ET54-2 | Palld | 122 kDa | 17 C | 1.745 | 0.09492 |
| Methionine aminopeptidase 1 | Q8BP48 | Metap1 | 43 kDa | 17 C | 1.741 | 0.33348 |
| Lysozyme C-2 | P08905 | Lyz2 | 17 kDa | 8 C | 1.733 | 0.07205 |
| COP9 signalosome complex subunit 3 | O88543 | Cops3 | 48 kDa | 10 C | 1.733 | 0.17353 |
| Glutamine synthetase | P15105 | Glul | 42 kDa | 13 C | 1.731 | 0.05615 |
| Actin-like protein 6A | Q9Z2N8 | Actl6a | 47 kDa | 9 C | 1.726 | 0.194 |
| Complement C1q tumor necrosis factor-related protein 3 | Q9ES30 | C1qtnf3 | 27 kDa | 5 C | 1.725 | 0.21916 |
| Isoleucine--tRNA ligase | Q8BIJ6 | Iars2 | 113 kDa | 20 C | 1.714 | 0.09461 |
| Actin-related protein 2 | P61161 | Actr2 | 45 kDa | 5 C | 1.714 | 0.10469 |
| Sorting nexin-3 | O70492 | Snx3 | 19 kDa | 1 C | 1.714 | 0.17638 |
| Tropomyosin alpha-4 chain | Q6IRU2 | Tpm4 | 28 kDa | 2 C | 1.714 | 0.01336 |
| 14 kDa phosphohistidine phosphatase | Q9DAK9 | Phpt1 | 14 kDa | 3 C | 1.714 | 0.02072 |
| Four and a half LIM domains protein 2 | O70433 | Fhl2 | 32 kDa | 35 C | 1.710 | 0.31289 |
| ADP-ribosylation factor-like protein 3 | Q9WUL7 | Arl3 | 20 kDa | 3 C | 1.705 | 0.21995 |
| Alcohol dehydrogenase 1 | P00329 | Adh1 | 40 kDa | 15 C | 1.705 | 0.25889 |
| Thioredoxin domain-containing protein 17 | Q9CQM5 | Txndc17 | 14 kDa | 6 C | 1.700 | 0.046 |
| Vinculin | Q64727 | Vcl | 117 kDa | 10 C | 1.696 | 0.00845 |
| Chromobox protein homolog 3 | P23198 | Cbx3 | 21 kDa | 3 C | 1.690 | 0.32033 |
| Glyoxalase domain-containing protein 5 | Q9D8I3 | Glod5 | 17 kDa | 5 C | 1.690 | 0.39481 |
| Exosome complex exonuclease RRP42 | Q9D0M0 | Exosc7 | 32 kDa | 12 C | 1.689 | 0.03569 |
| WD repeat domain phosphoinositide-interacting protein 3 | Q9CR39 | Wdr45b | 38 kDa | 15 C | 1.688 | 0.27599 |
| Ribonuclease inhibitor | Q91VI7 | Rnh1 | 50 kDa | 30 C | 1.686 | 0.12598 |
| 3-hydroxyisobutyrate dehydrogenase | Q99L13 | Hibadh | 35 kDa | 12 C | 1.680 | 0.03879 |
| Copper homeostasis protein cutC homolog | Q9D8X1 | Cutc | 29 kDa | 7 C | 1.677 | 0.22338 |
| Acetyl-CoA acetyltransferase | Q8CAY6 | Acat2 | 41 kDa | 8 C | 1.676 | 0.00934 |
| Gelsolin | P13020 | Gsn | 86 kDa | 7 C | 1.675 | 0.01042 |
| Aldose 1-epimerase | Q8K157 | Galm | 38 kDa | 4 C | 1.673 | 0.00122 |
| Glucosamine-6-phosphate isomerase 1 | O88958 | Gnpda1 | 33 kDa | 3 C | 1.667 | 0.01465 |
| Ester hydrolase C11orf54 homolog | Q91V76 | | 35 kDa | 8 C | 1.664 | 0.06205 |
| Filamin-A | Q8BTM8 | Flna | 281 kDa | 38 C | 1.655 | 0.15095 |
| Low-density lipoprotein receptor-related protein 2 | A2ARV4 | Lrp2 | 519 kDa | 329 C | 1.648 | 0.08607 |
| Heterogeneous nuclear ribonucleoproteins A2/B1 | O88569 | Hnrnpa2b1 | 37 kDa | 1 C | 1.626 | 0.0213 |
| Mitochondrial antiviral-signaling protein | Q8VCF0 | Mavs | 53 kDa | 8 C | 1.609 | 0.36587 |
| Pyruvate dehydrogenase protein X component | Q8BKZ9 | Pdhx | 54 kDa | 5 C | 1.600 | 0.15464 |
| Ig lambda-1 chain C region | P01843 | | 12 kDa | 3 C | 1.600 | 0.1674 |
| Arf-GAP with GTPase | Q8BXK8 | Agap1 | 94 kDa | 17 C | 1.600 | 0.20741 |
| Ubiquitin-conjugating enzyme E2 N | P61089 | Ube2n | 17 kDa | 1 C | 1.600 | 0.22745 |
| Splicing factor U2AF 65 kDa subunit | P26369 | U2af2 | 54 kDa | 6 C | 1.600 | 0.22745 |
| Serine/threonine-protein phosphatase CPPED1 | Q8BFS6 | Cpped1 | 35 kDa | 6 C | 1.600 | 0.22745 |
| Glycerol-3-phosphate dehydrogenase [NAD(+)] | P13707 | Gpd1 | 38 kDa | 11 C | 1.600 | 0.25834 |
| Isoform 2 of Peptidyl-prolyl cis-trans isomerase H | Q9D868-2 | Ppih | 19 kDa | 5 C | 1.600 | 0.43477 |
| Methylosome protein 50 | Q99J09 | Wdr77 | 37 kDa | 12 C | 1.600 | 0.25507 |
| Talin-2 | Q71LX4 | Tln2 | 254 kDa | 40 C | 1.600 | 0.29114 |
| Copine-1 | Q8C166 | Cpne1 | 59 kDa | 13 C | 1.600 | 0.04794 |
| Isoform 5 of Peripheral plasma membrane protein CASK | O70589-5 | Cask | 104 kDa | 16 C | 1.600 | 0.50876 |
| Isoform 2 of Clathrin interactor 1 | Q99KN9-2 | Clint1 | 51 kDa | 1 C | 1.600 | 0.5965 |
| Dihydropyrimidine dehydrogenase [NADP(+)] | Q8CHR6 | Dpyd | 111 kDa | 35 C | 1.584 | 0.24715 |
| L-lactate dehydrogenase B chain | P16125 | Ldhb | 37 kDa | 5 C | 1.584 | 0.1697 |
| Cytochrome c oxidase subunit 4 isoform 1 | P19783 | Cox4i1 | 20 kDa | 1 C | 1.580 | 0.3946 |
| Cleavage and polyadenylation specificity factor subunit 5 | Q9CQF3 | Nudt21 | 26 kDa | 1 C | 1.580 | 0.44242 |
| Hepatocyte growth factor-regulated tyrosine kinase substrate | Q99LI8 | Hgs | 86 kDa | 11 C | 1.575 | 0.70094 |
| Four and a half LIM domains protein 3 | Q9R059 | Fhl3 | 32 kDa | 35 C | 1.574 | 0.35748 |

| | | | | | | |
|---|---|---|---|---|---|---|
| KN motif and ankyrin repeat domain-containing protein 4 | Q6P9J5 | Kank4 | 110 kDa | 13 C | 1.574 | 0.35748 |
| Propionyl-CoA carboxylase alpha chain | Q91ZA3 | Pcca | 80 kDa | 11 C | 1.569 | 0.47729 |
| Inosine-5'-monophosphate dehydrogenase 2 | P24547 | Impdh2 | 56 kDa | 7 C | 1.565 | 0.21581 |
| Isoamyl acetate-hydrolyzing esterase 1 homolog | Q9DB29 | Iah1 | 28 kDa | 8 C | 1.564 | 0.13601 |
| Toll-like receptor 3 | Q99MB1 | Tlr3 | 104 kDa | 15 C | 1.564 | 0.66117 |
| 14-3-3 protein theta | P68254 | Ywhaq | 28 kDa | 5 C | 1.563 | 0.20772 |
| Ubiquitin-conjugating enzyme E2 variant 1 | Q9CZY3 | Ube2v1 | 16 kDa | 2 C | 1.561 | 0.34459 |
| Isoform Smooth muscle of Myosin light polypeptide 6 | Q60605-2 | Myl6 | 17 kDa | 3 C | 1.558 | 0.10691 |
| Na(+)/H(+) exchange regulatory cofactor NHE-RF2 | Q9JHL1 | Slc9a3r2 | 37 kDa | 6 C | 1.554 | 0.49899 |
| Filamin-B | Q80X90 | Flnb | 278 kDa | 43 C | 1.550 | 0.2674 |
| L-xylulose reductase | Q91X52 | Dcxr | 26 kDa | 4 C | 1.550 | 0.57674 |
| Isoform 2 of Inter alpha-trypsin inhibitor | A6X935-2 | Itih4 | 100 kDa | 3 C | 1.550 | 0.57674 |
| Alpha-centractin | P61164 | Actr1a | 43 kDa | 2 C | 1.550 | 0.57674 |
| Unconventional myosin-VI | Q64331 | Myo6 | 146 kDa | 25 C | 1.548 | 0.20057 |
| Clusterin | Q06890 | Clu | 52 kDa | 11 C | 1.548 | 0.20057 |
| Fibroblast growth factor 1 | P61148 | Fgf1 | 17 kDa | 3 C | 1.548 | 0.20057 |
| GrpE protein homolog 1 | Q99LP6 | Grpel1 | 24 kDa | 4 C | 1.548 | 0.20057 |
| Malignant T-cell-amplified sequence 1 | Q9DB27 | Mcts1 | 21 kDa | 4 C | 1.543 | 0.24087 |
| V-type proton ATPase subunit E 1 | P50518 | Atp6v1e1 | 26 kDa | 1 C | 1.538 | 0.11536 |
| Fatty acid-binding protein | P04117 | Fabp4 | 15 kDa | 2 C | 1.533 | 0.06969 |
| Catenin alpha-1 | P26231 | Ctnna1 | 100 kDa | 12 C | 1.532 | 0.22829 |
| Phosphotriesterase-related protein | Q60866 | Pter | 39 kDa | 6 C | 1.528 | 0.02192 |
| Calpain small subunit 1 | O88456 | Capns1 | 28 kDa | 2 C | 1.527 | 0.07594 |
| Transforming protein RhoA | Q9QUI0 | Rhoa | 22 kDa | 6 C | 1.527 | 0.25775 |
| RNA-binding protein 12 | Q8R4X3 | Rbm12 | 103 kDa | 4 C | 1.527 | 0.54905 |
| Galectin-3 | P16110 | Lgals3 | 28 kDa | 1 C | 1.527 | 0.54905 |
| DnaJ homolog subfamily A member 2 | Q9QYJ0 | Dnaja2 | 46 kDa | 11 C | 1.527 | 0.54905 |
| E3 ubiquitin-protein ligase RNF181 | Q9CY62 | Rnf181 | 19 kDa | 7 C | 1.527 | 0.54905 |
| Beta-ureidopropionase | Q8VC97 | Upb1 | 44 kDa | 10 C | 1.525 | 0.48409 |
| Isoaspartyl peptidase/L-asparaginase | Q8C0M9 | Asrgl1 | 34 kDa | 8 C | 1.520 | 0.42198 |
| Ferritin light chain 1 | P29391 | Ftl1 | 21 kDa | 1 C | 1.520 | 0.17917 |
| Prelamin-A/C | P48678 | Lmna | 74 kDa | 5 C | 1.520 | 0.48246 |
| Inorganic pyrophosphatase | Q9D819 | Ppa1 | 33 kDa | 8 C | 1.511 | 0.26398 |
| Glia maturation factor beta | Q9CQI3 | Gmfb | 17 kDa | 3 C | 1.511 | 0.0141 |
| Lysosomal protective protein | P16675 | Ctsa | 54 kDa | 11 C | 1.510 | 0.30826 |
| Adenylate kinase isoenzyme 1 | Q9R0Y5 | Ak1 | 22 kDa | 2 C | 1.506 | 0.20018 |
| Valacyclovir hydrolase | Q8R164 | Bphl | 33 kDa | 4 C | 1.503 | 0.6592 |
| Density-regulated protein | Q9CQJ6 | Denr | 22 kDa | 7 C | 1.503 | 0.6592 |
| Ig mu chain C region | P01872 | Ighm | 50 kDa | 19 C | 1.494 | 0.08102 |
| L-lactate dehydrogenase A chain | P06151 | Ldha | 36 kDa | 6 C | 1.493 | 0.25695 |
| Isoform 5 of Kinectin | Q61595-5 | Ktn1 | 146 kDa | 10 C | 1.491 | 0.44654 |
| Alpha-1-antitrypsin 1-1 | P07758 | Serpina1a | 46 kDa | 3 C | 1.491 | 0.29316 |
| 60 kDa heat shock protein | P63038 | Hspd1 | 61 kDa | 3 C | 1.490 | 0.09256 |
| Isoform 2 of CUGBP Elav-like family member 1 | P28659-2 | Celf1 | 52 kDa | 7 C | 1.477 | 0.75324 |
| Isoform 2 of Glucosidase 2 subunit beta | O08795-2 | Prkcsh | 60 kDa | 17 C | 1.477 | 0.08841 |
| Putative adenosylhomocysteinase 3 | Q68FL4 | Ahcyl2 | 67 kDa | 18 c | 1.477 | 0.48121 |
| Oxysterol-binding protein 1 | Q3B7Z2 | Osbp | 89 kDa | 15 C | 1.477 | 0.75324 |
| Maleylacetoacetate isomerase | Q9WVL0 | Gstz1 | 24 kDa | 3 C | 1.477 | 0.75324 |
| V-type proton ATPase subunit H | Q8BVE3 | Atp6v1h | 56 kDa | 8 C | 1.477 | 0.75324 |
| UDP-glucuronosyltransferase 3A2 | Q8JZZ0 | Ugt3a2 | 60 kDa | 3 C | 1.477 | 0.75324 |
| Glutathione S-transferase theta-2 | Q61133 | Gstt2 | 28 kDa | 2 C | 1.477 | 0.75324 |
| Beta-glucuronidase | P12265 | Gusb | 74 kDa | 8 C | 1.477 | 0.75324 |
| Costars family protein ABRACL | Q4KML4 | Abracl | 9 kDa | 1 C | 1.477 | 0.75324 |
| Cytochrome b-c1 complex subunit 6 | P99028 | Uqcrh | 10 kDa | 5 C | 1.477 | 0.75324 |
| Isoform 2 of Protein transport protein Sec31A | Q3UPL0-2 | Sec31a | 130 kDa | 18 C | 1.477 | 0.75324 |
| Carboxylesterase 1C | P23953 | Ces1c | 61 kDa | 5 C | 1.475 | 0.39479 |
| Tubulin alpha-4A chain | P68368 | Tuba4a | 50 kDa | 13 C | 1.475 | 0.49712 |
| Nucleolin | P09405 | Ncl | 77 kDa | 1 C | 1.474 | 0.46637 |
| Collagen alpha-1(XVIII) chain | P39061 | Col18a1 | 182 kDa | 8 C | 1.467 | 0.07862 |
| Nucleoside diphosphate kinase A | P15532 | Nme1 | 17 kDa | 2 C | 1.467 | 0.17236 |

| | | | | | | |
|---|---|---|---|---|---|---|
| Isoform 2 of Cordon-bleu protein-like 1 | Q3UMF0-2 | Cobll1 | 133 kDa | 10 C | 1.467 | 0.19962 |
| Polyribonucleotide nucleotidyltransferase 1 | Q8K1R3 | Pnpt1 | 86 kDa | 16 C | 1.467 | 0.248 |
| Basal cell adhesion molecule | Q9R069 | Bcam | 68 kDa | 14 C | 1.467 | 0.28931 |
| Cytochrome c oxidase subunit 5B | P19536 | Cox5b | 14 kDa | 5 C | 1.467 | 0.40708 |
| Galectin-1 | P16045 | Lgals1 | 15 kDa | 6 C | 1.467 | 0.09334 |
| m7GpppX diphosphatase | Q9DAR7 | Dcps | 39 kDa | 2 C | 1.456 | 0.35696 |
| 1-phosphatidylinositol 4 | Q8R3B1 | Plcd1 | 86 kDa | 14 C | 1.456 | 0.39313 |
| 14-3-3 protein gamma | P61982 | Ywhag | 28 kDa | 3 C | 1.455 | 0.0514 |
| N-acetylglucosamine-6-phosphate deacetylase | Q8JZV7 | Amdhd2 | 44 kDa | 8 C | 1.455 | 0.20985 |
| Copper chaperone for superoxide dismutase | Q9WU84 | Ccs | 29 kDa | 10 C | 1.455 | 0.37223 |
| Actin-related protein 2/3 complex subunit 2 | Q9CVB6 | Arpc2 | 34 kDa | 2 C | 1.450 | 0.12208 |
| NADH dehydrogenase [ubiquinone] iron-sulfur protein 3 | Q9DCT2 | Ndufs3 | 30 kDa | 3 C | 1.450 | 0.16352 |
| Heat shock protein 105 kDa | Q61699 | Hsph1 | 96 kDa | 17 C | 1.448 | 0.62222 |
| Isoform 2 of F-actin-capping protein subunit beta | P47757-2 | Capzb | 31 kDa | 5 C | 1.446 | 0.09138 |
| NADH dehydrogenase [ubiquinone] iron-sulfur protein 8 | Q8K3J1 | Ndufs8 | 24 kDa | 8 C | 1.443 | 0.36361 |
| Proteasome subunit beta type-5 | O55234 | Psmb5 | 29 kDa | 3 C | 1.440 | 0.13763 |
| Aldose reductase | P45376 | Akr1b1 | 36 kDa | 6 C | 1.440 | 0.21648 |
| Dynein light chain 2 | Q9D0M5 | Dynll2 | 10 kDa | 2 C | 1.440 | 0.28741 |
| WD repeat-containing protein 1 | O88342 | Wdr1 | 66 kDa | 12 C | 1.440 | 0.45575 |
| 14-3-3 protein beta/alpha | Q9CQV8 | Ywhab | 28 kDa | 2 C | 1.437 | 0.06218 |
| Quinone oxidoreductase-like protein 2 | Q3UNZ8 | | 38 kDa | 9 C | 1.432 | 0.64836 |
| Uromodulin | Q91X17 | Umod | 71 kDa | 48 C | 1.429 | 0.21684 |
| Thioredoxin | P10639 | Txn | 12 kDa | 6 C | 1.426 | 0.32556 |
| ATP synthase subunit gamma | Q91VR2 | Atp5c1 | 33 kDa | 2 C | 1.426 | 0.48893 |
| S-methyl-5'-thioadenosine phosphorylase | Q9CQ65 | Mtap | 31 kDa | 10 C | 1.422 | 0.32757 |
| Uridine diphosphate glucose pyrophosphatase | Q9D142 | Nudt14 | 24 kDa | 4 C | 1.420 | 0.36526 |
| Dual specificity protein phosphatase 3 | Q9D7X3 | Dusp3 | 20 kDa | 4 C | 1.420 | 0.47985 |
| NADH dehydrogenase [ubiquinone] flavoprotein 2 | Q9D6J6 | Ndufv2 | 27 kDa | 6 C | 1.419 | 0.10296 |
| 14-3-3 protein sigma | O70456 | Sfn | 28 kDa | 4 C | 1.412 | 0.23893 |
| Isoform 2 of Carboxypeptidase Q | Q9WVJ3-2 | Cpq | 50 kDa | 1 C | 1.412 | 0.39863 |
| Tubulin alpha-1C chain | P68373 | Tuba1c | 50 kDa | 12 C | 1.412 | 0.43894 |
| Vesicle-associated membrane protein-associated protein B | Q9QY76 | Vapb | 27 kDa | 4 C | 1.405 | 0.59116 |
| Adseverin | Q60604 | Scin | 80 kDa | 4 C | 1.404 | 0.13604 |
| Transgelin | P37804 | Tagln | 23 kDa | 1 C | 1.400 | 0.10133 |
| Protein NDRG1 | Q62433 | Ndrg1 | 43 kDa | 8 C | 1.400 | 0.19301 |
| Elongation factor 1-delta | P57776 | Eef1d | 31 kDa | 2 C | 1.400 | 0.22745 |
| Dihydropteridine reductase | Q8BVI4 | Qdpr | 26 kDa | 4 C | 1.400 | 0.19301 |
| Spectrin beta chain | Q62261 | Sptbn1 | 274 kDa | 15 C | 1.400 | 0.25425 |
| Isocitrate dehydrogenase [NAD] subunit alpha | Q9D6R2 | Idh3a | 40 kDa | 8 C | 1.400 | 0.33525 |
| Triokinase/FMN cyclase | Q8VC30 | Tkfc | 60 kDa | 5 C | 1.393 | 0.65028 |
| Cytochrome P450 4B1 | Q64462 | Cyp4b1 | 59 kDa | 9 C | 1.393 | 0.67039 |
| Methylmalonyl-CoA mutase | P16332 | Mut | 83 kDa | 8 C | 1.391 | 0.10513 |
| 6-pyruvoyl tetrahydrobiopterin synthase | Q9R1Z7 | Pts | 16 kDa | 1 C | 1.385 | 0.49723 |
| Receptor-type tyrosine-protein phosphatase kappa | P35822 | Ptprk | 164 kDa | 35 C | 1.383 | 0.44714 |
| Cathepsin B | P10605 | Ctsb | 37 kDa | 16 C | 1.383 | 0.57069 |
| Protein-glutamine gamma-glutamyltransferase K | Q9JLF6 | Tgm1 | 90 kDa | 16 C | 1.382 | 0.23273 |
| Isochorismatase domain-containing protein 2A | P85094 | Isoc2a | 22 kDa | 6 C | 1.374 | 0.08862 |
| Carboxymethylenebutenolidase homolog | Q8R1G2 | Cmbl | 28 kDa | 6 C | 1.371 | 0.22435 |
| Mitochondrial intermembrane space import and assembly protein 40 | Q8VEA4 | Chchd4 | 16 kDa | 7 C | 1.371 | 0.32819 |
| Exosome complex component RRP43 | Q9D753 | Exosc8 | 30 kDa | 10 C | 1.371 | 0.58396 |
| 4-trimethylaminobutyraldehyde dehydrogenase | Q9JLJ2 | Aldh9a1 | 54 kDa | 17 C | 1.370 | 0.04591 |
| Actin | P60710 | Actb | 42 kDa | 6 C | 1.369 | 0.10709 |
| Enoyl-CoA hydratase | Q8BH95 | Echs1 | 31 kDa | 7 C | 1.367 | 0.06574 |
| Isoform Short of Heterogeneous nuclear ribonucleoprotein A1 | P49312-2 | Hnrnpa1 | 29 kDa | 2 C | 1.367 | 0.09884 |
| Acyl-coenzyme A amino acid N-acyltransferase 1 | A2AKK5 | Acnat1 | 46 kDa | 7 C | 1.367 | 0.14994 |
| UDP-N-acetylhexosamine pyrophosphorylase-like protein 1 | Q3TW96 | Uap1l1 | 57 kDa | 13 C | 1.363 | 0.42398 |
| Beta-arrestin-1 | Q8BWG8 | Arrb1 | 47 kDa | 8 C | 1.360 | 0.18066 |
| Mitochondrial proton/calcium exchanger protein | Q9Z2I0 | Letm1 | 83 kDa | 12 C | 1.360 | 0.30309 |
| SH3 domain-binding glutamic acid-rich-like protein | Q9JJU8 | Sh3bgrl | 13 kDa | 2 C | 1.360 | 0.37757 |
| Ig kappa chain C region | P01837 | | 12 kDa | 3 C | 1.360 | 0.03855 |

| | | | | | | |
|---|---|---|---|---|---|---|
| Calreticulin | P14211 | Calr | 48 kDa | 6 C | 1.356 | 0.06338 |
| Histidine triad nucleotide-binding protein 1 | P70349 | Hint1 | 14 kDa | 2 C | 1.354 | 0.23088 |
| Tetratricopeptide repeat protein 38 | A3KMP2 | Ttc38 | 52 kDa | 9 C | 1.354 | 0.04463 |
| GMP reductase 1 | Q9DCZ1 | Gmpr | 37 kDa | 9 C | 1.352 | 0.05394 |
| NADH dehydrogenase [ubiquinone] 1 alpha subcomplex subunit 10 | Q99LC3 | Ndufa10 | 41 kDa | 5 C | 1.352 | 0.66663 |
| Aquaporin-1 | Q02013 | Aqp1 | 29 kDa | 4 C | 1.352 | 0.66663 |
| Carboxylesterase 1D | Q8VCT4 | Ces1d | 62 kDa | 5 C | 1.350 | 0.28067 |
| Isoform 2 of Cystathionine beta-synthase | Q91WT9-2 | Cbs | 60 kDa | 13 C | 1.349 | 0.24194 |
| Serine/threonine-protein phosphatase PP1-beta catalytic subunit | P62141 | Ppp1cb | 37 kDa | 14 C | 1.349 | 0.10426 |
| Alpha-actinin-4 | P57780 | Actn4 | 105 kDa | 8 C | 1.348 | 0.04432 |
| Plastin-3 | Q99K51 | Pls3 | 71 kDa | 9 C | 1.347 | 0.37861 |
| Heat shock 70 kDa protein 4 | Q61316 | Hspa4 | 94 kDa | 14 C | 1.346 | 0.09856 |
| Methylthioribulose-1-phosphate dehydratase | Q9WVQ5 | Apip | 27 kDa | 11 C | 1.345 | 0.24087 |
| Poly(rC)-binding protein 1 | P60335 | Pcbp1 | 37 kDa | 9 C | 1.344 | 0.23179 |
| 14-3-3 protein eta | P68510 | Ywhah | 28 kDa | 3 C | 1.342 | 0.28616 |
| Immunoglobulin-binding protein 1 | Q61249 | Igbp1 | 39 kDa | 3 C | 1.342 | 0.62652 |
| Endonuclease G | O08600 | Endog | 32 kDa | 2 C | 1.342 | 0.62652 |
| 39S ribosomal protein L49 | Q9CQ40 | Mrpl49 | 19 kDa | 1 C | 1.333 | 0.54906 |
| Methylglutaconyl-CoA hydratase | Q9JLZ3 | Auh | 33 kDa | 5 C | 1.333 | 0.05815 |
| Clathrin light chain A | O08585 | Clta | 26 kDa | 1 C | 1.333 | 0.08922 |
| Ig kappa chain V-V region K2 (Fragment) | P01635 | | 13 kDa | 3 C | 1.333 | 0.08922 |
| Histidine-rich glycoprotein | Q9ESB3 | Hrg | 59 kDa | 17 C | 1.333 | 0.09956 |
| Endoplasmic reticulum resident protein 44 | Q9D1Q6 | Erp44 | 47 kDa | 7 C | 1.333 | 0.11765 |
| Proteasome subunit alpha type-3 | O70435 | Psma3 | 28 kDa | 4 C | 1.333 | 0.13952 |
| Acyl-coenzyme A thioesterase THEM4 | Q3UUI3 | Them4 | 26 kDa | 4 C | 1.333 | 0.17047 |
| Macrophage-capping protein | P24452 | Capg | 39 kDa | 5 C | 1.333 | 0.19446 |
| Beta-mannosidase | Q8K2I4 | Manba | 101 kDa | 12 C | 1.333 | 0.21684 |
| Extracellular superoxide dismutase [Cu-Zn] | O09164 | Sod3 | 27 kDa | 6 C | 1.333 | 0.2675 |
| Tropomodulin-3 | Q9JHJ0 | Tmod3 | 40 kDa | 2 C | 1.333 | 0.29235 |
| Isoform 2 of Methionine adenosyltransferase 2 subunit beta | Q99LB6-2 | Mat2b | 36 kDa | 7 C | 1.333 | 0.29235 |
| Sorting nexin-12 | O70493 | Snx12 | 19 kDa | 3 C | 1.333 | 0.29235 |
| Complement component C8 beta chain | Q8BH35 | C8b | 66 kDa | 32 C | 1.333 | 0.35062 |
| Isoform 1 of Afadin | Q9QZQ1-2 | Afdn | 205 kDa | 11 C | 1.333 | 0.48033 |
| Peroxisomal sarcosine oxidase | Q9D826 | Pipox | 44 kDa | 11 C | 1.333 | 0.02267 |
| Alpha-actinin-1 | Q7TPR4 | Actn1 | 103 kDa | 11 C | 1.327 | 0.20734 |
| 39S ribosomal protein L39 | Q9JKF7 | Mrpl39 | 39 kDa | 7 C | 1.325 | 0.79102 |
| Lipoamide acyltransferase component of branched-chain alpha-keto acid dehydrogenase | P53395 | Dbt | 53 kDa | 6 C | 1.325 | 0.62175 |
| Multifunctional protein ADE2 | Q9DCL9 | Paics | 47 kDa | 13 C | 1.325 | 0.47196 |
| Superoxide dismutase [Mn] | P09671 | Sod2 | 25 kDa | 4 C | 1.324 | 0.03149 |
| Isoform 2 of Heterogeneous nuclear ribonucleoprotein K | P61979-2 | Hnrnpk | 51 kDa | 5 C | 1.319 | 0.29472 |
| 2-oxoglutarate dehydrogenase | Q60597 | Ogdh | 116 kDa | 21 C | 1.319 | 0.08504 |
| Isoform 2 of Poly(rC)-binding protein 2 | Q61990-2 | Pcbp2 | 35 kDa | 7 C | 1.318 | 0.19702 |
| Microtubule-associated protein RP/EB family member 1 | Q61166 | Mapre1 | 30 kDa | 3 C | 1.316 | 0.55382 |
| Ig heavy chain V region 36-65 | P01747 | | 13 kDa | 2 C | 1.316 | 0.55382 |
| Elongation factor 2 | P58252 | Eef2 | 95 kDa | 7 C | 1.315 | 0.29458 |
| Protein disulfide-isomerase A3 | P27773 | Pdia3 | 57 kDa | 8 C | 1.314 | 0.14897 |
| Actin-related protein 2/3 complex subunit 1A | Q9R0Q6 | Arpc1a | 42 kDa | 10 C | 1.314 | 0.22637 |
| 55 kDa erythrocyte membrane protein | P70290 | Mpp1 | 52 kDa | 5 C | 1.314 | 0.28741 |
| Histidine triad nucleotide-binding protein 2 | Q9D0S9 | Hint2 | 17 kDa | 1 C | 1.314 | 0.44086 |
| Beta-1 | Q09324 | Gcnt1 | 50 kDa | 11 C | 1.311 | 0.41649 |
| 14-3-3 protein epsilon | P62259 | Ywhae | 29 kDa | 3 C | 1.311 | 0.13461 |
| Secernin-2 | Q8VCA8 | Scrn2 | 47 kDa | 10 C | 1.309 | 0.20402 |
| Ubiquitin carboxyl-terminal hydrolase 5 | P56399 | Usp5 | 96 kDa | 16 C | 1.309 | 0.49127 |
| Aldehyde dehydrogenase | P47738 | Aldh2 | 57 kDa | 9 C | 1.309 | 0.03568 |
| Serum albumin | P07724 | Alb | 69 kDa | 36 C | 1.308 | 0.02005 |
| Stress-70 protein | P38647 | Hspa9 | 73 kDa | 5 C | 1.307 | 0.22541 |
| Isoform 2 of Ig gamma-2B chain C region | P01867-2 | Igh-3 | 37 kDa | 13 C | 1.303 | 0.72702 |
| Fructose-bisphosphate aldolase A | P05064 | Aldoa | 39 kDa | 8 C | 1.302 | 0.07152 |
| Complement C3 | P01027 | C3 | 186 kDa | 27 C | 1.302 | 0.17632 |
| Methylcrotonoyl-CoA carboxylase beta chain | Q3ULD5 | Mccc2 | 61 kDa | 10 C | 1.301 | 0.04308 |

| | | | | | | |
|---|---|---|---|---|---|---|
| Cell division control protein 42 homolog | P60766 | Cdc42 | 21 kDa | 7 C | 1.300 | 0.06758 |
| Cytospin-B | Q5SXY1 | Specc1 | 118 kDa | 12 C | 1.300 | 0.71228 |
| 3-mercaptopyruvate sulfurtransferase | Q99J99 | Mpst | 33 kDa | 4 C | 1.300 | 0.71228 |
| Protein DEK | Q7TNV0 | Dek | 43 kDa | 4 C | 1.300 | 0.71228 |
| Epidermal growth factor receptor kinase substrate 8-like protein 2 | Q99K30 | Eps8l2 | 82 kDa | 9 C | 1.300 | 0.74876 |
| Electron transfer flavoprotein subunit beta | Q9DCW4 | Etfb | 28 kDa | 4 C | 1.297 | 0.04063 |
| Extracellular matrix protein 1 | Q61508 | Ecm1 | 63 kDa | 29 C | 1.296 | 0.46922 |
| Molybdenum cofactor sulfurase | Q14CH1 | Mocos | 95 kDa | 24 C | 1.296 | 0.46922 |
| Eukaryotic translation initiation factor 5A-1 | P63242 | Eif5a | 17 kDa | 4 C | 1.295 | 0.2315 |
| Selenide | Q8BH69 | Sephs1 | 43 kDa | 9 C | 1.292 | 0.20107 |
| Glutaredoxin-3 | Q9CQM9 | Glrx3 | 38 kDa | 5 C | 1.292 | 0.26467 |
| UPF0160 protein MYG1 | Q9JK81 | Myg1 | 43 kDa | 7 C | 1.292 | 0.34262 |
| Calbindin | P12658 | Calb1 | 30 kDa | 4 C | 1.292 | 0.49843 |
| von Willebrand factor A domain-containing protein 5A | Q99KC8 | Vwa5a | 87 kDa | 13 C | 1.290 | 0.49047 |
| NADH dehydrogenase [ubiquinone] flavoprotein 1 | Q91YT0 | Ndufv1 | 51 kDa | 12 C | 1.289 | 0.18517 |
| Dihydropyrimidinase-related protein 2 | O08553 | Dpysl2 | 62 kDa | 7 C | 1.288 | 0.14712 |
| Phosphate carrier protein | Q8VEM8 | Slc25a3 | 40 kDa | 8 C | 1.288 | 0.68531 |
| Tubulin beta-4B chain | P68372 | Tubb4b | 50 kDa | 8 C | 1.287 | 0.6227 |
| NADH dehydrogenase [ubiquinone] 1 alpha subcomplex subunit 9 | Q9DC69 | Ndufa9 | 43 kDa | 2 C | 1.286 | 0.763 |
| GDP-mannose 4 | Q8K0C9 | Gmds | 42 kDa | 6 C | 1.286 | 0.10533 |
| Delta-aminolevulinic acid dehydratase | P10518 | Alad | 36 kDa | 8 C | 1.283 | 0.11406 |
| NADH dehydrogenase [ubiquinone] 1 beta subcomplex subunit 10 | Q9DCS9 | Ndufb10 | 21 kDa | 5 C | 1.282 | 0.57699 |
| Serine/threonine-protein phosphatase PP1-alpha catalytic subunit | P62137 | Ppp1ca | 38 kDa | 13 C | 1.280 | 0.08674 |
| Actin-related protein 2/3 complex subunit 3 | Q9JM76 | Arpc3 | 21 kDa | 4 C | 1.280 | 0.28931 |
| DAZ-associated protein 1 | Q9JII5 | Dazap1 | 43 kDa | 4 C | 1.280 | 0.32488 |
| TAR DNA-binding protein 43 | Q921F2 | Tardbp | 45 kDa | 7 C | 1.280 | 0.35582 |
| Cytidine deaminase | P56389 | Cda | 16 kDa | 7 C | 1.280 | 0.3736 |
| Serine beta-lactamase-like protein LACTB | Q9EP89 | Lactb | 61 kDa | 4 C | 1.280 | 0.38014 |
| V-type proton ATPase subunit G 1 | Q9CR51 | Atp6v1g1 | 14 kDa | 2 C | 1.280 | 0.50972 |
| Ig kappa chain V-II region 7S34.1 | P01630 | | 12 kDa | 3 C | 1.280 | 0.50972 |
| 3'(2') | Q9Z0S1 | Bpnt1 | 33 kDa | 6 C | 1.277 | 0.01772 |
| Nucleoside diphosphate kinase B | Q01768 | Nme2 | 17 kDa | 2 C | 1.277 | 0.21116 |
| Scaffold attachment factor B1 | D3YXK2 | Safb | 105 kDa | 9 C | 1.275 | 0.68707 |
| Isoform CW17E of Splicing factor 1 | Q64213-2 | Sf1 | 60 kDa | 4 C | 1.275 | 0.68707 |
| Branched-chain-amino-acid aminotransferase | P24288 | Bcat1 | 43 kDa | 10 C | 1.275 | 0.68707 |
| Isoform 2 of Leukocyte surface antigen CD47 | Q61735-2 | Cd47 | 35 kDa | 11 C | 1.275 | 0.68707 |
| 14-3-3 protein zeta/delta | P63101 | Ywhaz | 28 kDa | 3 C | 1.274 | 0.1477 |
| Beta-2-glycoprotein 1 | Q01339 | Apoh | 39 kDa | 23 C | 1.273 | 0.58567 |
| Adenylyl cyclase-associated protein 1 | P40124 | Cap1 | 52 kDa | 6 C | 1.273 | 0.51649 |
| S-formylglutathione hydrolase | Q9R0P3 | Esd | 31 kDa | 10 C | 1.273 | 0.51649 |
| Transketolase | P40142 | Tkt | 68 kDa | 12 C | 1.272 | 0.03958 |
| Aldehyde dehydrogenase family 8 member A1 | Q8BH00 | Aldh8a1 | 54 kDa | 13 C | 1.271 | 0.41934 |
| ADP/ATP translocase 2 | P51881 | Slc25a5 | 33 kDa | 4 C | 1.271 | 0.61419 |
| Vitamin D-binding protein | P21614 | Gc | 54 kDa | 28 C | 1.269 | 0.22591 |
| Isoform 3 of Kininogen-1 | O08677-3 | Kng1 | 53 kDa | 19 C | 1.269 | 0.29196 |
| V-type proton ATPase subunit B | P62814 | Atp6v1b2 | 57 kDa | 6 C | 1.269 | 0.43313 |
| Adenosine kinase | P55264 | Adk | 40 kDa | 6 C | 1.267 | 0.19301 |
| Acyl-coenzyme A synthetase ACSM1 | Q91VA0 | Acsm1 | 65 kDa | 14 C | 1.267 | 0.21285 |
| Fetuin-B | Q9QXC1 | Fetub | 43 kDa | 15 C | 1.267 | 0.24904 |
| 40S ribosomal protein SA | P14206 | Rpsa | 33 kDa | 2 C | 1.267 | 0.0278 |
| Fibrinogen gamma chain | Q8VCM7 | Fgg | 49 kDa | 12 C | 1.265 | 0.08108 |
| Protein farnesyltransferase/geranylgeranyltransferase type-1 subunit alpha | Q61239 | Fnta | 44 kDa | 3 C | 1.263 | 0.58099 |
| Annexin A2 | P07356 | Anxa2 | 39 kDa | 5 C | 1.262 | 0.11406 |
| Hydroxyacid oxidase 2 | Q9NYQ2 | Hao2 | 39 kDa | 8 C | 1.262 | 0.37817 |
| Acyl-coenzyme A thioesterase 1 | O55137 | Acot1 | 46 kDa | 4 C | 1.262 | 0.73793 |
| Inorganic pyrophosphatase 2 | Q91VM9 | Ppa2 | 38 kDa | 8 C | 1.257 | 0.16708 |
| Selenide | P97364 | Sephs2 | 48 kDa | 7 C | 1.257 | 0.28283 |
| Septin-7 | O55131 | 43350 | 51 kDa | 6 C | 1.257 | 0.5091 |
| Aflatoxin B1 aldehyde reductase member 2 | Q8CG76 | Akr7a2 | 41 kDa | 8 C | 1.252 | 0.21447 |
| Thiosulfate sulfurtransferase | P52196 | Tst | 33 kDa | 4 C | 1.251 | 0.63842 |

| | | | | | | |
|---|---|---|---|---|---|---|
| Proteasome subunit alpha type-7 | Q9Z2U0 | Psma7 | 28 kDa | 3 C | 1.250 | 0.08922 |
| DnaJ homolog subfamily C member 12 | Q9R022 | Dnajc12 | 23 kDa | 4 C | 1.250 | 0.46455 |
| Peroxisomal bifunctional enzyme | Q9DBM2 | Ehhadh | 78 kDa | 10 C | 1.250 | 0.60479 |
| UMP-CMP kinase | Q9DBP5 | Cmpk1 | 22 kDa | 6 C | 1.249 | 0.17133 |
| Transitional endoplasmic reticulum ATPase | Q01853 | Vcp | 89 kDa | 12 C | 1.245 | 0.12388 |
| Serine/threonine-protein phosphatase 5 | Q60676 | Ppp5c | 57 kDa | 11 C | 1.244 | 0.42092 |
| UPF0598 protein C8orf82 homolog | Q8VE95 | | 24 kDa | 6 C | 1.244 | 0.45575 |
| Harmonin | Q9ES64 | Ush1c | 102 kDa | 7 C | 1.244 | 0.45575 |
| Succinate dehydrogenase [ubiquinone] flavoprotein subunit | Q8K2B3 | Sdha | 73 kDa | 19 C | 1.244 | 0.08065 |
| Kinesin-1 heavy chain | Q61768 | Kif5b | 110 kDa | 13 C | 1.244 | 0.72801 |
| Aspartyl aminopeptidase | Q9Z2W0 | Dnpep | 52 kDa | 10 C | 1.243 | 0.31929 |
| Polymerase delta-interacting protein 2 | Q91VA6 | Poldip2 | 42 kDa | 4 C | 1.240 | 0.11239 |
| Isoform 3 of Disks large homolog 1 | Q811D0-3 | Dlg1 | 103 kDa | 7 C | 1.240 | 0.37357 |
| 2-aminoethanethiol dioxygenase | Q6PDY2 | Ado | 28 kDa | 7 C | 1.240 | 0.63307 |
| Ribosome-binding protein 1 | Q99PL5 | Rrbp1 | 173 kDa | 8 C | 1.238 | 0.58446 |
| Ig alpha chain C region | P01878 | | 37 kDa | 13 C | 1.236 | 0.79497 |
| Na(+)/H(+) exchange regulatory cofactor NHE-RF1 | P70441 | Slc9a3r1 | 39 kDa | 5 C | 1.236 | 0.20255 |
| Mannose-binding protein A | P39039 | Mbl1 | 25 kDa | 8 C | 1.236 | 0.39346 |
| Regulator of microtubule dynamics protein 3 | Q3UJU9 | Rmdn3 | 52 kDa | 6 C | 1.236 | 0.6537 |
| Triosephosphate isomerase | P17751 | Tpi1 | 32 kDa | 9 C | 1.236 | 0.22609 |
| Aspartoacylase | Q8R3P0 | Aspa | 35 kDa | 8 C | 1.232 | 0.28619 |
| Isoform 3 of Heterogeneous nuclear ribonucleoprotein D0 | Q60668-3 | Hnrnpd | 33 kDa | 3 C | 1.232 | 0.35076 |
| Proteasome subunit alpha type-5 | Q9Z2U1 | Psma5 | 26 kDa | 3 C | 1.231 | 0.11765 |
| Pyridoxal kinase | Q8K183 | Pdxk | 35 kDa | 5 C | 1.231 | 0.27605 |
| Glutaryl-CoA dehydrogenase | Q60759 | Gcdh | 49 kDa | 9 C | 1.231 | 0.65715 |
| S-adenosylhomocysteine hydrolase-like protein 1 | Q80SW1 | Ahcyl1 | 59 kDa | 19 C | 1.231 | 0.6783 |
| Plasmalemma vesicle-associated protein | Q91VC4 | Plvap | 50 kDa | 10 C | 1.229 | 0.79592 |
| Succinate-semialdehyde dehydrogenase | Q8BWF0 | Aldh5a1 | 56 kDa | 10 C | 1.223 | 0.12466 |
| Aspartate aminotransferase | P05201 | Got1 | 46 kDa | 5 C | 1.222 | 0.27314 |
| Isoform 2 of Adenylate kinase 2 | Q9WTP6-2 | Ak2 | 26 kDa | 5 C | 1.222 | 0.19798 |
| Peptidyl-prolyl cis-trans isomerase A | P17742 | Ppia | 18 kDa | 3 C | 1.219 | 0.40283 |
| Cystathionine gamma-lyase | Q8VCN5 | Cth | 44 kDa | 11 C | 1.216 | 0.1674 |
| Aconitate hydratase | Q99KI0 | Aco2 | 85 kDa | 13 C | 1.216 | 0.0742 |
| NADH-ubiquinone oxidoreductase 75 kDa subunit | Q91VD9 | Ndufs1 | 80 kDa | 18 C | 1.215 | 0.29682 |
| 2-iminobutanoate/2-iminopropanoate deaminase | P52760 | Rida | 14 kDa | 1 C | 1.213 | 0.28741 |
| Isoform 2 of Heterogeneous nuclear ribonucleoprotein A3 | Q8BG05-2 | Hnrnpa3 | 37 kDa | 4 C | 1.213 | 0.28002 |
| Cytosolic 10-formyltetrahydrofolate dehydrogenase | Q8R0Y6 | Aldh1l1 | 99 kDa | 15 C | 1.211 | 0.40589 |
| Fructose-1 | P70695 | Fbp2 | 37 kDa | 5 C | 1.210 | 0.1932 |
| Polyadenylate-binding protein 1 | P29341 | Pabpc1 | 71 kDa | 4 C | 1.209 | 0.35105 |
| Transaldolase | Q93092 | Taldo1 | 37 kDa | 3 C | 1.208 | 0.36446 |
| Fumarylacetoacetase | P35505 | Fah | 46 kDa | 6 C | 1.207 | 0.17938 |
| Isoform 2 of Tropomyosin alpha-3 chain | P21107-2 | Tpm3 | 29 kDa | 1 C | 1.205 | 0.30844 |
| 3-ketoacyl-CoA thiolase | Q8BWT1 | Acaa2 | 42 kDa | 8 C | 1.200 | 0.63174 |
| Dihydropyrimidinase | Q9EQF5 | Dpys | 57 kDa | 9 C | 1.200 | 0.10901 |
| Fructose-bisphosphate aldolase C | P05063 | Aldoc | 39 kDa | 7 C | 1.200 | 0.11203 |
| Cytochrome c | P62897 | Cycs | 12 kDa | 2 C | 1.200 | 0.11406 |
| Protein phosphatase 1 regulatory subunit 7 | Q3UM45 | Ppp1r7 | 41 kDa | 2 C | 1.200 | 0.13939 |
| Twinfilin-1 | Q91YR1 | Twf1 | 40 kDa | 4 C | 1.200 | 0.22745 |
| Insulin-degrading enzyme | Q9JHR7 | Ide | 118 kDa | 13 C | 1.200 | 0.26183 |
| Malate dehydrogenase | P14152 | Mdh1 | 37 kDa | 3 C | 1.200 | 0.28611 |
| Protein disulfide-isomerase A4 | P08003 | Pdia4 | 72 kDa | 6 C | 1.200 | 0.28741 |
| Probable aminopeptidase NPEPL1 | Q6NSR8 | Npepl1 | 56 kDa | 16 C | 1.200 | 0.34741 |
| Phosphoglycerate kinase 1 | P09411 | Pgk1 | 45 kDa | 7 C | 1.200 | 0.38779 |
| Regulator of microtubule dynamics protein 1 | Q9DCV4 | Rmdn1 | 35 kDa | 5 C | 1.200 | 0.40708 |
| Signal transducing adapter molecule 1 | P70297 | Stam | 60 kDa | 7 C | 1.200 | 0.40708 |
| Proteasome subunit beta type-6 | Q60692 | Psmb6 | 25 kDa | 4 C | 1.200 | 0.40708 |
| Destrin | Q9R0P5 | Dstn | 19 kDa | 6 C | 1.200 | 0.45379 |
| Serine/threonine-protein phosphatase 6 catalytic subunit | Q9CQR6 | Ppp6c | 35 kDa | 12 C | 1.200 | 0.53562 |
| Isoform 2 of Neutral alpha-glucosidase AB | Q8BHN3-2 | Ganab | 109 kDa | 8 C | 1.200 | 0.54778 |
| Heme-binding protein 1 | Q9R257 | Hebp1 | 21 kDa | 2 C | 1.200 | 0.56303 |

| | | | | | | |
|---|---|---|---|---|---|---|
| Protein canopy homolog 2 | Q9QXT0 | Cnpy2 | 21 kDa | 6 C | 1.200 | 0.56303 |
| ADP-ribosylation factor-binding protein GGA1 | Q8R0H9 | Gga1 | 70 kDa | 6 C | 1.200 | 0.57083 |
| Heat shock protein HSP 90-alpha | P07901 | Hsp90aa1 | 85 kDa | 7 C | 1.200 | 0.64702 |
| Pro-cathepsin H | P49935 | Ctsh | 37 kDa | 10 C | 1.200 | 0.64702 |
| Ubiquitin carboxyl-terminal hydrolase isozyme L3 | Q9JKB1 | Uchl3 | 26 kDa | 3 C | 1.200 | 0.75939 |
| MAGUK p55 subfamily member 6 | Q9JLB0 | Mpp6 | 63 kDa | 5 C | 1.200 | 0.82713 |
| Cob(I)yrinic acid a | Q9D273 | Mmab | 26 kDa | 5 C | 1.200 | 0.82713 |
| Small nuclear ribonucleoprotein Sm D3 | P62320 | Snrpd3 | 14 kDa | 2 C | 1.200 | 0.82713 |
| Protein arginine methyltransferase NDUFAF7 | Q9CWG8 | Ndufaf7 | 48 kDa | 7 C | 1.200 | 0.82713 |
| Protein disulfide-isomerase | P09103 | P4hb | 57 kDa | 7 C | 1.198 | 0.0653 |
| Actin | P68033 | Actc1 | 42 kDa | 6 C | 1.194 | 0.23858 |
| DNA-directed RNA polymerases I | Q923G2 | Polr2h | 17 kDa | 1 C | 1.190 | 0.72398 |
| Ubiquitin-like protein ISG15 | Q64339 | Isg15 | 18 kDa | 3 C | 1.190 | 0.72398 |
| Adrenodoxin | P46656 | Fdx1 | 20 kDa | 8 C | 1.190 | 0.76953 |
| Heterogeneous nuclear ribonucleoprotein Q | Q7TMK9 | Syncrip | 70 kDa | 4 C | 1.190 | 0.54675 |
| Inositol monophosphatase 1 | O55023 | Impa1 | 30 kDa | 6 C | 1.188 | 0.40475 |
| Isoform 2 of Nitrilase homolog 1 | Q8VDK1-2 | Nit1 | 32 kDa | 13 C | 1.187 | 0.32774 |
| Cytochrome b-c1 complex subunit 1 | Q9CZ13 | Uqcrc1 | 53 kDa | 11 C | 1.187 | 0.248 |
| Fructose-bisphosphate aldolase B | Q91Y97 | Aldob | 40 kDa | 8 C | 1.185 | 0.0445 |
| Ig heavy chain V region AC38 205.12 | P06330 | | 13 kDa | 2 C | 1.184 | 0.15173 |
| ATP synthase subunit d | Q9DCX2 | Atp5h | 19 kDa | 1 C | 1.184 | 0.67276 |
| Villin-1 | Q62468 | Vil1 | 93 kDa | 9 C | 1.183 | 0.08073 |
| Peroxisomal acyl-coenzyme A oxidase 2 | Q9QXD1 | Acox2 | 77 kDa | 12 C | 1.183 | 0.89655 |
| Quinone oxidoreductase | P47199 | Cryz | 35 kDa | 5 C | 1.182 | 0.11415 |
| Heterogeneous nuclear ribonucleoprotein A3 | Q8BG05 | Hnrnpa3 | 40 kDa | 4 C | 1.181 | 0.27461 |
| Cathepsin Z | Q9WUU7 | Ctsz | 34 kDa | 12 C | 1.181 | 0.83451 |
| Quinone oxidoreductase-like protein 1 | Q921W4 | Cryzl1 | 39 kDa | 6 C | 1.179 | 0.36446 |
| Cadherin-16 | O88338 | Cdh16 | 90 kDa | 7 C | 1.179 | 0.45475 |
| Hydroxymethylglutaryl-CoA lyase | P38060 | Hmgcl | 34 kDa | 8 C | 1.177 | 0.13417 |
| Coactosin-like protein | Q9CQI6 | Cotl1 | 16 kDa | 2 C | 1.176 | 0.21684 |
| Isoform Gamma-2 of Serine/threonine-protein phosphatase PP1-gamma catalytic subunit | P63087-2 | Ppp1cc | 39 kDa | 13 C | 1.176 | 0.33734 |
| Heterogeneous nuclear ribonucleoprotein H2 | P70333 | Hnrnph2 | 49 kDa | 5 C | 1.176 | 0.55479 |
| Meprin A subunit alpha | P28825 | Mep1a | 84 kDa | 19 C | 1.176 | 0.23805 |
| Enoyl-CoA hydratase domain-containing protein 2 | Q3TLP5 | Echdc2 | 32 kDa | 6 C | 1.173 | 0.28704 |
| Glutathione peroxidase 3 | P46412 | Gpx3 | 25 kDa | 3 C | 1.173 | 0.48246 |
| Transcription elongation factor A protein 1 | P10711 | Tcea1 | 34 kDa | 8 C | 1.173 | 0.67321 |
| Trifunctional enzyme subunit beta | Q99JY0 | Hadhb | 51 kDa | 5 C | 1.172 | 0.8846 |
| Acyl-coenzyme A thioesterase 4 | Q8BWN8 | Acot4 | 46 kDa | 6 C | 1.171 | 0.28611 |
| Oxygen-dependent coproporphyrinogen-III oxidase | P36552 | Cpox | 50 kDa | 10 C | 1.171 | 0.41467 |
| Purine nucleoside phosphorylase | P23492 | Pnp | 32 kDa | 5 C | 1.169 | 0.44257 |
| Mitotic checkpoint protein BUB3 | Q9WVA3 | Bub3 | 37 kDa | 7 C | 1.169 | 0.28741 |
| Glutathione S-transferase P 1 | P19157 | Gstp1 | 24 kDa | 3 C | 1.168 | 0.41167 |
| Hydroxyacid-oxoacid transhydrogenase | Q8R0N6 | Adhfe1 | 50 kDa | 8 C | 1.167 | 0.35697 |
| Isoform Cytoplasmic of Fumarate hydratase | P97807-2 | Fh | 50 kDa | 4 C | 1.167 | 0.55265 |
| Dihydropyrimidinase-related protein 3 | Q62188 | Dpysl3 | 62 kDa | 7 C | 1.166 | 0.25775 |
| Secernin-3 | Q3TMH2 | Scrn3 | 48 kDa | 7 C | 1.164 | 0.73702 |
| Methyltransferase-like 26 | Q9DCS2 | Mettl26 | 23 kDa | 6 C | 1.164 | 0.19702 |
| Ribokinase | Q8R1Q9 | Rbks | 34 kDa | 9 C | 1.164 | 0.53892 |
| S-phase kinase-associated protein 1 | Q9WTX5 | Skp1 | 19 kDa | 3 C | 1.164 | 0.55146 |
| Splicing factor | Q8VIJ6 | Sfpq | 75 kDa | 7 C | 1.164 | 0.59304 |
| Dipeptidyl peptidase 4 | P28843 | Dpp4 | 87 kDa | 12 C | 1.164 | 0.66626 |
| Phospholipid hydroperoxide glutathione peroxidase | O70325 | Gpx4 | 22 kDa | 10 C | 1.163 | 0.31543 |
| Isoform 3 of Synaptic functional regulator FMR1 | P35922-3 | Fmr1 | 66 kDa | 5 C | 1.162 | 0.79513 |
| Acyl-CoA dehydrogenase family member 10 | Q8K370 | Acad10 | 119 kDa | 16 C | 1.160 | 0.36898 |
| Isoform Cytoplasmic+peroxisomal of Peroxiredoxin-5 | P99029-2 | Prdx5 | 17 kDa | 6 C | 1.159 | 0.10084 |
| Sodium/potassium-transporting ATPase subunit alpha-1 | Q8VDN2 | Atp1a1 | 113 kDa | 23 C | 1.159 | 0.66063 |
| Glyceraldehyde-3-phosphate dehydrogenase | P16858 | Gapdh | 36 kDa | 5 C | 1.157 | 0.39189 |
| Phosphoglucomutase-1 | Q9D0F9 | Pgm1 | 61 kDa | 10 C | 1.156 | 0.48156 |
| Thioredoxin reductase 1 | Q9JMH6 | Txnrd1 | 67 kDa | 21 C | 1.156 | 0.11536 |
| Actin-related protein 3 | Q99JY9 | Actr3 | 47 kDa | 8 C | 1.154 | 0.52284 |

| | | | | | | |
|---|---|---|---|---|---|---|
| Peroxiredoxin-2 | Q61171 | Prdx2 | 22 kDa | 3 C | 1.154 | 0.50561 |
| Heat shock 70 kDa protein 1B | P17879 | Hspa1b | 70 kDa | 5 C | 1.150 | 0.50235 |
| Peroxiredoxin-1 | P35700 | Prdx1 | 22 kDa | 4 C | 1.150 | 0.17762 |
| Isoform Cytoplasmic of Glutathione reductase | P47791-2 | Gsr | 51 kDa | 11 C | 1.148 | 0.30487 |
| UDP-glucose 6-dehydrogenase | O70475 | Ugdh | 55 kDa | 12 C | 1.148 | 0.89094 |
| DnaJ homolog subfamily C member 2 | P54103 | Dnajc2 | 72 kDa | 8 C | 1.148 | 0.89094 |
| Phosphoenolpyruvate carboxykinase | Q9Z2V4 | Pck1 | 69 kDa | 13 C | 1.145 | 0.88306 |
| 78 kDa glucose-regulated protein | P20029 | Hspa5 | 72 kDa | 1 C | 1.145 | 0.23526 |
| Malectin | Q6ZQI3 | Mlec | 32 kDa | 3 C | 1.143 | 0.11765 |
| Antithrombin-III | P32261 | Serpinc1 | 52 kDa | 9 C | 1.143 | 0.48033 |
| Heat shock cognate 71 kDa protein | P63017 | Hspa8 | 71 kDa | 4 C | 1.143 | 0.49394 |
| Plastin-2 | Q61233 | Lcp1 | 70 kDa | 11 C | 1.143 | 0.63258 |
| Coatomer subunit delta | Q5XJY5 | Arcn1 | 57 kDa | 8 C | 1.143 | 0.66466 |
| Cytochrome c oxidase subunit 5A | P12787 | Cox5a | 16 kDa | 4 C | 1.143 | 0.69323 |
| Caprin-1 | Q60865 | Caprin1 | 78 kDa | 3 C | 1.143 | 0.70625 |
| Vesicle-associated membrane protein-associated protein A | Q9WV55 | Vapa | 28 kDa | 4 C | 1.143 | 0.77236 |
| Thioredoxin-like protein 1 | Q8CDN6 | Txnl1 | 32 kDa | 7 C | 1.138 | 0.34334 |
| Alpha-1-antitrypsin 1-4 | Q00897 | Serpina1d | 46 kDa | 3 C | 1.138 | 0.67112 |
| Bifunctional glutamate/proline--tRNA ligase | Q8CGC7 | Eprs | 170 kDa | 31 C | 1.138 | 0.80375 |
| Calcyclin-binding protein | Q9CXW3 | Cacybp | 27 kDa | 2 C | 1.136 | 0.73691 |
| Palmitoyl-protein thioesterase 1 | O88531 | Ppt1 | 34 kDa | 10 C | 1.136 | 0.73691 |
| Threonine synthase-like 2 | Q80W22 | Thnsl2 | 54 kDa | 14 C | 1.133 | 0.72943 |
| Serine hydroxymethyltransferase | P50431 | Shmt1 | 53 kDa | 10 C | 1.132 | 0.39778 |
| Elongation factor 1-alpha 1 | P10126 | Eef1a1 | 50 kDa | 6 C | 1.132 | 0.47277 |
| Electron transfer flavoprotein subunit alpha | Q99LC5 | Etfa | 35 kDa | 6 C | 1.132 | 0.37757 |
| Transgelin-2 | Q9WVA4 | Tagln2 | 22 kDa | 3 C | 1.131 | 0.20107 |
| 28S ribosomal protein S22 | Q9CXW2 | Mrps22 | 41 kDa | 2 C | 1.129 | 0.50538 |
| Dihydrolipoyl dehydrogenase | O08749 | Dld | 54 kDa | 9 C | 1.126 | 0.53151 |
| Kynurenine--oxoglutarate transaminase 1 | Q8BTY1 | Kyat1 | 48 kDa | 7 C | 1.126 | 0.62861 |
| Stress-induced-phosphoprotein 1 | Q60864 | Stip1 | 63 kDa | 11 C | 1.125 | 0.69374 |
| ADP-ribosylation factor 3 | P61205 | Arf3 | 21 kDa | 1 C | 1.124 | 0.75409 |
| Far upstream element-binding protein 2 | Q3U0V1 | Khsrp | 77 kDa | 8 C | 1.120 | 0.4758 |
| 4-hydroxy-2-oxoglutarate aldolase | Q9DCU9 | Hoga1 | 35 kDa | 6 C | 1.120 | 0.55479 |
| Isoform 2 of Nucleoporin SEH1 | Q8R2U0-2 | Seh1l | 39 kDa | 10 C | 1.120 | 0.64441 |
| Complement factor I | Q61129 | Cfi | 67 kDa | 40 C | 1.120 | 0.68453 |
| Retinol-binding protein 4 | Q00724 | Rbp4 | 23 kDa | 6 C | 1.120 | 0.68453 |
| Heterogeneous nuclear ribonucleoprotein A0 | Q9CX86 | Hnrnpa0 | 31 kDa | 3 C | 1.120 | 0.68453 |
| Branched-chain-amino-acid aminotransferase | O35855 | Bcat2 | 44 kDa | 10 C | 1.120 | 0.71666 |
| Proteasome subunit beta type-7 | P70195 | Psmb7 | 30 kDa | 6 C | 1.120 | 0.78724 |
| Cofilin-1 | P18760 | Cfl1 | 19 kDa | 4 C | 1.118 | 0.60141 |
| Actin-related protein 2/3 complex subunit 5-like protein | Q9D898 | Arpc5l | 17 kDa | 1 C | 1.114 | 0.83985 |
| Isoform 2 of Pleckstrin homology-like domain family B member 2 | Q8K1N2-2 | Phldb2 | 147 kDa | 15 C | 1.113 | 0.89744 |
| DnaJ homolog subfamily C member 7 | Q9QYI3 | Dnajc7 | 56 kDa | 14 C | 1.113 | 0.89744 |
| Proteasome subunit beta type-2 | Q9R1P3 | Psmb2 | 23 kDa | 3 C | 1.111 | 0.62628 |
| Dimethylglycine dehydrogenase | Q9DBT9 | Dmgdh | 97 kDa | 4 C | 1.111 | 0.63181 |
| Cytochrome b-c1 complex subunit 2 | Q9DB77 | Uqcrc2 | 48 kDa | 1 C | 1.111 | 0.75683 |
| Mitochondrial peptide methionine sulfoxide reductase | Q9D6Y7 | Msra | 26 kDa | 4 C | 1.110 | 0.13707 |
| Isoform 2 of Galectin-9 | O08573-2 | Lgals9 | 37 kDa | 7 C | 1.110 | 0.8122 |
| Rho GDP-dissociation inhibitor 2 | Q61599 | Arhgdib | 23 kDa | 1 C | 1.110 | 0.90231 |
| Isoform 2 of Sorcin | Q6P069-2 | Sri | 20 kDa | 5 C | 1.110 | 0.90231 |
| Serotransferrin | Q921I1 | Tf | 77 kDa | 38 C | 1.109 | 0.13441 |
| Deoxynucleoside triphosphate triphosphohydrolase SAMHD1 | Q60710 | Samhd1 | 73 kDa | 17 C | 1.109 | 0.61042 |
| Endoribonuclease LACTB2 | Q99KR3 | Lactb2 | 33 kDa | 5 C | 1.109 | 0.24087 |
| Heterogeneous nuclear ribonucleoprotein F | Q9Z2X1 | Hnrnpf | 46 kDa | 6 C | 1.108 | 0.456 |
| Heat shock 70 kDa protein 4L | P48722 | Hspa4l | 94 kDa | 15 C | 1.108 | 0.73976 |
| 2-hydroxyacyl-CoA lyase 1 | Q9QXE0 | Hacl1 | 64 kDa | 17 C | 1.107 | 0.71092 |
| Protein phosphatase 1H | Q3UYC0 | Ppm1h | 56 kDa | 9 C | 1.105 | 0.78556 |
| Peroxisomal multifunctional enzyme type 2 | P51660 | Hsd17b4 | 79 kDa | 9 C | 1.104 | 0.35461 |
| Nucleoside diphosphate kinase 3 | Q9WV85 | Nme3 | 19 kDa | 3 C | 1.100 | 0.65169 |
| Growth factor receptor-bound protein 2 | Q60631 | Grb2 | 25 kDa | 2 C | 1.100 | 0.79797 |

| | | | | | | |
|---|---|---|---|---|---|---|
| Proteasome subunit alpha type-6 | Q9QUM9 | Psma6 | 27 kDa | 8 C | 1.100 | 0.3339 |
| Fumarylacetoacetate hydrolase domain-containing protein 2A | Q3TC72 | Fahd2 | 35 kDa | 6 C | 1.100 | 0.63824 |
| Na(+)/H(+) exchange regulatory cofactor NHE-RF3 | Q9JIL4 | Pdzk1 | 56 kDa | 7 C | 1.098 | 0.49354 |
| Peptidyl-prolyl cis-trans isomerase FKBP4 | P30416 | Fkbp4 | 52 kDa | 7 C | 1.097 | 0.71713 |
| Choline dehydrogenase | Q8BJ64 | Chdh | 66 kDa | 12 C | 1.095 | 0.56303 |
| Chloride intracellular channel protein 1 | Q9Z1Q5 | Clic1 | 27 kDa | 6 C | 1.095 | 0.61762 |
| Isoform 3 of Drebrin-like protein | Q62418-3 | Dbnl | 48 kDa | 5 C | 1.095 | 0.61762 |
| Glutathione S-transferase Mu 1 | P10649 | Gstm1 | 26 kDa | 2 C | 1.094 | 0.3764 |
| Isoform 2 of Cytosol aminopeptidase | Q9CPY7-2 | Lap3 | 53 kDa | 7 C | 1.092 | 0.07949 |
| High mobility group protein B1 | P63158 | Hmgb1 | 25 kDa | 3 C | 1.091 | 0.79042 |
| Pyridoxal phosphate homeostasis protein | Q9Z2Y8 | Prosc | 30 kDa | 3 C | 1.091 | 0.80476 |
| Isoform 3 of NAD kinase 2 | Q8C5H8-3 | Nadk2 | 48 kDa | 9 C | 1.088 | 0.70175 |
| Apoptosis-inducing factor 1 | Q9Z0X1 | Aifm1 | 67 kDa | 4 C | 1.088 | 0.20741 |
| Serine-threonine kinase receptor-associated protein | Q9Z1Z2 | Strap | 38 kDa | 6 C | 1.086 | 0.50235 |
| teoclast-stimulating factor 1 | Q62422 | Ostf1 | 24 kDa | 4 C | 1.086 | 0.56303 |
| Transthyretin | P07309 | Ttr | 16 kDa | 2 C | 1.086 | 0.79797 |
| Peroxisomal trans-2-enoyl-CoA reductase | Q99MZ7 | Pecr | 32 kDa | 5 C | 1.086 | 0.83459 |
| Sarcosine dehydrogenase | Q99LB7 | Sardh | 102 kDa | 18 C | 1.086 | 0.86616 |
| Eukaryotic translation initiation factor 2 subunit 3 | Q9Z0N1 | Eif2s3x | 51 kDa | 10 C | 1.084 | 0.88833 |
| Kinesin-like protein KIF21A | Q9QXL2 | Kif21a | 187 kDa | 25 C | 1.084 | 0.88833 |
| 4-aminobutyrate aminotransferase | P61922 | Abat | 56 kDa | 12 C | 1.083 | 0.57913 |
| Hemoglobin subunit beta-1 | P02088 | Hbb-b1 | 16 kDa | 2 C | 1.082 | 0.65247 |
| Alpha-methylacyl-CoA racemase | O09174 | Amacr | 42 kDa | 6 C | 1.081 | 0.57083 |
| Sorbitol dehydrogenase | Q64442 | Sord | 38 kDa | 10 C | 1.080 | 0.47891 |
| Adapter molecule crk | Q64010 | Crk | 34 kDa | 1 C | 1.080 | 0.72943 |
| NADH dehydrogenase [ubiquinone] iron-sulfur protein 4 | Q9CXZ1 | Ndufs4 | 20 kDa | 1 C | 1.080 | 0.87094 |
| Rab GDP dissociation inhibitor beta | Q61598 | Gdi2 | 51 kDa | 9 C | 1.075 | 0.91088 |
| F-box only protein 50 | G3X9C2 | Nccrp1 | 30 kDa | 3 C | 1.075 | 0.6059 |
| Glutathione S-transferase kappa 1 | Q9DCM2 | Gstk1 | 26 kDa | 2 C | 1.075 | 0.93019 |
| Trehalase | Q9JLT2 | Treh | 65 kDa | 5 C | 1.075 | 0.93019 |
| Isoform Mt-VDAC1 of Voltage-dependent anion-selective channel protein 1 | Q60932-2 | Vdac1 | 31 kDa | 2 C | 1.074 | 0.84117 |
| Acetyl-CoA acetyltransferase | Q8QZT1 | Acat1 | 45 kDa | 6 C | 1.074 | 0.22422 |
| Succinate dehydrogenase [ubiquinone] iron-sulfur subunit | Q9CQA3 | Sdhb | 32 kDa | 14 C | 1.073 | 0.46011 |
| Selenium-binding protein 1 | P17563 | Selenbp1 | 53 kDa | 10 C | 1.072 | 0.58993 |
| Proteasome subunit beta type-3 | Q9R1P1 | Psmb3 | 23 kDa | 5 C | 1.067 | 0.56303 |
| ELAV-like protein 1 | P70372 | Elavl1 | 36 kDa | 3 C | 1.067 | 0.68453 |
| Exosome complex component RRP45 | Q9JHI7 | Exosc9 | 49 kDa | 11 C | 1.067 | 0.68453 |
| Fibrinogen alpha chain | E9PV24 | Fga | 87 kDa | 13 C | 1.067 | 0.70009 |
| ES1 protein homolog | Q9D172 | D10Jhu81e | 28 kDa | 6 C | 1.067 | 0.71556 |
| NFU1 iron-sulfur cluster scaffold homolog | Q9QZ23 | Nfu1 | 29 kDa | 5 C | 1.067 | 0.7746 |
| Lactoylglutathione lyase | Q9CPU0 | Glo1 | 21 kDa | 3 C | 1.067 | 0.79042 |
| Ig lambda-2 chain C region | P01844 | Iglc2 | 11 kDa | 3 C | 1.067 | 0.79797 |
| Thioredoxin domain-containing protein 5 | Q91W90 | Txndc5 | 46 kDa | 12 C | 1.067 | 0.81951 |
| Succinate--CoA ligase [ADP/GDP-forming] subunit alpha | Q9WUM5 | Suclg1 | 36 kDa | 6 C | 1.067 | 0.89433 |
| Propionyl-CoA carboxylase beta chain | Q99MN9 | Pccb | 58 kDa | 11 C | 1.064 | 0.60417 |
| Hydroxyacyl-coenzyme A dehydrogenase | Q61425 | Hadh | 34 kDa | 5 C | 1.062 | 0.74517 |
| Hemopexin | Q91X72 | Hpx | 51 kDa | 13 C | 1.060 | 0.64254 |
| Homogentisate 1 | O09173 | Hgd | 50 kDa | 14 C | 1.060 | 0.80536 |
| Succinate--CoA ligase [ADP-forming] subunit beta | Q9Z2I9 | Sucla2 | 50 kDa | 6 C | 1.058 | 0.94927 |
| Selenoprotein F | Q9ERR7 | Selenof | 18 kDa | 7 C | 1.055 | 0.8623 |
| DnaJ homolog subfamily B member 11 | Q99KV1 | Dnajb11 | 41 kDa | 5 C | 1.055 | 0.89706 |
| Heterogeneous nuclear ribonucleoprotein H | O35737 | Hnrnph1 | 49 kDa | 1 C | 1.053 | 0.64702 |
| Hsp90 co-chaperone Cdc37 | Q61081 | Cdc37 | 45 kDa | 9 C | 1.053 | 0.86808 |
| Sialic acid synthase | Q99J77 | Nans | 40 kDa | 8 C | 1.050 | 0.94546 |
| Platelet glycoprotein 4 | Q08857 | Cd36 | 53 kDa | 10 C | 1.050 | 0.94546 |
| Peroxiredoxin-6 | O08709 | Prdx6 | 25 kDa | 2 C | 1.049 | 0.68453 |
| Septin-2 | P42208 | 43345 | 42 kDa | 8 C | 1.048 | 0.8373 |
| Heterogeneous nuclear ribonucleoprotein A/B | Q99020 | Hnrnpab | 31 kDa | 2 C | 1.044 | 0.67719 |
| Profilin-1 | P62962 | Pfn1 | 15 kDa | 3 C | 1.043 | 0.8348 |
| Receptor of activated protein C kinase 1 | P68040 | Rack1 | 35 kDa | 8 C | 1.043 | 0.88697 |

| | | | | | | |
|---|---|---|---|---|---|---|
| Ketohexokinase | P97328 | Khk | 33 kDa | 10 C | 1.040 | 0.71666 |
| Cytosolic purine 5'-nucleotidase | Q3V1L4 | Nt5c2 | 65 kDa | 8 C | 1.040 | 0.88979 |
| Probable ATP-dependent RNA helicase DDX17 | Q501J6 | Ddx17 | 72 kDa | 11 C | 1.040 | 0.93335 |
| Translationally-controlled tumor protein | P63028 | Tpt1 | 19 kDa | 2 C | 1.040 | 0.93656 |
| Glutamate--cysteine ligase catalytic subunit | P97494 | Gclc | 73 kDa | 14 C | 1.037 | 0.87622 |
| Xaa-Pro aminopeptidase 1 | Q6P1B1 | Xpnpep1 | 70 kDa | 12 C | 1.035 | 0.84452 |
| Hydroxyacylglutathione hydrolase | Q99KB8 | Hagh | 34 kDa | 8 C | 1.034 | 0.84443 |
| Annexin A5 | P48036 | Anxa5 | 36 kDa | 1 C | 1.033 | 0.92469 |
| Carbonic anhydrase 2 | P00920 | Ca2 | 29 kDa | 2 C | 1.032 | 0.77236 |
| Cytochrome c1 | Q9D0M3 | Cyc1 | 35 kDa | 14 C | 1.029 | 0.94475 |
| Protein DJ-1 | Q99LX0 | Park7 | 20 kDa | 4 C | 1.029 | 0.85998 |
| Bleomycin hydrolase | Q8R016 | Blmh | 53 kDa | 4 C | 1.029 | 0.87346 |
| Protein DDI1 homolog 2 | A2ADY9 | Ddi2 | 45 kDa | 8 C | 1.029 | 0.87862 |
| Oligoribonuclease | Q9D8S4 | Rexo2 | 27 kDa | 4 C | 1.029 | 0.91958 |
| 39S ribosomal protein L18 | Q9CQL5 | Mrpl18 | 21 kDa | 5 C | 1.029 | 0.91958 |
| Growth arrest-specific protein 2 | P11862 | Gas2 | 35 kDa | 11 C | 1.029 | 0.92618 |
| Actin-related protein 2/3 complex subunit 1B | Q9WV32 | Arpc1b | 41 kDa | 14 C | 1.029 | 0.93563 |
| Acyl-coenzyme A synthetase ACSM5 | Q8BGA8 | Acsm5 | 64 kDa | 15 C | 1.029 | 0.9509 |
| NADH dehydrogenase [ubiquinone] 1 beta subcomplex subunit 9 | Q9CQJ8 | Ndufb9 | 22 kDa | 5 C | 1.029 | 0.9509 |
| Fibrinogen beta chain | Q8K0E8 | Fgb | 55 kDa | 12 C | 1.029 | 0.82684 |
| Isoform 2 of Meprin A subunit beta | Q61847-2 | Mep1b | 80 kDa | 11 C | 1.026 | 0.80276 |
| Sushi domain-containing protein 2 | Q9DBX3 | Susd2 | 91 kDa | 28 C | 1.025 | 0.9619 |
| Cytochrome b5 type B | Q9CQX2 | Cyb5b | 16 kDa | 1 C | 1.025 | 0.96456 |
| Inositol-3-phosphate synthase 1 | Q9JHU9 | Isyna1 | 61 kDa | 11 C | 1.025 | 0.9095 |
| Crk-like protein | P47941 | Crkl | 34 kDa | 2 C | 1.025 | 0.96556 |
| 60S ribosomal protein L22 | P67984 | Rpl22 | 15 kDa | 1 C | 1.020 | 0.95486 |
| Creatine kinase U-type | P30275 | Ckmt1 | 47 kDa | 7 C | 1.018 | 0.85285 |
| Immunoglobulin J chain | P01592 | Jchain | 18 kDa | 8 C | 1.018 | 0.94479 |
| Stromal cell-derived factor 2 | Q9DCT5 | Sdf2 | 23 kDa | 4 C | 1.018 | 0.96151 |
| Aminoacylase-1 | Q99JW2 | Acy1 | 46 kDa | 4 C | 1.016 | 0.81532 |
| Ezrin | P26040 | Ezr | 69 kDa | 2 C | 1.016 | 0.9431 |
| Flavin reductase (NADPH) | Q923D2 | Blvrb | 22 kDa | 2 C | 1.015 | 0.87862 |
| Elongation factor 1-gamma | Q9D8N0 | Eef1g | 50 kDa | 6 C | 1.015 | 0.98043 |
| Platelet-activating factor acetylhydrolase IB subunit alpha | P63005 | Pafah1b1 | 47 kDa | 10 C | 1.015 | 0.98043 |
| Thioredoxin-dependent peroxide reductase | P20108 | Prdx3 | 28 kDa | 4 C | 1.014 | 0.86758 |
| tRNA-splicing ligase RtcB homolog | Q99LF4 | Rtcb | 55 kDa | 9 C | 1.014 | 0.97911 |
| FAD-linked sulfhydryl oxidase ALR | P56213 | Gfer | 23 kDa | 8 C | 1.013 | 0.94479 |
| Chloride intracellular channel protein 4 | Q9QYB1 | Clic4 | 29 kDa | 4 C | 1.013 | 0.95535 |
| Peroxiredoxin-4 | O08807 | Prdx4 | 31 kDa | 4 C | 1.011 | 0.9431 |
| S-adenosylmethionine synthase isoform type-2 | Q3THS6 | Mat2a | 44 kDa | 6 C | 1.011 | 0.96377 |
| Disabled homolog 2 | P98078 | Dab2 | 82 kDa | 3 C | 1.011 | 0.97586 |
| Methylmalonate-semialdehyde dehydrogenase [acylating] | Q9EQ20 | Aldh6a1 | 58 kDa | 8 C | 1.010 | 0.92703 |
| COP9 signalosome complex subunit 5 | O35864 | Cops5 | 38 kDa | 4 C | 1.010 | 0.98217 |
| N-acyl-aromatic-L-amino acid amidohydrolase (carboxylate-forming) | Q91XE4 | Acy3 | 35 kDa | 8 C | 1.010 | 0.92893 |
| Major urinary protein 6 | P02762 | Mup6 | 21 kDa | 5 C | 1.009 | 0.98637 |
| Protein ABHD14B | Q8VCR7 | Abhd14b | 22 kDa | 2 C | 1.009 | 0.95535 |
| Heat shock protein HSP 90-beta | P11499 | Hsp90ab1 | 83 kDa | 6 C | 1.009 | 0.98588 |
| Isoform M1 of Pyruvate kinase PKM | P52480-2 | Pkm | 58 kDa | 9 C | 1.007 | 0.97699 |
| Hypoxanthine-guanine phosphoribosyltransferase | P00493 | Hprt1 | 25 kDa | 4 C | 1.007 | 0.97925 |
| Exosome complex component MTR3 | Q8BTW3 | Exosc6 | 28 kDa | 6 C | 1.007 | 0.98341 |
| Malate dehydrogenase | P08249 | Mdh2 | 36 kDa | 8 C | 1.006 | 0.97738 |
| Glyoxylate reductase/hydroxypyruvate reductase | Q91Z53 | Grhpr | 35 kDa | 7 C | 1.004 | 0.98298 |
| BAG family molecular chaperone regulator 1 | Q60739 | Bag1 | 40 kDa | 5 C | 1.000 | 1 |
| Copine-3 | Q8BT60 | Cpne3 | 60 kDa | 13 C | 1.000 | 1 |
| Isoform 2 of Heterogeneous nuclear ribonucleoprotein U | Q8VEK3-2 | Hnrnpu | 87 kDa | 14 C | 1.000 | 1 |
| 40S ribosomal protein S20 | P60867 | Rps20 | 13 kDa | 2 C | 1.000 | 1 |
| Protein SGT1 homolog | Q9CX34 | Sugt1 | 38 kDa | 5 C | 1.000 | 1 |
| Serine/threonine-protein phosphatase 2A 55 kDa regulatory subunit B alpha isoform | Q6P1F6 | Ppp2r2a | 52 kDa | 9 C | 1.000 | 1 |
| Peroxisomal acyl-coenzyme A oxidase 1 | Q9R0H0 | Acox1 | 75 kDa | 7 C | 0.996 | 0.9706 |
| Gamma-glutamyltranspeptidase 1 | Q60928 | Ggt1 | 62 kDa | 6 C | 0.996 | 0.98827 |

| | | | | | | |
|---|---|---|---|---|---|---|
| Eukaryotic translation initiation factor 3 subunit G | Q9Z1D1 | Eif3g | 36 kDa | 5 C | 0.995 | 0.99201 |
| Eukaryotic translation initiation factor 4H | Q9WUK2 | Eif4h | 27 kDa | 1 C | 0.994 | 0.99424 |
| Xaa-Pro dipeptidase | Q11136 | Pepd | 55 kDa | 17 C | 0.993 | 0.964 |
| Alcohol dehydrogenase [NADP(+)] | Q9JII6 | Akr1a1 | 37 kDa | 4 C | 0.993 | 0.98447 |
| Rho GDP-dissociation inhibitor 1 | Q99PT1 | Arhgdia | 23 kDa | 1 C | 0.992 | 0.96151 |
| Cold shock domain-containing protein E1 | Q91W50 | Csde1 | 89 kDa | 15 C | 0.992 | 0.97699 |
| Proteasome subunit alpha type-2 | P49722 | Psma2 | 26 kDa | 2 C | 0.990 | 0.91958 |
| Plasminogen | P20918 | Plg | 91 kDa | 48 C | 0.988 | 0.97379 |
| Glutathione synthetase | P51855 | Gss | 52 kDa | 4 C | 0.987 | 0.83459 |
| Sulfite oxidase | Q8R086 | Suox | 61 kDa | 9 C | 0.982 | 0.90972 |
| Hemoglobin subunit alpha | P01942 | Hba | 15 kDa | 1 C | 0.982 | 0.9509 |
| NADP-dependent malic enzyme | P06801 | Me1 | 64 kDa | 11 C | 0.980 | 0.9245 |
| Endoplasmic reticulum resident protein 29 | P57759 | Erp29 | 29 kDa | 1 C | 0.978 | 0.65169 |
| Ribosyldihydronicotinamide dehydrogenase [quinone] | Q9JI75 | Nqo2 | 26 kDa | 4 C | 0.978 | 0.93563 |
| Carboxylesterase 1F | Q91WU0 | Ces1f | 62 kDa | 7 C | 0.976 | 0.85683 |
| 2-amino-3-carboxymuconate-6-semialdehyde decarboxylase | Q8R519 | Acmsd | 38 kDa | 7 C | 0.974 | 0.92197 |
| Myosin light chain kinase | Q6PDN3 | Mylk | 213 kDa | 46 C | 0.971 | 0.85285 |
| Hydroxymethylglutaryl-CoA synthase | Q8JZK9 | Hmgcs1 | 58 kDa | 11 C | 0.971 | 0.90279 |
| Glutathione S-transferase A1 | P13745 | Gsta1 | 26 kDa | 2 C | 0.968 | 0.93139 |
| Fructose-1 | Q9QXD6 | Fbp1 | 37 kDa | 7 C | 0.968 | 0.70756 |
| Glutamate carboxypeptidase 2 | O35409 | Folh1 | 85 kDa | 3 C | 0.967 | 0.97258 |
| Isoform Rpn10B of 26S proteasome non-ATPase regulatory subunit 4 | O35226-2 | Psmd4 | 41 kDa | 4 C | 0.967 | 0.95265 |
| Protein disulfide-isomerase A6 | Q922R8 | Pdia6 | 48 kDa | 7 C | 0.967 | 0.9002 |
| Peroxisomal carnitine O-octanoyltransferase | Q9DC50 | Crot | 70 kDa | 16 C | 0.966 | 0.84211 |
| Fucose mutarotase | Q8R2K1 | Fuom | 17 kDa | 3 C | 0.960 | 0.7746 |
| Adenosylhomocysteinase | P50247 | Ahcy | 48 kDa | 9 C | 0.960 | 0.74553 |
| Protein NipSnap homolog 3B | Q9CQE1 | Nipsnap3b | 28 kDa | 2 C | 0.960 | 0.87862 |
| Thioredoxin domain-containing protein 12 | Q9CQU0 | Txndc12 | 19 kDa | 3 C | 0.960 | 0.87862 |
| Arsenite methyltransferase | Q91WU5 | As3mt | 42 kDa | 12 C | 0.960 | 0.90519 |
| CD5 antigen-like | Q9QWK4 | Cd5l | 39 kDa | 26 C | 0.957 | 0.56303 |
| Protein SEC13 homolog | Q9D1M0 | Sec13 | 36 kDa | 9 C | 0.957 | 0.92599 |
| Bifunctional epoxide hydrolase 2 | P34914 | Ephx2 | 63 kDa | 10 C | 0.955 | 0.81547 |
| Cytosolic non-specific dipeptidase | Q9D1A2 | Cndp2 | 53 kDa | 8 C | 0.954 | 0.50457 |
| Acyl-coenzyme A synthetase ACSM2 | Q8K0L3 | Acsm2 | 64 kDa | 9 C | 0.954 | 0.53311 |
| Dihydrolipoyllysine-residue acetyltransferase component of pyruvate dehydrogenase com | Q8BMF4 | Dlat | 68 kDa | 10 C | 0.953 | 0.87217 |
| Aldehyde dehydrogenase | O35945 | Aldh1a7 | 55 kDa | 8 C | 0.952 | 0.66466 |
| Actin-related protein 2/3 complex subunit 4 | P59999 | Arpc4 | 20 kDa | 4 C | 0.952 | 0.77236 |
| Citrate synthase | Q9CZU6 | Cs | 52 kDa | 5 C | 0.952 | 0.93598 |
| Cadherin-1 | P09803 | Cdh1 | 98 kDa | 8 C | 0.950 | 0.77107 |
| Complement component C8 gamma chain | Q8VCG4 | C8g | 23 kDa | 3 C | 0.948 | 0.70156 |
| 3-hydroxyanthranilate 3 | Q78JT3 | Haao | 33 kDa | 3 C | 0.948 | 0.75192 |
| Isoform 3 of Cellular nucleic acid-binding protein | P53996-3 | Cnbp | 19 kDa | 22 C | 0.947 | 0.87801 |
| Adaptin ear-binding coat-associated protein 2 | Q9D1J1 | Necap2 | 29 kDa | 2 C | 0.947 | 0.87801 |
| Serine/threonine-protein phosphatase 2A catalytic subunit alpha isoform | P63330 | Ppp2ca | 36 kDa | 10 C | 0.945 | 0.74063 |
| Elongation factor 1-beta | O70251 | Eef1b | 25 kDa | 3 C | 0.945 | 0.7746 |
| Serine--tRNA ligase | P26638 | Sars | 58 kDa | 9 C | 0.945 | 0.87406 |
| Superoxide dismutase [Cu-Zn] | P08228 | Sod1 | 16 kDa | 3 C | 0.945 | 0.87862 |
| Phosphoglycerate mutase 1 | Q9DBJ1 | Pgam1 | 29 kDa | 2 C | 0.944 | 0.75731 |
| 40S ribosomal protein S12 | P63323 | Rps12 | 15 kDa | 7 C | 0.943 | 0.51133 |
| Non-specific lipid-transfer protein | P32020 | Scp2 | 59 kDa | 11 C | 0.943 | 0.79935 |
| V-type proton ATPase catalytic subunit A | P50516 | Atp6v1a | 68 kDa | 6 C | 0.942 | 0.82849 |
| Pyruvate dehydrogenase E1 component subunit alpha | P35486 | Pdha1 | 43 kDa | 12 C | 0.941 | 0.74864 |
| Proteasome subunit alpha type-1 | Q9R1P4 | Psma1 | 30 kDa | 5 C | 0.941 | 0.79042 |
| Serine protease inhibitor A3K | P07759 | Serpina3k | 47 kDa | 4 C | 0.940 | 0.6059 |
| Peroxisomal acyl-coenzyme A oxidase 3 | Q9EPL9 | Acox3 | 78 kDa | 14 C | 0.939 | 0.77923 |
| Omega-amidase NIT2 | Q9JHW2 | Nit2 | 31 kDa | 2 C | 0.938 | 0.61762 |
| Ribosome-recycling factor | Q9D6S7 | Mrrf | 29 kDa | 2 C | 0.933 | 0.84337 |
| Hepatoma-derived growth factor | P51859 | Hdgf | 26 kDa | 2 C | 0.933 | 0.85285 |
| Actin-related protein 2/3 complex subunit 5 | Q9CPW4 | Arpc5 | 16 kDa | 1 C | 0.933 | 0.87862 |
| NADH dehydrogenase [ubiquinone] iron-sulfur protein 7 | Q9DC70 | Ndufs7 | 25 kDa | 5 C | 0.931 | 0.86645 |

| | | | | | | |
|---|---|---|---|---|---|---|
| Isoform 2 of Septin-10 | Q8C650-2 | 43353 | 50 kDa | 14 C | 0.931 | 0.89762 |
| Trimethyllysine dioxygenase | Q91ZE0 | Tmlhe | 50 kDa | 11 C | 0.931 | 0.78047 |
| Phytanoyl-CoA dioxygenase | O35386 | Phyh | 39 kDa | 8 C | 0.926 | 0.79582 |
| Integrin beta-1 | P09055 | Itgb1 | 88 kDa | 57 C | 0.924 | 0.89445 |
| Delta(3 | O35459 | Ech1 | 36 kDa | 6 C | 0.921 | 0.85026 |
| Heterogeneous nuclear ribonucleoprotein L | Q8R081 | Hnrnpl | 64 kDa | 11 C | 0.921 | 0.64254 |
| Mitochondrial fission 1 protein | Q9CQ92 | Fis1 | 17 kDa | 1 C | 0.914 | 0.67463 |
| Endoplasmin | P08113 | Hsp90b1 | 92 kDa | 5 C | 0.914 | 0.83978 |
| Catalase | P24270 | Cat | 60 kDa | 5 C | 0.914 | 0.4309 |
| Ornithine aminotransferase | P29758 | Oat | 48 kDa | 7 C | 0.914 | 0.66626 |
| Adenylate kinase 4 | Q9WUR9 | Ak4 | 25 kDa | 2 C | 0.913 | 0.84678 |
| Hsc70-interacting protein | Q99L47 | St13 | 42 kDa | 3 C | 0.910 | 0.48246 |
| Succinyl-CoA:3-ketoacid coenzyme A transferase 1 | Q9D0K2 | Oxct1 | 56 kDa | 7 c | 0.910 | 0.54741 |
| Cytochrome c oxidase assembly factor 7 | Q921H9 | Coa7 | 26 kDa | 13 C | 0.910 | 0.78788 |
| Kynurenine/alpha-aminoadipate aminotransferase | Q9WVM8 | Aadat | 48 kDa | 5 C | 0.909 | 0.74146 |
| Cytochrome b-c1 complex subunit Rieske | Q9CR68 | Uqcrfs1 | 29 kDa | 5 C | 0.905 | 0.56303 |
| 60S ribosomal protein L12 | P35979 | Rpl12 | 18 kDa | 3 C | 0.904 | 0.50538 |
| Proteasome subunit alpha type-4 | Q9R1P0 | Psma4 | 29 kDa | 5 C | 0.900 | 0.19301 |
| Sodium/potassium-transporting ATPase subunit beta-1 | P14094 | Atp1b1 | 35 kDa | 7 C | 0.900 | 0.68274 |
| Iron-sulfur cluster assembly enzyme ISCU | Q9D7P6 | Iscu | 18 kDa | 4 C | 0.900 | 0.72943 |
| Glycerol-3-phosphate phosphatase | Q8CHP8 | Pgp | 35 kDa | 8 C | 0.900 | 0.83179 |
| Medium-chain specific acyl-CoA dehydrogenase | P45952 | Acadm | 46 kDa | 8 C | 0.898 | 0.59117 |
| Glutaredoxin-related protein 5 | Q80Y14 | Glrx5 | 16 kDa | 2 C | 0.898 | 0.7862 |
| Nucleoside diphosphate-linked moiety X motif 19 | P11930 | Nudt19 | 40 kDa | 9 C | 0.898 | 0.90108 |
| Long-chain specific acyl-CoA dehydrogenase | P51174 | Acadl | 48 kDa | 7 C | 0.895 | 0.31248 |
| Glycine amidinotransferase | Q9D964 | Gatm | 48 kDa | 9 C | 0.894 | 0.533 |
| D-amino-acid oxidase | P18894 | Dao | 39 kDa | 6 C | 0.889 | 0.80833 |
| Isoform 2 of Vacuolar protein sorting-associated protein 26A | P40336-2 | Vps26a | 42 kDa | 2 C | 0.889 | 0.66466 |
| Isovaleryl-CoA dehydrogenase | Q9JHI5 | Ivd | 46 kDa | 7 C | 0.886 | 0.67135 |
| Alpha-1-antitrypsin 1-2 | P22599 | Serpina1b | 46 kDa | 3 C | 0.886 | 0.69613 |
| Glutathione S-transferase Mu 2 | P15626 | Gstm2 | 26 kDa | 3 C | 0.883 | 0.32316 |
| 3-hydroxybutyrate dehydrogenase type 2 | Q8JZV9 | Bdh2 | 27 kDa | 6 C | 0.882 | 0.34369 |
| Phosphatidylethanolamine-binding protein 1 | P70296 | Pebp1 | 21 kDa | 3 C | 0.878 | 0.14072 |
| Alpha-enolase | P17182 | Eno1 | 47 kDa | 6 C | 0.877 | 0.40862 |
| Golgi resident protein GCP60 | Q8BMP6 | Acbd3 | 60 kDa | 4 C | 0.877 | 0.89387 |
| Haloacid dehalogenase-like hydrolase domain-containing protein 2 | Q3UGR5 | Hdhd2 | 29 kDa | 3 C | 0.873 | 0.88989 |
| Dolichyl-diphosphooligosaccharide--protein glycosyltransferase subunit 1 | Q91YQ5 | Rpn1 | 69 kDa | 3 C | 0.873 | 0.88989 |
| Protein kinase C and casein kinase substrate in neurons protein 2 | Q9WVE8 | Pacsin2 | 56 kDa | 6 C | 0.873 | 0.88989 |
| Glutamine amidotransferase-like class 1 domain-containing protein 1 | Q8BFQ8 | Gatd1 | 23 kDa | 7 C | 0.873 | 0.88989 |
| Peptidyl-prolyl cis-trans isomerase FKBP2 | P45878 | Fkbp2 | 15 kDa | 3 C | 0.873 | 0.88989 |
| Prostaglandin E synthase 3 | Q9R0Q7 | Ptges3 | 19 kDa | 5 C | 0.867 | 0.61902 |
| Delta-1-pyrroline-5-carboxylate dehydrogenase | Q8CHT0 | Aldh4a1 | 62 kDa | 8 C | 0.867 | 0.82847 |
| Phosphoglycerate mutase 2 | O70250 | Pgam2 | 29 kDa | 3 C | 0.865 | 0.71749 |
| Kynurenine--oxoglutarate transaminase 3 | Q71RI9 | Kyat3 | 51 kDa | 10 C | 0.863 | 0.49927 |
| Dihydrolipoyllysine-residue succinyltransferase component of 2-oxoglutarate dehydrogen | Q9D2G2 | Dlst | 49 kDa | 6 C | 0.859 | 0.26467 |
| Selenocysteine lyase | Q9JLI6 | Scly | 47 kDa | 8 C | 0.859 | 0.26467 |
| Glutathione peroxidase 1 | P11352 | Gpx1 | 22 kDa | 4 C | 0.858 | 0.31854 |
| Polypyrimidine tract-binding protein 1 | P17225 | Ptbp1 | 56 kDa | 3 C | 0.857 | 0.29235 |
| Enoyl-CoA delta isomerase 1 | P42125 | Eci1 | 32 kDa | 5 C | 0.857 | 0.40708 |
| Ig heavy chain V-III region J606 | P01801 | | 13 kDa | 2 C | 0.853 | 0.50538 |
| Argininosuccinate synthase | P16460 | Ass1 | 47 kDa | 5 C | 0.846 | 0.38966 |
| Tubulin-folding cofactor B | Q9D1E6 | Tbcb | 27 kDa | 5 C | 0.844 | 0.50972 |
| ATP synthase subunit O | Q9DB20 | Atp5o | 23 kDa | 1 C | 0.844 | 0.68219 |
| Isoform 2 of Enoyl-CoA delta isomerase 2 | Q9WUR2-2 | Eci2 | 40 kDa | 7 C | 0.839 | 0.82609 |
| Glycine dehydrogenase (decarboxylating) | Q91W43 | Gldc | 113 kDa | 24 C | 0.837 | 0.5907 |
| Isoform 2 of MICOS complex subunit Mic60 | Q8CAQ8-2 | Immt | 83 kDa | 7 C | 0.836 | 0.66626 |
| Acyl-CoA dehydrogenase family member 9 | Q8JZN5 | Acad9 | 69 kDa | 9 C | 0.835 | 0.87862 |
| Calcium-binding mitochondrial carrier protein Aralar1 | Q8BH59 | Slc25a12 | 75 kDa | 7 C | 0.835 | 0.87862 |
| Solute carrier family 22 member 6 | Q8VC69 | Slc22a6 | 60 kDa | 13 C | 0.835 | 0.87862 |
| ATP synthase subunit alpha | Q03265 | Atp5a1 | 60 kDa | 2 C | 0.829 | 0.57611 |

| | | | | | | |
|---|---|---|---|---|---|---|
| Four and a half LIM domains protein 1 | P97447 | Fhl1 | 32 kDa | 34 C | 0.827 | 0.20999 |
| Nodal modulator 1 | Q6GQT9 | Nomo1 | 133 kDa | 18 C | 0.820 | 0.73528 |
| Ras-related protein Rab-5C | P35278 | Rab5c | 23 kDa | 4 C | 0.820 | 0.73528 |
| Guanine nucleotide-binding protein G(I)/G(S)/G(T) subunit beta-1 | P62874 | Gnb1 | 37 kDa | 14 C | 0.819 | 0.81895 |
| Calcium-binding mitochondrial carrier protein Aralar2 | Q9QXX4 | Slc25a13 | 74 kDa | 7 C | 0.819 | 0.87862 |
| Zyxin | Q62523 | Zyx | 61 kDa | 23 C | 0.816 | 0.58831 |
| Phospholipase D3 | O35405 | Pld3 | 54 kDa | 8 C | 0.816 | 0.75448 |
| Aspartate aminotransferase | P05202 | Got2 | 47 kDa | 7 C | 0.816 | 0.22257 |
| Moesin | P26041 | Msn | 68 kDa | 2 C | 0.816 | 0.33998 |
| Isoform 3 of Protein scribble homolog | Q80U72-3 | Scrib | 180 kDa | 21 C | 0.816 | 0.719 |
| Ganglioside GM2 activator | Q60648 | Gm2a | 21 kDa | 8 C | 0.813 | 0.74505 |
| Ig kappa chain V-VI region NQ2-6.1 | P04945 | | 12 kDa | 2 C | 0.800 | 0.08922 |
| START domain-containing protein 10 | Q9JMD3 | Stard10 | 33 kDa | 7 C | 0.800 | 0.28741 |
| MICOS complex subunit Mic19 | Q9CRB9 | Chchd3 | 26 kDa | 4 C | 0.800 | 0.29235 |
| Cofilin-2 | P45591 | Cfl2 | 19 kDa | 2 C | 0.800 | 0.34334 |
| Hypoxia up-regulated protein 1 | Q9JKR6 | Hyou1 | 111 kDa | 4 C | 0.800 | 0.40708 |
| Exosome complex component RRP4 | Q8VBV3 | Exosc2 | 33 kDa | 4 C | 0.800 | 0.50538 |
| Short-chain specific acyl-CoA dehydrogenase | Q07417 | Acads | 45 kDa | 5 C | 0.800 | 0.50709 |
| Lambda-crystallin homolog | Q99KP3 | Cryl1 | 35 kDa | 7 C | 0.800 | 0.53892 |
| Glutamate dehydrogenase 1 | P26443 | Glud1 | 61 kDa | 6 C | 0.800 | 0.64157 |
| Golgi apparatus protein 1 | Q61543 | Glg1 | 134 kDa | 68 C | 0.800 | 0.64307 |
| Napsin-A | O09043 | Napsa | 46 kDa | 7 C | 0.800 | 0.66066 |
| Basigin | P18572 | Bsg | 42 kDa | 8 C | 0.800 | 0.66908 |
| N-fatty-acyl-amino acid synthase/hydrolase PM20D1 | Q8C165 | Pm20d1 | 56 kDa | 2 C | 0.800 | 0.68996 |
| 3-hydroxyacyl-CoA dehydrogenase type-2 | O08756 | Hsd17b10 | 27 kDa | 2 C | 0.800 | 0.70323 |
| Complement C4-B | P01029 | C4b | 193 kDa | 29 C | 0.800 | 0.72943 |
| NADH dehydrogenase [ubiquinone] 1 alpha subcomplex subunit 8 | Q9DCJ5 | Ndufa8 | 20 kDa | 8 C | 0.800 | 0.74723 |
| Properdin | P11680 | Cfp | 50 kDa | 44 C | 0.800 | 0.81985 |
| Probable D-lactate dehydrogenase | Q7TNG8 | Ldhd | 52 kDa | 12 C | 0.800 | 0.00231 |
| 3-oxoacyl-[acyl-carrier-protein] synthase | Q9D404 | Oxsm | 49 kDa | 11 C | 0.800 | 0.35582 |
| Cytoplasmic aconitate hydratase | P28271 | Aco1 | 98 kDa | 11 C | 0.800 | 0.462 |
| Cysteine desulfurase | Q9Z1J3 | Nfs1 | 51 kDa | 7 C | 0.800 | 0.46555 |
| Glutathione S-transferase A3 | P30115 | Gsta3 | 25 kDa | 1 C | 0.800 | 0.54968 |
| Electron transfer flavoprotein-ubiquinone oxidoreductase | Q921G7 | Etfdh | 68 kDa | 15 C | 0.800 | 0.69458 |
| Carbonic anhydrase 5A | P23589 | Ca5a | 34 kDa | 7 C | 0.800 | 0.74337 |
| D-dopachrome decarboxylase | O35215 | Ddt | 13 kDa | 2 C | 0.800 | 0.81985 |
| Alkaline phosphatase | P09242 | Alpl | 58 kDa | 6 C | 0.792 | 0.67565 |
| Eukaryotic translation initiation factor 2 subunit 1 | Q6ZWX6 | Eif2s1 | 36 kDa | 5 C | 0.789 | 0.55205 |
| Mitochondrial dicarboxylate carrier | Q9QZD8 | Slc25a10 | 32 kDa | 8 C | 0.785 | 0.82371 |
| Very long-chain specific acyl-CoA dehydrogenase | P50544 | Acadvl | 71 kDa | 7 C | 0.780 | 0.43295 |
| 3-ketoacyl-CoA thiolase B | Q8VCH0 | Acaa1b | 44 kDa | 9 C | 0.765 | 0.22422 |
| Isocitrate dehydrogenase [NADP] | P54071 | Idh2 | 51 kDa | 8 C | 0.763 | 0.46261 |
| Ras-related protein Rab-11B | P46638 | Rab11b | 24 kDa | 2 C | 0.758 | 0.54147 |
| Pyruvate dehydrogenase E1 component subunit beta | Q9D051 | Pdhb | 39 kDa | 6 C | 0.753 | 0.32853 |
| Ig kappa chain V-III region PC 2880/PC 1229 | P01654 | | 12 kDa | 2 C | 0.753 | 0.48493 |
| Citrate lyase subunit beta-like protein | Q8R4N0 | Clybl | 38 kDa | 6 C | 0.751 | 0.56954 |
| Alanine aminotransferase 1 | Q8QZR5 | Gpt | 55 kDa | 14 C | 0.750 | 0.44725 |
| Acylpyruvase FAHD1 | Q8R0F8 | Fahd1 | 25 kDa | 6 C | 0.750 | 0.5205 |
| Glutamate--cysteine ligase regulatory subunit | O09172 | Gclm | 31 kDa | 6 C | 0.747 | 0.3067 |
| 26S proteasome non-ATPase regulatory subunit 9 | Q9CR00 | Psmd9 | 25 kDa | 3 C | 0.745 | 0.45928 |
| Coronin-1C | Q9WUM4 | Coro1c | 53 kDa | 12 C | 0.743 | 0.25507 |
| NEDD8-conjugating enzyme Ubc12 | P61082 | Ube2m | 21 kDa | 5 C | 0.743 | 0.25507 |
| Persulfide dioxygenase ETHE1 | Q9DCM0 | Ethe1 | 28 kDa | 9 C | 0.740 | 0.18715 |
| Fatty aldehyde dehydrogenase | P47740 | Aldh3a2 | 54 kDa | 8 C | 0.720 | 0.52751 |
| Cysteine sulfinic acid decarboxylase | Q9DBE0 | Csad | 55 kDa | 11 C | 0.720 | 0.32488 |
| Phenazine biosynthesis-like domain-containing protein 1 | Q9DCG6 | Pbld1 | 32 kDa | 3 C | 0.716 | 0.49197 |
| Radixin | P26043 | Rdx | 69 kDa | 1 C | 0.716 | 0.18252 |
| Calnexin | P35564 | Canx | 67 kDa | 7 C | 0.712 | 0.49999 |
| Ig gamma-1 chain C region | P01869 | Ighg1 | 43 kDa | 12 C | 0.711 | 0.10901 |
| Ras-related protein Rab-1A | P62821 | Rab1A | 23 kDa | 4 C | 0.711 | 0.23806 |

| | | | | | | |
|---|---|---|---|---|---|---|
| Glucose-6-phosphate isomerase | P06745 | Gpi | 63 kDa | 4 C | 0.708 | 0.41326 |
| Glycine N-acyltransferase | Q91XE0 | Glyat | 34 kDa | 5 C | 0.701 | 0.67887 |
| Isochorismatase domain-containing protein 1 | Q91V64 | Isoc1 | 32 kDa | 5 C | 0.697 | 0.39483 |
| Glutaredoxin-1 | Q9QUH0 | Glrx | 12 kDa | 5 C | 0.690 | 0.6964 |
| Macrophage migration inhibitory factor | P34884 | Mif | 13 kDa | 3 C | 0.688 | 0.51697 |
| NADH dehydrogenase [ubiquinone] 1 alpha subcomplex subunit 2 | Q9CQ75 | Ndufa2 | 11 kDa | 2 C | 0.686 | 0.12498 |
| Lysosome-associated membrane glycoprotein 1 | P11438 | Lamp1 | 44 kDa | 8 C | 0.680 | 0.2958 |
| Pterin-4-alpha-carbinolamine dehydratase | P61458 | Pcbd1 | 12 kDa | 1 C | 0.680 | 0.41653 |
| N(G) | Q9CWS0 | Ddah1 | 31 kDa | 7 C | 0.680 | 0.35829 |
| Isocitrate dehydrogenase [NADP] cytoplasmic | O88844 | Idh1 | 47 kDa | 7 C | 0.678 | 0.1287 |
| Epithelial cell adhesion molecule | Q99JW5 | Epcam | 35 kDa | 12 C | 0.678 | 0.4281 |
| Argininosuccinate lyase | Q91YI0 | Asl | 52 kDa | 13 C | 0.678 | 0.46108 |
| Eukaryotic translation initiation factor 3 subunit I | Q9QZD9 | Eif3i | 36 kDa | 7 C | 0.677 | 0.32853 |
| Ig kappa chain V-III region PC 7043 | P01665 | | 12 kDa | 2 C | 0.674 | 0.03569 |
| Fatty acid-binding protein | P12710 | Fabp1 | 14 kDa | 1 C | 0.671 | 0.26701 |
| Proliferation-associated protein 2G4 | P50580 | Pa2g4 | 44 kDa | 6 C | 0.669 | 0.46078 |
| Junction plakoglobin | Q02257 | Jup | 82 kDa | 13 C | 0.667 | 0.29235 |
| Desmoplakin | E9Q557 | Dsp | 333 kDa | 43 C | 0.667 | 0.30829 |
| Isoform 4 of Thioredoxin reductase 2 | Q9JLT4-4 | Txnrd2 | 53 kDa | 11 C | 0.661 | 0.00468 |
| Legumain | O89017 | Lgmn | 49 kDa | 7 C | 0.655 | 0.15576 |
| Prohibitin | P67778 | Phb | 30 kDa | 1 C | 0.649 | 0.72108 |
| Isoform 2 of Alpha-aminoadipic semialdehyde dehydrogenase | Q9DBF1-2 | Aldh7a1 | 56 kDa | 9 C | 0.647 | 0.00015 |
| Cytochrome c oxidase subunit 6B1 | P56391 | Cox6b1 | 10 kDa | 4 C | 0.646 | 0.55909 |
| Acid sphingomyelinase-like phosphodiesterase 3a | P70158 | Smpdl3a | 50 kDa | 9 C | 0.644 | 0.5262 |
| Isoform 2 of Septin-4 | P28661-2 | 43347 | 53 kDa | 8 C | 0.640 | 0.19702 |
| Eukaryotic translation initiation factor 6 | O55135 | Eif6 | 27 kDa | 8 C | 0.640 | 0.04794 |
| Methionine aminopeptidase 2 | O08663 | Metap2 | 53 kDa | 15 C | 0.629 | 0.61331 |
| Glyoxalase domain-containing protein 4 | Q9CPV4 | Glod4 | 33 kDa | 5 C | 0.624 | 0.39269 |
| Carbonic anhydrase 1 | P13634 | Ca1 | 28 kDa | 1 C | 0.624 | 0.39269 |
| Phenylalanine-4-hydroxylase | P16331 | Pah | 52 kDa | 9 C | 0.610 | 0.32819 |
| Retinal dehydrogenase 1 | P24549 | Aldh1a1 | 54 kDa | 11 C | 0.606 | 0.02351 |
| Carnitine O-palmitoyltransferase 2 | P52825 | Cpt2 | 74 kDa | 10 C | 0.593 | 0.45986 |
| Ig kappa chain V-V region HP 124E1 | P01647 | | 12 kDa | 2 C | 0.582 | 0.1053 |
| Alpha-1-antitrypsin 1-5 | Q00898 | Serpina1e | 46 kDa | 4 C | 0.582 | 0.03339 |
| NADH dehydrogenase [ubiquinone] iron-sulfur protein 2 | Q91WD5 | Ndufs2 | 53 kDa | 7 C | 0.566 | 0.28879 |
| Gamma-glutamylcyclotransferase | Q9D7X8 | Ggct | 21 kDa | 5 C | 0.551 | 0.27475 |
| Acylcarnitine hydrolase | Q91WG0 | Ces2c | 62 kDa | 5 C | 0.550 | 0.17219 |
| Isoform 2 of Acyl-coenzyme A synthetase ACSM3 | Q3UNX5-2 | Acsm3 | 70 kDa | 10 C | 0.547 | 0.22207 |
| Complement component C8 alpha chain | Q8K182 | C8a | 66 kDa | 29 C | 0.533 | 0.03577 |
| Isoform Long of Estradiol 17-beta-dehydrogenase 8 | P50171-2 | Hsd17b8 | 28 kDa | 4 C | 0.523 | 0.32655 |
| Nascent polypeptide-associated complex subunit alpha | P70670 | Naca | 220 kDa | 16 C | 0.516 | 0.10022 |
| Creatine kinase B-type | Q04447 | Ckb | 43 kDa | 5 C | 0.516 | 0.13563 |
| Isoform 2 of 5'-3' exoribonuclease 2 | Q9DBR1-2 | Xrn2 | 108 kDa | 17 C | 0.512 | 0.1094 |
| Oxidoreductase HTATIP2 | Q9Z2G9 | Htatip2 | 27 kDa | 4 C | 0.487 | 0.54552 |
| Membrane-associated progesterone receptor component 1 | O55022 | Pgrmc1 | 22 kDa | 2 C | 0.487 | 0.54552 |
| 6-phosphogluconate dehydrogenase | Q9DCD0 | Pgd | 53 kDa | 9 C | 0.487 | 0.54552 |
| Isopentenyl-diphosphate Delta-isomerase 1 | P58044 | Idi1 | 26 kDa | 8 C | 0.480 | 0.16096 |
| Acetyl-coenzyme A synthetase 2-like | Q99NB1 | Acss1 | 75 kDa | 13 C | 0.469 | 0.08691 |
| Transmembrane emp24 domain-containing protein 10 | Q9D1D4 | Tmed10 | 25 kDa | 3 C | 0.468 | 0.35426 |
| Mitochondrial amidoxime reducing component 2 | Q922Q1 | 43161 | 38 kDa | 11 C | 0.467 | 0.09687 |
| 60S ribosomal protein L30 | P62889 | Rpl30 | 13 kDa | 3 C | 0.453 | 0.28279 |
| 60S ribosomal protein L5 | P47962 | Rpl5 | 34 kDa | 4 C | 0.449 | 0.22315 |
| Protein S100-A11 | P50543 | S100a11 | 11 kDa | 3 C | 0.438 | 0.313 |
| Arginase-1 | Q61176 | Arg1 | 35 kDa | 3 C | 0.438 | 0.39455 |
| Neutral cholesterol ester hydrolase 1 | Q8BLF1 | Nceh1 | 46 kDa | 4 C | 0.420 | 0.18643 |
| Voltage-dependent anion-selective channel protein 2 | Q60930 | Vdac2 | 32 kDa | 11 C | 0.415 | 0.14455 |
| Isoform 2 of Acylamino-acid-releasing enzyme | Q8R146-2 | Apeh | 80 kDa | 16 C | 0.367 | 0.33557 |
| ATP-binding cassette sub-family D member 3 | P55096 | Abcd3 | 75 kDa | 9 C | 0.350 | 0.28281 |
| Alkyldihydroxyacetonephosphate synthase | Q8C0I1 | Agps | 72 kDa | 13 C | 0.273 | 0.1029 |
| Peptidyl-prolyl cis-trans isomerase C | P30412 | Ppic | 23 kDa | 2 C | 0.267 | 0.26411 |

| | | | | | | |
|---|---|---|---|---|---|---|
| GTP cyclohydrolase 1 feedback regulatory protein | P99025 | Gchfr | 10 kDa | 2 C | 0.267 | 0.26411 |
| Glycerol-3-phosphate dehydrogenase | Q64521 | Gpd2 | 81 kDa | 8 C | 0.228 | 0.14917 |
| Aldehyde oxidase 3 | G3X982 | Aox3 | 147 kDa | 38 C | 0.218 | 0.06091 |
| Sulfide:quinone oxidoreductase | Q9R112 | Sqor | 50 kDa | 8 C | 0.174 | 0.29235 |
| Regucalcin | Q64374 | Rgn | 33 kDa | 9 C | 0.095 | 0.08922 |
| Isoform 2 of RNA-binding protein Musashi homolog 2 | Q920Q6-2 | Msi2 | 35 kDa | 3 C | 0.480 | 0.00653 |
| Isoform 2 of ATP-dependent (S)-NAD(P)H-hydrate dehydratase | Q9CZ42-2 | Naxd | 32 kDa | 8 C | 0.410 | 0.02434 |
| Glycine N-methyltransferase | Q9QXF8 | Gnmt | 33 kDa | 8 C | 0.354 | 0.00138 |
| Betaine--homocysteine S-methyltransferase 1 | O35490 | Bhmt | 45 kDa | 8 C | 0.350 | 0.00242 |
| Desmoglein-1-alpha | Q61495 | Dsg1a | 115 kDa | 18 C | 0.187 | 0.00718 |
