## Supplemental Table 3 for "Dietary restriction transforms the protein sulfhydrome in a tissue-specific and cystathionine γ-lyase-dependent manner"

| Supplemental Table 3: Dietary Impact on Skeletal Muscle (Quadriceps) Sulfhydrome | | | | | | |
|---|---|---|---|---|---|---|
| Protein Name | Accession Number | Alternate ID | Molecular Weight | Cysteine Residues | DR/AL Spectral Count Ratio | P-value |
| Nucleolin | P09405 | Ncl | 77 kDa | 1 C | 26.200 | 0.0269 |
| Ubiquitin carboxyl-terminal hydrolase isozyme L1 | Q9R0P9 | Uchl1 | 25 kDa | 6 C | 22.200 | 0.01288 |
| UV excision repair protein RAD23 homolog A | P54726 | Rad23a | 40 kDa | 1 C | 18.000 | 0.03694 |
| Fatty acid-binding protein, epidermal | Q05816 | Fabp5 | 15 kDa | 6 C | 12.200 | 0.02824 |
| Non-specific lipid-transfer protein | P32020 | Scp2 | 59 kDa | 11 C | 12.000 | 0.00186 |
| Myosin regulatory light chain 2, skeletal muscle isoform | P97457 | Mylpf | 19 kDa | 2 C | 8.769 | 0.04807 |
| Gephyrin | Q8BUV3 | Gphn | 83 kDa | 13 C | 6.215 | 0.03147 |
| Poly(rC)-binding protein 1 | P60335 | Pcbp1 | 37 kDa | 9 C | 6.215 | 0.03147 |
| Filamin-C | Q8VHX6 | Flnc | 291 kDa | 46 C | 5.100 | 0.00983 |
| Glucosidase 2 subunit beta | O08795 (+1) | Prkcsh | 59 kDa | 17 C | 4.923 | 0.00726 |
| Calreticulin | P14211 | Calr | 48 kDa | 6 C | 4.667 | 0.03677 |
| NADH dehydrogenase [ubiquinone] iron-sulfur protein 4, mitochondrial | Q9CXZ1 | Ndufs4 | 20 kDa | 1 C | 4.645 | 0.00925 |
| DAZ-associated protein 1 | Q9JII5 (+1) | Dazap1 | 43 kDa | 4 C | 4.364 | 0.00133 |
| Dihydropteridine reductase | Q8BVI4 | Qdpr | 26 kDa | 4 C | 4.174 | 0.02106 |
| Heterogeneous nuclear ribonucleoprotein K | P61979 (+2) | Hnrnpk | 51 kDa | 5 C | 4.000 | 0.0286 |
| Methylthioribulose-1-phosphate dehydratase | Q9WVQ5 | Apip | 27 kDa | 11 C | 3.840 | 0.04143 |
| S-formylglutathione hydrolase | Q9R0P3 | Esd | 31 kDa | 10 C | 3.707 | 0.03128 |
| Galectin-1 | P16045 | Lgals1 | 15 kDa | 6 C | 3.604 | 0.00142 |
| Elongin-B | P62869 | Elob | 13 kDa | 1 C | 3.273 | 0.0357 |
| Kininogen-1 | O08677 | Kng1 | 73 kDa | 19 C | 2.980 | 0.02106 |
| Serpin B6 | Q60854 | Serpinb6 | 43 kDa | 6 C | 2.930 | 0.01776 |
| Isoform 3 of Titin | A2ASS6-3 | Ttn | 619 kDa | 498 C | 2.830 | 0.01943 |
| Serine/threonine-protein phosphatase PP1-gamma catalytic subunit | P63087 (+1) | Ppp1cc | 37 kDa | 13 C | 2.691 | 0.02293 |
| Glutathione S-transferase P 1 | P19157 | Gstp1 | 24 kDa | 3 C | 2.629 | 0.01018 |
| Pregnancy zone protein | Q61838 | Pzp | 166 kDa | 24 C | 2.614 | 0.02776 |
| Serotransferrin | Q921I1 | Tf | 77 kDa | 38 C | 2.607 | 0.02948 |
| Titin | A2ASS6 | Ttn | 3906 kDa | 498 C | 2.404 | 0.02273 |
| Vimentin | P20152 | Vim | 54 kDa | 1 C | 2.286 | 0.01994 |
| Peroxiredoxin-5, mitochondrial | P99029 (+1) | Prdx5 | 22 kDa | 6 C | 2.232 | 0.03645 |
| Transthyretin | P07309 | Ttr | 16 kDa | 2 C | 2.133 | 0.00157 |
| Heat shock cognate 71 kDa protein | P63017 | Hspa8 | 71 kDa | 4 C | 2.117 | 0.03575 |
| 14 kDa phosphohistidine phosphatase | Q9DAK9 | Phpt1 | 14 kDa | 3 C | 2.067 | 0.02263 |
| Myristoylated alanine-rich C-kinase substrate | P26645 | Marcks | 30 kDa | 1 C | 2.065 | 0.04608 |
| Serine/threonine-protein phosphatase PP1-alpha catalytic subunit | P62137 | Ppp1ca | 38 kDa | 13 C | 2.027 | 0.03511 |
| Glucose-6-phosphate isomerase | P06745 | Gpi | 63 kDa | 4 C | 38.400 | 0.28352 |
| Periaxin | O55103 | Prx | 148 kDa | 4 C | 20.200 | 0.14732 |
| Carboxylesterase 1C | P23953 | Ces1c | 61 kDa | 5 C | 18.800 | 0.40708 |
| 60 kDa heat shock protein, mitochondrial | P63038 | Hspd1 | 61 kDa | 3 C | 18.600 | 0.19721 |
| Alpha-2-HS-glycoprotein | P29699 | Ahsg | 37 kDa | 14 C | 18.400 | 0.20477 |
| Dystrophin | P11531 | Dmd | 426 kDa | 37 C | 15.385 | 0.14529 |
| Isoform 3 of Cellular nucleic acid-binding protein | P53996-3 | Cnbp | 19 kDa | 22 C | 14.400 | 0.14941 |
| Beta-2-glycoprotein 1 | Q01339 | Apoh | 39 kDa | 23 C | 14.200 | 0.05145 |
| Four and a half LIM domains protein 3 | Q9R059 | Fhl3 | 32 kDa | 35 C | 14.200 | 0.12498 |
| Medium-chain specific acyl-CoA dehydrogenase, mitochondrial | P45952 | Acadm | 46 kDa | 8 C | 12.600 | 0.31777 |
| Troponin C, skeletal muscle | P20801 | Tnnc2 | 18 kDa | 1 C | 10.800 | 0.40708 |
| Fructose-1,6-bisphosphatase isozyme 2 | P70695 | Fbp2 | 37 kDa | 5 C | 10.600 | 0.29934 |
| Quinone oxidoreductase | P47199 | Cryz | 35 kDa | 5 C | 10.400 | 0.09207 |
| Epiplakin | Q8R0W0 | Eppk1 | 725 kDa | 49 C | 8.800 | 0.40708 |
| Tropomyosin beta chain | P58774 | Tpm2 | 33 kDa | 2 C | 8.800 | 0.40918 |
| Succinyl-CoA:3-ketoacid coenzyme A transferase 1, mitochondrial | Q9D0K2 | Oxct1 | 56 kDa | 7 c | 8.800 | 0.40708 |
| Hydroxyacylglutathione hydrolase, mitochondrial | Q99KB8 (+1) | Hagh | 34 kDa | 8 C | 8.800 | 0.40708 |
| Plectin | Q9QXS1 (+1) | Plec | 534 kDa | 34 C | 8.658 | 0.0675 |
| Inorganic pyrophosphatase 2, mitochondrial | Q91VM9 | Ppa2 | 38 kDa | 8 C | 8.400 | 0.10695 |
| Obscurin | A2AAJ9 | Obscn | 966 kDa | 207 C | 8.400 | 0.19436 |
| Isoform 2 of Reticulon-2 | O70622-2 | Rtn2 | 22 kDa | 4 C | 6.800 | 0.40708 |
| Ceruloplasmin | Q61147 | Cp | 121 kDa | 14 C | 6.800 | 0.40708 |
| NADH dehydrogenase [ubiquinone] 1 beta subcomplex subunit 10 | Q9DCS9 | Ndufb10 | 21 kDa | 5 C | 6.800 | 0.40708 |

| | | | | | | |
|---|---|---|---|---|---|---|
| Bifunctional purine biosynthesis protein PURH | Q9CWJ9 | Atic | 64 kDa | 9 C | 6.800 | 0.40708 |
| Lumican | P51885 | Lum | 38 kDa | 7 C | 6.769 | 0.06472 |
| NADH dehydrogenase [ubiquinone] 1 alpha subcomplex subunit 7 | Q9Z1P6 | Ndufa7 | 13 kDa | 1 C | 6.600 | 0.23224 |
| Aminoacyl tRNA synthase complex-interacting multifunctional protein 1 | P31230 | Aimp1 | 34 kDa | 7 C | 6.600 | 0.23224 |
| Calpain small subunit 1 | O88456 | Capns1 | 28 kDa | 2 C | 6.600 | 0.23224 |
| Serine/threonine-protein phosphatase 2A catalytic subunit alpha isoform | P63330 | Ppp2ca | 36 kDa | 10 C | 6.600 | 0.23224 |
| Inorganic pyrophosphatase | Q9D819 | Ppa1 | 33 kDa | 8 C | 6.600 | 0.23224 |
| Secernin-3 | Q3TMH2 | Scrn3 | 48 kDa | 7 C | 6.600 | 0.23224 |
| Selenoprotein F | Q9ERR7 | Selenof | 18 kDa | 7 C | 6.600 | 0.23224 |
| Endoribonuclease LACTB2 | Q99KR3 | Lactb2 | 33 kDa | 5 C | 6.600 | 0.23224 |
| von Willebrand factor A domain-containing protein 1 | Q8R2Z5 | Vwa1 | 45 kDa | 2 C | 6.600 | 0.23224 |
| Propionyl-CoA carboxylase beta chain, mitochondrial | Q99MN9 | Pccb | 58 kDa | 11 C | 6.215 | 0.1195 |
| 3-hydroxyisobutyrate dehydrogenase, mitochondrial | Q99L13 | Hibadh | 35 kDa | 12 C | 5.818 | 0.07775 |
| Tubulin beta-2A chain | Q7TMM9 | Tubb2a | 50 kDa | 7 C | 5.785 | 0.46828 |
| UV excision repair protein RAD23 homolog B | P54728 | Rad23b | 44 kDa | 1 C | 5.662 | 0.18708 |
| 14-3-3 protein sigma | O70456 | Sfn | 28 kDa | 2 C | 5.600 | 0.07479 |
| Cysteine and glycine-rich protein 3 | P50462 | Csrp3 | 21 kDa | 16 C | 5.108 | 0.26467 |
| Ribonuclease inhibitor | Q91VI7 | Rnh1 | 50 kDa | 30 C | 5.073 | 0.12372 |
| Thioredoxin domain-containing protein 5 | Q91W90 | Txndc5 | 46 kDa | 12 C | 4.923 | 0.11494 |
| Decorin | P28654 | Dcn | 40 kDa | 6 C | 4.800 | 0.40708 |
| NADH dehydrogenase [ubiquinone] 1 alpha subcomplex subunit 8 | Q9DCJ5 | Ndufa8 | 20 kDa | 8 C | 4.800 | 0.40708 |
| Troponin I, slow skeletal muscle | Q9WUZ5 | Tnni1 | 22 kDa | 3 C | 4.800 | 0.40708 |
| Electron transfer flavoprotein-ubiquinone oxidoreductase, mitochondrial | Q921G7 | Etfdh | 68 kDa | 15 C | 4.800 | 0.40708 |
| Platelet glycoprotein 4 | Q08857 | Cd36 | 53 kDa | 10 C | 4.800 | 0.40708 |
| NADH dehydrogenase [ubiquinone] iron-sulfur protein 8, mitochondrial | Q8K3J1 | Ndufs8 | 24 kDa | 8 C | 4.800 | 0.40708 |
| Ras-related protein Rab-11B | P46638 | Rab11b | 24 kDa | 2 C | 4.800 | 0.40708 |
| Calpain-1 catalytic subunit | O35350 | Capn1 | 82 kDa | 12 C | 4.800 | 0.40708 |
| NADH dehydrogenase [ubiquinone] 1 beta subcomplex subunit 8, mitochondrial | Q9D6J5 | Ndufb8 | 22 kDa | 1 C | 4.800 | 0.40708 |
| NADH dehydrogenase [ubiquinone] 1 beta subcomplex subunit 7 | Q9CR61 | Ndufb7 | 16 kDa | 4 C | 4.800 | 0.40708 |
| Desmoglein-4 | Q7TMD7 | Dsg4 | 114 kDa | 27 C | 4.800 | 0.40708 |
| cAMP-dependent protein kinase type II-alpha regulatory subunit | P12367 | Prkar2a | 45 kDa | 6 C | 4.800 | 0.40708 |
| Density-regulated protein | Q9CQJ6 | Denr | 22 kDa | 7 C | 4.800 | 0.40708 |
| Glycogen phosphorylase, muscle form | Q9WUB3 | Pygm | 97 kDa | 8 C | 4.000 | 0.06579 |
| Myosin-binding protein H | P70402 | Mybph | 53 kDa | 8 C | 3.902 | 0.08337 |
| Mitochondrial fission 1 protein | Q9CQ92 | Fis1 | 17 kDa | 1 C | 3.826 | 0.07625 |
| Dual specificity protein phosphatase 3 | Q9D7X3 | Dusp3 | 20 kDa | 4 C | 3.815 | 0.21303 |
| Histidine-rich glycoprotein | Q9ESB3 | Hrg | 59 kDa | 17 C | 3.754 | 0.08807 |
| NADH dehydrogenase [ubiquinone] iron-sulfur protein 3, mitochondrial | Q9DCT2 | Ndufs3 | 30 kDa | 3 C | 3.636 | 0.06454 |
| Cytochrome b-c1 complex subunit Rieske, mitochondrial | Q9CR68 | Uqcrfs1 | 29 kDa | 5 C | 3.554 | 0.10807 |
| Sulfite oxidase, mitochondrial | Q8R086 | Suox | 61 kDa | 9 C | 3.532 | 0.16798 |
| Carbonic anhydrase 2 | P00920 | Ca2 | 29 kDa | 2 C | 3.478 | 0.06661 |
| Isocitrate dehydrogenase [NADP], mitochondrial | P54071 | Idh2 | 51 kDa | 8 C | 3.432 | 0.50182 |
| Translationally-controlled tumor protein | P63028 | Tpt1 | 19 kDa | 2 C | 3.309 | 0.10533 |
| Isocitrate dehydrogenase [NAD] subunit alpha, mitochondrial | Q9D6R2 | Idh3a | 40 kDa | 8 C | 3.262 | 0.43068 |
| Murinoglobulin-1 | P28665 | Mug1 | 165 kDa | 25 C | 3.262 | 0.34006 |
| Desmin | P31001 | Des | 53 kDa | 1 C | 3.200 | 0.14074 |
| Pyridoxal kinase | Q8K183 | Pdxk | 35 kDa | 5 C | 3.200 | 0.2954 |
| Xaa-Pro dipeptidase | Q11136 | Pepd | 55 kDa | 17 C | 3.176 | 0.14833 |
| Superoxide dismutase [Cu-Zn] | P08228 | Sod1 | 16 kDa | 3 C | 3.138 | 0.12115 |
| Platelet-activating factor acetylhydrolase IB subunit beta | Q61206 | Pafah1b2 | 26 kDa | 3 C | 3.138 | 0.12115 |
| Eukaryotic translation initiation factor 5A-1 | P63242 | Eif5a | 17 kDa | 4 C | 3.072 | 0.14399 |
| 2-oxoglutarate dehydrogenase, mitochondrial | Q60597 | Ogdh | 116 kDa | 21 C | 3.055 | 0.05063 |
| Dihydrolipoyllysine-residue acetyltransferase component of pyruvate dehydrogenase comp | Q8BMF4 | Dlat | 68 kDa | 10 C | 2.857 | 0.05858 |
| Vesicle-associated membrane protein-associated protein B | Q9QY76 | Vapb | 27 kDa | 4 C | 2.839 | 0.07779 |
| Glutathione S-transferase Mu 2 | P15626 | Gstm2 | 26 kDa | 3 C | 2.775 | 0.21442 |
| Isoform 2 of Myomesin-1 | Q62234-2 | Myom1 | 175 kDa | 21 C | 2.767 | 0.10635 |
| Elongation factor 2 | P58252 | Eef2 | 95 kDa | 7 C | 2.743 | 0.11789 |
| Ig kappa chain V-III region PC 7043 | P01665 (+1) | | 12 kDa | 2 C | 2.715 | 0.34395 |
| Isochorismatase domain-containing protein 2A | P85094 | Isoc2a | 22 kDa | 6 C | 2.646 | 0.44946 |
| Filamin-A | Q8BTM8 | Flna | 281 kDa | 38 C | 2.618 | 0.42866 |
| Retinal dehydrogenase 1 | P24549 | Aldh1a1 | 54 kDa | 11 C | 2.595 | 0.06715 |

| | | | | | | |
|---|---|---|---|---|---|---|
| Hemopexin | Q91X72 | Hpx | 51 kDa | 13 C | 2.592 | 0.21135 |
| S-phase kinase-associated protein 1 | Q9WTX5 | Skp1 | 19 kDa | 3 C | 2.585 | 0.28648 |
| Far upstream element-binding protein 2 | Q3U0V1 | Khsrp | 77 kDa | 8 C | 2.585 | 0.28648 |
| Elongation factor 1-alpha 1 | P10126 | Eef1a1 | 50 kDa | 6 C | 2.582 | 0.30789 |
| Long-chain specific acyl-CoA dehydrogenase, mitochondrial | P51174 | Acadl | 48 kDa | 7 C | 2.568 | 0.12729 |
| Peptidyl-prolyl cis-trans isomerase A | P17742 | Ppia | 18 kDa | 3 C | 2.560 | 0.05807 |
| Methylmalonate-semialdehyde dehydrogenase [acylating], mitochondrial | Q9EQ20 | Aldh6a1 | 58 kDa | 8 C | 2.525 | 0.20855 |
| Adenylosuccinate synthetase isozyme 1 | P28650 (+1) | Adssl1 | 50 kDa | 5 C | 2.510 | 0.22978 |
| Electron transfer flavoprotein subunit alpha, mitochondrial | Q99LC5 | Etfa | 35 kDa | 6 C | 2.500 | 0.25332 |
| Carbonic anhydrase 3 | P16015 | Ca3 | 29 kDa | 5 C | 2.421 | 0.05267 |
| Cytosolic non-specific dipeptidase | Q9D1A2 | Cndp2 | 53 kDa | 8 C | 2.382 | 0.07813 |
| Alpha-actinin-3 | O88990 | Actn3 | 103 kDa | 11 C | 2.372 | 0.08231 |
| Complement component C8 gamma chain | Q8VCG4 | C8g | 23 kDa | 3 C | 2.366 | 0.06542 |
| Ubiquitin recognition factor in ER-associated degradation protein 1 | P70362 | Ufd1 | 34 kDa | 5 C | 2.348 | 0.10945 |
| Sarcoplasmic/endoplasmic reticulum calcium ATPase 1 | Q8R429 | Atp2a1 | 109 kDa | 24 C | 2.323 | 0.05857 |
| Isoform 3 of Elongation factor 1-delta | P57776-3 | Eef1d | 73 kDa | 2 C | 2.323 | 0.12569 |
| Beta-enolase | P21550 | Eno3 | 47 kDa | 6 C | 2.321 | 0.10287 |
| Myosin light chain 1/3, skeletal muscle isoform | P05977 | Myl1 | 21 kDa | 2 C | 2.275 | 0.05101 |
| Dihydrolipoyl dehydrogenase, mitochondrial | O08749 | Dld | 54 kDa | 9 C | 2.267 | 0.07807 |
| Complement C3 | P01027 | C3 | 186 kDa | 27 C | 2.230 | 0.31121 |
| Plakophilin-1 | P97350 | Pkp1 | 81 kDa | 25 C | 2.226 | 0.63039 |
| Glutaredoxin-1 | Q9QUH0 | Glrx | 12 kDa | 5 C | 2.218 | 0.19366 |
| 40S ribosomal protein S12 | P63323 | Rps12 | 15 kDa | 7 C | 2.218 | 0.29199 |
| Troponin I, fast skeletal muscle | P13412 | Tnni2 | 21 kDa | 3 C | 2.182 | 0.24325 |
| Proteasome subunit alpha type-4 | Q9R1P0 | Psma4 | 29 kDa | 5 C | 2.171 | 0.06406 |
| NADH dehydrogenase [ubiquinone] flavoprotein 1, mitochondrial | Q91YT0 | Ndufv1 | 51 kDa | 12 C | 2.171 | 0.1214 |
| Isocitrate dehydrogenase [NADP] cytoplasmic | O88844 | Idh1 | 47 kDa | 7 C | 2.169 | 0.28241 |
| Acetyl-CoA acetyltransferase, mitochondrial | Q8QZT1 | Acat1 | 45 kDa | 6 C | 2.166 | 0.27958 |
| Ankyrin repeat domain-containing protein 2 | Q9WV06 | Ankrd2 | 37 kDa | 3 C | 2.147 | 0.24938 |
| Endoplasmic reticulum resident protein 44 | Q9D1Q6 | Erp44 | 47 kDa | 7 C | 2.146 | 0.06771 |
| Histidine triad nucleotide-binding protein 1 | P70349 | Hint1 | 14 kDa | 2 C | 2.146 | 0.1144 |
| Four and a half LIM domains protein 1 | P97447 | Fhl1 | 32 kDa | 34 C | 2.145 | 0.09171 |
| DnaJ homolog subfamily B member 11 | Q99KV1 | Dnajb11 | 41 kDa | 5 C | 2.122 | 0.37702 |
| Radixin | P26043 | Rdx | 69 kDa | 1 C | 2.092 | 0.62111 |
| Histidine ammonia-lyase | P35492 | Hal | 72 kDa | 12 C | 2.092 | 0.62111 |
| Sarcoplasmic/endoplasmic reticulum calcium ATPase 2 | O55143 (+1) | Atp2a2 | 115 kDa | 29 C | 2.090 | 0.34813 |
| Insulin-degrading enzyme | Q9JHR7 | Ide | 118 kDa | 13 C | 2.087 | 0.18026 |
| Heat shock 70 kDa protein 4 | Q61316 | Hspa4 | 94 kDa | 14 C | 2.074 | 0.05833 |
| Vinculin | Q64727 | Vcl | 117 kDa | 10 C | 2.073 | 0.05484 |
| Methionine-R-sulfoxide reductase B3, mitochondrial | Q8BU85 | Msrb3 | 27 kDa | 8 C | 2.044 | 0.11166 |
| NADH dehydrogenase [ubiquinone] flavoprotein 2, mitochondrial | Q9D6J6 | Ndufv2 | 27 kDa | 6 C | 2.036 | 0.06312 |
| Enoyl-CoA delta isomerase 1, mitochondrial | P42125 | Eci1 | 32 kDa | 5 C | 2.031 | 0.50045 |
| Ferritin heavy chain | P09528 | Fth1 | 21 kDa | 3 C | 2.031 | 0.50045 |
| Target of Myb protein 1 | O88746 | Tom1 | 54 kDa | 4 C | 2.031 | 0.50045 |
| Glutamine synthetase | P15105 | Glul | 42 kDa | 13 C | 2.031 | 0.50045 |
| Osteoclast-stimulating factor 1 | Q62422 | Ostf1 | 24 kDa | 4 C | 2.031 | 0.50045 |
| Heterogeneous nuclear ribonucleoprotein Q | Q7TMK9 | Syncrip | 70 kDa | 4 C | 2.025 | 0.32529 |
| Cytochrome b-c1 complex subunit 2, mitochondrial | Q9DB77 | Uqcrc2 | 48 kDa | 1 C | 2.025 | 0.36901 |
| Ig kappa chain C region | P01837 | | 12 kDa | 3 C | 2.000 | 0.28292 |
| NADH-ubiquinone oxidoreductase 75 kDa subunit, mitochondrial | Q91VD9 | Ndufs1 | 80 kDa | 18 C | 2.000 | 0.046 |
| Isoform Mt-VDAC1 of Voltage-dependent anion-selective channel protein 1 | Q60932-2 | Vdac1 | 31 kDa | 2 C | 1.990 | 0.52046 |
| Peroxiredoxin-4 | O08807 | Prdx4 | 31 kDa | 4 C | 1.976 | 0.12803 |
| Isoform 2 of Heterogeneous nuclear ribonucleoprotein A3 | Q8BG05-2 | Hnrnpa3 | 37 kDa | 4 C | 1.964 | 0.11594 |
| Carboxylesterase 1D | Q8VCT4 | Ces1d | 62 kDa | 5 C | 1.964 | 0.16701 |
| Synaptopodin 2-like protein | Q8BWB1 | Synpo2l | 103 kDa | 5 C | 1.964 | 0.44764 |
| Proteasome subunit alpha type-5 | Q9Z2U1 | Psma5 | 26 kDa | 3 C | 1.956 | 0.01973 |
| Phosphatidylethanolamine-binding protein 1 | P70296 | Pebp1 | 21 kDa | 3 C | 1.943 | 0.07652 |
| Ig heavy chain V region AC38 205.12 | P06330 | | 13 kDa | 2 C | 1.943 | 0.30091 |
| 14-3-3 protein beta/alpha | Q9CQV8 | Ywhab | 28 kDa | 2 C | 1.929 | 0.0311 |
| Adapter molecule crk | Q64010 | Crk | 34 kDa | 1 C | 1.920 | 0.05646 |
| Thioredoxin-like protein 1 | Q8CDN6 | Txnl1 | 32 kDa | 7 C | 1.920 | 0.1053 |

| | | | | | | |
|---|---|---|---|---|---|---|
| Ubiquitin-conjugating enzyme E2 variant 2 | Q9D2M8 | Ube2v2 | 16 kDa | 1 C | 1.882 | 0.10564 |
| Ubiquitin carboxyl-terminal hydrolase isozyme L3 | Q9JKB1 | Uchl3 | 26 kDa | 3 C | 1.882 | 0.19743 |
| Adenylate kinase isoenzyme 1 | Q9R0Y5 | Ak1 | 22 kDa | 2 C | 1.871 | 0.09995 |
| Stress-70 protein, mitochondrial | P38647 | Hspa9 | 73 kDa | 5 C | 1.867 | 0.07504 |
| Isoform 2 of PDZ and LIM domain protein 7 | Q3TJD7-2 | Pdlim7 | 21 kDa | 19 C | 1.855 | 0.28959 |
| Nucleolysin TIAR | P70318 | Tial1 | 43 kDa | 6 C | 1.855 | 0.28959 |
| Alpha-actinin-4 | P57780 | Actn4 | 105 kDa | 8 C | 1.850 | 0.16594 |
| Profilin-1 | P62962 | Pfn1 | 15 kDa | 3 C | 1.849 | 0.25479 |
| L-lactate dehydrogenase A chain | P06151 | Ldha | 36 kDa | 6 C | 1.846 | 0.0999 |
| Thioredoxin reductase 1, cytoplasmic | Q9JMH6 | Txnrd1 | 67 kDa | 21 C | 1.846 | 0.14897 |
| Fibrinogen gamma chain | Q8VCM7 | Fgg | 49 kDa | 12 C | 1.846 | 0.03294 |
| Fructose-bisphosphate aldolase A | P05064 | Aldoa | 39 kDa | 8 C | 1.816 | 0.1236 |
| Serum albumin | P07724 | Alb | 69 kDa | 36 C | 1.812 | 0.03373 |
| Peroxiredoxin-6 | O08709 | Prdx6 | 25 kDa | 2 C | 1.811 | 0.13327 |
| Dihydrolipoyllysine-residue succinyltransferase component of 2-oxoglutarate dehydrogena | Q9D2G2 | Dlst | 49 kDa | 6 C | 1.811 | 0.25564 |
| Serine protease inhibitor A3K | P07759 | Serpina3k | 47 kDa | 4 C | 1.811 | 0.02049 |
| Cofilin-1 | P18760 | Cfl1 | 19 kDa | 4 C | 1.803 | 0.24054 |
| Isoform 2 of Voltage-dependent L-type calcium channel subunit beta-1 | Q8R3Z5-2 | Cacnb1 | 74 kDa | 3 C | 1.800 | 0.09012 |
| Tubulin alpha-1C chain | P68373 | Tuba1c | 50 kDa | 12 C | 1.800 | 0.26556 |
| Aldehyde dehydrogenase, mitochondrial | P47738 | Aldh2 | 57 kDa | 9 C | 1.800 | 0.04133 |
| Branched-chain-amino-acid aminotransferase, mitochondrial | O35855 | Bcat2 | 44 kDa | 10 C | 1.776 | 0.36014 |
| Hsc70-interacting protein | Q99L47 | St13 | 42 kDa | 3 C | 1.760 | 0.08707 |
| Septin-7 | O55131 | 43350 | 51 kDa | 6 C | 1.760 | 0.21492 |
| Septin-2 | P42208 | 43345 | 42 kDa | 8 C | 1.760 | 0.3067 |
| Malate dehydrogenase, mitochondrial | P08249 | Mdh2 | 36 kDa | 8 C | 1.760 | 0.19787 |
| Elongation factor 1-beta | O70251 | Eef1b | 25 kDa | 3 C | 1.756 | 0.10024 |
| Caveolae-associated protein 4 | A2AMM0 | Cavin4 | 41 kDa | 1 C | 1.756 | 0.27316 |
| Vesicle-associated membrane protein-associated protein A | Q9WV55 | Vapa | 28 kDa | 4 C | 1.756 | 0.33212 |
| Phosphoglucomutase-1 | Q9D0F9 | Pgm1 | 61 kDa | 10 C | 1.755 | 0.27633 |
| Serine/threonine-protein phosphatase PP1-beta catalytic subunit | P62141 | Ppp1cb | 37 kDa | 14 C | 1.752 | 0.0463 |
| Dihydropyrimidinase-related protein 2 | O08553 | Dpysl2 | 62 kDa | 7 C | 1.751 | 0.06527 |
| Fibrinogen beta chain | Q8K0E8 | Fgb | 55 kDa | 12 C | 1.750 | 0.13142 |
| Stress-induced-phosphoprotein 1 | Q60864 | Stip1 | 63 kDa | 11 C | 1.748 | 0.15201 |
| Phosphoglycerate kinase 1 | P09411 | Pgk1 | 45 kDa | 7 C | 1.744 | 0.1199 |
| Peptidyl-prolyl cis-trans isomerase FKBP3 | Q62446 | Fkbp3 | 25 kDa | 1 C | 1.733 | 0.07205 |
| 26S proteasome non-ATPase regulatory subunit 4 | O35226 | Psmd4 | 41 kDa | 4 C | 1.718 | 0.30744 |
| Proteasome subunit beta type-4 | P99026 | Psmb4 | 29 kDa | 2 C | 1.714 | 0.09956 |
| UMP-CMP kinase | Q9DBP5 | Cmpk1 | 22 kDa | 6 C | 1.714 | 0.02072 |
| 14-3-3 protein eta | P68510 | Ywhah | 28 kDa | 3 C | 1.700 | 0.07862 |
| Heterogeneous nuclear ribonucleoproteins A2/B1 | O88569 | Hnrnpa2b1 | 37 kDa | 1 C | 1.700 | 0.14295 |
| Proteasome subunit beta type-3 | Q9R1P1 | Psmb3 | 23 kDa | 5 C | 1.700 | 0.17236 |
| Catalase | P24270 | Cat | 60 kDa | 5 C | 1.694 | 0.04061 |
| Transitional endoplasmic reticulum ATPase | Q01853 | Vcp | 89 kDa | 12 C | 1.687 | 0.06037 |
| Probable aminopeptidase NPEPL1 | Q6NSR8 | Npepl1 | 56 kDa | 16 C | 1.680 | 0.22835 |
| Isoform M1 of Pyruvate kinase PKM | P52480-2 | Pkm | 58 kDa | 9 C | 1.679 | 0.05572 |
| Glyceraldehyde-3-phosphate dehydrogenase | P16858 | Gapdh | 36 kDa | 5 C | 1.677 | 0.05927 |
| Myosin-binding protein C, fast-type | Q5XKE0 | Mybpc2 | 127 kDa | 15 C | 1.673 | 0.12217 |
| Alpha-enolase | P17182 | Eno1 | 47 kDa | 6 C | 1.662 | 0.33785 |
| Succinate dehydrogenase [ubiquinone] iron-sulfur subunit, mitochondrial | Q9CQA3 | Sdhb | 32 kDa | 14 C | 1.659 | 0.0701 |
| Myoglobin | P04247 | Mb | 17 kDa | 1 C | 1.656 | 0.16763 |
| Cofilin-2 | P45591 | Cfl2 | 19 kDa | 2 C | 1.644 | 0.09138 |
| SH3 domain-binding glutamic acid-rich protein | Q9WUZ7 | Sh3bgr | 23 kDa | 1 C | 1.642 | 0.09041 |
| 40S ribosomal protein SA | P14206 | Rpsa | 33 kDa | 2 C | 1.600 | 0.10091 |
| Pyruvate dehydrogenase protein X component, mitochondrial | Q8BKZ9 | Pdhx | 54 kDa | 5 C | 1.600 | 0.11239 |
| Electron transfer flavoprotein subunit beta | Q9DCW4 | Etfb | 28 kDa | 4 C | 1.600 | 0.20003 |
| Transgelin | P37804 | Tagln | 23 kDa | 1 C | 1.600 | 0.37502 |
| Glutathione S-transferase Mu 1 | P10649 | Gstm1 | 26 kDa | 2 C | 1.600 | 0.16978 |
| Phospholipid hydroperoxide glutathione peroxidase, mitochondrial | O70325 (+1) | Gpx4 | 22 kDa | 10 C | 1.600 | 0.27173 |
| Vitamin D-binding protein | P21614 | Gc | 54 kDa | 28 C | 1.600 | 0.72854 |
| Isoform 4 of LIM domain-binding protein 3 | Q9JKS4-4 | Ldb3 | 67 kDa | 21 C | 1.585 | 0.05824 |
| UBX domain-containing protein 1 | Q922Y1 | Ubxn1 | 34 kDa | 2 C | 1.574 | 0.35748 |

| | | | | | | |
|---|---|---|---|---|---|---|
| Thioredoxin | P10639 | Txn | 12 kDa | 6 C | 1.574 | 0.19861 |
| Alpha-actinin-2 | Q9JI91 | Actn2 | 104 kDa | 10 C | 1.571 | 0.24237 |
| Pyruvate dehydrogenase E1 component subunit alpha, somatic form, mitochondrial | P35486 | Pdha1 | 43 kDa | 12 C | 1.569 | 0.11392 |
| Protein disulfide-isomerase | P09103 | P4hb | 57 kDa | 7 C | 1.567 | 0.13391 |
| Ig kappa chain V-II region 26-10 | P01631 | | 12 kDa | 2 C | 1.561 | 0.47453 |
| Phosphoglycerate mutase 2 | O70250 | Pgam2 | 29 kDa | 3 C | 1.549 | 0.19771 |
| 14-3-3 protein zeta/delta | P63101 | Ywhaz | 28 kDa | 3 C | 1.548 | 0.09012 |
| Heterogeneous nuclear ribonucleoprotein H2 | P70333 | Hnrnph2 | 49 kDa | 5 C | 1.548 | 0.20057 |
| Selenium-binding protein 1 | P17563 | Selenbp1 | 53 kDa | 10 C | 1.538 | 0.1735 |
| Tropomyosin alpha-1 chain | P58771 | Tpm1 | 33 kDa | 1 C | 1.535 | 0.70414 |
| Cadherin-13 | Q9WTR5 | Cdh13 | 78 kDa | 7 C | 1.527 | 0.10668 |
| Eukaryotic translation initiation factor 4H | Q9WUK2 | Eif4h | 27 kDa | 1 C | 1.527 | 0.54905 |
| Crk-like protein | P47941 | Crkl | 34 kDa | 2 C | 1.525 | 0.48409 |
| Glutathione peroxidase 1 | P11352 | Gpx1 | 22 kDa | 4 C | 1.520 | 0.04167 |
| Alpha-1-antitrypsin 1-1 | P07758 (+1) | Serpina1a | 46 kDa | 3 C | 1.514 | 0.34643 |
| Myosin light chain 3 | P09542 | Myl3 | 22 kDa | 2 C | 1.500 | 0.06043 |
| 14-3-3 protein epsilon | P62259 | Ywhae | 29 kDa | 3 C | 1.500 | 0.20481 |
| Hemoglobin subunit beta-1 | P02088 | Hbb-b1 | 16 kDa | 2 C | 1.500 | 0.25243 |
| Complement C1q and tumor necrosis factor-related protein 9 | Q4ZJN1 | C1qtnf9 | 35 kDa | 3 C | 1.491 | 0.44031 |
| Glycerol-3-phosphate phosphatase | Q8CHP8 | Pgp | 35 kDa | 8 C | 1.486 | 0.36446 |
| Complement factor I | Q61129 | Cfi | 67 kDa | 40 C | 1.477 | 0.75324 |
| Proteasome subunit alpha type-7 | Q9Z2U0 | Psma7 | 28 kDa | 3 C | 1.477 | 0.03569 |
| Heat shock protein beta-6 | Q5EBG6 | Hspb6 | 18 kDa | 1 C | 1.461 | 0.66058 |
| Protein argonaute-2 | Q8CJG0 | Ago2 | 97 kDa | 22 C | 1.461 | 0.66058 |
| Glutaredoxin-3 | Q9CQM9 | Glrx3 | 38 kDa | 5 C | 1.459 | 0.22591 |
| GMP reductase 1 | Q9DCZ1 | Gmpr | 37 kDa | 9 C | 1.455 | 0.28022 |
| Malate dehydrogenase, cytoplasmic | P14152 | Mdh1 | 37 kDa | 3 C | 1.450 | 0.14557 |
| Mammalian ependymin-related protein 1 | Q99M71 | Epdr1 | 25 kDa | 7 C | 1.440 | 0.12498 |
| Low molecular weight phosphotyrosine protein phosphatase | Q9D358 | Acp1 | 18 kDa | 8 C | 1.440 | 0.28741 |
| Aconitate hydratase, mitochondrial | Q99KI0 | Aco2 | 85 kDa | 13 C | 1.439 | 0.10189 |
| Alpha-1-antitrypsin 1-2 | P22599 | Serpina1b | 46 kDa | 3 C | 1.432 | 0.28845 |
| Striated muscle-specific serine/threonine-protein kinase | Q62407 | Speg | 354 kDa | 44 C | 1.429 | 0.14484 |
| High mobility group protein B1 | P63158 | Hmgb1 | 25 kDa | 3 C | 1.426 | 0.61414 |
| CD5 antigen-like | Q9QWK4 | Cd5l | 39 kDa | 26 C | 1.425 | 0.0975 |
| Superoxide dismutase [Mn], mitochondrial | P09671 | Sod2 | 25 kDa | 4 C | 1.422 | 0.26467 |
| Ubiquitin-conjugating enzyme E2 N | P61089 | Ube2n | 17 kDa | 1 C | 1.420 | 0.47985 |
| Proteasome subunit alpha type-3 | O70435 | Psma3 | 28 kDa | 4 C | 1.415 | 0.10091 |
| PDZ and LIM domain protein 3 | O70209 | Pdlim3 | 34 kDa | 8 C | 1.415 | 0.31264 |
| Proteasome subunit alpha type-1 | Q9R1P4 | Psma1 | 30 kDa | 5 C | 1.415 | 0.50685 |
| Glutathione peroxidase 3 | P46412 | Gpx3 | 25 kDa | 3 C | 1.400 | 0.14956 |
| Heterogeneous nuclear ribonucleoprotein A1 | P49312 | Hnrnpa1 | 34 kDa | 2 C | 1.400 | 0.56145 |
| Protein DJ-1 | Q99LX0 | Park7 | 20 kDa | 4 C | 1.387 | 0.0965 |
| Myopalladin | Q5DTJ9 | Mypn | 144 kDa | 21 C | 1.371 | 0.10901 |
| Junctophilin-1 | Q9ET80 | Jph1 | 72 kDa | 9 C | 1.366 | 0.42196 |
| NADH dehydrogenase [ubiquinone] 1 alpha subcomplex subunit 2 | Q9CQ75 | Ndufa2 | 11 kDa | 2 C | 1.366 | 0.42196 |
| Protein phosphatase 1 regulatory subunit 7 | Q3UM45 | Ppp1r7 | 41 kDa | 2 C | 1.364 | 0.21818 |
| Junctophilin-2 | Q9ET78 | Jph2 | 75 kDa | 3 C | 1.354 | 0.36524 |
| Fumarate hydratase, mitochondrial | P97807 (+1) | Fh | 54 kDa | 4 C | 1.352 | 0.71328 |
| Fatty acid-binding protein, adipocyte | P04117 | Fabp4 | 15 kDa | 2 C | 1.352 | 0.22655 |
| Mitochondrial peptide methionine sulfoxide reductase | Q9D6Y7 (+1) | Msra | 26 kDa | 4 C | 1.350 | 0.14295 |
| Peroxiredoxin-2 | Q61171 | Prdx2 | 22 kDa | 3 C | 1.345 | 0.25598 |
| Lactoylglutathione lyase | Q9CPU0 | Glo1 | 21 kDa | 3 C | 1.342 | 0.4991 |
| Selenoprotein P | P70274 | Selenop | 43 kDa | 18 C | 1.342 | 0.62652 |
| Alpha-1-antitrypsin 1-4 | Q00897 | Serpina1d | 46 kDa | 3 C | 1.338 | 0.50215 |
| Dihydropyrimidinase-related protein 3 | Q62188 | Dpysl3 | 62 kDa | 7 C | 1.333 | 0.23546 |
| Cytochrome b-c1 complex subunit 1, mitochondrial | Q9CZ13 | Uqcrc1 | 53 kDa | 11 C | 1.333 | 0.4282 |
| Carboxymethylenebutenolidase homolog | Q8R1G2 | Cmbl | 28 kDa | 6 C | 1.333 | 0.54906 |
| Succinate dehydrogenase [ubiquinone] flavoprotein subunit, mitochondrial | Q8K2B3 | Sdha | 73 kDa | 19 C | 1.331 | 0.09797 |
| Creatine kinase S-type, mitochondrial | Q6P8J7 | Ckmt2 | 47 kDa | 8 C | 1.327 | 0.25948 |
| Ig heavy chain V-III region A4 | P01796 (+3) | | 13 kDa | 2 C | 1.325 | 0.66207 |
| Aspartate aminotransferase, cytoplasmic | P05201 | Got1 | 46 kDa | 5 C | 1.320 | 0.05095 |

| | | | | | | |
|---|---|---|---|---|---|---|
| Thioredoxin domain-containing protein 17 | Q9CQM5 | Txndc17 | 14 kDa | 6 C | 1.316 | 0.55382 |
| Ubiquitin-conjugating enzyme E2 L3 | P68037 | Ube2l3 | 18 kDa | 3 C | 1.316 | 0.55382 |
| Creatine kinase M-type | P07310 | Ckm | 43 kDa | 8 C | 1.314 | 0.17329 |
| Peroxiredoxin-1 | P35700 | Prdx1 | 22 kDa | 4 C | 1.310 | 0.29778 |
| Proteasome subunit beta type-2 | Q9R1P3 | Psmb2 | 23 kDa | 3 C | 1.300 | 0.06758 |
| Proteasome subunit alpha type-2 | P49722 | Psma2 | 26 kDa | 2 C | 1.300 | 0.53562 |
| Ig gamma-3 chain C region | P03987 (+1) | | 44 kDa | 10 C | 1.300 | 0.74876 |
| Enoyl-CoA hydratase, mitochondrial | Q8BH95 | Echs1 | 31 kDa | 7 C | 1.295 | 0.16501 |
| Cytochrome c, somatic | P62897 | Cycs | 12 kDa | 2 C | 1.287 | 0.24216 |
| PDZ and LIM domain protein 1 | O70400 | Pdlim1 | 36 kDa | 8 C | 1.285 | 0.83201 |
| 78 kDa glucose-regulated protein | P20029 | Hspa5 | 72 kDa | 1 C | 1.280 | 0.22478 |
| Acylphosphatase-2 | P56375 | Acyp2 | 12 kDa | 1 C | 1.280 | 0.35582 |
| Serine hydroxymethyltransferase, cytosolic | P50431 | Shmt1 | 53 kDa | 10 C | 1.280 | 0.48156 |
| Ig mu chain C region | P01872 | Ighm | 50 kDa | 19 C | 1.273 | 0.39714 |
| Heterogeneous nuclear ribonucleoprotein A/B | Q99020 | Hnrnpab | 31 kDa | 2 C | 1.257 | 0.19702 |
| L-lactate dehydrogenase B chain | P16125 | Ldhb | 37 kDa | 5 C | 1.257 | 0.30309 |
| Serine/threonine-protein phosphatase 5 | Q60676 | Ppp5c | 57 kDa | 11 C | 1.257 | 0.4758 |
| Aspartate aminotransferase, mitochondrial | P05202 | Got2 | 47 kDa | 7 C | 1.252 | 0.1363 |
| Isoform 2 of PDZ and LIM domain protein 5 | Q8CI51-2 | Pdlim5 | 36 kDa | 22 C | 1.250 | 0.40708 |
| Triosephosphate isomerase | P17751 | Tpi1 | 32 kDa | 9 C | 1.247 | 0.29664 |
| 14-3-3 protein gamma | P61982 | Ywhag | 28 kDa | 3 C | 1.244 | 0.22598 |
| Serine/threonine-protein phosphatase 2A 55 kDa regulatory subunit B alpha isoform | Q6P1F6 | Ppp2r2a | 52 kDa | 9 C | 1.236 | 0.44313 |
| Aspartyl aminopeptidase | Q9Z2W0 | Dnpep | 52 kDa | 10 C | 1.236 | 0.4734 |
| Transketolase | P40142 | Tkt | 68 kDa | 12 C | 1.236 | 0.85715 |
| Septin-11 | Q8C1B7 (+2) | 43354 | 50 kDa | 6 C | 1.231 | 0.29235 |
| Rho GDP-dissociation inhibitor 1 | Q99PT1 | Arhgdia | 23 kDa | 1 C | 1.231 | 0.7027 |
| Myotilin | Q9JIF9 | Myot | 55 kDa | 8 C | 1.211 | 0.05394 |
| Isoform 2 of Alpha-aminoadipic semialdehyde dehydrogenase | Q9DBF1-2 | Aldh7a1 | 56 kDa | 9 C | 1.210 | 0.32418 |
| Myocilin | O70624 | Myoc | 55 kDa | 9 C | 1.200 | 0.40708 |
| Myosin light chain kinase 2, skeletal/cardiac muscle | Q8VCR8 | Mylk2 | 66 kDa | 12 C | 1.200 | 0.40708 |
| Mth938 domain-containing protein | Q8R0P4 | Aamdc | 13 kDa | 2 C | 1.200 | 0.56303 |
| Protein disulfide-isomerase A6 | Q922R8 | Pdia6 | 48 kDa | 7 C | 1.200 | 0.68453 |
| Annexin A2 | P07356 | Anxa2 | 39 kDa | 5 C | 1.200 | 0.8083 |
| Glia maturation factor beta | Q9CQI3 | Gmfb | 17 kDa | 3 C | 1.200 | 0.82713 |
| ES1 protein homolog, mitochondrial | Q9D172 | D10Jhu81e | 28 kDa | 6 C | 1.200 | 0.82713 |
| Macrophage migration inhibitory factor | P34884 | Mif | 13 kDa | 3 C | 1.190 | 0.72398 |
| Proteasome subunit alpha type-6 | Q9QUM9 | Psma6 | 27 kDa | 8 C | 1.183 | 0.35582 |
| Annexin A5 | P48036 | Anxa5 | 36 kDa | 1 C | 1.181 | 0.81895 |
| Sialic acid synthase | Q99J77 | Nans | 40 kDa | 8 C | 1.180 | 0.65798 |
| Cytochrome c oxidase subunit 5B, mitochondrial | P19536 | Cox5b | 14 kDa | 5 C | 1.169 | 0.38779 |
| 60S ribosomal protein L12 | P35979 | Rpl12 | 18 kDa | 3 C | 1.169 | 0.38779 |
| Fibrinogen alpha chain | E9PV24 | Fga | 87 kDa | 13 C | 1.164 | 0.53892 |
| CAP-Gly domain-containing linker protein 1 | Q922J3 (+1) | Clip1 | 156 kDa | 11 C | 1.148 | 0.51536 |
| Adenosylhomocysteinase | P50247 | Ahcy | 48 kDa | 9 C | 1.143 | 0.57083 |
| Deoxynucleoside triphosphate triphosphohydrolase SAMHD1 | Q60710 | Samhd1 | 73 kDa | 17 C | 1.135 | 0.88273 |
| ADP/ATP translocase 1 | P48962 | Slc25a4 | 33 kDa | 4 C | 1.133 | 0.61902 |
| Succinate-semialdehyde dehydrogenase, mitochondrial | Q8BWF0 | Aldh5a1 | 56 kDa | 10 C | 1.120 | 0.70009 |
| Selenide, water dikinase 2 | P97364 | Sephs2 | 48 kDa | 7 C | 1.120 | 0.68453 |
| NADH dehydrogenase [ubiquinone] iron-sulfur protein 6, mitochondrial | P52503 | Ndufs6 | 13 kDa | 3 C | 1.120 | 0.68453 |
| Alpha-1-antitrypsin 1-5 | Q00898 | Serpina1e | 46 kDa | 4 C | 1.114 | 0.85252 |
| Tubulin polymerization-promoting protein family member 3 | Q9CRB6 | Tppp3 | 19 kDa | 3 C | 1.114 | 0.83985 |
| Fumarylacetoacetate hydrolase domain-containing protein 2A | Q3TC72 | Fahd2 | 35 kDa | 6 C | 1.110 | 0.88436 |
| Proteasome subunit beta type-5 | O55234 | Psmb5 | 29 kDa | 3 C | 1.105 | 0.68071 |
| Hepatoma-derived growth factor | P51859 | Hdgf | 26 kDa | 2 C | 1.098 | 0.84238 |
| ATP synthase subunit alpha, mitochondrial | Q03265 | Atp5a1 | 60 kDa | 2 C | 1.091 | 0.64912 |
| Macrophage-capping protein | P24452 | Capg | 39 kDa | 5 C | 1.077 | 0.90157 |
| Triadin | E9Q9K5 | Trdn | 78 kDa | 3 C | 1.075 | 0.93862 |
| 26S proteasome non-ATPase regulatory subunit 9 | Q9CR00 | Psmd9 | 25 kDa | 3 C | 1.067 | 0.77107 |
| Delta-aminolevulinic acid dehydratase | P10518 | Alad | 36 kDa | 8 C | 1.067 | 0.7746 |
| Isoform 2 of Cytosol aminopeptidase | Q9CPY7-2 | Lap3 | 53 kDa | 7 C | 1.054 | 0.79648 |
| Adiponectin | Q60994 | Adipoq | 27 kDa | 2 C | 1.053 | 0.71556 |

| | | | | | | |
|---|---|---|---|---|---|---|
| Nucleolar protein 3 | Q9D1X0 | Nol3 | 25 kDa | 4 C | 1.049 | 0.91917 |
| Transaldolase | Q93092 | Taldo1 | 37 kDa | 3 C | 1.040 | 0.79797 |
| Transgelin-2 | Q9WVA4 | Tagln2 | 22 kDa | 3 C | 1.040 | 0.91958 |
| Cardiomyopathy-associated protein 5 | Q70KF4 | Cmya5 | 413 kDa | 26 C | 1.040 | 0.92095 |
| O-acetyl-ADP-ribose deacetylase MACROD1 | Q922B1 | Macrod1 | 35 kDa | 10 C | 1.029 | 0.91958 |
| Thioredoxin-dependent peroxide reductase, mitochondrial | P20108 | Prdx3 | 28 kDa | 4 C | 1.022 | 0.91327 |
| Desmoplakin | E9Q557 | Dsp | 333 kDa | 43 C | 1.017 | 0.98516 |
| Polyubiquitin-C | P0CG50 | Ubc | 83 kDa | 6 C | 1.000 | 1 |
| Proteasome subunit beta type-6 | Q60692 | Psmb6 | 25 kDa | 4 C | 1.000 | 1 |
| Isoform 3 of Heterogeneous nuclear ribonucleoprotein D0 | Q60668-3 | Hnrnpd | 33 kDa | 3 C | 1.000 | #DIV/0! |
| Synaptopodin-2 | Q91YE8 | Synpo2 | 117 kDa | 12 C | 0.998 | 0.99768 |
| Glyoxylate reductase/hydroxypyruvate reductase | Q91Z53 | Grhpr | 35 kDa | 7 C | 0.995 | 0.99201 |
| Isoform Cytoplasmic of Glutathione reductase, mitochondrial | P47791-2 | Gsr | 51 kDa | 11 C | 0.978 | 0.94074 |
| Myc box-dependent-interacting protein 1 | O08539 | Bin1 | 64 kDa | 4 C | 0.960 | 0.81532 |
| NEDD8-conjugating enzyme Ubc12 | P61082 | Ube2m | 21 kDa | 5 C | 0.945 | 0.83978 |
| Selenide, water dikinase 1 | Q8BH69 | Sephs1 | 43 kDa | 9 C | 0.923 | 0.66466 |
| Hemoglobin subunit alpha | P01942 | Hba | 15 kDa | 1 C | 0.900 | 0.79797 |
| Serine-threonine kinase receptor-associated protein | Q9Z1Z2 | Strap | 38 kDa | 6 C | 0.899 | 0.8354 |
| Protein disulfide-isomerase A3 | P27773 | Pdia3 | 57 kDa | 8 C | 0.884 | 0.68071 |
| Persulfide dioxygenase ETHE1, mitochondrial | Q9DCM0 | Ethe1 | 28 kDa | 9 C | 0.880 | 0.40708 |
| Phosphotriesterase-related protein | Q60866 | Pter | 39 kDa | 6 C | 0.880 | 0.56303 |
| Nucleoside diphosphate kinase B | Q01768 | Nme2 | 17 kDa | 2 C | 0.867 | 0.67719 |
| Sarcalumenin | Q7TQ48 | Srl | 99 kDa | 7 C | 0.850 | 0.88033 |
| Nexilin | Q7TPW1 | Nexn | 72 kDa | 5 C | 0.840 | 0.70126 |
| Annexin A1 | P10107 | Anxa1 | 39 kDa | 5 C | 0.835 | 0.87862 |
| Isoform 3 of Drebrin-like protein | Q62418-3 | Dbnl | 48 kDa | 5 C | 0.832 | 0.7035 |
| Nascent polypeptide-associated complex subunit alpha, muscle-specific form | P70670 | Naca | 220 kDa | 16 C | 0.830 | 0.42669 |
| Plasminogen activator inhibitor 1 RNA-binding protein | Q9CY58 (+1) | Serbp1 | 45 kDa | 2 C | 0.820 | 0.73528 |
| Methionine adenosyltransferase 2 subunit beta | Q99LB6 | Mat2b | 37 kDa | 7 C | 0.816 | 0.719 |
| Apoptotic chromatin condensation inducer in the nucleus | Q9JIX8 (+1) | Acin1 | 151 kDa | 7 C | 0.816 | 0.719 |
| Immunoglobulin J chain | P01592 | Jchain | 18 kDa | 8 C | 0.806 | 0.59537 |
| Lamina-associated polypeptide 2, isoforms alpha/zeta | Q61033 | Tmpo | 75 kDa | 1 C | 0.800 | 0.48246 |
| Growth factor receptor-bound protein 2 | Q60631 | Grb2 | 25 kDa | 2 C | 0.800 | 0.53892 |
| Endoplasmic reticulum resident protein 29 | P57759 | Erp29 | 29 kDa | 1 C | 0.800 | 0.66908 |
| Regulator of microtubule dynamics protein 1 | Q9DCV4 | Rmdn1 | 35 kDa | 5 C | 0.800 | 0.70094 |
| Transforming growth factor beta-2 | P27090 | Tgfb2 | 48 kDa | 15 C | 0.800 | 0.81985 |
| Peptidyl-prolyl cis-trans isomerase D | Q9CR16 | Ppid | 41 kDa | 7 C | 0.800 | 0.81985 |
| Methylcrotonoyl-CoA carboxylase beta chain, mitochondrial | Q3ULD5 | Mccc2 | 61 kDa | 10 C | 0.800 | 0.21922 |
| Tubulin-folding cofactor B | Q9D1E6 | Tbcb | 27 kDa | 5 C | 0.800 | 0.40708 |
| Desmoglein-1-alpha | Q61495 (+1) | Dsg1a | 115 kDa | 18 C | 0.800 | 0.74337 |
| Protein phosphatase 1 regulatory subunit 12B | Q8BG95 | Ppp1r12b | 109 kDa | 6 C | 0.780 | 0.35062 |
| BAG family molecular chaperone regulator 3 | Q9JLV1 | Bag3 | 62 kDa | 4 C | 0.740 | 0.51737 |
| Apoptosis-inducing factor 1, mitochondrial | Q9Z0X1 | Aifm1 | 67 kDa | 4 C | 0.733 | 0.26556 |
| Junction plakoglobin | Q02257 | Jup | 82 kDa | 13 C | 0.711 | 0.59288 |
| Inositol monophosphatase 1 | O55023 | Impa1 | 30 kDa | 6 C | 0.686 | 0.2542 |
| Actin, aortic smooth muscle | P62737 (+1) | Acta2 | 42 kDa | 7 C | 0.686 | 0.04574 |
| Synaptopodin | Q8CC35 | Synpo | 100 kDa | 5 C | 0.682 | 0.57028 |
| Hydroxyacyl-coenzyme A dehydrogenase, mitochondrial | Q61425 | Hadh | 34 kDa | 5 C | 0.680 | 0.64142 |
| Actin, cytoplasmic 1 | P60710 | Actb | 42 kDa | 6 C | 0.672 | 0.07949 |
| Prelamin-A/C | P48678 | Lmna | 74 kDa | 5 C | 0.669 | 0.5585 |
| Haptoglobin | Q61646 | Hp | 39 kDa | 9 C | 0.667 | 0.08922 |
| Microtubule-associated protein 4 | P27546 (+2) | Map4 | 117 kDa | 10 C | 0.656 | 0.1946 |
| Myosin-6 | Q02566 | Myh6 | 224 kDa | 14 C | 0.649 | 0.02786 |
| Cdc42-interacting protein 4 | Q8CJ53 (+1) | Trip10 | 68 kDa | 4 C | 0.643 | 0.34602 |
| Proteasome subunit beta type-1 | O09061 | Psmb1 | 26 kDa | 5 C | 0.631 | 0.21188 |
| Protein disulfide-isomerase A4 | P08003 | Pdia4 | 72 kDa | 6 C | 0.608 | 0.30168 |
| 5-phosphohydroxy-L-lysine phospho-lyase | Q8R1K4 (+1) | Phykpl | 52 kDa | 7 C | 0.575 | 0.49263 |
| Isoform 2 of Prostaglandin reductase 2 | Q8VDQ1-2 | Ptgr2 | 34 kDa | 8 C | 0.560 | 0.20626 |
| Aldose reductase | P45376 | Akr1b1 | 36 kDa | 6 C | 0.551 | 0.34752 |
| Ig gamma-1 chain C region secreted form | P01868 (+1) | Ighg1 | 36 kDa | 12 C | 0.449 | 0.22315 |
| Ran-specific GTPase-activating protein | P34022 | Ranbp1 | 24 kDa | 3 C | 0.438 | 0.313 |

| | | | | | | |
|---|---|---|---|---|---|---|
| Sorbin and SH3 domain-containing protein 1 | Q62417 | Sorbs1 | 143 kDa | 4 C | 0.350 | 0.28281 |
| Microtubule-associated protein tau | P10637 | Mapt | 76 kDa | 2 C | 0.349 | 0.05671 |
| 3-ketoacyl-CoA thiolase, mitochondrial | Q8BWT1 | Acaa2 | 42 kDa | 8 C | 0.320 | 0.08069 |
| Protein phosphatase 1 regulatory subunit 3A | Q99MR9 | Ppp1r3a | 121 kDa | 19 C | 0.273 | 0.1029 |
| Gelsolin | P13020 | Gsn | 86 kDa | 7 C | 0.211 | 0.37175 |
| 5'-nucleotidase domain-containing protein 3 | Q3UHB1 | Nt5dc3 | 63 kDa | 6 C | 0.174 | 0.29235 |
| CapZ-interacting protein | Q3UZA1 (+1) | Rcsd1 | 44 kDa | 5 C | 0.125 | 0.12176 |
| Proteasome subunit beta type-8 | P28063 | Psmb8 | 30 kDa | 5 C | 0.125 | 0.12176 |
