## Supplemental Tabe 4 for "Dietary restriction transforms the protein sulfhydrome in a tissue-specific and cystathionine γ-lyase-dependent manner"

| Supplemental Table 4: Dietary Impact on the Brain Sulfhydrome | | | | | | |
|---|---|---|---|---|---|---|
| Protein Name | Accession Number | Alternate ID | Molecular Weight | Cysteine Residues | DR/AL Spectral Count Ratio | P-value |
| Pyruvate kinase PKM | P52480 | Pkm | 58 kDa | 9 C | 530.000 | 6.33735E-06 |
| Far upstream element-binding protein 2 | Q3U0V1 | Khsrp | 77 kDa | 8 C | 98.000 | 0.0265567 |
| Cytosolic non-specific dipeptidase | Q9D1A2 | Cndp2 | 53 kDa | 8 C | 76.200 | 0.01272141 |
| Catalase | P24270 | Cat | 60 kDa | 5 C | 60.000 | 0.001635444 |
| Epsin-1 | Q80VP1 (+1) | Epn1 | 60 kDa | 2 C | 50.000 | 0.014633577 |
| Neuronal pentraxin-1 | Q62443 | Nptx1 | 47 kDa | 9 C | 50.000 | 0.024708148 |
| 3-hydroxyisobutyrate dehydrogenase, mitochondrial | Q99L13 | Hibadh | 35 kDa | 12 C | 46.000 | 0.017809078 |
| Alpha-soluble NSF attachment protein | Q9DB05 | Napa | 33 kDa | 8 C | 44.000 | 0.000823367 |
| Isoform 3 of Neuronal cell adhesion molecule | Q810U4-3 | Nrcam | 139 kDa | 15 C | 44.000 | 0.003354847 |
| S-phase kinase-associated protein 1 | Q9WTX5 | Skp1 | 19 kDa | 3 C | 40.000 | 0.004510464 |
| Hydroxymethylglutaryl-CoA synthase, cytoplasmic | Q11136 | Hmgcs1 | 58 kDa | 11 C | 38.000 | 0.000819238 |
| Xaa-Pro dipeptidase | Q8JZK9 | Pepd | 55 kDa | 17 C | 38.000 | 0.00171501 |
| Plasma membrane calcium-transporting ATPase 1 | G5E829 | Atp2b1 | 135 kDa | 17 C | 36.200 | 0.018139944 |
| Inorganic pyrophosphatase 2, mitochondrial | Q9CQI3 | Ppa2 | 38 kDa | 8 C | 36.000 | 0.001332063 |
| Glia maturation factor beta | Q91VM9 | Gmfb | 17 kDa | 3 C | 36.000 | 0.00263263 |
| Aspartyl aminopeptidase | Q9Z2W0 | Dnpep | 52 kDa | 10 C | 36.000 | 0.00263263 |
| Heterogeneous nuclear ribonucleoprotein L | Q8R081 | Hnrnpl | 64 kDa | 11 C | 34.000 | 6.79657E-06 |
| Aspartoacylase | Q8R3P0 | Aspa | 35 kDa | 8 C | 34.000 | 0.003606068 |
| Hsp90 co-chaperone Cdc37 | Q61081 | Cdc37 | 45 kDa | 9 C | 34.000 | 0.008947867 |
| Neuroplastin | Q8C0M9 | Nptn | 44 kDa | 4 C | 32.000 | 0.000161817 |
| Isoaspartyl peptidase/L-asparaginase | Q9Z0E0 | Asrgl1 | 34 kDa | 8 C | 32.000 | 0.002237977 |
| Peptidyl-prolyl cis-trans isomerase D | Q9CR16 | Ppid | 41 kDa | 7 C | 32.000 | 0.004450765 |
| Neurochondrin | P97300 | Ncdn | 79 kDa | 25 C | 32.000 | 0.015530712 |
| TOM1-like protein 2 | Q5SRX1 | Tom1l2 | 56 kDa | 6 C | 30.000 | 8.5013E-05 |
| Copine-4 | Q8BLR2 | Cpne4 | 62 kDa | 15 C | 30.000 | 8.5013E-05 |
| Fibrinogen beta chain | Q8K0E8 | Fgb | 55 kDa | 12 C | 30.000 | 0.00490883 |
| Syntaxin-1A | O35526 | Stx1a | 33 kDa | 3 C | 30.000 | 0.01840437 |
| Isocitrate dehydrogenase [NADP] cytoplasmic | O88844 | Idh1 | 47 kDa | 7 C | 28.364 | 3.80852E-05 |
| Isoform 3 of Calcium/calmodulin-dependent protein kinase type II subunit gamma | Q923T9-3 | Camk2g | 56 kDa | 12 C | 28.200 | 0.040318335 |
| GRIP1-associated protein 1 | Q8VD04 | Gripap1 | 93 kDa | 5 C | 28.200 | 0.048589874 |
| Dynactin subunit 2 | Q9QUR7 | Dctn2 | 44 kDa | 2 C | 28.000 | 0.001833986 |
| Peptidyl-prolyl cis-trans isomerase NIMA-interacting 1 | Q99KJ8 | Pin1 | 18 kDa | 2 C | 28.000 | 0.00466796 |
| Glucose-6-phosphate isomerase | P06745 | Gpi | 63 kDa | 4 C | 27.692 | 4.41634E-05 |
| Vesicle-associated membrane protein 2 | P63044 | Vamp2 | 13 kDa | 1 C | 26.000 | 0.000895113 |
| Protein FAM49B | Q921M7 | Fam49b | 37 kDa | 5 C | 26.000 | 0.003463377 |
| Proline-rich transmembrane protein 2 | E9PUL5 | Prrt2 | 36 kDa | 3 C | 26.000 | 0.01404195 |
| Stress-induced-phosphoprotein 1 | Q60864 | Stip1 | 63 kDa | 11 C | 25.000 | 0.002886446 |
| 182 kDa tankyrase-1-binding protein | P58871 | Tnks1bp1 | 182 kDa | 23 C | 24.000 | 0.001445441 |
| Isoform 2 of Mitochondrial import receptor subunit TOM34 | Q9CYG7-2 | Tomm34 | 34 kDa | 7 C | 24.000 | 0.005337243 |
| Pyridoxal kinase | Q8K183 | Pdxk | 35 kDa | 5 C | 22.500 | 0.002973951 |
| Hsc70-interacting protein | P54728 | St13 | 42 kDa | 3 C | 22.200 | 0.012880809 |
| Cytosolic purine 5'-nucleotidase | Q3V1L4 | Nt5c2 | 65 kDa | 8 C | 22.200 | 0.012880809 |
| Serine--tRNA ligase, cytoplasmic | Q99L47 | Sars | 58 kDa | 9 C | 22.200 | 0.023300765 |
| Pyridoxal phosphate phosphatase | P26638 | Pdxp | 32 kDa | 7 C | 22.200 | 0.049058816 |
| UV excision repair protein RAD23 homolog B | P60487 | Rad23b | 44 kDa | 1 C | 22.200 | 0.049058816 |
| Dual specificity protein phosphatase 3 | Q8BTV2 (+1) | Dusp3 | 20 kDa | 4 C | 22.000 | 0.001656725 |
| Cysteine-rich protein 2 | Q9DCT8 | Crip2 | 23 kDa | 14 C | 22.000 | 0.001656725 |
| Cleavage and polyadenylation specificity factor subunit 7 | Q9D7X3 | Cpsf7 | 52 kDa | 3 C | 22.000 | 0.006888389 |
| Beta-centractin | Q8R5C5 | Actr1b | 42 kDa | 2 C | 22.000 | 0.015572842 |
| Radixin | P26043 | Rdx | 69 kDa | 1 C | 22.000 | 0.039793779 |
| Ubiquitin carboxyl-terminal hydrolase 5 | P56399 | Usp5 | 96 kDa | 16 C | 20.923 | 0.002614655 |
| Poly(rC)-binding protein 2 | Q61990 (+2) | Pcbp2 | 38 kDa | 7 C | 20.200 | 0.015113676 |
| Sulfite oxidase, mitochondrial | Q8R086 | Suox | 61 kDa | 9 C | 20.200 | 0.044560936 |
| Leucine-rich glioma-inactivated protein 1 | Q9JIA1 | Lgi1 | 64 kDa | 13 C | 20.200 | 0.044560936 |
| Matrin-3 | Q8K310 | Matr3 | 95 kDa | 9 C | 20.000 | 0.001124775 |
| Ezrin | P26040 | Ezr | 69 kDa | 2 C | 20.000 | 0.001124775 |
| Fascin | Q61553 | Fscn1 | 55 kDa | 13 C | 20.000 | 0.018341734 |
| ATP synthase subunit d, mitochondrial | Q9DCX2 | Atp5h | 19 kDa | 1 C | 20.000 | 0.018341734 |
| Cytochrome c1, heme protein, mitochondrial | Q9JJK2 | Cyc1 | 35 kDa | 14 C | 18.200 | 0.031929179 |
| Leukocyte surface antigen CD47 | Q61735 (+1) | Cd47 | 33 kDa | 11 C | 18.000 | 0.000137733 |

| | | | | | | |
|---|---|---|---|---|---|---|
| 60S ribosomal protein L12 | P35979 | Rpl12 | 18 kDa | 3 C | 18.000 | 0.005144296 |
| Neuronal pentraxin receptor | Q99J85 | Nptxr | 52 kDa | 8 C | 18.000 | 0.005144296 |
| Arf-GAP with GTPase, ANK repeat and PH domain-containing protein 3 | Q8VHH5 (+1) | Agap3 | 98 kDa | 5 C | 18.000 | 0.005144296 |
| Isoform 3 of Protein RUFY3 | Q9D394-3 | Rufy3 | 55 kDa | 5 C | 18.000 | 0.018314417 |
| Synaptosomal-associated protein 25 | P60879 | Snap25 | 23 kDa | 4 C | 17.846 | 0.001749834 |
| NEDD8-conjugating enzyme Ubc12 | O08677 | Ube2m | 21 kDa | 5 C | 16.200 | 0.009627925 |
| Heterogeneous nuclear ribonucleoproteins C1/C2 | P61082 | Hnrnpc | 34 kDa | 1 C | 16.200 | 0.009627925 |
| Paralemmin-2 | Q8BR92 | Palm2 | 42 kDa | 4 C | 16.200 | 0.009627925 |
| Platelet-activating factor acetylhydrolase IB subunit alpha | P63005 | Pafah1b1 | 47 kDa | 10 C | 16.200 | 0.030130552 |
| Protein Shroom2 | Q9Z204 | Shroom2 | 165 kDa | 24 C | 16.200 | 0.030130552 |
| Alcohol dehydrogenase [NADP(+)] | A2ALU4 | Akr1a1 | 37 kDa | 4 C | 16.200 | 0.030130552 |
| Kininogen-1 | Q3UYC0 | Kng1 | 73 kDa | 19 C | 16.200 | 0.030130552 |
| Protein phosphatase 1H | Q9JII6 | Ppm1h | 56 kDa | 9 C | 16.200 | 0.030130552 |
| COP9 signalosome complex subunit 5 | Q8CBW3 | Cops5 | 38 kDa | 4 C | 16.000 | 0.001007816 |
| Cytoplasmic protein NCK2 | O55033 | Nck2 | 43 kDa | 4 C | 16.000 | 0.001007816 |
| Abl interactor 1 | Q9Z218 | Abi1 | 52 kDa | 2 C | 16.000 | 0.01299165 |
| DnaJ homolog subfamily B member 11 | Q99KV1 | Dnajb11 | 41 kDa | 5 C | 16.000 | 0.01299165 |
| DCN1-like protein 1 | Q9QZ73 | Dcun1d1 | 30 kDa | 4 C | 16.000 | 0.01299165 |
| Dipeptidyl aminopeptidase-like protein 6 | Q9DAR7 | Dpp6 | 91 kDa | 12 C | 16.000 | 0.01299165 |
| m7GpppX diphosphatase | O35864 | Dcps | 39 kDa | 2 C | 16.000 | 0.01299165 |
| TAR DNA-binding protein 43 | Q921F2 | Tardbp | 45 kDa | 7 C | 16.000 | 0.01299165 |
| Dihydropteridine reductase | Q8BVI4 | Qdpr | 26 kDa | 4 C | 15.385 | 0.000410159 |
| Isoform 3 of 2-oxoglutarate dehydrogenase, mitochondrial | Q60597-3 | Ogdh | 118 kDa | 21 C | 15.273 | 0.004779282 |
| Proteasome subunit beta type-1 | O09061 | Psmb1 | 26 kDa | 5 C | 14.769 | 2.87334E-05 |
| 3'(2'),5'-bisphosphate nucleotidase 1 | Q9Z0S1 | Bpnt1 | 33 kDa | 6 C | 14.200 | 0.018780516 |
| Interleukin enhancer-binding factor 2 | Q9CXY6 | Ilf2 | 43 kDa | 4 C | 14.200 | 0.018780516 |
| Isoform 10 of Disintegrin and metalloproteinase domain-containing protein 22 | Q9R1V6-12 | Adam22 | 108 kDa | 47 C | 14.154 | 0.000212204 |
| Ubiquitin-conjugating enzyme E2 variant 2 | Q9D2M8 | Ube2v2 | 16 kDa | 1 C | 14.000 | 0.002259411 |
| Carbonic anhydrase 1 | P68404 (+1) | Ca1 | 28 kDa | 1 C | 14.000 | 0.002259411 |
| Cell cycle exit and neuronal differentiation protein 1 | Q9JKC6 | Cend1 | 15 kDa | 1 C | 14.000 | 0.002259411 |
| Fucose mutarotase | Q8R2K1 | Fuom | 17 kDa | 3 C | 14.000 | 0.024121664 |
| Protein kinase C beta type | P13634 | Prkcb | 77 kDa | 20 C | 14.000 | 0.024121664 |
| UMP-CMP kinase | Q9DBP5 | Cmpk1 | 22 kDa | 6 C | 13.538 | 6.52249E-06 |
| Serine/threonine-protein phosphatase 5 | Q60676 | Ppp5c | 57 kDa | 11 C | 13.455 | 0.004734457 |
| Vinculin | Q64727 | Vcl | 117 kDa | 10 C | 13.091 | 0.000424124 |
| Transketolase | P40142 | Tkt | 68 kDa | 12 C | 12.571 | 0.000507922 |
| Sorbin and SH3 domain-containing protein 1 | Q9D0F9 | Sorbs1 | 143 kDa | 4 C | 12.308 | 0.001674392 |
| Phosphoglucomutase-1 | Q62417 | Pgm1 | 61 kDa | 10 C | 12.308 | 0.006781791 |
| Inositol-3-phosphate synthase 1 | P46660 | Isyna1 | 61 kDa | 11 C | 12.308 | 0.006781791 |
| Cell adhesion molecule 3 | Q60710 | Cadm3 | 43 kDa | 7 C | 12.200 | 0.028236713 |
| Catenin delta-2 | Q99N28 | Ctnnd2 | 135 kDa | 19 C | 12.200 | 0.028236713 |
| Isoform 2 of Low molecular weight phosphotyrosine protein phosphatase | Q9WVQ5 | Acp1 | 18 kDa | 8 C | 12.200 | 0.028236713 |
| Protein NipSnap homolog 3B | Q9DAK9 | Nipsnap3b | 28 kDa | 2 C | 12.200 | 0.028236713 |
| Deoxynucleoside triphosphate triphosphohydrolase SAMHD1 | Q9CQE1 | Samhd1 | 73 kDa | 17 C | 12.200 | 0.028236713 |
| Isoform 4 of Serine/threonine-protein kinase LMTK1 | O35927 | Aatk | 128 kDa | 30 C | 12.200 | 0.028236713 |
| Prostaglandin E synthase 3 | P97797-2 | Ptges3 | 19 kDa | 5 C | 12.200 | 0.028236713 |
| Isoform 2 of Tyrosine-protein phosphatase non-receptor type substrate 1 | Q6NVF9 | Sirpa | 56 kDa | 10 C | 12.200 | 0.028236713 |
| Solute carrier family 2, facilitated glucose transporter member 1 | P17809 | Slc2a1 | 54 kDa | 6 C | 12.200 | 0.028236713 |
| Vesicle-trafficking protein SEC22b | O08547 | Sec22b | 25 kDa | 3 C | 12.200 | 0.028236713 |
| Ubiquitin-associated protein 2-like | Q80X50 (+2) | Ubap2l | 117 kDa | 2 C | 12.000 | 0.001855162 |
| Serine/threonine-protein phosphatase 2A activator | P58389 | Ptpa | 37 kDa | 5 C | 12.000 | 0.001855162 |
| Signal transducing adapter molecule 1 | P70297 | Stam | 60 kDa | 7 C | 12.000 | 0.001855162 |
| Copine-6 | Q9Z140 | Cpne6 | 62 kDa | 15 C | 11.754 | 0.027452445 |
| SH3 and multiple ankyrin repeat domains protein 2 | Q80Z38 (+1) | Shank2 | 159 kDa | 9 C | 11.692 | 0.000603211 |
| DAZ-associated protein 1 | Q9JII5 (+1) | Dazap1 | 43 kDa | 4 C | 11.692 | 0.017055117 |
| Isoform 2 of Glial fibrillary acidic protein | P03995-2 | Gfap | 49 kDa | 1 C | 11.138 | 0.025425941 |
| AP-2 complex subunit alpha-2 | P17427 | Ap2a2 | 104 kDa | 16 C | 11.130 | 0.044085505 |
| Apolipoprotein E | P08226 | Apoe | 36 kDa | 1 C | 10.200 | 0.030703132 |
| Ras-related protein Rab-11B | P46638 | Rab11b | 24 kDa | 2 C | 10.200 | 0.030703132 |
| Coiled-coil domain-containing protein 6 | D3YZP9 | Ccdc6 | 53 kDa | 3 C | 10.200 | 0.030703132 |
| Charged multivesicular body protein 5 | Q9D7S9 | Chmp5 | 25 kDa | 1 C | 10.200 | 0.030703132 |
| Multifunctional protein ADE2 | Q9DCL9 | Paics | 47 kDa | 13 C | 10.200 | 0.030703132 |
| Glutamate decarboxylase 1 | P48318 | Gad1 | 67 kDa | 13 C | 10.200 | 0.030703132 |
| F-box only protein 2 | Q80UW2 | Fbxo2 | 34 kDa | 5 C | 10.200 | 0.030703132 |

| | | | | | | |
|---|---|---|---|---|---|---|
| MAGUK p55 subfamily member 2 | Q9WV34 (+1) | Mpp2 | 62 kDa | 5 C | 10.200 | 0.030703132 |
| Prefoldin subunit 3 | P61759 | Vbp1 | 22 kDa | 3 C | 10.200 | 0.030703132 |
| Glutaredoxin-related protein 5, mitochondrial | Q80Y14 | Glrx5 | 16 kDa | 2 C | 10.200 | 0.030703132 |
| Rab GDP dissociation inhibitor beta | Q61598 (+1) | Gdi2 | 51 kDa | 9 C | 10.087 | 0.042720211 |
| ATP-dependent 6-phosphofructokinase, platelet type | Q9WUA3 | Pfkp | 85 kDa | 20 C | 9.292 | 0.04073788 |
| Heterogeneous nuclear ribonucleoprotein Q | Q7TMK9 | Syncrip | 70 kDa | 4 C | 9.231 | 0.0045479 |
| GMP reductase 1 | Q9DCZ1 | Gmpr | 37 kDa | 9 C | 9.231 | 0.008785733 |
| Ubiquitin carboxyl-terminal hydrolase isozyme L1 | Q9R0P9 | Uchl1 | 25 kDa | 6 C | 8.941 | 0.002005364 |
| Septin-4 | P28661 (+1) | 43347 | 55 kDa | 8 C | 8.727 | 3.32802E-06 |
| Secernin-1 | Q9CZC8 | Scrn1 | 46 kDa | 11 C | 8.727 | 0.000502701 |
| Thioredoxin reductase 1, cytoplasmic | Q9JMH6 | Txnrd1 | 67 kDa | 21 C | 8.615 | 0.008897492 |
| Heterogeneous nuclear ribonucleoprotein U | P97765 | Hnrnpu | 88 kDa | 14 C | 8.615 | 7.6145E-05 |
| WW domain-binding protein 2 | Q61838 | Wbp2 | 28 kDa | 2 C | 8.615 | 0.001137978 |
| Pregnancy zone protein | Q8VEK3 (+1) | Pzp | 166 kDa | 24 C | 8.615 | 0.008897492 |
| Dihydropyrimidinase-related protein 5 | Q9EQF6 | Dpysl5 | 62 kDa | 10 C | 8.364 | 0.00148195 |
| Spectrin alpha chain, non-erythrocytic 1 | P16546 | Sptan1 | 285 kDa | 14 C | 8.333 | 0.001099083 |
| Heterogeneous nuclear ribonucleoprotein A3 | Q8BG05 | Hnrnpa3 | 40 kDa | 4 C | 8.157 | 0.001731548 |
| Serine/threonine-protein kinase PAK 1 | O88643 | Pak1 | 61 kDa | 5 C | 8.000 | 0.000447895 |
| V-type proton ATPase subunit E 1 | Q11011 | Atp6v1e1 | 26 kDa | 1 C | 8.000 | 0.002488969 |
| T-complex protein 1 subunit delta | P50518 | Cct4 | 58 kDa | 9 C | 8.000 | 0.046450948 |
| Puromycin-sensitive aminopeptidase | P80315 | Npepps | 103 kDa | 12 C | 8.000 | 0.046450948 |
| Septin-6 | Q9R1T4 (+2) | 43349 | 50 kDa | 8 C | 7.686 | 0.00021835 |
| Isochorismatase domain-containing protein 2A | P85094 | Isoc2a | 22 kDa | 6 C | 7.673 | 0.034959728 |
| Ermin | Q5EBJ4 | Ermn | 32 kDa | 3 C | 7.500 | 0.00038844 |
| Centrosomal protein of 170 kDa | Q6A065 | Cep170 | 175 kDa | 10 C | 7.500 | 0.000656704 |
| Clathrin coat assembly protein AP180 | Q61548 (+2) | Snap91 | 92 kDa | 5 C | 7.415 | 0.001001648 |
| Eukaryotic translation initiation factor 5A-1 | P63242 | Eif5a | 17 kDa | 4 C | 7.415 | 0.00127294 |
| Ornithine aminotransferase, mitochondrial | P29758 | Oat | 48 kDa | 7 C | 7.385 | 0.048762123 |
| Glutathione S-transferase P 1 | P19157 | Gstp1 | 24 kDa | 3 C | 7.373 | 0.002108034 |
| Polyadenylate-binding protein 1 | P29341 | Pabpc1 | 71 kDa | 4 C | 7.273 | 0.021818262 |
| Neurofascin | Q810U3 | Nfasc | 138 kDa | 14 C | 7.250 | 0.001482823 |
| Beta-soluble NSF attachment protein | P28663 | Napb | 34 kDa | 6 C | 7.250 | 0.008225165 |
| Oxidation resistance protein 1 | Q4KMM3 | Oxr1 | 96 kDa | 8 C | 7.211 | 2.25316E-05 |
| 4-aminobutyrate aminotransferase, mitochondrial | P61922 | Abat | 56 kDa | 12 C | 7.000 | 0.026460902 |
| Serine/threonine-protein phosphatase PP1-gamma catalytic subunit | P63087 (+1) | Ppp1cc | 37 kDa | 13 C | 6.982 | 0.000550944 |
| Proteasome subunit beta type-7 | Q91W90 | Psmb7 | 30 kDa | 6 C | 6.831 | 0.041309888 |
| Microtubule-associated protein RP/EB family member 1 | Q9CQR6 | Mapre1 | 30 kDa | 3 C | 6.769 | 0.015487704 |
| Serine/threonine-protein phosphatase 6 catalytic subunit | Q61166 | Ppp6c | 35 kDa | 12 C | 6.769 | 0.029889918 |
| Glycerol-3-phosphate dehydrogenase, mitochondrial | Q6RHR9 | Gpd2 | 81 kDa | 8 C | 6.769 | 0.005149524 |
| Glucosidase 2 subunit beta | P14231 | Prkcsh | 59 kDa | 17 C | 6.769 | 0.005149524 |
| Membrane-associated guanylate kinase, WW and PDZ domain-containing protein 1 | Q64521 | Magi1 | 162 kDa | 16 C | 6.769 | 0.015487704 |
| Sodium/potassium-transporting ATPase subunit beta-2 | O08795 (+1) | Atp1b2 | 33 kDa | 7 C | 6.769 | 0.029889918 |
| Tubulin beta-6 chain | Q922F4 | Tubb6 | 50 kDa | 8 C | 6.687 | 0.046263575 |
| Serine/threonine-protein phosphatase 2A catalytic subunit alpha isoform | P63330 | Ppp2ca | 36 kDa | 10 C | 6.609 | 0.00443087 |
| Isoform Tau-A of Microtubule-associated protein tau | P10637-2 | Mapt | 45 kDa | 2 C | 6.601 | 0.000649637 |
| Ubiquitin-like modifier-activating enzyme 1 | Q02053 | Uba1 | 118 kDa | 21 C | 6.557 | 0.0109152 |
| Tropomodulin-2 | Q9JKK7 | Tmod2 | 40 kDa | 1 C | 6.545 | 0.025240612 |
| Septin-5 | Q61644 | 43348 | 43 kDa | 8 C | 6.275 | 0.001055301 |
| Protein kinase C and casein kinase substrate in neurons protein 1 | Q9Z2Q6 | Pacsin1 | 51 kDa | 6 C | 6.275 | 0.002747831 |
| Actin-related protein 2/3 complex subunit 1A | Q9R0Q6 | Arpc1a | 42 kDa | 10 C | 6.261 | 0.003827219 |
| Epsin-2 | P42208 | Epn2 | 63 kDa | 3 C | 6.250 | 0.000328709 |
| Septin-2 | Q8CHU3 | 43345 | 42 kDa | 8 C | 6.250 | 0.008047106 |
| Nucleolin | P09405 | Ncl | 77 kDa | 1 C | 6.244 | 0.011532038 |
| Proteasome subunit beta type-3 | Q9R1P1 | Psmb3 | 23 kDa | 5 C | 6.182 | 0.002528366 |
| Splicing factor, proline- and glutamine-rich | Q8VIJ6 | Sfpq | 75 kDa | 7 C | 6.182 | 0.040229743 |
| Destrin | Q9R0P5 | Dstn | 19 kDa | 6 C | 6.154 | 6.29127E-05 |
| Transcription elongation factor A protein-like 5 | Q8CCT4 | Tceal5 | 22 kDa | 1 C | 6.154 | 6.29127E-05 |
| Actin-related protein 2/3 complex subunit 5-like protein | Q9D898 | Arpc5l | 17 kDa | 1 C | 6.154 | 6.29127E-05 |
| Translationally-controlled tumor protein | P63028 | Tpt1 | 19 kDa | 2 C | 6.154 | 0.017873283 |
| A-kinase anchor protein 12 | Q9WTQ5 | Akap12 | 181 kDa | 10 C | 6.057 | 0.007028895 |
| Heat shock 70 kDa protein 4L | P48722 | Hspa4l | 94 kDa | 15 C | 6.000 | 0.001585229 |
| Phospholipid hydroperoxide glutathione peroxidase, mitochondrial | O70325 | Gpx4 | 22 kDa | 10 C | 6.000 | 0.002204732 |
| Fibrinogen alpha chain | E9PV24 | Fga | 87 kDa | 13 C | 5.913 | 0.002531249 |
| Calbindin | P12658 | Calb1 | 30 kDa | 4 C | 5.818 | 0.033163632 |

| | | | | | | |
|---|---|---|---|---|---|---|
| Actin-related protein 3 | Q99JY9 | Actr3 | 47 kDa | 8 C | 5.770 | 0.006282391 |
| Peroxiredoxin-6 | O08709 | Prdx6 | 25 kDa | 2 C | 5.689 | 0.00034855 |
| Small nuclear ribonucleoprotein Sm D2 | P62317 | Snrpd2 | 14 kDa | 2 C | 5.538 | 0.016118365 |
| Ester hydrolase C11orf54 homolog | Q91V76 | | 35 kDa | 8 C | 5.538 | 0.001746044 |
| Coactosin-like protein | Q9CQI6 | Cotl1 | 16 kDa | 2 C | 5.538 | 0.016118365 |
| COP9 signalosome complex subunit 3 | O88543 | Cops3 | 48 kDa | 10 C | 5.538 | 0.016118365 |
| Elongation factor Tu, mitochondrial | Q8BFR5 | Tufm | 50 kDa | 7 C | 5.538 | 0.040889938 |
| Spectrin beta chain, non-erythrocytic 1 | Q62261 | Sptbn1 | 274 kDa | 15 C | 5.500 | 0.004251782 |
| Isoform 2 of CUGBP Elav-like family member 2 | Q9Z0H4-2 (+1 | Celf2 | 55 kDa | 7 C | 5.500 | 0.008881104 |
| Cytosolic 10-formyltetrahydrofolate dehydrogenase | Q8R0Y6 | Aldh1l1 | 99 kDa | 15 C | 5.490 | 0.04370562 |
| Gephyrin | Q9CRB6 | Gphn | 83 kDa | 13 C | 5.250 | 0.003131209 |
| Tubulin polymerization-promoting protein family member 3 | Q8BUV3 | Tppp3 | 19 kDa | 3 C | 5.250 | 0.007833144 |
| Protein NDRG2 | Q9QYG0 (+1) | Ndrg2 | 41 kDa | 6 C | 5.250 | 0.007833144 |
| Guanine nucleotide-binding protein G(I)/G(S)/G(T) subunit beta-1 | P62874 | Gnb1 | 37 kDa | 14 C | 5.171 | 0.007508644 |
| Aldehyde dehydrogenase, mitochondrial | P47738 | Aldh2 | 57 kDa | 9 C | 5.143 | 0.000544963 |
| Glutamate dehydrogenase 1, mitochondrial | P26443 | Glud1 | 61 kDa | 6 C | 5.032 | 0.01731047 |
| A-kinase anchor protein 5 | D3YVF0 | Akap5 | 79 kDa | 2 C | 4.959 | 0.000307934 |
| Neural cell adhesion molecule 1 | P13595 | Ncam1 | 119 kDa | 14 C | 4.952 | 0.012890279 |
| Proteasome subunit beta type-2 | O08585 | Psmb2 | 23 kDa | 3 C | 4.923 | 0.007259302 |
| Far upstream element-binding protein 1 | Q91WJ8 | Fubp1 | 69 kDa | 3 C | 4.904 | 0.043466436 |
| Serine-threonine kinase receptor-associated protein | Q9Z1Z2 | Strap | 38 kDa | 6 C | 4.903 | 0.018860224 |
| Glutathione peroxidase 1 | P11352 | Gpx1 | 22 kDa | 4 C | 4.900 | 0.009256267 |
| 14-3-3 protein theta | P68254 | Ywhaq | 28 kDa | 5 C | 4.867 | 0.000761708 |
| Microtubule-associated protein 4 | P27546 (+1) | Map4 | 117 kDa | 10 C | 4.848 | 0.027345193 |
| Serine/threonine-protein phosphatase PP1-alpha catalytic subunit | P62137 | Ppp1ca | 38 kDa | 13 C | 4.800 | 0.000639764 |
| Endophilin-A1 | Q62420 | Sh3gl2 | 40 kDa | 3 C | 4.800 | 0.002551294 |
| Endoplasmic reticulum resident protein 29 | P57759 | Erp29 | 29 kDa | 1 C | 4.762 | 0.003922479 |
| Serine/threonine-protein phosphatase PP1-beta catalytic subunit | P62141 | Ppp1cb | 37 kDa | 14 C | 4.716 | 0.000310245 |
| Protein DJ-1 | Q99LX0 | Park7 | 20 kDa | 4 C | 4.683 | 0.013490819 |
| Alpha-actinin-4 | P57780 | Actn4 | 105 kDa | 8 C | 4.667 | 0.010836891 |
| Calreticulin | P14211 | Calr | 48 kDa | 6 C | 4.620 | 0.000129545 |
| Isoform 2 of Ankyrin-2 | Q8C8R3-2 | Ank2 | 429 kDa | 19 C | 4.565 | 0.001094258 |
| Tubulin alpha-1C chain | P68373 | Tuba1c | 50 kDa | 12 C | 4.474 | 0.006304732 |
| Dihydropyrimidinase-related protein 4 | O35098 | Dpysl4 | 62 kDa | 10 C | 4.373 | 0.001582172 |
| Heat shock protein 105 kDa | Q8CJ19 | Hsph1 | 96 kDa | 17 C | 4.364 | 0.001329065 |
| [F-actin]-methionine sulfoxide oxidase MICAL3 | P49722 | Mical3 | 224 kDa | 21 C | 4.364 | 0.008246549 |
| Rab GDP dissociation inhibitor alpha | P50396 | Gdi1 | 51 kDa | 10 C | 4.348 | 0.027537603 |
| Mitogen-activated protein kinase 1 | P63085 | Mapk1 | 41 kDa | 7 C | 4.308 | 0.016070239 |
| MAGUK p55 subfamily member 6 | Q99MN9 | Mpp6 | 63 kDa | 5 C | 4.308 | 0.016070239 |
| Propionyl-CoA carboxylase beta chain, mitochondrial | P68037 | Pccb | 58 kDa | 11 C | 4.308 | 0.016070239 |
| Brain-specific angiogenesis inhibitor 1-associated protein 2 | Q8BKX1 (+2) | Baiap2 | 59 kDa | 4 C | 4.308 | 0.016070239 |
| Syntaxin-1B | P61264 | Stx1b | 33 kDa | 4 C | 4.267 | 0.000337226 |
| Isoform 2 of Disks large homolog 4 | Q62108-2 | Dlg4 | 85 kDa | 6 C | 4.250 | 0.001052638 |
| Sodium/potassium-transporting ATPase subunit alpha-3 | Q6PIC6 | Atp1a3 | 112 kDa | 25 C | 4.235 | 0.017776997 |
| Drebrin-like protein | Q62418 (+2) | Dbnl | 49 kDa | 5 C | 4.229 | 1.00155E-05 |
| Calretinin | Q08331 | Calb2 | 31 kDa | 2 C | 4.200 | 0.010537241 |
| A-kinase anchor protein 2 | O54931 (+1) | Akap2 | 99 kDa | 9 C | 4.174 | 0.037569028 |
| Immunoglobulin superfamily member 8 | Q8R366 | Igsf8 | 65 kDa | 11 C | 4.174 | 0.037569028 |
| Phosphoglycerate mutase 1 | Q9DBJ1 | Pgam1 | 29 kDa | 2 C | 4.160 | 0.005086699 |
| V-type proton ATPase catalytic subunit A | P50516 | Atp6v1a | 68 kDa | 6 C | 4.129 | 0.011406756 |
| NAD-dependent protein deacetylase sirtuin-2 | Q8VDQ8 | Sirt2 | 43 kDa | 11 C | 4.121 | 0.027961543 |
| Contactin-1 | P12960 | Cntn1 | 113 kDa | 15 C | 4.099 | 0.002279311 |
| ATP-dependent 6-phosphofructokinase, muscle type | Q64010 | Pfkm | 85 kDa | 15 C | 4.098 | 0.039557146 |
| 14-3-3 protein eta | P68510 | Ywhah | 28 kDa | 3 C | 4.028 | 0.000679054 |
| Guanine nucleotide-binding protein G(I)/G(S)/G(T) subunit beta-2 | P62880 | Gnb2 | 37 kDa | 13 C | 4.000 | 0.00930365 |
| AP-2 complex subunit mu | P84091 | Ap2m1 | 50 kDa | 6 C | 4.000 | 0.025740696 |
| Carbonic anhydrase 2 | P00920 | Ca2 | 29 kDa | 2 C | 3.951 | 0.001447652 |
| Tubulin beta-3 chain | Q9ERD7 | Tubb3 | 50 kDa | 8 C | 3.930 | 0.019562549 |
| Sodium/potassium-transporting ATPase subunit alpha-2 | Q6PIE5 | Atp1a2 | 112 kDa | 23 C | 3.926 | 0.028936299 |
| Creatine kinase B-type | Q04447 | Ckb | 43 kDa | 5 C | 3.925 | 0.00216121 |
| Tenascin-R | Q8BYI9 | Tnr | 150 kDa | 41 C | 3.920 | 0.003353242 |
| Isoform 4 of Disks large homolog 2 | Q91XM9-4 | Dlg2 | 98 kDa | 8 C | 3.913 | 0.007970009 |
| Synapsin-1 | O88935 | Syn1 | 74 kDa | 3 C | 3.907 | 0.016283602 |
| 2',3'-cyclic-nucleotide 3'-phosphodiesterase | P16330 | Cnp | 47 kDa | 7 C | 3.883 | 0.024031581 |

| | | | | | | |
|---|---|---|---|---|---|---|
| RNA-binding protein FUS | P56959 | Fus | 53 kDa | 4 C | 3.871 | 0.01122771 |
| Beta-arrestin-1 | Q8BWG8 | Arrb1 | 47 kDa | 8 C | 3.867 | 0.000989766 |
| Tubulin alpha-4A chain | P68368 | Tuba4a | 50 kDa | 13 C | 3.852 | 0.008550958 |
| Serine/threonine-protein phosphatase 2B catalytic subunit alpha isoform | P63328 | Ppp3ca | 59 kDa | 12 C | 3.831 | 0.007994056 |
| Retinal dehydrogenase 1 | P24549 | Aldh1a1 | 54 kDa | 11 C | 3.829 | 0.000702329 |
| Ketimine reductase mu-crystallin | O54983 | Crym | 34 kDa | 3 C | 3.826 | 0.029297102 |
| Isocitrate dehydrogenase [NAD] subunit gamma 1, mitochondrial | P70404 | Idh3g | 43 kDa | 7 C | 3.826 | 0.029297102 |
| Nucleolysin TIAR | P70318 | Tial1 | 43 kDa | 6 C | 3.826 | 0.029297102 |
| Reticulon-3 | Q9ES97 | Rtn3 | 104 kDa | 11 C | 3.753 | 0.009602527 |
| Microtubule-associated protein 1B | P14873 | Map1b | 270 kDa | 22 C | 3.728 | 3.3724E-05 |
| Adenylyl cyclase-associated protein 1 | P40124 | Cap1 | 52 kDa | 6 C | 3.707 | 0.031281003 |
| Isoform A of Cytosolic acyl coenzyme A thioester hydrolase | Q91V12-2 | Acot7 | 38 kDa | 8 C | 3.692 | 0.000740884 |
| Neurofilament medium polypeptide | P08553 | Nefm | 96 kDa | 2 C | 3.692 | 0.015885968 |
| Protein phosphatase 1 regulatory subunit 1B | Q60829 | Ppp1r1b | 22 kDa | 2 C | 3.692 | 0.02269657 |
| S-adenosylmethionine synthase isoform type-2 | Q3THS6 | Mat2a | 44 kDa | 6 C | 3.692 | 0.02269657 |
| Latexin | P70202 | Lxn | 25 kDa | 3 C | 3.692 | 0.02269657 |
| Tubulin beta-4A chain | Q9D6F9 | Tubb4a | 50 kDa | 8 C | 3.638 | 0.022225584 |
| Serotransferrin | Q921I1 | Tf | 77 kDa | 38 C | 3.621 | 0.002344105 |
| Tubulin beta-5 chain | P99024 | Tubb5 | 50 kDa | 8 C | 3.616 | 0.009519619 |
| Acetyl-CoA acetyltransferase, mitochondrial | Q8QZT1 | Acat1 | 45 kDa | 6 C | 3.600 | 0.010313311 |
| Tubulin beta-2A chain | Q7TMM9 | Tubb2a | 50 kDa | 7 C | 3.579 | 0.013295382 |
| Cofilin-1 | P18760 | Cfl1 | 19 kDa | 4 C | 3.576 | 0.001390721 |
| Isoform 2 of Neutral alpha-glucosidase AB | Q8BHN3-2 | Ganab | 109 kDa | 8 C | 3.538 | 0.021393807 |
| Isoform 7 of SH3-containing GRB2-like protein 3-interacting protein 1 | Q8VD37-8 | Sgip1 | 89 kDa | 8 C | 3.520 | 0.002280958 |
| Diphosphoinositol polyphosphate phosphohydrolase 1 | Q9JI46 | Nudt3 | 19 kDa | 4 C | 3.500 | 0.003278774 |
| Elongation factor 1-beta | O70251 | Eef1b | 25 kDa | 3 C | 3.500 | 0.035830179 |
| Creatine kinase U-type, mitochondrial | P30275 | Ckmt1 | 47 kDa | 7 C | 3.497 | 0.003328425 |
| Protein piccolo | Q9QYX7 | Pclo | 551 kDa | 33 C | 3.451 | 0.003385892 |
| Transitional endoplasmic reticulum ATPase | Q01853 | Vcp | 89 kDa | 12 C | 3.434 | 0.001590723 |
| 14-3-3 protein beta/alpha | Q9CQV8 | Ywhab | 28 kDa | 2 C | 3.433 | 0.001100437 |
| Crk-like protein | P47941 | Crkl | 34 kDa | 2 C | 3.410 | 0.023995575 |
| Tubulin beta-4B chain | P68372 | Tubb4b | 50 kDa | 8 C | 3.380 | 0.019355425 |
| Reticulon-4 | Q99P72 | Rtn4 | 127 kDa | 9 C | 3.280 | 0.004469541 |
| 14-3-3 protein epsilon | P62259 | Ywhae | 29 kDa | 3 C | 3.278 | 0.001569101 |
| Heat shock 70 kDa protein 4 | Q61316 | Hspa4 | 94 kDa | 14 C | 3.270 | 0.001311471 |
| Isoform M1 of Pyruvate kinase PKM | P52480-2 | Pkm | 58 kDa | 9 C | 3.265 | 9.92085E-05 |
| Rho GDP-dissociation inhibitor 1 | Q99PT1 | Arhgdia | 23 kDa | 1 C | 3.259 | 0.003151828 |
| Phosphoglycerate kinase 1 | P09411 | Pgk1 | 45 kDa | 7 C | 3.257 | 0.017763322 |
| D-3-phosphoglycerate dehydrogenase | Q61753 | Phgdh | 57 kDa | 14 C | 3.250 | 0.037274381 |
| ATP synthase subunit O, mitochondrial | Q9DB20 | Atp5o | 23 kDa | 1 C | 3.238 | 0.025809632 |
| Reticulon-1 | Q8K0T0 | Rtn1 | 84 kDa | 6 C | 3.231 | 0.027726814 |
| Brevican core protein | Q61361 | Bcan | 96 kDa | 27 C | 3.226 | 0.028336401 |
| Dihydrolipoyllysine-residue succinyltransferase component of 2-oxoglutarate dehydrogena | Q9D2G2 | Dlst | 49 kDa | 6 C | 3.179 | 4.88558E-05 |
| Dihydropyrimidinase-related protein 2 | O08553 | Dpysl2 | 62 kDa | 7 C | 3.145 | 0.000175952 |
| Synaptojanin-1 | Q8CHC4 | Synj1 | 173 kDa | 19 C | 3.137 | 0.001459719 |
| Peptidyl-prolyl cis-trans isomerase A | P17742 | Ppia | 18 kDa | 3 C | 3.127 | 0.015041791 |
| Microtubule-associated protein 2 | P20357 | Map2 | 199 kDa | 7 C | 3.103 | 0.00014762 |
| Drebrin | Q9QXS6 | Dbn1 | 77 kDa | 13 C | 3.100 | 0.010417675 |
| Actin-related protein 2/3 complex subunit 4 | P59999 | Arpc4 | 20 kDa | 4 C | 3.097 | 0.002042503 |
| Actin-related protein 2 | P61161 | Actr2 | 45 kDa | 5 C | 3.086 | 0.011476134 |
| Myristoylated alanine-rich C-kinase substrate | P26645 | Marcks | 30 kDa | 1 C | 3.067 | 0.009986912 |
| Heterogeneous nuclear ribonucleoprotein K | P61979 (+1) | Hnrnpk | 51 kDa | 5 C | 3.059 | 0.001868464 |
| Delta-aminolevulinic acid dehydratase | P10518 | Alad | 36 kDa | 8 C | 3.048 | 0.022211907 |
| Versican core protein | Q62059 | Vcan | 367 kDa | 36 C | 3.040 | 8.56523E-05 |
| Succinate-semialdehyde dehydrogenase, mitochondrial | Q8BWF0 | Aldh5a1 | 56 kDa | 10 C | 3.017 | 0.000261892 |
| Septin-7 | O55131 | 43350 | 51 kDa | 6 C | 2.963 | 0.001976333 |
| Heterogeneous nuclear ribonucleoprotein H | O35737 | Hnrnph1 | 49 kDa | 1 C | 2.960 | 0.000515366 |
| Mitotic checkpoint protein BUB3 | Q9WVA3 | Bub3 | 37 kDa | 7 C | 2.927 | 0.005243098 |
| Phosphatidylethanolamine-binding protein 1 | P70296 | Pebp1 | 21 kDa | 3 C | 2.900 | 0.009453117 |
| 14-3-3 protein zeta/delta | P63101 | Ywhaz | 28 kDa | 3 C | 2.889 | 0.000276194 |
| Protein PRRC2A | Q7TSC1 | Prrc2a | 229 kDa | 1 C | 2.857 | 0.014113719 |
| Serine/threonine-protein phosphatase 2A 55 kDa regulatory subunit B alpha isoform | Q6P1F6 | Ppp2r2a | 52 kDa | 9 C | 2.844 | 0.000274863 |
| Ras-related protein Rab-3A | P63011 | Rab3a | 25 kDa | 4 C | 2.839 | 0.024864943 |
| Disks large homolog 3 | P70175 | Dlg3 | 93 kDa | 10 C | 2.829 | 0.019429949 |

| | | | | | | | |
|---|---|---|---|---|---|---|---|
| L-lactate dehydrogenase A chain | P06151 | Ldha | 36 kDa | 6 C | | 2.813 | 0.003056449 |
| 14-3-3 protein gamma | P61982 | Ywhag | 28 kDa | 3 C | | 2.787 | 1.72385E-05 |
| Heat shock protein HSP 90-beta | P11499 | Hsp90ab1 | 83 kDa | 6 C | | 2.786 | 0.040441745 |
| F-actin-capping protein subunit beta | P34022 | Capzb | 31 kDa | 5 C | | 2.750 | 0.046941612 |
| Guanine deaminase | Q9R111 | Gda | 51 kDa | 9 C | | 2.750 | 0.046941612 |
| Protein disulfide-isomerase A6 | Q922R8 | Pdia6 | 48 kDa | 7 C | | 2.750 | 0.046941612 |
| Platelet-activating factor acetylhydrolase IB subunit beta | Q9Z0P4 | Pafah1b2 | 26 kDa | 3 C | | 2.732 | 0.003415227 |
| Paralemmin-1 | Q61206 | Palm | 42 kDa | 3 C | | 2.732 | 0.013836984 |
| Glutathione S-transferase Mu 1 | P10649 | Gstm1 | 26 kDa | 2 C | | 2.711 | 0.043154896 |
| Isoform Cytoplasmic of Glutathione reductase, mitochondrial | P47791-2 | Gsr | 51 kDa | 11 C | | 2.691 | 0.008337877 |
| CAP-Gly domain-containing linker protein 2 | Q9Z0H8 | Clip2 | 116 kDa | 9 C | | 2.667 | 0.011726692 |
| Protein bassoon | O88737 | Bsn | 419 kDa | 35 C | | 2.639 | 0.009776436 |
| Thioredoxin-like protein 1 | Q8CDN6 | Txnl1 | 32 kDa | 7 C | | 2.629 | 0.032793293 |
| 40S ribosomal protein SA | P14206 | Rpsa | 33 kDa | 2 C | | 2.623 | 0.028987402 |
| Profilin-2 | Q80TK0 | Pfn2 | 15 kDa | 6 C | | 2.600 | 0.009627925 |
| Uncharacterized protein KIAA1107 | Q9JJV2 | Kiaa1107 | 149 kDa | 23 C | | 2.600 | 0.02780078 |
| AP2-associated protein kinase 1 | Q3UHJ0 | Aak1 | 103 kDa | 13 C | | 2.596 | 0.025626851 |
| Insulin-degrading enzyme | Q9JHR7 | Ide | 118 kDa | 13 C | | 2.588 | 0.045420776 |
| Heat shock cognate 71 kDa protein | P63017 | Hspa8 | 71 kDa | 4 C | | 2.583 | 0.001532018 |
| Sodium/potassium-transporting ATPase subunit alpha-1 | Q8VDN2 | Atp1a1 | 113 kDa | 23 C | | 2.557 | 0.03920612 |
| WD repeat-containing protein 44 | Q6NVE8 | Wdr44 | 102 kDa | 10 C | | 2.545 | 0.049999391 |
| Hypoxanthine-guanine phosphoribosyltransferase | P00493 | Hprt1 | 25 kDa | 4 C | | 2.533 | 0.033386415 |
| Fructose-bisphosphate aldolase C | P05063 | Aldoc | 39 kDa | 7 C | | 2.517 | 0.006032647 |
| Protein disulfide-isomerase | P09103 | P4hb | 57 kDa | 7 C | | 2.514 | 0.000329988 |
| Fatty acid-binding protein, epidermal | Q05816 | Fabp5 | 15 kDa | 6 C | | 2.510 | 0.026198745 |
| Septin-8 | Q8CHH9 (+1) | 43351 | 50 kDa | 6 C | | 2.507 | 0.034056087 |
| Isoform 1B of Synaptogyrin-1 | O55100-2 | Syngr1 | 21 kDa | 4 C | | 2.500 | 0.019453132 |
| Band 4.1-like protein 3 | Q9WV92 | Epb41l3 | 103 kDa | 8 C | | 2.496 | 0.023650201 |
| Isoform 3 of Heterogeneous nuclear ribonucleoprotein D0 | Q60668-3 | Hnrnpd | 33 kDa | 3 C | | 2.494 | 0.006157187 |
| Proteasome subunit beta type-4 | P99026 | Psmb4 | 29 kDa | 2 C | | 2.492 | 0.033179331 |
| Serine protease inhibitor A3K | P07759 | Serpina3k | 47 kDa | 4 C | | 2.479 | 0.018878715 |
| Dihydropyrimidinase-related protein 1 | P97427 | Crmp1 | 62 kDa | 6 C | | 2.456 | 0.003941207 |
| Proteasome subunit alpha type-3 | O70435 | Psma3 | 28 kDa | 4 C | | 2.400 | 0.027238682 |
| Glutamine synthetase | P15105 | Glul | 42 kDa | 13 C | | 2.400 | 0.012477816 |
| Profilin-1 | P62962 | Pfn1 | 15 kDa | 3 C | | 2.400 | 0.035769575 |
| Malate dehydrogenase, mitochondrial | P08249 | Mdh2 | 36 kDa | 8 C | | 2.365 | 0.000147951 |
| Tubulin-folding cofactor B | Q6P9K8 (+1) | Tbcb | 27 kDa | 5 C | | 2.361 | 0.049486 |
| Serum albumin | P07724 | Alb | 69 kDa | 36 C | | 2.356 | 0.001339586 |
| Heterogeneous nuclear ribonucleoprotein A/B | Q99020 | Hnrnpab | 31 kDa | 2 C | | 2.343 | 0.004210672 |
| Na(+)/H(+) exchange regulatory cofactor NHE-RF1 | P70441 | Slc9a3r1 | 39 kDa | 5 C | | 2.300 | 0.018217288 |
| Gelsolin | P13020 | Gsn | 86 kDa | 7 C | | 2.255 | 0.035819656 |
| Proteasome subunit alpha type-4 | Q9R1P0 | Psma4 | 29 kDa | 5 C | | 2.254 | 0.019209488 |
| Aconitate hydratase, mitochondrial | Q99KI0 | Aco2 | 85 kDa | 13 C | | 2.225 | 0.00270688 |
| Proteasome subunit alpha type-5 | Q99MI1-2 | Psma5 | 26 kDa | 3 C | | 2.196 | 0.034874556 |
| LIM and SH3 domain protein 1 | Q61792 | Lasp1 | 30 kDa | 7 C | | 2.182 | 0.012461121 |
| Enoyl-CoA hydratase, mitochondrial | Q8BH95 | Echs1 | 31 kDa | 7 C | | 2.171 | 0.010955099 |
| Superoxide dismutase [Mn], mitochondrial | P09671 | Sod2 | 25 kDa | 4 C | | 2.133 | 0.008067765 |
| Isoform 2 of Septin-11 | Q8C1B7-2 (+1 | 43354 | 49 kDa | 6 C | | 2.116 | 0.00132971 |
| Peroxiredoxin-5, mitochondrial | P99029 | Prdx5 | 22 kDa | 6 C | | 2.116 | 0.005788574 |
| Isoform 2 of Tropomyosin alpha-3 chain | P21107-2 | Tpm3 | 29 kDa | 1 C | | 2.080 | 0.014245753 |
| Protein phosphatase 1 regulatory subunit 7 | Q3UM45 | Ppp1r7 | 41 kDa | 2 C | | 2.055 | 0.004747769 |
| Cofilin-2 | P45591 | Cfl2 | 19 kDa | 2 C | | 2.022 | 0.030498309 |
| Isoform 2 of Microtubule-associated protein 1A | Q9QYR6-2 | Map1a | 325 kDa | 24 C | | 2.004 | 0.002423615 |
| Isoform 3 of Dynamin-1 | P39053-3 | Dnm1 | 96 kDa | 7 C | | 78.600 | 0.210157817 |
| Plasma membrane calcium-transporting ATPase 2 | Q9R0K7 | Atp2b2 | 133 kDa | 21 C | | 58.000 | 0.05427114 |
| AP-1 complex subunit beta-1 | O35643 | Ap1b1 | 104 kDa | 15 C | | 36.400 | 0.113894147 |
| Gamma-soluble NSF attachment protein | Q9CWZ7 | Napg | 35 kDa | 6 C | | 34.000 | 0.05260414 |
| Vacuolar protein sorting-associated protein 35 | Q9EQH3 | Vps35 | 92 kDa | 24 C | | 26.200 | 0.105887376 |
| Cullin-associated NEDD8-dissociated protein 1 | Q6ZQ38 | Cand1 | 136 kDa | 30 C | | 24.000 | 0.077189785 |
| Importin subunit beta-1 | P70168 | Kpnb1 | 97 kDa | 22 C | | 22.400 | 0.100598557 |
| T-complex protein 1 subunit beta | P80314 | Cct2 | 57 kDa | 6 C | | 22.400 | 0.154296542 |
| T-complex protein 1 subunit eta | P80313 | Cct7 | 60 kDa | 9 C | | 20.400 | 0.171369386 |
| Alpha-centractin | P61164 | Actr1a | 43 kDa | 2 C | | 20.200 | 0.096788477 |
| Protein farnesyltransferase/geranylgeranyltransferase type-1 subunit alpha | Q61239 | Fnta | 44 kDa | 3 C | | 20.200 | 0.147323995 |

| | | | | | | |
|---|---|---|---|---|---|---|
| Tetraspanin-2 | Q922J6 | Tspan2 | 24 kDa | 11 C | 20.000 | #DIV/0! |
| Acetyl-CoA acetyltransferase, cytosolic | Q8CAY6 | Acat2 | 41 kDa | 8 C | 18.400 | 0.089027282 |
| ATP-citrate synthase | Q91V92 | Acly | 120 kDa | 16 C | 18.400 | 0.089027282 |
| Isoform 2 of Solute carrier family 12 member 5 | Q91V14-2 | Slc12a5 | 124 kDa | 26 C | 18.400 | 0.130866563 |
| Synaptosomal-associated protein 47 | Q9D0M3 | Snap47 | 47 kDa | 4 C | 18.200 | 0.052307671 |
| LanC-like protein 2 | Q8R570 | Lancl2 | 51 kDa | 13 C | 18.200 | 0.117483358 |
| Ubiquitin carboxyl-terminal hydrolase isozyme L3 | Q9JKB1 | Uchl3 | 26 kDa | 3 C | 16.400 | 0.103977532 |
| Isoform 2 of Dynamin-like 120 kDa protein, mitochondrial | P58281-2 | Opa1 | 116 kDa | 9 C | 16.400 | 0.130436227 |
| Isoform 3 of 1-phosphatidylinositol 4,5-bisphosphate phosphodiesterase eta-2 | A2AP18-3 | Plch2 | 138 kDa | 29 C | 16.400 | 0.130436227 |
| Protein SGT1 homolog | Q9CX34 | Sugt1 | 38 kDa | 5 C | 16.200 | 0.084257374 |
| Phosphatidylinositol transfer protein alpha isoform | P53810 | Pitpna | 32 kDa | 4 C | 14.400 | 0.079631514 |
| Isoform 3 of Elongation factor 1-delta | P57776-3 | Eef1d | 73 kDa | 2 C | 14.400 | 0.079631514 |
| Visinin-like protein 1 | P62761 | Vsnl1 | 22 kDa | 3 C | 14.400 | 0.115431007 |
| Cell adhesion molecule 2 | Q8BLQ9 (+2) | Cadm2 | 48 kDa | 8 C | 14.400 | 0.149411093 |
| Amyloid beta A4 precursor protein-binding family A member 1 | B2RUJ5 (+1) | Apba1 | 93 kDa | 8 C | 14.200 | 0.05144906 |
| Haloacid dehalogenase-like hydrolase domain-containing protein 2 | Q3UGR5 | Hdhd2 | 29 kDa | 3 C | 14.200 | 0.05144906 |
| Dynamin-1-like protein | Q8K1M6 | Dnm1l | 83 kDa | 9 C | 13.600 | 0.092814765 |
| Elongation factor 2 | P58252 | Eef2 | 95 kDa | 7 C | 12.923 | 0.066196142 |
| CaM kinase-like vesicle-associated protein | Q3UHL1 | Camkv | 55 kDa | 8 C | 12.800 | 0.407083822 |
| Isoform 2 of Calcium-dependent secretion activator 1 | Q80TJ1-2 | Cadps | 154 kDa | 19 C | 12.600 | 0.193006088 |
| Glutaminase kidney isoform, mitochondrial | D3Z7P3 | Gls | 74 kDa | 15 C | 12.600 | 0.193006088 |
| V-type proton ATPase subunit H | Q8BVE3 | Atp6v1h | 56 kDa | 8 C | 12.600 | 0.229730754 |
| Alanine--tRNA ligase, cytoplasmic | Q8BGQ7 | Aars | 107 kDa | 15 C | 12.600 | 0.229730754 |
| Target of Myb protein 1 | O88746 | Tom1 | 54 kDa | 4 C | 12.400 | 0.067583294 |
| Isoform 2 of Guanine nucleotide-binding protein subunit beta-5 | P62881-2 | Gnb5 | 39 kDa | 19 C | 12.400 | 0.117123764 |
| Hippocalcin-like protein 1 | P62748 | Hpcal1 | 22 kDa | 2 C | 12.400 | 0.117123764 |
| BTB/POZ domain-containing protein KCTD12 | Q6WVG3 | Kctd12 | 36 kDa | 5 C | 12.400 | 0.117123764 |
| Coronin-1C | Q9WUM4 | Coro1c | 53 kDa | 12 C | 12.400 | 0.117123764 |
| 60S ribosomal protein L6 | P47911 | Rpl6 | 34 kDa | 1 C | 12.400 | 0.117123764 |
| cAMP-dependent protein kinase type II-alpha regulatory subunit | P12367 | Prkar2a | 45 kDa | 6 C | 12.400 | 0.067583294 |
| Scaffold attachment factor B1 | D3YXK2 | Safb | 105 kDa | 9 C | 12.400 | 0.117123764 |
| Protein arginine N-methyltransferase 5 | Q8CIG8 | Prmt5 | 73 kDa | 12 C | 12.400 | 0.117123764 |
| Cell adhesion molecule 4 | Q8R464 | Cadm4 | 43 kDa | 8 C | 12.400 | 0.117123764 |
| Hydroxyacylglutathione hydrolase-like protein | Q9DB32 | Haghl | 31 kDa | 7 C | 12.400 | 0.117123764 |
| Neurofilament light polypeptide | P08551 | Nefl | 62 kDa | 1 C | 12.369 | 0.096810667 |
| Alpha-internexin | Q9JHU9 | Ina | 55 kDa | 3 C | 12.308 | 0.066599131 |
| 14 kDa phosphohistidine phosphatase | Q9I Z31 (+1) | Phpt1 | 14 kDa | 3 C | 12.200 | 0.077582781 |
| Methylthioribulose-1-phosphate dehydratase | Q80YE4-4 | Apip | 27 kDa | 11 C | 12.200 | 0.077582781 |
| Polypyrimidine tract-binding protein 2 | Q9R0Q7 | Ptbp2 | 57 kDa | 2 C | 12.200 | 0.077582781 |
| Cleavage and polyadenylation specificity factor subunit 6 | Q9CYH2 | Cpsf6 | 59 kDa | 3 C | 12.200 | 0.077582781 |
| Redox-regulatory protein FAM213A | Q9D358-2 | Fam213a | 24 kDa | 2 C | 12.200 | 0.077582781 |
| Glucose 1,6-bisphosphate synthase | O35295 | Pgm2l1 | 70 kDa | 16 C | 10.600 | 0.206912274 |
| Sideroflexin-1 | Q99JR1 | Sfxn1 | 36 kDa | 5 C | 10.600 | 0.206912274 |
| Transcriptional activator protein Pur-beta | Q8CAA7 | Purb | 34 kDa | 2 C | 10.600 | 0.299341419 |
| AP-2 complex subunit beta | Q9DBG3 (+1) | Ap2b1 | 105 kDa | 16 C | 10.435 | 0.057328573 |
| Isoform 2 of SRC kinase signaling inhibitor 1 | Q9QWI6-2 | Srcin1 | 131 kDa | 5 C | 10.400 | 0.092068749 |
| 26S proteasome non-ATPase regulatory subunit 4 | O35226 | Psmd4 | 41 kDa | 4 C | 10.400 | 0.092068749 |
| Isoform APP695 of Amyloid beta A4 protein | P12023-2 | App | 78 kDa | 18 C | 10.400 | 0.092068749 |
| Alanine aminotransferase 1 | Q8QZR5 | Gpt | 55 kDa | 14 C | 10.400 | 0.092068749 |
| Synaptic vesicle glycoprotein 2B | Q8BG39 | Sv2b | 77 kDa | 14 C | 10.400 | 0.092068749 |
| Mitochondrial fission 1 protein | Q9CQ92 | Fis1 | 17 kDa | 1 C | 10.400 | 0.092068749 |
| ADP-ribosylation factor-binding protein GGA1 | Q8R0H9 | Gga1 | 70 kDa | 6 C | 10.400 | 0.092068749 |
| Asparagine--tRNA ligase, cytoplasmic | Q8BP47 | Nars | 64 kDa | 16 C | 10.400 | 0.092068749 |
| Actin-related protein 2/3 complex subunit 2 | Q9CVB6 | Arpc2 | 34 kDa | 2 C | 10.400 | 0.161564153 |
| Carboxylesterase 1C | P23953 | Ces1c | 61 kDa | 5 C | 10.400 | 0.161564153 |
| Osteoclast-stimulating factor 1 | Q62422 | Ostf1 | 24 kDa | 4 C | 10.400 | 0.161564153 |
| Poly(U)-binding-splicing factor PUF60 | Q3UEB3 | Puf60 | 60 kDa | 3 C | 10.400 | 0.092068749 |
| Polyribonucleotide nucleotidyltransferase 1, mitochondrial | Q8K1R3 | Pnpt1 | 86 kDa | 16 C | 10.400 | 0.092068749 |
| Adenylosuccinate synthetase isozyme 1 | P28650 (+1) | Adssl1 | 50 kDa | 5 C | 10.400 | 0.092068749 |
| Excitatory amino acid transporter 1 | P56564 | Slc1a3 | 60 kDa | 3 C | 10.087 | 0.077689981 |
| Clathrin heavy chain 1 | Q68FD5 | Cltc | 192 kDa | 31 C | 10.000 | 0.076128679 |
| Neuronal-specific septin-3 | Q9Z1S5 (+1) | 43346 | 40 kDa | 4 C | 9.292 | 0.066717351 |
| ADP-ribosylation factor 5 | P84084 | Arf5 | 21 kDa | 2 C | 8.800 | 0.407083822 |
| Ras-related protein Rab-3C | P62823 | Rab3c | 26 kDa | 4 C | 8.800 | 0.407083822 |

| | | | | | | |
|---|---|---|---|---|---|---|
| AP-2 complex subunit alpha-1 | P17426 (+1) | Ap2a1 | 108 kDa | 19 C | 8.696 | 0.060083898 |
| Carbonic anhydrase-related protein | P28651 | Ca8 | 33 kDa | 5 C | 8.677 | 0.057802225 |
| Sprouty-related, EVH1 domain-containing protein 1 | Q924S8 | Spred1 | 51 kDa | 23 C | 8.600 | 0.193006088 |
| Peptidyl-prolyl cis-trans isomerase FKBP3 | Q62446 | Fkbp3 | 25 kDa | 1 C | 8.600 | 0.193006088 |
| GTPase NRas | P08556 (+1) | Nras | 21 kDa | 5 C | 8.600 | 0.193006088 |
| ADP-ribosylation factor 3 | P61205 (+1) | Arf3 | 21 kDa | 1 C | 8.600 | 0.272392948 |
| Ribose-phosphate pyrophosphokinase 1 | Q9D7G0 | Prps1 | 35 kDa | 9 C | 8.600 | 0.272392948 |
| Band 4.1-like protein 2 | Q9CZW5 | Epb41l2 | 110 kDa | 9 C | 8.400 | 0.106948222 |
| Glyoxalase domain-containing protein 4 | O70318 | Glod4 | 33 kDa | 5 C | 8.400 | 0.106948222 |
| Heterogeneous nuclear ribonucleoprotein A0 | Q9CPV4 (+1) | Hnrnpa0 | 31 kDa | 3 C | 8.400 | 0.106948222 |
| Mitochondrial import receptor subunit TOM70 | Q91WC3 (+1) | Tomm70 | 68 kDa | 13 C | 8.400 | 0.106948222 |
| Long-chain-fatty-acid--CoA ligase 6 | P68040 | Acsl6 | 78 kDa | 18 C | 8.400 | 0.106948222 |
| Receptor of activated protein C kinase 1 | Q9CX86 | Rack1 | 35 kDa | 8 C | 8.400 | 0.106948222 |
| cAMP-dependent protein kinase type II-beta regulatory subunit | P31324 | Prkar2b | 46 kDa | 7 C | 8.400 | 0.106948222 |
| Cytoplasmic aconitate hydratase | P28271 | Aco1 | 98 kDa | 11 C | 8.400 | 0.106948222 |
| Isoform 2 of Rap1 GTPase-activating protein 1 | A2ALS5-2 | Rap1gap | 81 kDa | 9 C | 8.400 | 0.106948222 |
| Syntaxin-12 | Q9ER00 | Stx12 | 31 kDa | 12 C | 8.400 | 0.106948222 |
| Glycogen synthase kinase-3 beta | Q9WV60 | Gsk3b | 47 kDa | 9 C | 8.400 | 0.106948222 |
| Cellular nucleic acid-binding protein | P53996 (+2) | Cnbp | 20 kDa | 22 C | 8.400 | 0.106948222 |
| Gamma-adducin | Q9QYB5 (+1) | Add3 | 79 kDa | 6 C | 8.400 | 0.106948222 |
| Eukaryotic translation initiation factor 6 | O55135 | Eif6 | 27 kDa | 8 C | 8.400 | 0.106948222 |
| Complement C3 | Q99LI8 | C3 | 186 kDa | 27 C | 8.400 | 0.106948222 |
| Hepatocyte growth factor-regulated tyrosine kinase substrate | P01027 | Hgs | 86 kDa | 11 C | 8.400 | 0.106948222 |
| MARCKS-related protein | P28667 | Marcksl1 | 20 kDa | 1 C | 8.400 | 0.106948222 |
| Alpha-2-HS-glycoprotein | P29699 | Ahsg | 37 kDa | 14 C | 8.400 | 0.106948222 |
| Excitatory amino acid transporter 2 | P43006 | Slc1a2 | 62 kDa | 9 C | 8.185 | 0.251386983 |
| Transcriptional activator protein Pur-alpha | P42669 | Pura | 35 kDa | 21 C | 7.446 | 0.116772966 |
| WASH complex subunit 2 | Q6PGL7 | Washc2 | 145 kDa | 2 C | 6.957 | 0.094149263 |
| Homer protein homolog 3 | Q99JP6 (+1) | Homer3 | 40 kDa | 1 C | 6.892 | 0.122849885 |
| Thioredoxin domain-containing protein 5 | P70195 | Txndc5 | 46 kDa | 12 C | 6.831 | 0.076664102 |
| Adenylosuccinate synthetase isozyme 2 | P46664 | Adss | 50 kDa | 7 C | 6.831 | 0.094733665 |
| Elongin-B | P63318 | Elob | 13 kDa | 1 C | 6.800 | 0.407083822 |
| Protein kinase C gamma type | P62869 | Prkcg | 78 kDa | 22 C | 6.800 | 0.407083822 |
| Synaptic vesicle membrane protein VAT-1 homolog | Q62465 | Vat1 | 43 kDa | 4 C | 6.800 | 0.407083822 |
| Annexin A6 | P14824 | Anxa6 | 76 kDa | 8 C | 6.800 | 0.407083822 |
| 1-phosphatidylinositol 4,5-bisphosphate phosphodiesterase beta-1 | Q9Z1B3 | Plcb1 | 138 kDa | 15 C | 6.800 | 0.407083822 |
| Thioredoxin | Q9DCM0 | Txn | 12 kDa | 6 C | 6.600 | 0.232236752 |
| Ras-related protein Rab-5C | P10639 | Rab5c | 23 kDa | 4 C | 6.600 | 0.232236752 |
| Persulfide dioxygenase ETHE1, mitochondrial | P28738 | Ethe1 | 28 kDa | 9 C | 6.600 | 0.232236752 |
| ATP-dependent RNA helicase DDX3Y | P21279 | Ddx3y | 73 kDa | 8 C | 6.600 | 0.232236752 |
| Methionine aminopeptidase 2 | P24547 | Metap2 | 53 kDa | 15 C | 6.600 | 0.232236752 |
| Glycine--tRNA ligase | O08663 | Gars | 82 kDa | 14 C | 6.600 | 0.232236752 |
| Kinesin heavy chain isoform 5C | Q9CZD3 | Kif5c | 109 kDa | 10 C | 6.600 | 0.232236752 |
| Guanine nucleotide-binding protein G(q) subunit alpha | Q62095 | Gnaq | 42 kDa | 5 C | 6.600 | 0.232236752 |
| Inosine-5'-monophosphate dehydrogenase 2 | Q640R3 | Impdh2 | 56 kDa | 7 C | 6.600 | 0.232236752 |
| Hepatocyte cell adhesion molecule | P23492 | Hepacam | 46 kDa | 4 C | 6.600 | 0.232236752 |
| Purine nucleoside phosphorylase | P14685 | Pnp | 32 kDa | 5 C | 6.600 | 0.232236752 |
| 26S proteasome non-ATPase regulatory subunit 3 | P35278 | Psmd3 | 61 kDa | 2 C | 6.600 | 0.232236752 |
| Inositol polyphosphate 1-phosphatase | P49442 | Inpp1 | 43 kDa | 8 C | 6.600 | 0.232236752 |
| Coatomer subunit gamma-2 | Q9QXK3 | Copg2 | 98 kDa | 21 C | 6.600 | 0.232236752 |
| Calponin-3 | Q9DAW9 | Cnn3 | 36 kDa | 3 C | 6.600 | 0.232236752 |
| Ribosome-recycling factor, mitochondrial | Q9D6S7 | Mrrf | 29 kDa | 2 C | 6.600 | 0.232236752 |
| Phosphatidate cytidylyltransferase 2 | Q99L43 | Cds2 | 51 kDa | 10 C | 6.600 | 0.232236752 |
| EH domain-containing protein 3 | Q9QXY6 | Ehd3 | 61 kDa | 1 C | 6.600 | 0.232236752 |
| N(G),N(G)-dimethylarginine dimethylaminohydrolase 2 | Q99LD8 | Ddah2 | 30 kDa | 6 C | 6.600 | 0.232236752 |
| Phosphatidylinositol 5-phosphate 4-kinase type-2 beta | Q80XI4 | Pip4k2b | 47 kDa | 7 C | 6.600 | 0.232236752 |
| Secretory carrier-associated membrane protein 1 | Q8K021 | Scamp1 | 38 kDa | 7 C | 6.600 | 0.232236752 |
| Coronin-1B | Q9WUM3 | Coro1b | 54 kDa | 11 C | 6.600 | 0.232236752 |
| Oxysterol-binding protein-related protein 1 | Q91XL9 | Osbpl1a | 108 kDa | 26 C | 6.600 | 0.232236752 |
| Isoform 2 of Paraspeckle component 1 | Q8K0S0 | Pspc1 | 53 kDa | 2 C | 6.600 | 0.232236752 |
| Phytanoyl-CoA hydroxylase-interacting protein | Q8R326-2 | Phyhip | 38 kDa | 10 C | 6.600 | 0.232236752 |
| Uncharacterized protein C1orf198 homolog | Q8C3W1 | | 35 kDa | 2 C | 6.600 | 0.232236752 |
| Eukaryotic translation initiation factor 2 subunit 1 | Q6ZWX6 | Eif2s1 | 36 kDa | 5 C | 6.600 | 0.232236752 |
| T-complex protein 1 subunit gamma | P80318 | Cct3 | 61 kDa | 10 C | 6.277 | 0.153110977 |

| | | | | | | |
|---|---|---|---|---|---|---|
| Hexokinase-1 | P17710 | Hk1 | 108 kDa | 21 C | 6.218 | 0.106382162 |
| Microtubule-associated protein RP/EB family member 3 | Q6PER3 | Mapre3 | 32 kDa | 5 C | 6.215 | 0.074790988 |
| Phosphoserine aminotransferase | Q99K85 | Psat1 | 40 kDa | 5 C | 6.215 | 0.074790988 |
| T-complex protein 1 subunit epsilon | P80316 | Cct5 | 60 kDa | 8 C | 5.600 | 0.090897218 |
| Guanine nucleotide-binding protein G(i) subunit alpha-2 | P08752 | Gnai2 | 40 kDa | 10 C | 5.565 | 0.160497706 |
| NIF3-like protein 1 | Q9EQ80 | Nif3l1 | 42 kDa | 8 C | 5.217 | 0.051851993 |
| ADP-ribosylation factor-like protein 3 | Q9WUL7 | Arl3 | 20 kDa | 3 C | 4.985 | 0.065761219 |
| Leukotriene A-4 hydrolase | P24527 | Lta4h | 69 kDa | 11 C | 4.985 | 0.065761219 |
| Twinfilin-1 | Q99PU5 | Twf1 | 40 kDa | 4 C | 4.985 | 0.065761219 |
| LanC-like protein 1 | Q91YR1 | Lancl1 | 45 kDa | 13 C | 4.985 | 0.065761219 |
| Long-chain-fatty-acid--CoA ligase ACSBG1 | P26369 | Acsbg1 | 80 kDa | 15 C | 4.985 | 0.141385212 |
| Splicing factor U2AF 65 kDa subunit | O89112 | U2af2 | 54 kDa | 6 C | 4.985 | 0.141385212 |
| Clathrin light chain A | Q9R1P3 | Clta | 26 kDa | 1 C | 4.923 | 0.114937112 |
| Glucose-6-phosphate 1-dehydrogenase X | Q00612 | G6pdx | 59 kDa | 8 C | 4.800 | 0.407083822 |
| Ig gamma-2B chain C region | P01867 (+1) | Igh-3 | 44 kDa | 13 C | 4.800 | 0.407083822 |
| Protein DEK | Q7TNV0 | Dek | 43 kDa | 4 C | 4.800 | 0.407083822 |
| Neutral amino acid transporter A | O35874 | Slc1a4 | 56 kDa | 5 C | 4.800 | 0.407083822 |
| NAD-dependent malic enzyme, mitochondrial | Q99KE1 | Me2 | 66 kDa | 12 C | 4.800 | 0.407083822 |
| Ribonuclease inhibitor | Q91VI7 | Rnh1 | 50 kDa | 30 C | 4.800 | 0.407083822 |
| Isoform 3 of Septin-9 | Q80UG5-3 | 43352 | 65 kDa | 6 C | 4.800 | 0.407083822 |
| Protein capicua homolog | Q924A2 | Cic | 258 kDa | 25 C | 4.800 | 0.407083822 |
| Cytoplasmic FMR1-interacting protein 2 | Q5SQX6 | Cyfip2 | 146 kDa | 33 C | 4.800 | 0.407083822 |
| Nck-associated protein 1 | P28660 (+1) | Nckap1 | 129 kDa | 23 C | 4.800 | 0.407083822 |
| Protein-L-isoaspartate(D-aspartate) O-methyltransferase | P23506 (+1) | Pcmt1 | 25 kDa | 2 C | 4.800 | 0.407083822 |
| Isoform 2 of ATP-dependent (S)-NAD(P)H-hydrate dehydratase | Q9CZ42-2 | Naxd | 32 kDa | 8 C | 4.800 | 0.407083822 |
| Neuroligin-2 | Q69ZK9 | Nlgn2 | 91 kDa | 10 C | 4.800 | 0.407083822 |
| 60S ribosomal protein L11 | Q9CXW4 | Rpl11 | 20 kDa | 4 C | 4.800 | 0.407083822 |
| Calcium/calmodulin-dependent protein kinase type II subunit beta | P28652 | Camk2b | 60 kDa | 13 C | 4.462 | 0.095709717 |
| Actin-related protein 2/3 complex subunit 3 | Q501J6 | Arpc3 | 21 kDa | 4 C | 4.431 | 0.14666401 |
| Probable ATP-dependent RNA helicase DDX17 | Q9JM76 | Ddx17 | 72 kDa | 11 C | 4.431 | 0.14666401 |
| Ras-related protein Rab-14 | Q91V41 | Rab14 | 24 kDa | 4 C | 4.431 | 0.324967724 |
| 40S ribosomal protein S3 | P62908 | Rps3 | 27 kDa | 3 C | 4.369 | 0.10904607 |
| Sideroflexin-3 | Q9JIS5 | Sfxn3 | 35 kDa | 5 C | 4.369 | 0.10904607 |
| ATP-dependent 6-phosphofructokinase, liver type | Q9CWS0 | Pfkl | 85 kDa | 6 C | 4.369 | 0.10904607 |
| Ras-related protein Rab-2A | Q99J77 | Rab2a | 24 kDa | 3 C | 4.369 | 0.10904607 |
| Sialic acid synthase | P12382 | Nans | 40 kDa | 8 C | 4.369 | 0.10904607 |
| Synaptic vesicle glycoprotein 2A | P53994 | Sv2a | 83 kDa | 14 C | 4.369 | 0.204650715 |
| N(G),N(G)-dimethylarginine dimethylaminohydrolase 1 | Q91V61 | Ddah1 | 31 kDa | 7 C | 4.369 | 0.204650715 |
| Proteasome subunit alpha type-2 | Q61699 | Psma2 | 26 kDa | 2 C | 4.364 | 0.1109889 |
| Ubiquitin-conjugating enzyme E2 L3 | Q9JLB0 (+1) | Ube2l3 | 18 kDa | 3 C | 4.308 | 0.066336557 |
| Amphiphysin | Q7TQF7 | Amph | 75 kDa | 2 C | 4.293 | 0.104625972 |
| Serine/threonine-protein phosphatase 2A 65 kDa regulatory subunit A alpha isoform | Q76MZ3 | Ppp2r1a | 65 kDa | 14 C | 4.275 | 0.141913359 |
| Calcium/calmodulin-dependent protein kinase type II subunit alpha | P11798 | Camk2a | 54 kDa | 10 C | 4.258 | 0.083624371 |
| Glutaredoxin-3 | Q9CQM9 | Glrx3 | 38 kDa | 5 C | 4.250 | 0.101729309 |
| Isoform 2 of Calcium/calmodulin-dependent protein kinase type II subunit delta | Q6PHZ2-2 (+1 | Camk2d | 54 kDa | 11 C | 4.243 | 0.213943758 |
| Adapter molecule crk | P47857 | Crk | 34 kDa | 1 C | 4.098 | 0.051750172 |
| Isocitrate dehydrogenase [NAD] subunit alpha, mitochondrial | Q9D6R2 | Idh3a | 40 kDa | 8 C | 4.078 | 0.117627449 |
| Selenide, water dikinase 1 | Q8BH69 | Sephs1 | 43 kDa | 9 C | 3.826 | 0.076246855 |
| WD repeat-containing protein 1 | O88342 | Wdr1 | 66 kDa | 12 C | 3.815 | 0.213028511 |
| Cysteine and glycine-rich protein 1 | P97315 | Csrp1 | 21 kDa | 15 C | 3.754 | 0.088066548 |
| ADP-ribosylation factor-binding protein GGA3 | Q9Z1N5 | Gga3 | 78 kDa | 9 C | 3.754 | 0.088066548 |
| Neuronal membrane glycoprotein M6-b | P35803 (+1) | Gpm6b | 36 kDa | 14 C | 3.754 | 0.088066548 |
| Spliceosome RNA helicase Ddx39b | Q8BMI3 | Ddx39b | 49 kDa | 8 C | 3.754 | 0.164688775 |
| Glutathione S-transferase Mu 5 | P48774 | Gstm5 | 27 kDa | 7 C | 3.673 | 0.11551585 |
| Isoform 4 of Dynamin-1 | P39053-4 | Dnm1 | 97 kDa | 7 C | 3.549 | 0.069876081 |
| Nucleolar protein 3 | Q9D1X0 | Nol3 | 25 kDa | 4 C | 3.513 | 0.088656004 |
| Syntaxin-binding protein 1 | O08599 | Stxbp1 | 68 kDa | 7 C | 3.442 | 0.054894353 |
| V-type proton ATPase subunit B, brain isoform | P62814 | Atp6v1b2 | 57 kDa | 6 C | 3.333 | 0.088305247 |
| ATP synthase subunit gamma, mitochondrial | Q91VR2 | Atp5c1 | 33 kDa | 2 C | 3.317 | 0.103200096 |
| T-complex protein 1 subunit theta | P42932 | Cct8 | 60 kDa | 10 C | 3.309 | 0.179112714 |
| Toll-interacting protein | Q9QZ06 | Tollip | 30 kDa | 4 C | 3.238 | 0.081699092 |
| Heat shock 70 kDa protein 12A | Q8K0U4 | Hspa12a | 75 kDa | 6 C | 3.200 | 0.249386756 |
| Isoform 3 of Serine/arginine repetitive matrix protein 2 | Q8BTI8-3 | Srrm2 | 285 kDa | 14 C | 3.200 | 0.212166407 |
| Catenin beta-1 | Q02248 | Ctnnb1 | 85 kDa | 11 C | 3.200 | 0.295404396 |

| | | | | | | |
|---|---|---|---|---|---|---|
| Eukaryotic translation initiation factor 3 subunit I | Q9QZD9 | Eif3i | 36 kDa | 7 C | 3.200 | 0.295404396 |
| Vesicle-fusing ATPase | P46460 | Nsf | 83 kDa | 10 C | 3.155 | 0.056249871 |
| Splicing factor 1 | Q64213 (+2) | Sf1 | 70 kDa | 4 C | 3.138 | 0.121146511 |
| Sorting nexin-1 | Q9WV80 | Snx1 | 59 kDa | 2 C | 3.138 | 0.121146511 |
| ELAV-like protein 1 | P70372 | Elavl1 | 36 kDa | 3 C | 3.138 | 0.121146511 |
| Synaptic vesicle membrane protein VAT-1 homolog-like | Q9JLM8 | Vat1l | 46 kDa | 6 C | 3.138 | 0.121146511 |
| Leukocyte elastase inhibitor A | Q80TB8 | Serpinb1a | 43 kDa | 3 C | 3.138 | 0.121146511 |
| Serine/threonine-protein kinase DCLK1 | Q9D154 | Dclk1 | 84 kDa | 1 C | 3.138 | 0.121146511 |
| Phosphate carrier protein, mitochondrial | Q8VEM8 | Slc25a3 | 40 kDa | 8 C | 3.016 | 0.075775315 |
| Guanine nucleotide-binding protein G(o) subunit alpha | P18872 | Gnao1 | 40 kDa | 9 C | 2.885 | 0.055362948 |
| Transcription elongation factor A protein 1 | P10711 | Tcea1 | 34 kDa | 8 C | 2.865 | 0.067822066 |
| Sodium/potassium-transporting ATPase subunit beta-1 | P14094 | Atp1b1 | 35 kDa | 7 C | 2.839 | 0.093687894 |
| Synaptotagmin-1 | P46096 | Syt1 | 47 kDa | 6 C | 2.769 | 0.054419044 |
| Synapsin-2 | Q64332 | Syn2 | 63 kDa | 5 C | 2.754 | 0.068263222 |
| Ran-specific GTPase-activating protein | P47757 (+1) | Ranbp1 | 24 kDa | 3 C | 2.750 | 0.112465387 |
| PC4 and SFRS1-interacting protein | Q99JF8 | Psip1 | 60 kDa | 2 C | 2.710 | 0.075902966 |
| Glycogen phosphorylase, brain form | Q8CI94 | Pygb | 97 kDa | 12 C | 2.667 | 0.126178675 |
| Hydroxyacylglutathione hydrolase, mitochondrial | Q99KB8 (+1) | Hagh | 34 kDa | 8 C | 2.667 | 0.147723001 |
| Synaptophysin | Q62277 | Syp | 34 kDa | 5 C | 2.646 | 0.449460363 |
| Tubulin polymerization-promoting protein | Q7TQD2 | Tppp | 24 kDa | 3 C | 2.646 | 0.115468476 |
| T-complex protein 1 subunit zeta | P80317 | Cct6a | 58 kDa | 8 C | 2.646 | 0.374028936 |
| Hepatoma-derived growth factor-related protein 3 | Q9JMG7 (+1) | Hdgfl3 | 22 kDa | 1 C | 2.585 | 0.286477374 |
| Membrane-associated guanylate kinase, WW and PDZ domain-containing protein 2 | Q9WVQ1 | Magi2 | 141 kDa | 14 C | 2.585 | 0.286477374 |
| Galectin-1 | P62702 | Lgals1 | 15 kDa | 6 C | 2.585 | 0.286477374 |
| 40S ribosomal protein S4, X isoform | P24452 | Rps4x | 30 kDa | 4 C | 2.585 | 0.286477374 |
| Macrophage-capping protein | P16045 | Capg | 39 kDa | 5 C | 2.585 | 0.286477374 |
| Polymerase delta-interacting protein 2 | Q9WTP6 (+1) | Poldip2 | 42 kDa | 4 C | 2.585 | 0.286477374 |
| Adenylate kinase 2, mitochondrial | Q91VA6 | Ak2 | 26 kDa | 5 C | 2.585 | 0.286477374 |
| Receptor-type tyrosine-protein phosphatase zeta | Q9QUR6 | Ptprz1 | 254 kDa | 22 C | 2.582 | 0.119202463 |
| Adenylyl cyclase-associated protein 2 | B9EKR1 | Cap2 | 53 kDa | 13 C | 2.582 | 0.19511802 |
| Moesin | P26041 | Msn | 68 kDa | 2 C | 2.582 | 0.19511802 |
| Prolyl endopeptidase | Q7TQI3 | Prep | 81 kDa | 17 C | 2.582 | 0.19511802 |
| Ubiquitin thioesterase OTUB1 | Q9CYT6 | Otub1 | 31 kDa | 4 C | 2.582 | 0.19511802 |
| Beta-adducin | Q9QYB8 | Add2 | 81 kDa | 8 C | 2.537 | 0.05134265 |
| Isoform 2 of 4F2 cell-surface antigen heavy chain | P10852-2 | Slc3a2 | 62 kDa | 2 C | 2.504 | 0.345130314 |
| S-adenosylhomocysteine hydrolase-like protein 1 | Q80SW1 | Ahcyl1 | 59 kDa | 19 C | 2.504 | 0.381686567 |
| Poly(rC)-binding protein 1 | P60335 | Pcbp1 | 37 kDa | 9 C | 2.476 | 0.051645791 |
| Sepiapterin reductase | Q64105 | Spr | 28 kDa | 10 C | 2.470 | 0.26695978 |
| High mobility group protein B1 | P63158 | Hmgb1 | 25 kDa | 3 C | 2.462 | 0.146502425 |
| Oxygen-dependent coproporphyrinogen-III oxidase, mitochondrial | Q9DB27-2 | Cpox | 50 kDa | 10 C | 2.435 | 0.14374529 |
| Isoform 2 of Malignant T-cell-amplified sequence 1 | P36552 | Mcts1 | 21 kDa | 4 C | 2.435 | 0.22249858 |
| Myelin-oligodendrocyte glycoprotein | Q61885 | Mog | 28 kDa | 7 C | 2.424 | 0.151903428 |
| Elongation factor 1-alpha 1 | P10126 | Eef1a1 | 50 kDa | 6 C | 2.380 | 0.133604851 |
| Caskin-1 | Q9D1E6 | Caskin1 | 150 kDa | 10 C | 2.361 | 0.094373094 |
| Dematin | Q9WV69 (+1) | Dmtn | 45 kDa | 3 C | 2.348 | 0.255078181 |
| Elongation factor 1-alpha 2 | P62631 | Eef1a2 | 50 kDa | 6 C | 2.314 | 0.077311748 |
| Isoform 2 of Eukaryotic initiation factor 4A-II | P10630-2 | Eif4a2 | 46 kDa | 4 C | 2.305 | 0.223100575 |
| Elongation factor 1-gamma | Q9D8N0 | Eef1g | 50 kDa | 6 C | 2.300 | 0.284933121 |
| Histidine triad nucleotide-binding protein 1 | P70349 | Hint1 | 14 kDa | 2 C | 2.286 | 0.153564249 |
| Tripartite motif-containing protein 2 | Q9ESN6 | Trim2 | 81 kDa | 18 C | 2.272 | 0.051705867 |
| Ig kappa chain V-III region PC 2880/PC 1229 | P01654 (+1) | | 12 kDa | 2 C | 2.255 | 0.34242599 |
| NADH dehydrogenase [ubiquinone] flavoprotein 2, mitochondrial | Q3UUI3 | Ndufv2 | 27 kDa | 6 C | 2.250 | 0.129175413 |
| Acyl-coenzyme A thioesterase THEM4 | Q9D6J6 | Them4 | 26 kDa | 4 C | 2.250 | 0.129175413 |
| Adaptin ear-binding coat-associated protein 1 | Q9CR95 | Necap1 | 30 kDa | 1 C | 2.230 | 0.204282153 |
| Ig heavy chain V region AC38 205.12 | P06330 | | 13 kDa | 2 C | 2.218 | 0.193661573 |
| Heat shock protein HSP 90-alpha | P07901 | Hsp90aa1 | 85 kDa | 7 C | 2.200 | 0.062606719 |
| Isoform 2 of ELKS/Rab6-interacting/CAST family member 1 | Q9Z2U1 | Erc1 | 112 kDa | 4 C | 2.196 | 0.089293991 |
| Lactoylglutathione lyase | Q9CPU0 | Glo1 | 21 kDa | 3 C | 2.182 | 0.083077963 |
| Endonuclease domain-containing 1 protein | Q8C522 | Endod1 | 55 kDa | 12 C | 2.182 | 0.083077963 |
| F-actin-capping protein subunit alpha-2 | P47754 | Capza2 | 33 kDa | 3 C | 2.166 | 0.183059155 |
| Guanine nucleotide-binding protein G(i) subunit alpha-1 | B2RSH2 | Gnai1 | 40 kDa | 10 C | 2.114 | 0.336224353 |
| Cytochrome c, somatic | P62897 | Cycs | 12 kDa | 2 C | 2.100 | 0.085470245 |
| Isoleucine--tRNA ligase, mitochondrial | Q8BIJ6 | Iars2 | 113 kDa | 20 C | 2.098 | 0.178376649 |
| Small nuclear ribonucleoprotein-associated protein B | P27048 | Snrpb | 24 kDa | 3 C | 2.065 | 0.139342663 |

| | | | | | | |
|---|---|---|---|---|---|---|
| T-complex protein 1 subunit alpha | P11983 | Tcp1 | 60 kDa | 6 C | 2.050 | 0.421105136 |
| Ig kappa chain C region | P01837 | | 12 kDa | 3 C | 2.044 | 0.134170023 |
| Serine/threonine-protein kinase LMTK3 | P00405 | Lmtk3 | 151 kDa | 19 C | 2.031 | 0.500453236 |
| Branched-chain-amino-acid aminotransferase, cytosolic | P24288 | Bcat1 | 43 kDa | 10 C | 2.031 | 0.500453236 |
| 60S acidic ribosomal protein P0 | Q05920 | Rplp0 | 34 kDa | 3 C | 2.031 | 0.500453236 |
| Pyruvate carboxylase, mitochondrial | Q5XJV6 | Pc | 130 kDa | 13 C | 2.031 | 0.500453236 |
| Cytochrome c oxidase subunit 2 | Q8VCG4 | Mtco2 | 26 kDa | 3 C | 2.031 | 0.500453236 |
| Complement component C8 gamma chain | P14869 | C8g | 23 kDa | 3 C | 2.031 | 0.500453236 |
| PITH domain-containing protein 1 | Q8BWR2 | Pithd1 | 24 kDa | 4 C | 2.025 | 0.271700588 |
| Bleomycin hydrolase | Q8R016 | Blmh | 53 kDa | 4 C | 2.025 | 0.271700588 |
| Dihydropyrimidinase-related protein 3 | Q62188 | Dpysl3 | 62 kDa | 7 C | 1.988 | 0.001979015 |
| Triosephosphate isomerase | P17751 | Tpi1 | 32 kDa | 9 C | 1.979 | 0.001973066 |
| MAP7 domain-containing protein 2 | A2AG50 | Map7d2 | 86 kDa | 7 C | 1.964 | 0.007586217 |
| Protein PRRC2C | Q3TLH4 | Prrc2c | 311 kDa | 11 C | 1.956 | 0.043913765 |
| RNA-binding protein EWS | Q61545 | Ewsr1 | 68 kDa | 5 C | 1.951 | 0.08939479 |
| Long-chain specific acyl-CoA dehydrogenase, mitochondrial | P51174 | Acadl | 48 kDa | 7 C | 1.927 | 0.583059492 |
| WD repeat domain phosphoinositide-interacting protein 4 | Q91VM3 (+1) | Wdr45 | 40 kDa | 14 C | 1.924 | 0.359412393 |
| Myelin proteolipid protein | P60202 | Plp1 | 30 kDa | 14 C | 1.920 | 0.06724376 |
| Astrocytic phosphoprotein PEA-15 | P30416 | Pea15 | 15 kDa | 1 C | 1.920 | 0.056457972 |
| Hemopexin | Q91X72 | Hpx | 51 kDa | 13 C | 1.920 | 0.056457972 |
| Peptidyl-prolyl cis-trans isomerase FKBP4 | Q62048 | Fkbp4 | 52 kDa | 7 C | 1.920 | 0.239525231 |
| Actin, cytoplasmic 1 | P60710 | Actb | 42 kDa | 6 C | 1.915 | 0.011532209 |
| Alpha-1-antitrypsin 1-1 | P07758 (+1) | Serpina1a | 46 kDa | 3 C | 1.905 | 0.216538162 |
| Peroxiredoxin-2 | Q61171 | Prdx2 | 22 kDa | 3 C | 1.898 | 0.001147394 |
| Eukaryotic translation initiation factor 4B | Q8BGD9 | Eif4b | 69 kDa | 3 C | 1.882 | 0.053637574 |
| Annexin A5 | P48036 | Anxa5 | 36 kDa | 1 C | 1.882 | 0.197430576 |
| Heterogeneous nuclear ribonucleoprotein D-like | Q9Z130 | Hnrnpdl | 34 kDa | 3 C | 1.867 | 0.000455139 |
| Peroxiredoxin-4 | O08807 | Prdx4 | 31 kDa | 4 C | 1.843 | 0.025703067 |
| Selenium-binding protein 1 | P17563 | Selenbp1 | 53 kDa | 10 C | 1.829 | 0.106675452 |
| Src substrate cortactin | Q60598 | Cttn | 61 kDa | 3 C | 1.829 | 0.013587476 |
| Myc box-dependent-interacting protein 1 | O08539 | Bin1 | 64 kDa | 4 C | 1.822 | 0.020189936 |
| Cytochrome b-c1 complex subunit 2, mitochondrial | Q9DB77 | Uqcrc2 | 48 kDa | 1 C | 1.813 | 0.090408688 |
| Methylcrotonoyl-CoA carboxylase beta chain, mitochondrial | Q3ULD5 | Mccc2 | 61 kDa | 10 C | 1.807 | 0.000955006 |
| CB1 cannabinoid receptor-interacting protein 1 | Q5M8N0 | Cnrip1 | 19 kDa | 2 C | 1.806 | 0.246932657 |
| Plasminogen activator inhibitor 1 RNA-binding protein | Q9CY58 | Serbp1 | 45 kDa | 2 C | 1.802 | 0.218805979 |
| Apoptotic chromatin condensation inducer in the nucleus | Q9JIX8 (+1) | Acin1 | 151 kDa | 7 C | 1.800 | 0.149564746 |
| Pyruvate dehydrogenase E1 component subunit alpha, somatic form, mitochondrial | P35486 | Pdha1 | 43 kDa | 12 C | 1.800 | 0.219191547 |
| Proteasome subunit alpha type-1 | Q9R1P4 | Psma1 | 30 kDa | 5 C | 1.778 | 0.094507828 |
| Alpha-adducin | Q9QYC0 | Add1 | 81 kDa | 7 C | 1.778 | 0.094507828 |
| Cleavage and polyadenylation specificity factor subunit 5 | Q9CQF3 | Nudt21 | 26 kDa | 1 C | 1.774 | 0.43607882 |
| Neuronal membrane glycoprotein M6-a | P35802 | Gpm6a | 31 kDa | 13 C | 1.774 | 0.43607882 |
| Disks large homolog 1 | Q811D0 | Dlg1 | 100 kDa | 7 C | 1.748 | 0.061950407 |
| Isoform 4 of Microtubule-associated protein 4 | P27546-4 | Map4 | 95 kDa | 10 C | 1.741 | 0.070438185 |
| Fibrinogen gamma chain | P51880 | Fgg | 49 kDa | 12 C | 1.725 | 0.157306561 |
| Fatty acid-binding protein, brain | Q8VCM7 | Fabp7 | 15 kDa | 5 C | 1.725 | 0.219164384 |
| Transaldolase | Q93092 | Taldo1 | 37 kDa | 3 C | 1.714 | 0.058153609 |
| Fumarylacetoacetase | P35505 | Fah | 46 kDa | 6 C | 1.714 | 0.139519583 |
| Eukaryotic translation initiation factor 5 | P59325 | Eif5 | 49 kDa | 8 C | 1.714 | 0.020721124 |
| Clathrin light chain B | Q6IRU5 | Cltb | 25 kDa | 2 C | 1.708 | 0.488424448 |
| Dynein light chain 2, cytoplasmic | Q9D0M5 | Dynll2 | 10 kDa | 2 C | 1.705 | 0.124604379 |
| Alpha-actinin-1 | Q7TPR4 | Actn1 | 103 kDa | 11 C | 1.680 | 0.103059278 |
| Nucleophosmin | Q61937 | Npm1 | 33 kDa | 3 C | 1.680 | 0.114064092 |
| Calcium-binding mitochondrial carrier protein Aralar1 | Q8BH59 | Slc25a12 | 75 kDa | 7 C | 1.671 | 0.537731021 |
| Peroxiredoxin-1 | P35700 | Prdx1 | 22 kDa | 4 C | 1.638 | 0.062623886 |
| 78 kDa glucose-regulated protein | P20029 | Hspa5 | 72 kDa | 1 C | 1.617 | 0.100586248 |
| Heterogeneous nuclear ribonucleoprotein A1 | P49312 | Hnrnpa1 | 34 kDa | 2 C | 1.600 | 0.147236457 |
| Probable G-protein coupled receptor 158 | Q8C419 | Gpr158 | 134 kDa | 26 C | 1.600 | 0.227452818 |
| Methylglutaconyl-CoA hydratase, mitochondrial | Q9JLZ3 | Auh | 33 kDa | 5 C | 1.600 | 0.063924123 |
| Proteasome subunit alpha type-7 | Q9Z2U0 | Psma7 | 28 kDa | 3 C | 1.600 | 0.122182492 |
| Pyruvate dehydrogenase protein X component, mitochondrial | Q923D2 | Pdhx | 54 kDa | 5 C | 1.600 | 0.343336751 |
| Flavin reductase (NADPH) | P38060 | Blvrb | 22 kDa | 2 C | 1.600 | 0.508755163 |
| Hydroxymethylglutaryl-CoA lyase, mitochondrial | Q8BKZ9 | Hmgcl | 34 kDa | 8 C | 1.600 | 0.596503285 |
| Heterogeneous nuclear ribonucleoproteins A2/B1 | O88569 | Hnrnpa2b1 | 37 kDa | 1 C | 1.600 | 0.017722124 |
| Succinate dehydrogenase [ubiquinone] iron-sulfur subunit, mitochondrial | Q9CQA3 | Sdhb | 32 kDa | 14 C | 1.600 | 0.029660167 |

| | | | | | | |
|---|---|---|---|---|---|---|
| Isoform 2 of Hepatoma-derived growth factor-related protein 2 | Q3UMU9-2 | Hdgfl2 | 74 kDa | 3 C | 1.600 | 0.43647815 |
| Formin-binding protein 1-like | Q8K012 | Fnbp1l | 70 kDa | 8 C | 1.580 | 0.394596718 |
| Protein phosphatase 1 regulatory subunit 12C | P51855 | Ppp1r12c | 85 kDa | 8 C | 1.574 | 0.357478808 |
| Immunoglobulin J chain | Q3UMT1 | Jchain | 18 kDa | 8 C | 1.574 | 0.357478808 |
| Glutathione synthetase | P01592 | Gss | 52 kDa | 4 C | 1.574 | 0.357478808 |
| Wiskott-Aldrich syndrome protein family member 1 | Q8R5H6 | Wasf1 | 62 kDa | 5 C | 1.561 | 0.232283407 |
| Ubiquitin-40S ribosomal protein S27a | P62983 | Rps27a | 18 kDa | 6 C | 1.561 | 0.34458907 |
| Rab-like protein 6 | Q5U3K5 | Rabl6 | 80 kDa | 5 C | 1.543 | 0.465826074 |
| Rabphilin-3A | P47708 | Rph3a | 75 kDa | 15 C | 1.533 | 0.061986861 |
| Annexin A2 | Q9JM52 | Anxa2 | 39 kDa | 5 C | 1.527 | 0.54904678 |
| Misshapen-like kinase 1 | P07356 | Mink1 | 147 kDa | 12 C | 1.527 | 0.54904678 |
| Isoform 4 of Actin-binding LIM protein 2 | Q8BL65-4 | Ablim2 | 69 kDa | 3 C | 1.527 | 0.54904678 |
| Ras-related protein Rab-1A | P62821 | Rab1A | 23 kDa | 4 C | 1.525 | 0.484089872 |
| Protein disulfide-isomerase A3 | P27773 | Pdia3 | 57 kDa | 8 C | 1.500 | 0.06042742 |
| Nascent polypeptide-associated complex subunit alpha, muscle-specific form | P70670 | Naca | 220 kDa | 16 C | 1.486 | 0.135503222 |
| Fructose-bisphosphate aldolase A | P05064 | Aldoa | 39 kDa | 8 C | 1.486 | 0.014106752 |
| SH3 domain-binding glutamic acid-rich-like protein 3 | Q91VW3 | Sh3bgrl3 | 10 kDa | 1 C | 1.477 | 0.753244047 |
| Proteasome subunit alpha type-6 | Q9QUM9 | Psma6 | 27 kDa | 8 C | 1.477 | 0.265756364 |
| Fatty-acid amide hydrolase 1 | O08914 | Faah | 63 kDa | 18 C | 1.477 | 0.753244047 |
| Ig mu chain C region | P01872 | Ighm | 50 kDa | 19 C | 1.475 | 0.128352379 |
| Microtubule-associated protein tau | P10637 | Mapt | 76 kDa | 2 C | 1.463 | 0.022534771 |
| Transgelin-3 | Q9R1Q8 | Tagln3 | 22 kDa | 3 C | 1.461 | 0.043026248 |
| Isoform 2 of Synaptopodin | Q8CC35-2 | Synpo | 96 kDa | 5 C | 1.459 | 0.263453276 |
| Neuromodulin | P06837 | Gap43 | 24 kDa | 2 C | 1.447 | 0.011264539 |
| Phospholipase D3 | O35405 | Pld3 | 54 kDa | 8 C | 1.443 | 0.363605629 |
| Vacuolar protein sorting-associated protein 26B | Q8C0E2 | Vps26b | 39 kDa | 1 C | 1.440 | 0.45574536 |
| Aspartate aminotransferase, cytoplasmic | P05201 | Got1 | 46 kDa | 5 C | 1.404 | 0.029123156 |
| D-tyrosyl-tRNA(Tyr) deacylase 1 | Q9DD18 | Dtd1 | 23 kDa | 3 C | 1.400 | 0.193006088 |
| Protein disulfide-isomerase A4 | P08003 | Pdia4 | 72 kDa | 6 C | 1.382 | 0.063924123 |
| Glycerol-3-phosphate phosphatase | Q8CHP8 | Pgp | 35 kDa | 8 C | 1.371 | 0.23806046 |
| LIM zinc-binding domain-containing Nebulette | Q9DC07 | Nebl | 31 kDa | 7 C | 1.366 | 0.421963137 |
| Dynein light chain 1, cytoplasmic | P63168 | Dynll1 | 10 kDa | 3 C | 1.333 | 0.613173826 |
| Microtubule-associated protein 6 | Q7TSJ2 | Map6 | 96 kDa | 5 C | 1.321 | 0.12233082 |
| Alpha-aminoadipic semialdehyde dehydrogenase | Q9DBF1 (+1) | Aldh7a1 | 59 kDa | 9 C | 1.314 | 0.245220773 |
| Hyaluronan and proteoglycan link protein 1 | Q9QUP5 | Hapln1 | 40 kDa | 11 C | 1.311 | 0.64012481 |
| Mammalian ependymin-related protein 1 | Q99M71 | Epdr1 | 25 kDa | 7 C | 1.309 | 0.497432213 |
| Malate dehydrogenase, cytoplasmic | P14152 | Mdh1 | 37 kDa | 3 C | 1.300 | 0.087622829 |
| Isoform 2 of Methionine adenosyltransferase 2 subunit beta | Q9Z2I0 | Mat2b | 36 kDa | 7 C | 1.300 | 0.712278087 |
| Mitochondrial proton/calcium exchanger protein | Q99LB6-2 | Letm1 | 83 kDa | 12 C | 1.300 | 0.712278087 |
| Protein phosphatase 1 regulatory subunit 12B | Q8BG95 | Ppp1r12b | 109 kDa | 6 C | 1.292 | 0.342618418 |
| Carbonic anhydrase-related protein 10 | P61215 | Ca10 | 38 kDa | 5 C | 1.280 | 0.289311364 |
| ELAV-like protein 4 | Q61701 | Elavl4 | 42 kDa | 5 C | 1.280 | 0.50971724 |
| Coronin-1A | Q9Z2H5 | Coro1a | 51 kDa | 12 C | 1.275 | 0.687074079 |
| Band 4.1-like protein 1 | O89053 | Epb41l1 | 98 kDa | 11 C | 1.275 | 0.687074079 |
| Protein phosphatase 1 regulatory subunit 11 | Q8K1L5 | Ppp1r11 | 15 kDa | 5 C | 1.275 | 0.687074079 |
| Tropomyosin alpha-1 chain | P58771 | Tpm1 | 33 kDa | 1 C | 1.267 | 0.368981333 |
| Protein phosphatase 1 regulatory subunit 12A | Q9DBR7 | Ppp1r12a | 115 kDa | 8 C | 1.221 | 0.439621189 |
| Cathepsin L1 | P06797 | Ctsl | 38 kDa | 10 C | 1.220 | 0.604985939 |
| Aspartate aminotransferase, mitochondrial | P05202 | Got2 | 47 kDa | 7 C | 1.207 | 0.277282141 |
| Chromobox protein homolog 3 | P23198 | Cbx3 | 21 kDa | 3 C | 1.200 | 0.078619235 |
| Hemoglobin subunit beta-1 | P02088 | Hbb-b1 | 16 kDa | 2 C | 1.200 | 0.388896231 |
| Vitamin D-binding protein | P21614 | Gc | 54 kDa | 28 C | 1.200 | 0.827125331 |
| Thioredoxin-dependent peroxide reductase, mitochondrial | P20108 | Prdx3 | 28 kDa | 4 C | 1.159 | 0.193624159 |
| Adenosylhomocysteinase | P50247 | Ahcy | 48 kDa | 9 C | 1.156 | 0.589123466 |
| Stress-70 protein, mitochondrial | P38647 | Hspa9 | 73 kDa | 5 C | 1.152 | 0.214916452 |
| Electron transfer flavoprotein-ubiquinone oxidoreductase, mitochondrial | Q921G7 | Etfdh | 68 kDa | 15 C | 1.148 | 0.890942227 |
| G protein-regulated inducer of neurite outgrowth 1 | Q3UNH4 | Gprin1 | 95 kDa | 11 C | 1.137 | 0.564208541 |
| CD5 antigen-like | Q9QWK4 | Cd5l | 39 kDa | 26 C | 1.127 | 0.50971724 |
| Isoform 2 of Cytosol aminopeptidase | Q9CPY7-2 | Lap3 | 53 kDa | 7 C | 1.120 | 0.573323692 |
| Mitochondrial peptide methionine sulfoxide reductase | Q9D6Y7 (+1) | Msra | 26 kDa | 4 C | 1.120 | 0.684528336 |
| Succinate dehydrogenase [ubiquinone] flavoprotein subunit, mitochondrial | Q8K2B3 | Sdha | 73 kDa | 19 C | 1.111 | 0.345420222 |
| 26S proteasome non-ATPase regulatory subunit 9 | Q9CR00 | Psmd9 | 25 kDa | 3 C | 1.091 | 0.790416234 |
| Neutral cholesterol ester hydrolase 1 | Q8BLF1 | Nceh1 | 46 kDa | 4 C | 1.075 | 0.938623127 |
| Proteasome subunit beta type-6 | Q60692 | Psmb6 | 25 kDa | 4 C | 1.067 | 0.852853872 |

| | | | | | | |
|---|---|---|---|---|---|---|
| Endoplasmic reticulum resident protein 44 | Q9D1Q6 | Erp44 | 47 kDa | 7 C | 1.042 | 0.970111629 |
| Hepatoma-derived growth factor | A2AJI0 | Hdgf | 26 kDa | 2 C | 1.029 | 0.932529844 |
| MAP7 domain-containing protein 1 | P51859 | Map7d1 | 93 kDa | 6 C | 1.029 | 0.935632269 |
| Dihydrolipoyl dehydrogenase, mitochondrial | O08749 | Dld | 54 kDa | 9 C | 1.026 | 0.848858543 |
| Betaine--homocysteine S-methyltransferase 1 | O35490 | Bhmt | 45 kDa | 8 C | 1.015 | 0.983277979 |
| Nucleoside diphosphate kinase A | Q80YN3 | Nme1 | 17 kDa | 2 C | 1.000 | 1 |
| Breast carcinoma-amplified sequence 1 homolog | P15532 | Bcas1 | 67 kDa | 8 C | 1.000 | 1 |
| Lamina-associated polypeptide 2, isoforms alpha/zeta | Q61033 | Tmpo | 75 kDa | 1 C | 0.992 | 0.967302017 |
| Thyroid hormone receptor-associated protein 3 | Q569Z6 | Thrap3 | 108 kDa | 1 C | 0.968 | 0.952137997 |
| Growth factor receptor-bound protein 2 | Q60631 | Grb2 | 25 kDa | 2 C | 0.967 | 0.931762061 |
| Inositol monophosphatase 1 | O55023 | Impa1 | 30 kDa | 6 C | 0.925 | 0.424789699 |
| Proteasome subunit beta type-5 | O55234 | Psmb5 | 29 kDa | 3 C | 0.923 | 0.772355126 |
| Bcl-2-associated transcription factor 1 | Q8K019 (+1) | Bclaf1 | 106 kDa | 5 C | 0.914 | 0.684528336 |
| GMP reductase 2 | Q99L27 | Gmpr2 | 38 kDa | 9 C | 0.914 | 0.774595202 |
| Fumarylacetoacetate hydrolase domain-containing protein 2A | Q3TC72 | Fahd2 | 35 kDa | 6 C | 0.900 | 0.729434311 |
| CAP-Gly domain-containing linker protein 1 | Q922J3 | Clip1 | 156 kDa | 11 C | 0.880 | 0.586114133 |
| Mitochondrial 2-oxoglutarate/malate carrier protein | Q9CR62 | Slc25a11 | 34 kDa | 3 C | 0.876 | 0.722165328 |
| Metastasis suppressor protein 1 | Q8R1S4 | Mtss1 | 82 kDa | 11 C | 0.873 | 0.889887267 |
| Vesicular glutamate transporter 1 | Q3TXX4 | Slc17a7 | 62 kDa | 12 C | 0.873 | 0.889887267 |
| Pyruvate dehydrogenase E1 component subunit beta, mitochondrial | Q9D051 | Pdhb | 39 kDa | 6 C | 0.844 | 0.682188682 |
| Dihydrolipoyllysine-residue acetyltransferase component of pyruvate dehydrogenase comp | Q8BMF4 | Dlat | 68 kDa | 10 C | 0.835 | 0.587930925 |
| NADH dehydrogenase [ubiquinone] 1 alpha subcomplex subunit 9, mitochondrial | Q9DC69 | Ndufa9 | 43 kDa | 2 C | 0.825 | 0.817523442 |
| L-lactate dehydrogenase B chain | P16125 | Ldhb | 37 kDa | 5 C | 0.814 | 0.4150258 |
| Endoplasmin | P08113 | Hsp90b1 | 92 kDa | 5 C | 0.810 | 0.630589837 |
| Glycogen phosphorylase, muscle form | Q9WUB3 | Pygm | 97 kDa | 8 C | 0.808 | 0.749949259 |
| Methylmalonate-semialdehyde dehydrogenase [acylating], mitochondrial | P01942 | Aldh6a1 | 58 kDa | 8 C | 0.800 | 0.292351992 |
| Hemoglobin subunit alpha | Q9EQ20 | Hba | 15 kDa | 1 C | 0.800 | 0.303090534 |
| Ran-binding protein 3 | Q9CT10 | Ranbp3 | 53 kDa | 5 C | 0.800 | 0.407083822 |
| Neurocan core protein | P55066 | Ncan | 137 kDa | 34 C | 0.800 | 0.819850613 |
| Macrophage migration inhibitory factor | P34884 | Mif | 13 kDa | 3 C | 0.800 | 0.743372989 |
| Electron transfer flavoprotein subunit beta | Q9DCW4 | Etfb | 28 kDa | 4 C | 0.789 | 0.647782186 |
| Glyceraldehyde-3-phosphate dehydrogenase | P16858 | Gapdh | 36 kDa | 5 C | 0.740 | 0.028682866 |
| Gamma-enolase | P17183 | Eno2 | 47 kDa | 6 C | 0.740 | 0.00595225 |
| Adenylate kinase isoenzyme 1 | Q9R0Y5 (+1) | Ak1 | 22 kDa | 2 C | 0.706 | 0.099559166 |
| Succinyl-CoA:3-ketoacid coenzyme A transferase 1, mitochondrial | Q9D0K2 | Oxct1 | 56 kDa | 7 c | 0.695 | 0.295154117 |
| Citrate synthase, mitochondrial | Q9CZU6 | Cs | 52 kDa | 5 C | 0.686 | 0.358984804 |
| Glycerol-3-phosphate dehydrogenase [NAD(+)], cytoplasmic | P13707 | Gpd1 | 38 kDa | 11 C | 0.675 | 0.5990798 |
| Cytochrome b-c1 complex subunit 1, mitochondrial | Q9CZ13 | Uqcrc1 | 53 kDa | 11 C | 0.656 | 0.205679918 |
| Serpin B6 | Q60854 | Serpinb6 | 43 kDa | 6 C | 0.629 | 0.564208541 |
| Creatine kinase S-type, mitochondrial | Q6P8J7 | Ckmt2 | 47 kDa | 8 C | 0.600 | 0.151399763 |
| Alpha-enolase | P17182 | Eno1 | 47 kDa | 6 C | 0.593 | 0.001824122 |
| 60 kDa heat shock protein, mitochondrial | P63038 | Hspd1 | 61 kDa | 3 C | 0.577 | 0.067750018 |
| Malectin | Q6ZQI3 | Mlec | 32 kDa | 3 C | 0.575 | 0.492633575 |
| Thioredoxin reductase 2, mitochondrial | Q9JLT4 (+1) | Txnrd2 | 57 kDa | 11 C | 0.560 | 0.206259148 |
| D-beta-hydroxybutyrate dehydrogenase, mitochondrial | Q80XN0 | Bdh1 | 38 kDa | 6 C | 0.542 | 0.461165939 |
| Fumarate hydratase, mitochondrial | P97807 | Fh | 54 kDa | 4 C | 0.533 | 0.100398681 |
| NADH-ubiquinone oxidoreductase 75 kDa subunit, mitochondrial | Q91VD9 | Ndufs1 | 80 kDa | 18 C | 0.533 | 0.00122918 |
| Cytochrome b-c1 complex subunit Rieske, mitochondrial | Q9CR68 | Uqcrfs1 | 29 kDa | 5 C | 0.522 | 0.092120273 |
| ADP/ATP translocase 2 | P51881 | Slc25a5 | 33 kDa | 4 C | 0.519 | 0.122413092 |
| Isocitrate dehydrogenase [NADP], mitochondrial | P54071 | Idh2 | 51 kDa | 8 C | 0.492 | 0.089938561 |
| Talin-1 | P26039 | Tln1 | 270 kDa | 38 C | 0.487 | 0.545515313 |
| WAS/WASL-interacting protein family member 3 | P0C7L0 | Wipf3 | 49 kDa | 4 C | 0.480 | 0.08646656 |
| 2,4-dienoyl-CoA reductase, mitochondrial | Q9CQ62 | Decr1 | 36 kDa | 5 C | 0.466 | 0.482288821 |
| NADH dehydrogenase [ubiquinone] 1 alpha subcomplex subunit 10, mitochondrial | Q99LC3 | Ndufa10 | 41 kDa | 5 C | 0.444 | 0.292351992 |
| ADP/ATP translocase 1 | P48962 | Slc25a4 | 33 kDa | 4 C | 0.441 | 0.107356845 |
| Desmoplakin | E9Q557 | Dsp | 333 kDa | 43 C | 0.438 | 0.394547626 |
| Electron transfer flavoprotein subunit alpha, mitochondrial | Q99LC5 | Etfa | 35 kDa | 6 C | 0.400 | 0.063924123 |
| 3-hydroxyacyl-CoA dehydrogenase type-2 | O08756 | Hsd17b10 | 27 kDa | 2 C | 0.383 | 0.303632367 |
| Myosin-9 | Q8VDD5 | Myh9 | 226 kDa | 21 C | 0.350 | 0.282810056 |
| Succinate--CoA ligase [ADP-forming] subunit beta, mitochondrial | Q9Z2I9 | Sucla2 | 50 kDa | 6 C | 0.295 | 0.058413578 |
| Alpha-actinin-2 | Q9JI91 | Actn2 | 104 kDa | 10 C | 0.210 | 0.050003617 |
| Ig heavy chain V-III region A4 | P01796 (+3) | | 13 kDa | 2 C | 0.174 | 0.292351992 |
| Collagen alpha-1(VI) chain | Q04857 | Col6a1 | 108 kDa | 20 C | 0.174 | 0.292351992 |
| Short/branched chain specific acyl-CoA dehydrogenase, mitochondrial | Q9DBL1 | Acadsb | 48 kDa | 8 C | 0.174 | 0.292351992 |

| | | | | | | |
|---|---|---|---|---|---|---|
| Very long-chain specific acyl-CoA dehydrogenase, mitochondrial | P50544 | Acadvl | 71 kDa | 7 C | 0.151 | 0.065561447 |
| NADH dehydrogenase [ubiquinone] flavoprotein 1, mitochondrial | Q91YT0 | Ndufv1 | 51 kDa | 12 C | 0.138 | 0.111106307 |
| Desmin | P31001 | Des | 53 kDa | 1 C | 0.131 | 0.064671855 |
| PDZ and LIM domain protein 5 | Q8CI51 | Pdlim5 | 63 kDa | 22 C | 0.077 | 0.183043643 |
| Junction plakoglobin | Q02257 | Jup | 82 kDa | 13 C | 0.077 | 0.100434009 |
| Obscurin | A2AAJ9 | Obscn | 966 kDa | 207 C | 0.066 | 0.098532637 |
| Long-chain-fatty-acid--CoA ligase 1 | P41216 | Acsl1 | 78 kDa | 18 C | 0.066 | 0.098532637 |
| Atypical kinase COQ8A, mitochondrial | Q60936 | Coq8a | 72 kDa | 8 C | 0.056 | 0.138226369 |
| Protein S100-A1 | P56565 | S100a1 | 11 kDa | 1 C | 0.049 | 0.106157348 |
| Perilipin-4 | O88492 | Plin4 | 139 kDa | 17 C | 0.048 | 0.292351992 |
| Troponin I, cardiac muscle | P48787 | Tnni3 | 24 kDa | 2 C | 0.036 | 0.087343362 |
| LIM domain-binding protein 3 | Q9JKS4 (+1) | Ldb3 | 76 kDa | 21 C | 0.033 | 0.196780115 |
| Medium-chain specific acyl-CoA dehydrogenase, mitochondrial | P45952 | Acadm | 46 kDa | 8 C | 0.031 | 0.082090741 |
| NADH dehydrogenase [ubiquinone] iron-sulfur protein 2, mitochondrial | Q91WD5 | Ndufs2 | 53 kDa | 7 C | 0.026 | 0.099839072 |
| Myoglobin | P04247 | Mb | 17 kDa | 1 C | 0.409 | 0.030377445 |
| Actin, alpha cardiac muscle 1 | P68033 | Actc1 | 42 kDa | 6 C | 0.385 | 0.027287892 |
| ATP synthase subunit alpha, mitochondrial | Q03265 | Atp5a1 | 60 kDa | 2 C | 0.375 | 0.034633095 |
| Isovaleryl-CoA dehydrogenase, mitochondrial | Q9JHI5 | Ivd | 46 kDa | 7 C | 0.325 | 0.029157721 |
| Vimentin | P20152 | Vim | 54 kDa | 1 C | 0.320 | 0.000761185 |
| Cytochrome c oxidase subunit 5A, mitochondrial | P12787 | Cox5a | 16 kDa | 4 C | 0.307 | 0.022310136 |
| Fructose-1,6-bisphosphatase 1 | Q9QXD6 | Fbp1 | 37 kDa | 7 C | 0.296 | 0.049896549 |
| Ras-related protein Rap-1b | Q99JI6 | Rap1b | 21 kDa | 4 C | 0.284 | 0.013276947 |
| Enoyl-CoA delta isomerase 1, mitochondrial | P42125 | Eci1 | 32 kDa | 5 C | 0.267 | 0.000434525 |
| 3-ketoacyl-CoA thiolase, mitochondrial | Q8BWT1 | Acaa2 | 42 kDa | 8 C | 0.264 | 0.009025458 |
| Beta-enolase | P21550 | Eno3 | 47 kDa | 6 C | 0.253 | 0.000140417 |
| Hydroxyacyl-coenzyme A dehydrogenase, mitochondrial | Q61425 | Hadh | 34 kDa | 5 C | 0.238 | 0.00230501 |
| Creatine kinase M-type | P07310 | Ckm | 43 kDa | 8 C | 0.234 | 0.013682437 |
| Voltage-dependent anion-selective channel protein 1 | Q60932 (+1) | Vdac1 | 32 kDa | 2 C | 0.222 | 0.014647192 |
| Trifunctional enzyme subunit beta, mitochondrial | Q99JY0 | Hadhb | 51 kDa | 5 C | 0.215 | 0.010844009 |
| Trifunctional enzyme subunit alpha, mitochondrial | Q8BMS1 | Hadha | 83 kDa | 12 C | 0.206 | 0.007752288 |
| Short-chain specific acyl-CoA dehydrogenase, mitochondrial | Q07417 | Acads | 45 kDa | 5 C | 0.184 | 0.001175837 |
| Myosin light chain 4 | P09541 | Myl4 | 21 kDa | 2 C | 0.176 | 0.009000722 |
| Myosin light chain 3 | P09542 | Myl3 | 22 kDa | 2 C | 0.172 | 0.008873964 |
| MICOS complex subunit Mic60 | Q8CAQ8 (+1) | Immt | 84 kDa | 7 C | 0.146 | 0.008962564 |
| Myosin-6 | Q02566 | Myh6 | 224 kDa | 14 C | 0.136 | 0.009906061 |
| ES1 protein homolog, mitochondrial | Q9D172 | D10Jhu81e | 28 kDa | 6 C | 0.124 | 0.003655179 |
| Voltage-dependent anion-selective channel protein 3 | Q60931 | Vdac3 | 31 kDa | 6 C | 0.123 | 0.017756615 |
| Delta-1-pyrroline-5-carboxylate dehydrogenase, mitochondrial | Q8CHT0 | Aldh4a1 | 62 kDa | 8 C | 0.113 | 0.009464046 |
| Voltage-dependent anion-selective channel protein 2 | Q60930 | Vdac2 | 32 kDa | 11 C | 0.108 | 0.040932146 |
| NADH dehydrogenase [ubiquinone] 1 alpha subcomplex subunit 5 | Q9CPP6 | Ndufa5 | 13 kDa | 1 C | 0.101 | 0.005124211 |
| Mannose-6-phosphate isomerase | Q924M7 | Mpi | 47 kDa | 12 C | 0.098 | 0.030042362 |
| Isoform 2 of Sarcoplasmic/endoplasmic reticulum calcium ATPase 2 | O55143-2 | Atp2a2 | 110 kDa | 29 C | 0.079 | 0.034005986 |
| Succinate--CoA ligase [ADP/GDP-forming] subunit alpha, mitochondrial | Q9WUM5 | Suclg1 | 36 kDa | 6 C | 0.074 | 0.027452445 |
| Phosphoglycerate mutase 2 | O70250 | Pgam2 | 29 kDa | 3 C | 0.066 | 0.003665612 |
| Titin | A2ASS6 | Ttn | 3906 kDa | 498 C | 0.053 | 0.005034806 |
| MICOS complex subunit Mic19 | Q9CRB9 | Chchd3 | 26 kDa | 4 C | 0.050 | 0.018341734 |
| MICOS complex subunit MIC13 | Q8R404 | Mic13 | 13 kDa | 1 C | 0.044 | 0.020624692 |
| Prohibitin | P67778 | Phb | 30 kDa | 1 C | 0.033 | 0.008544952 |
| Aldose reductase | P45376 | Akr1b1 | 36 kDa | 6 C | 0.033 | 0.013088357 |
| 2-oxoisovalerate dehydrogenase subunit beta, mitochondrial | Q6P3A8 | Bckdhb | 43 kDa | 14 C | 0.031 | 0.029450286 |
| Hydroxysteroid dehydrogenase-like protein 2 | Q2TPA8 | Hsdl2 | 54 kDa | 2 C | 0.027 | 0.031319381 |
| Acyl-coenzyme A thioesterase 13 | Q9CQR4 | Acot13 | 15 kDa | 2 C | 0.023 | 0.04895283 |
| Dehydrogenase/reductase SDR family member 4 | Q99LB2 | Dhrs4 | 30 kDa | 4 C | 0.022 | 0.001209297 |
| Myomesin-1 | Q62234 | Myom1 | 185 kDa | 21 C | 0.017 | 0.005251368 |
| Delta(3,5)-Delta(2,4)-dienoyl-CoA isomerase, mitochondrial | O35459 | Ech1 | 36 kDa | 6 C | 0.011 | 0.024190983 |
| Myosin-binding protein C, cardiac-type | O70468 | Mybpc3 | 141 kDa | 21 C | 0.005 | 0.013366939 |
