## Supplemental Table 5 for "Dietary restriction transforms the protein sulfhydrome in a tissue-specific and cystathionine γ-lyase-dependent manner"

| Supplemental Table 5: Accession Numbers for the 209 Common Sulfhydrated Proteins in Liver, Kidney, Muscle, and Brain |
|---|
| P48036 |
| Q03265 |
| Q02257 |
| Q921G7 |
| Q9D172 |
| P01942 |
| P20152 |
| P23953 |
| E9Q557 |
| P42125 |
| P63028 |
| P61082 |
| P51174 |
| Q9DB77 |
| P02088 |
| P46638 |
| P70349 |
| P45952 |
| Q9D6J6 |
| Q61838 |
| Q60854 |
| P06745 |
| P29699 |
| Q8BWT1 |
| P05202 |
| Q3TC72 |
| Q9WTX5 |
| P60335 |
| P57780 |
| Q9JKB1 |
| Q9D0F9 |
| P49722 |
| P06151 |
| P10639 |
| P08249 |
| Q9QWK4 |
| P01592 |
| Q9DAK9 |
| P24549 |
| Q91YT0 |
| Q9DBP5 |
| O70251 |
| Q99PT1 |
| Q9EQ20 |
| Q91VI7 |
| O70435 |

| |
|---|
| P17182 |
| P45591 |
| P50247 |
| Q9D1Q6 |
| P09411 |
| Q8BVI4 |
| P16045 |
| P63038 |
| Q9DCW4 |
| Q61425 |
| P68510 |
| Q91VD9 |
| Q9Z2U1 |
| Q61171 |
| P07724 |
| P26043 |
| P34884 |
| O08749 |
| Q99020 |
| Q99MN9 |
| Q9CQV8 |
| P16858 |
| P35700 |
| P34022 |
| O55234 |
| Q9Z2U0 |
| P14211 |
| Q9R1P0 |
| P00920 |
| P68037 |
| P11352 |
| P07356 |
| P05064 |
| Q9CZ13 |
| O08709 |
| O09061 |
| Q9DCM0 |
| Q9WVQ5 |
| P10126 |
| P38647 |
| Q91VM9 |
| P27773 |
| Q60676 |
| P70318 |
| P62259 |
| P14206 |
| Q99L47 |

| |
|---|
| Q8R086 |
| Q8QZT1 |
| Q60668-3 |
| P20108 |
| Q9R1P4 |
| P40142 |
| P62137 |
| P10518 |
| O88569 |
| P61982 |
| Q921I1 |
| P09671 |
| P62897 |
| Q8BWF0 |
| P70296 |
| Q62188 |
| P14152 |
| Q8K0E8 |
| Q9CQM9 |
| Q99L13 |
| Q8CDN6 |
| P20029 |
| Q9D1A2 |
| Q9R1P1 |
| P08003 |
| Q93092 |
| P35979 |
| P63017 |
| Q99KI0 |
| P47738 |
| Q9CPU0 |
| P09103 |
| P01837 |
| P47941 |
| P01872 |
| P10649 |
| P62962 |
| Q3ULD5 |
| Q9CR68 |
| Q60692 |
| P63330 |
| Q61206 |
| Q8BH69 |
| O88844 |
| Q8K2B3 |
| Q9R1P3 |
| P62141 |

| |
|---|
| Q99LC5 |
| P17742 |
| P42208 |
| P57759 |
| Q9CPY7-2 |
| P21614 |
| Q8K183 |
| Q8VCG4 |
| Q922R8 |
| P99026 |
| P24270 |
| P18760 |
| Q01853 |
| Q60864 |
| Q9CR00 |
| Q64727 |
| E9PV24 |
| P05201 |
| Q99LX0 |
| Q8BH95 |
| Q99J77 |
| Q9Z2W0 |
| P17563 |
| Q60710 |
| Q60631 |
| Q9D2G2 |
| Q99KV1 |
| Q9CQA3 |
| P01027 |
| P17751 |
| Q6P1F6 |
| Q3U0V1 |
| Q91X72 |
| Q9CR16 |
| P19157 |
| Q3UM45 |
| P63242 |
| P07759 |
| O08553 |
| Q91W90 |
| Q9Z1Z2 |
| Q9JMH6 |
| Q8VCM7 |
| O08807 |
| P63101 |
| P24452 |
| Q9JHR7 |

| |
|---|
| P58252 |
| Q9D1E6 |
| Q64010 |
| Q62422 |
| O55131 |
| P85094 |
| Q61316 |
| Q9QUM9 |
| O55023 |
| Q9CQ92 |
| P63158 |
| P06330 |
| Q9CQI3 |
| P51859 |
| P70670 |
| Q9D7X3 |
| Q11136 |
| Q8BUV3 |
| P09405 |
| P15105 |
| Q05816 |
| Q9DCZ1 |
