## Supplemental Table 6 for "Dietary restriction transforms the protein sulfhydrome in a tissue-specific and cystathionine γ-lyase-dependent manner"

| Supplemental Table 6: Sulfhydrated Protein Pathway Enrichment in Liver | | | |
|---|---|---|---|
| KEGG Pathway (DR) | p-val (adj) | -LOG10(p-val adj) | Involved Genes/Proteins |
| Metabolic pathways | 0.0345 | 1.46 | NDUFS6,HSD17B10,ACADL,NDUFA8,HMGCS2,UROD,ACSM1,PKLR,CES1C,PTGES3 |
| KEGG Pathway (Unchanged) | p-val (adj) | -LOG10(p-val adj) | Involved Genes/Proteins |
| Pentose phosphate pathway | 0.00639 | 2.19 | ALDOC,TKT,RGN,TALDO1,ALDOB,RBKS,PGM1,GPI,FBP1 |
| Bacterial invasion of epithelial cells | 0.00658 | 2.18 | CRKL,ARPC2,CDC42,RHOA,ARPC5,CRK,VCL,CLTA,ARPC3,ARPC1A,ARPC1B,CTTN,ACTG1,ARPC4 |
| Peroxisome | 0.000000196 | 6.71 | SOD2,ACAA1B,PIPOX,ACOX1,HACL1,EPHX2,AMACR,SOD1,XDH,HSD17B4,PRDX5,IDH1,AGXT,PHYH,CAT,HAO1,SCP2,HMGCL,PRDX1,NUDT7,ACAA1A,ECH1 |
| Glycine, serine and threonine metabolism | 0.00000344 | 5.46 | GNMT,SARDH,PGAM1,CHDH,PIPOX,SHMT1,DLD,CBS,GLDC,AGXT,CTH,GRHPR,DMGDH,ALDH7A1 |
| Glutathione metabolism | 0.00000136 | 5.87 | GSTM3,SRM,MGST1,GPX3,OPLAH,GSTO1,GSTA3,IDH1,GSS,TXNDC12,GSR,GCLC,LAP3,GSTM2,GSTM1,GPX1,GSTA1,GPX4 |
| Lysine degradation | 0.00156 | 2.81 | GCDH,DLST,PIPOX,OGDH,ACAT2,ECHS1,HADHA,ALDH9A1,HADH,ALDH2,ACAT1,ALDH7A1,TMLHE |
| Protein processing in endoplasmic reticulum | 0.00447 | 2.35 | PRKCSH,RAD23A,CALR,DNAJB11,UFD1,HSPA8,CANX,PDIA6,EIF2S1,HSP90AB1,P4HB,PDIA4,HSPA5,PDIA3,RRBP1,VCP,ERP29,SEC13,SEC31A,SKP1,TXNDC5,NPLOC4,GANAB |
| Glycolysis / Gluconeogenesis | 0.0000141 | 4.85 | PGAM1,ALDOC,DLD,TPI1,ALDH9A1,ALDOB,PGM1,ALDH2,GALM,GPI,ALDH7A1,GAPDH,PGK1,LDHA,ENO1,FBP1,ADH1 |
| Arginine and proline metabolism | 0.0000883 | 4.05 | CKB,SRM,ARG1,PYCR3,CNDP2,HOGA1,GOT1,ALDH9A1,ALDH2,OAT,GOT2,LAP3,AGMAT,ALDH7A1 |
| Histidine metabolism | 0.00466 | 2.33 | AMDHD1,HAL,ASPA,CNDP2,ALDH9A1,ALDH2,UROC1,ALDH7A1 |
| Fatty acid degradation | 0.00000553 | 5.26 | GCDH,ACAA1B,ACOX1,ACAT2,ECHS1,HADHA,ALDH9A1,HADH,ALDH2,ACAT1,ACAA1A,ACAA2,ALDH7A1,ACADM,ADH1 |
| Biosynthesis of amino acids | 4.29E-14 | 13.37 | PGAM1,ALDOC,ARG1,SHMT1,TKT,ACO2,GPT,PYCR3,ACY1,TPI1,CBS,GOT1,TALDO1,ASL,IDH1,CPS1,GLUL,CTH,ALDOB,OTC,GOT2,GPT2,MAT1A,MAT2B,GAPDH,PGK1,ENO1,ASS1 |
| Nitrogen metabolism | 0.0238 | 1.62 | GLUD1,CA5A,CPS1,GLUL,CA3,CA2 |
| Folate biosynthesis | 0.047 | 1.33 | MOCS2,QDPR,PCBD1,GCH1,GPHN,CBR1,MOCS1 |
| Citrate cycle (TCA cycle) | 0.000647 | 3.19 | DLST,SDHB,MDH2,MDH1,OGDH,DLD,SDHA,ACO2,IDH1,FH |
| Tryptophan metabolism | 1.02E-09 | 8.99 | HAAO,INMT,GCDH,OGDH,ACAT2,ECHS1,HADHA,ALDH9A1,KYNU,CAT,HADH,ALDH2,ACAT1,KYAT1,KYAT3,ALDH7A1,AOX1,AOX3 |
| Arginine biosynthesis | 3.28E-09 | 8.48 | ARG1,GLUD1,GPT,ACY1,GOT1,ASL,CPS1,GLUL,OTC,GOT2,GPT2,ASS1 |
| Selenocompound metabolism | 0.000268 | 3.57 | INMT,TXNRD1,SCLY,SEPHS1,CTH,KYAT1,KYAT3,SEPHS2 |
| Proteasome | 0.00000155 | 5.81 | PSMD4,PSMB1,PSMA2,PSMB6,PSMA6,PSME1,PSMB8,PSMB7,PSMB2,PSMA1,PSMA4,PSMA3,PSMA5,PSMB3,PSME2 |
| Glyoxylate and dicarboxylate metabolism | 2.75E-12 | 11.56 | MDH2,MDH1,SHMT1,DLD,ACO2,ACAT2,MUT,GLDC,HOGA1,AGXT,GLUL,CAT,HAO1,ACAT1,PCCB,MCEE,GRHPR |
| Alanine, aspartate and glutamate metabolism | 0.000000338 | 6.47 | ASPA,GLUD1,GPT,NIT2,GOT1,ASL,CPS1,AGXT,GLUL,GOT2,GPT2,ALDH5A1,ABAT,ASS1 |
| 2-Oxocarboxylic acid metabolism | 0.00507 | 2.29 | ACO2,GPT,ACY1,GOT1,IDH1,GOT2,GPT2 |
| Metabolic pathways | 5.94E-33 | 32.23 | COX5A,HAAO,CKB,PAFAH1B2,GCDH,DLST,SRM,SARDH,SDHB,ACAA1B,APIP,PGAM1,NDUFA2,MOCS2,QDPR,LTA4H,AMDHD1,CHDH,ALDOC,PIPOX,PNPO,MDH2,ETNPPL,ARG1,HAL,PCBD1,MDH1,OGDH,SHMT1,ADI1,DLD,ASPA,ACOX1,NME2,POLR2H,PYGL,ALDH6A1,AUH,SDHA,MCCC2,GLUD1,TKT,EPHX2,CMBL,AMACR,DPYS,ACO2,GPT,OPLAH,PYCR3,PMM2,CPOX,HGD,CSAD,RGN,ACY1,TPI1,ACAT2,MUT,CBS,XDH,NDUFV2,HSD17B4,CNDP2,ALDH1A7,GLDC,NT5C2,HOGA1,GOT1,ECHS1,URAH,TALDO1,ASL,GUSB,HPRT1,HADHA,IDH1,NDUFS1,CPS1,AGXT,SCLY,GLUL,FH,BPNT1,SEPHS1,ALDH9A1,PRDX6,KYNU,SORD,HAO1,IMPA1,AHCY,GSS,HADH,CTH,UOX,ALDOB,NANS,ALAD,SCP2,HMGCL,CMPK1,AK2,KHK,PGM1,UGDH,PAICS,HPD,ALDH2,HIBADH,FAH,OAT,OTC,GOT2,GPT2,CES1F,ACAT1,GCLC,ME1,PCCB,PDXK,DPYD,UPB1,MCEE,TKFC,UROC1,ATP5H,GALM,GRHPR,ALDH5A1,GALT,ACAA1A,GPI,ACAA2,GCH1,NME1,MAT1A,NDUFV1,UQCRFS1,ADK,KYAT1,LAP3,NDUFB10,KYAT3,BLVRB,AGMAT,NDUFA7,MAT2B,DMGDH,BHMT2,ACP1,TST,GPHN,SEPHS2,SUOX,CBR1,ACOT4,ALDH1A1,ALDH7A1,CES1D,GAPDH,ABAT,IDI1,GDA,FDPS,PGK1,IMPDH2,ACADM,MTAP,LDHA,ENO1,AOX1,PPCDC,MOCS1,AOX3,FBP1,GANAB,NME3,ADH1,URAD,ASS1,HMGCS1,PNP |
| Pyruvate metabolism | 7.12E-08 | 7.15 | MDH2,MDH1,DLD,ACAT2,GLO1,HAGH,FH,ALDH9A1,ALDH2,LDHD,ACAT1,ME1,GRHPR,ALDH7A1,LDHA |
| beta-Alanine metabolism | 0.00000104 | 5.98 | SRM,ALDH6A1,DPYS,CNDP2,ECHS1,HADHA,ALDH9A1,ALDH2,DPYD,UPB1,ALDH7A1,ABAT,ACADM |
| Drug metabolism - other enzymes | 0.000000247 | 6.61 | GSTM3,MGST1,NME2,DPYS,XDH,GSTO1,GUSB,HPRT1,GSTA3,CMPK1,CES1F,CES2E,DPYD,UPB1,NME1,GSTM2,CES1D,GSTM1,CES2C,IMPDH2,NME3,GSTA1 |
| Cysteine and methionine metabolism | 3.73E-09 | 8.43 | SRM,APIP,MDH2,MDH1,ADI1,CBS,GOT1,AHCY,GSS,CTH,GOT2,GCLC,MAT1A,MAT2B,BHMT2,TST,MTAP,LDHA |
| Tyrosine metabolism | 0.0319 | 1.50 | HGD,GOT1,HPD,FAH,GOT2,MIF,AOX1,AOX3,ADH1 |
| Propanoate metabolism | 0.000000415 | 6.38 | ECHDC1,DLD,ALDH6A1,ACAT2,MUT,ECHS1,HADHA,ACAT1,PCCB,MCEE,ABAT,ACADM,LDHA |
| Carbon metabolism | 3.53E-24 | 23.45 | DLST,SDHB,PGAM1,ALDOC,MDH2,MDH1,OGDH,SHMT1,DLD,ALDH6A1,SDHA,GLUD1,TKT,ESD,ACO2,GPT,RGN,TPI1,ACAT2,MUT,GLDC,GOT1,ECHS1,TALDO1,HADHA,IDH1,CPS1,AGXT,FH,CAT,HAO1,ALDOB,GOT2,GPT2,ACAT1,ME1,PCCB,MCEE,TKFC,GPI,GAPDH,PGK1,ACADM,ENO1,FBP1 |
| Valine, leucine and isoleucine degradation | 2.05E-15 | 14.69 | ACAA1B,DLD,ALDH6A1,AUH,MCCC2,ACAT2,MUT,ECHS1,HADHA,ALDH9A1,HADH,HMGCL,ALDH2,HIBADH,ACAT1,PCCB,MCEE,ACAA1A,ACAA2,ALDH7A1,ABAT,ACADM,AOX1,AOX3,HMGCS1 |
| Complement and coagulation cascades | 0.0386 | 1.41 | C8G,VTN,KNG1,C3,SERPINC1,FGA,FGB,FGG,C8A,MBL1,PLG,SERPINA1A,SERPINA1D,SERPINA1B |
| Butanoate metabolism | 0.00105 | 2.98 | ACAT2,ECHS1,HADHA,HADH,HMGCL,ACAT1,ALDH5A1,ABAT,HMGCS1 |
| Fructose and mannose metabolism | 0.0106 | 1.97 | ALDOC,PMM2,TPI1,PFKFB1,SORD,ALDOB,KHK,TKFC,FBP1 |
| KEGG Pathway (AL) | p-val (adj) | -LOG10(p-val adj) | Involved Genes/Proteins |
| Terpenoid backbone biosynthesis | 0.0241 | 1.62 | FNTA,FNTB |
| Selenocompound metabolism | 0.016 | 1.80 | TXNRD3,SEPSECS |
