## Supplemental Table 7 for "Dietary restriction transforms the protein sulfhydrome in a tissue-specific and cystathionine γ-lyase-dependent manner"

| Supplemental Table 7: Sulfhydrated Protein Pathway Enrichment in Kidney | | | |
|---|---|---|---|
| KEGG Pathway (DR) | p-val (adj) | -LOG10(p-val adj) | Involved Genes |
| Nitrogen metabolism | 0.0248 | 1.61 | CA3,CA5B |
| KEGG Pathway (Unchanged) | p-val (adj) | -LOG10(p-val adj) | Involved Genes |
| Butanoate metabolism | 6.49E-12 | 11.19 | OXCT1,EHHADH,ACAT2,ECHS1,HADH,BDH2,HMGCL,ACADS,ACSM3,ACSM2,ACSM5,ACAT1,ACSM1,ALDH5A1,BDH1,ABAT,HMGCS1 |
| Metabolic pathways | 5.12E-50 | 49.29 | COX5A,DLAT,CKMT1,DBT,NDUFA9,HAAO,CKB,AKR1B1,FOLH1,GGCT,PLD3,GCDH,DLST,CS,ATP6V1B2,GGT1,SARDH,SDHB,ALDH3A2,ACAA1B,PLCD1,APIP,PDHX,PGAM1,NDUFS2,NDUFA2,QDPR,LTA4H,CHDH,ALDOC,PIPOX,ACADVL,MDH2,ATP6V1E1,PAH,PCBD1,NDUFS7,MDH1,OGDH,PGAM2,SHMT1,DLD,PAFAH1B1,ASPA,ACOX1,NME2,POLR2H,GSTZ1,ALDH6A1,AUH,SDHA,MCCC2,PDHB,ACOX2,NDUFS4,OXSM,GLUD1,CRYL1,TKT,EPHX2,SUCLA2,CMBL,AMACR,DPYS,NDUFB9,ACO2,GPT,CYC1,CPOX,HGD,EHHADH,CSAD,ACY1,TPI1,ACAT2,MUT,CBS,NDUFV2,HSD17B4,CNDP2,ALDH1A7,GLDC,NT5C2,GK,HOGA1,GOT1,HSD17B10,ECHS1,TALDO1,ASL,GUSB,HPRT1,ATP5C1,IDH1,NDUFS1,ACADL,NDUFA10,SCLY,ACMSD,GLUL,FH,EPRS,BPNT1,SEPHS1,ALDH9A1,PRDX6,AK1,NDUFA8,UAP1L1,SORD,IVD,PCK1,AHCY,GSS,NFS1,HAO2,AHCYL1,HADH,GCLM,BDH2,CTH,ALDOB,NANS,ALAD,ACO1,AK4,SCP2,PPT1,HMGCL,AKR1A1,CMPK1,ALDH4A1,CDA,ALPL,AK2,ACOX3,KHK,PGM1,UGDH,PAICS,HPD,ALDH2,ACADS,MMAB,AHCYL2,RPN1,LDHB,BCAT1,IDH2,FAH,BCAT2,UQCRC2,OAT,ACSM3,ACSM2,ACSM5,PDHA1,GOT2,CES1F,COX4I1,ACAT1,PTS,TREH,IDH3A,PKM,GCLC,ME1,PCCB,PDXK,DPYD,UPB1,ACSM1,ATP6V1H,TKFC,ATP5H,NAGK,GALM,GRHPR,ALDH5A1,GPI,COX6B1,ACAA2,NME1,NDUFV1,GMDS,UQCRFS1,GCNT1,ATP6V1G1,ADK,DCXR,KYAT1,LAP3,NDUFB10,KYAT3,BLVRB,PCCA,MAT2B,DAO,DMGDH,PGP,TST,FAHD1,BDH1,GPHN,SEPHS2,SUOX,GNPDA1,ACOT4,ATP6V1A,SUCLG1,ALDH7A1,CES1D,AADAT,CES1C,GAPDH,ABAT,HADHB,NDUFS8,COX5B,SUCLG2,PGK1,IMPDH2,ACADM,MTAP,ENO1,UQCRH,MTCO2,FBP1,PTGES3,GANAB,MPST,ACOT1,HSD17B8,NME3,ADH1,ASS1,HMGCS1,PNP |
| Arginine biosynthesis | 0.00202 | 2.69 | GLUD1,GPT,ACY1,GOT1,ASL,GLUL,GOT2,ASS1 |
| Alzheimer disease | 0.000385 | 3.41 | COX5A,NDUFA9,SDHB,NDUFS2,NDUFA2,NDUFS7,SDHA,NDUFS4,NDUFB9,CYC1,NDUFV2,HSD17B10,ATP5C1,NDUFS1,NDUFA10,NDUFA8,UQCRC2,FADD,COX4I1,ATP5H,COX6B1,NDUFV1,UQCRFS1,NDUFB10,GAPDH,NDUFS8,COX5B,CYCS,UQCRH,MTCO2 |
| Selenocompound metabolism | 0.00124 | 2.91 | TXNRD1,SCLY,SEPHS1,CTH,SEPSECS,KYAT1,KYAT3,SEPHS2 |
| Biosynthesis of amino acids | 1.06E-14 | 13.97 | CS,PGAM1,ALDOC,PAH,PGAM2,SHMT1,TKT,ACO2,GPT,ACY1,TPI1,CBS,GOT1,TALDO1,ASL,IDH1,GLUL,CTH,ALDOB,ACO1,BCAT1,IDH2,BCAT2,GOT2,IDH3A,PKM,MAT2B,GAPDH,PGK1,ENO1,ASS1 |
| Tryptophan metabolism | 0.000000264 | 6.58 | HAAO,GCDH,ALDH3A2,OGDH,EHHADH,ACAT2,ECHS1,ACMSD,ALDH9A1,CAT,HADH,ALDH2,ACAT1,KYAT1,KYAT3,ALDH7A1,AADAT |
| Propanoate metabolism | 2.51E-09 | 8.60 | DBT,DLD,ALDH6A1,SUCLA2,EHHADH,ACAT2,MUT,ECHS1,LDHB,ACAT1,PCCB,PCCA,SUCLG1,ABAT,SUCLG2,ACADM |
| Sulfur metabolism | 0.0168 | 1.77 | BPNT1,TST,SUOX,ETHE1,MPST |
| Drug metabolism - other enzymes | 0.0000446 | 4.35 | GSTM5,NME2,DPYS,GUSB,HPRT1,GSTA3,CMPK1,CDA,CES1F,DPYD,GSTT2,UPB1,NME1,GSTM2,CES1D,CES1C,GSTM1,CES2C,IMPDH2,NME3,GSTA1 |
| Huntington disease | 0.0000137 | 4.86 | COX5A,NDUFA9,SOD2,SDHB,NDUFS2,NDUFA2,SLC25A5,NDUFS7,VDAC1,POLR2H,SDHA,NDUFS4,NDUFB9,CYC1,SOD1,NDUFV2,ATP5C1,NDUFS1,NDUFA10,NDUFA8,CLTA,UQCRC2,COX4I1,ATP5H,COX6B1,TGM2,NDUFV1,UQCRFS1,NDUFB10,NDUFS8,COX5B,CYCS,GPX1,UQCRH,MTCO2 |
| Proximal tubule bicarbonate reclamation | 0.00711 | 2.15 | AQP1,MDH1,GLUD1,SLC25A10,ATP1B1,PCK1,CA2,ATP1A1 |
| Valine, leucine and isoleucine degradation | 3.96E-19 | 18.40 | DBT,ALDH3A2,ACAA1B,DLD,ALDH6A1,AUH,MCCC2,OXCT1,EHHADH,ACAT2,MUT,HSD17B10,ECHS1,ALDH9A1,IVD,HADH,HMGCL,ALDH2,ACADS,BCAT1,BCAT2,ACAT1,PCCB,ACAA2,PCCA,ALDH7A1,ABAT,HADHB,ACADM,HMGCS1 |
| Glyoxylate and dicarboxylate metabolism | 1.74E-13 | 12.76 | CS,MDH2,MDH1,SHMT1,DLD,ACO2,ACAT2,MUT,GLDC,HOGA1,GLUL,CAT,HAO2,ACO1,ACAT1,PCCB,GRHPR,PCCA,PGP |
| Citrate cycle (TCA cycle) | 1.81E-15 | 14.74 | DLAT,DLST,CS,SDHB,MDH2,MDH1,OGDH,DLD,SDHA,PDHB,SUCLA2,ACO2,IDH1,FH,PCK1,ACO1,IDH2,PDHA1,IDH3A,SUCLG1,SUCLG2 |
| Peroxisome | 3.24E-11 | 10.49 | CROT,SOD2,ACAA1B,PIPOX,ACOX1,ECI2,ACOX2,HACL1,EPHX2,AMACR,EHHADH,SOD1,HSD17B4,PRDX5,IDH1,PECR,PHYH,CAT,SLC27A2,HAO2,SCP2,HMGCL,PRDX1,ACOX3,GSTK1,IDH2,NUDT19,DAO,ECH1 |
| beta-Alanine metabolism | 0.0000117 | 4.93 | ALDH3A2,ALDH6A1,DPYS,EHHADH,CNDP2,ECHS1,ALDH9A1,ALDH2,DPYD,UPB1,ALDH7A1,ABAT,ACADM |
| Bacterial invasion of epithelial cells | 0.00102 | 2.99 | CDH1,CRKL,CDC42,CTNNB1,RHOA,ARPC5,CRK,VCL,ITGB1,ARPC5L,CLTA,ARPC3,ACTB,ARPC1A,ARPC1B,CTNNA1,ARPC4 |
| Proteasome | 0.00000314 | 5.50 | PSMD4,PSMB4,PSMB1,PSMA2,PSMB6,PSMA6,PSMB5,PSME1,PSMB7,PSMA7,PSMB2,PSMA1,PSMA4,PSMA3,PSMA5,PSMB3 |
| Pyruvate metabolism | 2.73E-13 | 12.56 | DLAT,ALDH3A2,MDH2,MDH1,DLD,PDHB,ACAT2,GLO1,HAGH,FH,ALDH9A1,PCK1,ALDH2,LDHB,PDHA1,LDHD,ACAT1,PKM,ME1,GRHPR,ALDH7A1 |
| Synthesis and degradation of ketone bodies | 0.000201 | 3.70 | OXCT1,ACAT2,BDH2,HMGCL,ACAT1,BDH1,HMGCS1 |
| Cysteine and methionine metabolism | 1.17E-09 | 8.93 | APIP,MDH2,MDH1,CBS,GOT1,AHCY,GSS,AHCYL1,GCLM,CTH,AHCYL2,LDHB,BCAT1,BCAT2,GOT2,GCLC,MAT2B,TST,MTAP,MPST |
| Arginine and proline metabolism | 0.000967 | 3.01 | CKMT1,CKB,ALDH3A2,CNDP2,HOGA1,GOT1,ALDH9A1,ALDH4A1,ALDH2,OAT,GOT2,LAP3,DAO,ALDH7A1 |
| Glycine, serine and threonine metabolism | 0.000043 | 4.37 | SARDH,PGAM1,CHDH,PIPOX,PGAM2,SHMT1,DLD,CBS,GLDC,CTH,GRHPR,DAO,DMGDH,ALDH7A1 |
| Protein processing in endoplasmic reticulum | 0.000632 | 3.20 | PRKCSH,CALR,DNAJB11,HSPA8,HSP90B1,CANX,PDIA6,EIF2S1,HSP90AA1,HSP90AB1,P4HB,PDIA4,HSPA5,PDIA3,RRBP1,BAG1,VCP,ERP29,HSPH1,RPN1,SEC13,DNAJA2,HYOU1,SEC31A,SKP1,TXNDC5,GANAB,HSPA1B |
| Glutathione metabolism | 0.000000881 | 6.06 | GGCT,GSTM5,GGT1,GPX3,GSTA3,IDH1,GSS,GCLM,TXNDC12,GSTK1,IDH2,GSR,GCLC,GSTT2,LAP3,GSTM2,GSTM1,GPX1,GSTA1,GPX4 |
| Complement and coagulation cascades | 0.00254 | 2.60 | C8G,CLU,KNG1,C3,SERPINC1,FGA,C8B,FGB,FGG,C8A,MBL1,CFI,PLG,SERPINA1A,SERPINA1D,SERPINA1B,SERPINA1E,C4B |
| Folate biosynthesis | 0.0272 | 1.57 | AKR1B1,QDPR,PAH,PCBD1,ALPL,PTS,MOCOS,GPHN |
| Fatty acid metabolism | 0.00000046 | 6.34 | ACAA1B,ACADVL,ACOX1,OXSM,EHHADH,ACAT2,ECHS1,ACADL,PECR,HADH,CPT2,PPT1,ACOX3,ACADS,ACAT1,ACAA2,HADHB,ACADM |
| Glycolysis / Gluconeogenesis | 8.7E-11 | 10.06 | DLAT,ALDH3A2,PGAM1,ALDOC,PGAM2,DLD,PDHB,TPI1,ALDH9A1,PCK1,ALDOB,AKR1A1,PGM1,ALDH2,LDHB,PDHA1,PKM,GALM,GPI,ALDH7A1,GAPDH,PGK1,ENO1,FBP1,ADH1 |
| Oxidative phosphorylation | 1.15E-08 | 7.94 | COX5A,NDUFA9,ATP6V1B2,SDHB,NDUFS2,NDUFA2,ATP6V1E1,PPA1,NDUFS7,SDHA,NDUFS4,NDUFB9,CYC1,NDUFV2,ATP5C1,NDUFS1,NDUFA10,NDUFA8,PPA2,UQCRC2,COX4I1,ATP6V1H,ATP5H,COX6B1,NDUFV1,UQCRFS1,ATP6V1G1,NDUFB10,ATP6V1A,NDUFS8,COX5B,UQCRH,MTCO2 |
| Fructose and mannose metabolism | 0.05 | 1.30 | AKR1B1,ALDOC,TPI1,SORD,ALDOB,KHK,TKFC,GMDS,FBP1 |
| Non-alcoholic fatty liver disease (NAFLD) | 0.000489 | 3.31 | COX5A,NDUFA9,CDC42,SDHB,NDUFS2,NDUFA2,NDUFS7,EIF2S1,SDHA,NDUFS4,NDUFB9,CYC1,NDUFV2,NDUFS1,NDUFA10,NDUFA8,UQCRC2,COX4I1,COX6B1,NDUFV1,UQCRFS1,NDUFB10,NDUFS8,COX5B,CYCS,UQCRH,MTCO2 |
| 2-Oxocarboxylic acid metabolism | 1.36E-09 | 8.87 | CS,ACO2,GPT,ACY1,GOT1,IDH1,ACO1,BCAT1,IDH2,BCAT2,GOT2,IDH3A,AADAT |
| PPAR signaling pathway | 0.00154 | 2.81 | CD36,ACAA1B,FABP7,ACOX1,ACOX2,EHHADH,GK,ACADL,SLC27A2,PCK1,FABP5,SCP2,CPT2,ACOX3,ME1,FABP1,FABP4,ACADM |
| Parkinson disease | 0.0000052 | 5.28 | COX5A,NDUFA9,SDHB,NDUFS2,NDUFA2,SLC25A5,NDUFS7,VDAC1,SDHA,NDUFS4,NDUFB9,CYC1,NDUFV2,ATP5C1,NDUFS1,NDUFA10,NDUFA8,PARK7,UQCRC2,COX4I1,ATP5H,COX6B1,NDUFV1,UQCRFS1,NDUFB10,NDUFS8,COX5B,CYCS,UQCRH,MTCO2 |
| Lysine degradation | 0.000614 | 3.21 | GCDH,DLST,ALDH3A2,PIPOX,OGDH,EHHADH,ACAT2,ECHS1,ALDH9A1,HADH,ALDH2,ACAT1,ALDH7A1,AADAT,TMLHE |
| Alanine, aspartate and glutamate metabolism | 0.0000389 | 4.41 | FOLH1,ASPA,GLUD1,GPT,NIT2,GOT1,ASL,GLUL,ALDH4A1,GOT2,ALDH5A1,ABAT,ASS1 |
| Fatty acid degradation | 2.33E-12 | 11.63 | GCDH,ALDH3A2,ACAA1B,ACADVL,ACOX1,ECI2,EHHADH,ACAT2,ECI1,ECHS1,ACADL,ALDH9A1,HADH,CPT2,ACOX3,ALDH2,ACADS,ACAT1,ACAA2,ALDH7A1,HADHB,ACADM,ADH1 |
| Thermogenesis | 0.0141 | 1.85 | COX5A,NDUFA9,SDHB,NDUFS2,NDUFA2,NDUFS7,SDHA,NDUFS4,NDUFB9,CYC1,NDUFAF7,NDUFV2,ATP5C1,NDUFS1,NDUFA10,NDUFA8,ACTL6A,CPT2,ACTB,UQCRC2,COX4I1,ATP5H,COX6B1,NDUFV1,UQCRFS1,NDUFB10,COA7,NDUFS8,GRB2,COX5B,UQCRH,MTCO2 |
| Carbon metabolism | 8.76E-31 | 30.06 | DLAT,DLST,CS,SDHB,PGAM1,ALDOC,MDH2,MDH1,OGDH,PGAM2,SHMT1,DLD,ALDH6A1,SDHA,PDHB,GLUD1,TKT,ESD,SUCLA2,ACO2,GPT,EHHADH,TPI1,ACAT2,MUT,GLDC,GOT1,ECHS1,TALDO1,IDH1,FH,CAT,HAO2,ALDOB,ACO1,ACADS,IDH2,PDHA1,GOT2,ACAT1,IDH3A,PKM,ME1,PCCB,TKFC,GPI,PCCA,PGP,SUCLG1,GAPDH,SUCLG2,PGK1,ACADM,ENO1,FBP1 |
| KEGG Pathway (AL) | p-val (adj) | -LOG10(p-val adj) | Involved Genes |
| Glycine, serine and threonine metabolism | 0.00748 | 2.13 | GNMT,BHMT |
