## Supplemental Table 8 for "Dietary restriction transforms the protein sulfhydrome in a tissue-specific and cystathionine γ-lyase-dependent manner"

| Supplemental Table 8: Sulfhydrated Protein Pathway Enrichment in Muscle | | | |
|---|---|---|---|
| KEGG Pathway (DR) | p-val (adj) | -LOG10(p-val adj) | Involved Gene |
| No KEGG | | | |
| KEGG Pathway (Unchanged) | p-val (adj) | -LOG10(p-val adj) | Involved Gene |
| Metabolic pathways | 0.0000139 | 4.86 | AKR1B1,DLST,PDHX,NDUFA2,MDH1,PHYKPL,PGAM2,SHMT1,DLD,NME2,SDHA,NDUFS6,CKMT2,MCCC2,TKT,CMBL,TPI1,GOT1,ECHS1,TALDO1,NDUFS1,ACADL,FH,SEPHS1,PRDX6,AK1,IMPA1,AHCY,HADH,NANS,ALAD,CMPK1,PGM1,ALDH2,LDHB,CKM,BCAT2,PDHA1,GOT2,GRHPR,ALDH5A1,NDUFV1,LAP3,MAT2B,PGP,ACP1,SEPHS2,ALDH7A1,CES1D,GAPDH,ENO3,COX5B,PGK1,LDHA |
| Cysteine and methionine metabolism | 0.000397 | 3.40 | MDH1,GOT1,AHCY,LDHB,BCAT2,GOT2,MAT2B,LDHA |
| Glyoxylate and dicarboxylate metabolism | 0.00223 | 2.65 | MDH1,SHMT1,DLD,CAT,GRHPR,PGP |
| Biosynthesis of amino acids | 0.00000369 | 5.43 | PGAM2,SHMT1,TKT,TPI1,GOT1,TALDO1,BCAT2,GOT2,MAT2B,GAPDH,ENO3,PGK1 |
| Valine, leucine and isoleucine degradation | 0.00977 | 2.01 | DLD,MCCC2,ECHS1,HADH,ALDH2,BCAT2,ALDH7A1 |
| Hypertrophic cardiomyopathy (HCM) | 0.0317 | 1.50 | CACNB1,LMNA,ACTB,TPM1,TGFB2,MYH6,TTN,MYL3 |
| Parkinson disease | 0.016 | 1.80 | NDUFA2,VDAC1,SDHA,NDUFS6,NDUFS1,PARK7,SLC25A4,NDUFV1,UBE2L3,COX5B,CYCS |
| Carbon metabolism | 1.87E-10 | 9.73 | DLST,MDH1,PGAM2,SHMT1,DLD,SDHA,TKT,TPI1,GOT1,ECHS1,TALDO1,FH,CAT,PDHA1,GOT2,PGP,GAPDH,ENO3,PGK1 |
| Glycolysis / Gluconeogenesis | 0.000000736 | 6.13 | PGAM2,DLD,TPI1,PGM1,ALDH2,LDHB,PDHA1,ALDH7A1,GAPDH,ENO3,PGK1,LDHA |
| Proteasome | 2.85E-10 | 9.55 | PSMD4,PSMB4,PSMB1,PSMA2,PSMB6,PSMA6,PSMB5,PSMA7,PSMB2,PSMA1,PSMA3,PSMA5,PSMB3 |
| Citrate cycle (TCA cycle) | 0.00329 | 2.48 | DLST,MDH1,DLD,SDHA,FH,PDHA1 |
| Complement and coagulation cascades | 0.0342 | 1.47 | FGA,FGB,FGG,CFI,SERPINA1A,SERPINA1D,SERPINA1B,SERPINA1E |
| Arginine and proline metabolism | 0.00768 | 2.11 | CKMT2,GOT1,ALDH2,CKM,GOT2,LAP3,ALDH7A1 |
| Pyruvate metabolism | 1.07E-08 | 7.97 | MDH1,DLD,GLO1,FH,ALDH2,LDHB,PDHA1,GRHPR,ALDH7A1,ACYP2,LDHA |
| Dilated cardiomyopathy (DCM) | 0.043 | 1.37 | CACNB1,LMNA,ACTB,TPM1,TGFB2,MYH6,TTN,MYL3 |
| KEGG Pathway (AL) | p-val (adj) | -LOG10(p-val adj) | Involved Gene |
| No KEGG | | | |
