## Supplemental Table 9 for "Dietary restriction transforms the protein sulfhydrome in a tissue-specific and cystathionine γ-lyase-dependent manner"

| Supplemental Table 9: Sulfhydrated Protein Pathway Enrichment in Brain | | | |
|---|---|---|---|
| KEGG Pathway (DR) | p-val (adj) | -LOG10(p-val adj) | Involved Genes |
| Carbon metabolism | 7.87E-09 | 8.10 | IDH3G,DLST,PGAM1,ALDOC,MDH2,OGDH,PFKP,GLUD1,TKT,ACO2,ECHS1,IDH1,CAT,ACAT1,PKM,PCCB,PFKM,GPI,PHGDH,PGK1 |
| Regulation of actin cytoskeleton | 0.0102 | 1.99 | PPP1CC,CRKL,PPP1CB,PFN1,VCL,BAIAP2,ARPC5L,PFN2,ARPC1A,PAK1,RDX,EZR,ACTN4,CFL1,CFL2,MAPK1,ARPC4 |
| Synaptic vesicle cycle | 0.00658 | 2.18 | AP2A2,NAPA,STX1A,ATP6V1E1,VAMP2,AP2M1,SNAP25,STX1B,ATP6V1A |
| Pentose phosphate pathway | 0.024 | 1.62 | ALDOC,PFKP,TKT,PGM1,PFKM,GPI |
| Insulin secretion | 0.00276 | 2.56 | ATP1A2,STX1A,VAMP2,CAMK2G,SNAP25,SLC2A1,ATP1A1,ATP1A3,ATP1B2,PRKCB,PCLC |
| Biosynthesis of amino acids | 0.000129 | 3.89 | IDH3G,PGAM1,ALDOC,PFKP,TKT,ACO2,IDH1,GLUL,PKM,PFKM,PHGDH,PGK1 |
| Fc gamma R-mediated phagocytosis | 0.0144 | 1.84 | CRKL,ARPC5L,ARPC1A,PAK1,PRKCB,CFL1,CFL2,MAPK1,MARCKS,ARPC4 |
| Glyoxylate and dicarboxylate metabolism | 0.0138 | 1.86 | MDH2,ACO2,GLUL,CAT,ACAT1,PCCB |
| Citrate cycle (TCA cycle) | 0.0201 | 1.70 | IDH3G,DLST,MDH2,OGDH,ACO2,IDH1 |
| Glycolysis / Gluconeogenesis | 0.000216 | 3.67 | PGAM1,ALDOC,PFKP,AKR1A1,PGM1,ALDH2,PKM,PFKM,GPI,PGK1,LDHA |
| Proximal tubule bicarbonate reclamation | 0.00209 | 2.68 | ATP1A2,GLUD1,CA2,ATP1A1,ATP1A3,ATP1B2 |
| Oocyte meiosis | 0.00228 | 2.64 | PPP1CC,PPP1CB,YWHAB,YWHAH,PPP2CA,YWHAE,CAMK2G,YWHAZ,PPP3CA,SKP1,YWHAG,MAPK1,YWHAC |
| Gap junction | 0.0131 | 1.88 | TUBB6,TUBB5,TUBA4A,TUBB4B,TUBA1C,PRKCB,TUBB2A,TUBB3,TUBB4A,MAPK1 |
| Protein processing in endoplasmic reticulum | 0.0171 | 1.77 | PRKCSH,CALR,DNAJB11,HSPA8,PDIA6,HSP90AB1,P4HB,RAD23B,VCP,ERP29,HSPH1,SKP1,FBXO2,GANAB |
| Alanine, aspartate and glutamate metabolism | 0.0394 | 1.40 | ASPA,GLUD1,GLUL,ALDH5A1,ABAT,GAD1 |
| Butanoate metabolism | 0.00738 | 2.13 | ECHS1,ACAT1,ALDH5A1,ABAT,GAD1,HMGCS1 |
| Proteasome | 0.00312 | 2.51 | PSMB4,PSMB1,PSMB7,PSMB2,PSMA4,PSMA3,PSMA5,PSMB3 |
| Endocrine and other factor-regulated calcium reabsorption | 0.00167 | 2.78 | AP2A2,ATP1A2,ATP2B1,AP2M1,CALB1,ATP1A1,ATP1A3,ATP1B2,PRKCE |
| Metabolic pathways | 0.0157 | 1.80 | CKMT1,CKB,IDH3G,PAFAH1B2,DLST,PGAM1,QDPR,ALDOC,ISYNA1,MDH2,ATP6V1E1,OGDH,PAFAH1B1,ASPA,PFKP,GLUD1,TKT,ACO2,CYC1,SYNJ1,CNDP2,NT5C2,ECHS1,HPRT1,IDH1,GLUL,BPNT1,PRDX6,ALAD,AKR1A1,CMPK1,PGM1, PAICS,ALDH2,HIBADH,OAT,ACAT1,PKM,PCCB,PDXK,PFKM,ATP5H,ALDH5A1,GPI,ACP1,GPHN,SUOX,ATP6V1A,ALDH1A1,PHGDH,ABAT,GDA,PGK1,LDHA,GAD1,PTGES3,GANAB,HMGCS1,PDXP |
| KEGG Pathway (Unchanged) | p-val (adj) | -LOG10(p-val adj) | Involved Genes |
| Biosynthesis of amino acids | 0.0000537 | 4.27 | ENO2,CS,ACO2,GOT1,TALDO1,GOT2,MAT2B,GAPDH,ENO1 |
| Synthesis and degradation of ketone bodies | 0.0278 | 1.56 | OXCT1,HMGCL,BDH1 |
| Valine, leucine and isoleucine degradation | 0.000521 | 3.28 | DLD,ALDH6A1,AUH,MCCC2,OXCT1,HMGCL,ALDH7A1 |
| Pyruvate metabolism | 0.0000388 | 4.41 | DLAT,MDH1,DLD,PDHB,LDHB,PDHA1,ALDH7A1 |
| Arginine and proline metabolism | 0.0498 | 1.30 | CKMT2,GOT1,GOT2,LAP3,ALDH7A1 |
| Proteasome | 0.0256 | 1.59 | PSMB6,PSMA6,PSMB5,PSMA7,PSMA1 |
| Metabolic pathways | 0.000000726 | 6.14 | DLAT,NDUFA9,PLD3,ENO2,CS,SDHB,PDHX,MDH1,DLD,ALDH6A1,AUH,SDHA,CKMT2,MCCC2,PDHB,ACO2,GOT1,TALDO1,NDUFS1,ACADL,AK1,IMPA1,AHCY,GSS,HMGCL,LDHB,FAH,UQCRC2,PDHA1,GOT2,PYGB,NME1,UQCRFS1,LAP3, LVRB,MAT2B,PGP,BDH1,ALDH7A1,GAPDH,ENO1,BHMT |
| Cysteine and methionine metabolism | 0.0000127 | 4.90 | MDH1,GOT1,AHCY,GSS,LDHB,GOT2,MAT2B,BHMT |
| Carbon metabolism | 1.15E-11 | 10.94 | DLAT,ENO2,CS,SDHB,MDH1,DLD,ALDH6A1,SDHA,PDHB,ACO2,GOT1,TALDO1,PDHA1,GOT2,PGP,GAPDH,ENO1 |
| Glyoxylate and dicarboxylate metabolism | 0.00313 | 2.50 | CS,MDH1,DLD,ACO2,PGP |
| Citrate cycle (TCA cycle) | 1.35E-08 | 7.87 | DLAT,CS,SDHB,MDH1,DLD,SDHA,PDHB,ACO2,PDHA1 |
| Glycolysis / Gluconeogenesis | 0.0000162 | 4.79 | DLAT,ENO2,DLD,PDHB,LDHB,PDHA1,ALDH7A1,GAPDH,ENO1 |
| 2-Oxocarboxylic acid metabolism | 0.00763 | 2.12 | CS,ACO2,GOT1,GOT2 |
| KEGG Pathway (AL) | p-val (adj) | -LOG10(p-val adj) | Involved Genes |
| Cardiac muscle contraction | 0.0000458 | 4.34 | COX5A,ATP2A2,MYH6,MYL3,MYL4,ACTC1 |
| Butanoate metabolism | 0.0108 | 1.97 | HADHA,HADH,ACADS |
| Carbon metabolism | 0.00043 | 3.37 | PGAM2,HADHA,ACADS,SUCLG1,ENO3,FBP1 |
| Parkinson disease | 0.0175 | 1.76 | COX5A,VDAC3,VDAC1,VDAC2,NDUFA5 |
| Fatty acid metabolism | 0.000111 | 3.95 | HADHA,HADH,ACADS,ACAA2,HADHB |
| Fructose and mannose metabolism | 0.0237 | 1.63 | AKR1B1,MPI,FBP1 |
| Dilated cardiomyopathy (DCM) | 0.0000851 | 4.07 | MYBPC3,ATP2A2,MYH6,TTN,MYL3,ACTC1 |
| Metabolic pathways | 0.0000271 | 4.57 | COX5A,AKR1B1,PGAM2,DHRS4,NDUFA5,HADHA,IVD,HADH,ALDH4A1,ACADS,CKM,BCKDHB,MPI,ACAA2,SUCLG1,HADHB,ENO3,FBP1 |
| Fatty acid elongation | 0.000468 | 3.33 | HADHA,HADH,ACAA2,HADHB |
| Adrenergic signaling in cardiomyocytes | 0.021 | 1.68 | ATP2A2,MYH6,MYL3,MYL4,ACTC1 |
| Fatty acid degradation | 0.00000227 | 5.64 | ECI1,HADHA,HADH,ACADS,ACAA2,HADHB |
| Hypertrophic cardiomyopathy (HCM) | 0.0000652 | 4.19 | MYBPC3,ATP2A2,MYH6,TTN,MYL3,ACTC1 |
| Propanoate metabolism | 0.0164 | 1.79 | HADHA,BCKDHB,SUCLG1 |
| Valine, leucine and isoleucine degradation | 9.21E-08 | 7.04 | HADHA,IVD,HADH,ACADS,BCKDHB,ACAA2,HADHB |
