## Supplemental Table 10 for "Dietary restriction transforms the protein sulfhydrome in a tissue-specific and cystathionine γ-lyase-dependent manner"

| Supplemental Table 10: Dietary Impact on the Heart Sulfhydrome | | | | | | |
|---|---|---|---|---|---|---|
| Protein Name | Accession Number | Alternate ID | Molecular Weight | Cysteine Residues | DR/AL Spectral Count Ratio | P-value |
| Ig lambda-2 chain C region | P01844 | Iglc2 | 11 kDa | 3 C | 16.000 | 0.00101 |
| Gelsolin | P13020 (+1) | Gsn | 86 kDa | 7 C | 11.130 | 0.00133 |
| Glutamate--cysteine ligase regulatory subunit | O09172 | Gclm | 31 kDa | 6 C | 10.200 | 0.0307 |
| Ig gamma-3 chain C region | P03987 | | 44 kDa | 10 C | 7.636 | 0.0005 |
| Ferritin heavy chain | P09528 | Fth1 | 21 kDa | 3 C | 6.182 | 0.02617 |
| Antithrombin-III | P32261 | Serpinc1 | 52 kDa | 9 C | 5.333 | 0.03116 |
| Bisphosphoglycerate mutase | P15327 | Bpgm | 30 kDa | 3 C | 4.645 | 0.01998 |
| Vitamin D-binding protein | Q9QVP4 | Gc | 54 kDa | 28 C | 4.541 | 0.0206 |
| Properdin | P11680 | Cfp | 50 kDa | 44 C | 3.692 | 0.0227 |
| Complement factor B | P01867 (+1) | Cfb | 85 kDa | 20 C | 3.636 | 0.01126 |
| Transforming growth factor beta-1 | P04202 | Tgfb1 | 44 kDa | 12 C | 3.273 | 0.00601 |
| Ferritin light chain 1 | P29391 | Ftl1 | 21 kDa | 1 C | 3.250 | 0.0204 |
| Ig lambda-1 chain C region | Q9CPV4-2 | | 12 kDa | 3 C | 2.844 | 0.02618 |
| Kininogen-1 | Q8K182 | Kng1 | 73 kDa | 19 C | 2.840 | 0.01359 |
| Beta-2-glycoprotein 1 | Q01339 | Apoh | 39 kDa | 23 C | 2.691 | 0.00579 |
| Complement C3 | P01027 | C3 | 186 kDa | 27 C | 2.556 | 0.00991 |
| Complement factor I | P02088 | Cfi | 67 kDa | 40 C | 2.324 | 0.02636 |
| Ig heavy chain V region 102 | P01750 | | 13 kDa | 3 C | 16.200 | 0.1642 |
| Afamin | O89020 (+1) | Afm | 69 kDa | 34 C | 14.400 | 0.07963 |
| Dehydrogenase/reductase SDR family member 11 | Q3U0B3 | Dhrs11 | 28 kDa | 8 C | 10.400 | 0.09207 |
| Myosin light chain 4 | P09541 | Myl4 | 21 kDa | 2 C | 9.908 | 0.23919 |
| Myeloperoxidase | P11247 | Mpo | 81 kDa | 16 C | 8.800 | 0.40708 |
| Myosin regulatory light chain 2, skeletal muscle isoform | P97457 | Mylpf | 19 kDa | 2 C | 8.800 | 0.40708 |
| Plasminogen | P20918 | Plg | 91 kDa | 48 C | 8.800 | 0.40708 |
| Filamin-A | Q8BTM8 | Flna | 281 kDa | 38 C | 8.600 | 0.19301 |
| Fetuin-B | Q9QXC1 | Fetub | 43 kDa | 15 C | 8.400 | 0.10695 |
| Fatty acid-binding protein, epidermal | Q05816 | Fabp5 | 15 kDa | 6 C | 8.400 | 0.10695 |
| Major urinary protein 17 | B5X0G2 (+2) | Mup17 | 21 kDa | 5 C | 6.800 | 0.40708 |
| Na(+)/H(+) exchange regulatory cofactor NHE-RF2 | Q9JHL1 | Slc9a3r2 | 37 kDa | 6 C | 6.600 | 0.23224 |
| Myosin light chain 1/3, skeletal muscle isoform | P05977 | Myl1 | 21 kDa | 2 C | 6.338 | 0.26168 |
| Complement factor H | P06909 | Cfh | 139 kDa | 82 C | 5.913 | 0.15149 |
| Murinoglobulin-1 | P28665 | Mug1 | 165 kDa | 25 C | 4.904 | 0.05584 |
| Crk-like protein | Q08857 | Crkl | 34 kDa | 2 C | 4.800 | 0.40708 |
| Platelet glycoprotein 4 | P47941 | Cd36 | 53 kDa | 10 C | 4.800 | 0.40708 |
| Carbonic anhydrase 3 | P16015 | Ca3 | 29 kDa | 5 C | 4.800 | 0.40708 |
| Myotilin | Q9JIF9 | Myot | 55 kDa | 8 C | 4.800 | 0.40708 |
| Myosin regulatory light chain 2, atrial isoform | P21614 | Myl7 | 19 kDa | 3 C | 4.492 | 0.41732 |
| Hepatocyte growth factor activator | Q9R098 | Hgfac | 71 kDa | 41 C | 3.879 | 0.05793 |
| Clusterin | Q06890 | Clu | 52 kDa | 11 C | 3.815 | 0.14846 |
| Mannose-binding protein C | P41317 | Mbl2 | 26 kDa | 7 C | 3.754 | 0.08807 |
| Ig gamma-2B chain C region | P04186 | Igh-3 | 44 kDa | 13 C | 3.636 | 0.15411 |
| Methylmalonyl-CoA mutase, mitochondrial | P16332 | Mut | 83 kDa | 8 C | 3.262 | 0.43068 |
| Chromobox protein homolog 3 | P23198 | Cbx3 | 21 kDa | 3 C | 3.200 | 0.21217 |
| Heterogeneous nuclear ribonucleoprotein F | Q9Z2X1 (+1) | Hnrnpf | 46 kDa | 6 C | 3.200 | 0.2954 |
| Isochorismatase domain-containing protein 2A | P85094 | Isoc2a | 22 kDa | 6 C | 2.698 | 0.24065 |
| Vimentin | P20152 | Vim | 54 kDa | 1 C | 2.686 | 0.16574 |
| Isoform 2 of Tropomyosin alpha-3 chain | P21107-2 | Tpm3 | 29 kDa | 1 C | 2.600 | 0.17192 |
| Caspase-3 | P70677 | Casp3 | 31 kDa | 8 C | 2.585 | 0.28648 |
| Haptoglobin | Q61646 | Hp | 39 kDa | 9 C | 2.581 | 0.18143 |
| Band 3 anion transport protein | P04919 | Slc4a1 | 103 kDa | 6 C | 2.550 | 0.34828 |
| Complement component C8 alpha chain | Q9R0P5 | C8a | 66 kDa | 29 C | 2.509 | 0.05098 |
| Vitronectin | P29788 | Vtn | 55 kDa | 14 C | 2.435 | 0.14375 |
| Adenosylhomocysteinase | P50247 | Ahcy | 48 kDa | 9 C | 2.400 | 0.07034 |
| Alpha-2-HS-glycoprotein | P29699 | Ahsg | 37 kDa | 14 C | 2.400 | 0.12218 |
| Hemoglobin subunit beta-1 | P01843 | Hbb-b1 | 16 kDa | 2 C | 2.307 | 0.09462 |
| Transketolase | P40142 | Tkt | 68 kDa | 12 C | 2.279 | 0.39077 |

| | | | | | | |
|---|---|---|---|---|---|---|
| Aspartyl aminopeptidase | Q9Z2W0 | Dnpep | 52 kDa | 10 C | 2.255 | 0.44176 |
| Proteasome subunit alpha type-2 | P49722 | Psma2 | 26 kDa | 2 C | 2.230 | 0.10612 |
| Proteasome subunit beta type-7 | P70195 | Psmb7 | 30 kDa | 6 C | 2.116 | 0.29727 |
| Carboxylesterase 1C | P23953 | Ces1c | 61 kDa | 5 C | 2.098 | 0.24819 |
| SH3 domain-binding glutamic acid-rich-like protein | Q9JJU8 | Sh3bgrl | 13 kDa | 2 C | 2.092 | 0.62111 |
| Purine nucleoside phosphorylase | P23492 | Pnp | 32 kDa | 5 C | 2.055 | 0.26527 |
| Isoform 2 of Glyoxalase domain-containing protein 4 | Q8BND5 (+1) | Glod4 | 32 kDa | 5 C | 2.031 | 0.50045 |
| Sulfhydryl oxidase 1 | P70274 | Qsox1 | 83 kDa | 14 C | 2.031 | 0.50045 |
| Serum albumin | P01872 | Alb | 69 kDa | 36 C | 1.966 | 0.0124 |
| Selenoprotein P | Q9JLT4-2 | Selenop | 43 kDa | 18 C | 1.951 | 0.08939 |
| Isoform 2 of Thioredoxin reductase 2, mitochondrial | Q8R4N0 | Txnrd2 | 53 kDa | 11 C | 1.943 | 0.04293 |
| Ig gamma-2A chain C region secreted form | O55023 | | 37 kDa | 9 C | 1.920 | 0.1053 |
| Inositol monophosphatase 1 | P01635 | Impa1 | 30 kDa | 6 C | 1.920 | 0.00858 |
| Pregnancy zone protein | Q61838 | Pzp | 166 kDa | 24 C | 1.851 | 0.07842 |
| Ig kappa chain V-VI region NQ2-17.4.1 | P04940 (+2) | | 12 kDa | 2 C | 1.843 | 0.62636 |
| Endoplasmic reticulum resident protein 44 | Q9D1Q6 | Erp44 | 47 kDa | 7 C | 1.832 | 0.21814 |
| Alpha-1-antitrypsin 1-1 | Q9ESB3 | Serpina1a | 46 kDa | 3 C | 1.809 | 0.09739 |
| Histidine-rich glycoprotein | Q99KB8 | Hrg | 59 kDa | 17 C | 1.803 | 0.07459 |
| Hydroxyacylglutathione hydrolase, mitochondrial | P01787 (+4) | Hagh | 34 kDa | 8 C | 1.800 | 0.47363 |
| Ig heavy chain V regions TEPC 15/S107/HPCM1/HPCM2/HPCM3 | P01942 | | 14 kDa | 2 C | 1.800 | 0.47363 |
| Hemoglobin subunit alpha | Q8BGD9 | Hba | 15 kDa | 1 C | 1.778 | 0.1735 |
| Eukaryotic translation initiation factor 4B | P39039 | Eif4b | 69 kDa | 3 C | 1.776 | 0.31128 |
| Mannose-binding protein A | P24270 | Mbl1 | 25 kDa | 8 C | 1.733 | 0.2542 |
| Catalase | P97371 | Cat | 60 kDa | 5 C | 1.714 | 0.22028 |
| Ig kappa chain V-II region 7S34.1 | P01630 | | 12 kDa | 3 C | 1.705 | 0.06913 |
| Ficolin-1 | O70165 | Fcn1 | 36 kDa | 10 C | 1.657 | 0.1053 |
| Titin | A2ASS6 | Ttn | 3906 kDa | 498 C | 1.652 | 0.71992 |
| Lactoylglutathione lyase | Q9CPU0 | Glo1 | 21 kDa | 3 C | 1.600 | 0.28611 |
| Annexin A5 | P48036 | Anxa5 | 36 kDa | 1 C | 1.587 | 0.36761 |
| Alpha-1-antitrypsin 1-2 | P22599 | Serpina1b | 46 kDa | 3 C | 1.571 | 0.14401 |
| Alpha-1-antitrypsin 1-4 | Q61129 | Serpina1d | 46 kDa | 3 C | 1.570 | 0.12078 |
| Ig heavy chain V region AC38 205.12 | P06330 | | 13 kDa | 2 C | 1.570 | 0.18001 |
| cAMP-dependent protein kinase type I-alpha regulatory subunit | Q00897 | Prkar1a | 43 kDa | 5 C | 1.564 | 0.66117 |
| Cofilin-1 | Q60854 | Cfl1 | 19 kDa | 4 C | 1.527 | 0.23691 |
| Destrin | P17563 | Dstn | 19 kDa | 6 C | 1.527 | 0.54905 |
| Serpin B6 | O70435 | Serpinb6 | 43 kDa | 6 C | 1.527 | 0.54905 |
| Proteasome subunit alpha type-3 | P01029 | Psma3 | 28 kDa | 4 C | 1.511 | 0.31989 |
| Selenium-binding protein 1 | Q8CC35-2 | Selenbp1 | 53 kDa | 10 C | 1.511 | 0.37637 |
| Complement C4-B | Q9JKB1 | C4b | 193 kDa | 29 C | 1.500 | 0.05688 |
| Ubiquitin carboxyl-terminal hydrolase isozyme L3 | P11352 | Uchl3 | 26 kDa | 3 C | 1.496 | 0.68453 |
| Glutathione peroxidase 1 | Q8VCG4 | Gpx1 | 22 kDa | 4 C | 1.492 | 0.08725 |
| Myosin light chain 3 | P09542 | Myl3 | 22 kDa | 2 C | 1.488 | 0.20619 |
| Adenylyl cyclase-associated protein 1 | P40124 | Cap1 | 52 kDa | 6 C | 1.477 | 0.75324 |
| Heat shock protein beta-6 | Q5EBG6 | Hspb6 | 18 kDa | 1 C | 1.477 | 0.75324 |
| Alpha-actinin-1 | Q7TPR4 | Actn1 | 103 kDa | 11 C | 1.461 | 0.66058 |
| Delta-aminolevulinic acid dehydratase | Q60631 | Alad | 36 kDa | 8 C | 1.440 | 0.18479 |
| Citrate lyase subunit beta-like protein, mitochondrial | P10518 | Clybl | 38 kDa | 6 C | 1.440 | 0.28741 |
| Immunoglobulin J chain | P01592 | Jchain | 18 kDa | 8 C | 1.429 | 0.19446 |
| Carbonic anhydrase 1 | P07758 (+1) | Ca1 | 28 kDa | 1 C | 1.426 | 0.34119 |
| Ig lambda-1 chain V region | P01723 (+1) | | 12 kDa | 2 C | 1.426 | 0.61414 |
| Ig mu chain C region | Q0II04 | Ighm | 50 kDa | 19 C | 1.416 | 0.08377 |
| Hemopexin | P13634 | Hpx | 51 kDa | 13 C | 1.415 | 0.21091 |
| Ig kappa chain C region | O08677 | | 12 kDa | 3 C | 1.393 | 0.06778 |
| Proteasome activator complex subunit 1 | Q91X72 | Psme1 | 29 kDa | 3 C | 1.383 | 0.52037 |
| Carbonic anhydrase 2 | P00920 | Ca2 | 29 kDa | 2 C | 1.371 | 0.3371 |
| 6-pyruvoyl tetrahydrobiopterin synthase | Q9R1Z7 | Pts | 16 kDa | 1 C | 1.366 | 0.42196 |
| Triosephosphate isomerase | Q9DBD0 | Tpi1 | 32 kDa | 9 C | 1.354 | 0.36558 |
| Inhibitor of carbonic anhydrase | P07759 | Ica | 77 kDa | 35 C | 1.347 | 0.54889 |
| Serine protease inhibitor A3K | Q921I1 | Serpina3k | 47 kDa | 4 C | 1.333 | 0.2816 |
| Serotransferrin | O08749 | Tf | 77 kDa | 38 C | 1.325 | 0.09052 |

| | | | | | | |
|---|---|---|---|---|---|---|
| CD5 antigen-like | P49312 | Cd5l | 39 kDa | 26 C | 1.301 | 0.0062 |
| Heterogeneous nuclear ribonucleoprotein A1 | Q923D2 | Hnrnpa1 | 34 kDa | 2 C | 1.300 | 0.71228 |
| Flavin reductase (NADPH) | O35855 | Blvrb | 22 kDa | 2 C | 1.287 | 0.33745 |
| Complement component C8 gamma chain | P56375 | C8g | 23 kDa | 3 C | 1.284 | 0.37402 |
| Branched-chain-amino-acid aminotransferase, mitochondrial | P47791-2 | Bcat2 | 44 kDa | 10 C | 1.267 | 0.3228 |
| Isoform Cytoplasmic of Glutathione reductase, mitochondrial | Q9R0Y5 (+1) | Gsr | 51 kDa | 11 C | 1.267 | 0.4666 |
| Adenylate kinase isoenzyme 1 | Q61171 | Ak1 | 22 kDa | 2 C | 1.265 | 0.37098 |
| Peroxiredoxin-2 | Q9JKS4-3 | Prdx2 | 22 kDa | 3 C | 1.261 | 0.31905 |
| Ig heavy chain V region J558 | P01757 | | 13 kDa | 2 C | 1.257 | 0.43285 |
| Isoform 2 of Plasminogen activator inhibitor 1 RNA-binding protein | Q9CY58-2 | Serbp1 | 43 kDa | 2 C | 1.255 | 0.60821 |
| Rho GDP-dissociation inhibitor 1 | Q99PT1 | Arhgdia | 23 kDa | 1 C | 1.255 | 0.60821 |
| Isoform 2 of Synaptopodin | P62141 | Synpo | 96 kDa | 5 C | 1.246 | 0.71663 |
| Serine/threonine-protein phosphatase PP1-beta catalytic subunit | P01878 | Ppp1cb | 37 kDa | 14 C | 1.240 | 0.61243 |
| Transthyretin | P07309 | Ttr | 16 kDa | 2 C | 1.237 | 0.69688 |
| Ig kappa chain V-VI region NQ2-6.1 | P04945 | | 12 kDa | 2 C | 1.236 | 0.44313 |
| Ig kappa chain V-II region 26-10 | P01631 | | 12 kDa | 2 C | 1.231 | 0.49326 |
| Ig kappa chain V-V region K2 (Fragment) | P17182 | | 13 kDa | 3 C | 1.200 | 0.40708 |
| Peptidyl-prolyl cis-trans isomerase A | Q00898 | Ppia | 18 kDa | 3 C | 1.200 | 0.64702 |
| Nebulette | P62137 | Nebl | 52 kDa | 7 C | 1.200 | 0.82713 |
| Serine/threonine-protein phosphatase PP1-alpha catalytic subunit | Q9DCX2 | Ppp1ca | 38 kDa | 13 C | 1.183 | 0.71405 |
| Alpha-enolase | Q9WTR5 | Eno1 | 47 kDa | 6 C | 1.164 | 0.45557 |
| Alpha-1-antitrypsin 1-5 | P08228 | Serpina1e | 46 kDa | 4 C | 1.160 | 0.67135 |
| Ig gamma-1 chain C region secreted form | Q9D1X0 | Ighg1 | 36 kDa | 12 C | 1.156 | 0.77681 |
| Cadherin-13 | Q9D172 | Cdh13 | 78 kDa | 7 C | 1.154 | 0.7235 |
| Superoxide dismutase [Cu-Zn] | P70296 | Sod1 | 16 kDa | 3 C | 1.148 | 0.89094 |
| Nucleolar protein 3 | Q8K0E8 | Nol3 | 25 kDa | 4 C | 1.148 | 0.89094 |
| ES1 protein homolog, mitochondrial | Q8VCM7 | D10Jhu81e | 28 kDa | 6 C | 1.140 | 0.82575 |
| Phosphatidylethanolamine-binding protein 1 | Q9DBC7 | Pebp1 | 21 kDa | 3 C | 1.139 | 0.68623 |
| Fibrinogen gamma chain | O35459 | Fgg | 49 kDa | 12 C | 1.120 | 0.70009 |
| Fibrinogen beta chain | P48787 | Fgb | 55 kDa | 12 C | 1.120 | 0.7682 |
| Delta(3,5)-Delta(2,4)-dienoyl-CoA isomerase, mitochondrial | P01837 | Ech1 | 36 kDa | 6 C | 1.120 | 0.80276 |
| Troponin I, cardiac muscle | Q99KI0 | Tnni3 | 24 kDa | 2 C | 1.091 | 0.86479 |
| Acylphosphatase-2 | O08807 | Acyp2 | 12 kDa | 1 C | 1.084 | 0.88833 |
| Growth factor receptor-bound protein 2 | Q93092 | Grb2 | 25 kDa | 2 C | 1.075 | 0.93862 |
| Transaldolase | P46412 | Taldo1 | 37 kDa | 3 C | 1.067 | 0.79797 |
| Peroxiredoxin-4 | P63260 | Prdx4 | 31 kDa | 4 C | 1.067 | 0.85069 |
| Thioredoxin-dependent peroxide reductase, mitochondrial | Q3ULD5 | Prdx3 | 28 kDa | 4 C | 1.055 | 0.83689 |
| Septin-2 | Q9DBP5 | 43345 | 42 kDa | 8 C | 1.048 | 0.95823 |
| Actin, cytoplasmic 2 | Q9QWK4 | Actg1 | 42 kDa | 6 C | 1.048 | 0.76138 |
| Glutathione peroxidase 3 | P17751 | Gpx3 | 25 kDa | 3 C | 1.046 | 0.81532 |
| Complement component C8 beta chain | P15532 | C8b | 66 kDa | 32 C | 1.040 | 0.93253 |
| UMP-CMP kinase | P01804 | Cmpk1 | 22 kDa | 6 C | 1.040 | 0.93656 |
| Nucleoside diphosphate kinase A | Q9QUM9 | Nme1 | 17 kDa | 2 C | 1.033 | 0.84337 |
| Ig heavy chain V-III region HPC76 (Fragment) | P21550 | | 12 kDa | 2 C | 1.032 | 0.95886 |
| Dihydrolipoyl dehydrogenase, mitochondrial | P36552 | Dld | 54 kDa | 9 C | 1.029 | 0.92417 |
| Proteasome subunit alpha type-6 | Q9R1P3 | Psma6 | 27 kDa | 8 C | 1.022 | 0.91958 |
| Beta-enolase | P14211 | Eno3 | 47 kDa | 6 C | 1.013 | 0.95758 |
| Oxygen-dependent coproporphyrinogen-III oxidase, mitochondrial | P01639 | Cpox | 50 kDa | 10 C | 1.010 | 0.98553 |
| Isoform 3 of LIM domain-binding protein 3 | Q8BH35 | Ldb3 | 71 kDa | 21 C | 1.000 | 1 |
| Proteasome subunit beta type-2 | Q9Z2U1 | Psmb2 | 23 kDa | 3 C | 1.000 | 1 |
| Ig kappa chain V-V region MOPC 41 | Q9Z2U0 | Gm5571 | 14 kDa | 3 C | 1.000 | 1 |
| Proteasome subunit alpha type-5 | Q9JI91 | Psma5 | 26 kDa | 3 C | 0.987 | 0.97268 |
| Proteasome subunit alpha type-7 | P09671 | Psma7 | 28 kDa | 3 C | 0.985 | 0.87862 |
| Alpha-actinin-2 | Q60597-3 | Actn2 | 104 kDa | 10 C | 0.977 | 0.95313 |
| Superoxide dismutase [Mn], mitochondrial | P60335 | Sod2 | 25 kDa | 4 C | 0.975 | 0.91813 |
| Peroxiredoxin-5, mitochondrial | P37804 | Prdx5 | 22 kDa | 6 C | 0.967 | 0.84581 |
| Isoform 3 of 2-oxoglutarate dehydrogenase, mitochondrial | P01868 (+1) | Ogdh | 118 kDa | 21 C | 0.962 | 0.94758 |
| Poly(rC)-binding protein 1 | O55234 | Pcbp1 | 37 kDa | 9 C | 0.957 | 0.92599 |
| Transgelin | Q62188 | Tagln | 23 kDa | 1 C | 0.937 | 0.89101 |
| Thioredoxin | Q9CQ62 | Txn | 12 kDa | 6 C | 0.933 | 0.40708 |

| | | | | | | |
|---|---|---|---|---|---|---|
| Proteasome subunit beta type-5 | Q9R1P1 | Psmb5 | 29 kDa | 3 C | 0.933 | 0.86758 |
| Dihydropyrimidinase-related protein 3 | Q9CZ13 | Dpysl3 | 62 kDa | 7 C | 0.929 | 0.84448 |
| 2,4-dienoyl-CoA reductase, mitochondrial | P08249 | Decr1 | 36 kDa | 5 C | 0.923 | 0.77236 |
| Proteasome subunit beta type-3 | O08709 | Psmb3 | 23 kDa | 5 C | 0.923 | 0.79042 |
| Aconitate hydratase, mitochondrial | Q91YT0 | Aco2 | 85 kDa | 13 C | 0.916 | 0.64109 |
| Myosin-binding protein C, cardiac-type | Q9DCT2 | Mybpc3 | 141 kDa | 21 C | 0.914 | 0.7746 |
| Malate dehydrogenase, mitochondrial | P20108 | Mdh2 | 36 kDa | 8 C | 0.914 | 0.7958 |
| Peroxiredoxin-6 | P10639 | Prdx6 | 25 kDa | 2 C | 0.908 | 0.67293 |
| Ig alpha chain C region | P70349 | | 37 kDa | 13 C | 0.903 | 0.86707 |
| NADH dehydrogenase [ubiquinone] iron-sulfur protein 3, mitochondrial | P99026 | Ndufs3 | 30 kDa | 3 C | 0.898 | 0.82894 |
| Proteasome subunit beta type-4 | Q9CR68 | Psmb4 | 29 kDa | 2 C | 0.889 | 0.66466 |
| Histidine triad nucleotide-binding protein 1 | O08553 | Hint1 | 14 kDa | 2 C | 0.889 | 0.77236 |
| Glutathione S-transferase Mu 2 | P12787 | Gstm2 | 26 kDa | 3 C | 0.888 | 0.74676 |
| Calreticulin | Q01768 | Calr | 48 kDa | 6 C | 0.888 | 0.81803 |
| Nucleoside diphosphate kinase B | P07724 | Nme2 | 17 kDa | 2 C | 0.884 | 0.44086 |
| NADH dehydrogenase [ubiquinone] flavoprotein 1, mitochondrial | Q9JHU2 | Ndufv1 | 51 kDa | 12 C | 0.884 | 0.65449 |
| Cytochrome b-c1 complex subunit Rieske, mitochondrial | P68033 | Uqcrfs1 | 29 kDa | 5 C | 0.883 | 0.62861 |
| Dihydropyrimidinase-related protein 2 | P15626 | Dpysl2 | 62 kDa | 7 C | 0.881 | 0.77152 |
| Cytochrome c oxidase subunit 5A, mitochondrial | Q9R1P0 | Cox5a | 16 kDa | 4 C | 0.880 | 0.68453 |
| Actin, alpha cardiac muscle 1 | O70251 | Actc1 | 42 kDa | 6 C | 0.869 | 0.46071 |
| Ig kappa chain V-V region HP R16.7 | P24549 | | 12 kDa | 2 C | 0.867 | 0.40708 |
| Proteasome subunit alpha type-4 | O09061 | Psma4 | 29 kDa | 5 C | 0.867 | 0.61902 |
| Methylglutaconyl-CoA hydratase, mitochondrial | P35979 | Auh | 33 kDa | 5 C | 0.862 | 0.37757 |
| Elongation factor 1-beta | P10649 | Eef1b | 25 kDa | 3 C | 0.860 | 0.79843 |
| Proteasome subunit beta type-1 | P03977 | Psmb1 | 26 kDa | 5 C | 0.857 | 0.48033 |
| 60S ribosomal protein L12 | Q9D358 | Rpl12 | 18 kDa | 3 C | 0.857 | 0.64702 |
| Glutathione S-transferase Mu 1 | P01644 (+1) | Gstm1 | 26 kDa | 2 C | 0.856 | 0.39515 |
| BAG family molecular chaperone regulator 3 | P05201 | Bag3 | 62 kDa | 4 C | 0.846 | 0.55011 |
| Ig kappa chain V-III region 50S10.1 | Q04447 | | 12 kDa | 2 C | 0.842 | 0.46637 |
| Low molecular weight phosphotyrosine protein phosphatase | Q3UTJ2-2 | Acp1 | 18 kDa | 8 C | 0.839 | 0.82609 |
| Adiponectin | Q99LX0 | Adipoq | 27 kDa | 2 C | 0.833 | 0.32686 |
| Cytochrome b-c1 complex subunit 1, mitochondrial | Q99L13 | Uqcrc1 | 53 kDa | 11 C | 0.833 | 0.61312 |
| Aspartate aminotransferase, cytoplasmic | Q64010 | Got1 | 46 kDa | 5 C | 0.832 | 0.08166 |
| Isoform 2 of Sorbin and SH3 domain-containing protein 2 | Q6P8J7 | Sorbs2 | 145 kDa | 3 C | 0.830 | 0.79325 |
| Protein DJ-1 | O08600 | Park7 | 20 kDa | 4 C | 0.830 | 0.36524 |
| Creatine kinase B-type | P42125 | Ckb | 43 kDa | 5 C | 0.828 | 0.5112 |
| Adapter molecule crk | P35700 | Crk | 34 kDa | 1 C | 0.825 | 0.81752 |
| Creatine kinase S-type, mitochondrial | P26041 | Ckmt2 | 47 kDa | 8 C | 0.820 | 0.32643 |
| Peroxiredoxin-1 | P63158 | Prdx1 | 22 kDa | 4 C | 0.819 | 0.38844 |
| Endonuclease G, mitochondrial | P19783 | Endog | 32 kDa | 2 C | 0.819 | 0.81895 |
| Moesin | Q9WUZ7 | Msn | 68 kDa | 2 C | 0.816 | 0.58831 |
| Cytochrome c oxidase subunit 4 isoform 1, mitochondrial | Q9CQ92 | Cox4i1 | 20 kDa | 1 C | 0.816 | 0.58831 |
| SH3 domain-binding glutamic acid-rich protein | P07310 | Sh3bgr | 23 kDa | 1 C | 0.816 | 0.75448 |
| Mitochondrial fission 1 protein | Q9D1A2 | Fis1 | 17 kDa | 1 C | 0.811 | 0.70298 |
| Creatine kinase M-type | P51174 | Ckm | 43 kDa | 8 C | 0.809 | 0.31 |
| Cytosolic non-specific dipeptidase | P09103 | Cndp2 | 53 kDa | 8 C | 0.804 | 0.62267 |
| ATP synthase subunit d, mitochondrial | P19536 | Atp5h | 19 kDa | 1 C | 0.800 | 0.84228 |
| Long-chain specific acyl-CoA dehydrogenase, mitochondrial | P61982 | Acadl | 48 kDa | 7 C | 0.800 | 0.3454 |
| Protein disulfide-isomerase | O88569 | P4hb | 57 kDa | 7 C | 0.800 | 0.46011 |
| Glycerol-3-phosphate phosphatase | P19157 | Pgp | 35 kDa | 8 C | 0.800 | 0.4758 |
| Glutathione S-transferase P 1 | Q8CHP8 | Gstp1 | 24 kDa | 3 C | 0.800 | 0.53127 |
| Heterogeneous nuclear ribonucleoproteins A2/B1 | P07356 | Hnrnpa2b1 | 37 kDa | 1 C | 0.800 | 0.61656 |
| Annexin A2 | Q9D0S9 | Anxa2 | 39 kDa | 5 C | 0.800 | 0.61902 |
| Histidine triad nucleotide-binding protein 2, mitochondrial | Q9QXD6 | Hint2 | 17 kDa | 1 C | 0.800 | 0.64307 |
| Cytochrome c oxidase subunit 5B, mitochondrial | Q64339 | Cox5b | 14 kDa | 5 C | 0.800 | 0.74616 |
| Fructose-1,6-bisphosphatase 1 | P63038 | Fbp1 | 37 kDa | 7 C | 0.800 | 0.74723 |
| Ubiquitin-like protein ISG15 | Q9D6Y7-2 | Isg15 | 18 kDa | 3 C | 0.800 | 0.81985 |
| Isoform 2 of Mitochondrial peptide methionine sulfoxide reductase | P01796 (+3) | Msra | 24 kDa | 4 C | 0.800 | 0.27712 |
| Proteasome subunit alpha type-1 | P57776-3 | Psma1 | 30 kDa | 5 C | 0.800 | 0.50235 |
| 60 kDa heat shock protein, mitochondrial | Q9CPY7 | Hspd1 | 61 kDa | 3 C | 0.800 | 0.59511 |

| | | | | | | |
|---|---|---|---|---|---|---|
| Ig heavy chain V-III region A4 | Q01853 | | 13 kDa | 2 C | 0.800 | 0.67938 |
| Isoform 3 of Elongation factor 1-delta | Q9R1P4 | Eef1d | 73 kDa | 2 C | 0.800 | 0.77464 |
| Cytosol aminopeptidase | P18760 | Lap3 | 56 kDa | 7 C | 0.789 | 0.31167 |
| Transitional endoplasmic reticulum ATPase | Q9EQ20 | Vcp | 89 kDa | 12 C | 0.775 | 0.48624 |
| Malate dehydrogenase, cytoplasmic | Q60692 | Mdh1 | 37 kDa | 3 C | 0.763 | 0.05709 |
| Profilin-1 | P62962 | Pfn1 | 15 kDa | 3 C | 0.756 | 0.12498 |
| Electron transfer flavoprotein subunit beta | P62897 | Etfb | 28 kDa | 4 C | 0.755 | 0.26657 |
| Succinate dehydrogenase [ubiquinone] iron-sulfur subunit, mitochondrial | Q9JLZ3 | Sdhb | 32 kDa | 14 C | 0.751 | 0.42244 |
| Succinate-semialdehyde dehydrogenase, mitochondrial | Q8BKZ9 | Aldh5a1 | 56 kDa | 10 C | 0.741 | 0.18272 |
| Enoyl-CoA hydratase, mitochondrial | P09411 | Echs1 | 31 kDa | 7 C | 0.740 | 0.44313 |
| Pyruvate dehydrogenase protein X component, mitochondrial | Q8BFR5 | Pdhx | 54 kDa | 5 C | 0.738 | 0.32316 |
| Methylcrotonoyl-CoA carboxylase beta chain, mitochondrial | O70468 | Mccc2 | 61 kDa | 10 C | 0.736 | 0.43102 |
| Elongation factor Tu, mitochondrial | P16045 | Tufm | 50 kDa | 7 C | 0.728 | 0.48832 |
| Cofilin-2 | Q9CZU6 | Cfl2 | 19 kDa | 2 C | 0.727 | 0.29235 |
| NADH dehydrogenase [ubiquinone] flavoprotein 2, mitochondrial | Q3UM45 | Ndufv2 | 27 kDa | 6 C | 0.720 | 0.09517 |
| Galectin-1 | Q8K2B3 | Lgals1 | 15 kDa | 6 C | 0.720 | 0.4498 |
| Citrate synthase, mitochondrial | Q9WVA4 | Cs | 52 kDa | 5 C | 0.716 | 0.363 |
| Phosphoglycerate kinase 1 | Q9D0K2 | Pgk1 | 45 kDa | 7 C | 0.716 | 0.42277 |
| Protein phosphatase 1 regulatory subunit 7 | P63101 | Ppp1r7 | 41 kDa | 2 C | 0.709 | 0.14143 |
| Succinate dehydrogenase [ubiquinone] flavoprotein subunit, mitochondrial | Q9DBF1 (+1) | Sdha | 73 kDa | 19 C | 0.709 | 0.27701 |
| Transgelin-2 | P45952 | Tagln2 | 22 kDa | 3 C | 0.704 | 0.49458 |
| Succinyl-CoA:3-ketoacid coenzyme A transferase 1, mitochondrial | Q9DCT8 | Oxct1 | 56 kDa | 7 c | 0.703 | 0.37927 |
| 14-3-3 protein zeta/delta | Q9JHR7 | Ywhaz | 28 kDa | 3 C | 0.703 | 0.35582 |
| Medium-chain specific acyl-CoA dehydrogenase, mitochondrial | Q61425 | Acadm | 46 kDa | 8 C | 0.700 | 0.06758 |
| Cysteine-rich protein 2 | P01642 | Crip2 | 23 kDa | 14 C | 0.700 | 0.28611 |
| Insulin-degrading enzyme | Q9D0M3-2 | Ide | 118 kDa | 13 C | 0.697 | 0.46483 |
| Hydroxyacyl-coenzyme A dehydrogenase, mitochondrial | P14206 | Hadh | 34 kDa | 5 C | 0.696 | 0.2533 |
| Ig kappa chain V-V region L7 (Fragment) | Q8CI51-3 | Gm10881 | 13 kDa | 2 C | 0.693 | 0.42615 |
| Retinal dehydrogenase 1 | P04117 | Aldh1a1 | 54 kDa | 11 C | 0.691 | 0.41087 |
| Isoform 2 of Cytochrome c1, heme protein, mitochondrial | Q9D6R2 | Cyc1 | 29 kDa | 14 C | 0.689 | 0.6224 |
| Palmdelphin | Q149B8 | Palmd | 63 kDa | 3 C | 0.688 | 0.51697 |
| 40S ribosomal protein SA | Q99M71 | Rpsa | 33 kDa | 2 C | 0.683 | 0.50305 |
| Isoform 3 of PDZ and LIM domain protein 5 | P20029 | Pdlim5 | 26 kDa | 22 C | 0.678 | 0.24788 |
| Fatty acid-binding protein, adipocyte | O35737 | Fabp4 | 15 kDa | 2 C | 0.677 | 0.22208 |
| Isocitrate dehydrogenase [NAD] subunit alpha, mitochondrial | P63028 | Idh3a | 40 kDa | 8 C | 0.677 | 0.29767 |
| PGC-1 and ERR-induced regulator in muscle protein 1 | P63242 | Perm1 | 85 kDa | 13 C | 0.674 | 0.5302 |
| Mammalian ependymin-related protein 1 | P16125 | Epdr1 | 25 kDa | 7 C | 0.672 | 0.39897 |
| 78 kDa glucose-regulated protein | Q60994 | Hspa5 | 72 kDa | 1 C | 0.667 | 0.02265 |
| Heterogeneous nuclear ribonucleoprotein H | Q8BH69 | Hnrnph1 | 49 kDa | 1 C | 0.663 | 0.65554 |
| High mobility group protein B1 | Q91VD9 | Hmgb1 | 25 kDa | 3 C | 0.663 | 0.65554 |
| Translationally-controlled tumor protein | P52503 | Tpt1 | 19 kDa | 2 C | 0.659 | 0.46799 |
| Eukaryotic translation initiation factor 5A-1 | Q99020 | Eif5a | 17 kDa | 4 C | 0.649 | 0.51957 |
| L-lactate dehydrogenase B chain | Q9CR00 | Ldhb | 37 kDa | 5 C | 0.649 | 0.09688 |
| Selenide, water dikinase 1 | Q64727 | Sephs1 | 43 kDa | 9 C | 0.644 | 0.5262 |
| Protein phosphatase 1 regulatory subunit 12B | O09131 | Ppp1r12b | 109 kDa | 6 C | 0.640 | 0.34334 |
| NADH-ubiquinone oxidoreductase 75 kDa subunit, mitochondrial | Q8BMS1 | Ndufs1 | 80 kDa | 18 C | 0.638 | 0.20882 |
| NADH dehydrogenase [ubiquinone] iron-sulfur protein 6, mitochondrial | P63017 | Ndufs6 | 13 kDa | 3 C | 0.631 | 0.27602 |
| Heterogeneous nuclear ribonucleoprotein A/B | Q8BWT1 | Hnrnpab | 31 kDa | 2 C | 0.630 | 0.51468 |
| 26S proteasome non-ATPase regulatory subunit 9 | Q9WVQ5 | Psmd9 | 25 kDa | 3 C | 0.629 | 0.56421 |
| Vinculin | P52480-2 | Vcl | 117 kDa | 10 C | 0.627 | 0.06693 |
| Alpha-aminoadipic semialdehyde dehydrogenase | Q9DC69 | Aldh7a1 | 59 kDa | 9 C | 0.625 | 0.30063 |
| Glutathione S-transferase omega-1 | Q8VCT4 | Gsto1 | 27 kDa | 4 C | 0.624 | 0.39269 |
| Heat shock cognate 71 kDa protein | Q8BH95 | Hspa8 | 71 kDa | 4 C | 0.622 | 0.07444 |
| 3-ketoacyl-CoA thiolase, mitochondrial | P99029 | Acaa2 | 42 kDa | 8 C | 0.620 | 0.15439 |
| Methylthioribulose-1-phosphate dehydratase | Q9CQ75 | Apip | 27 kDa | 11 C | 0.607 | 0.22865 |
| Isoform M1 of Pyruvate kinase PKM | Q8BWF0 | Pkm | 58 kDa | 9 C | 0.604 | 0.0779 |
| 14-3-3 protein gamma | P70670 | Ywhag | 28 kDa | 3 C | 0.603 | 0.22802 |
| NADH dehydrogenase [ubiquinone] 1 alpha subcomplex subunit 9, mitochondrial | P05202 | Ndufa9 | 43 kDa | 2 C | 0.600 | 0.60218 |
| Carboxylesterase 1D | Q9D2G2 | Ces1d | 62 kDa | 5 C | 0.597 | 0.50124 |
| 14-3-3 protein epsilon | P14152 | Ywhae | 29 kDa | 3 C | 0.594 | 0.26823 |

| | | | | | | |
|---|---|---|---|---|---|---|
| Fructose-bisphosphate aldolase A | Q9CQA3 | Aldoa | 39 kDa | 8 C | 0.593 | 0.09987 |
| Cytochrome c, somatic | Q91WD5 | Cycs | 12 kDa | 2 C | 0.589 | 0.10481 |
| NADH dehydrogenase [ubiquinone] 1 alpha subcomplex subunit 2 | P50462 | Ndufa2 | 11 kDa | 2 C | 0.586 | 0.34006 |
| Nascent polypeptide-associated complex subunit alpha, muscle-specific form | Q9WTX5 | Naca | 220 kDa | 16 C | 0.579 | 0.19906 |
| Aspartate aminotransferase, mitochondrial | Q9JJW5 | Got2 | 47 kDa | 7 C | 0.577 | 0.05592 |
| Proteasome subunit beta type-6 | P04247 | Psmb6 | 25 kDa | 4 C | 0.571 | 0.11765 |
| ATP synthase subunit alpha, mitochondrial | Q9JLV1 | Atp5a1 | 60 kDa | 2 C | 0.559 | 0.05087 |
| Dihydrolipoyllysine-residue succinyltransferase component of 2-oxoglutarate dehydrogena | Q91Z53 | Dlst | 49 kDa | 6 C | 0.559 | 0.01747 |
| NADH dehydrogenase [ubiquinone] iron-sulfur protein 2, mitochondrial | P97807 | Ndufs2 | 53 kDa | 7 C | 0.558 | 0.61016 |
| Cysteine and glycine-rich protein 3 | P17742 | Csrp3 | 21 kDa | 16 C | 0.558 | 0.24709 |
| Enoyl-CoA delta isomerase 1, mitochondrial | Q8QZT1 | Eci1 | 32 kDa | 5 C | 0.552 | 0.06983 |
| Myoglobin | Q9DCB8 | Mb | 17 kDa | 1 C | 0.544 | 0.07516 |
| Fumarate hydratase, mitochondrial | P47738 | Fh | 54 kDa | 4 C | 0.542 | 0.0767 |
| Acetyl-CoA acetyltransferase, mitochondrial | P45591 | Acat1 | 45 kDa | 6 C | 0.541 | 0.10901 |
| Methylmalonate-semialdehyde dehydrogenase [acylating], mitochondrial | Q9DCW4 | Aldh6a1 | 58 kDa | 8 C | 0.533 | 0.05236 |
| Iron-sulfur cluster assembly 2 homolog, mitochondrial | P50544 | Isca2 | 17 kDa | 4 C | 0.528 | 0.25981 |
| Aldehyde dehydrogenase, mitochondrial | Q91VM9 | Aldh2 | 57 kDa | 9 C | 0.526 | 0.0429 |
| Very long-chain specific acyl-CoA dehydrogenase, mitochondrial | P38647 | Acadvl | 71 kDa | 7 C | 0.524 | 0.10121 |
| Inorganic pyrophosphatase 2, mitochondrial | P48962 | Ppa2 | 38 kDa | 8 C | 0.520 | 0.15401 |
| Cytochrome c oxidase subunit 2 | Q9CXZ1 | Mtco2 | 26 kDa | 3 C | 0.520 | 0.30322 |
| NADH dehydrogenase [ubiquinone] iron-sulfur protein 4, mitochondrial | P06151 | Ndufs4 | 20 kDa | 1 C | 0.518 | 0.40096 |
| ADP/ATP translocase 1 | Q02566 | Slc25a4 | 33 kDa | 4 C | 0.510 | 0.08186 |
| Myosin-6 | Q8BG95 | Myh6 | 224 kDa | 14 C | 0.509 | 0.05604 |
| L-lactate dehydrogenase A chain | Q9DCS9 | Ldha | 36 kDa | 6 C | 0.509 | 0.03158 |
| NADH dehydrogenase [ubiquinone] 1 beta subcomplex subunit 10 | P54071 | Ndufb10 | 21 kDa | 5 C | 0.508 | 0.43065 |
| Isocitrate dehydrogenase [NADP], mitochondrial | Q99JY0 | Idh2 | 51 kDa | 8 C | 0.500 | 0.01269 |
| Trifunctional enzyme subunit beta, mitochondrial | P42208 | Hadhb | 51 kDa | 5 C | 0.496 | 0.04329 |
| Isoform 3 of Heterogeneous nuclear ribonucleoprotein D0 | Q60668-3 | Hnrnpd | 33 kDa | 3 C | 0.491 | 0.22867 |
| Microtubule-associated protein tau | P10637 | Mapt | 76 kDa | 2 C | 0.491 | 0.15899 |
| S-phase kinase-associated protein 1 | Q9CRB6 | Skp1 | 19 kDa | 3 C | 0.487 | 0.54552 |
| Tubulin polymerization-promoting protein family member 3 | Q8C1B7 | Tppp3 | 19 kDa | 3 C | 0.487 | 0.54552 |
| Septin-11 | O70250 | 43354 | 50 kDa | 6 C | 0.485 | 0.26171 |
| Phosphoglycerate mutase 2 | Q9Z0X1 | Pgam2 | 29 kDa | 3 C | 0.480 | 0.09212 |
| Heat shock 70 kDa protein 4 | Q9WUM5 | Hspa4 | 94 kDa | 14 C | 0.480 | 0.15201 |
| Succinate--CoA ligase [ADP/GDP-forming] subunit alpha, mitochondrial | P51125 (+1) | Suclg1 | 36 kDa | 6 C | 0.480 | 0.27632 |
| Calpastatin | P38060 | Cast | 85 kDa | 4 C | 0.473 | 0.30086 |
| Hydroxymethylglutaryl-CoA lyase, mitochondrial | Q9DBJ1 | Hmgcl | 34 kDa | 8 C | 0.471 | 0.30719 |
| CapZ-interacting protein | P27773 | Rcsd1 | 44 kDa | 5 C | 0.468 | 0.35426 |
| Phosphoglycerate mutase 1 | Q4ZJN1 | Pgam1 | 29 kDa | 2 C | 0.468 | 0.35426 |
| Protein disulfide-isomerase A3 | O55126 | Pdia3 | 57 kDa | 8 C | 0.467 | 0.09687 |
| Complement C1q and tumor necrosis factor-related protein 9 | Q9DB77 | C1qtnf9 | 35 kDa | 3 C | 0.462 | 0.14286 |
| Protein NipSnap homolog 2 | Q01279 | Nipsnap2 | 33 kDa | 3 C | 0.453 | 0.21332 |
| Myozenin-2 | E9PV24 | Myoz2 | 30 kDa | 1 C | 0.449 | 0.22315 |
| Epidermal growth factor receptor | Q921G7 | Egfr | 135 kDa | 60 C | 0.449 | 0.22315 |
| ATP synthase subunit gamma, mitochondrial | Q9DCM0 | Atp5c1 | 33 kDa | 2 C | 0.444 | 0.05353 |
| Persulfide dioxygenase ETHE1, mitochondrial | Q9D8X1 | Ethe1 | 28 kDa | 9 C | 0.440 | 0.13526 |
| Electron transfer flavoprotein-ubiquinone oxidoreductase, mitochondrial | Q9D051 | Etfdh | 68 kDa | 15 C | 0.440 | 0.1946 |
| Copper homeostasis protein cutC homolog | Q99LC5 | Cutc | 29 kDa | 7 C | 0.438 | 0.313 |
| Aminoacylase-1 | Q07417 | Acy1 | 46 kDa | 4 C | 0.438 | 0.313 |
| Short-chain specific acyl-CoA dehydrogenase, mitochondrial | Q9D6J6 | Acads | 45 kDa | 5 C | 0.436 | 0.07489 |
| MICOS complex subunit Mic19 | P62983 | Chchd3 | 26 kDa | 4 C | 0.428 | 0.51277 |
| Ubiquitin-40S ribosomal protein S27a | Q8CDN6 | Rps27a | 18 kDa | 6 C | 0.427 | 0.13256 |
| Thioredoxin-like protein 1 | O70325 | Txnl1 | 32 kDa | 7 C | 0.420 | 0.11563 |
| Phospholipid hydroperoxide glutathione peroxidase, mitochondrial | Q9DC70 | Gpx4 | 22 kDa | 10 C | 0.420 | 0.18643 |
| NADH dehydrogenase [ubiquinone] iron-sulfur protein 7, mitochondrial | Q60932 (+1) | Ndufs7 | 25 kDa | 5 C | 0.420 | 0.2989 |
| Voltage-dependent anion-selective channel protein 1 | O55143-2 | Vdac1 | 32 kDa | 2 C | 0.410 | 0.14693 |
| Tropomyosin alpha-1 chain | Q9DCZ1 | Tpm1 | 33 kDa | 1 C | 0.408 | 0.36392 |
| Glyoxylate reductase/hydroxypyruvate reductase | Q6NSR8 | Grhpr | 35 kDa | 7 C | 0.389 | 0.16668 |
| Betaine--homocysteine S-methyltransferase 1 | Q7TQ48 | Bhmt | 45 kDa | 8 C | 0.376 | 0.28676 |
| Sarcalumenin | E9Q557 | Srl | 99 kDa | 7 C | 0.371 | 0.09217 |

| | | | | | | |
|---|---|---|---|---|---|---|
| Desmoplakin | Q63918 | Dsp | 333 kDa | 43 C | 0.369 | 0.32042 |
| Ran-specific GTPase-activating protein | Q9WTP6 (+1) | Ranbp1 | 24 kDa | 3 C | 0.361 | 0.12851 |
| Acylpyruvase FAHD1, mitochondrial | Q8R0F8 | Fahd1 | 25 kDa | 6 C | 0.350 | 0.28281 |
| Elongation factor 2 | Q78IK4 | Eef2 | 95 kDa | 7 C | 0.350 | 0.28281 |
| Isovaleryl-CoA dehydrogenase, mitochondrial | Q9JHI5 | Ivd | 46 kDa | 7 C | 0.325 | 0.09788 |
| Regulator of microtubule dynamics protein 1 | Q9DCV4 | Rmdn1 | 35 kDa | 5 C | 0.315 | 0.17844 |
| Nexilin | P51881 | Nexn | 72 kDa | 5 C | 0.302 | 0.12135 |
| Glucose-6-phosphate isomerase | Q7TPW1 | Gpi | 63 kDa | 4 C | 0.293 | 0.13286 |
| Acetyl-coenzyme A synthetase 2-like, mitochondrial | Q78J03 | Acss1 | 75 kDa | 13 C | 0.273 | 0.1029 |
| Methionine-R-sulfoxide reductase B2, mitochondrial | Q99NB1 | Msrb2 | 19 kDa | 9 C | 0.273 | 0.1029 |
| Heterogeneous nuclear ribonucleoprotein A3 | Q8BG05 | Hnrnpa3 | 40 kDa | 4 C | 0.273 | 0.1029 |
| Voltage-dependent anion-selective channel protein 2 | Q60930 | Vdac2 | 32 kDa | 11 C | 0.267 | 0.26411 |
| ATP synthase F(0) complex subunit B1, mitochondrial | Q9CQQ7 | Atp5f1 | 29 kDa | 2 C | 0.267 | 0.26411 |
| O-acetyl-ADP-ribose deacetylase MACROD1 | Q922B1 | Macrod1 | 35 kDa | 10 C | 0.267 | 0.18457 |
| Protein NDRG2 | Q9QYG0 (+1) | Ndrg2 | 41 kDa | 6 C | 0.261 | 0.10264 |
| Heterogeneous nuclear ribonucleoprotein K | P61979 (+2) | Hnrnpk | 51 kDa | 5 C | 0.259 | 0.22757 |
| Dihydropyrimidinase-related protein 1 | P01864 | Crmp1 | 62 kDa | 6 C | 0.256 | 0.41604 |
| Elongation factor 1-alpha 2 | P97427 | Eef1a2 | 50 kDa | 6 C | 0.236 | 0.05934 |
| Acyl-coenzyme A thioesterase 2, mitochondrial | Q9QYR9 | Acot2 | 50 kDa | 4 C | 0.220 | 0.12756 |
| Sorbin and SH3 domain-containing protein 1 | Q62417 | Sorbs1 | 143 kDa | 4 C | 0.220 | 0.06287 |
| ATP-dependent 6-phosphofructokinase, muscle type | P47857 | Pfkm | 85 kDa | 15 C | 0.215 | 0.16972 |
| Glycogen phosphorylase, muscle form | Q9WUB3 | Pygm | 97 kDa | 8 C | 0.204 | 0.10418 |
| Polymeric immunoglobulin receptor | O70570 | Pigr | 85 kDa | 23 C | 0.174 | 0.29235 |
| Fructose-bisphosphate aldolase B | Q91Y97 | Aldob | 40 kDa | 8 C | 0.174 | 0.29235 |
| Succinate--CoA ligase [GDP-forming] subunit beta, mitochondrial | P52825 | Suclg2 | 47 kDa | 5 C | 0.174 | 0.29235 |
| NADH dehydrogenase [ubiquinone] 1 alpha subcomplex subunit 8 | Q9Z2I8 | Ndufa8 | 20 kDa | 8 C | 0.174 | 0.29235 |
| Carnitine O-palmitoyltransferase 2, mitochondrial | O88342 | Cpt2 | 74 kDa | 10 C | 0.174 | 0.29235 |
| WD repeat-containing protein 1 | Q9DCJ5 | Wdr1 | 66 kDa | 12 C | 0.174 | 0.29235 |
| Estradiol 17-beta-dehydrogenase 8 | Q62446 | Hsd17b8 | 27 kDa | 4 C | 0.174 | 0.29235 |
| Eukaryotic initiation factor 4A-II | P50171 (+1) | Eif4a2 | 46 kDa | 4 C | 0.174 | 0.29235 |
| [Pyruvate dehydrogenase (acetyl-transferring)] kinase isozyme 1, mitochondrial | O88935 (+1) | Pdk1 | 49 kDa | 6 C | 0.174 | 0.29235 |
| Monocarboxylate transporter 1 | P10630 (+1) | Slc16a1 | 53 kDa | 11 C | 0.174 | 0.29235 |
| Peptidyl-prolyl cis-trans isomerase FKBP3 | Q8CHT0 | Fkbp3 | 25 kDa | 1 C | 0.174 | 0.29235 |
| Synapsin-1 | Q8BGH2 | Syn1 | 74 kDa | 3 C | 0.174 | 0.29235 |
| Delta-1-pyrroline-5-carboxylate dehydrogenase, mitochondrial | Q02053 | Aldh4a1 | 62 kDa | 8 C | 0.174 | 0.29235 |
| Sorting and assembly machinery component 50 homolog | Q8BVI4 | Samm50 | 52 kDa | 7 C | 0.174 | 0.29235 |
| Ubiquitin-like modifier-activating enzyme 1 | Q8BFP9 | Uba1 | 118 kDa | 21 C | 0.174 | 0.29235 |
| Dihydropteridine reductase | P53986 | Qdpr | 26 kDa | 4 C | 0.174 | 0.29235 |
| Ribosyldihydronicotinamide dehydrogenase [quinone] | P05063 | Nqo2 | 26 kDa | 4 C | 0.125 | 0.12176 |
| Aflatoxin B1 aldehyde reductase member 2 | Q9JI75 | Akr7a2 | 41 kDa | 8 C | 0.125 | 0.12176 |
| Methylcrotonoyl-CoA carboxylase subunit alpha, mitochondrial | Q8CG76 | Mccc1 | 79 kDa | 8 C | 0.125 | 0.12176 |
| Ribosome-recycling factor, mitochondrial | Q99MR8 | Mrrf | 29 kDa | 2 C | 0.125 | 0.12176 |
| Caveolae-associated protein 4 | Q9D6S7 | Cavin4 | 41 kDa | 1 C | 0.125 | 0.12176 |
| Glutamate dehydrogenase 1, mitochondrial | A2AMM0 | Glud1 | 61 kDa | 6 C | 0.121 | 0.29235 |
| Carnitine O-acetyltransferase | P47934 | Crat | 71 kDa | 8 C | 0.121 | 0.29235 |
| Prohibitin | P67778 | Phb | 30 kDa | 1 C | 0.121 | 0.29235 |
| Glutathione S-transferase kappa 1 | P26443 | Gstk1 | 26 kDa | 2 C | 0.121 | 0.29235 |
| NADH dehydrogenase [ubiquinone] 1 beta subcomplex subunit 7 | Q9DCM2 | Ndufb7 | 16 kDa | 4 C | 0.121 | 0.29235 |
| Calcium-binding mitochondrial carrier protein Aralar1 | Q8BH59 | Slc25a12 | 75 kDa | 7 C | 0.101 | 0.15842 |
| Junction plakoglobin | Q02257 | Jup | 82 kDa | 13 C | 0.095 | 0.15768 |
| MICOS complex subunit Mic27 | Q8QZS1 | Apool | 29 kDa | 2 C | 0.095 | 0.15768 |
| 3-hydroxyisobutyryl-CoA hydrolase, mitochondrial | Q64105 | Hibch | 43 kDa | 5 C | 0.095 | 0.15768 |
| Sepiapterin reductase | P41216 | Spr | 28 kDa | 10 C | 0.095 | 0.15768 |
| Glycogen phosphorylase, brain form | Q924X2 | Pygb | 97 kDa | 12 C | 0.085 | 0.05633 |
| D-beta-hydroxybutyrate dehydrogenase, mitochondrial | P68372 | Bdh1 | 38 kDa | 6 C | 0.077 | 0.10043 |
| Tubulin beta-4B chain | Q9CR61 | Tubb4b | 50 kDa | 8 C | 0.073 | 0.05223 |
| Syntaxin-binding protein 1 | O08599 (+1) | Stxbp1 | 68 kDa | 7 C | 0.063 | 0.29235 |
| MICOS complex subunit Mic60 | Q8CAQ8 (+1) | Immt | 84 kDa | 7 C | 0.053 | 0.10046 |
| Calcium-binding mitochondrial carrier protein Aralar2 | Q9QXX4 | Slc25a13 | 74 kDa | 7 C | 0.031 | 0.10565 |
| Apoptosis-inducing factor 1, mitochondrial | Q61316 | Aifm1 | 67 kDa | 4 C | 0.480 | 0.01175 |

| | | | | | | |
|---|---|---|---|---|---|---|
| Trifunctional enzyme subunit alpha, mitochondrial | Q3UZA1 | Hadha | 83 kDa | 12 C | 0.472 | 0.01384 |
| Stress-70 protein, mitochondrial | P05064 | Hspa9 | 73 kDa | 5 C | 0.469 | 0.02493 |
| Cytochrome b-c1 complex subunit 2, mitochondrial | Q91VR2 | Uqcrc2 | 48 kDa | 1 C | 0.451 | 0.02038 |
| Fibrinogen alpha chain | Q99JW2 | Fga | 87 kDa | 13 C | 0.443 | 0.03421 |
| Pyruvate dehydrogenase E1 component subunit beta, mitochondrial | Q9CRB9 | Pdhb | 39 kDa | 6 C | 0.436 | 0.01621 |
| Electron transfer flavoprotein subunit alpha, mitochondrial | P00405 | Etfa | 35 kDa | 6 C | 0.436 | 0.04467 |
| GMP reductase 1 | P19123 | Gmpr | 37 kDa | 9 C | 0.400 | 0.00492 |
| Isoform 2 of Sarcoplasmic/endoplasmic reticulum calcium ATPase 2 | Q8VEM8 | Atp2a2 | 110 kDa | 29 C | 0.400 | 0.04886 |
| Troponin C, slow skeletal and cardiac muscles | P35486 | Tnnc1 | 18 kDa | 2 C | 0.388 | 0.00406 |
| 3-hydroxyisobutyrate dehydrogenase, mitochondrial | Q8BMF4 | Hibadh | 35 kDa | 12 C | 0.386 | 0.01912 |
| Pyruvate dehydrogenase E1 component subunit alpha, somatic form, mitochondrial | Q03265 | Pdha1 | 43 kDa | 12 C | 0.384 | 0.01725 |
| Probable aminopeptidase NPEPL1 | O35490 | Npepl1 | 56 kDa | 16 C | 0.384 | 0.04168 |
| Dihydrolipoyllysine-residue acetyltransferase component of pyruvate dehydrogenase com | P62259 | Dlat | 68 kDa | 10 C | 0.383 | 0.03211 |
| Acyl-coenzyme A thioesterase 13 | Q9CQR4 | Acot13 | 15 kDa | 2 C | 0.368 | 0.04929 |
| Caveolae-associated protein 2 | P16858 | Cavin2 | 47 kDa | 1 C | 0.368 | 0.04929 |
| Glyceraldehyde-3-phosphate dehydrogenase | P34022 | Gapdh | 36 kDa | 5 C | 0.365 | 0.02427 |
| Adenylate kinase 2, mitochondrial | P58252 | Ak2 | 26 kDa | 5 C | 0.360 | 0.0247 |
| Phosphoglucomutase-1 | Q9D0F9 | Pgm1 | 61 kDa | 10 C | 0.331 | 0.0478 |
| ADP/ATP translocase 2 | P58771 | Slc25a5 | 33 kDa | 4 C | 0.308 | 0.00313 |
| Phosphate carrier protein, mitochondrial | P06745 | Slc25a3 | 40 kDa | 8 C | 0.289 | 0.00109 |
| FAD-linked sulfhydryl oxidase ALR | P56213 | Gfer | 23 kDa | 8 C | 0.263 | 0.03356 |
| von Willebrand factor A domain-containing protein 1 | Q8R2Z5 | Vwa1 | 45 kDa | 2 C | 0.263 | 0.03356 |
| Heat shock protein HSP 90-beta | P11499 | Hsp90ab1 | 83 kDa | 6 C | 0.259 | 0.00711 |
| UPF0160 protein MYG1, mitochondrial | Q9JK81 | Myg1 | 43 kDa | 7 C | 0.224 | 0.01434 |
| Voltage-dependent anion-selective channel protein 3 | Q60931 | Vdac3 | 31 kDa | 6 C | 0.213 | 0.02614 |
| ATP synthase subunit O, mitochondrial | Q9DB20 | Atp5o | 23 kDa | 1 C | 0.189 | 0.01551 |
| NADH dehydrogenase [ubiquinone] 1 alpha subcomplex subunit 10, mitochondrial | Q99LC3 | Ndufa10 | 41 kDa | 5 C | 0.187 | 0.03985 |
| Nucleoside diphosphate-linked moiety X motif 8 | Q9CR24 | Nudt8 | 23 kDa | 4 C | 0.187 | 0.03985 |
| Tubulin alpha-1B chain | P05213 | Tuba1b | 50 kDa | 12 C | 0.184 | 0.04019 |
| Isocitrate dehydrogenase [NAD] subunit gamma 1, mitochondrial | P70404 | Idh3g | 43 kDa | 7 C | 0.167 | 0.00226 |
| Mitochondrial 2-oxoglutarate/malate carrier protein | P62631 | Slc25a11 | 34 kDa | 3 C | 0.154 | 0.00389 |
| CAP-Gly domain-containing linker protein 1 | Q9CR62 | Clip1 | 156 kDa | 11 C | 0.147 | 0.03927 |
| 3-hydroxyacyl-CoA dehydrogenase type-2 | Q922J3 | Hsd17b10 | 27 kDa | 2 C | 0.142 | 0.02079 |
| Tubulin alpha-4A chain | O08756 | Tuba4a | 50 kDa | 13 C | 0.132 | 0.00718 |
| Fructose-bisphosphate aldolase C | P68368 | Aldoc | 39 kDa | 7 C | 0.131 | 0.00174 |
| Ubiquinone biosynthesis protein COQ9, mitochondrial | Q8K1Z0 | Coq9 | 35 kDa | 3 C | 0.102 | 0.01554 |
| Lipoamide acyltransferase component of branched-chain alpha-keto acid dehydrogenase c | P53395 | Dbt | 53 kDa | 6 C | 0.098 | 0.03004 |
| Sodium/potassium-transporting ATPase subunit beta-1 | P14094 | Atp1b1 | 35 kDa | 7 C | 0.098 | 0.03004 |
| Long-chain-fatty-acid--CoA ligase 1 | Q8CI94 | Acsl1 | 78 kDa | 18 C | 0.093 | 0.00991 |
| Carnitine O-palmitoyltransferase 1, muscle isoform | Q9Z2I9 | Cpt1b | 88 kDa | 16 C | 0.080 | 0.00464 |
| Succinate--CoA ligase [ADP-forming] subunit beta, mitochondrial | Q80XN0 | Sucla2 | 50 kDa | 6 C | 0.080 | 0.02866 |
| Propionyl-CoA carboxylase beta chain, mitochondrial | Q99MN9 | Pccb | 58 kDa | 11 C | 0.044 | 0.01379 |
| Sodium/potassium-transporting ATPase subunit alpha-1 | Q8VDN2 | Atp1a1 | 113 kDa | 23 C | 0.025 | 0.00183 |
