## Supplemental Table 11 for "Dietary restriction transforms the protein sulfhydrome in a tissue-specific and cystathionine γ-lyase-dependent manner"

| Supplemental Table 11: Dietary Impact on the Serum Sulfhydrome | | | | | | |
|---|---|---|---|---|---|---|
| Protein Name | Accession Number | Alternate ID | Molecular Weight | Cysteine Residues | DR/AL Spectral Count Ratio | P-value |
| Cytochrome c, somatic | P62897 | Cycs | 12 kDa | 2 C | 8.600 | 0.272392948 |
| Ig heavy chain V region 3 | P01749 | Ighv1-61 | 13 kDa | 3 C | 2.571 | 0.350829714 |
| Afamin | O89020 | Afm | 69 kDa | 34 C | 1.829 | 0.013587476 |
| Fetuin-B | Q9QXC1 | Fetub | 43 kDa | 15 C | 1.802 | 0.052756178 |
| Thyroxine-binding globulin | P61939 | Serpina7 | 47 kDa | 5 C | 1.774 | 0.43607882 |
| Triosephosphate isomerase | P17751 | Tpi1 | 32 kDa | 9 C | 1.723 | 0.013353797 |
| Coagulation factor XIII B chain | Q07968 | F13b | 76 kDa | 40 C | 1.689 | 0.101847512 |
| Coagulation factor XIII A chain | Q8BH61 | F13a1 | 83 kDa | 2 C | 1.569 | 0.359574378 |
| Transforming growth factor-beta-induced protein ig-h3 | P82198 | Tgfbi | 75 kDa | 11 C | 1.527 | 0.54904678 |
| Lysosomal protective protein | P16675 | Ctsa | 54 kDa | 11 C | 1.477 | 0.753244047 |
| Histidine-rich glycoprotein | Q9ESB3 | Hrg | 59 kDa | 17 C | 1.456 | 0.041393898 |
| Ig kappa chain V-VI region NQ2-17.4.1 | P04940 | | 12 kDa | 2 C | 1.422 | 0.570069638 |
| Zinc-alpha-2-glycoprotein | Q64726 | Azgp1 | 35 kDa | 4 C | 1.400 | 0.439152536 |
| Angiotensinogen | P11859 | Agt | 52 kDa | 5 C | 1.378 | 0.156036858 |
| Beta-2-glycoprotein 1 | Q01339 | Apoh | 39 kDa | 23 C | 1.352 | 0.053936283 |
| Complement factor H | P06909 | Cfh | 139 kDa | 82 C | 1.331 | 0.019914874 |
| Isoform 3 of Interleukin-1 receptor accessory protein | Q61730-3 | Il1rap | 79 kDa | 12 C | 1.318 | 0.126870367 |
| Protein Z-dependent protease inhibitor | Q8R121 | Serpina10 | 52 kDa | 2 C | 1.316 | 0.553817402 |
| Murinoglobulin-1 | P28665 | Mug1 | 165 kDa | 25 C | 1.310 | 0.031849989 |
| Ig kappa chain V-II region 26-10 | P01631 | | 12 kDa | 2 C | 1.300 | 0.460841443 |
| Hemoglobin subunit beta-1 | P02088 | Hbb-b1 | 16 kDa | 2 C | 1.297 | 0.286190589 |
| Insulin-like growth factor-binding protein complex acid labile subunit | P70389 | Igfals | 67 kDa | 14 C | 1.292 | 0.019674848 |
| Clusterin | Q06890 | Clu | 52 kDa | 11 C | 1.255 | 0.172364527 |
| Ig gamma-2B chain C region | P01867 | Igh-3 | 44 kDa | 13 C | 1.247 | 0.326021828 |
| Inter-alpha-trypsin inhibitor heavy chain H2 | Q61703 | Itih2 | 106 kDa | 7 C | 1.236 | 0.308473704 |
| Hepatocyte growth factor activator | Q9R098 | Hgfac | 71 kDa | 41 C | 1.236 | 0.013776484 |
| Protein AMBP | Q07456 | Ambp | 39 kDa | 16 C | 1.233 | 0.199618666 |
| Ig heavy chain V region 102 | P01750 | | 13 kDa | 3 C | 1.231 | 0.396934567 |
| Inter alpha-trypsin inhibitor, heavy chain 4 | A6X935 | Itih4 | 105 kDa | 3 C | 1.231 | 0.046416158 |
| Ig gamma-2A chain C region secreted form | P01864 | | 37 kDa | 9 C | 1.217 | 0.20984574 |
| Complement factor B | P04186 | Cfb | 85 kDa | 20 C | 1.200 | 0.197022072 |
| Ig heavy chain V region B1-8/186-2 | P01751 | Ighv1-72 | 15 kDa | 3 C | 1.200 | 0.563027804 |
| Inter-alpha-trypsin inhibitor heavy chain H1 | Q61702 | Itih1 | 101 kDa | 8 C | 1.200 | 0.263653334 |
| Inter-alpha-trypsin inhibitor heavy chain H3 | Q61704 | Itih3 | 99 kDa | 6 C | 1.200 | 0.616704191 |
| Cholinesterase | Q03311 | Bche | 68 kDa | 9 C | 1.200 | 0.827125331 |
| Fibrinogen alpha chain | E9PV24 | Fga | 87 kDa | 13 C | 1.182 | 0.380361602 |
| Vitamin K-dependent protein C | P33587 | Proc | 52 kDa | 24 C | 1.171 | 0.681904416 |
| Serum albumin | P07724 | Alb | 69 kDa | 36 C | 1.156 | 0.02549729 |
| Fibrinogen gamma chain | Q8VCM7 | Fgg | 49 kDa | 12 C | 1.150 | 0.466588428 |
| Complement C1q subcomponent subunit B | P14106 | C1qb | 27 kDa | 4 C | 1.148 | 0.890942227 |
| Adiponectin | Q60994 | Adipoq | 27 kDa | 2 C | 1.143 | 0.292351992 |
| Selenoprotein P | P70274 | Selenop | 43 kDa | 18 C | 1.123 | 0.443290017 |
| Glutathione peroxidase 3 | P46412 | Gpx3 | 25 kDa | 3 C | 1.113 | 0.346380429 |
| Prothrombin | P19221 | F2 | 70 kDa | 25 C | 1.112 | 0.263976603 |
| Plasma kallikrein | P26262 | Klkb1 | 71 kDa | 37 C | 1.107 | 0.265556347 |
| Carboxypeptidase N subunit 2 | Q9DBB9 | Cpn2 | 60 kDa | 15 C | 1.103 | 0.373670035 |
| EGF-containing fibulin-like extracellular matrix protein 1 | Q8BPB5 | Efemp1 | 55 kDa | 40 C | 1.080 | 0.870943338 |
| Ig kappa chain V-VI region NQ2-6.1 | P04945 | | 12 kDa | 2 C | 1.078 | 0.644407158 |
| C4b-binding protein | P08607 | C4bpa | 52 kDa | 26 C | 1.077 | 0.531972046 |
| Ig mu chain C region | P01872 | Ighm | 50 kDa | 19 C | 1.074 | 0.771637737 |
| Inhibitor of carbonic anhydrase | Q9DBD0 | Ica | 77 kDa | 35 C | 1.073 | 0.341185392 |
| Fibrinogen beta chain | Q8K0E8 | Fgb | 55 kDa | 12 C | 1.072 | 0.689914861 |
| Kininogen-1 | O08677 | Kng1 | 73 kDa | 19 C | 1.071 | 0.333899141 |
| Ig kappa chain V-V region L6 (Fragment) | P01638 | | 13 kDa | 3 C | 1.067 | 0.563027804 |
| Lumican | P51885 | Lum | 38 kDa | 7 C | 1.067 | 0.684528336 |
| Ig heavy chain V region MOPC 173 | P01812 | | 13 kDa | 2 C | 1.067 | 0.852853872 |
| Pregnancy zone protein | Q61838 | Pzp | 166 kDa | 24 C | 1.056 | 0.591465068 |
| Mannose-binding protein C | P41317 | Mbl2 | 26 kDa | 7 C | 1.056 | 0.639396881 |
| Serotransferrin | Q921I1 | Tf | 77 kDa | 38 C | 1.055 | 0.426656862 |

| | | | | | | |
|---|---|---|---|---|---|---|
| Complement C3 | P01027 | C3 | 186 kDa | 27 C | 1.051 | 0.573709556 |
| Serum paraoxonase/arylesterase 1 | P52430 | Pon1 | 40 kDa | 3 C | 1.050 | 0.752840563 |
| Ig kappa chain V-II region 2S1.3 | P01629 | | 12 kDa | 2 C | 1.042 | 0.970111629 |
| Complement factor I | Q61129 | Cfi | 67 kDa | 40 C | 1.040 | 0.725035956 |
| Alpha-2-HS-glycoprotein | P29699 | Ahsg | 37 kDa | 14 C | 1.036 | 0.680712116 |
| Peroxiredoxin-1 | P35700 | Prdx1 | 22 kDa | 4 C | 1.029 | 0.919578039 |
| Ig heavy chain V region AC38 205.12 | P06330 | | 13 kDa | 2 C | 1.025 | 0.887127985 |
| Complement component C8 gamma chain | Q8VCG4 | C8g | 23 kDa | 3 C | 1.020 | 0.890978708 |
| Ceruloplasmin | Q61147 | Cp | 121 kDa | 14 C | 1.020 | 0.938968396 |
| Heparin cofactor 2 | P49182 | Serpind1 | 54 kDa | 5 C | 1.018 | 0.950902942 |
| Serine protease inhibitor A3K | P07759 | Serpina3k | 47 kDa | 4 C | 1.016 | 0.737197039 |
| Corticosteroid-binding globulin | Q06770 | Serpina6 | 45 kDa | 3 C | 1.009 | 0.97925306 |
| Plasminogen | P20918 | Plg | 91 kDa | 48 C | 0.983 | 0.86124277 |
| Vitamin D-binding protein | P21614 | Gc | 54 kDa | 28 C | 0.978 | 0.790082752 |
| Alpha-1-antitrypsin 1-2 | P22599 | Serpina1b | 46 kDa | 3 C | 0.975 | 0.882951638 |
| Carboxylesterase 1C | P23953 | Ces1c | 61 kDa | 5 C | 0.972 | 0.832737964 |
| Antithrombin-III | P32261 | Serpinc1 | 52 kDa | 9 C | 0.964 | 0.771257542 |
| Mannose-binding protein A | P39039 | Mbl1 | 25 kDa | 8 C | 0.960 | 0.664658218 |
| Phosphatidylcholine-sterol acyltransferase | P16301 | Lcat | 50 kDa | 6 C | 0.960 | 0.878616768 |
| Ig kappa chain C region | P01837 | | 12 kDa | 3 C | 0.954 | 0.845082003 |
| Gelsolin | P13020 (+1) | Gsn | 86 kDa | 7 C | 0.953 | 0.505375212 |
| Peroxiredoxin-2 | Q61171 | Prdx2 | 22 kDa | 3 C | 0.952 | 0.790416234 |
| Haptoglobin | Q61646 | Hp | 39 kDa | 9 C | 0.948 | 0.921785709 |
| Collagen alpha-2(I) chain | Q01149 | Col1a2 | 130 kDa | 9 C | 0.945 | 0.684528336 |
| Ig heavy chain V region 345 | P18526 | | 13 kDa | 3 C | 0.945 | 0.684528336 |
| Ig lambda-1 chain C region | P01843 | | 12 kDa | 3 C | 0.943 | 0.729434311 |
| Serine protease inhibitor A3N | Q91WP6 | Serpina3n | 47 kDa | 3 C | 0.936 | 0.700492989 |
| Mannan-binding lectin serine protease 2 | Q91WP0 | Masp2 | 76 kDa | 26 C | 0.933 | 0.729434311 |
| Properdin | P11680 | Cfp | 50 kDa | 44 C | 0.928 | 0.475797239 |
| Alpha-2-antiplasmin | Q61247 | Serpinf2 | 55 kDa | 4 C | 0.923 | 0.516489552 |
| Alpha-1-antitrypsin 1-4 | Q00897 | Serpina1d | 46 kDa | 3 C | 0.911 | 0.646830139 |
| Sulfhydryl oxidase 1 | Q8BND5 (+2) | Qsox1 | 83 kDa | 14 C | 0.910 | 0.583955183 |
| Serine protease inhibitor A3M | Q03734 | Serpina3m | 47 kDa | 3 C | 0.903 | 0.51406374 |
| Plasma protease C1 inhibitor | P97290 | Serping1 | 56 kDa | 5 C | 0.900 | 0.193006088 |
| Vitronectin | P29788 | Vtn | 55 kDa | 14 C | 0.896 | 0.431842872 |
| Hemoglobin subunit alpha | P01942 | Hba | 15 kDa | 1 C | 0.880 | 0.227452818 |
| Ficolin-1 | O70165 | Fcn1 | 36 kDa | 10 C | 0.880 | 0.502354017 |
| Carbonic anhydrase 2 | P00920 | Ca2 | 29 kDa | 2 C | 0.868 | 0.782660754 |
| Complement C1r-A subcomponent | Q8CG16 | C1ra | 80 kDa | 26 C | 0.863 | 0.463860997 |
| Complement component C8 beta chain | Q8BH35 | C8b | 66 kDa | 32 C | 0.863 | 0.482777903 |
| Alpha-1-antitrypsin 1-1 | P07758 | Serpina1a | 46 kDa | 3 C | 0.862 | 0.483828653 |
| Coagulation factor XII | Q80YC5 | F12 | 66 kDa | 40 C | 0.853 | 0.574178826 |
| CD5 antigen-like | Q9QWK4 | Cd5l | 39 kDa | 26 C | 0.850 | 0.196696052 |
| Ig gamma-3 chain C region | P03987 (+1) | | 44 kDa | 10 C | 0.848 | 0.350795203 |
| Complement component C8 alpha chain | Q8K182 | C8a | 66 kDa | 29 C | 0.845 | 0.292351992 |
| Alpha-1-antitrypsin 1-5 | Q00898 | Serpina1e | 46 kDa | 4 C | 0.839 | 0.451997051 |
| Annexin A2 | P07356 | Anxa2 | 39 kDa | 5 C | 0.825 | 0.817523442 |
| Phospholipid transfer protein | P55065 | Pltp | 54 kDa | 4 C | 0.800 | 0.292351992 |
| Collagen alpha-1(I) chain | P11087 | Col1a1 | 138 kDa | 18 C | 0.800 | 0.328187341 |
| Ig kappa chain V-V region MOPC 41 | P01639 | Gm5571 | 14 kDa | 3 C | 0.800 | 0.368981333 |
| Ig heavy chain V regions TEPC 15/S107/HPCM1/HPCM2/HPCM3 | P01787 | | 14 kDa | 2 C | 0.800 | 0.49325971 |
| Fibronectin | P11276 | Fn1 | 273 kDa | 64 C | 0.800 | 0.67284459 |
| Ig kappa chain V-III region ABPC 22/PC 9245 | P01662 | | 12 kDa | 2 C | 0.800 | 0.704617585 |
| Ig heavy chain V region 93G7 | P01746 | | 16 kDa | 2 C | 0.800 | 0.407083822 |
| Ig kappa chain V-IV region S107B | P01680 | | 14 kDa | 2 C | 0.800 | 0.407083822 |
| Pyrethroid hydrolase Ces2e | Q8BK48 | Ces2e | 62 kDa | 7 C | 0.800 | 0.481558033 |
| Complement C4-B | P01029 | C4b | 193 kDa | 29 C | 0.792 | 0.158859618 |
| Complement C5 | P06684 | C5 | 189 kDa | 30 C | 0.789 | 0.200523726 |
| Hemopexin | Q91X72 | Hpx | 51 kDa | 13 C | 0.772 | 0.011911246 |
| Myosin-6 | Q02566 | Myh6 | 224 kDa | 14 C | 0.764 | 0.407083822 |
| Coagulation factor X | O88947 | F10 | 54 kDa | 25 C | 0.762 | 0.240110429 |
| Ig kappa chain V-VI region XRPC 44 | P01675 (+1) | | 12 kDa | 2 C | 0.753 | 0.063924123 |
| Ig gamma-1 chain C region secreted form | P01868 (+1) | Ighg1 | 36 kDa | 12 C | 0.750 | 0.412577511 |

| | | | | | | |
|---|---|---|---|---|---|---|
| Immunoglobulin J chain | P01592 | Jchain | 18 kDa | 8 C | 0.745 | 0.195150657 |
| Ig kappa chain V-V region MOPC 173 | P01643 | | 12 kDa | 2 C | 0.738 | 0.010844009 |
| Transthyretin | P07309 | Ttr | 16 kDa | 2 C | 0.727 | 0.144839256 |
| Ig alpha chain C region | P01878 | | 37 kDa | 13 C | 0.724 | 0.214465003 |
| Ig kappa chain V-V region HP R16.7 | P01644 (+1) | | 12 kDa | 2 C | 0.670 | 0.055657465 |
| Myosin-7 | Q91Z83 | Myh7 | 223 kDa | 14 C | 0.667 | 0.257166808 |
| Complement factor D | P03953 (+1) | Cfd | 28 kDa | 9 C | 0.667 | 0.292351992 |
| Phosphatidylinositol-glycan-specific phospholipase D | O70362 | Gpld1 | 93 kDa | 10 C | 0.661 | 0.026562152 |
| Actin, cytoplasmic 1 | P60710 | Actb | 42 kDa | 6 C | 0.659 | 0.214465003 |
| Ig heavy chain V-III region J606 | P01801 | | 13 kDa | 2 C | 0.655 | 0.201631928 |
| H-2 class I histocompatibility antigen, Q10 alpha chain | P01898 | H2-Q10 | 37 kDa | 4 C | 0.653 | 0.089938561 |
| Ig kappa chain V-III region PC 7043 | P01665 | | 12 kDa | 2 C | 0.648 | 0.082204885 |
| Complement C1s-A subcomponent | Q8CG14 | C1sa | 77 kDa | 27 C | 0.631 | 0.276020719 |
| Leukemia inhibitory factor receptor | P42703 | Lifr | 123 kDa | 23 C | 0.617 | 0.098036165 |
| Ig lambda-1 chain V region S43 | P01727 | | 14 kDa | 2 C | 0.610 | 0.193221181 |
| Extracellular matrix protein 1 | Q61508 | Ecm1 | 63 kDa | 29 C | 0.586 | 0.444898457 |
| Ig kappa chain V-V region K2 (Fragment) | P01635 | | 13 kDa | 3 C | 0.554 | 0.106675452 |
| Ig lambda-2 chain C region | P01844 | Iglc2 | 11 kDa | 3 C | 0.554 | 0.002503335 |
| Ig kappa chain V-V region MOPC 149 | P01636 | | 12 kDa | 2 C | 0.533 | 0.012654499 |
| Thrombospondin-1 | P35441 | Thbs1 | 130 kDa | 70 C | 0.528 | 0.259806123 |
| Polymeric immunoglobulin receptor | O70570 | Pigr | 85 kDa | 23 C | 0.518 | 0.279340105 |
| Ig kappa chain V-III region PC 2880/PC 1229 | P01654 | | 12 kDa | 2 C | 0.504 | 0.210673935 |
| Ig kappa chain V-V region L7 (Fragment) | P01642 | Gm10881 | 13 kDa | 2 C | 0.500 | 0.151399763 |
| Creatine kinase M-type | P07310 | Ckm | 43 kDa | 8 C | 0.480 | 0.157024205 |
| Ig heavy chain V region 3-6 | P18531 | Ighv3-6 | 13 kDa | 2 C | 0.480 | 0.136589638 |
| Mannan-binding lectin serine protease 1 | P98064 | Masp1 | 80 kDa | 29 C | 0.448 | 0.229165553 |
| Epidermal growth factor receptor | Q01279 | Egfr | 135 kDa | 60 C | 0.415 | 0.220991111 |
| Apolipoprotein E | P08226 | Apoe | 36 kDa | 1 C | 0.310 | 0.105650339 |
| Major urinary protein 1 | P11588 | Mup1 | 21 kDa | 5 C | 0.302 | 0.121352611 |
| Fructose-bisphosphate aldolase A | P05064 | Aldoa | 39 kDa | 8 C | 0.284 | 0.13623991 |
| Desmoplakin | E9Q557 | Dsp | 333 kDa | 43 C | 0.222 | 0.38744688 |
| Fructose-bisphosphate aldolase B | Q91Y97 | Aldob | 40 kDa | 8 C | 0.174 | 0.292351992 |
| Beta-enolase | P21550 | Eno3 | 47 kDa | 6 C | 0.174 | 0.292351992 |
| Cytosol aminopeptidase | Q9CPY7 | Lap3 | 56 kDa | 7 C | 0.140 | 0.257370431 |
| Isoform 3 of Periostin | Q62009-3 | Postn | 90 kDa | 12 C | 0.125 | 0.121758646 |
| Complement component C9 | P06683 | C9 | 62 kDa | 22 C | 0.485 | 0.001380299 |
| Vascular cell adhesion protein 1 | P29533 | Vcam1 | 81 kDa | 20 C | 0.427 | 0.046405029 |
| Betaine--homocysteine S-methyltransferase 1 | O35490 | Bhmt | 45 kDa | 8 C | 0.240 | 0.016593483 |
