## Supplemental Table 12 for "Dietary restriction transforms the protein sulfhydrome in a tissue-specific and cystathionine γ-lyase-dependent manner"

| Supplemental Table 12: Accession Numbers for the 78 Common Sulfhydrated Proteins in Heart and Serum |
|---|
| P01750 |
| P01844 |
| P13020 (+1) |
| P20918 |
| Q9QXC1 |
| P06909 |
| P32261 |
| P28665 |
| P21614 |
| Q9R098 |
| Q06890 |
| P41317 |
| P11680 |
| P04186 |
| Q8K182 |
| Q01339 |
| Q61646 |
| P01027 |
| P29788 |
| P29699 |
| P02088 |
| P01843 |
| P23953 |
| P70274 |
| P01872 |
| P01635 |
| Q61838 |
| Q9ESB3 |
| P01942 |
| P39039 |
| O70165 |
| P22599 |
| Q61129 |
| P06330 |
| Q00897 |
| P01029 |
| Q8VCG4 |
| P01592 |
| O08677 |
| Q91X72 |
| P00920 |
| Q9DBD0 |
| P07759 |
| Q921I1 |
| Q61171 |
| P01878 |

| |
|---|
| P07309 |
| P04945 |
| P01631 |
| Q00898 |
| Q8K0E8 |
| Q8VCM7 |
| P01837 |
| P46412 |
| Q9QWK4 |
| P17751 |
| P21550 |
| P01639 |
| Q8BH35 |
| P01868 (+1) |
| P07724 |
| P01644 (+1) |
| P35700 |
| P07310 |
| P07356 |
| Q9CPY7 |
| P62897 |
| P01642 |
| Q60994 |
| Q02566 |
| P05064 |
| Q01279 |
| E9PV24 |
| O35490 |
| E9Q557 |
| P01864 |
| O70570 |
| Q91Y97 |
