## Supplemental Table 13 for "Dietary restriction transforms the protein sulfhydrome in a tissue-specific and cystathionine γ-lyase-dependent manner"

| Supplemental Table 13: Sulfhydrated Protein Pathway Enrichment in Heart | | | |
|---|---|---|---|
| KEGG Pathway (DR) | p-val (adj) | -LOG10(p-val adj) | Involved Genes |
| Ferroptosis | 0.00173 | 2.76 | FTH1,GCLM,FTL1 |
| Complement and coagulation cascades | 0.0000084 | 5.08 | C3,SERPINC1,CFI,SERPINA1D,CFB |
| Staphylococcus aureus infection | 0.00421 | 2.38 | C3,CFI,CFB |
| KEGG Pathway (Unchanged) | p-val (adj) | -LOG10(p-val adj) | Involved Genes |
| Carbon metabolism | 4.45E-20 | 19.35 | DLAT,CS,SDHB,MDH2,MDH1,OGDH,DLD,ALDH6A1,SDHA,ACO2,TPI1,GOT1,ECHS1,TALDO1,FH,CAT,ACADS,IDH2,PDHA1,ACAT1,IDH3A,PKM,PGP,ENO3,PGK1,ACADM,ENO1,FBP1 |
| Glutathione metabolism | 0.00638 | 2.20 | GPX3,GSTO1,IDH2,GSR,LAP3,GSTM2,GSTM1,GPX1 |
| Fatty acid degradation | 0.000000469 | 6.33 | ECI1,ECHS1,ACADL,HADH,ALDH2,ACADS,ACAT1,ACAA2,ALDH7A1,HADHB,ACADM |
| Metabolic pathways | 3.36E-15 | 14.47 | COX5A,DLAT,NDUFA9,CKB,CS,SDHB,APIP,PDHX,NDUFS2,NDUFA2,MDH2,MDH1,OGDH,DLD,NME2,ALDH6A1,AUH,SDHA,NDUFS6,CKMT2,MCCC2,NDUFS4,ACO2,CYC1,CPOX,TPI1,NDUFV2,CNDP2,GOT1,ECHS1,TALDO1,ACADL,FH,SEPHS1,PRDX6,AK1,IMPA1,HADH,ALAD,CMPK1,ALDH2,ACADS,LDHB,CKM,IDH2,BCAT2,PDHA1,COX4I1,ACAT1,PTS,IDH3A,PKM,ATP5H,ALDH5A1,ACAA2,NME1,NDUFV1,UQCRFS1,LAP3,NDUFB10,BLVRB,PGP,ACP1,ALDH1A1,ALDH7A1,CES1D,HADHB,ENO3,COX5B,PGK1,ACADM,LDHA,ENO1,MTCO2,FBP1 |
| Arginine and proline metabolism | 0.00124 | 2.91 | CKB,CKMT2,CNDP2,GOT1,ALDH2,CKM,LAP3,ALDH7A1 |
| Propanoate metabolism | 0.000266 | 3.58 | DLD,ALDH6A1,ECHS1,LDHB,ACAT1,ACADM,LDHA |
| Huntington disease | 1.05E-08 | 7.98 | COX5A,NDUFA9,SOD2,SDHB,NDUFS2,NDUFA2,SDHA,NDUFS6,NDUFS4,CYC1,SOD1,NDUFV2,SLC25A4,COX4I1,ATP5H,NDUFV1,UQCRFS1,NDUFB10,COX5B,CYCS,GPX1,MTCO2 |
| Biosynthesis of amino acids | 0.00000658 | 5.18 | CS,ACO2,TPI1,GOT1,TALDO1,IDH2,BCAT2,IDH3A,PKM,ENO3,PGK1,ENO1 |
| Thermogenesis | 0.0000415 | 4.38 | COX5A,NDUFA9,SDHB,NDUFS2,NDUFA2,SDHA,NDUFS6,NDUFS4,CYC1,NDUFV2,COX4I1,ATP5H,NDUFV1,UQCRFS1,NDUFB10,GRB2,COX5B,ACTG1,MTCO2 |
| Glyoxylate and dicarboxylate metabolism | 0.0000124 | 4.91 | CS,MDH2,MDH1,DLD,ACO2,CAT,ACAT1,PGP |
| Cardiac muscle contraction | 0.000737 | 3.13 | COX5A,CYC1,COX4I1,TNNI3,UQCRFS1,MYH6,MYL3,COX5B,MTCO2,ACTC1 |
| Butanoate metabolism | 0.00156 | 2.81 | OXCT1,ECHS1,HADH,ACADS,ACAT1,ALDH5A1 |
| Glycolysis / Gluconeogenesis | 0.000000115 | 6.94 | DLAT,DLD,TPI1,ALDH2,LDHB,PDHA1,PKM,ALDH7A1,ENO3,PGK1,LDHA,ENO1,FBP1 |
| Parkinson disease | 2.32E-09 | 8.63 | COX5A,NDUFA9,SDHB,NDUFS2,NDUFA2,SDHA,NDUFS6,NDUFS4,CYC1,NDUFV2,PARK7,SLC25A4,COX4I1,ATP5H,NDUFV1,UQCRFS1,NDUFB10,COX5B,CYCS,MTCO2 |
| Valine, leucine and isoleucine degradation | 2.19E-11 | 10.66 | DLD,ALDH6A1,AUH,MCCC2,OXCT1,ECHS1,HADH,ALDH2,ACADS,BCAT2,ACAT1,ACAA2,ALDH7A1,HADHB,ACADM |
| Non-alcoholic fatty liver disease (NAFLD) | 0.000000271 | 6.57 | COX5A,NDUFA9,SDHB,NDUFS2,NDUFA2,SDHA,NDUFS6,NDUFS4,CYC1,ADIPOQ,NDUFV2,COX4I1,NDUFV1,UQCRFS1,NDUFB10,COX5B,CYCS,MTCO2 |
| Tryptophan metabolism | 0.00358 | 2.45 | OGDH,ECHS1,CAT,HADH,ALDH2,ACAT1,ALDH7A1 |
| Cysteine and methionine metabolism | 0.00551 | 2.26 | APIP,MDH2,MDH1,GOT1,LDHB,BCAT2,LDHA |
| Pyruvate metabolism | 3.9E-14 | 13.41 | DLAT,MDH2,MDH1,DLD,GLO1,HAGH,FH,ALDH2,LDHB,PDHA1,ACAT1,PKM,ALDH7A1,ACYP2,LDHA |
| Alzheimer disease | 0.00000304 | 5.52 | COX5A,NDUFA9,SDHB,NDUFS2,NDUFA2,SDHA,NDUFS6,NDUFS4,CYC1,NDUFV2,COX4I1,ATP5H,NDUFV1,UQCRFS1,NDUFB10,COX5B,CYCS,MTCO2 |
| Citrate cycle (TCA cycle) | 2.46E-12 | 11.61 | DLAT,CS,SDHB,MDH2,MDH1,OGDH,DLD,SDHA,ACO2,FH,IDH2,PDHA1,IDH3A |
| Proteasome | 5.57E-10 | 9.25 | PSMB4,PSMB1,PSMB6,PSMA6,PSMB5,PSME1,PSMA7,PSMB2,PSMA1,PSMA4,PSMA3,PSMA5,PSMB3 |
| beta-Alanine metabolism | 0.00524 | 2.28 | ALDH6A1,CNDP2,ECHS1,ALDH2,ALDH7A1,ACADM |
| Oxidative phosphorylation | 4.39E-08 | 7.36 | COX5A,NDUFA9,SDHB,NDUFS2,NDUFA2,SDHA,NDUFS6,NDUFS4,CYC1,NDUFV2,PPA2,COX4I1,ATP5H,NDUFV1,UQCRFS1,NDUFB10,COX5B,MTCO2 |
| Complement and coagulation cascades | 0.00139 | 2.86 | C8G,C8B,FGB,FGG,C8A,MBL1,SERPINA1A,SERPINA1B,SERPINA1E,C4B |
| Fatty acid metabolism | 0.00108 | 2.97 | ECHS1,ACADL,HADH,ACADS,ACAT1,ACAA2,HADHB,ACADM |
| 2-Oxocarboxylic acid metabolism | 0.000164 | 3.79 | CS,ACO2,GOT1,IDH2,BCAT2,IDH3A |
| KEGG Pathway (AL) | p-val (adj) | -LOG10(p-val adj) | Involved Genes |
| Metabolic pathways | 0.000049 | 4.31 | DLAT,DBT,IDH3G,ALDOC,ACSL1,PDHB,SUCLA2,HSD17B10,HADHA,NDUFA10,AK2,PGM1,HIBADH,UQCRC2,PDHA1,PCCB,GAPDH,HADHB |
| cGMP-PKG signaling pathway | 0.0385 | 1.41 | VDAC3,SLC25A5,ATP1B1,ATP2A2,ATP1A1 |
| Fatty acid metabolism | 0.00367 | 2.44 | ACSL1,HADHA,HADHB,CPT1B |
| Glycolysis / Gluconeogenesis | 0.0000176 | 4.75 | DLAT,ALDOC,PDHB,PGM1,PDHA1,GAPDH |
| Alzheimer disease | 0.0475 | 1.32 | HSD17B10,NDUFA10,ATP2A2,UQCRC2,GAPDH |
| Citrate cycle (TCA cycle) | 0.0000104 | 4.98 | DLAT,IDH3G,PDHB,SUCLA2,PDHA1 |
| Fatty acid degradation | 0.00314 | 2.50 | ACSL1,HADHA,HADHB,CPT1B |
| Pyruvate metabolism | 0.0323 | 1.49 | DLAT,PDHB,PDHA1 |
| Cardiac muscle contraction | 0.00122 | 2.91 | ATP1B1,ATP2A2,UQCRC2,ATP1A1,TNNC1 |
| Valine, leucine and isoleucine degradation | 0.00000483 | 5.32 | DBT,HSD17B10,HADHA,HIBADH,PCCB,HADHB |
| Propanoate metabolism | 0.000453 | 3.34 | DBT,SUCLA2,HADHA,PCCB |
| Carbon metabolism | 4.89E-08 | 7.31 | DLAT,IDH3G,ALDOC,PDHB,SUCLA2,HADHA,PDHA1,PCCB,GAPDH |
