## Supplemental Table 14 for "Dietary restriction transforms the protein sulfhydrome in a tissue-specific and cystathionine γ-lyase-dependent manner"

| Supplemental Table 14: Sulfhydrated Protein Pathway Enrichment in Serum | | | |
|---|---|---|---|
| KEGG Pathway (Unchanged) | p-val (adj) | -LOG10(p-val adj) | Involved Gene |
| Staphylococcus aureus infection | 3.59E-14 | 13.44 | MBL2,CFH,C5,MASP2,FGG,C1QB,MBL1,C1RA,CFI,PLG,CFD,C4B,CFB |
| Prion diseases | 0.0000268 | 4.57 | C8G,C9,C5,C8B,C8A,C1QB |
| Pertussis | 0.00328 | 2.48 | SERPING1,C4BPA,C5,C1QB,C1RA,C4B |
| Amoebiasis | 0.00251 | 2.60 | COL1A1,C8G,C9,FN1,C8B,COL1A2,C8A |
| Systemic lupus erythematosus | 0.00198 | 2.70 | C8G,C9,C5,C8B,C8A,C1QB,C1RA,C4B |
| Complement and coagulation cascades | 2.85E-53 | 52.55 | C8G,VTN,F12,C9,SERPIND1,KNG1,SERPING1,PROC,MBL2,CFH,F13B,C4BPA,SERPINC1,C5,F2,FGA,MASP2,C8B,F10,FGB,FGG,C8A,C1QB,MBL1,SERPINF2,F13A1,C1RA,CFI,PLG,CFD,SERPINA1A,SERPINA1D,SERPINA1E,C4B,CFB,KLKB1 |
