## Supplemental Table 15 for "Dietary restriction transforms the protein sulfhydrome in a tissue-specific and cystathionine γ-lyase-dependent manner"

| Supplemental Table 15: Accession Numbers for the 28 Common Sulfhydrated Proteins in all 6 tested WT Tissues |
|---|
| P01942 |
| P23953 |
| E9Q557 |
| P02088 |
| Q61838 |
| P29699 |
| Q9QWK4 |
| P01592 |
| Q61171 |
| P07724 |
| P35700 |
| P00920 |
| P07356 |
| P05064 |
| Q921I1 |
| P62897 |
| Q8K0E8 |
| P01837 |
| P01872 |
| P21614 |
| Q8VCG4 |
| E9PV24 |
| P01027 |
| P17751 |
| Q91X72 |
| P07759 |
| Q8VCM7 |
| P06330 |
