## Supplemental Table 16 for "Dietary restriction transforms the protein sulfhydrome in a tissue-specific and cystathionine γ-lyase-dependent manner"

| Supplemental Table 16: Dietary Impact on the CGL KO Liver Sulfhydrome | | | | | | |
|---|---|---|---|---|---|---|
| Protein Name | Accession Number | Alternate ID | Molecular Weight | Cysteine Residues | DR/AL Spectral Count Ratio | P-value |
| Glucosidase 2 subunit beta | O08795 (+1) | Prkcsh | 59 kDa | 17 C | 23.333 | 0.00258 |
| ADP/ATP translocase 2 | P51881 | Slc25a5 | 33 kDa | 4 C | 20.000 | 4.26931 |
| Calponin-3 | Q9DAW9 | Cnn3 | 36 kDa | 3 C | 16.667 | 0.00930 |
| Adenylyl cyclase-associated protein 1 | P40124 | Cap1 | 52 kDa | 6 C | 16.667 | 0.00930 |
| Prolow-density lipoprotein receptor-related protein 1 | Q91ZX7 | Lrp1 | 505 kDa | 332 C | 13.333 | 0.02083 |
| Microtubule-associated protein RP/EB family member 1 | Q61166 | Mapre1 | 30 kDa | 3 C | 13.333 | 0.02083 |
| Tetratricopeptide repeat protein 36 | Q8VBW8 | Ttc36 | 20 kDa | 1 C | 13.333 | 0.02083 |
| Tubulin alpha-1C chain | P68373 | Tuba1c | 50 kDa | 12 C | 13.333 | 0.02083 |
| Fumarate hydratase, mitochondrial | P97807 | Fh | 54 kDa | 4 C | 5.833 | 0.01253 |
| Protein canopy homolog 2 | Q9QXT0 | Cnpy2 | 21 kDa | 6 C | 5.833 | 0.01253 |
| DCN1-like protein 1 | Q9QZ73 | Dcun1d1 | 30 kDa | 4 C | 5.000 | 0.00595 |
| Phytanoyl-CoA dioxygenase, peroxisomal | O35386 | Phyh | 39 kDa | 8 C | 4.167 | 0.04760 |
| cAMP-dependent protein kinase type I-alpha regulatory subunit | Q9DBC7 | Prkar1a | 43 kDa | 5 C | 4.167 | 0.04760 |
| Retinol-binding protein 4 | Q00724 | Rbp4 | 23 kDa | 6 C | 4.167 | 0.04760 |
| Exosome complex component MTR3 | Q8BTW3 | Exosc6 | 28 kDa | 6 C | 3.333 | 0.02192 |
| Short-chain specific acyl-CoA dehydrogenase, mitochondrial | Q07417 | Acads | 45 kDa | 5 C | 3.333 | 0.02192 |
| Hydroxyacid-oxoacid transhydrogenase, mitochondrial | Q8R0N6 | Adhfe1 | 50 kDa | 8 C | 3.226 | 0.02312 |
| Enoyl-CoA hydratase domain-containing protein 3, mitochondrial | Q9D7J9 | Echdc3 | 32 kDa | 5 C | 3.137 | 0.02824 |
| Isoform 2 of CAP-Gly domain-containing linker protein 1 | Q922J3-2 | Clip1 | 147 kDa | 11 C | 2.903 | 0.02308 |
| Cytochrome c oxidase subunit 5A, mitochondrial | P12787 | Cox5a | 16 kDa | 4 C | 2.857 | 0.01231 |
| UPF0598 protein C8orf82 homolog | Q8VE95 | | 24 kDa | 6 C | 2.857 | 0.01231 |
| Acyl-coenzyme A thioesterase 4 | Q8BWN8 | Acot4 | 46 kDa | 6 C | 2.833 | 0.00532 |
| DnaJ homolog subfamily B member 11 | Q99KV1 | Dnajb11 | 41 kDa | 5 C | 2.625 | 0.00796 |
| 4-hydroxy-2-oxoglutarate aldolase, mitochondrial | Q9DCU9 | Hoga1 | 35 kDa | 6 C | 2.300 | 0.02548 |
| Thioredoxin | P10639 | Txn | 12 kDa | 6 C | 2.231 | 0.00849 |
| Fatty aldehyde dehydrogenase | P47740 | Aldh3a2 | 54 kDa | 8 C | 27.000 | 0.20915 |
| Very long-chain specific acyl-CoA dehydrogenase, mitochondrial | P50544 | Acadvl | 71 kDa | 7 C | 26.667 | 0.09957 |
| 40S ribosomal protein S17 | P63276 | Rps17 | 16 kDa | 1 C | 17.000 | 0.24627 |
| Electron transfer flavoprotein-ubiquinone oxidoreductase, mitochondrial | Q921G7 | Etfdh | 68 kDa | 15 C | 16.667 | 0.07851 |
| Selenoprotein P | P70274 | Selenop | 43 kDa | 18 C | 13.667 | 0.21347 |
| Alcohol dehydrogenase [NADP(+)] | Q9JII6 | Akr1a1 | 37 kDa | 4 C | 10.333 | 0.1641 |
| N(G),N(G)-dimethylarginine dimethylaminohydrolase 1 | Q9CWS0 | Ddah1 | 31 kDa | 7 C | 10.333 | 0.1641 |
| Eukaryotic translation initiation factor 5 | P59325 | Eif5 | 49 kDa | 8 C | 10.333 | 0.1641 |
| Calnexin | P35564 | Canx | 67 kDa | 7 C | 10.333 | 0.1641 |
| 60S ribosomal protein L7a | P12970 | Rpl7a | 30 kDa | 3 C | 10.333 | 0.1641 |
| Cytochrome b5 type B | Q9CQX2 | Cyb5b | 16 kDa | 1 C | 10.333 | 0.1641 |
| Membrane-associated progesterone receptor component 1 | O55022 | Pgrmc1 | 22 kDa | 2 C | 10.000 | 0.10994 |
| Radixin | P26043 | Rdx | 69 kDa | 1 C | 8.143 | 0.18923 |
| Putative hydroxypyruvate isomerase | Q8R1F5 | Hyi | 30 kDa | 2 c | 7.333 | 0.37390 |
| Apolipoprotein E | P08226 | Apoe | 36 kDa | 1 C | 7.333 | 0.37390 |
| Acyl-CoA dehydrogenase family member 10 | Q8K370 | Acad10 | 119 kDa | 16 C | 7.333 | 0.37390 |
| 60S ribosomal protein L5 | P47962 | Rpl5 | 34 kDa | 4 C | 7.333 | 0.37390 |
| Cathepsin Z | Q9WUU7 | Ctsz | 34 kDa | 12 C | 7.333 | 0.37390 |
| Polyadenylate-binding protein 2 | Q8CCS6 | Pabpn1 | 32 kDa | 2 C | 7.333 | 0.37390 |
| Leukocyte elastase inhibitor A | Q9D154 | Serpinb1a | 43 kDa | 3 C | 7.333 | 0.37390 |
| Acylpyruvase FAHD1, mitochondrial | Q8R0F8 | Fahd1 | 25 kDa | 6 C | 7.333 | 0.37390 |
| G-rich sequence factor 1 | Q8C5Q4 | Grsf1 | 53 kDa | 9 C | 7.333 | 0.37390 |
| ES1 protein homolog, mitochondrial | Q9D172 | D10Jhu81e | 28 kDa | 6 C | 5.833 | 0.10657 |
| Exosome complex exonuclease RRP42 | Q9D0M0 | Exosc7 | 32 kDa | 12 C | 4.333 | 0.46703 |
| N-fatty-acyl-amino acid synthase/hydrolase PM20D1 | Q8C165 | Pm20d1 | 56 kDa | 2 C | 4.250 | 0.22289 |
| Ig gamma-1 chain C region secreted form | P01868 | Ighg1 | 36 kDa | 12 C | 4.167 | 0.1581 |
| Tumor protein D54 | Q9CYZ2 | Tpd52l2 | 24 kDa | 1 C | 4.167 | 0.1581 |
| Annexin A6 | P14824 | Anxa6 | 76 kDa | 8 C | 4.091 | 0.24234 |
| Glucose-6-phosphate isomerase | P06745 | Gpi | 63 kDa | 4 C | 3.857 | 0.32033 |
| Small glutamine-rich tetratricopeptide repeat-containing protein alpha | Q8BJU0 (+1) | Sgta | 34 kDa | 4 C | 3.500 | 0.49521 |
| GMP reductase 1 | Q9DCZ1 | Gmpr | 37 kDa | 9 C | 3.417 | 0.34704 |
| 60S ribosomal protein L14 | Q9CR57 | Rpl14 | 24 kDa | 2 C | 3.417 | 0.23987 |
| Mitotic checkpoint protein BUB3 | Q9WVA3 | Bub3 | 37 kDa | 7 C | 3.333 | 0.10587 |
| Acyl-CoA-binding domain-containing protein 5 | Q5XG73 (+1) | Acbd5 | 57 kDa | 9 C | 3.333 | 0.10587 |
| ATP synthase subunit alpha, mitochondrial | Q03265 | Atp5a1 | 60 kDa | 2 C | 3.279 | 0.31177 |
| 2,4-dienoyl-CoA reductase, mitochondrial | Q9CQ62 | Decr1 | 36 kDa | 5 C | 3.258 | 0.32354 |
| Voltage-dependent anion-selective channel protein 1 | Q60932 (+1) | Vdac1 | 32 kDa | 2 C | 3.227 | 0.44751 |
| Medium-chain specific acyl-CoA dehydrogenase, mitochondrial | P45952 | Acadm | 46 kDa | 8 C | 3.226 | 0.09118 |
| Kynureninase | Q9CXF0 | Kynu | 52 kDa | 8 C | 3.182 | 0.08905 |
| 1,2-dihydroxy-3-keto-5-methylthiopentene dioxygenase | Q99JT9 | Adi1 | 22 kDa | 1 C | 3.182 | 0.15683 |
| Translationally-controlled tumor protein | P63028 | Tpt1 | 19 kDa | 2 C | 3.171 | 0.05231 |
| Long-chain specific acyl-CoA dehydrogenase, mitochondrial | P51174 | Acadl | 48 kDa | 7 C | 3.099 | 0.07125 |
| Leucine-rich repeat-containing protein 59 | Q922Q8 | Lrrc59 | 35 kDa | 8 C | 2.857 | 0.11637 |
| PITH domain-containing protein 1 | Q8BWR2 | Pithd1 | 24 kDa | 4 C | 2.857 | 0.11637 |
| Trifunctional enzyme subunit alpha, mitochondrial | Q8BMS1 | Hadha | 83 kDa | 12 C | 2.813 | 0.16110 |
| NIF3-like protein 1 | Q9EQ80 | Nif3l1 | 42 kDa | 8 C | 2.800 | 0.05333 |
| Phenazine biosynthesis-like domain-containing protein 2 | Q9CXN7 | Pbld2 | 32 kDa | 4 C | 2.745 | 0.20434 |
| Glutathione peroxidase 3 | P46412 | Gpx3 | 25 kDa | 3 C | 2.727 | 0.21347 |
| Mitochondrial intermembrane space import and assembly protein 40 | Q8VEA4 | Chchd4 | 16 kDa | 7 C | 2.667 | 0.06676 |

| | | | | | | |
|---|---|---|---|---|---|---|
| Cytochrome c oxidase assembly factor 7 | Q921H9 | Coa7 | 26 kDa | 13 C | 2.583 | 0.36848 |
| Elongin-B | P62869 | Elob | 13 kDa | 1 C | 2.583 | 0.36848 |
| Cathepsin B | P10605 | Ctsb | 37 kDa | 16 C | 2.583 | 0.36848 |
| Na(+)/H(+) exchange regulatory cofactor NHE-RF2 | Q9JHL1 | Slc9a3r2 | 37 kDa | 6 C | 2.583 | 0.36848 |
| Isoform 3 of Afamin | O89020-3 | Afm | 70 kDa | 34 C | 2.583 | 0.36848 |
| Rho GDP-dissociation inhibitor 1 | Q99PT1 | Arhgdia | 23 kDa | 1 C | 2.581 | 0.28399 |
| Glutathione S-transferase A1 | P13745 | Gsta1 | 26 kDa | 2 C | 2.432 | 0.05580 |
| Nucleolin | P09405 | Ncl | 77 kDa | 1 C | 2.429 | 0.32967 |
| BolA-like protein 1 | Q9D8S9 | Bola1 | 14 kDa | 3 C | 2.429 | 0.32967 |
| Hemoglobin subunit beta-1 | P02088 | Hbb-b1 | 16 kDa | 2 C | 2.407 | 0.06945 |
| Stromal cell-derived factor 2-like protein 1 | Q9ESP1 | Sdf2l1 | 24 kDa | 4 C | 2.400 | 0.05723 |
| Protein-glucosylgalactosylhydroxylysine glucosidase | Q8BP56 | Pgghg | 76 kDa | 8 C | 2.381 | 0.09737 |
| Cytochrome b-c1 complex subunit 2, mitochondrial | Q9DB77 | Uqcrc2 | 48 kDa | 1 C | 2.381 | 0.09737 |
| OTU domain-containing protein 6B | Q8K2H2 | Otud6b | 34 kDa | 4 C | 2.353 | 0.05385 |
| Isoform 2 of Gelsolin | P13020-2 | Gsn | 81 kDa | 7 C | 2.353 | 0.08887 |
| Phosphatidylinositol transfer protein beta isoform | P53811 | Pitpnb | 31 kDa | 5 C | 2.333 | 0.11611 |
| Thioredoxin reductase 2, mitochondrial | Q9JLT4 (+1) | Txnrd2 | 57 kDa | 11 C | 2.286 | 0.09444 |
| Methylosome protein 50 | Q99J09 | Wdr77 | 37 kDa | 12 C | 2.273 | 0.36750 |
| Myosin light polypeptide 6 | Q60605 (+1) | Myl6 | 17 kDa | 3 C | 2.250 | 0.06676 |
| Phenazine biosynthesis-like domain-containing protein 1 | Q9DCG6 | Pbld1 | 32 kDa | 3 C | 2.222 | 0.15697 |
| Far upstream element-binding protein 1 | Q91WJ8 | Fubp1 | 69 kDa | 3 C | 2.195 | 0.12935 |
| Sarcosine dehydrogenase, mitochondrial | Q99LB7 | Sardh | 102 kDa | 18 C | 2.195 | 0.28267 |
| Actin-related protein 2/3 complex subunit 2 | Q9CVB6 | Arpc2 | 34 kDa | 2 C | 2.195 | 0.3194 |
| Glycerol-3-phosphate dehydrogenase [NAD(+)], cytoplasmic | P13707 | Gpd1 | 38 kDa | 11 C | 2.195 | 0.40385 |
| Alpha-actinin-4 | P57780 | Actn4 | 105 kDa | 8 C | 2.157 | 0.09765 |
| Purine nucleoside phosphorylase | P23492 | Pnp | 32 kDa | 5 C | 2.125 | 0.07392 |
| Proteasome subunit alpha type-3 | O70435 | Psma3 | 28 kDa | 4 C | 2.100 | 0.05137 |
| Acyl-coenzyme A synthetase ACSM1, mitochondrial | Q91VA0 | Acsm1 | 65 kDa | 14 C | 2.000 | 0.06858 |
| Interferon-activable protein 204 | P0DOV2 | Ifi204 | 69 kDa | 13 C | 2.000 | 0.23019 |
| Serine/threonine-protein phosphatase 6 catalytic subunit | Q9CQR6 | Ppp6c | 35 kDa | 12 C | 1.967 | 0.19509 |
| Isoform 3 of 2-oxoglutarate dehydrogenase, mitochondrial | Q60597-3 | Ogdh | 118 kDa | 21 C | 1.967 | 0.35857 |
| Coactosin-like protein | Q9CQI6 | Cotl1 | 16 kDa | 2 C | 1.952 | 0.50353 |
| Isoform Peroxisomal of Serine--pyruvate aminotransferase, mitochondrial | O35423-2 | Agxt | 44 kDa | 7 C | 1.951 | 0.29728 |
| Protein NipSnap homolog 3B | Q9CQE1 | Nipsnap3b | 28 kDa | 2 C | 1.947 | 0.01989 |
| Proteasome activator complex subunit 1 | P97371 | Psme1 | 29 kDa | 3 C | 1.944 | 0.1007 |
| Optineurin | Q8K3K8 | Optn | 67 kDa | 12 C | 1.935 | 0.44446 |
| Proteasome subunit alpha type-7 | Q9Z2U0 | Psma7 | 28 kDa | 3 C | 1.923 | 0.05504 |
| S-phase kinase-associated protein 1 | Q9WTX5 | Skp1 | 19 kDa | 3 C | 1.909 | 0.08347 |
| Glutamate dehydrogenase 1, mitochondrial | P26443 | Glud1 | 61 kDa | 6 C | 1.909 | 0.1614 |
| 60 kDa heat shock protein, mitochondrial | P63038 | Hspd1 | 61 kDa | 3 C | 1.909 | 0.01438 |
| D-dopachrome decarboxylase | O35215 | Ddt | 13 kDa | 2 C | 1.909 | 0.66873 |
| Trans-1,2-dihydrobenzene-1,2-diol dehydrogenase | Q9DBB8 | Dhdh | 36 kDa | 6 C | 1.905 | 0.23073 |
| Galectin-3-binding protein | Q07797 | Lgals3bp | 64 kDa | 16 C | 1.905 | 0.23073 |
| Isoform 2 of Low molecular weight phosphotyrosine protein phosphatase | Q9D358-2 | Acp1 | 18 kDa | 8 C | 1.905 | 0.23073 |
| Dihydrolipoyl dehydrogenase, mitochondrial | O08749 | Dld | 54 kDa | 9 C | 1.881 | 0.32830 |
| Delta(3,5)-Delta(2,4)-dienoyl-CoA isomerase, mitochondrial | O35459 | Ech1 | 36 kDa | 6 C | 1.864 | 0.51851 |
| Synapse-associated protein 1 | Q9D5V6 | Syap1 | 41 kDa | 1 C | 1.864 | 0.58427 |
| Glutathione S-transferase A3 | P30115 | Gsta3 | 25 kDa | 1 C | 1.861 | 0.10861 |
| Phosphoribosyl pyrophosphate synthase-associated protein 1 | Q9D0M1 | Prpsap1 | 39 kDa | 6 C | 1.852 | 0.33234 |
| Shootin-1 | Q8K2Q9 | Shtn1 | 71 kDa | 9 C | 1.833 | 0.65913 |
| Isoform 5 of Ubiquitin-associated protein 2-like | Q80X50-5 | Ubap2l | 117 kDa | 2 C | 1.833 | 0.65913 |
| 5'-3' exoribonuclease 2 | Q9DBR1 (+1) | Xrn2 | 109 kDa | 17 C | 1.833 | 0.65913 |
| Protein farnesyltransferase/geranylgeranyltransferase type-1 subunit alpha | Q61239 | Fnta | 44 kDa | 3 C | 1.833 | 0.65913 |
| BAG family molecular chaperone regulator 5 | Q8CI32 | Bag5 | 51 kDa | 10 C | 1.833 | 0.65913 |
| Ribose-phosphate pyrophosphokinase 1 | Q9D7G0 | Prps1 | 35 kDa | 9 C | 1.833 | 0.3464 |
| Uricase | P25688 | Uox | 35 kDa | 4 C | 1.833 | 0.3464 |
| Glutathione S-transferase omega-1 | O09131 | Gsto1 | 27 kDa | 4 C | 1.833 | 0.65913 |
| Tight junction protein ZO-1 | P39447 | Tjp1 | 195 kDa | 7 C | 1.833 | 0.65913 |
| Prostaglandin reductase 2 | Q8VDQ1 | Ptgr2 | 38 kDa | 8 C | 1.833 | 0.65913 |
| Endoplasmin | P08113 | Hsp90b1 | 92 kDa | 5 C | 1.833 | 0.65913 |
| Hsp90 co-chaperone Cdc37 | Q61081 | Cdc37 | 45 kDa | 9 C | 1.818 | 0.1144 |
| Nicotinate-nucleotide pyrophosphorylase [carboxylating] | Q91X91 | Qprt | 32 kDa | 6 C | 1.818 | 0.44901 |
| Malate dehydrogenase, mitochondrial | P08249 | Mdh2 | 36 kDa | 8 C | 1.813 | 0.37289 |
| Eukaryotic translation initiation factor 4B | Q8BGD9 | Eif4b | 69 kDa | 3 C | 1.800 | 0.18484 |
| COP9 signalosome complex subunit 5 | O35864 | Cops5 | 38 kDa | 4 C | 1.784 | 0.58155 |
| Actin-related protein 2/3 complex subunit 1A | Q9R0Q6 | Arpc1a | 42 kDa | 10 C | 1.778 | 0.33046 |
| Serine--tRNA ligase, cytoplasmic | P26638 | Sars | 58 kDa | 9 C | 1.775 | 0.44335 |
| Protein disulfide-isomerase A6 | Q922R8 | Pdia6 | 48 kDa | 7 C | 1.769 | 0.13688 |
| Isoform 2 of Alpha-aminoadipic semialdehyde dehydrogenase | Q9DBF1-2 | Aldh7a1 | 56 kDa | 9 C | 1.768 | 0.0871 |
| Histidine triad nucleotide-binding protein 1 | P70349 | Hint1 | 14 kDa | 2 C | 1.765 | 0.02264 |
| Glutaredoxin-related protein 5, mitochondrial | Q80Y14 | Glrx5 | 16 kDa | 2 C | 1.750 | 0.05504 |
| Eukaryotic translation initiation factor 6 | O55135 | Eif6 | 27 kDa | 8 C | 1.750 | 0.10119 |
| Probable D-lactate dehydrogenase, mitochondrial | Q7TNG8 | Ldhd | 52 kDa | 12 C | 1.750 | 0.50715 |
| Ribonuclease inhibitor | Q91VI7 | Rnh1 | 50 kDa | 30 C | 1.739 | 0.20301 |
| Fructose-bisphosphate aldolase A | P05064 | Aldoa | 39 kDa | 8 C | 1.737 | 0.01779 |
| Enoyl-CoA hydratase domain-containing protein 2, mitochondrial | Q3TLP5 | Echdc2 | 32 kDa | 6 C | 1.727 | 0.14814 |
| 3-ketoacyl-CoA thiolase, mitochondrial | Q8BWT1 | Acaa2 | 42 kDa | 8 C | 1.714 | 0.29000 |
| ATP synthase subunit d, mitochondrial | Q9DCX2 | Atp5h | 19 kDa | 1 C | 1.707 | 0.35245 |

| | | | | | | |
|---|---|---|---|---|---|---|
| Alpha-2-HS-glycoprotein | P29699 | Ahsg | 37 kDa | 14 C | 1.707 | 0.35245 |
| Ataxin-2 | O70305 | Atxn2 | 136 kDa | 15 C | 1.707 | 0.42361 |
| Carboxymethylenebutenolidase homolog | Q8R1G2 | Cmbl | 28 kDa | 6 C | 1.700 | 0.23019 |
| Proteasome subunit beta type-1 | O09061 | Psmb1 | 26 kDa | 5 C | 1.696 | 0.02699 |
| Septin-11 | Q8C1B7 (+1) | 11-Sep | 50 kDa | 6 C | 1.692 | 0.186 |
| Phosphoglycerate kinase 1 | P09411 | Pgk1 | 45 kDa | 7 C | 1.684 | 0.19004 |
| Clathrin light chain A | O08585 | Clta | 26 kDa | 1 C | 1.667 | 0.11611 |
| Cytochrome b-c1 complex subunit 1, mitochondrial | Q9CZ13 | Uqcrc1 | 53 kDa | 11 C | 1.667 | 0.11611 |
| Ficolin-1 | O70165 | Fcn1 | 36 kDa | 10 C | 1.667 | 0.11611 |
| RNA-binding protein EWS | Q61545 | Ewsr1 | 68 kDa | 5 C | 1.667 | 0.11611 |
| Eukaryotic translation initiation factor 1 | P48024 | Eif1 | 13 kDa | 2 C | 1.667 | 0.27457 |
| Ig alpha chain C region | P01878 | | 37 kDa | 13 C | 1.667 | 0.28786 |
| Phosphoglucomutase-1 | Q9D0F9 | Pgm1 | 61 kDa | 10 C | 1.667 | 0.00377 |
| Ferritin light chain 1 | P29391 | Ftl1 | 21 kDa | 1 C | 1.667 | 0.01944 |
| Proteasome subunit beta type-2 | Q9R1P3 | Psmb2 | 23 kDa | 3 C | 1.636 | 0.09126 |
| Quinone oxidoreductase | P47199 | Cryz | 35 kDa | 5 C | 1.634 | 0.01350 |
| Calreticulin | P14211 | Calr | 48 kDa | 6 C | 1.630 | 0.00578 |
| NADH dehydrogenase [ubiquinone] flavoprotein 2, mitochondrial | Q9D6J6 | Ndufv2 | 27 kDa | 6 C | 1.625 | 0.15183 |
| Maleylacetoacetate isomerase | Q9WVL0 | Gstz1 | 24 kDa | 3 C | 1.613 | 0.37976 |
| Peroxiredoxin-2 | Q61171 | Prdx2 | 22 kDa | 3 C | 1.612 | 0.00131 |
| Hydroxyacyl-coenzyme A dehydrogenase, mitochondrial | Q61425 | Hadh | 34 kDa | 5 C | 1.609 | 0.13470 |
| Carbonic anhydrase 1 | P13634 | Ca1 | 28 kDa | 1 C | 1.600 | 0.10119 |
| Elongation factor 1-beta | O70251 | Eef1b | 25 kDa | 3 C | 1.600 | 0.21930 |
| 3'(2'),5'-bisphosphate nucleotidase 1 | Q9Z0S1 | Bpnt1 | 33 kDa | 6 C | 1.600 | 0.2372 |
| Immunoglobulin J chain | P01592 | Jchain | 18 kDa | 8 C | 1.600 | 0.2508 |
| Eukaryotic translation initiation factor 4H | Q9WUK2 | Eif4h | 27 kDa | 1 C | 1.600 | 0.2508 |
| Proteasome subunit beta type-8 | P28063 | Psmb8 | 30 kDa | 5 C | 1.600 | 0.34864 |
| Hemoglobin subunit alpha | P01942 | Hba | 15 kDa | 1 C | 1.600 | 0.45746 |
| Persulfide dioxygenase ETHE1, mitochondrial | Q9DCM0 | Ethe1 | 28 kDa | 9 C | 1.600 | 0.013 |
| Regucalcin | Q64374 | Rgn | 33 kDa | 9 C | 1.571 | 0.38539 |
| Ig heavy chain V region X44 | P01807 (+1) | | 13 kDa | 2 C | 1.569 | 0.51483 |
| Actin, cytoplasmic 1 | P60710 | Actb | 42 kDa | 6 C | 1.569 | 0.1108 |
| Vitamin D-binding protein | P21614 | Gc | 54 kDa | 28 C | 1.566 | 0.00131 |
| Tetratricopeptide repeat protein 38 | A3KMP2 | Ttc38 | 52 kDa | 9 C | 1.563 | 0.1144 |
| Triokinase/FMN cyclase | Q8VC30 | Tkfc | 60 kDa | 5 C | 1.563 | 0.63628 |
| Nascent polypeptide-associated complex subunit alpha, muscle-specific form | P70670 | Naca | 220 kDa | 16 C | 1.556 | 0.06676 |
| Malate dehydrogenase, cytoplasmic | P14152 | Mdh1 | 37 kDa | 3 C | 1.545 | 0.10119 |
| Sorting nexin-12 | O70493 | Snx12 | 19 kDa | 3 C | 1.524 | 0.73548 |
| Agmatinase, mitochondrial | A2AS89 | Agmat | 38 kDa | 9 C | 1.520 | 0.11217 |
| Eukaryotic translation initiation factor 5A-1 | P63242 | Eif5a | 17 kDa | 4 C | 1.516 | 0.01610 |
| Serum albumin | P07724 | Alb | 69 kDa | 36 C | 1.508 | 0.03033 |
| ELKS/Rab6-interacting/CAST family member 1 | Q99MI1 | Erc1 | 128 kDa | 4 C | 1.500 | 0.37390 |
| Omega-amidase NIT2 | Q9JHW2 | Nit2 | 31 kDa | 2 C | 1.500 | 0.38665 |
| Secernin-2 | Q8VCA8 | Scrn2 | 47 kDa | 10 C | 1.500 | 0.39820 |
| 2-aminoethanethiol dioxygenase | Q6PDY2 | Ado | 28 kDa | 7 C | 1.500 | 0.56143 |
| Fructose-1,6-bisphosphatase 1 | Q9QXD6 | Fbp1 | 37 kDa | 7 C | 1.489 | 0.05022 |
| Actin-related protein 2/3 complex subunit 3 | Q9JM76 | Arpc3 | 21 kDa | 4 C | 1.476 | 0.62226 |
| Pyridoxine-5'-phosphate oxidase | Q91XF0 | Pnpo | 30 kDa | 6 C | 1.476 | 0.62226 |
| Alpha-soluble NSF attachment protein | Q9DB05 | Napa | 33 kDa | 8 C | 1.476 | 0.62226 |
| Glucokinase regulatory protein | Q91X44 | Gckr | 65 kDa | 8 C | 1.476 | 0.62226 |
| Malignant T-cell-amplified sequence 1 | Q9DB27 | Mcts1 | 21 kDa | 4 C | 1.467 | 0.05723 |
| GMP reductase 2 | Q99L27 | Gmpr2 | 38 kDa | 9 C | 1.463 | 0.50091 |
| Regulator of microtubule dynamics protein 3 | Q3UJU9 | Rmdn3 | 52 kDa | 6 C | 1.463 | 0.50091 |
| Caspase-6 | O08738 | Casp6 | 32 kDa | 10 C | 1.463 | 0.57307 |
| Protein phosphatase 1 regulatory subunit 7 | Q3UM45 | Ppp1r7 | 41 kDa | 2 C | 1.462 | 0.00653 |
| 5-oxoprolinase | Q8K010 | Oplah | 138 kDa | 25 C | 1.457 | 0.77043 |
| Elongation factor 1-delta | P57776 | Eef1d | 31 kDa | 2 C | 1.444 | 0.20510 |
| Neutral alpha-glucosidase AB | Q8BHN3 | Ganab | 107 kDa | 8 C | 1.440 | 0.30602 |
| NADH dehydrogenase [ubiquinone] flavoprotein 1, mitochondrial | Q91YT0 | Ndufv1 | 51 kDa | 12 C | 1.429 | 0.19224 |
| Transthyretin | P07309 | Ttr | 16 kDa | 2 C | 1.429 | 0.40178 |
| Aldehyde dehydrogenase family 8 member A1 | Q8BH00 | Aldh8a1 | 54 kDa | 13 C | 1.429 | 0.69484 |
| Proteasome subunit alpha type-2 | P49722 | Psma2 | 26 kDa | 2 C | 1.429 | 0.03994 |
| NADH-ubiquinone oxidoreductase 75 kDa subunit, mitochondrial | Q91VD9 | Ndufs1 | 80 kDa | 18 C | 1.421 | 0.27457 |
| Carboxylesterase 3A | Q63880 | Ces3a | 63 kDa | 7 C | 1.417 | 0.52623 |
| Proteasome subunit beta type-5 | O55234 | Psmb5 | 29 kDa | 3 C | 1.409 | 0.1144 |
| Protein arginine N-methyltransferase 5 | Q8CIG8 | Prmt5 | 73 kDa | 12 C | 1.409 | 0.73843 |
| Alanine--tRNA ligase, cytoplasmic | Q8BGQ7 | Aars | 107 kDa | 15 C | 1.409 | 0.73843 |
| High mobility group protein B1 | P63158 | Hmgb1 | 25 kDa | 3 C | 1.407 | 0.22213 |
| Proteasome subunit beta type-6 | Q60692 | Psmb6 | 25 kDa | 4 C | 1.400 | 0.14814 |
| Hydroxyacylglutathione hydrolase, mitochondrial | Q99KB8 | Hagh | 34 kDa | 8 C | 1.400 | 0.41350 |
| Dynein light chain 2, cytoplasmic | Q9D0M5 | Dynll2 | 10 kDa | 2 C | 1.400 | 0.42164 |
| Microsomal triglyceride transfer protein large subunit | O08601 | Mttp | 99 kDa | 11 C | 1.400 | 0.47662 |
| SH3 domain-binding glutamic acid-rich-like protein | Q9JJU8 | Sh3bgrl | 13 kDa | 2 C | 1.400 | 0.51851 |
| Ubiquitin carboxyl-terminal hydrolase 5 | P56399 | Usp5 | 96 kDa | 16 C | 1.386 | 0.7080 |
| Cordon-bleu protein-like 1 | Q3UMF0 | Cobll1 | 137 kDa | 10 C | 1.375 | 0.65195 |
| Argininosuccinate lyase | Q91YI0 | Asl | 52 kDa | 13 C | 1.373 | 0.63247 |
| NEDD8-conjugating enzyme Ubc12 | P61082 | Ube2m | 21 kDa | 5 C | 1.373 | 0.63247 |
| Dipeptidyl peptidase 1 | P97821 | Ctsc | 52 kDa | 14 C | 1.373 | 0.63247 |

| | | | | | | |
|---|---|---|---|---|---|---|
| Fatty acid-binding protein, liver | P12710 | Fabp1 | 14 kDa | 1 C | 1.370 | 0.02471 |
| Glutathione synthetase | P51855 | Gss | 52 kDa | 4 C | 1.368 | 0.31956 |
| Ferritin heavy chain | P09528 | Fth1 | 21 kDa | 3 C | 1.367 | 0.69078 |
| GrpE protein homolog 1, mitochondrial | Q99LP6 | Grpel1 | 24 kDa | 4 C | 1.364 | 0.69535 |
| Proteasome subunit alpha type-1 | Q9R1P4 | Psma1 | 30 kDa | 5 C | 1.360 | 0.13380 |
| Ran-specific GTPase-activating protein | P34022 | Ranbp1 | 24 kDa | 3 C | 1.357 | 0.15183 |
| Isoform Short of Heterogeneous nuclear ribonucleoprotein A1 | P49312-2 | Hnrnpa1 | 29 kDa | 2 C | 1.353 | 0.14470 |
| 2-iminobutanoate/2-iminopropanoate deaminase | P52760 | Rida | 14 kDa | 1 C | 1.353 | 0.14470 |
| Dihydropteridine reductase | Q8BVI4 | Qdpr | 26 kDa | 4 C | 1.353 | 0.21930 |
| Carboxylesterase 1D | Q8VCT4 | Ces1d | 62 kDa | 5 C | 1.348 | 0.03004 |
| Peptidyl-prolyl cis-trans isomerase D | Q9CR16 | Ppid | 41 kDa | 7 C | 1.333 | 0.05723 |
| Proteasome subunit alpha type-4 | Q9R1P0 | Psma4 | 29 kDa | 5 C | 1.333 | 0.05723 |
| Kininogen-1 | O08677 | Kng1 | 73 kDa | 19 C | 1.333 | 0.27457 |
| Acyl-coenzyme A thioesterase THEM4 | Q3UUI3 | Them4 | 26 kDa | 4 C | 1.333 | 0.28786 |
| Plastin-2 | Q61233 | Lcp1 | 70 kDa | 11 C | 1.333 | 0.29618 |
| Transcription elongation factor A protein 1 | P10711 | Tcea1 | 34 kDa | 8 C | 1.333 | 0.44182 |
| Major urinary protein 1 | P11588 | Mup1 | 21 kDa | 5 C | 1.333 | 0.6408 |
| Selenide, water dikinase 2 | P97364 | Sephs2 | 48 kDa | 7 C | 1.329 | 0.08003 |
| Succinate dehydrogenase [ubiquinone] iron-sulfur subunit, mitochondrial | Q9CQA3 | Sdhb | 32 kDa | 14 C | 1.324 | 0.39783 |
| Carbamoyl-phosphate synthase [ammonia], mitochondrial | Q8C196 | Cps1 | 165 kDa | 21 C | 1.319 | 0.52373 |
| Serine/threonine-protein phosphatase PP1-beta catalytic subunit | P62141 | Ppp1cb | 37 kDa | 14 C | 1.318 | 0.21817 |
| Isoamyl acetate-hydrolyzing esterase 1 homolog | Q9DB29 | Iah1 | 28 kDa | 8 C | 1.316 | 0.4168 |
| Macrophage-capping protein | P24452 | Capg | 39 kDa | 5 C | 1.311 | 0.65368 |
| Proteasome subunit beta type-7 | P70195 | Psmb7 | 30 kDa | 6 C | 1.308 | 0.34527 |
| Heme-binding protein 1 | Q9R257 | Hebp1 | 21 kDa | 2 C | 1.308 | 0.67786 |
| Copper chaperone for superoxide dismutase | Q9WU84 | Ccs | 29 kDa | 10 C | 1.304 | 0.56370 |
| Prostaglandin E synthase 3 | Q9R0Q7 | Ptges3 | 19 kDa | 5 C | 1.300 | 0.10119 |
| 14-3-3 protein gamma | P61982 | Ywhag | 28 kDa | 3 C | 1.294 | 0.35601 |
| Alpha-enolase | P17182 | Eno1 | 47 kDa | 6 C | 1.293 | 0.05764 |
| Phosphotriesterase-related protein | Q60866 | Pter | 39 kDa | 6 C | 1.293 | 0.44393 |
| Proliferation-associated protein 2G4 | P50580 | Pa2g4 | 44 kDa | 6 C | 1.290 | 0.6646 |
| Methylglutaconyl-CoA hydratase, mitochondrial | Q9JLZ3 | Auh | 33 kDa | 5 C | 1.290 | 0.6646 |
| Liver carboxylesterase 1 | Q8VCC2 | Ces1 | 63 kDa | 7 C | 1.286 | 0.06467 |
| Sorbitol dehydrogenase | Q64442 | Sord | 38 kDa | 10 C | 1.286 | 0.18874 |
| Isoform 2 of Heterogeneous nuclear ribonucleoprotein A3 | Q8BG05-2 | Hnrnpa3 | 37 kDa | 4 C | 1.286 | 0.36938 |
| Protein ABHD14B | Q8VCR7 | Abhd14b | 22 kDa | 2 C | 1.286 | 0.37390 |
| Acidic leucine-rich nuclear phosphoprotein 32 family member A | O35381 | Anp32a | 29 kDa | 3 C | 1.286 | 0.56143 |
| Transitional endoplasmic reticulum ATPase | Q01853 | Vcp | 89 kDa | 12 C | 1.282 | 0.25997 |
| Glutaredoxin-3 | Q9CQM9 | Glrx3 | 38 kDa | 5 C | 1.281 | 0.01579 |
| Major urinary protein 20 | Q5FW60 | Mup20 | 21 kDa | 4 C | 1.281 | 0.827 |
| Isoform 2 of Nitrilase homolog 1 | Q8VDK1-2 | Nit1 | 32 kDa | 13 C | 1.278 | 0.50189 |
| Thioredoxin domain-containing protein 5 | Q91W90 | Txndc5 | 46 kDa | 12 C | 1.273 | 0.18407 |
| Peroxisomal sarcosine oxidase | Q9D826 | Pipox | 44 kDa | 11 C | 1.273 | 0.5879 |
| Hydroxymethylglutaryl-CoA lyase, mitochondrial | P38060 | Hmgcl | 34 kDa | 8 C | 1.273 | 0.02239 |
| Thioredoxin-dependent peroxide reductase, mitochondrial | P20108 | Prdx3 | 28 kDa | 4 C | 1.269 | 0.1835 |
| Glutathione peroxidase 1 | P11352 | Gpx1 | 22 kDa | 4 C | 1.268 | 0.18600 |
| Endoplasmic reticulum resident protein 44 | Q9D1Q6 | Erp44 | 47 kDa | 7 C | 1.267 | 0.44182 |
| Glyceraldehyde-3-phosphate dehydrogenase | P16858 | Gapdh | 36 kDa | 5 C | 1.256 | 0.05137 |
| Hsc70-interacting protein | Q99L47 | St13 | 42 kDa | 3 C | 1.250 | 0.15183 |
| Nucleoside diphosphate kinase B | Q01768 | Nme2 | 17 kDa | 2 C | 1.250 | 0.26176 |
| Heterogeneous nuclear ribonucleoprotein H | O35737 | Hnrnph1 | 49 kDa | 1 C | 1.250 | 0.28786 |
| Heterogeneous nuclear ribonucleoprotein A3 | Q8BG05 | Hnrnpa3 | 40 kDa | 4 C | 1.250 | 0.38231 |
| 3-hydroxyisobutyrate dehydrogenase, mitochondrial | Q99L13 | Hibadh | 35 kDa | 12 C | 1.250 | 0.47950 |
| Cleavage and polyadenylation specificity factor subunit 5 | Q9CQF3 | Nudt21 | 26 kDa | 1 C | 1.250 | 0.51851 |
| Myotrophin | P62774 | Mtpn | 13 kDa | 3 C | 1.250 | 0.51851 |
| Splicing factor 1 | Q64213 | Sf1 | 70 kDa | 4 C | 1.250 | 0.54842 |
| Growth factor receptor-bound protein 2 | Q60631 | Grb2 | 25 kDa | 2 C | 1.250 | 0.67786 |
| Pterin-4-alpha-carbinolamine dehydratase | P61458 | Pcbd1 | 12 kDa | 1 C | 1.250 | 0.67786 |
| Electron transfer flavoprotein subunit beta | Q9DCW4 | Etfb | 28 kDa | 4 C | 1.243 | 0.24636 |
| Carbonic anhydrase 3 | P16015 | Ca3 | 29 kDa | 5 C | 1.243 | 0.10262 |
| 14-3-3 protein beta/alpha | Q9CQV8 | Ywhab | 28 kDa | 2 C | 1.235 | 0.54671 |
| Carboxylesterase 1F | Q91WU0 | Ces1f | 62 kDa | 7 C | 1.226 | 0.0101 |
| Oxygen-dependent coproporphyrinogen-III oxidase, mitochondrial | P36552 | Cpox | 50 kDa | 10 C | 1.224 | 0.27558 |
| Protein disulfide-isomerase | P09103 | P4hb | 57 kDa | 7 C | 1.221 | 0.04865 |
| Glia maturation factor beta | Q9CQI3 | Gmfb | 17 kDa | 3 C | 1.220 | 0.69662 |
| Eukaryotic translation initiation factor 3 subunit I | Q9QZD9 | Eif3i | 36 kDa | 7 C | 1.220 | 0.69662 |
| Ig kappa chain V-V region K2 (Fragment) | P01635 | | 13 kDa | 3 C | 1.220 | 0.69662 |
| Isoform Short of Adenosine kinase | P55264-2 | Adk | 38 kDa | 6 C | 1.219 | 0.47499 |
| 3-hydroxyanthranilate 3,4-dioxygenase | Q78JT3 | Haao | 33 kDa | 3 C | 1.217 | 0.22762 |
| Heterogeneous nuclear ribonucleoprotein K | P61979 (+1) | Hnrnpk | 51 kDa | 5 C | 1.217 | 0.25240 |
| Vinculin | Q64727 | Vcl | 117 kDa | 10 C | 1.216 | 0.25877 |
| Destrin | Q9R0P5 | Dstn | 19 kDa | 6 C | 1.214 | 0.2508 |
| Proteasome subunit alpha type-5 | Q9Z2U1 | Psma5 | 26 kDa | 3 C | 1.214 | 0.46760 |
| Copper homeostasis protein cutC homolog | Q9D8X1 | Cutc | 29 kDa | 7 C | 1.214 | 0.73441 |
| Serine/threonine-protein phosphatase PP1-alpha catalytic subunit | P62137 | Ppp1ca | 38 kDa | 13 C | 1.208 | 0.37390 |
| Flavin reductase (NADPH) | Q923D2 | Blvrb | 22 kDa | 2 C | 1.205 | 0.09073 |
| Glutathione reductase, mitochondrial | P47791 | Gsr | 54 kDa | 11 C | 1.204 | 0.07892 |
| Platelet-activating factor acetylhydrolase IB subunit beta | Q61206 | Pafah1b2 | 26 kDa | 3 C | 1.200 | 0.37390 |

| | | | | | | |
|---|---|---|---|---|---|---|
| Aldose 1-epimerase | Q8K157 | Galm | 38 kDa | 4 C | 1.200 | 0.37390 |
| Glyoxylate reductase/hydroxypyruvate reductase | Q91Z53 | Grhpr | 35 kDa | 7 C | 1.200 | 0.47499 |
| Ubiquitin-conjugating enzyme E2 N | P61089 | Ube2n | 17 kDa | 1 C | 1.200 | 0.64332 |
| Selenoprotein F | Q9ERR7 | Selenof | 18 kDa | 7 C | 1.200 | 0.64332 |
| Adrenodoxin, mitochondrial | P46656 | Fdx1 | 20 kDa | 8 C | 1.200 | 0.64332 |
| Elongation factor 2 | P58252 | Eef2 | 95 kDa | 7 C | 1.197 | 0.62197 |
| Complement factor I | Q61129 | Cfi | 67 kDa | 40 C | 1.196 | 0.84797 |
| 78 kDa glucose-regulated protein | P20029 | Hspa5 | 72 kDa | 1 C | 1.195 | 0.11552 |
| Glutathione S-transferase Mu 3 | P19639 | Gstm3 | 26 kDa | 4 C | 1.192 | 0.5062 |
| Methylcrotonoyl-CoA carboxylase beta chain, mitochondrial | Q3ULD5 | Mccc2 | 61 kDa | 10 C | 1.192 | 0.04218 |
| Peroxiredoxin-5, mitochondrial | P99029 | Prdx5 | 22 kDa | 6 C | 1.190 | 0.02538 |
| Isochorismatase domain-containing protein 2A | P85094 | Isoc2a | 22 kDa | 6 C | 1.188 | 0.18407 |
| Phosphoglycerate mutase 1 | Q9DBJ1 | Pgam1 | 29 kDa | 2 C | 1.185 | 0.62226 |
| Cytosolic non-specific dipeptidase | Q9D1A2 | Cndp2 | 53 kDa | 8 C | 1.184 | 0.39867 |
| 26S proteasome non-ATPase regulatory subunit 9 | Q9CR00 | Psmd9 | 25 kDa | 3 C | 1.182 | 0.57901 |
| Ornithine carbamoyltransferase, mitochondrial | P11725 | Otc | 40 kDa | 2 C | 1.182 | 0.60395 |
| Caprin-1 | Q60865 | Caprin1 | 78 kDa | 3 C | 1.182 | 0.79133 |
| Galactose-1-phosphate uridylyltransferase | Q03249 | Galt | 43 kDa | 7 C | 1.182 | 0.80248 |
| Lactoylglutathione lyase | Q9CPU0 | Glo1 | 21 kDa | 3 C | 1.179 | 0.52623 |
| Nucleoside diphosphate kinase 3 | Q9WV85 | Nme3 | 19 kDa | 3 C | 1.176 | 0.83044 |
| Plastin-3 | Q99K51 | Pls3 | 71 kDa | 9 C | 1.167 | 0.64332 |
| Glutathione S-transferase P 1 | P19157 | Gstp1 | 24 kDa | 3 C | 1.164 | 0.43060 |
| Succinate dehydrogenase [ubiquinone] flavoprotein subunit, mitochondrial | Q8K2B3 | Sdha | 73 kDa | 19 C | 1.162 | 0.07854 |
| Drebrin-like protein | Q62418 (+2) | Dbnl | 49 kDa | 5 C | 1.161 | 0.00749 |
| Methylmalonyl-CoA mutase, mitochondrial | P16332 | Mut | 83 kDa | 8 C | 1.158 | 0.56553 |
| Aspartate aminotransferase, cytoplasmic | P05201 | Got1 | 46 kDa | 5 C | 1.156 | 0.51851 |
| Cofilin-1 | P18760 | Cfl1 | 19 kDa | 4 C | 1.154 | 0.46852 |
| Succinate-semialdehyde dehydrogenase, mitochondrial | Q8BWF0 | Aldh5a1 | 56 kDa | 10 C | 1.151 | 0.2839 |
| Inositol monophosphatase 1 | O55023 | Impa1 | 30 kDa | 6 C | 1.150 | 0.2508 |
| Selenide, water dikinase 1 | Q8BH69 | Sephs1 | 43 kDa | 9 C | 1.150 | 0.59788 |
| Caspase-3 | P70677 | Casp3 | 31 kDa | 8 C | 1.148 | 0.81095 |
| Adenosylhomocysteinase | P50247 | Ahcy | 48 kDa | 9 C | 1.146 | 0.28198 |
| Thioredoxin-like protein 1 | Q8CDN6 | Txnl1 | 32 kDa | 7 C | 1.143 | 0.29425 |
| Plasminogen activator inhibitor 1 RNA-binding protein | Q9CY58 | Serbp1 | 45 kDa | 2 C | 1.143 | 0.51851 |
| FAD-linked sulfhydryl oxidase ALR | P56213 | Gfer | 23 kDa | 8 C | 1.143 | 0.57901 |
| Polymeric immunoglobulin receptor | O70570 | Pigr | 85 kDa | 23 C | 1.143 | 0.67786 |
| Estradiol 17 beta-dehydrogenase 5 | P70694 | Akr1c6 | 37 kDa | 7 C | 1.143 | 0.74152 |
| Electron transfer flavoprotein subunit alpha, mitochondrial | Q99LC5 | Etfa | 35 kDa | 6 C | 1.143 | 0.74152 |
| Delta-aminolevulinic acid dehydratase | P10518 | Alad | 36 kDa | 8 C | 1.141 | 0.38675 |
| Argininosuccinate synthase | P16460 | Ass1 | 47 kDa | 5 C | 1.140 | 0.85510 |
| Thioredoxin reductase 1, cytoplasmic | Q9JMH6 | Txnrd1 | 67 kDa | 21 C | 1.133 | 0.61301 |
| Proteasome subunit beta type-3 | Q9R1P1 | Psmb3 | 23 kDa | 5 C | 1.133 | 0.64332 |
| Aldehyde oxidase 3 | G3X982 | Aox3 | 147 kDa | 38 C | 1.130 | 0.65894 |
| Phosphatidylethanolamine-binding protein 1 | P70296 | Pebp1 | 21 kDa | 3 C | 1.125 | 0.34864 |
| Serine protease inhibitor A3K | P07759 | Serpina3k | 47 kDa | 4 C | 1.125 | 0.7174 |
| Hepatoma-derived growth factor | P51859 | Hdgf | 26 kDa | 2 C | 1.125 | 0.72465 |
| Glucosamine 6-phosphate N-acetyltransferase | Q9JK38 | Gnpnat1 | 21 kDa | 6 C | 1.125 | 0.72465 |
| Superoxide dismutase [Cu-Zn] | P08228 | Sod1 | 16 kDa | 3 C | 1.125 | 0.81490 |
| Early endosome antigen 1 | Q8BL66 | Eea1 | 161 kDa | 20 C | 1.125 | 0.86012 |
| Ig mu chain C region | P01872 | Ighm | 50 kDa | 19 C | 1.124 | 0.50041 |
| Retinal dehydrogenase 1 | P24549 | Aldh1a1 | 54 kDa | 11 C | 1.121 | 0.44436 |
| Ubiquitin-conjugating enzyme E2 L3 | P68037 | Ube2l3 | 18 kDa | 3 C | 1.115 | 0.62395 |
| S-methylmethionine--homocysteine S-methyltransferase BHMT2 | Q91WS4 | Bhmt2 | 40 kDa | 10 C | 1.115 | 0.66558 |
| UMP-CMP kinase | Q9DBP5 | Cmpk1 | 22 kDa | 6 C | 1.111 | 0.53714 |
| Receptor of activated protein C kinase 1 | P68040 | Rack1 | 35 kDa | 8 C | 1.111 | 0.67017 |
| Eukaryotic translation initiation factor 1A, X-chromosomal | Q8BMJ3 | Eif1ax | 16 kDa | 2 C | 1.111 | 0.72465 |
| GTP cyclohydrolase 1 | Q05915 | Gch1 | 27 kDa | 3 C | 1.111 | 0.84159 |
| Heterogeneous nuclear ribonucleoprotein A/B | Q99020 | Hnrnpab | 31 kDa | 2 C | 1.107 | 0.60716 |
| 4-hydroxyphenylpyruvate dioxygenase | P49429 | Hpd | 45 kDa | 4 C | 1.107 | 0.10616 |
| Isoform 2 of Cytosol aminopeptidase | Q9CPY7-2 (+ | Lap3 | 53 kDa | 7 C | 1.106 | 0.13857 |
| Endoribonuclease LACTB2 | Q99KR3 | Lactb2 | 33 kDa | 5 C | 1.106 | 0.42668 |
| Histidine ammonia-lyase | P35492 | Hal | 72 kDa | 12 C | 1.106 | 0.57247 |
| Hydroxymethylglutaryl-CoA synthase, mitochondrial | P54869 | Hmgcs2 | 57 kDa | 9 C | 1.105 | 0.60865 |
| Superoxide dismutase [Mn], mitochondrial | P09671 | Sod2 | 25 kDa | 4 C | 1.105 | 0.82281 |
| Complement component C8 gamma chain | Q8VCG4 | C8g | 23 kDa | 3 C | 1.100 | 0.51851 |
| Dimethylglycine dehydrogenase, mitochondrial | Q9DBT9 | Dmgdh | 97 kDa | 4 C | 1.100 | 0.73382 |
| Inorganic pyrophosphatase 2, mitochondrial | Q91VM9 | Ppa2 | 38 kDa | 8 C | 1.100 | 0.86249 |
| Polyadenylate-binding protein 1 | P29341 | Pabpc1 | 71 kDa | 4 C | 1.098 | 0.72111 |
| Golgi reassembly-stacking protein 2 | Q99JX3 | Gorasp2 | 47 kDa | 4 C | 1.091 | 0.82979 |
| Fructose-bisphosphate aldolase B | Q91Y97 | Aldob | 40 kDa | 8 C | 1.087 | 0.41081 |
| Protein disulfide-isomerase A3 | P27773 | Pdia3 | 57 kDa | 8 C | 1.086 | 0.47837 |
| S-adenosylmethionine synthase isoform type-1 | Q91X83 | Mat1a | 44 kDa | 10 C | 1.083 | 0.81490 |
| Stress-70 protein, mitochondrial | P38647 | Hspa9 | 73 kDa | 5 C | 1.083 | 0.86022 |
| Isoform 3 of Heterogeneous nuclear ribonucleoprotein D0 | Q60668-3 | Hnrnpd | 33 kDa | 3 C | 1.077 | 0.69806 |
| Cysteine-rich protein 2 | Q9DCT8 | Crip2 | 23 kDa | 14 C | 1.077 | 0.74152 |
| Alpha-1-antitrypsin 1-1 | P07758 | Serpina1a | 46 kDa | 3 C | 1.077 | 0.87899 |
| Leucine-rich repeat flightless-interacting protein 1 | Q3UZ39 | Lrrfip1 | 79 kDa | 12 C | 1.077 | 0.89108 |
| 14-3-3 protein theta | P68254 | Ywhaq | 28 kDa | 5 C | 1.077 | 0.9027 |

| | | | | | | |
|---|---|---|---|---|---|---|
| Transgelin-2 | Q9WVA4 | Tagln2 | 22 kDa | 3 C | 1.074 | 0.49176 |
| UV excision repair protein RAD23 homolog B | P54728 | Rad23b | 44 kDa | 1 C | 1.074 | 0.7488 |
| Peroxisomal multifunctional enzyme type 2 | P51660 | Hsd17b4 | 79 kDa | 9 C | 1.072 | 0.77405 |
| 26S proteasome non-ATPase regulatory subunit 4 | O35226 | Psmd4 | 41 kDa | 4 C | 1.071 | 0.64332 |
| Endoplasmic reticulum resident protein 29 | P57759 | Erp29 | 29 kDa | 1 C | 1.071 | 0.64332 |
| Methylmalonate-semialdehyde dehydrogenase [acylating], mitochondrial | Q9EQ20 | Aldh6a1 | 58 kDa | 8 C | 1.069 | 0.52654 |
| Profilin-1 | P62962 | Pfn1 | 15 kDa | 3 C | 1.067 | 0.76764 |
| Propionyl-CoA carboxylase beta chain, mitochondrial | Q99MN9 | Pccb | 58 kDa | 11 C | 1.067 | 0.80074 |
| Mannose-binding protein A | P39039 | Mbl1 | 25 kDa | 8 C | 1.067 | 0.87215 |
| 14-3-3 protein zeta/delta | P63101 | Ywhaz | 28 kDa | 3 C | 1.065 | 0.44182 |
| Alcohol dehydrogenase 1 | P00329 | Adh1 | 40 kDa | 15 C | 1.059 | 0.84159 |
| Acetyl-CoA acetyltransferase, cytosolic | Q8CAY6 | Acat2 | 41 kDa | 8 C | 1.059 | 0.0161 |
| Serine hydroxymethyltransferase, cytosolic | P50431 | Shmt1 | 53 kDa | 10 C | 1.056 | 0.45862 |
| Arginase-1 | Q61176 | Arg1 | 35 kDa | 3 C | 1.056 | 0.73624 |
| Carbonic anhydrase 2 | P00920 | Ca2 | 29 kDa | 2 C | 1.056 | 0.81490 |
| Fibrinogen gamma chain | Q8VCM7 | Fgg | 49 kDa | 12 C | 1.055 | 0.62395 |
| Fumarylacetoacetase | P35505 | Fah | 46 kDa | 6 C | 1.055 | 0.73714 |
| Bifunctional epoxide hydrolase 2 | P34914 | Ephx2 | 63 kDa | 10 C | 1.050 | 0.87485 |
| Methylthioribulose-1-phosphate dehydratase | Q9WVQ5 | Apip | 27 kDa | 11 C | 1.047 | 0.82641 |
| Apoptosis-inducing factor 1, mitochondrial | Q9Z0X1 | Aifm1 | 67 kDa | 4 C | 1.045 | 0.80248 |
| Protein disulfide-isomerase A4 | P08003 | Pdia4 | 72 kDa | 6 C | 1.044 | 0.62395 |
| Aldehyde dehydrogenase, mitochondrial | P47738 | Aldh2 | 57 kDa | 9 C | 1.037 | 0.82090 |
| Hydroxyacid oxidase 1 | Q9WU19 | Hao1 | 41 kDa | 5 C | 1.037 | 0.79525 |
| Heterogeneous nuclear ribonucleoproteins A2/B1 | O88569 | Hnrnpa2b1 | 37 kDa | 1 C | 1.033 | 0.88202 |
| Protein DEK | Q7TNV0 | Dek | 43 kDa | 4 C | 1.033 | 0.95447 |
| Ig kappa chain C region | P01837 | | 12 kDa | 3 C | 1.031 | 0.88202 |
| Acetyl-CoA acetyltransferase, mitochondrial | Q8QZT1 | Acat1 | 45 kDa | 6 C | 1.030 | 0.81898 |
| Fibrinogen alpha chain | E9PV24 | Fga | 87 kDa | 13 C | 1.029 | 0.82979 |
| Non-specific lipid-transfer protein | P32020 | Scp2 | 59 kDa | 11 C | 1.027 | 0.80581 |
| Ester hydrolase C11orf54 homolog | Q91V76 | | 35 kDa | 8 C | 1.025 | 0.93864 |
| Protein transport protein Sec31A | Q3UPL0 | Sec31a | 134 kDa | 18 C | 1.025 | 0.96508 |
| PDZ and LIM domain protein 5 | Q8CI51 | Pdlim5 | 63 kDa | 22 C | 1.025 | 0.96508 |
| Sorbin and SH3 domain-containing protein 1 | Q62417 | Sorbs1 | 143 kDa | 4 C | 1.025 | 0.96508 |
| Na(+)/H(+) exchange regulatory cofactor NHE-RF1 | P70441 | Slc9a3r1 | 39 kDa | 5 C | 1.024 | 0.92281 |
| Serotransferrin | Q921I1 | Tf | 77 kDa | 38 C | 1.024 | 0.70959 |
| Gamma-butyrobetaine dioxygenase | Q924Y0 | Bbox1 | 45 kDa | 9 C | 1.023 | 0.88746 |
| Mitochondrial peptide methionine sulfoxide reductase | Q9D6Y7 | Msra | 26 kDa | 4 C | 1.021 | 0.69176 |
| Adenylate kinase 2, mitochondrial | Q9WTP6 (+1 | Ak2 | 26 kDa | 5 C | 1.021 | 0.93780 |
| Triosephosphate isomerase | P17751 | Tpi1 | 32 kDa | 9 C | 1.019 | 0.89809 |
| Sulfite oxidase, mitochondrial | Q8R086 | Suox | 61 kDa | 9 C | 1.018 | 0.82979 |
| Far upstream element-binding protein 2 | Q3U0V1 | Khsrp | 77 kDa | 8 C | 1.017 | 0.93381 |
| Heat shock 70 kDa protein 4 | Q61316 | Hspa4 | 94 kDa | 14 C | 1.015 | 0.93715 |
| Selenium-binding protein 1 | P17563 | Selenbp1 | 53 kDa | 10 C | 1.013 | 0.9126 |
| Peroxiredoxin-1 | P35700 | Prdx1 | 22 kDa | 4 C | 1.005 | 0.96647 |
| Selenium-binding protein 2 | Q63836 | Selenbp2 | 53 kDa | 10 C | 1.000 | |
| Heat shock cognate 71 kDa protein | P63017 | Hspa8 | 71 kDa | 4 C | 1.000 | |
| Pyruvate kinase PKLR | P53657 | Pklr | 62 kDa | 6 C | 1.000 | |
| Urocanate hydratase | Q8VC12 | Uroc1 | 75 kDa | 12 C | 1.000 | |
| Inorganic pyrophosphatase | Q9D819 | Ppa1 | 33 kDa | 8 C | 1.000 | |
| Hemopexin | Q91X72 | Hpx | 51 kDa | 13 C | 1.000 | |
| Proteasome subunit beta type-4 | P99026 | Psmb4 | 29 kDa | 2 C | 1.000 | |
| Ethanolamine-phosphate cytidylyltransferase | Q922E4 | Pcyt2 | 45 kDa | 8 C | 1.000 | |
| DAZ-associated protein 1 | Q9JII5 (+1) | Dazap1 | 43 kDa | 4 C | 1.000 | |
| NAD kinase 2, mitochondrial | Q8C5H8 | Nadk2 | 51 kDa | 9 C | 1.000 | |
| Complement factor H | P06909 | Cfh | 139 kDa | 82 C | 1.000 | |
| Osteoclast-stimulating factor 1 | Q62422 | Ostf1 | 24 kDa | 4 C | 1.000 | |
| Galectin-1 | P16045 | Lgals1 | 15 kDa | 6 C | 1.000 | |
| Ketohexokinase | P97328 | Khk | 33 kDa | 10 C | 1.000 | |
| Endonuclease G, mitochondrial | O08600 | Endog | 32 kDa | 2 C | 1.000 | |
| UBX domain-containing protein 1 | Q922Y1 | Ubxn1 | 34 kDa | 2 C | 1.000 | |
| Extracellular matrix protein 1 | Q61508 | Ecm1 | 63 kDa | 29 C | 1.000 | |
| Isoform 3 of Peroxisomal coenzyme A diphosphatase NUDT7 | Q99P30-3 | Nudt7 | 25 kDa | 4 C | 1.000 | |
| Serine/threonine-protein phosphatase 5 | Q60676 | Ppp5c | 57 kDa | 11 C | 1.000 | |
| Heterogeneous nuclear ribonucleoprotein F | Q9Z2X1 | Hnrnpf | 46 kDa | 6 C | 1.000 | |
| Oligoribonuclease, mitochondrial | Q9D8S4 | Rexo2 | 27 kDa | 4 C | 1.000 | |
| Aldo-keto reductase family 1 member C13 | Q8VC28 | Akr1c13 | 37 kDa | 10 C | 1.000 | |
| Eukaryotic translation initiation factor 2 subunit 1 | Q6ZWX6 | Eif2s1 | 36 kDa | 5 C | 1.000 | |
| Tubulin-folding cofactor B | Q9D1E6 | Tbcb | 27 kDa | 5 C | 1.000 | |
| Gamma-soluble NSF attachment protein | Q9CWZ7 | Napg | 35 kDa | 6 C | 1.000 | |
| Serine/threonine-protein phosphatase 2A 55 kDa regulatory subunit B alpha isoform | Q6P1F6 | Ppp2r2a | 52 kDa | 9 C | 1.000 | |
| Probable ATP-dependent RNA helicase DDX17 | Q501J6 | Ddx17 | 72 kDa | 11 C | 1.000 | |
| Isoform 2 of Galectin-9 | O08573-2 (+ | Lgals9 | 37 kDa | 7 C | 1.000 | |
| Thioredoxin domain-containing protein 17 | Q9CQM5 | Txndc17 | 14 kDa | 6 C | 1.000 | |
| WW domain-binding protein 2 | P97765 | Wbp2 | 28 kDa | 2 C | 1.000 | |
| Inhibitor of carbonic anhydrase | Q9DBD0 | Ica | 77 kDa | 35 C | 1.000 | |
| RNA-binding protein FUS | P56959 | Fus | 53 kDa | 4 C | 1.000 | |
| Plasminogen | P20918 | Plg | 91 kDa | 48 C | 1.000 | |
| Isoform 2 of Cysteine--tRNA ligase, cytoplasmic | Q9ER72-2 | Cars | 86 kDa | 11 C | 1.000 | |

| | | | | | | |
|---|---|---|---|---|---|---|
| Actin-related protein 2/3 complex subunit 4 | P59999 | Arpc4 | 20 kDa | 4 C | 1.000 | |
| Methionine aminopeptidase 1 | Q8BP48 | Metap1 | 43 kDa | 17 C | 1.000 | |
| Catenin alpha-1 | P26231 | Ctnna1 | 100 kDa | 12 C | 1.000 | |
| Enoyl-CoA delta isomerase 1, mitochondrial | P42125 | Eci1 | 32 kDa | 5 C | 1.000 | |
| 2-oxo-4-hydroxy-4-carboxy-5-ureidoimidazoline decarboxylase | Q283N4 | Urad | 20 kDa | 3 C | 1.000 | |
| Clathrin interactor 1 | Q99KN9 (+1) | Clint1 | 69 kDa | 1 C | 1.000 | |
| 14-3-3 protein eta | P68510 | Ywhah | 28 kDa | 3 C | 1.000 | |
| Transgelin | P37804 | Tagln | 23 kDa | 1 C | 1.000 | |
| Ig lambda-1 chain C region | P01843 | | 12 kDa | 3 C | 1.000 | |
| Deoxynucleoside triphosphate triphosphohydrolase SAMHD1 | Q60710 | Samhd1 | 73 kDa | 17 C | 1.000 | |
| Transketolase | P40142 | Tkt | 68 kDa | 12 C | 0.988 | 0.88202 |
| Peroxiredoxin-4 | O08807 | Prdx4 | 31 kDa | 4 C | 0.988 | 0.92321 |
| Glycine dehydrogenase (decarboxylating), mitochondrial | Q91W43 | Gldc | 113 kDa | 24 C | 0.984 | 0.96901 |
| Peroxiredoxin-6 | O08709 | Prdx6 | 25 kDa | 2 C | 0.981 | 0.92155 |
| Glutathione S-transferase Mu 1 | P10649 | Gstm1 | 26 kDa | 2 C | 0.981 | 0.86827 |
| Isoform 2 of F-actin-capping protein subunit beta | P47757-2 | Capzb | 31 kDa | 5 C | 0.980 | 0.97263 |
| Ribokinase | Q8R1Q9 | Rbks | 34 kDa | 9 C | 0.980 | 0.97959 |
| Epsin-1 | Q80VP1 (+1) | Epn1 | 60 kDa | 2 C | 0.980 | 0.98154 |
| Ethanolamine-phosphate phospho-lyase | Q8BWU8 | Etnppl | 55 kDa | 10 C | 0.979 | 0.9106 |
| 14-3-3 protein epsilon | P62259 | Ywhae | 29 kDa | 3 C | 0.977 | 0.91713 |
| 3-ketoacyl-CoA thiolase A, peroxisomal | Q921H8 | Acaa1a | 44 kDa | 8 C | 0.976 | 0.91246 |
| Peptidyl-prolyl cis-trans isomerase FKBP3 | Q62446 | Fkbp3 | 25 kDa | 1 C | 0.976 | 0.96508 |
| Scaffold attachment factor B1 | D3YXK2 | Safb | 105 kDa | 9 C | 0.969 | 0.97751 |
| Isoform 2 of Tropomyosin alpha-3 chain | P21107-2 | Tpm3 | 29 kDa | 1 C | 0.968 | 0.92624 |
| FAS-associated death domain protein | Q61160 | Fadd | 23 kDa | 3 C | 0.968 | 0.95447 |
| Homogentisate 1,2-dioxygenase | O09173 | Hgd | 50 kDa | 14 C | 0.963 | 0.87564 |
| Nucleoside diphosphate kinase A | P15532 | Nme1 | 17 kDa | 2 C | 0.963 | 0.82979 |
| Proteasome subunit alpha type-6 | Q9QUM9 | Psma6 | 27 kDa | 8 C | 0.963 | 0.84159 |
| Selenocysteine lyase | Q9JLI6 | Scly | 47 kDa | 8 C | 0.963 | 0.85930 |
| Enoyl-CoA hydratase, mitochondrial | Q8BH95 | Echs1 | 31 kDa | 7 C | 0.963 | 0.89640 |
| 40S ribosomal protein SA | P14206 | Rpsa | 33 kDa | 2 C | 0.957 | 0.85196 |
| Glycerol-3-phosphate phosphatase | Q8CHP8 | Pgp | 35 kDa | 8 C | 0.955 | 0.96434 |
| Aldehyde dehydrogenase, cytosolic 1 | O35945 | Aldh1a7 | 55 kDa | 8 C | 0.954 | 0.66870 |
| Catalase | P24270 | Cat | 60 kDa | 5 C | 0.953 | 0.51487 |
| Actin-related protein 3 | Q99JY9 | Actr3 | 47 kDa | 8 C | 0.952 | 0.82979 |
| Serine beta-lactamase-like protein LACTB, mitochondrial | Q9EP89 | Lactb | 61 kDa | 4 C | 0.938 | 0.82979 |
| Isoleucine--tRNA ligase, mitochondrial | Q8BIJ6 | Iars2 | 113 kDa | 20 C | 0.938 | 0.89640 |
| Betaine--homocysteine S-methyltransferase 1 | O35490 | Bhmt | 45 kDa | 8 C | 0.937 | 0.73554 |
| Methyltransferase-like 26 | Q9DCS2 | Mettl26 | 23 kDa | 6 C | 0.933 | 0.84159 |
| Haptoglobin | Q61646 | Hp | 39 kDa | 9 C | 0.931 | 0.37390 |
| Cofilin-2 | P45591 | Cfl2 | 19 kDa | 2 C | 0.923 | 0.64332 |
| Fucose mutarotase | Q8R2K1 | Fuom | 17 kDa | 3 C | 0.923 | 0.72465 |
| Isoform 2 of Peroxisomal acyl-coenzyme A oxidase 1 | Q9R0H0-2 | Acox1 | 75 kDa | 7 C | 0.917 | 0.61044 |
| 40S ribosomal protein S20 | P60867 | Rps20 | 13 kDa | 2 C | 0.917 | 0.64332 |
| Transaldolase | Q93092 | Taldo1 | 37 kDa | 3 C | 0.917 | 0.7113 |
| Polymerase delta-interacting protein 2 | Q91VA6 | Poldip2 | 42 kDa | 4 C | 0.917 | 0.81490 |
| Mitochondrial antiviral-signaling protein | Q8VCF0 | Mavs | 53 kDa | 8 C | 0.914 | 0.75416 |
| Aconitate hydratase, mitochondrial | Q99KI0 | Aco2 | 85 kDa | 13 C | 0.912 | 0.68309 |
| Ubiquitin-60S ribosomal protein L40 | P62984 | Uba52 | 15 kDa | 5 C | 0.909 | 0.51851 |
| Glutamate--cysteine ligase regulatory subunit | O09172 | Gclm | 31 kDa | 6 C | 0.909 | 0.51851 |
| StAR-related lipid transfer protein 5 | Q9EPQ7 | Stard5 | 24 kDa | 8 C | 0.909 | 0.83401 |
| Glycine N-methyltransferase | Q9QXF8 | Gnmt | 33 kDa | 8 C | 0.900 | 0.47342 |
| Inosine-5'-monophosphate dehydrogenase 2 | P24547 | Impdh2 | 56 kDa | 7 C | 0.900 | 0.87739 |
| CD5 antigen-like | Q9QWK4 | Cd5l | 39 kDa | 26 C | 0.896 | 0.52623 |
| Carbonic anhydrase 5A, mitochondrial | P23589 | Ca5a | 34 kDa | 7 C | 0.895 | 0.77804 |
| Pyridoxal kinase | Q8K183 | Pdxk | 35 kDa | 5 C | 0.891 | 0.65842 |
| Fibrinogen beta chain | Q8K0E8 | Fgb | 55 kDa | 12 C | 0.890 | 0.39239 |
| Kynurenine--oxoglutarate transaminase 1 | Q8BTY1 | Kyat1 | 48 kDa | 7 C | 0.889 | 0.60865 |
| 60S ribosomal protein L30 | P62889 | Rpl30 | 13 kDa | 3 C | 0.889 | 0.64332 |
| Stress-induced-phosphoprotein 1 | Q60864 | Stip1 | 63 kDa | 11 C | 0.888 | 0.54190 |
| Protein DJ-1 | Q99LX0 | Park7 | 20 kDa | 4 C | 0.882 | 0.44182 |
| Cytochrome b-c1 complex subunit Rieske, mitochondrial | Q9CR68 | Uqcrfs1 | 29 kDa | 5 C | 0.882 | 0.70959 |
| Ornithine aminotransferase, mitochondrial | P29758 | Oat | 48 kDa | 7 C | 0.882 | 0.75146 |
| Ribosyldihydronicotinamide dehydrogenase [quinone] | Q9JI75 | Nqo2 | 26 kDa | 4 C | 0.882 | 0.77557 |
| Peptidyl-prolyl cis-trans isomerase A | P17742 | Ppia | 18 kDa | 3 C | 0.877 | 0.52250 |
| Presequence protease, mitochondrial | Q8K411 (+1) | Pitrm1 | 117 kDa | 20 C | 0.875 | 0.62909 |
| Src substrate cortactin | Q60598 | Cttn | 61 kDa | 3 C | 0.875 | 0.67786 |
| Actin-related protein 2 | P61161 | Actr2 | 45 kDa | 5 C | 0.875 | 0.67786 |
| 4-aminobutyrate aminotransferase, mitochondrial | P61922 | Abat | 56 kDa | 12 C | 0.875 | 0.82029 |
| O-phosphoseryl-tRNA(Sec) selenium transferase | Q6P6M7 | Sepsecs | 55 kDa | 13 C | 0.872 | 0.68882 |
| 4-trimethylaminobutyraldehyde dehydrogenase | Q9JLJ2 | Aldh9a1 | 54 kDa | 17 C | 0.867 | 0.26057 |
| Probable aminopeptidase NPEPL1 | Q6NSR8 | Npepl1 | 56 kDa | 16 C | 0.867 | 0.52654 |
| Dihydrolipoyllysine-residue succinyltransferase component of 2-oxoglutarate dehydrogenase complex, mitochondrial | Q9D2G2 | Dlst | 49 kDa | 6 C | 0.865 | 0.52433 |
| Protein DDI1 homolog 2 | A2ADY9 | Ddi2 | 45 kDa | 8 C | 0.864 | 0.674 |
| Pregnancy zone protein | Q61838 | Pzp | 166 kDa | 24 C | 0.862 | 0.47662 |
| Cytosolic 10-formyltetrahydrofolate dehydrogenase | Q8R0Y6 | Aldh1l1 | 99 kDa | 15 C | 0.859 | 0.70206 |
| S-formylglutathione hydrolase | Q9R0P3 | Esd | 31 kDa | 10 C | 0.857 | 0.67017 |
| BAG family molecular chaperone regulator 3 | Q9JLV1 | Bag3 | 62 kDa | 4 C | 0.857 | 0.70959 |

| | | | | | | |
|---|---|---|---|---|---|---|
| Aminoacylase-1 | Q99JW2 | Acy1 | 46 kDa | 4 C | 0.855 | 0.14814 |
| Ig heavy chain V regions TEPC 15/S107/HPCM1/HPCM2/HPCM3 | P01787 (+4) | | 14 kDa | 2 C | 0.850 | 0.7849 |
| Ribosome-recycling factor, mitochondrial | Q9D6S7 | Mrrf | 29 kDa | 2 C | 0.846 | 0.51851 |
| 3-ketoacyl-CoA thiolase B, peroxisomal | Q8VCH0 | Acaa1b | 44 kDa | 9 C | 0.846 | 0.52654 |
| Isoform 3 of Calpastatin | P51125-3 | Cast | 80 kDa | 4 C | 0.846 | 0.74152 |
| Dihydropyrimidinase-related protein 2 | O08553 | Dpysl2 | 62 kDa | 7 C | 0.838 | 0.48215 |
| Iron-sulfur cluster assembly enzyme ISCU, mitochondrial | Q9D7P6 | Iscu | 18 kDa | 4 C | 0.833 | 0.37390 |
| START domain-containing protein 10 | Q9JMD3 | Stard10 | 33 kDa | 7 C | 0.833 | 0.43533 |
| Mesencephalic astrocyte-derived neurotrophic factor | Q9CXI5 | Manf | 20 kDa | 8 C | 0.833 | 0.64332 |
| Copine-3 | Q8BT60 | Cpne3 | 60 kDa | 13 C | 0.833 | 0.72465 |
| L-lactate dehydrogenase A chain | P06151 | Ldha | 36 kDa | 6 C | 0.833 | 0.72465 |
| Septin-7 | O55131 | 7-Sep | 51 kDa | 6 C | 0.824 | 0.75416 |
| Peptidyl-prolyl cis-trans isomerase FKBP4 | P30416 | Fkbp4 | 52 kDa | 7 C | 0.823 | 0.36298 |
| 60S ribosomal protein L12 | P35979 | Rpl12 | 18 kDa | 3 C | 0.818 | 0.33485 |
| Methionine adenosyltransferase 2 subunit beta | Q99LB6 | Mat2b | 37 kDa | 7 C | 0.818 | 0.37390 |
| Histidine-rich glycoprotein | Q9ESB3 | Hrg | 59 kDa | 17 C | 0.818 | 0.56143 |
| Aspartyl aminopeptidase | Q9Z2W0 | Dnpep | 52 kDa | 10 C | 0.817 | 0.569 |
| Cytochrome c, somatic | P62897 | Cycs | 12 kDa | 2 C | 0.815 | 0.06676 |
| Protein phosphatase 1B | P36993 | Ppm1b | 43 kDa | 12 C | 0.813 | 0.63876 |
| Septin-10 | Q8C650 | 10-Sep | 52 kDa | 14 C | 0.810 | 0.79990 |
| Xaa-Pro dipeptidase | Q11136 | Pepd | 55 kDa | 17 C | 0.808 | 0.27943 |
| 2-hydroxyacyl-CoA lyase 1 | Q9QXE0 | Hacl1 | 64 kDa | 17 C | 0.806 | 0.39124 |
| Beta-ureidopropionase | Q8VC97 | Upb1 | 44 kDa | 10 C | 0.800 | 0.46563 |
| Serine/threonine-protein phosphatase CPPED1 | Q8BFS6 | Cpped1 | 35 kDa | 6 C | 0.800 | 0.51851 |
| LIM and SH3 domain protein 1 | Q61792 | Lasp1 | 30 kDa | 7 C | 0.800 | 0.53418 |
| Antithrombin-III | P32261 | Serpinc1 | 52 kDa | 9 C | 0.800 | 0.65299 |
| von Willebrand factor A domain-containing protein 5A | Q99KC8 | Vwa5a | 87 kDa | 13 C | 0.794 | 0.5681 |
| Glutamine synthetase | P15105 | Glul | 42 kDa | 13 C | 0.789 | 0.56879 |
| Gephyrin | Q8BUV3 | Gphn | 83 kDa | 13 C | 0.788 | 0.7627 |
| Phospholipid hydroperoxide glutathione peroxidase, mitochondrial | O70325 (+1) | Gpx4 | 22 kDa | 10 C | 0.787 | 0.28263 |
| G protein-regulated inducer of neurite outgrowth 3 | Q8BWS5 | Gprin3 | 80 kDa | 16 C | 0.784 | 0.7796 |
| N-myc-interactor | O35309 | Nmi | 35 kDa | 8 C | 0.780 | 0.827 |
| Target of Myb protein 1 | O88746 | Tom1 | 54 kDa | 4 C | 0.780 | 0.827 |
| Probable imidazolonepropionase | Q9DBA8 | Amdhd1 | 46 kDa | 11 C | 0.779 | 0.34540 |
| Ig kappa chain V-II region 26-10 | P01631 | | 12 kDa | 2 C | 0.778 | 0.56143 |
| Serine/threonine-protein phosphatase 2A catalytic subunit alpha isoform | P63330 | Ppp2ca | 36 kDa | 10 C | 0.778 | 0.60865 |
| Toll-interacting protein | Q9QZ06 | Tollip | 30 kDa | 4 C | 0.775 | 0.6646 |
| Isoaspartyl peptidase/L-asparaginase | Q8C0M9 | Asrgl1 | 34 kDa | 8 C | 0.773 | 0.47850 |
| Elongation factor 1-alpha 1 | P10126 | Eef1a1 | 50 kDa | 6 C | 0.771 | 0.46852 |
| Insulin-degrading enzyme | Q9JHR7 | Ide | 118 kDa | 13 C | 0.767 | 0.46951 |
| Glutathione S-transferase Mu 2 | P15626 | Gstm2 | 26 kDa | 3 C | 0.767 | 0.15501 |
| Poly(rC)-binding protein 1 | P60335 | Pcbp1 | 37 kDa | 9 C | 0.765 | 0.23019 |
| Xanthine dehydrogenase/oxidase | Q00519 | Xdh | 147 kDa | 37 C | 0.764 | 0.52619 |
| Isoform 6 of Palladin | Q9ET54-6 | Palld | 151 kDa | 17 C | 0.763 | 0.68078 |
| Fatty acid-binding protein, adipocyte | P04117 | Fabp4 | 15 kDa | 2 C | 0.756 | 0.71110 |
| 40S ribosomal protein S21 | Q9CQR2 | Rps21 | 9 kDa | 2 C | 0.756 | 0.75967 |
| GTP cyclohydrolase 1 feedback regulatory protein | P99025 | Gchfr | 10 kDa | 2 C | 0.756 | 0.75967 |
| Dihydropyrimidinase | Q9EQF5 | Dpys | 57 kDa | 9 C | 0.750 | 0.20065 |
| Actin-related protein 2/3 complex subunit 5 | Q9CPW4 | Arpc5 | 16 kDa | 1 C | 0.750 | 0.37390 |
| Protein LSM12 homolog | Q9D0R8 | Lsm12 | 22 kDa | 5 C | 0.750 | 0.37390 |
| Glycogen phosphorylase, liver form | Q9ET01 | Pygl | 97 kDa | 8 C | 0.747 | 0.17733 |
| Cystathionine beta-synthase | Q91WT9 (+1 | Cbs | 62 kDa | 13 C | 0.740 | 0.50569 |
| Uncharacterized protein C1orf50 homolog | Q5EBG8 | | 22 kDa | 3 C | 0.732 | 0.69078 |
| Prolyl endopeptidase | Q9QUR6 | Prep | 81 kDa | 17 C | 0.729 | 0.70928 |
| Isoform 2 of Cellular nucleic acid-binding protein | P53996-2 (+1 | Cnbp | 19 kDa | 22 C | 0.727 | 0.10119 |
| Secernin-3 | Q3TMH2 | Scrn3 | 48 kDa | 7 C | 0.727 | 0.34864 |
| PDZ and LIM domain protein 1 | O70400 | Pdlim1 | 36 kDa | 8 C | 0.727 | 0.34864 |
| 5-hydroxyisourate hydrolase | Q9CRB3 | Urah | 14 kDa | 3 C | 0.727 | 0.46760 |
| Vigilin | Q8VDJ3 | Hdlbp | 142 kDa | 10 C | 0.719 | 0.46090 |
| Microsomal glutathione S-transferase 1 | Q91VS7 | Mgst1 | 18 kDa | 1 C | 0.718 | 0.70861 |
| Sorting nexin-3 | O70492 | Snx3 | 19 kDa | 1 C | 0.714 | 0.23019 |
| Ig heavy chain V region AC38 205.12 | P06330 | | 13 kDa | 2 C | 0.714 | 0.23019 |
| Crk-like protein | P47941 | Crkl | 34 kDa | 2 C | 0.714 | 0.23019 |
| Aspartoacylase | Q8R3P0 | Aspa | 35 kDa | 8 C | 0.714 | 0.51851 |
| 14 kDa phosphohistidine phosphatase | Q9DAK9 | Phpt1 | 14 kDa | 3 C | 0.714 | 0.51851 |
| Na(+)/H(+) exchange regulatory cofactor NHE-RF3 | Q9JIL4 | Pdzk1 | 56 kDa | 7 C | 0.712 | 0.02294 |
| Septin-9 | Q80UG5 | 9-Sep | 66 kDa | 6 C | 0.710 | 0.73843 |
| Ribosome-binding protein 1 | Q99PL5 | Rrbp1 | 173 kDa | 8 C | 0.696 | 0.31218 |
| Dihydropyrimidine dehydrogenase [NADP(+)] | Q8CHR6 | Dpyd | 111 kDa | 35 C | 0.693 | 0.26366 |
| Cold shock domain-containing protein E1 | Q91W50 | Csde1 | 89 kDa | 15 C | 0.683 | 0.57307 |
| Isoform 2 of Heterogeneous nuclear ribonucleoprotein M | Q9D0E1-2 | Hnrnpm | 74 kDa | 6 C | 0.677 | 0.62226 |
| Caspase-7 | P97864 | Casp7 | 34 kDa | 11 C | 0.677 | 0.62226 |
| 3-hydroxybutyrate dehydrogenase type 2 | Q8JZV9 | Bdh2 | 27 kDa | 6 C | 0.677 | 0.62226 |
| Mitochondrial fission 1 protein | Q9CQ92 | Fis1 | 17 kDa | 1 C | 0.667 | 0.13177 |
| Serine/arginine-rich splicing factor 3 | P84104 (+1) | Srsf3 | 19 kDa | 4 C | 0.667 | 0.28786 |
| TAR DNA-binding protein 43 | Q921F2 | Tardbp | 45 kDa | 7 C | 0.667 | 0.28786 |
| Aldehyde oxidase 1 | O54754 | Aox1 | 147 kDa | 41 C | 0.667 | 0.33625 |
| Phosphopantothenoylcysteine decarboxylase | Q8BZB2 | Ppcdc | 22 kDa | 6 C | 0.667 | 0.43533 |

| | | | | | | |
|---|---|---|---|---|---|---|
| Isocitrate dehydrogenase [NADP] cytoplasmic | Q88844 | Idh1 | 47 kDa | 7 C | 0.647 | 0.01336 |
| Alanine aminotransferase 1 | Q8QZR5 | Gpt | 55 kDa | 14 C | 0.646 | 0.02452 |
| Fatty acid-binding protein, epidermal | Q05816 | Fabp5 | 15 kDa | 6 C | 0.641 | 0.01779 |
| S-adenosylmethionine synthase isoform type-2 | Q3THS6 | Mat2a | 44 kDa | 6 C | 0.636 | 0.42164 |
| Pyrethroid hydrolase Ces2e | Q8BK48 | Ces2e | 62 kDa | 7 C | 0.625 | 0.2508 |
| Stromal cell-derived factor 2 | Q9DCT5 | Sdf2 | 23 kDa | 4 C | 0.625 | 0.4168 |
| Isoform 2 of Heterogeneous nuclear ribonucleoprotein Q | Q7TMK9-2 | Syncrip | 63 kDa | 4 C | 0.621 | 0.18015 |
| Splicing factor, proline- and glutamine-rich | Q8VIJ6 | Sfpq | 75 kDa | 7 C | 0.620 | 0.37976 |
| NADH dehydrogenase [ubiquinone] iron-sulfur protein 6, mitochondrial | P52503 | Ndufs6 | 13 kDa | 3 C | 0.620 | 0.37976 |
| Aflatoxin B1 aldehyde reductase member 2 | Q8CG76 | Akr7a2 | 41 kDa | 8 C | 0.620 | 0.50393 |
| Trimethyllysine dioxygenase, mitochondrial | Q91ZE0 | Tmlhe | 50 kDa | 11 C | 0.619 | 0.43450 |
| Hypoxanthine-guanine phosphoribosyltransferase | P00493 | Hprt1 | 25 kDa | 4 C | 0.615 | 0.07420 |
| Molybdenum cofactor biosynthesis protein 1 | Q5RKZ7 | Mocs1 | 70 kDa | 15 C | 0.610 | 0.61538 |
| Filamin-B | Q80X90 | Flnb | 278 kDa | 43 C | 0.608 | 0.54609 |
| 40S ribosomal protein S12 | P63323 | Rps12 | 15 kDa | 7 C | 0.606 | 0.03137 |
| Multifunctional protein ADE2 | Q9DCL9 | Paics | 47 kDa | 13 C | 0.605 | 0.26226 |
| Complement C3 | P01027 | C3 | 186 kDa | 27 C | 0.600 | 0.1614 |
| Cytosolic purine 5'-nucleotidase | Q3V1L4 | Nt5c2 | 65 kDa | 8 C | 0.600 | 0.37390 |
| Ig kappa chain V-V region MOPC 41 | P01639 | Gm5571 | 14 kDa | 3 C | 0.600 | 0.37390 |
| S-methyl-5'-thioadenosine phosphorylase | Q9CQ65 | Mtap | 31 kDa | 10 C | 0.600 | 0.38640 |
| Aspartate aminotransferase, mitochondrial | P05202 | Got2 | 47 kDa | 7 C | 0.600 | 0.0161 |
| Alpha-mannosidase 2C1 | Q91W89 | Man2c1 | 116 kDa | 24 C | 0.595 | 0.71604 |
| Glutaredoxin-1 | Q9QUH0 | Glrx | 12 kDa | 5 C | 0.586 | 0.35245 |
| 6-phosphofructo-2-kinase/fructose-2,6-bisphosphatase 1 | P70266 | Pfkfb1 | 55 kDa | 10 C | 0.575 | 0.15607 |
| Beta-arrestin-1 | Q8BWG8 (+1 | Arrb1 | 47 kDa | 8 C | 0.556 | 0.11611 |
| Eukaryotic translation initiation factor 2 subunit 3, X-linked | Q9Z0N1 | Eif2s3x | 51 kDa | 10 C | 0.556 | 0.46852 |
| Complement C4-B | P01029 | C4b | 193 kDa | 29 C | 0.550 | 0.44901 |
| Eukaryotic translation initiation factor 4 gamma 3 | Q80XI3 (+1) | Eif4g3 | 175 kDa | 23 C | 0.545 | 0.65913 |
| Protein argonaute-2 | Q8CJG0 | Ago2 | 97 kDa | 22 C | 0.545 | 0.65913 |
| Protein farnesyltransferase subunit beta | Q8K2I1 | Fntb | 49 kDa | 17 C | 0.545 | 0.65913 |
| Delta-1-pyrroline-5-carboxylate dehydrogenase, mitochondrial | Q8CHT0 | Aldh4a1 | 62 kDa | 8 C | 0.545 | 0.65913 |
| Choline dehydrogenase, mitochondrial | Q8BJ64 | Chdh | 66 kDa | 12 C | 0.542 | 0.06510 |
| Splicing factor U2AF 65 kDa subunit | P26369 | U2af2 | 54 kDa | 6 C | 0.537 | 0.51851 |
| Interferon-induced 35 kDa protein homolog | Q9D8C4 | Ifi35 | 32 kDa | 4 C | 0.537 | 0.58427 |
| WD repeat domain phosphoinositide-interacting protein 3 | Q9CR39 | Wdr45b | 38 kDa | 15 C | 0.537 | 0.58427 |
| Histidine triad nucleotide-binding protein 2, mitochondrial | Q9D0S9 | Hint2 | 17 kDa | 1 C | 0.533 | 0.45374 |
| UDP-glucose 6-dehydrogenase | O70475 | Ugdh | 55 kDa | 12 C | 0.532 | 0.27251 |
| Formimidoyltransferase-cyclodeaminase | Q91XD4 | Ftcd | 59 kDa | 11 C | 0.530 | 0.66229 |
| Myeloperoxidase | P11247 | Mpo | 81 kDa | 16 C | 0.525 | 0.51386 |
| NADP-dependent malic enzyme | P06801 | Me1 | 64 kDa | 11 C | 0.522 | 0.01529 |
| Septin-2 | P42208 | 2-Sep | 42 kDa | 4 C | 0.522 | 0.10616 |
| Calcium-regulated heat stable protein 1 | Q9CR86 | Carhsp1 | 16 kDa | 4 C | 0.513 | 0.29728 |
| Calcyclin-binding protein | Q9CXW3 | Cacybp | 27 kDa | 2 C | 0.512 | 0.39531 |
| Ig kappa chain V-V region MOPC 149 | P01636 | | 12 kDa | 2 C | 0.512 | 0.39531 |
| Glutaryl-CoA dehydrogenase, mitochondrial | Q60759 | Gcdh | 49 kDa | 9 C | 0.512 | 0.50353 |
| Molybdenum cofactor sulfurase | Q14CH1 | Mocos | 95 kDa | 24 C | 0.500 | 0.219 |
| Ig heavy chain V-III region J606 | P01801 | | 13 kDa | 2 C | 0.500 | 0.26349 |
| Guanine deaminase | Q9R111 | Gda | 51 kDa | 9 C | 0.500 | 0.3098 |
| Hydroxymethylglutaryl-CoA synthase, cytoplasmic | Q8JZK9 | Hmgcs1 | 58 kDa | 11 C | 0.481 | 0.13987 |
| 182 kDa tankyrase-1-binding protein | P58871 | Tnks1bp1 | 182 kDa | 23 C | 0.437 | 0.52469 |
| 2-amino-3-carboxymuconate-6-semialdehyde decarboxylase | Q8R519 | Acmsd | 38 kDa | 7 C | 0.431 | 0.51008 |
| Tyrosine--tRNA ligase, cytoplasmic | Q91WQ3 | Yars | 59 kDa | 7 C | 0.420 | 0.09737 |
| Kynurenine--oxoglutarate transaminase 3 | Q71RI9 (+1) | Kyat3 | 51 kDa | 10 C | 0.412 | 0.45725 |
| Ubiquitin-fold modifier-conjugating enzyme 1 | Q9CR09 | Ufc1 | 19 kDa | 3 C | 0.400 | 0.2446 |
| Beta-2-glycoprotein 1 | Q01339 | Apoh | 39 kDa | 23 C | 0.392 | 0.14820 |
| Bleomycin hydrolase | Q8R016 | Blmh | 53 kDa | 4 C | 0.388 | 0.19094 |
| Beta-glucuronidase | P12265 | Gusb | 74 kDa | 8 C | 0.388 | 0.19094 |
| Phospholipase B-like 1 | Q8VCI0 | Plbd1 | 63 kDa | 7 C | 0.387 | 0.36848 |
| Alanine aminotransferase 2 | Q8BGT5 | Gpt2 | 58 kDa | 14 C | 0.387 | 0.36848 |
| Sulfatase-modifying factor 1 | Q8R0F3 | Sumf1 | 41 kDa | 11 C | 0.375 | 0.54609 |
| Acyl-coenzyme A amino acid N-acyltransferase 1 | A2AKK5 (+1) | Acnat1 | 46 kDa | 7 C | 0.350 | 0.11637 |
| Ethylmalonyl-CoA decarboxylase | Q9D9V3 | Echdc1 | 35 kDa | 6 C | 0.235 | 0.22289 |
| Microtubule-associated protein 4 | P27546 (+1) | Map4 | 117 kDa | 10 C | 0.197 | 0.1818 |
| Ig kappa chain V19-17 | P01633 | Igk-V19-17 | 16 kDa | 2 C | 0.197 | 0.1818 |
| Kinectin | Q61595 (+1) | Ktn1 | 153 kDa | 10 C | 0.136 | 0.37390 |
| Complement component C8 alpha chain | Q8K182 | C8a | 66 kDa | 29 C | 0.136 | 0.37390 |
| Mitochondrial proton/calcium exchanger protein | Q9Z2I0 | Letm1 | 83 kDa | 12 C | 0.136 | 0.37390 |
| Fructose-bisphosphate aldolase C | P05063 | Aldoc | 39 kDa | 7 C | 0.136 | 0.37390 |
| 6-pyruvoyl tetrahydrobiopterin synthase | Q9R1Z7 | Pts | 16 kDa | 1 C | 0.136 | 0.37390 |
| Estradiol 17-beta-dehydrogenase 8 | P50171 (+1) | Hsd17b8 | 27 kDa | 4 C | 0.136 | 0.37390 |
| Molybdopterin synthase catalytic subunit | Q9Z223 | Mocs2 | 21 kDa | 4 C | 0.097 | 0.1641 |
| Complement component C8 beta chain | Q8BH35 | C8b | 66 kDa | 32 C | 0.097 | 0.1641 |
| Pyruvate carboxylase, mitochondrial | Q05920 | Pc | 130 kDa | 13 C | 0.071 | 0.37390 |
| Nucleoprotein TPR | F6ZDS4 | Tpr | 274 kDa | 7 C | 0.059 | 0.1330 |
| Glutamate--cysteine ligase catalytic subunit | P97494 | Gclc | 73 kDa | 14 C | 0.390 | 0.02411 |
| Properdin | P11680 | Cfp | 50 kDa | 44 C | 0.300 | 0.02192 |
| Heterogeneous nuclear ribonucleoprotein U | Q8VEK3 (+1) | Hnrnpu | 88 kDa | 14 C | 0.240 | 0.04760 |
| Adapter molecule crk | Q64010 | Crk | 34 kDa | 1 C | 0.240 | 0.04760 |
