## Supplemental Table 17 for "Dietary restriction transforms the protein sulfhydrome in a tissue-specific and cystathionine γ-lyase-dependent manner"

| Supplemental Table 17: Sulfhydrated Protein Pathway Enrichment in CGL KO Liver | | | |
|---|---|---|---|
| KEGG Pathway (DR) | p-val (adj) | -LOG10(p-val adj) | Involved Gene |
| No KEGG | | | |
| KEGG Pathway (Unchanged) | p-val (adj) | -LOG10(p-val adj) | Involved Gene |
| Vitamin B6 metabolism | 0.0442 | 1.35 | PNPO,PDXK,AOX1,AOX3 |
| Peroxisome | 1.26E-07 | 6.90 | SOD2,ACAA1B,PIPOX,ACOX1,HACL1,EPHX2,SOD1,XDH,HSD17B4,PRDX5,IDH1,AGXT,CAT,HAO1,SCP2,HMGCL,PRDX1,NUDT7,ACAA1A,ECH1 |
| Proteasome | 5.84E-09 | 8.23 | PSMD4,PSMB4,PSMB1,PSMA2,PSMB6,PSMA6,PSMB5,PSME1,PSMB8,PSMB7,PSMA7,PSMB2,PSMA1,PSMA4,PSMA5,PSMB3 |
| Arginine and proline metabolism | 0.00186 | 2.73 | ARG1,CNDP2,GOT1,ALDH9A1,ALDH4A1,ALDH2,OAT,GOT2,LAP3,AGMAT,ALDH7A1 |
| Histidine metabolism | 0.0000753 | 4.12 | FTCD,AMDHD1,HAL,ASPA,CNDP2,ALDH9A1,ALDH2,UROC1,ALDH7A1 |
| Carbon metabolism | 8.76E-21 | 20.06 | DLST,SDHB,PGAM1,MDH2,MDH1,OGDH,SHMT1,DLD,ALDH6A1,SDHA,GLUD1,TKT,ESD,ACO2,GPT,TPI1,ACAT2,MUT,GLDC,GOT1,ECHS1,TALDO1,IDH1,CPS1,AGXT,CAT,HAO1,ALDOB,PRPS1,GOT2,ACAT1,PCCB,TKFC,PKLR,PGP,GAPDH,PGK1,ENO1 |
| 2-Oxocarboxylic acid metabolism | 0.0136 | 1.87 | ACO2,GPT,ACY1,GOT1,IDH1,GOT2 |
| Selenocompound metabolism | 0.00964 | 2.02 | TXNRD1,SCLY,SEPHS1,SEPSECS,KYAT1,SEPHS2 |
| Cysteine and methionine metabolism | 0.0000013 | 5.89 | APIP,MDH2,MDH1,CBS,GOT1,AHCY,GSS,GCLM,GOT2,MAT1A,MAT2B,BHMT2,LDHA,BHMT |
| Butanoate metabolism | 0.00016 | 3.80 | ACAT2,ECHS1,HMGCS2,HADH,BDH2,HMGCL,ACAT1,ALDH5A1,ABAT |
| Propanoate metabolism | 0.000585 | 3.23 | DLD,ALDH6A1,ACAT2,MUT,ECHS1,ACAT1,PCCB,ABAT,LDHA |
| Glycine, serine and threonine metabolism | 0.00000196 | 5.71 | GNMT,PGAM1,CHDH,PIPOX,SHMT1,DLD,CBS,GLDC,AGXT,GRHPR,DMGDH,ALDH7A1,BHMT |
| Pyruvate metabolism | 3.94E-08 | 7.40 | MDH2,MDH1,DLD,ACAT2,GLO1,HAGH,ALDH9A1,ALDH2,LDHD,ACAT1,GRHPR,PKLR,ALDH7A1,LDHA |
| Tryptophan metabolism | 0.00000051 | 6.29 | HAAO,GCDH,OGDH,ACAT2,ECHS1,ALDH9A1,CAT,HADH,ALDH2,ACAT1,KYAT1,ALDH7A1,AOX1,AOX3 |
| Valine, leucine and isoleucine degradation | 4.06E-14 | 13.39 | ACAA1B,DLD,ALDH6A1,AUH,MCCC2,ACAT2,MUT,ECHS1,ALDH9A1,HMGCS2,HADH,HMGCL,ALDH2,HIBADH,ACAT1,PCCB,ACAA1A,ACAA2,ALDH7A1,ABAT,AOX1,AOX3 |
| Drug metabolism - other enzymes | 1.56E-07 | 6.81 | GSTM3,MGST1,MPO,NME2,DPYS,XDH,GSTO1,HPRT1,GSTA3,CMPK1,CES1F,CES2E,DPYD,UPB1,NME1,GSTM2,CES1D,GSTM1,IMPDH2,NME3 |
| Fatty acid degradation | 2.51E-07 | 6.60 | GCDH,ACAA1B,ACOX1,ACAT2,ECI1,ECHS1,ACADL,ALDH9A1,HADH,ALDH2,ACAT1,ACAA1A,ACAA2,ALDH7A1,ADH1 |
| Fatty acid metabolism | 0.0464 | 1.33 | ACAA1B,ACOX1,ACAT2,ECHS1,ACADL,HADH,ACAT1,ACAA1A,ACAA2 |
| Glyoxylate and dicarboxylate metabolism | 3.94E-11 | 10.40 | MDH2,MDH1,SHMT1,DLD,ACO2,ACAT2,MUT,GLDC,AGXT,CAT,HAO1,ACAT1,PCCB,GRHPR,PGP |
| Glutathione metabolism | 0.00318 | 2.50 | GSTM3,MGST1,GSTO1,GSTA3,IDH1,GSS,GCLM,GSR,LAP3,GSTM2,GSTM1,GPX1 |
| Arginine biosynthesis | 1.97E-07 | 6.71 | ARG1,GLUD1,GPT,ACY1,GOT1,ASL,CPS1,OTC,GOT2,ASS1 |
| Alanine, aspartate and glutamate metabolism | 2.27E-07 | 6.64 | ASPA,GLUD1,GPT,NIT2,GOT1,ASL,CPS1,AGXT,ALDH4A1,GOT2,ALDH5A1,ABAT,ASS1 |
| Tyrosine metabolism | 0.00555 | 2.26 | GSTZ1,HGD,GOT1,HPD,FAH,GOT2,AOX1,AOX3,ADH1 |
| Synthesis and degradation of ketone bodies | 0.00686 | 2.16 | ACAT2,HMGCS2,BDH2,HMGCL,ACAT1 |
| Biosynthesis of amino acids | 1.75E-13 | 12.76 | PGAM1,ARG1,SHMT1,TKT,ACO2,GPT,ACY1,TPI1,CBS,GOT1,TALDO1,ASL,IDH1,CPS1,ALDOB,OTC,PRPS1,GOT2,MAT1A,PKLR,MAT2B,GAPDH,PGK1,ENO1,ASS1 |
| beta-Alanine metabolism | 0.000112 | 3.95 | ALDH6A1,DPYS,CNDP2,ECHS1,ALDH9A1,ALDH2,DPYD,UPB1,ALDH7A1,ABAT |
| Glycolysis / Gluconeogenesis | 0.0000237 | 4.63 | PGAM1,DLD,TPI1,ALDH9A1,ALDOB,PGM1,ALDH2,GALM,PKLR,ALDH7A1,GAPDH,PGK1,LDHA,ENO1,ADH1 |
| Lysine degradation | 0.000796 | 3.10 | GCDH,DLST,PIPOX,OGDH,ACAT2,ECHS1,ALDH9A1,HADH,ALDH2,ACAT1,BBOX1,ALDH7A1 |
| Metabolic pathways | 3.31E-31 | 30.48 | HAAO,FTCD,PAFAH1B2,GCDH,DLST,SDHB,ACAA1B,APIP,PGAM1,QDPR,AMDHD1,CHDH,PIPOX,PNPO,MDH2,ETNPPL,ARG1,HAL,PCBD1,MDH1,OGDH,SHMT1,DLD,ASPA,ACOX1,NME2,GSTZ1,PYGL,ALDH6A1,AUH,SDHA,NDUFS6,MCCC2,GLUD1,TKT,EPHX2,CMBL,DPYS,ACO2,GPT,CPOX,HGD,ACY1,TPI1,ACAT2,MUT,CBS,XDH,NDUFV2,HSD17B4,CNDP2,GLDC,NT5C2,PCYT2,GOT1,ECHS1,URAH,TALDO1,ASL,HPRT1,IDH1,NDUFS1,CPS1,ACADL,AGXT,SCLY,BPNT1,SEPHS1,ALDH9A1,PRDX6,SORD,HAO1,IMPA1,AHCY,GSS,HMGCS2,HADH,GCLM,BDH2,UOX,ALDOB,ALAD,SCP2,HMGCL,CMPK1,ALDH4A1,AK2,KHK,PGM1,UGDH,PAICS,HPD,ALDH2,HIBADH,FAH,QPRT,OAT,OTC,PRPS1,GOT2,CES1F,ACAT1,PCCB,PDXK,DPYD,UPB1,TKFC,UROC1,ATP5H,GALM,GRHPR,ALDH5A1,GALT,ACAA1A,ACAA2,GCH1,NME1,MAT1A,NDUFV1,UQCRFS1,ADK,KYAT1,LAP3,BLVRB,AGMAT,PKLR,MAT2B,DMGDH,BHMT2,PGP,ACP1,GPHN,SEPHS2,SUOX,ALDH1A1,ALDH7A1,CES1D,GAPDH,ABAT,GDA,PGK1,IMPDH2,LDHA,ENO1,AOX1,PPCDC,MOCS1,AOX3,PTGES3,GANAB,NME3,ADH1,BHMT,URAD,ASS1 |
| Citrate cycle (TCA cycle) | 0.000782 | 3.11 | DLST,SDHB,MDH2,MDH1,OGDH,DLD,SDHA,ACO2,IDH1 |
| KEGG Pathway (AL) | p-val (adj) | -LOG10(p-val adj) | Involved Gene |
| No KEGG | | | |
