## Supplemental Table 18 for "Dietary restriction transforms the protein sulfhydrome in a tissue-specific and cystathionine γ-lyase-dependent manner"

| Supplemental Table 18: Dietary Impact on the CGL KO Kidney Sulfhydrome | | | | | | |
|---|---|---|---|---|---|---|
| Protein Name | Accession Number | Alternate ID | Molecular Weight | Cysteine Residues | DR/AL Spectral Count Ratio | P-value |
| Ig gamma-2A chain C region, A allele | P01863 (+1) | Ighg | 36 kDa | 10 C | 5.952 | 0.03767 |
| Heterogeneous nuclear ribonucleoprotein M | Q9D0E1 (+1) | Hnrnpm | 78 kDa | 6 C | 5.000 | 0.00595 |
| Phospholipase D3 | O35405 | Pld3 | 54 kDa | 8 C | 4.167 | 0.04761 |
| Ig kappa chain V-V region L7 (Fragment) | P01642 | Gm10881 | 13 kDa | 2 C | 2.857 | 0.01232 |
| UPF0160 protein MYG1, mitochondrial | Q9JK81 | Myg1 | 43 kDa | 7 C | 2.333 | 0.01613 |
| Copper homeostasis protein cutC homolog | Q9D8X1 | Cutc | 29 kDa | 7 C | 10.333 | 0.16419 |
| Corticosteroid-binding globulin | Q06770 | Serpina6 | 45 kDa | 3 C | 10.333 | 0.16419 |
| 28S ribosomal protein S22, mitochondrial | Q9CXW2 | Mrps22 | 41 kDa | 2 C | 7.333 | 0.3739 |
| Isoform 3 of Agrin | A2ASQ1-3 | Agrn | 198 kDa | 2 C | 7.333 | 0.3739 |
| 3-oxoacyl-[acyl-carrier-protein] synthase, mitochondrial | Q9D404 | Oxsm | 49 kDa | 11 C | 7.333 | 0.3739 |
| Cordon-bleu protein-like 1 | Q3UMF0 (+3) | Cobll1 | 137 kDa | 10 C | 5.833 | 0.10658 |
| ADP-sugar pyrophosphatase | Q9JKX6 | Nudt5 | 24 kDa | 5 C | 4.167 | 0.15819 |
| Complement C4-B | P01029 | C4b | 193 kDa | 29 C | 3.381 | 0.23959 |
| Protein-glutamine gamma-glutamyltransferase 2 | P21981 | Tgm2 | 77 kDa | 20 C | 3.381 | 0.23959 |
| Isochorismatase domain-containing protein 1 | Q91V64 | Isoc1 | 32 kDa | 5 C | 3.333 | 0.10588 |
| Serpin B8 | O08800 | Serpinb8 | 42 kDa | 11 C | 2.903 | 0.06902 |
| Heterogeneous nuclear ribonucleoprotein A0 | Q9CX86 | Hnrnpa0 | 31 kDa | 3 C | 2.667 | 0.5461 |
| Proteasome subunit beta type-8 | P28063 | Psmb8 | 30 kDa | 5 C | 2.583 | 0.36848 |
| Ig kappa chain V-V region MOPC 149 | P01636 | | 12 kDa | 2 C | 2.583 | 0.36848 |
| Ubiquitin-conjugating enzyme E2 variant 2 | Q9D2M8 | Ube2v2 | 16 kDa | 1 C | 2.583 | 0.36848 |
| Small nuclear ribonucleoprotein Sm D2 | P62317 | Snrpd2 | 14 kDa | 2 C | 2.583 | 0.36848 |
| Serine/threonine-protein phosphatase CPPED1 | Q8BFS6 | Cpped1 | 35 kDa | 6 C | 2.583 | 0.36848 |
| Ubiquitin-40S ribosomal protein S27a | P62983 | Rps27a | 18 kDa | 6 C | 2.581 | 0.06371 |
| Arsenite methyltransferase | Q91WU5 | As3mt | 42 kDa | 12 C | 2.581 | 0.19094 |
| Isocitrate dehydrogenase [NADP] cytoplasmic | O88844 | Idh1 | 47 kDa | 7 C | 2.393 | 0.23033 |
| Beta-1,3-galactosyl-O-glycosyl-glycoprotein beta-1,6-N-acetylglucosaminyltransferase | Q09324 | Gcnt1 | 50 kDa | 11 C | 2.381 | 0.09737 |
| Periplakin | Q9R269 | Ppl | 204 kDa | 11 C | 2.318 | 0.51008 |
| Complement factor I | Q61129 | Cfi | 67 kDa | 40 C | 2.258 | 0.11288 |
| Peroxidasin homolog | Q3UQ28 | Pxdn | 165 kDa | 46 C | 2.200 | 0.06661 |
| Tropomyosin alpha-4 chain | Q6IRU2 | Tpm4 | 28 kDa | 2 C | 2.195 | 0.3792 |
| ADP-ribosylation factor-like protein 3 | Q9WUL7 | Arl3 | 20 kDa | 3 C | 1.935 | 0.15293 |
| Growth arrest-specific protein 2 | P11862 | Gas2 | 35 kDa | 11 C | 1.905 | 0.23073 |
| Glycerol-3-phosphate phosphatase | Q8CHP8 | Pgp | 35 kDa | 8 C | 1.864 | 0.58428 |
| Septin-6 | Q9R1T4 (+2) | 6-Sep | 50 kDa | 8 C | 1.864 | 0.58428 |
| Ig heavy chain V region 441 | P01806 (+2) | | 13 kDa | 3 C | 1.831 | 0.3708 |
| Nucleolysin TIAR | P70318 | Tial1 | 43 kDa | 6 C | 1.765 | 0.27466 |
| Dynein light chain 1, cytoplasmic | P63168 | Dynll1 | 10 kDa | 3 C | 1.750 | 0.25082 |
| Glutathione S-transferase A1 | P13745 | Gsta1 | 26 kDa | 2 C | 1.727 | 0.12434 |
| 2-amino-3-carboxymuconate-6-semialdehyde decarboxylase | Q8R519 | Acmsd | 38 kDa | 7 C | 1.714 | 0.01944 |
| Phosphopantothenoylcysteine decarboxylase | Q8BZB2 | Ppcdc | 22 kDa | 6 C | 1.707 | 0.42361 |
| Rab GDP dissociation inhibitor beta | Q61598 | Gdi2 | 51 kDa | 9 C | 1.667 | 0.5844 |
| Glutathione peroxidase 3 | P46412 | Gpx3 | 25 kDa | 3 C | 1.667 | 0.44182 |
| Aldose reductase | P45376 | Akr1b1 | 36 kDa | 6 C | 1.645 | 0.63521 |
| Serine/threonine-protein phosphatase 6 catalytic subunit | Q9CQR6 | Ppp6c | 35 kDa | 12 C | 1.613 | 0.50393 |
| Ribokinase | Q8R1Q9 | Rbks | 34 kDa | 9 C | 1.613 | 0.50393 |
| m7GpppX diphosphatase | Q9DAR7 | Dcps | 39 kDa | 2 C | 1.613 | 0.50393 |
| Isoform 1 of Harmonin | Q9ES64-3 | Ush1c | 68 kDa | 7 C | 1.600 | 0.34864 |
| Quinone oxidoreductase-like protein 1 | Q921W4 | Cryzl1 | 39 kDa | 6 C | 1.600 | 0.34864 |
| Succinate--CoA ligase [ADP-forming] subunit beta, mitochondrial | Q9Z2I9 | Sucla2 | 50 kDa | 6 C | 1.594 | 0.64857 |
| Lysosomal protective protein | P16675 | Ctsa | 54 kDa | 11 C | 1.558 | 0.67847 |
| Tubulin beta-4B chain | P68372 | Tubb4b | 50 kDa | 8 C | 1.520 | 0.55161 |
| Myosin light chain kinase, smooth muscle | Q6PDN3 | Mylk | 213 kDa | 46 C | 1.500 | 0.11612 |
| NEDD8-conjugating enzyme Ubc12 | P61082 | Ube2m | 21 kDa | 5 C | 1.500 | 0.1583 |
| Complement component C8 beta chain | Q8BH35 | C8b | 66 kDa | 32 C | 1.500 | 0.28786 |
| Serine/arginine-rich splicing factor 1 | Q6PDM2 | Srsf1 | 28 kDa | 2 C | 1.476 | 0.62227 |
| Ig kappa chain V-V region MOPC 41 | P01639 | Gm5571 | 14 kDa | 3 C | 1.463 | 0.62097 |

| | | | | | | |
|---|---|---|---|---|---|---|
| Thioredoxin domain-containing protein 5 | Q91W90 | Txndc5 | 46 kDa | 12 C | 1.462 | 0.05504 |
| Inter alpha-trypsin inhibitor, heavy chain 4 | A6X935 (+1) | Itih4 | 105 kDa | 3 C | 1.455 | 0.78734 |
| Valacyclovir hydrolase | Q8R164 | Bphl | 33 kDa | 4 C | 1.452 | 0.67247 |
| Complement C3 | P01027 | C3 | 186 kDa | 27 C | 1.419 | 0.12158 |
| Glycerol kinase | Q64516 (+2) | Gk | 61 kDa | 15 C | 1.409 | 0.73844 |
| NADH dehydrogenase [ubiquinone] 1 alpha subcomplex subunit 10, mitochondrial | Q99LC3 | Ndufa10 | 41 kDa | 5 C | 1.409 | 0.73844 |
| Protein phosphatase 1 regulatory subunit 7 | Q3UM45 | Ppp1r7 | 41 kDa | 2 C | 1.393 | 0.10617 |
| Fumarylacetoacetate hydrolase domain-containing protein 2A | Q3TC72 | Fahd2 | 35 kDa | 6 C | 1.391 | 0.21807 |
| Fetuin-B | Q9QXC1 | Fetub | 43 kDa | 15 C | 1.379 | 0.04239 |
| Ribosyldihydronicotinamide dehydrogenase [quinone] | Q9JI75 | Nqo2 | 26 kDa | 4 C | 1.367 | 0.59366 |
| 55 kDa erythrocyte membrane protein | P70290 | Mpp1 | 52 kDa | 5 C | 1.367 | 0.59366 |
| Nucleoporin SEH1 | Q8R2U0 (+1) | Seh1l | 40 kDa | 10 C | 1.367 | 0.69079 |
| Ataxin-2 | O70305 | Atxn2 | 136 kDa | 15 C | 1.364 | 0.69536 |
| Dynein light chain 2, cytoplasmic | Q9D0M5 | Dynll2 | 10 kDa | 2 C | 1.333 | 0.1583 |
| Astrocytic phosphoprotein PEA-15 | Q62048 (+1) | Pea15 | 15 kDa | 1 C | 1.333 | 0.3739 |
| Ig gamma-1 chain C region, membrane-bound form | P01869 | Ighg1 | 43 kDa | 12 C | 1.333 | 0.64333 |
| Annexin A2 | P07356 | Anxa2 | 39 kDa | 5 C | 1.320 | 0.06468 |
| Ig kappa chain V-III region PC 7043 | P01665 (+1) | | 12 kDa | 2 C | 1.308 | 0.55179 |
| Carbonic anhydrase 2 | P00920 | Ca2 | 29 kDa | 2 C | 1.300 | 0.19225 |
| Ig alpha chain C region | P01878 | | 37 kDa | 13 C | 1.300 | 0.65196 |
| Tyrosine--tRNA ligase, cytoplasmic | Q91WQ3 | Yars | 59 kDa | 7 C | 1.290 | 0.66463 |
| Ran-binding protein 3 | Q9CT10 | Ranbp3 | 53 kDa | 5 C | 1.290 | 0.66463 |
| Inositol monophosphatase 1 | O55023 | Impa1 | 30 kDa | 6 C | 1.286 | 0.27458 |
| Pyruvate dehydrogenase E1 component subunit beta, mitochondrial | Q9D051 | Pdhb | 39 kDa | 6 C | 1.286 | 0.56144 |
| Very long-chain acyl-CoA synthetase | O35488 | Slc27a2 | 70 kDa | 14 C | 1.286 | 0.71359 |
| Aldo-keto reductase family 1 member C21 | Q91WR5 | Akr1c21 | 37 kDa | 9 C | 1.281 | 0.82776 |
| Fatty acid-binding protein, adipocyte | P04117 | Fabp4 | 15 kDa | 2 C | 1.267 | 0.11612 |
| Bleomycin hydrolase | Q8R016 | Blmh | 53 kDa | 4 C | 1.267 | 0.44182 |
| F-box only protein 50 | G3X9C2 | Nccrp1 | 30 kDa | 3 C | 1.261 | 0.10119 |
| Glucosamine-6-phosphate isomerase 1 | O88958 | Gnpda1 | 33 kDa | 3 C | 1.250 | 0.14815 |
| Aldehyde dehydrogenase, cytosolic 1 | O35945 | Aldh1a7 | 55 kDa | 8 C | 1.250 | 0.1583 |
| Plasminogen | P20918 | Plg | 91 kDa | 48 C | 1.250 | 0.51852 |
| Chloride intracellular channel protein 1 | Q9Z1Q5 | Clic1 | 27 kDa | 6 C | 1.250 | 0.51852 |
| COP9 signalosome complex subunit 3 | O88543 | Cops3 | 48 kDa | 10 C | 1.250 | 0.51852 |
| Kinesin-1 heavy chain | Q61768 | Kif5b | 110 kDa | 13 C | 1.250 | 0.51852 |
| Signal transducing adapter molecule 1 | P70297 | Stam | 60 kDa | 7 C | 1.250 | 0.51852 |
| Osteoclast-stimulating factor 1 | Q62422 | Ostf1 | 24 kDa | 4 C | 1.250 | 0.65299 |
| Immunoglobulin J chain | P01592 | Jchain | 18 kDa | 8 C | 1.250 | 0.74152 |
| Isoform 4 of Thioredoxin reductase 2, mitochondrial | Q9JLT4-4 | Txnrd2 | 53 kDa | 11 C | 1.235 | 0.27458 |
| Tubulin alpha-1B chain | P05213 | Tuba1b | 50 kDa | 12 C | 1.235 | 0.74056 |
| Ig heavy chain V regions TEPC 15/S107/HPCM1/HPCM2/HPCM3 | P01787 (+3) | | 14 kDa | 2 C | 1.231 | 0.62396 |
| High mobility group protein B1 | P63158 | Hmgb1 | 25 kDa | 3 C | 1.226 | 0.42423 |
| Glutaredoxin-related protein 5, mitochondrial | Q80Y14 | Glrx5 | 16 kDa | 2 C | 1.222 | 0.3739 |
| Basement membrane-specific heparan sulfate proteoglycan core protein | Q05793 | Hspg2 | 398 kDa | 188 C | 1.222 | 0.7096 |
| Splicing factor U2AF 65 kDa subunit | P26369 | U2af2 | 54 kDa | 6 C | 1.220 | 0.76059 |
| Heterogeneous nuclear ribonucleoprotein U | Q8VEK3 (+1) | Hnrnpu | 88 kDa | 14 C | 1.200 | 0.3739 |
| Exosome complex component MTR3 | Q8BTW3 | Exosc6 | 28 kDa | 6 C | 1.200 | 0.64333 |
| Succinate--CoA ligase [ADP/GDP-forming] subunit alpha, mitochondrial | Q9WUM5 | Suclg1 | 36 kDa | 6 C | 1.200 | 0.72466 |
| Glycine N-acyltransferase | Q91XE0 | Glyat | 34 kDa | 5 C | 1.190 | 0.86405 |
| DnaJ homolog subfamily B member 11 | Q99KV1 | Dnajb11 | 41 kDa | 5 C | 1.188 | 0.68427 |
| Proteasome subunit beta type-4 | P99026 | Psmb4 | 29 kDa | 2 C | 1.182 | 0.2302 |
| Serine beta-lactamase-like protein LACTB, mitochondrial | Q9EP89 | Lactb | 61 kDa | 4 C | 1.182 | 0.74152 |
| Methylthioribulose-1-phosphate dehydratase | Q9WVQ5 | Apip | 27 kDa | 11 C | 1.176 | 0.34864 |
| Heterogeneous nuclear ribonucleoprotein A/B | Q99020 | Hnrnpab | 31 kDa | 2 C | 1.167 | 0.27458 |
| Complement C1q tumor necrosis factor-related protein 3 | Q9ES30 | C1qtnf3 | 27 kDa | 5 C | 1.167 | 0.72466 |
| Polyribonucleotide nucleotidyltransferase 1, mitochondrial | Q8K1R3 | Pnpt1 | 86 kDa | 16 C | 1.167 | 0.8149 |
| Glutathione peroxidase 1 | P11352 | Gpx1 | 22 kDa | 4 C | 1.162 | 0.54304 |
| Kininogen-1 | O08677 (+1) | Kng1 | 73 kDa | 19 C | 1.156 | 0.32616 |
| Peroxiredoxin-6 | O08709 | Prdx6 | 25 kDa | 2 C | 1.152 | 0.21817 |
| Secernin-2 | Q8VCA8 | Scrn2 | 47 kDa | 10 C | 1.150 | 0.50716 |

| | | | | | | |
|---|---|---|---|---|---|---|
| GDP-mannose 4,6 dehydratase | Q8K0C9 | Gmds | 42 kDa | 6 C | 1.143 | 0.28786 |
| Ficolin-1 | O70165 | Fcn1 | 36 kDa | 10 C | 1.143 | 0.74152 |
| Splicing factor, proline- and glutamine-rich | Q8VIJ6 | Sfpq | 75 kDa | 7 C | 1.133 | 0.7588 |
| Actin-related protein 2/3 complex subunit 3 | Q9JM76 | Arpc3 | 21 kDa | 4 C | 1.125 | 0.3739 |
| Proteasome subunit alpha type-6 | Q9QUM9 | Psma6 | 27 kDa | 8 C | 1.125 | 0.43533 |
| Inhibitor of carbonic anhydrase | Q9DBD0 | Ica | 77 kDa | 35 C | 1.125 | 0.72466 |
| Proteasome subunit beta type-1 | O09061 | Psmb1 | 26 kDa | 5 C | 1.120 | 0.53908 |
| Acyl-coenzyme A amino acid N-acyltransferase 1 | A2AKK5 (+1) | Acnat1 | 46 kDa | 7 C | 1.111 | 0.3739 |
| 3'(2'),5'-bisphosphate nucleotidase 1 | Q9Z0S1 | Bpnt1 | 33 kDa | 6 C | 1.111 | 0.60865 |
| Heterogeneous nuclear ribonucleoprotein L | Q8R081 | Hnrnpl | 64 kDa | 11 C | 1.111 | 0.60865 |
| S-adenosylmethionine synthase isoform type-2 | Q3THS6 | Mat2a | 44 kDa | 6 C | 1.111 | 0.72466 |
| Dihydrolipoyllysine-residue acetyltransferase component of pyruvate dehydrogenase com | Q8BMF4 | Dlat | 68 kDa | 10 C | 1.111 | 0.72466 |
| Nascent polypeptide-associated complex subunit alpha, muscle-specific form | P70670 | Naca | 220 kDa | 16 C | 1.111 | 0.8149 |
| Glutathione S-transferase Mu 1 | P10649 | Gstm1 | 26 kDa | 2 C | 1.105 | 0.24636 |
| 14-3-3 protein beta/alpha | Q9CQV8 | Ywhab | 28 kDa | 2 C | 1.103 | 0.44182 |
| Isoform Crk-I of Adapter molecule crk | Q64010-2 | Crk | 23 kDa | 1 C | 1.100 | 0.67787 |
| Thioredoxin reductase 3 | Q99MD6 | Txnrd3 | 71 kDa | 18 C | 1.100 | 0.84501 |
| Serpin B6 | Q60854 | Serpinb6 | 43 kDa | 6 C | 1.099 | 0.89987 |
| Actin-related protein 2/3 complex subunit 4 | P59999 | Arpc4 | 20 kDa | 4 C | 1.091 | 0.64333 |
| Tetratricopeptide repeat protein 38 | A3KMP2 | Ttc38 | 52 kDa | 9 C | 1.091 | 0.72466 |
| Septin-8 | Q8CHH9 | 8-Sep | 50 kDa | 6 C | 1.091 | 0.79526 |
| Histidine-rich glycoprotein | Q9ESB3 | Hrg | 59 kDa | 17 C | 1.091 | 0.8149 |
| Plastin-3 | Q99K51 | Pls3 | 71 kDa | 9 C | 1.091 | 0.87216 |
| Hemoglobin subunit beta-1 | P02088 | Hbb-b1 | 16 kDa | 2 C | 1.083 | 0.72 |
| Trifunctional enzyme subunit alpha, mitochondrial | Q8BMS1 | Hadha | 83 kDa | 12 C | 1.081 | 0.91304 |
| Haptoglobin | Q61646 | Hp | 39 kDa | 9 C | 1.080 | 0.72466 |
| Aldehyde dehydrogenase, mitochondrial | P47738 | Aldh2 | 57 kDa | 9 C | 1.079 | 0.75589 |
| Glutamate--cysteine ligase catalytic subunit | P97494 | Gclc | 73 kDa | 14 C | 1.075 | 0.74487 |
| Polymerase delta-interacting protein 2 | Q91VA6 | Poldip2 | 42 kDa | 4 C | 1.071 | 0.64333 |
| Glutamate--cysteine ligase regulatory subunit | O09172 | Gclm | 31 kDa | 6 C | 1.071 | 0.8149 |
| Isocitrate dehydrogenase [NADP], mitochondrial | P54071 | Idh2 | 51 kDa | 8 C | 1.069 | 0.82404 |
| Sodium/potassium-transporting ATPase subunit alpha-1 | Q8VDN2 | Atp1a1 | 113 kDa | 23 C | 1.069 | 0.89277 |
| Septin-11 | Q8C1B7 (+2) | 11-Sep | 50 kDa | 6 C | 1.067 | 0.74887 |
| Cytochrome c, somatic | P62897 | Cycs | 12 kDa | 2 C | 1.067 | 0.78928 |
| Glutamine synthetase | P15105 | Glul | 42 kDa | 13 C | 1.053 | 0.72918 |
| Cytochrome b-c1 complex subunit 1, mitochondrial | Q9CZ13 | Uqcrc1 | 53 kDa | 11 C | 1.050 | 0.92516 |
| Septin-2 | P42208 | 2-Sep | 42 kDa | 4 C | 1.048 | 0.64333 |
| 39S ribosomal protein L39, mitochondrial | Q9JKF7 | Mrpl39 | 39 kDa | 7 C | 1.048 | 0.96434 |
| Golgin subfamily A member 2 | Q921M4 (+1) | Golga2 | 113 kDa | 7 C | 1.048 | 0.96434 |
| Aldo-keto reductase family 1 member C13 | Q8VC28 | Akr1c13 | 37 kDa | 10 C | 1.048 | 0.96434 |
| DnaJ homolog subfamily C member 2 | P54103 | Dnajc2 | 72 kDa | 8 C | 1.048 | 0.96434 |
| Disabled homolog 2 | P98078 (+1) | Dab2 | 82 kDa | 3 C | 1.045 | 0.88747 |
| Complement factor H | P06909 | Cfh | 139 kDa | 82 C | 1.045 | 0.89437 |
| Isoleucine--tRNA ligase, mitochondrial | Q8BIJ6 | Iars2 | 113 kDa | 20 C | 1.043 | 0.85931 |
| Aspartate aminotransferase, cytoplasmic | P05201 | Got1 | 46 kDa | 5 C | 1.036 | 0.82029 |
| Septin-7 | O55131 | 7-Sep | 51 kDa | 6 C | 1.036 | 0.86172 |
| Vesicle-associated membrane protein-associated protein B | Q9QY76 | Vapb | 27 kDa | 4 C | 1.033 | 0.95448 |
| Thioredoxin domain-containing protein 12 | Q9CQU0 | Txndc12 | 19 kDa | 3 C | 1.033 | 0.95448 |
| Proteasome subunit beta type-5 | O55234 | Psmb5 | 29 kDa | 3 C | 1.031 | 0.8298 |
| Fructose-bisphosphate aldolase A | P05064 | Aldoa | 39 kDa | 8 C | 1.030 | 0.67452 |
| UMP-CMP kinase | Q9DBP5 | Cmpk1 | 22 kDa | 6 C | 1.029 | 0.76764 |
| Ig mu chain C region | P01872 | Ighm | 50 kDa | 19 C | 1.027 | 0.92346 |
| Protein SGT1 homolog | Q9CX34 | Sugt1 | 38 kDa | 5 C | 1.025 | 0.97282 |
| NFU1 iron-sulfur cluster scaffold homolog, mitochondrial | Q9QZ23 | Nfu1 | 29 kDa | 5 C | 1.025 | 0.97282 |
| Phosphotriesterase-related protein | Q60866 | Pter | 39 kDa | 6 C | 1.025 | 0.82143 |
| Glyoxalase domain-containing protein 4 | Q9CPV4 | Glod4 | 33 kDa | 5 C | 1.014 | 0.97945 |
| Aconitate hydratase, mitochondrial | Q99KI0 | Aco2 | 85 kDa | 13 C | 1.014 | 0.70108 |
| Actin, alpha cardiac muscle 1 | P68033 | Actc1 | 42 kDa | 6 C | 1.013 | 0.86827 |
| Serum albumin | P07724 | Alb | 69 kDa | 36 C | 1.008 | 0.92625 |
| Plastin-2 | Q61233 | Lcp1 | 70 kDa | 11 C | 1.000 | 1 |

| | | | | | | |
|---|---|---|---|---|---|---|
| Serine hydroxymethyltransferase, cytosolic | P50431 | Shmt1 | 53 kDa | 10 C | 1.000 | 1 |
| Alpha-actinin-4 | P57780 | Actn4 | 105 kDa | 8 C | 1.000 | 1 |
| Adseverin | Q60604 | Scin | 80 kDa | 4 C | 1.000 | 1 |
| Pregnancy zone protein | Q61838 | Pzp | 166 kDa | 24 C | 1.000 | 1 |
| CD5 antigen-like | Q9QWK4 | Cd5l | 39 kDa | 26 C | 1.000 | 1 |
| Selenide, water dikinase 2 | P97364 | Sephs2 | 48 kDa | 7 C | 1.000 | 1 |
| Adenylate kinase 2, mitochondrial | Q9WTP6 (+1) | Ak2 | 26 kDa | 5 C | 1.000 | 1 |
| Serine/threonine-protein phosphatase PP1-alpha catalytic subunit | P62137 | Ppp1ca | 38 kDa | 13 C | 1.000 | 1 |
| Choline dehydrogenase, mitochondrial | Q8BJ64 | Chdh | 66 kDa | 12 C | 1.000 | 1 |
| Ribonuclease inhibitor | Q91VI7 | Rnh1 | 50 kDa | 30 C | 1.000 | 1 |
| Adenylate kinase isoenzyme 1 | Q9R0Y5 (+1) | Ak1 | 22 kDa | 2 C | 1.000 | 1 |
| Extracellular superoxide dismutase [Cu-Zn] | O09164 | Sod3 | 27 kDa | 6 C | 1.000 | 1 |
| Acylpyruvase FAHD1, mitochondrial | Q8R0F8 | Fahd1 | 25 kDa | 6 C | 1.000 | 1 |
| Prolow-density lipoprotein receptor-related protein 1 | Q91ZX7 | Lrp1 | 505 kDa | 332 C | 1.000 | 1 |
| Isoform 2 of Disks large homolog 1 | Q811D0-2 | Dlg1 | 100 kDa | 7 C | 1.000 | 1 |
| Far upstream element-binding protein 2 | Q3U0V1 | Khsrp | 77 kDa | 8 C | 1.000 | 1 |
| Protein NDRG1 | Q62433 | Ndrg1 | 43 kDa | 8 C | 1.000 | 1 |
| 14-3-3 protein eta | P68510 | Ywhah | 28 kDa | 3 C | 1.000 | 1 |
| Four and a half LIM domains protein 1 | P97447 | Fhl1 | 32 kDa | 34 C | 1.000 | 1 |
| Cofilin-2 | P45591 | Cfl2 | 19 kDa | 2 C | 1.000 | 1 |
| Proteasome activator complex subunit 1 | P97371 | Psme1 | 29 kDa | 3 C | 1.000 | 1 |
| Hepatoma-derived growth factor | P51859 | Hdgf | 26 kDa | 2 C | 1.000 | 1 |
| Carbonic anhydrase 5B, mitochondrial | Q9QZA0 | Ca5b | 37 kDa | 7 C | 1.000 | 1 |
| Eukaryotic translation initiation factor 3 subunit I | Q9QZD9 | Eif3i | 36 kDa | 7 C | 1.000 | 1 |
| Palladin | Q9ET54 (+2) | Palld | 152 kDa | 17 C | 1.000 | 1 |
| Fatty acid-binding protein, epidermal | Q05816 | Fabp5 | 15 kDa | 6 C | 1.000 | 1 |
| Murinoglobulin-1 | P28665 | Mug1 | 165 kDa | 25 C | 1.000 | 1 |
| Polypyrimidine tract-binding protein 1 | P17225 | Ptbp1 | 56 kDa | 3 C | 1.000 | 1 |
| ATP synthase subunit O, mitochondrial | Q9DB20 | Atp5o | 23 kDa | 1 C | 1.000 | 1 |
| Eukaryotic translation initiation factor 2 subunit 1 | Q6ZWX6 | Eif2s1 | 36 kDa | 5 C | 1.000 | 1 |
| Ig heavy chain V region 3 | P01749 | Ighv1-61 | 13 kDa | 3 C | 1.000 | 1 |
| Acylphosphatase-2 | P56375 | Acyp2 | 12 kDa | 1 C | 1.000 | 1 |
| Isoform 1 of Afadin | Q9QZQ1-2 | Afdn | 205 kDa | 11 C | 1.000 | 1 |
| Heme-binding protein 1 | Q9R257 | Hebp1 | 21 kDa | 2 C | 1.000 | 1 |
| Arf-GAP with GTPase, ANK repeat and PH domain-containing protein 1 | Q8BXK8 | Agap1 | 94 kDa | 17 C | 1.000 | 1 |
| Cadherin-1 | P09803 | Cdh1 | 98 kDa | 8 C | 1.000 | 1 |
| Molybdenum cofactor biosynthesis protein 1 | Q5RKZ7 | Mocs1 | 70 kDa | 15 C | 1.000 | 1 |
| Platelet-activating factor acetylhydrolase IB subunit beta | Q61206 | Pafah1b2 | 26 kDa | 3 C | 1.000 | 1 |
| Legumain | O89017 | Lgmn | 49 kDa | 7 C | 1.000 | 1 |
| Ig heavy chain V region 345 | P18526 | | 13 kDa | 3 C | 1.000 | 1 |
| Insulin-like growth factor-binding protein 7 | Q61581 | Igfbp7 | 29 kDa | 18 C | 1.000 | 1 |
| Protein DEK | Q7TNV0 | Dek | 43 kDa | 4 C | 1.000 | 1 |
| Isoform 2 of Acyl-protein thioesterase 1 | P97823-2 | Lypla1 | 23 kDa | 6 C | 1.000 | 1 |
| SH3 domain-binding glutamic acid-rich-like protein | Q9JJU8 | Sh3bgrl | 13 kDa | 2 C | 1.000 | 1 |
| Fatty acid-binding protein, brain | P51880 | Fabp7 | 15 kDa | 5 C | 1.000 | 1 |
| Elongation factor 1-alpha 1 | P10126 | Eef1a1 | 50 kDa | 6 C | 1.000 | 1 |
| Peroxiredoxin-2 | Q61171 | Prdx2 | 22 kDa | 3 C | 1.000 | 1 |
| Electron transfer flavoprotein subunit beta | Q9DCW4 | Etfb | 28 kDa | 4 C | 1.000 | 1 |
| Adenylyl cyclase-associated protein 1 | P40124 | Cap1 | 52 kDa | 6 C | 1.000 | 1 |
| Cleavage and polyadenylation specificity factor subunit 5 | Q9CQF3 | Nudt21 | 26 kDa | 1 C | 0.988 | 0.98409 |
| Sarcosine dehydrogenase, mitochondrial | Q99LB7 | Sardh | 102 kDa | 18 C | 0.988 | 0.98897 |
| Aldehyde dehydrogenase family 8 member A1 | Q8BH00 | Aldh8a1 | 54 kDa | 13 C | 0.986 | 0.98509 |
| Glutaryl-CoA dehydrogenase, mitochondrial | Q60759 | Gcdh | 49 kDa | 9 C | 0.986 | 0.9872 |
| Insulin-degrading enzyme | Q9JHR7 | Ide | 118 kDa | 13 C | 0.978 | 0.90187 |
| Isoform 2 of Tropomyosin alpha-3 chain | P21107-2 | Tpm3 | 29 kDa | 1 C | 0.978 | 0.93844 |
| Alpha-actinin-1 | Q7TPR4 | Actn1 | 103 kDa | 11 C | 0.977 | 0.9043 |
| Chromobox protein homolog 3 | P23198 | Cbx3 | 21 kDa | 3 C | 0.976 | 0.96508 |
| Glutathione synthetase | P51855 | Gss | 52 kDa | 4 C | 0.973 | 0.67787 |
| Aminoacylase-1 | Q99JW2 | Acy1 | 46 kDa | 4 C | 0.973 | 0.81202 |
| D-beta-hydroxybutyrate dehydrogenase, mitochondrial | Q80XN0 | Bdh1 | 38 kDa | 6 C | 0.969 | 0.97751 |

| | | | | | | |
|---|---|---|---|---|---|---|
| Mitochondrial dicarboxylate carrier | Q9QZD8 | Slc25a10 | 32 kDa | 8 C | 0.969 | 0.97751 |
| Xylulose kinase | Q3TNA1 | Xylb | 60 kDa | 14 C | 0.969 | 0.97751 |
| Sodium/glucose cotransporter 2 | Q923I7 | Slc5a2 | 73 kDa | 17 C | 0.969 | 0.97751 |
| Phenylalanine-4-hydroxylase | P16331 | Pah | 52 kDa | 9 C | 0.968 | 0.97561 |
| Heterogeneous nuclear ribonucleoprotein F | Q9Z2X1 | Hnrnpf | 46 kDa | 6 C | 0.967 | 0.88203 |
| Thioredoxin-like protein 1 | Q8CDN6 | Txnl1 | 32 kDa | 7 C | 0.964 | 0.76764 |
| Retinal dehydrogenase 1 | P24549 | Aldh1a1 | 54 kDa | 11 C | 0.962 | 0.60865 |
| Flavin reductase (NADPH) | Q923D2 | Blvrb | 22 kDa | 2 C | 0.962 | 0.83402 |
| 14-3-3 protein zeta/delta | P63101 | Ywhaz | 28 kDa | 3 C | 0.961 | 0.59698 |
| Methylmalonate-semialdehyde dehydrogenase [acylating], mitochondrial | Q9EQ20 | Aldh6a1 | 58 kDa | 8 C | 0.961 | 0.65799 |
| Methylmalonyl-CoA mutase, mitochondrial | P16332 | Mut | 83 kDa | 8 C | 0.960 | 0.8149 |
| Spectrin beta chain, non-erythrocytic 1 | Q62261 | Sptbn1 | 274 kDa | 15 C | 0.957 | 0.67787 |
| Heterogeneous nuclear ribonucleoprotein A3 | Q8BG05 | Hnrnpa3 | 40 kDa | 4 C | 0.956 | 0.82282 |
| Transthyretin | P07309 | Ttr | 16 kDa | 2 C | 0.955 | 0.72466 |
| Eukaryotic translation initiation factor 5 | P59325 | Eif5 | 49 kDa | 8 C | 0.955 | 0.96434 |
| Eukaryotic initiation factor 4A-II | P10630 | Eif4a2 | 46 kDa | 4 C | 0.955 | 0.96434 |
| Peroxiredoxin-1 | P35700 | Prdx1 | 22 kDa | 4 C | 0.954 | 0.33908 |
| GMP reductase 2 | Q99L27 | Gmpr2 | 38 kDa | 9 C | 0.952 | 0.8298 |
| Fructose-bisphosphate aldolase B | Q91Y97 | Aldob | 40 kDa | 8 C | 0.951 | 0.4382 |
| Peroxiredoxin-4 | O08807 | Prdx4 | 31 kDa | 4 C | 0.950 | 0.48182 |
| Actin-related protein 2 | P61161 | Actr2 | 45 kDa | 5 C | 0.949 | 0.85338 |
| Selenide, water dikinase 1 | Q8BH69 | Sephs1 | 43 kDa | 9 C | 0.949 | 0.85931 |
| Fructose-1,6-bisphosphatase isozyme 2 | P70695 | Fbp2 | 37 kDa | 5 C | 0.947 | 0.69176 |
| Dihydropyrimidinase | Q9EQF5 | Dpys | 57 kDa | 9 C | 0.947 | 0.76764 |
| Complement component C8 gamma chain | Q8VCG4 | C8g | 23 kDa | 3 C | 0.947 | 0.8416 |
| 14-3-3 protein theta | P68254 | Ywhaq | 28 kDa | 5 C | 0.946 | 0.86977 |
| Cytosolic non-specific dipeptidase | Q9D1A2 | Cndp2 | 53 kDa | 8 C | 0.945 | 0.50231 |
| Proteasome subunit alpha type-2 | P49722 | Psma2 | 26 kDa | 2 C | 0.944 | 0.85124 |
| Tubulin alpha-4A chain | P68368 | Tuba4a | 50 kDa | 13 C | 0.944 | 0.93276 |
| Fructose-1,6-bisphosphatase 1 | Q9QXD6 | Fbp1 | 37 kDa | 7 C | 0.941 | 0.54128 |
| Serotransferrin | Q921I1 | Tf | 77 kDa | 38 C | 0.939 | 0.05888 |
| Isoform 2 of Cytosol aminopeptidase | Q9CPY7-2 (+1 | Lap3 | 53 kDa | 7 C | 0.939 | 0.36851 |
| Isoform Gamma-2 of Serine/threonine-protein phosphatase PP1-gamma catalytic subunit | P63087-2 | Ppp1cc | 39 kDa | 13 C | 0.938 | 0.60865 |
| 14-3-3 protein gamma | P61982 | Ywhag | 28 kDa | 3 C | 0.938 | 0.75118 |
| Aspartoacylase | Q8R3P0 | Aspa | 35 kDa | 8 C | 0.935 | 0.71489 |
| Isoform 2 of Meprin A subunit beta | Q61847-2 | Mep1b | 80 kDa | 11 C | 0.935 | 0.3739 |
| Heterogeneous nuclear ribonucleoprotein D0 | Q60668 (+1) | Hnrnpd | 38 kDa | 3 C | 0.933 | 0.60865 |
| Succinate-semialdehyde dehydrogenase, mitochondrial | Q8BWF0 | Aldh5a1 | 56 kDa | 10 C | 0.932 | 0.39313 |
| Endoribonuclease LACTB2 | Q99KR3 | Lactb2 | 33 kDa | 5 C | 0.932 | 0.56873 |
| Superoxide dismutase [Mn], mitochondrial | P09671 | Sod2 | 25 kDa | 4 C | 0.931 | 0.56144 |
| Na(+)/H(+) exchange regulatory cofactor NHE-RF1 | P70441 | Slc9a3r1 | 39 kDa | 5 C | 0.930 | 0.74887 |
| Electron transfer flavoprotein subunit alpha, mitochondrial | Q99LC5 | Etfa | 35 kDa | 6 C | 0.926 | 0.78928 |
| Glycine amidinotransferase, mitochondrial | Q9D964 | Gatm | 48 kDa | 9 C | 0.923 | 0.72466 |
| Drebrin-like protein | Q62418 | Dbnl | 49 kDa | 5 C | 0.923 | 0.76764 |
| Apoptosis-inducing factor 1, mitochondrial | Q9Z0X1 | Aifm1 | 67 kDa | 4 C | 0.921 | 0.74965 |
| Actin-related protein 2/3 complex subunit 1A | Q9R0Q6 | Arpc1a | 42 kDa | 10 C | 0.920 | 0.65299 |
| Mitochondrial peptide methionine sulfoxide reductase | Q9D6Y7 (+1) | Msra | 26 kDa | 4 C | 0.917 | 0.27944 |
| Selenium-binding protein 2 | Q63836 | Selenbp2 | 53 kDa | 10 C | 0.917 | 0.41703 |
| 4-trimethylaminobutyraldehyde dehydrogenase | Q9JLJ2 | Aldh9a1 | 54 kDa | 17 C | 0.917 | 0.5019 |
| Fructose-bisphosphate aldolase C | P05063 | Aldoc | 39 kDa | 7 C | 0.917 | 0.72466 |
| Actin-like protein 6A | Q9Z2N8 | Actl6a | 47 kDa | 9 C | 0.917 | 0.8149 |
| Low-density lipoprotein receptor-related protein 2 | A2ARV4 | Lrp2 | 519 kDa | 329 C | 0.916 | 0.57159 |
| Eukaryotic translation initiation factor 5A-1 | P63242 | Eif5a | 17 kDa | 4 C | 0.915 | 0.27707 |
| Serine/threonine-protein phosphatase PP1-beta catalytic subunit | P62141 | Ppp1cb | 37 kDa | 14 C | 0.914 | 0.34864 |
| Actin, cytoplasmic 2 | P63260 | Actg1 | 42 kDa | 6 C | 0.912 | 0.39091 |
| Isochorismatase domain-containing protein 2A | P85094 | Isoc2a | 22 kDa | 6 C | 0.911 | 0.3153 |
| 3-hydroxybutyrate dehydrogenase type 2 | Q8JZV9 | Bdh2 | 27 kDa | 6 C | 0.909 | 0.51852 |
| Isoform Rpn10B of 26S proteasome non-ATPase regulatory subunit 4 | O35226-2 (+1 | Psmd4 | 41 kDa | 4 C | 0.909 | 0.67787 |
| Translationally-controlled tumor protein | P63028 | Tpt1 | 19 kDa | 2 C | 0.909 | 0.67787 |
| Calreticulin | P14211 | Calr | 48 kDa | 6 C | 0.907 | 0.30355 |

| | | | | | | |
|---|---|---|---|---|---|---|
| Vinculin | Q64727 | Vcl | 117 kDa | 10 C | 0.907 | 0.52158 |
| Meprin A subunit alpha | P28825 | Mep1a | 84 kDa | 19 C | 0.906 | 0.20961 |
| Non-specific lipid-transfer protein | P32020 | Scp2 | 59 kDa | 11 C | 0.905 | 0.48182 |
| Hemoglobin subunit alpha | P01942 | Hba | 15 kDa | 1 C | 0.905 | 0.70108 |
| Gelsolin | P13020 | Gsn | 86 kDa | 7 C | 0.904 | 0.45863 |
| Thioredoxin-dependent peroxide reductase, mitochondrial | P20108 | Prdx3 | 28 kDa | 4 C | 0.900 | 0.3153 |
| Heterogeneous nuclear ribonucleoprotein K | P61979 (+1) | Hnrnpk | 51 kDa | 5 C | 0.900 | 0.43533 |
| FAD-linked sulfhydryl oxidase ALR | P56213 | Gfer | 23 kDa | 8 C | 0.900 | 0.64333 |
| Malate dehydrogenase, cytoplasmic | P14152 | Mdh1 | 37 kDa | 3 C | 0.900 | 0.7376 |
| Ketohexokinase | P97328 | Khk | 33 kDa | 10 C | 0.900 | 0.76764 |
| Acyl-coenzyme A thioesterase THEM4 | Q3UUI3 | Them4 | 26 kDa | 4 C | 0.900 | 0.76764 |
| Creatine kinase B-type | Q04447 | Ckb | 43 kDa | 5 C | 0.900 | 0.79526 |
| ATP synthase subunit gamma, mitochondrial | Q91VR2 | Atp5c1 | 33 kDa | 2 C | 0.900 | 0.88835 |
| Nucleoside diphosphate kinase B | Q01768 | Nme2 | 17 kDa | 2 C | 0.896 | 0.62227 |
| Isoform Cytoplasmic+peroxisomal of Peroxiredoxin-5, mitochondrial | P99029-2 | Prdx5 | 17 kDa | 6 C | 0.894 | 0.17949 |
| Protein disulfide-isomerase A3 | P27773 | Pdia3 | 57 kDa | 8 C | 0.892 | 0.42689 |
| Clathrin light chain A | O08585 | Clta | 26 kDa | 1 C | 0.889 | 0.64333 |
| N-acetylglucosamine-6-phosphate deacetylase | Q8JZV7 | Amdhd2 | 44 kDa | 8 C | 0.889 | 0.72466 |
| Glia maturation factor beta | Q9CQI3 | Gmfb | 17 kDa | 3 C | 0.889 | 0.76764 |
| Aldose 1-epimerase | Q8K157 | Galm | 38 kDa | 4 C | 0.889 | 0.8149 |
| Isoform 2 of Alpha-aminoadipic semialdehyde dehydrogenase | Q9DBF1-2 | Aldh7a1 | 56 kDa | 9 C | 0.886 | 0.09126 |
| Nucleoside diphosphate kinase A | P15532 | Nme1 | 17 kDa | 2 C | 0.885 | 0.60717 |
| Heterogeneous nuclear ribonucleoprotein A1 | P49312 (+1) | Hnrnpa1 | 34 kDa | 2 C | 0.885 | 0.63877 |
| Enoyl-CoA hydratase, mitochondrial | Q8BH95 | Echs1 | 31 kDa | 7 C | 0.884 | 0.28948 |
| Spectrin alpha chain, non-erythrocytic 1 | P16546 | Sptan1 | 285 kDa | 14 C | 0.883 | 0.42112 |
| Methylcrotonoyl-CoA carboxylase beta chain, mitochondrial | Q3ULD5 | Mccc2 | 61 kDa | 10 C | 0.883 | 0.13561 |
| 3-hydroxyacyl-CoA dehydrogenase type-2 | O08756 | Hsd17b10 | 27 kDa | 2 C | 0.882 | 0.7376 |
| Villin-1 | Q62468 | Vil1 | 93 kDa | 9 C | 0.881 | 0.04742 |
| Adenosylhomocysteinase | P50247 | Ahcy | 48 kDa | 9 C | 0.880 | 0.18407 |
| Antithrombin-III | P32261 | Serpinc1 | 52 kDa | 9 C | 0.880 | 0.34864 |
| Homogentisate 1,2-dioxygenase | O09173 | Hgd | 50 kDa | 14 C | 0.880 | 0.24424 |
| GMP reductase 1 | Q9DCZ1 | Gmpr | 37 kDa | 9 C | 0.879 | 0.77744 |
| Selenium-binding protein 1 | P17563 | Selenbp1 | 53 kDa | 10 C | 0.878 | 0.30456 |
| Hemopexin | Q91X72 | Hpx | 51 kDa | 13 C | 0.875 | 0.20615 |
| Talin-2 | Q71LX4 | Tln2 | 254 kDa | 40 C | 0.875 | 0.74152 |
| Prostaglandin E synthase 3 | Q9R0Q7 | Ptges3 | 19 kDa | 5 C | 0.875 | 0.74152 |
| Quinone oxidoreductase | P47199 | Cryz | 35 kDa | 5 C | 0.874 | 0.08365 |
| Prostaglandin reductase 2 | Q8VDQ1 | Ptgr2 | 38 kDa | 8 C | 0.871 | 0.87186 |
| Proteasome subunit alpha type-4 | Q9R1P0 | Psma4 | 29 kDa | 5 C | 0.870 | 0.62396 |
| 14-3-3 protein epsilon | P62259 | Ywhae | 29 kDa | 3 C | 0.867 | 0.25891 |
| Isoform 2 of Heterogeneous nuclear ribonucleoprotein A3 | Q8BG05-2 | Hnrnpa3 | 37 kDa | 4 C | 0.865 | 0.49701 |
| ATP synthase subunit alpha, mitochondrial | Q03265 | Atp5a1 | 60 kDa | 2 C | 0.860 | 0.74872 |
| DAZ-associated protein 1 | Q9JII5 | Dazap1 | 43 kDa | 4 C | 0.857 | 0.56144 |
| ADP/ATP translocase 2 | P51881 | Slc25a5 | 33 kDa | 4 C | 0.857 | 0.57339 |
| Endoplasmic reticulum resident protein 44 | Q9D1Q6 | Erp44 | 47 kDa | 7 C | 0.857 | 0.57339 |
| Transgelin | P37804 | Tagln | 23 kDa | 1 C | 0.857 | 0.62909 |
| Transcription elongation factor A protein 1 | P10711 | Tcea1 | 34 kDa | 8 C | 0.857 | 0.64333 |
| Threonine synthase-like 2 | Q80W22 | Thnsl2 | 54 kDa | 14 C | 0.857 | 0.72466 |
| Isoform 2 of Cellular nucleic acid-binding protein | P53996-2 (+1 | Cnbp | 19 kDa | 22 C | 0.857 | 0.76764 |
| Ig kappa chain V-VI region NQ2-6.1 | P04945 | | 12 kDa | 2 C | 0.857 | 0.76764 |
| Alpha-1-antitrypsin 1-2 | P22599 | Serpina1b | 46 kDa | 3 C | 0.857 | 0.8069 |
| Carbonic anhydrase 1 | P13634 | Ca1 | 28 kDa | 1 C | 0.857 | 0.8298 |
| Phosphatidylethanolamine-binding protein 1 | P70296 | Pebp1 | 21 kDa | 3 C | 0.855 | 0.27944 |
| Vitamin D-binding protein | P21614 | Gc | 54 kDa | 28 C | 0.852 | 0.28664 |
| Galectin-1 | P16045 | Lgals1 | 15 kDa | 6 C | 0.850 | 0.53908 |
| Glyceraldehyde-3-phosphate dehydrogenase | P16858 | Gapdh | 36 kDa | 5 C | 0.850 | 0.65196 |
| Ran-specific GTPase-activating protein | P34022 | Ranbp1 | 24 kDa | 3 C | 0.850 | 0.78492 |
| Succinate dehydrogenase [ubiquinone] iron-sulfur subunit, mitochondrial | Q9CQA3 | Sdhb | 32 kDa | 14 C | 0.848 | 0.29801 |
| Protein ABHD14B | Q8VCR7 | Abhd14b | 22 kDa | 2 C | 0.846 | 0.29426 |
| Protein disulfide-isomerase A4 | P08003 | Pdia4 | 72 kDa | 6 C | 0.846 | 0.29426 |

| | | | | | | |
|---|---|---|---|---|---|---|
| 14 kDa phosphohistidine phosphatase | Q9DAK9 | Phpt1 | 14 kDa | 3 C | 0.846 | 0.51852 |
| Probable ATP-dependent RNA helicase DDX17 | Q501J6 | Ddx17 | 72 kDa | 11 C | 0.846 | 0.62131 |
| Serine/threonine-protein phosphatase 5 | Q60676 | Ppp5c | 57 kDa | 11 C | 0.846 | 0.62131 |
| Dihydropyrimidinase-related protein 3 | Q62188 | Dpysl3 | 62 kDa | 7 C | 0.844 | 0.2378 |
| Heterogeneous nuclear ribonucleoproteins A2/B1 | O88569 | Hnrnpa2b1 | 37 kDa | 1 C | 0.844 | 0.51452 |
| von Willebrand factor A domain-containing protein 5A | Q99KC8 | Vwa5a | 87 kDa | 13 C | 0.842 | 0.34864 |
| Rho GDP-dissociation inhibitor 1 | Q99PT1 | Arhgdia | 23 kDa | 1 C | 0.842 | 0.53908 |
| Heat shock 70 kDa protein 4 | Q61316 | Hspa4 | 94 kDa | 14 C | 0.841 | 0.4995 |
| Fibrinogen alpha chain | E9PV24 | Fga | 87 kDa | 13 C | 0.840 | 0.49177 |
| Alcohol dehydrogenase [NADP(+)] | Q9JII6 | Akr1a1 | 37 kDa | 4 C | 0.840 | 0.76764 |
| Enoyl-CoA hydratase domain-containing protein 2, mitochondrial | Q3TLP5 | Echdc2 | 32 kDa | 6 C | 0.839 | 0.38232 |
| Succinate dehydrogenase [ubiquinone] flavoprotein subunit, mitochondrial | Q8K2B3 | Sdha | 73 kDa | 19 C | 0.837 | 0.1416 |
| Oxygen-dependent coproporphyrinogen-III oxidase, mitochondrial | P36552 | Cpox | 50 kDa | 10 C | 0.837 | 0.24769 |
| Malate dehydrogenase, mitochondrial | P08249 | Mdh2 | 36 kDa | 8 C | 0.837 | 0.42165 |
| Fumarylacetoacetase | P35505 | Fah | 46 kDa | 6 C | 0.835 | 0.24112 |
| Isoform 2 of Vacuolar protein sorting-associated protein 26A | P40336-2 | Vps26a | 42 kDa | 2 C | 0.833 | 0.3739 |
| UPF0598 protein C8orf82 homolog | Q8VE95 | | 24 kDa | 6 C | 0.833 | 0.64333 |
| Properdin | P11680 | Cfp | 50 kDa | 44 C | 0.833 | 0.64333 |
| Calpain small subunit 1 | O88456 | Capns1 | 28 kDa | 2 C | 0.833 | 0.69176 |
| Ig kappa chain V-II region 7S34.1 | P01630 | | 12 kDa | 3 C | 0.833 | 0.72466 |
| Branched-chain-amino-acid aminotransferase, mitochondrial | O35855 | Bcat2 | 44 kDa | 10 C | 0.833 | 0.79526 |
| Heat shock protein HSP 90-alpha | P07901 | Hsp90aa1 | 85 kDa | 7 C | 0.833 | 0.8416 |
| Actin-related protein 3 | Q99JY9 | Actr3 | 47 kDa | 8 C | 0.830 | 0.52094 |
| Transaldolase | Q93092 | Taldo1 | 37 kDa | 3 C | 0.829 | 0.58593 |
| Profilin-1 | P62962 | Pfn1 | 15 kDa | 3 C | 0.826 | 0.29426 |
| Pyridoxal kinase | Q8K183 | Pdxk | 35 kDa | 5 C | 0.824 | 0.48631 |
| Heterogeneous nuclear ribonucleoprotein H | O35737 | Hnrnph1 | 49 kDa | 1 C | 0.821 | 0.32616 |
| 3-hydroxyanthranilate 3,4-dioxygenase | Q78JT3 | Haao | 33 kDa | 3 C | 0.820 | 0.29542 |
| COP9 signalosome complex subunit 5 | O35864 | Cops5 | 38 kDa | 4 C | 0.820 | 0.69662 |
| Exosome complex component RRP4 | Q8VBV3 | Exosc2 | 33 kDa | 4 C | 0.820 | 0.69662 |
| Cell division control protein 42 homolog | P60766 | Cdc42 | 21 kDa | 7 C | 0.820 | 0.69662 |
| 2-oxoglutarate dehydrogenase, mitochondrial | Q60597 | Ogdh | 116 kDa | 21 C | 0.820 | 0.31943 |
| Glutathione reductase, mitochondrial | P47791 | Gsr | 54 kDa | 11 C | 0.819 | 0.20326 |
| Isoform 2 of Methionine adenosyltransferase 2 subunit beta | Q99LB6-2 | Mat2b | 36 kDa | 7 C | 0.818 | 0.11612 |
| Malectin | Q6ZQI3 | Mlec | 32 kDa | 3 C | 0.818 | 0.3739 |
| Ig kappa chain V-V region HP R16.7 | P01644 (+1) | | 12 kDa | 2 C | 0.818 | 0.3739 |
| Coactosin-like protein | Q9CQI6 | Cotl1 | 16 kDa | 2 C | 0.818 | 0.56144 |
| Protein disulfide-isomerase | P09103 | P4hb | 57 kDa | 7 C | 0.818 | 0.10866 |
| Thioredoxin reductase 1, cytoplasmic | Q9JMH6 | Txnrd1 | 67 kDa | 21 C | 0.816 | 0.27204 |
| Pyruvate carboxylase, mitochondrial | Q05920 | Pc | 130 kDa | 13 C | 0.815 | 0.80054 |
| Carboxymethylenebutenolidase homolog | Q8R1G2 | Cmbl | 28 kDa | 6 C | 0.813 | 0.34864 |
| Isoform M1 of Pyruvate kinase PKM | P52480-2 | Pkm | 58 kDa | 9 C | 0.813 | 0.41687 |
| UDP-N-acetylhexosamine pyrophosphorylase-like protein 1 | Q3TW96 | Uap1l1 | 57 kDa | 13 C | 0.813 | 0.60717 |
| Filamin-B | Q80X90 | Flnb | 278 kDa | 43 C | 0.811 | 0.54893 |
| Proteasome subunit beta type-3 | Q9R1P1 | Psmb3 | 23 kDa | 5 C | 0.810 | 0.11612 |
| Hydroxymethylglutaryl-CoA lyase, mitochondrial | P38060 | Hmgcl | 34 kDa | 8 C | 0.804 | 0.13178 |
| Polyadenylate-binding protein 1 | P29341 | Pabpc1 | 71 kDa | 4 C | 0.804 | 0.25942 |
| Eukaryotic translation initiation factor 6 | O55135 | Eif6 | 27 kDa | 8 C | 0.804 | 0.76897 |
| Transgelin-2 | Q9WVA4 | Tagln2 | 22 kDa | 3 C | 0.800 | 0.0936 |
| Cytidine deaminase | P56389 | Cda | 16 kDa | 7 C | 0.800 | 0.11612 |
| Isoform 2 of Nitrilase homolog 1 | Q8VDK1-2 | Nit1 | 32 kDa | 13 C | 0.800 | 0.19174 |
| Macrophage-capping protein | P24452 | Capg | 39 kDa | 5 C | 0.800 | 0.2524 |
| Beta-arrestin-1 | Q8BWG8 (+1 | Arrb1 | 47 kDa | 8 C | 0.800 | 0.28786 |
| Cadherin-16 | O88338 | Cdh16 | 90 kDa | 7 C | 0.800 | 0.35081 |
| Delta-aminolevulinic acid dehydratase | P10518 | Alad | 36 kDa | 8 C | 0.800 | 0.41351 |
| Aflatoxin B1 aldehyde reductase member 2 | Q8CG76 | Akr7a2 | 41 kDa | 8 C | 0.800 | 0.43533 |
| Glutaredoxin-3 | Q9CQM9 | Glrx3 | 38 kDa | 5 C | 0.800 | 0.47662 |
| Hypoxanthine-guanine phosphoribosyltransferase | P00493 | Hprt1 | 25 kDa | 4 C | 0.800 | 0.48182 |
| Heterogeneous nuclear ribonucleoprotein Q | Q7TMK9 | Syncrip | 70 kDa | 4 C | 0.800 | 0.49292 |
| Protein NipSnap homolog 3B | Q9CQE1 | Nipsnap3b | 28 kDa | 2 C | 0.800 | 0.51852 |

| | | | | | | |
|---|---|---|---|---|---|---|
| Protein-glucosylgalactosylhydroxylysine glucosidase | Q8BP56 | Pgghg | 76 kDa | 8 C | 0.800 | 0.51852 |
| NADH dehydrogenase [ubiquinone] 1 alpha subcomplex subunit 2 | Q9CQ75 | Ndufa2 | 11 kDa | 2 C | 0.800 | 0.51852 |
| 6-pyruvoyl tetrahydrobiopterin synthase | Q9R1Z7 | Pts | 16 kDa | 1 C | 0.800 | 0.51852 |
| StAR-related lipid transfer protein 5 | Q9EPQ7 | Stard5 | 24 kDa | 8 C | 0.800 | 0.51852 |
| S-methyl-5'-thioadenosine phosphorylase | Q9CQ65 | Mtap | 31 kDa | 10 C | 0.800 | 0.61301 |
| Xaa-Pro aminopeptidase 1 | Q6P1B1 | Xpnpep1 | 70 kDa | 12 C | 0.800 | 0.65605 |
| Dihydropyrimidinase-related protein 2 | O08553 | Dpysl2 | 62 kDa | 7 C | 0.800 | 0.00211 |
| N-acyl-aromatic-L-amino acid amidohydrolase (carboxylate-forming) | Q91XE4 | Acy3 | 35 kDa | 8 C | 0.798 | 0.14932 |
| Persulfide dioxygenase ETHE1, mitochondrial | Q9DCM0 | Ethe1 | 28 kDa | 9 C | 0.795 | 0.20511 |
| Triosephosphate isomerase | P17751 | Tpi1 | 32 kDa | 9 C | 0.794 | 0.17882 |
| Deoxynucleoside triphosphate triphosphohydrolase SAMHD1 | Q60710 | Samhd1 | 73 kDa | 17 C | 0.792 | 0.39419 |
| Succinyl-CoA:3-ketoacid coenzyme A transferase 1, mitochondrial | Q9D0K2 | Oxct1 | 56 kDa | 7 c | 0.792 | 0.4213 |
| Serine protease inhibitor A3K | P07759 | Serpina3k | 47 kDa | 4 C | 0.792 | 0.29801 |
| L-lactate dehydrogenase A chain | P06151 | Ldha | 36 kDa | 6 C | 0.792 | 0.65262 |
| Omega-amidase NIT2 | Q9JHW2 | Nit2 | 31 kDa | 2 C | 0.789 | 0.11612 |
| Proteasome subunit alpha type-5 | Q9Z2U1 | Psma5 | 26 kDa | 3 C | 0.789 | 0.41072 |
| Glucosidase 2 subunit beta | O08795 (+1) | Prkcsh | 59 kDa | 17 C | 0.786 | 0.25082 |
| NAD kinase 2, mitochondrial | Q8C5H8 (+1) | Nadk2 | 51 kDa | 9 C | 0.786 | 0.34864 |
| Thioredoxin domain-containing protein 17 | Q9CQM5 | Txndc17 | 14 kDa | 6 C | 0.786 | 0.53908 |
| Vimentin | P20152 | Vim | 54 kDa | 1 C | 0.786 | 0.56553 |
| ES1 protein homolog, mitochondrial | Q9D172 | D10Jhu81e | 28 kDa | 6 C | 0.786 | 0.62396 |
| Peroxisomal sarcosine oxidase | Q9D826 | Pipox | 44 kDa | 11 C | 0.784 | 0.15059 |
| 60S ribosomal protein L12 | P35979 | Rpl12 | 18 kDa | 3 C | 0.783 | 0.2378 |
| Catenin alpha-1 | P26231 | Ctnna1 | 100 kDa | 12 C | 0.781 | 0.45283 |
| Inositol oxygenase | Q9QXN5 | Miox | 33 kDa | 6 C | 0.780 | 0.68833 |
| Hsc70-interacting protein | Q99L47 | St13 | 42 kDa | 3 C | 0.778 | 0.07048 |
| Malignant T-cell-amplified sequence 1 | Q9DB27 | Mcts1 | 21 kDa | 4 C | 0.778 | 0.20511 |
| Protein disulfide-isomerase A6 | Q922R8 | Pdia6 | 48 kDa | 7 C | 0.778 | 0.27458 |
| Glyoxalase domain-containing protein 5 | Q9D8I3 | Glod5 | 17 kDa | 5 C | 0.778 | 0.3739 |
| Elongation factor 1-beta | O70251 | Eef1b | 25 kDa | 3 C | 0.778 | 0.3739 |
| Twinfilin-1 | Q91YR1 | Twf1 | 40 kDa | 4 C | 0.778 | 0.3739 |
| Probable D-lactate dehydrogenase, mitochondrial | Q7TNG8 | Ldhd | 52 kDa | 12 C | 0.778 | 0.46552 |
| Mammalian ependymin-related protein 1 | Q99M71 | Epdr1 | 25 kDa | 7 C | 0.775 | 0.66463 |
| Exosome complex component RRP43 | Q9D753 | Exosc8 | 30 kDa | 10 C | 0.775 | 0.66463 |
| Phytanoyl-CoA dioxygenase, peroxisomal | O35386 | Phyh | 39 kDa | 8 C | 0.775 | 0.66463 |
| Purine nucleoside phosphorylase | P23492 | Pnp | 32 kDa | 5 C | 0.775 | 0.03347 |
| Carbonic anhydrase 3 | P16015 | Ca3 | 29 kDa | 5 C | 0.769 | 0.25082 |
| Proteasome subunit beta type-7 | P70195 | Psmb7 | 30 kDa | 6 C | 0.769 | 0.25082 |
| Argininosuccinate synthase | P16460 | Ass1 | 47 kDa | 5 C | 0.769 | 0.64389 |
| Phosphate carrier protein, mitochondrial | Q8VEM8 | Slc25a3 | 40 kDa | 8 C | 0.769 | 0.78945 |
| Stress-70 protein, mitochondrial | P38647 | Hspa9 | 73 kDa | 5 C | 0.769 | 0.30411 |
| Ezrin | P26040 | Ezr | 69 kDa | 2 C | 0.767 | 0.2889 |
| Acyl-coenzyme A synthetase ACSM2, mitochondrial | Q8K0L3 (+1) | Acsm2 | 64 kDa | 9 C | 0.766 | 0.01709 |
| Histidine triad nucleotide-binding protein 2, mitochondrial | Q9D0S9 | Hint2 | 17 kDa | 1 C | 0.765 | 0.14815 |
| Fibrinogen beta chain | Q8K0E8 | Fgb | 55 kDa | 12 C | 0.763 | 0.02564 |
| Peptidyl-prolyl cis-trans isomerase A | P17742 | Ppia | 18 kDa | 3 C | 0.763 | 0.13988 |
| 60 kDa heat shock protein, mitochondrial | P63038 | Hspd1 | 61 kDa | 3 C | 0.761 | 0.19543 |
| Succinate--CoA ligase [GDP-forming] subunit beta, mitochondrial | Q9Z2I8 | Suclg2 | 47 kDa | 5 C | 0.761 | 0.81805 |
| Alpha-1-antitrypsin 1-1 | P07758 (+1) | Serpina1a | 46 kDa | 3 C | 0.759 | 0.43425 |
| Serine/arginine-rich splicing factor 3 | P84104 (+1) | Srsf3 | 19 kDa | 4 C | 0.756 | 0.75967 |
| Ig gamma-3 chain C region | P03987 (+1) | | 44 kDa | 10 C | 0.756 | 0.75967 |
| Calponin-3 | Q9DAW9 | Cnn3 | 36 kDa | 3 C | 0.756 | 0.75967 |
| Transitional endoplasmic reticulum ATPase | Q01853 | Vcp | 89 kDa | 12 C | 0.751 | 0.20293 |
| Retinol-binding protein 4 | Q00724 | Rbp4 | 23 kDa | 6 C | 0.750 | 0.11612 |
| Methylglutaconyl-CoA hydratase, mitochondrial | Q9JLZ3 | Auh | 33 kDa | 5 C | 0.750 | 0.20511 |
| Mannose-binding protein A | P39039 | Mbl1 | 25 kDa | 8 C | 0.750 | 0.20511 |
| Isoform Smooth muscle of Myosin light polypeptide 6 | Q60605-2 | Myl6 | 17 kDa | 3 C | 0.750 | 0.2378 |
| Peptidyl-prolyl cis-trans isomerase FKBP4 | P30416 | Fkbp4 | 52 kDa | 7 C | 0.750 | 0.25643 |
| Proteasome subunit beta type-2 | Q9R1P3 | Psmb2 | 23 kDa | 3 C | 0.750 | 0.27458 |
| Talin-1 | P26039 | Tln1 | 270 kDa | 38 C | 0.750 | 0.35004 |

| | | | | | | | |
|---|---|---|---|---|---|---|---|
| Afamin | O89020 (+1) | Afm | 69 kDa | 34 C | | 0.750 | 0.3739 |
| Elongation factor 1-delta | P57776 | Eef1d | 31 kDa | 2 C | | 0.750 | 0.3739 |
| Sorting nexin-3 | O70492 | Snx3 | 19 kDa | 1 C | | 0.750 | 0.56144 |
| Nucleoprotein TPR | F6ZDS4 | Tpr | 274 kDa | 7 C | | 0.750 | 0.60865 |
| 4-hydroxyphenylpyruvate dioxygenase | P49429 | Hpd | 45 kDa | 4 C | | 0.750 | 0.64333 |
| Ig kappa chain C region | P01837 | | 12 kDa | 3 C | | 0.750 | 0.0453 |
| Transketolase | P40142 | Tkt | 68 kDa | 12 C | | 0.748 | 0.04863 |
| Carboxylesterase 1F | Q91WU0 | Ces1f | 62 kDa | 7 C | | 0.747 | 0.19429 |
| Glutathione S-transferase P 1 | P19157 | Gstp1 | 24 kDa | 3 C | | 0.744 | 0.02061 |
| Phospholipid hydroperoxide glutathione peroxidase, mitochondrial | O70325 | Gpx4 | 22 kDa | 10 C | | 0.743 | 0.1338 |
| Fibrinogen gamma chain | Q8VCM7 | Fgg | 49 kDa | 12 C | | 0.742 | 0.01754 |
| Triokinase/FMN cyclase | Q8VC30 | Tkfc | 60 kDa | 5 C | | 0.741 | 0.78377 |
| Propionyl-CoA carboxylase beta chain, mitochondrial | Q99MN9 | Pccb | 58 kDa | 11 C | | 0.740 | 0.23391 |
| Proteasome subunit alpha type-3 | O70435 | Psma3 | 28 kDa | 4 C | | 0.737 | 0.08901 |
| Ig heavy chain V-III region J606 | P01801 | | 13 kDa | 2 C | | 0.737 | 0.35601 |
| Cofilin-1 | P18760 | Cfl1 | 19 kDa | 4 C | | 0.736 | 0.09126 |
| Sorbitol dehydrogenase | Q64442 | Sord | 38 kDa | 10 C | | 0.734 | 0.13424 |
| Palmitoyl-protein thioesterase 1 | O88531 | Ppt1 | 34 kDa | 10 C | | 0.733 | 0.69536 |
| Mitochondrial intermembrane space import and assembly protein 40 | Q8VEA4 | Chchd4 | 16 kDa | 7 C | | 0.733 | 0.11612 |
| Ig kappa chain V-II region 26-10 | P01631 | | 12 kDa | 2 C | | 0.733 | 0.32946 |
| 4-aminobutyrate aminotransferase, mitochondrial | P61922 | Abat | 56 kDa | 12 C | | 0.733 | 0.41072 |
| L-lactate dehydrogenase B chain | P16125 | Ldhb | 37 kDa | 5 C | | 0.733 | 0.4481 |
| Kynurenine/alpha-aminoadipate aminotransferase, mitochondrial | Q9WVM8 | Aadat | 48 kDa | 5 C | | 0.728 | 0.58622 |
| Serine--tRNA ligase, cytoplasmic | P26638 | Sars | 58 kDa | 9 C | | 0.727 | 0.25082 |
| Multifunctional protein ADE2 | Q9DCL9 | Paics | 47 kDa | 13 C | | 0.727 | 0.4103 |
| Tubulin-folding cofactor B | Q9D1E6 | Tbcb | 27 kDa | 5 C | | 0.727 | 0.4676 |
| Filamin-A | Q8BTM8 | Flna | 281 kDa | 38 C | | 0.727 | 0.14481 |
| NADH-ubiquinone oxidoreductase 75 kDa subunit, mitochondrial | Q91VD9 | Ndufs1 | 80 kDa | 18 C | | 0.722 | 0.20947 |
| Methyltransferase-like 26 | Q9DCS2 | Mettl26 | 23 kDa | 6 C | | 0.722 | 0.2378 |
| Long-chain specific acyl-CoA dehydrogenase, mitochondrial | P51174 | Acadl | 48 kDa | 7 C | | 0.721 | 0.18407 |
| 40S ribosomal protein SA | P14206 | Rpsa | 33 kDa | 2 C | | 0.720 | 0.25643 |
| Alpha-methylacyl-CoA racemase | O09174 | Amacr | 42 kDa | 6 C | | 0.720 | 0.29618 |
| 3-hydroxyisobutyrate dehydrogenase, mitochondrial | Q99L13 | Hibadh | 35 kDa | 12 C | | 0.717 | 0.24353 |
| NADH dehydrogenase [ubiquinone] flavoprotein 1, mitochondrial | Q91YT0 | Ndufv1 | 51 kDa | 12 C | | 0.717 | 0.30383 |
| Elongation factor 2 | P58252 | Eef2 | 95 kDa | 7 C | | 0.716 | 0.25809 |
| Endoplasmic reticulum resident protein 29 | P57759 | Erp29 | 29 kDa | 1 C | | 0.714 | 0.0993 |
| Protein arginine N-methyltransferase 5 | Q8CIG8 | Prmt5 | 73 kDa | 12 C | | 0.714 | 0.47662 |
| Pyruvate dehydrogenase E1 component subunit alpha, somatic form, mitochondrial | P35486 | Pdha1 | 43 kDa | 12 C | | 0.714 | 0.60183 |
| 78 kDa glucose-regulated protein | P20029 | Hspa5 | 72 kDa | 1 C | | 0.713 | 0.01234 |
| Carboxylesterase 1D | Q8VCT4 | Ces1d | 62 kDa | 5 C | | 0.709 | 0.28214 |
| Caprin-1 | Q60865 | Caprin1 | 78 kDa | 3 C | | 0.708 | 0.09126 |
| Sulfite oxidase, mitochondrial | Q8R086 | Suox | 61 kDa | 9 C | | 0.708 | 0.12738 |
| Catalase | P24270 | Cat | 60 kDa | 5 C | | 0.708 | 0.06726 |
| Proteasome subunit alpha type-7 | Q9Z2U0 | Psma7 | 28 kDa | 3 C | | 0.706 | 0.3153 |
| Fumarate hydratase, mitochondrial | P97807 | Fh | 54 kDa | 4 C | | 0.706 | 0.33908 |
| Radixin | P26043 | Rdx | 69 kDa | 1 C | | 0.706 | 0.42997 |
| NADH dehydrogenase [ubiquinone] flavoprotein 2, mitochondrial | Q9D6J6 | Ndufv2 | 27 kDa | 6 C | | 0.706 | 0.47343 |
| Dimethylglycine dehydrogenase, mitochondrial | Q9DBT9 | Dmgdh | 97 kDa | 4 C | | 0.706 | 0.51452 |
| Superoxide dismutase [Cu-Zn] | P08228 | Sod1 | 16 kDa | 3 C | | 0.704 | 0.65767 |
| Histidine triad nucleotide-binding protein 1 | P70349 | Hint1 | 14 kDa | 2 C | | 0.700 | 0.10119 |
| Beta-2-glycoprotein 1 | Q01339 | Apoh | 39 kDa | 23 C | | 0.700 | 0.10119 |
| Inositol-3-phosphate synthase 1 | Q9JHU9 | Isyna1 | 61 kDa | 11 C | | 0.700 | 0.36785 |
| Ubiquitin-conjugating enzyme E2 N | P61089 | Ube2n | 17 kDa | 1 C | | 0.700 | 0.41687 |
| Glutathione S-transferase A3 | P30115 | Gsta3 | 25 kDa | 1 C | | 0.700 | 0.4676 |
| Adenosine kinase | P55264 | Adk | 40 kDa | 6 C | | 0.700 | 0.50716 |
| S-phase kinase-associated protein 1 | Q9WTX5 | Skp1 | 19 kDa | 3 C | | 0.696 | 0.35081 |
| Vigilin | Q8VDJ3 | Hdlbp | 142 kDa | 10 C | | 0.692 | 0.06468 |
| Hydroxyacylglutathione hydrolase, mitochondrial | Q99KB8 | Hagh | 34 kDa | 8 C | | 0.692 | 0.12705 |
| Glycerol-3-phosphate dehydrogenase [NAD(+)], cytoplasmic | P13707 | Gpd1 | 38 kDa | 11 C | | 0.692 | 0.20511 |
| Serine/threonine-protein phosphatase 2A 55 kDa regulatory subunit B alpha isoform | Q6P1F6 | Ppp2r2a | 52 kDa | 9 C | | 0.692 | 0.3739 |

| | | | | | | |
|---|---|---|---|---|---|---|
| Ester hydrolase C11orf54 homolog | Q91V76 | | 35 kDa | 8 C | 0.690 | 0.16512 |
| Probable aminopeptidase NPEPL1 | Q6NSR8 | Npepl1 | 56 kDa | 16 C | 0.690 | 0.17851 |
| Proteasome subunit alpha type-1 | Q9R1P4 | Psma1 | 30 kDa | 5 C | 0.690 | 0.20284 |
| Hydroxyacyl-coenzyme A dehydrogenase, mitochondrial | Q61425 | Hadh | 34 kDa | 5 C | 0.688 | 0.0869 |
| Fucose mutarotase | Q8R2K1 | Fuom | 17 kDa | 3 C | 0.688 | 0.20615 |
| Chloride intracellular channel protein 4 | Q9QYB1 | Clic4 | 29 kDa | 4 C | 0.688 | 0.78734 |
| Aspartyl aminopeptidase | Q9Z2W0 | Dnpep | 52 kDa | 10 C | 0.684 | 0.16492 |
| Isovaleryl-CoA dehydrogenase, mitochondrial | Q9JHI5 | Ivd | 46 kDa | 7 C | 0.684 | 0.53908 |
| Hydroxyacid oxidase 2 | Q9NYQ2 | Hao2 | 39 kDa | 8 C | 0.683 | 0.03113 |
| Exosome complex exonuclease RRP42 | Q9D0M0 | Exosc7 | 32 kDa | 12 C | 0.683 | 0.50092 |
| Iron-sulfur cluster assembly enzyme ISCU, mitochondrial | Q9D7P6 | Iscu | 18 kDa | 4 C | 0.683 | 0.50092 |
| Enoyl-CoA delta isomerase 1, mitochondrial | P42125 | Eci1 | 32 kDa | 5 C | 0.683 | 0.65571 |
| Betaine--homocysteine S-methyltransferase 1 | O35490 | Bhmt | 45 kDa | 8 C | 0.683 | 0.23464 |
| Poly(rC)-binding protein 1 | P60335 | Pcbp1 | 37 kDa | 9 C | 0.679 | 0.05334 |
| V-type proton ATPase subunit E 1 | P50518 | Atp6v1e1 | 26 kDa | 1 C | 0.678 | 0.439 |
| Mitochondrial amidoxime reducing component 2 | Q922Q1 | 2-Mar | 38 kDa | 11 C | 0.677 | 0.62227 |
| E3 ubiquitin-protein ligase NEDD4 | P46935 | Nedd4 | 103 kDa | 9 C | 0.677 | 0.62227 |
| Leucine-rich repeat-containing protein 59 | Q922Q8 | Lrrc59 | 35 kDa | 8 C | 0.677 | 0.62227 |
| Calcyclin-binding protein | Q9CXW3 | Cacybp | 27 kDa | 2 C | 0.677 | 0.62227 |
| Pyridoxine-5'-phosphate oxidase | Q91XF0 | Pnpo | 30 kDa | 6 C | 0.677 | 0.62227 |
| ADP/ATP translocase 1 | P48962 | Slc25a4 | 33 kDa | 4 C | 0.677 | 0.79137 |
| Inorganic pyrophosphatase 2, mitochondrial | Q91VM9 | Ppa2 | 38 kDa | 8 C | 0.672 | 0.05223 |
| Protein DJ-1 | Q99LX0 | Park7 | 20 kDa | 4 C | 0.667 | 0.05198 |
| Adrenodoxin, mitochondrial | P46656 | Fdx1 | 20 kDa | 8 C | 0.667 | 0.11612 |
| Ig kappa chain V-V region K2 (Fragment) | P01635 | | 13 kDa | 3 C | 0.667 | 0.11612 |
| Ubiquitin-conjugating enzyme E2 L3 | P68037 | Ube2l3 | 18 kDa | 3 C | 0.667 | 0.11612 |
| Isoform 2 of F-actin-capping protein subunit beta | P47757-2 | Capzb | 31 kDa | 5 C | 0.667 | 0.15502 |
| Serine-threonine kinase receptor-associated protein | Q9Z1Z2 | Strap | 38 kDa | 6 C | 0.667 | 0.20511 |
| Lactoylglutathione lyase | Q9CPU0 | Glo1 | 21 kDa | 3 C | 0.667 | 0.20511 |
| Stromal cell-derived factor 2 | Q9DCT5 | Sdf2 | 23 kDa | 4 C | 0.667 | 0.20511 |
| Actin-related protein 2/3 complex subunit 2 | Q9CVB6 | Arpc2 | 34 kDa | 2 C | 0.667 | 0.26962 |
| Hypoxia up-regulated protein 1 | Q9JKR6 | Hyou1 | 111 kDa | 4 C | 0.667 | 0.28786 |
| 2-iminobutanoate/2-iminopropanoate deaminase | P52760 | Rida | 14 kDa | 1 C | 0.667 | 0.30355 |
| Isoform Cytoplasmic of Cysteine desulfurase, mitochondrial | Q9Z1J3-2 | Nfs1 | 44 kDa | 7 C | 0.667 | 0.32946 |
| Receptor of activated protein C kinase 1 | P68040 | Rack1 | 35 kDa | 8 C | 0.667 | 0.46853 |
| Heat shock protein HSP 90-beta | P11499 | Hsp90ab1 | 83 kDa | 6 C | 0.667 | 0.5225 |
| Acidic leucine-rich nuclear phosphoprotein 32 family member A | O35381 | Anp32a | 29 kDa | 3 C | 0.667 | 0.56144 |
| Alcohol dehydrogenase 1 | P00329 | Adh1 | 40 kDa | 15 C | 0.667 | 0.64333 |
| Hydroxymethylglutaryl-CoA synthase, cytoplasmic | Q8JZK9 | Hmgcs1 | 58 kDa | 11 C | 0.662 | 0.61845 |
| Na(+)/H(+) exchange regulatory cofactor NHE-RF3 | Q9JIL4 | Pdzk1 | 56 kDa | 7 C | 0.658 | 0.01762 |
| Aspartate aminotransferase, mitochondrial | P05202 | Got2 | 47 kDa | 7 C | 0.657 | 0.11242 |
| Small glutamine-rich tetratricopeptide repeat-containing protein alpha | Q8BJU0 (+1) | Sgta | 34 kDa | 4 C | 0.656 | 0.73548 |
| Alpha-enolase | P17182 | Eno1 | 47 kDa | 6 C | 0.655 | 0.08712 |
| Creatine kinase U-type, mitochondrial | P30275 | Ckmt1 | 47 kDa | 7 C | 0.654 | 0.20284 |
| 4-hydroxy-2-oxoglutarate aldolase, mitochondrial | Q9DCU9 | Hoga1 | 35 kDa | 6 C | 0.650 | 0.26349 |
| Gephyrin | Q8BUV3 | Gphn | 83 kDa | 13 C | 0.650 | 0.61982 |
| Acetyl-CoA acetyltransferase, mitochondrial | Q8QZT1 | Acat1 | 45 kDa | 6 C | 0.646 | 0.0365 |
| ADP-ribosylation factor-binding protein GGA1 | Q8R0H9 | Gga1 | 70 kDa | 6 C | 0.645 | 0.34529 |
| Malonyl-CoA-acyl carrier protein transacylase, mitochondrial | Q8R3F5 | Mcat | 42 kDa | 12 C | 0.643 | 0.34649 |
| Glyoxylate reductase/hydroxypyruvate reductase | Q91Z53 | Grhpr | 35 kDa | 7 C | 0.643 | 0.01235 |
| Isoform 2 of Neutral alpha-glucosidase AB | Q8BHN3-2 | Ganab | 109 kDa | 8 C | 0.642 | 0.22536 |
| Ig heavy chain V region AC38 205.12 | P06330 | | 13 kDa | 2 C | 0.640 | 0.20284 |
| Ig gamma-2A chain C region secreted form | P01864 | | 37 kDa | 9 C | 0.638 | 0.4216 |
| Destrin | Q9R0P5 | Dstn | 19 kDa | 6 C | 0.632 | 0.09126 |
| Stress-induced-phosphoprotein 1 | Q60864 | Stip1 | 63 kDa | 11 C | 0.632 | 0.13988 |
| Protein-glutamine gamma-glutamyltransferase K | Q9JLF6 | Tgm1 | 90 kDa | 16 C | 0.632 | 0.25643 |
| Ubiquitin carboxyl-terminal hydrolase 5 | P56399 | Usp5 | 96 kDa | 16 C | 0.629 | 0.25316 |
| Actin-related protein 2/3 complex subunit 5 | Q9CPW4 | Arpc5 | 16 kDa | 1 C | 0.625 | 0.10119 |
| Acyl-coenzyme A thioesterase 4 | Q8BWN8 | Acot4 | 46 kDa | 6 C | 0.625 | 0.18407 |
| Bifunctional epoxide hydrolase 2 | P34914 | Ephx2 | 63 kDa | 10 C | 0.625 | 0.29214 |

| | | | | | | |
|---|---|---|---|---|---|---|
| Acetyl-CoA acetyltransferase, cytosolic | Q8CAY6 | Acat2 | 41 kDa | 8 C | 0.622 | 0.01316 |
| WD repeat domain phosphoinositide-interacting protein 3 | Q9CR39 | Wdr45b | 38 kDa | 15 C | 0.620 | 0.37977 |
| START domain-containing protein 10 | Q9JMD3 | Stard10 | 33 kDa | 7 C | 0.620 | 0.37977 |
| Selenoprotein P | P70274 | Selenop | 43 kDa | 18 C | 0.620 | 0.50393 |
| Endoplasmin | P08113 | Hsp90b1 | 92 kDa | 5 C | 0.620 | 0.50393 |
| Serine/threonine-protein phosphatase 2A catalytic subunit alpha isoform | P63330 | Ppp2ca | 36 kDa | 10 C | 0.620 | 0.50393 |
| Hsp90 co-chaperone Cdc37 | Q61081 | Cdc37 | 45 kDa | 9 C | 0.619 | 0.22412 |
| Heat shock cognate 71 kDa protein | P63017 | Hspa8 | 71 kDa | 4 C | 0.618 | 0.00275 |
| Peroxisomal bifunctional enzyme | Q9DBM2 | Ehhadh | 78 kDa | 10 C | 0.617 | 0.69763 |
| Ras GTPase-activating-like protein IQGAP1 | Q9JKF1 | Iqgap1 | 189 kDa | 14 C | 0.615 | 0.74312 |
| S-formylglutathione hydrolase | Q9R0P3 | Esd | 31 kDa | 10 C | 0.611 | 0.20988 |
| Phosphoglycerate kinase 1 | P09411 | Pgk1 | 45 kDa | 7 C | 0.611 | 0.32644 |
| Hydroxyacid-oxoacid transhydrogenase, mitochondrial | Q8R0N6 | Adhfe1 | 50 kDa | 8 C | 0.609 | 0.09445 |
| 40S ribosomal protein S12 | P63323 | Rps12 | 15 kDa | 7 C | 0.609 | 0.1338 |
| Isoform 2 of Peroxisomal acyl-coenzyme A oxidase 1 | Q9R0H0-2 | Acox1 | 75 kDa | 7 C | 0.606 | 0.04086 |
| Very long-chain specific acyl-CoA dehydrogenase, mitochondrial | P50544 | Acadvl | 71 kDa | 7 C | 0.600 | 0.07565 |
| Src substrate cortactin | Q60598 | Cttn | 61 kDa | 3 C | 0.600 | 0.11612 |
| Kynurenine--oxoglutarate transaminase 3 | Q71RI9 | Kyat3 | 51 kDa | 10 C | 0.600 | 0.11612 |
| Isocitrate dehydrogenase [NAD] subunit alpha, mitochondrial | Q9D6R2 | Idh3a | 40 kDa | 8 C | 0.600 | 0.27458 |
| Protein phosphatase 1B | P36993 | Ppm1b | 43 kDa | 12 C | 0.600 | 0.27458 |
| Isoamyl acetate-hydrolyzing esterase 1 homolog | Q9DB29 | Iah1 | 28 kDa | 8 C | 0.600 | 0.27944 |
| Ras-related protein Rab-1A | P62821 | Rab1A | 23 kDa | 4 C | 0.600 | 0.3739 |
| Cytosolic 10-formyltetrahydrofolate dehydrogenase | Q8R0Y6 | Aldh1l1 | 99 kDa | 15 C | 0.596 | 0.06736 |
| DnaJ homolog subfamily C member 12 | Q9R022 | Dnajc12 | 23 kDa | 4 C | 0.592 | 0.34832 |
| TAR DNA-binding protein 43 | Q921F2 | Tardbp | 45 kDa | 7 C | 0.591 | 0.20284 |
| Calbindin | P12658 | Calb1 | 30 kDa | 4 C | 0.588 | 0.15003 |
| Protein S100-A11 | P50543 | S100a11 | 11 kDa | 3 C | 0.588 | 0.45671 |
| Cytochrome b-c1 complex subunit 2, mitochondrial | Q9DB77 | Uqcrc2 | 48 kDa | 1 C | 0.588 | 0.54619 |
| Glutaredoxin-1 | Q9QUH0 | Glrx | 12 kDa | 5 C | 0.586 | 0.35246 |
| Copine-3 | Q8BT60 | Cpne3 | 60 kDa | 13 C | 0.583 | 0.27944 |
| Cytochrome b-c1 complex subunit Rieske, mitochondrial | Q9CR68 | Uqcrfs1 | 29 kDa | 5 C | 0.583 | 0.39821 |
| Glycine N-methyltransferase | Q9QXF8 | Gnmt | 33 kDa | 8 C | 0.581 | 0.25732 |
| Peroxisomal multifunctional enzyme type 2 | P51660 | Hsd17b4 | 79 kDa | 9 C | 0.581 | 0.00864 |
| Peroxisomal acyl-coenzyme A oxidase 3 | Q9EPL9 | Acox3 | 78 kDa | 14 C | 0.579 | 0.20674 |
| 3-ketoacyl-CoA thiolase, mitochondrial | Q8BWT1 | Acaa2 | 42 kDa | 8 C | 0.579 | 0.4442 |
| Moesin | P26041 | Msn | 68 kDa | 2 C | 0.576 | 0.26695 |
| Haloacid dehalogenase-like hydrolase domain-containing protein 3 | Q9CYW4 | Hdhd3 | 28 kDa | 4 C | 0.571 | 0.10119 |
| Selenocysteine lyase | Q9JLI6 | Scly | 47 kDa | 8 C | 0.571 | 0.14007 |
| Citrate lyase subunit beta-like protein, mitochondrial | Q8R4N0 | Clybl | 38 kDa | 6 C | 0.571 | 0.27908 |
| Presequence protease, mitochondrial | Q8K411 | Pitrm1 | 117 kDa | 20 C | 0.569 | 0.13911 |
| Phosphoglycerate mutase 1 | Q9DBJ1 | Pgam1 | 29 kDa | 2 C | 0.568 | 0.11612 |
| Oligoribonuclease, mitochondrial | Q9D8S4 | Rexo2 | 27 kDa | 4 C | 0.563 | 0.15502 |
| Cytochrome c oxidase subunit 5A, mitochondrial | P12787 | Cox5a | 16 kDa | 4 C | 0.556 | 0.06468 |
| Nucleoside diphosphate kinase 3 | Q9WV85 | Nme3 | 19 kDa | 3 C | 0.556 | 0.11612 |
| Peptidyl-prolyl cis-trans isomerase D | Q9CR16 | Ppid | 41 kDa | 7 C | 0.556 | 0.20511 |
| Proteasome subunit beta type-6 | Q60692 | Psmb6 | 25 kDa | 4 C | 0.556 | 0.20511 |
| Copper chaperone for superoxide dismutase | Q9WU84 | Ccs | 29 kDa | 10 C | 0.556 | 0.20511 |
| Heat shock 70 kDa protein 4L | P48722 | Hspa4l | 94 kDa | 15 C | 0.555 | 0.31717 |
| Clathrin heavy chain 1 | Q68FD5 | Cltc | 192 kDa | 31 C | 0.554 | 0.70728 |
| Dihydropyrimidine dehydrogenase [NADP(+)] | Q8CHR6 | Dpyd | 111 kDa | 35 C | 0.550 | 0.2627 |
| Dihydrolipoyllysine-residue succinyltransferase component of 2-oxoglutarate dehydrogena | Q9D2G2 | Dlst | 49 kDa | 6 C | 0.548 | 0.19696 |
| Early endosome antigen 1 | Q8BL66 | Eea1 | 161 kDa | 20 C | 0.547 | 0.22548 |
| Phosphoenolpyruvate carboxykinase, cytosolic [GTP] | Q9Z2V4 | Pck1 | 69 kDa | 13 C | 0.545 | 0.65913 |
| Eukaryotic translation initiation factor 2 subunit 3, X-linked | Q9Z0N1 | Eif2s3x | 51 kDa | 10 C | 0.545 | 0.65913 |
| UDP-glucuronosyltransferase 3A2 | Q8JZZ0 | Ugt3a2 | 60 kDa | 3 C | 0.545 | 0.65913 |
| Dimethylaniline monooxygenase [N-oxide-forming] 1 | P50285 | Fmo1 | 60 kDa | 10 C | 0.545 | 0.65913 |
| Bifunctional coenzyme A synthase | Q9DBL7 | Coasy | 62 kDa | 4 C | 0.545 | 0.65913 |
| 60S ribosomal protein L14 | Q9CR57 | Rpl14 | 24 kDa | 2 C | 0.545 | 0.65913 |
| Vesicle-associated membrane protein-associated protein A | Q9WV55 | Vapa | 28 kDa | 4 C | 0.545 | 0.189 |
| Cysteine sulfinic acid decarboxylase | Q9DBE0 | Csad | 55 kDa | 11 C | 0.545 | 0.39821 |

| | | | | | | |
|---|---|---|---|---|---|---|
| Prolyl endopeptidase | Q9QUR6 | Prep | 81 kDa | 17 C | 0.545 | 0.65913 |
| Methionine aminopeptidase 2 | O08663 | Metap2 | 53 kDa | 15 C | 0.545 | 0.65913 |
| Isoform 3 of Programmed cell death 6-interacting protein | Q9WU78-3 | Pdcd6ip | 97 kDa | 10 C | 0.545 | 0.65913 |
| Xaa-Pro dipeptidase | Q11136 | Pepd | 55 kDa | 17 C | 0.543 | 0.05136 |
| Enoyl-CoA hydratase domain-containing protein 3, mitochondrial | Q9D7J9 | Echdc3 | 32 kDa | 5 C | 0.537 | 0.51852 |
| Ribose-phosphate pyrophosphokinase 1 | Q9D7G0 | Prps1 | 35 kDa | 9 C | 0.537 | 0.58428 |
| Dynein light chain Tctex-type 3 | P56387 | Dynlt3 | 13 kDa | 6 C | 0.537 | 0.58428 |
| Acyl-coenzyme A synthetase ACSM1, mitochondrial | Q91VA0 | Acsm1 | 65 kDa | 14 C | 0.533 | 0.09126 |
| Ig lambda-1 chain C region | P01843 | | 12 kDa | 3 C | 0.533 | 0.33046 |
| Indolethylamine N-methyltransferase | P40936 | Inmt | 29 kDa | 11 C | 0.530 | 0.58899 |
| Thioredoxin | P10639 | Txn | 12 kDa | 6 C | 0.526 | 0.09445 |
| Uromodulin | Q91X17 | Umod | 71 kDa | 48 C | 0.526 | 0.21807 |
| NADP-dependent malic enzyme | P06801 | Me1 | 64 kDa | 11 C | 0.526 | 0.56176 |
| Bifunctional glutamate/proline--tRNA ligase | Q8CGC7 | Eprs | 170 kDa | 31 C | 0.525 | 0.23073 |
| Density-regulated protein | Q9CQJ6 | Denr | 22 kDa | 7 C | 0.525 | 0.23073 |
| Ankyrin repeat and SAM domain-containing protein 4B | Q8K3X6 | Anks4b | 48 kDa | 6 C | 0.525 | 0.23073 |
| Heat shock protein 75 kDa, mitochondrial | Q9CQN1 | Trap1 | 80 kDa | 20 C | 0.524 | 0.66874 |
| Nucleolin | P09405 | Ncl | 77 kDa | 1 C | 0.524 | 0.06677 |
| Cytosolic purine 5'-nucleotidase | Q3V1L4 | Nt5c2 | 65 kDa | 8 C | 0.524 | 0.13178 |
| Ribosome-binding protein 1 | Q99PL5 | Rrbp1 | 173 kDa | 8 C | 0.519 | 0.09211 |
| Glutamate dehydrogenase 1, mitochondrial | P26443 | Glud1 | 61 kDa | 6 C | 0.519 | 0.34545 |
| Secernin-3 | Q3TMH2 | Scrn3 | 48 kDa | 7 C | 0.517 | 0.29165 |
| Inosine-5'-monophosphate dehydrogenase 2 | P24547 | Impdh2 | 56 kDa | 7 C | 0.517 | 0.29165 |
| Caspase-3 | P70677 | Casp3 | 31 kDa | 8 C | 0.517 | 0.29165 |
| Lupus La protein homolog | P32067 | Ssb | 48 kDa | 3 C | 0.517 | 0.29165 |
| Mitochondrial antiviral-signaling protein | Q8VCF0 | Mavs | 53 kDa | 8 C | 0.517 | 0.29165 |
| Delta-1-pyrroline-5-carboxylate dehydrogenase, mitochondrial | Q8CHT0 | Aldh4a1 | 62 kDa | 8 C | 0.513 | 0.50632 |
| Calnexin | P35564 | Canx | 67 kDa | 7 C | 0.512 | 0.39532 |
| Methionine-R-sulfoxide reductase B2, mitochondrial | Q78J03 | Msrb2 | 19 kDa | 9 C | 0.512 | 0.50353 |
| Dihydrolipoyl dehydrogenase, mitochondrial | O08749 | Dld | 54 kDa | 9 C | 0.511 | 0.05143 |
| Methylosome protein 50 | Q99J09 | Wdr77 | 37 kDa | 12 C | 0.510 | 0.36006 |
| Collagen alpha-1(XVIII) chain | P39061 | Col18a1 | 182 kDa | 8 C | 0.508 | 0.46937 |
| Peroxisomal carnitine O-octanoyltransferase | Q9DC50 | Crot | 70 kDa | 16 C | 0.508 | 0.16508 |
| Alpha-aminoadipic semialdehyde synthase, mitochondrial | Q99K67 | Aass | 103 kDa | 13 C | 0.500 | 0.62754 |
| Kynurenine--oxoglutarate transaminase 1 | Q8BTY1 | Kyat1 | 48 kDa | 7 C | 0.500 | 0.05935 |
| Acyl-CoA dehydrogenase family member 10 | Q8K370 | Acad10 | 119 kDa | 16 C | 0.500 | 0.06017 |
| Regulator of microtubule dynamics protein 3 | Q3UJU9 | Rmdn3 | 52 kDa | 6 C | 0.500 | 0.14815 |
| Myotrophin | P62774 | Mtpn | 13 kDa | 3 C | 0.500 | 0.1583 |
| Sodium/potassium-transporting ATPase subunit beta-1 | P14094 | Atp1b1 | 35 kDa | 7 C | 0.500 | 0.19225 |
| Glycine N-acyltransferase-like protein Keg1 | Q9DCY0 | Keg1 | 34 kDa | 8 C | 0.500 | 0.51852 |
| Annexin A5 | P48036 | Anxa5 | 36 kDa | 1 C | 0.492 | 0.3453 |
| Nucleoside diphosphate-linked moiety X motif 19 | P11930 | Nudt19 | 40 kDa | 9 C | 0.490 | 0.62887 |
| Medium-chain specific acyl-CoA dehydrogenase, mitochondrial | P45952 | Acadm | 46 kDa | 8 C | 0.485 | 0.05478 |
| 3-ketoacyl-CoA thiolase A, peroxisomal | Q921H8 | Acaa1a | 44 kDa | 8 C | 0.476 | 0.15698 |
| Dihydropteridine reductase | Q8BVI4 | Qdpr | 26 kDa | 4 C | 0.474 | 0.06677 |
| Delta(3,5)-Delta(2,4)-dienoyl-CoA isomerase, mitochondrial | O35459 | Ech1 | 36 kDa | 6 C | 0.469 | 0.23505 |
| 2,4-dienoyl-CoA reductase, mitochondrial | Q9CQ62 | Decr1 | 36 kDa | 5 C | 0.462 | 0.13209 |
| ATP synthase subunit d, mitochondrial | Q9DCX2 | Atp5h | 19 kDa | 1 C | 0.455 | 0.18407 |
| Phosphoglucomutase-1 | Q9D0F9 | Pgm1 | 61 kDa | 10 C | 0.455 | 0.41687 |
| Cytoplasmic aconitate hydratase | P28271 | Aco1 | 98 kDa | 11 C | 0.444 | 0.22632 |
| Beta-ureidopropionase | Q8VC97 | Upb1 | 44 kDa | 10 C | 0.444 | 0.23275 |
| FAS-associated death domain protein | Q61160 | Fadd | 23 kDa | 3 C | 0.443 | 0.11288 |
| Lipoamide acyltransferase component of branched-chain alpha-keto acid dehydrogenase c | P53395 | Dbt | 53 kDa | 6 C | 0.443 | 0.27892 |
| F-actin-capping protein subunit alpha-2 | P47754 | Capza2 | 33 kDa | 3 C | 0.443 | 0.27892 |
| Beta-mannosidase | Q8K2I4 | Manba | 101 kDa | 12 C | 0.443 | 0.27892 |
| Cystathionine beta-synthase | Q91WT9 (+1) | Cbs | 62 kDa | 13 C | 0.437 | 0.05583 |
| cAMP-dependent protein kinase type I-alpha regulatory subunit | Q9DBC7 | Prkar1a | 43 kDa | 5 C | 0.431 | 0.41355 |
| Ras-related protein Rab-11B | P46638 | Rab11b | 24 kDa | 2 C | 0.431 | 0.41355 |
| Golgi reassembly-stacking protein 2 | Q99JX3 | Gorasp2 | 47 kDa | 4 C | 0.429 | 0.11612 |
| Annexin A6 | P14824 | Anxa6 | 76 kDa | 8 C | 0.425 | 0.21103 |

| | | | | | | |
|---|---|---|---|---|---|---|
| V-type proton ATPase catalytic subunit A | P50516 | Atp6v1a | 68 kDa | 6 C | 0.421 | 0.29765 |
| F-BAR domain only protein 2 | Q3UQN2 | Fcho2 | 89 kDa | 9 C | 0.420 | 0.09737 |
| Polyadenylate-binding protein-interacting protein 1 | Q8VE62 | Paip1 | 46 kDa | 6 C | 0.420 | 0.09737 |
| Isoform 5 of Kinectin | Q61595-5 | Ktn1 | 146 kDa | 10 C | 0.412 | 0.45725 |
| ADP-ribosylation factor 1 | P84078 | Arf1 | 21 kDa | 1 C | 0.412 | 0.45725 |
| Cold shock domain-containing protein E1 | Q91W50 | Csde1 | 89 kDa | 15 C | 0.407 | 0.13969 |
| E3 SUMO-protein ligase RanBP2 | Q9ERU9 | Ranbp2 | 341 kDa | 67 C | 0.387 | 0.36848 |
| Ig lambda-1 chain V region | P01723 (+1) | | 12 kDa | 2 C | 0.387 | 0.36848 |
| MIP18 family protein FAM96A | Q9DCL2 | Fam96a | 18 kDa | 5 C | 0.387 | 0.36848 |
| V-type proton ATPase subunit G 1 | Q9CR51 | Atp6v1g1 | 14 kDa | 2 C | 0.387 | 0.36848 |
| Endonuclease G, mitochondrial | O08600 | Endog | 32 kDa | 2 C | 0.387 | 0.36848 |
| CAP-Gly domain-containing linker protein 1 | Q922J3 | Clip1 | 156 kDa | 11 C | 0.387 | 0.36848 |
| Low molecular weight phosphotyrosine protein phosphatase | Q9D358 | Acp1 | 18 kDa | 8 C | 0.387 | 0.36848 |
| Synaptic functional regulator FMR1 | P35922 (+5) | Fmr1 | 69 kDa | 5 C | 0.387 | 0.36848 |
| Protein farnesyltransferase/geranylgeranyltransferase type-1 subunit alpha | Q61239 | Fnta | 44 kDa | 3 C | 0.387 | 0.36848 |
| Mitochondrial-processing peptidase subunit alpha | Q9DC61 | Pmpca | 58 kDa | 9 C | 0.383 | 0.30298 |
| Complement component C8 alpha chain | Q8K182 | C8a | 66 kDa | 29 C | 0.375 | 0.5461 |
| Sideroflexin-1 | Q99JR1 | Sfxn1 | 36 kDa | 5 C | 0.375 | 0.5461 |
| UPF0568 protein C14orf166 homolog | Q9CQE8 | | 28 kDa | 2 C | 0.375 | 0.5461 |
| N(G),N(G)-dimethylarginine dimethylaminohydrolase 1 | Q9CWS0 | Ddah1 | 31 kDa | 7 C | 0.373 | 0.22037 |
| Cytochrome c oxidase assembly factor 7 | Q921H9 | Coa7 | 26 kDa | 13 C | 0.367 | 0.21347 |
| Isoform 3 of Protein arginine N-methyltransferase 1 | Q9JIF0-3 | Prmt1 | 40 kDa | 11 C | 0.350 | 0.11638 |
| Cathepsin B | P10605 | Ctsb | 37 kDa | 16 C | 0.344 | 0.25812 |
| Cytochrome c oxidase subunit 5B, mitochondrial | P19536 | Cox5b | 14 kDa | 5 C | 0.344 | 0.4342 |
| Ribosome-recycling factor, mitochondrial | Q9D6S7 | Mrrf | 29 kDa | 2 C | 0.342 | 0.06341 |
| Ferritin light chain 1 | P29391 | Ftl1 | 21 kDa | 1 C | 0.333 | 0.09998 |
| Major urinary protein 1 | P11588 | Mup1 | 21 kDa | 5 C | 0.333 | 0.11612 |
| Fatty acid-binding protein, liver | P12710 | Fabp1 | 14 kDa | 1 C | 0.333 | 0.189 |
| Aldehyde oxidase 3 | G3X982 | Aox3 | 147 kDa | 38 C | 0.333 | 0.34515 |
| Putative adenosylhomocysteinase 3 | Q68FL4 (+1) | Ahcyl2 | 67 kDa | 18 c | 0.315 | 0.37498 |
| Alanine aminotransferase 1 | Q8QZR5 | Gpt | 55 kDa | 14 C | 0.314 | 0.08906 |
| Alpha-2-HS-glycoprotein | P29699 | Ahsg | 37 kDa | 14 C | 0.314 | 0.21459 |
| Nuclear mitotic apparatus protein 1 | E9Q7G0 | Numa1 | 236 kDa | 19 C | 0.310 | 0.21275 |
| GrpE protein homolog 1, mitochondrial | Q99LP6 | Grpel1 | 24 kDa | 4 C | 0.300 | 0.10588 |
| Lysozyme C-2 | P08905 | Lyz2 | 17 kDa | 8 C | 0.300 | 0.05724 |
| BH3-interacting domain death agonist | P70444 | Bid | 22 kDa | 2 C | 0.300 | 0.10588 |
| Isoform 2 of Cysteine--tRNA ligase, cytoplasmic | Q9ER72-2 | Cars | 86 kDa | 11 C | 0.300 | 0.10588 |
| Glutamyl aminopeptidase | P16406 | Enpep | 108 kDa | 9 C | 0.300 | 0.29805 |
| Peroxisomal trans-2-enoyl-CoA reductase | Q99MZ7 | Pecr | 32 kDa | 5 C | 0.294 | 0.16492 |
| Pterin-4-alpha-carbinolamine dehydratase | P61458 | Pcbd1 | 12 kDa | 1 C | 0.293 | 0.23987 |
| 60S ribosomal protein L30 | P62889 | Rpl30 | 13 kDa | 3 C | 0.293 | 0.23987 |
| Voltage-dependent anion-selective channel protein 2 | Q60930 | Vdac2 | 32 kDa | 11 C | 0.293 | 0.34704 |
| Protein canopy homolog 2 | Q9QXT0 | Cnpy2 | 21 kDa | 6 C | 0.293 | 0.23987 |
| Citrate synthase, mitochondrial | Q9CZU6 | Cs | 52 kDa | 5 C | 0.293 | 0.34704 |
| L-xylulose reductase | Q91X52 | Dcxr | 26 kDa | 4 C | 0.293 | 0.34704 |
| 6-phosphogluconate dehydrogenase, decarboxylating | Q9DCD0 | Pgd | 53 kDa | 9 C | 0.286 | 0.49521 |
| Maleylacetoacetate isomerase | Q9WVL0 | Gstz1 | 24 kDa | 3 C | 0.286 | 0.49521 |
| Calcium-binding mitochondrial carrier protein Aralar1 | Q8BH59 | Slc25a12 | 75 kDa | 7 C | 0.286 | 0.49521 |
| Trimethyllysine dioxygenase, mitochondrial | Q91ZE0 | Tmlhe | 50 kDa | 11 C | 0.283 | 0.119 |
| V-type proton ATPase subunit B, brain isoform | P62814 | Atp6v1b2 | 57 kDa | 6 C | 0.283 | 0.23999 |
| Gamma-glutamyltranspeptidase 1 | Q60928 | Ggt1 | 62 kDa | 6 C | 0.261 | 0.19272 |
| Quinone oxidoreductase-like protein 2 | Q3UNZ8 | | 38 kDa | 9 C | 0.259 | 0.32034 |
| Acylamino-acid-releasing enzyme | Q8R146 (+1) | Apeh | 82 kDa | 16 C | 0.259 | 0.41206 |
| G-rich sequence factor 1 | Q8C5Q4 | Grsf1 | 53 kDa | 9 C | 0.240 | 0.15819 |
| O-phosphoseryl-tRNA(Sec) selenium transferase | Q6P6M7 | Sepsecs | 55 kDa | 13 C | 0.239 | 0.48469 |
| 5-oxoprolinase | Q8K010 | Oplah | 138 kDa | 25 C | 0.238 | 0.19127 |
| Eukaryotic peptide chain release factor GTP-binding subunit ERF3A | Q8R050 (+1) | Gspt1 | 69 kDa | 14 C | 0.235 | 0.2229 |
| Sorting nexin-12 | O70493 | Snx12 | 19 kDa | 3 C | 0.233 | 0.12613 |
| Cytochrome c oxidase subunit 2 | P00405 | Mtco2 | 26 kDa | 3 C | 0.231 | 0.46703 |
| Acylcarnitine hydrolase | Q91WG0 | Ces2c | 62 kDa | 5 C | 0.221 | 0.19052 |

| | | | | | | |
|---|---|---|---|---|---|---|
| Glycine dehydrogenase (decarboxylating), mitochondrial | Q91W43 | Gldc | 113 kDa | 24 C | 0.220 | 0.12817 |
| Isoform HK1 of Hexokinase-1 | P17710-3 | Hk1 | 102 kDa | 21 C | 0.210 | 0.12628 |
| Carboxylesterase 1C | P23953 | Ces1c | 61 kDa | 5 C | 0.210 | 0.15057 |
| Aminopeptidase N | P97449 | Anpep | 110 kDa | 8 C | 0.210 | 0.23391 |
| Cingulin-like protein 1 | Q6AW69 (+2 | Cgnl1 | 148 kDa | 13 C | 0.200 | 0.06976 |
| Isoform 3 of ATP-dependent (S)-NAD(P)H-hydrate dehydratase | Q9CZ42-3 | Naxd | 35 kDa | 8 C | 0.197 | 0.18188 |
| Acyl-coenzyme A synthetase ACSM3, mitochondrial | Q3UNX5 (+1) | Acsm3 | 66 kDa | 10 C | 0.195 | 0.15049 |
| Integrin beta-1 | P09055 (+1) | Itgb1 | 88 kDa | 57 C | 0.194 | 0.0504 |
| Cytochrome P450 4B1 | Q64462 | Cyp4b1 | 59 kDa | 9 C | 0.191 | 0.45851 |
| Isoform Mt-VDAC1 of Voltage-dependent anion-selective channel protein 1 | Q60932-2 | Vdac1 | 31 kDa | 2 C | 0.191 | 0.09004 |
| Glucose-6-phosphate isomerase | P06745 | Gpi | 63 kDa | 4 C | 0.174 | 0.18287 |
| Ornithine aminotransferase, mitochondrial | P29758 | Oat | 48 kDa | 7 C | 0.169 | 0.22112 |
| Inorganic pyrophosphatase | Q9D819 | Ppa1 | 33 kDa | 8 C | 0.136 | 0.3739 |
| GTP-binding nuclear protein Ran | P62827 | Ran | 24 kDa | 3 C | 0.136 | 0.3739 |
| C-1-tetrahydrofolate synthase, cytoplasmic | Q922D8 | Mthfd1 | 101 kDa | 12 C | 0.136 | 0.3739 |
| Carbamoyl-phosphate synthase [ammonia], mitochondrial | Q8C196 | Cps1 | 165 kDa | 21 C | 0.136 | 0.3739 |
| Thiosulfate sulfurtransferase | P52196 | Tst | 33 kDa | 4 C | 0.136 | 0.3739 |
| Enoyl-CoA delta isomerase 2, mitochondrial | Q9WUR2 | Eci2 | 43 kDa | 7 C | 0.136 | 0.3739 |
| Alkaline phosphatase, tissue-nonspecific isozyme | P09242 | Alpl | 58 kDa | 6 C | 0.136 | 0.3739 |
| DNA-directed RNA polymerases I, II, and III subunit RPABC3 | Q923G2 | Polr2h | 17 kDa | 1 C | 0.136 | 0.3739 |
| Sulfide:quinone oxidoreductase, mitochondrial | Q9R112 | Sqor | 50 kDa | 8 C | 0.136 | 0.3739 |
| Complement component 1 Q subcomponent-binding protein, mitochondrial | O35658 | C1qbp | 31 kDa | 6 C | 0.136 | 0.3739 |
| ATP-binding cassette sub-family D member 3 | P55096 | Abcd3 | 75 kDa | 9 C | 0.136 | 0.3739 |
| Lamin-B1 | P14733 | Lmnb1 | 67 kDa | 4 C | 0.136 | 0.3739 |
| Annexin A11 | P97384 | Anxa11 | 54 kDa | 6 C | 0.136 | 0.3739 |
| 60S acidic ribosomal protein P0 | P14869 | Rplp0 | 34 kDa | 3 C | 0.136 | 0.3739 |
| Protein FAM151A | Q8QZW3 | Fam151a | 67 kDa | 6 C | 0.136 | 0.3739 |
| NADH dehydrogenase [ubiquinone] iron-sulfur protein 7, mitochondrial | Q9DC70 | Ndufs7 | 25 kDa | 5 C | 0.136 | 0.3739 |
| Acyl-CoA dehydrogenase family member 11 | Q80XL6 | Acad11 | 87 kDa | 13 C | 0.136 | 0.3739 |
| 60S ribosomal protein L7a | P12970 | Rpl7a | 30 kDa | 3 C | 0.136 | 0.3739 |
| ATP-dependent RNA helicase DDX3X | Q62167 | Ddx3x | 73 kDa | 7 C | 0.136 | 0.3739 |
| Beta-2-microglobulin | P01887 | B2m | 14 kDa | 2 C | 0.136 | 0.3739 |
| Galectin-3-binding protein | Q07797 | Lgals3bp | 64 kDa | 16 C | 0.136 | 0.3739 |
| Prefoldin subunit 2 | O70591 | Pfdn2 | 17 kDa | 1 C | 0.136 | 0.3739 |
| NADH dehydrogenase [ubiquinone] 1 alpha subcomplex subunit 9, mitochondrial | Q9DC69 | Ndufa9 | 43 kDa | 2 C | 0.136 | 0.3739 |
| 40S ribosomal protein S8 | P62242 | Rps8 | 24 kDa | 5 C | 0.136 | 0.3739 |
| Catechol O-methyltransferase | O88587 (+1) | Comt | 29 kDa | 4 C | 0.136 | 0.3739 |
| NADH dehydrogenase [ubiquinone] iron-sulfur protein 3, mitochondrial | Q9DCT2 | Ndufs3 | 30 kDa | 3 C | 0.132 | 0.1592 |
| Electron transfer flavoprotein-ubiquinone oxidoreductase, mitochondrial | Q921G7 | Etfdh | 68 kDa | 15 C | 0.132 | 0.20252 |
| Argininosuccinate lyase | Q91YI0 | Asl | 52 kDa | 13 C | 0.132 | 0.27412 |
| Heat shock protein 105 kDa | Q61699 (+1) | Hsph1 | 96 kDa | 17 C | 0.120 | 0.09838 |
| Dipeptidase 1 | P31428 | Dpep1 | 46 kDa | 8 C | 0.119 | 0.16681 |
| WD repeat-containing protein 1 | O88342 | Wdr1 | 66 kDa | 12 C | 0.097 | 0.16419 |
| Sushi domain-containing protein 2 | Q9DBX3 | Susd2 | 91 kDa | 28 C | 0.097 | 0.16419 |
| Eukaryotic translation initiation factor 1A, X-chromosomal | Q8BMJ3 | Eif1ax | 16 kDa | 2 C | 0.097 | 0.16419 |
| D-dopachrome decarboxylase | O35215 | Ddt | 13 kDa | 2 C | 0.097 | 0.16419 |
| Carboxypeptidase Q | Q9WVJ3 (+1) | Cpq | 52 kDa | 1 C | 0.097 | 0.16419 |
| Coronin-1C | Q9WUM4 | Coro1c | 53 kDa | 12 C | 0.097 | 0.16419 |
| Apolipoprotein E | P08226 | Apoe | 36 kDa | 1 C | 0.097 | 0.16419 |
| Adaptin ear-binding coat-associated protein 2 | Q9D1J1 | Necap2 | 29 kDa | 2 C | 0.097 | 0.16419 |
| Transmembrane emp24 domain-containing protein 10 | Q9D1D4 | Tmed10 | 25 kDa | 3 C | 0.097 | 0.16419 |
| UDP-glucose 6-dehydrogenase | O70475 | Ugdh | 55 kDa | 12 C | 0.094 | 0.3739 |
| T-complex protein 1 subunit theta | P42932 | Cct8 | 60 kDa | 10 C | 0.094 | 0.3739 |
| Ubiquitin-like modifier-activating enzyme 1 | Q02053 | Uba1 | 118 kDa | 21 C | 0.094 | 0.3739 |
| NADH dehydrogenase [ubiquinone] iron-sulfur protein 2, mitochondrial | Q91WD5 | Ndufs2 | 53 kDa | 7 C | 0.094 | 0.3739 |
| Probable 2-oxoglutarate dehydrogenase E1 component DHKTD1, mitochondrial | A2ATU0 | Dhtkd1 | 103 kDa | 13 C | 0.094 | 0.3739 |
| ER membrane protein complex subunit 8 | O70378 | Emc8 | 23 kDa | 8 C | 0.094 | 0.3739 |
| NADH dehydrogenase [ubiquinone] iron-sulfur protein 8, mitochondrial | Q8K3J1 | Ndufs8 | 24 kDa | 8 C | 0.094 | 0.3739 |
| Aquaporin-1 | Q02013 | Aqp1 | 29 kDa | 4 C | 0.094 | 0.3739 |
| 3-mercaptopyruvate sulfurtransferase | Q99J99 | Mpst | 33 kDa | 4 C | 0.094 | 0.3739 |

| | | | | | | |
|---|---|---|---|---|---|---|
| NADH dehydrogenase [ubiquinone] 1 alpha subcomplex subunit 8 | Q9DCJ5 | Ndufa8 | 20 kDa | 8 C | 0.073 | 0.11612 |
| Lysosome-associated membrane glycoprotein 1 | P11438 | Lamp1 | 44 kDa | 8 C | 0.073 | 0.11612 |
| Membrane-associated progesterone receptor component 1 | O55022 | Pgrmc1 | 22 kDa | 2 C | 0.073 | 0.11612 |
| Estradiol 17-beta-dehydrogenase 8 | P50171 (+1) | Hsd17b8 | 27 kDa | 4 C | 0.073 | 0.21347 |
| Ornithine carbamoyltransferase, mitochondrial | P11725 | Otc | 40 kDa | 2 C | 0.073 | 0.21347 |
| Golgin subfamily A member 4 | Q91VW5 | Golga4 | 258 kDa | 25 C | 0.071 | 0.3739 |
| ATP synthase F(0) complex subunit B1, mitochondrial | Q9CQQ7 | Atp5f1 | 29 kDa | 2 C | 0.071 | 0.3739 |
| Ras-related protein Rab-5C | P35278 | Rab5c | 23 kDa | 4 C | 0.071 | 0.3739 |
| Prohibitin | P67778 | Phb | 30 kDa | 1 C | 0.071 | 0.3739 |
| Isoform 2 of Cadherin-related family member 5 | Q8VHF2-2 | Cdhr5 | 73 kDa | 7 C | 0.071 | 0.3739 |
| Catenin beta-1 | Q02248 | Ctnnb1 | 85 kDa | 11 C | 0.060 | 0.07852 |
| Cytochrome b5 type B | Q9CQX2 | Cyb5b | 16 kDa | 1 C | 0.060 | 0.07852 |
| Pyridoxal phosphate homeostasis protein | Q9Z2Y8 | Prosc | 30 kDa | 3 C | 0.049 | 0.1611 |
| Beta-glucuronidase | P12265 | Gusb | 74 kDa | 8 C | 0.049 | 0.1611 |
| Ceruloplasmin | Q61147 | Cp | 121 kDa | 14 C | 0.049 | 0.26852 |
| Calcium-binding mitochondrial carrier protein Aralar2 | Q9QXX4 | Slc25a13 | 74 kDa | 7 C | 0.048 | 0.3739 |
| MICOS complex subunit Mic60 | Q8CAQ8 (+3) | Immt | 84 kDa | 7 C | 0.048 | 0.3739 |
| UDP-glucuronosyltransferase 1-7C | Q6ZQM8 | Ugt1a7c | 60 kDa | 13 C | 0.048 | 0.3739 |
| Arginase-1 | Q61176 | Arg1 | 35 kDa | 3 C | 0.042 | 0.12465 |
| D-amino-acid oxidase | P18894 | Dao | 39 kDa | 6 C | 0.042 | 0.28437 |
| Phenazine biosynthesis-like domain-containing protein 1 | Q9DCG6 | Pbld1 | 32 kDa | 3 C | 0.030 | 0.13241 |
| S-adenosylhomocysteine hydrolase-like protein 1 | Q80SW1 | Ahcyl1 | 59 kDa | 19 C | 0.027 | 0.17807 |
| Regucalcin | Q64374 | Rgn | 33 kDa | 9 C | 0.025 | 0.11612 |
| Mitochondrial fission 1 protein | Q9CQ92 | Fis1 | 17 kDa | 1 C | 0.464 | 0.01064 |
| 2-hydroxyacyl-CoA lyase 1 | Q9QXE0 | Hacl1 | 64 kDa | 17 C | 0.435 | 0.00225 |
| Macrophage migration inhibitory factor | P34884 | Mif | 13 kDa | 3 C | 0.350 | 0.01232 |
| Protein phosphatase 1 regulatory subunit 12A | Q9DBR7 | Ppp1r12a | 115 kDa | 8 C | 0.313 | 0.00793 |
| Clusterin | Q06890 | Clu | 52 kDa | 11 C | 0.300 | 0.0249 |
| Isoform 2 of ELKS/Rab6-interacting/CAST family member 1 | Q99MI1-2 | Erc1 | 112 kDa | 4 C | 0.240 | 0.04761 |
| Golgi apparatus protein 1 | Q61543 | Glg1 | 134 kDa | 68 C | 0.233 | 0.02412 |
| Short-chain specific acyl-CoA dehydrogenase, mitochondrial | Q07417 | Acads | 45 kDa | 5 C | 0.231 | 0.01301 |
| Mitochondrial proton/calcium exchanger protein | Q9Z2I0 | Letm1 | 83 kDa | 12 C | 0.221 | 0.02494 |
| Transforming protein RhoA | Q9QUI0 | Rhoa | 22 kDa | 6 C | 0.100 | 0.0261 |
