## Supplemental Table 19 for "Dietary restriction transforms the protein sulfhydrome in a tissue-specific and cystathionine γ-lyase-dependent manner"

| Supplemental Table 19: Sulfhydrated Protein Pathway Enrichment in CGL KO Kidney | | | |
|---|---|---|---|
| KEGG Pathway (DR) | p-val (adj) | -LOG10(p-val adj) | Involved Gene |
| No KEGG | | | |
| KEGG Pathway (Unchanged) | p-val (adj) | -LOG10(p-val adj) | Involved Gene |
| Tryptophan metabolism | 2.5E-09 | 8.60 | HAAO,INMT,GCDH,OGDH,ACAT2,ECHS1,HADHA,ACMSD,ALDH9A1,CAT,HADH,ALDH2,ACAT1,KYAT1,KYAT3,ALDH7A1,AADAT |
| Alanine, aspartate and glutamate metabolism | 0.00634 | 2.20 | ASPA,GLUD1,NIT2,GLUL,ALDH4A1,GOT2,ALDH5A1,ABAT,ASS1 |
| Butanoate metabolism | 7.72E-10 | 9.11 | OXCT1,ACAT2,ECHS1,HADHA,HADH,BDH2,HMGCL,ACSM2,ACAT1,ACSM1,ALDH5A1,BDH1,ABAT,HMGCS1 |
| Cysteine and methionine metabolism | 0.0000069 | 5.16 | APIP,MDH2,MDH1,AHCY,GSS,GCLM,AHCYL2,LDHB,BCAT2,GOT2,GCLC,MAT2B,MTAP,BHMT |
| Biosynthesis of amino acids | 0.0000297 | 4.53 | ALDOC,PAH,SHMT1,ACO2,ACY1,TPI1,GLUL,ACO1,IDH2,BCAT2,PRPS1,GOT2,IDH3A,PKM,MAT2B,GAPDH,ASS1 |
| Carbon metabolism | 1.01E-19 | 19.00 | DLAT,DLST,SDHB,ALDOC,MDH2,MDH1,OGDH,SHMT1,DLD,ALDH6A1,FBP2,SDHA,PDHB,GLUD1,ESD,SUCLA2,ACO2,TPI1,ACAT2,MUT,ECHS1,HADHA,CAT,HAO2,ACO1,IDH2,PDHA1,PRPS1,GOT2,ACAT1,IDH3A,PKM,ME1,TKFC,PGP,SUCLG1,GAPDH,SUCLG2,FBP1 |
| Lysine degradation | 0.0000137 | 4.86 | GCDH,DLST,PIPOX,OGDH,ACAT2,ECHS1,HADHA,ALDH9A1,HADH,ALDH2,AASS,ACAT1,ALDH7A1,AADAT,TMLHE |
| Thyroid hormone synthesis | 0.0352 | 1.45 | GPX3,HSP90B1,CANX,PDIA4,ATP1B1,HSPA5,LRP2,ALB,GSR,ATP1A1,TTR,GPX1 |
| Fatty acid degradation | 0.0000122 | 4.91 | GCDH,ACAT2,ECI1,ECHS1,HADHA,ACADL,ALDH9A1,HADH,ACOX3,ALDH2,ACAT1,ACAA2,ALDH7A1,ADH1 |
| beta-Alanine metabolism | 0.000371 | 3.43 | ALDH6A1,DPYS,CNDP2,ECHS1,HADHA,ALDH9A1,ALDH2,DPYD,ALDH7A1,ABAT |
| Fructose and mannose metabolism | 0.00498 | 2.30 | AKR1B1,ALDOC,FBP2,TPI1,SORD,KHK,TKFC,GMDS,FBP1 |
| Fatty acid metabolism | 0.00521 | 2.28 | ACAT2,ECHS1,HADHA,ACADL,PECR,HADH,PPT1,ACOX3,ACAT1,ACAA2,MCAT |
| Citrate cycle (TCA cycle) | 1.02E-13 | 12.99 | DLAT,DLST,SDHB,MDH2,MDH1,OGDH,DLD,SDHA,PDHB,SUCLA2,ACO2,PCK1,ACO1,IDH2,PDHA1,IDH3A,SUCLG1,SUCLG2 |
| Arginine and proline metabolism | 0.0312 | 1.51 | CKMT1,CKB,CNDP2,HOGA1,ALDH9A1,ALDH4A1,ALDH2,GOT2,LAP3,ALDH7A1 |
| Metabolic pathways | 1.85E-28 | 27.73 | COX5A,DLAT,CKMT1,HAAO,CKB,AKR1B1,COASY,PAFAH1B2,GCDH,DLST,SARDH,SDHB,APIP,NDUFA2,CHDH,ALDOC,PIPOX,PNPO,ISYNA1,MDH2,ATP6V1E1,PAH,MDH1,OGDH,SHMT1,DLD,ASPA,NME2,ALDH6A1,FBP2,AUH,SDHA,MCCC2,PDHB,GLUD1,EPHX2,SUCLA2,CMBL,AMACR,DPYS,ACO2,CPOX,HGD,CSAD,ACY1,TPI1,ACAT2,MUT,NDUFV2,HSD17B4,CNDP2,ALDH1A7,NT5C2,GK,HOGA1,HSD17B10,ECHS1,HPRT1,HADHA,ATP5C1,NDUFS1,ACADL,NDUFA10,SCLY,ACMSD,GLUL,EPRS,BPNT1,SEPHS1,ALDH9A1,PRDX6,AK1,UAP1L1,SORD,IVD,PCK1,IMPA1,AHCY,GSS,NFS1,HAO2,HADH,GCLM,BDH2,ALAD,ACO1,SCP2,PPT1,HMGCL,AKR1A1,CMPK1,ALDH4A1,CDA,AK2,ACOX3,KHK,PAICS,HPD,ALDH2,AASS,AHCYL2,HIBADH,LDHB,IDH2,FAH,BCAT2,UQCRC2,ACSM2,PDHA1,PRPS1,GOT2,CES1F,ACAT1,PTS,IDH3A,PKM,GCLC,ME1,DPYD,ACSM1,TKFC,GALM,XYLB,ALDH5A1,ACAA2,NME1,GMDS,UQCRFS1,ADK,KYAT1,LAP3,KYAT3,BLVRB,MAT2B,DMGDH,PGP,FAHD1,BDH1,GPHN,MCAT,SEPHS2,SUOX,GNPDA1,ACOT4,SUCLG1,ALDH1A1,ALDH7A1,CES1D,AADAT,GAPDH,ABAT,SUCLG2,IMPDH2,MTAP,PPCDC,MOCS1,FBP1,PTGES3,GANAB,NME3,ADH1,BHMT,ASS1,HMGCS1 |
| Synthesis and degradation of ketone bodies | 0.0000254 | 4.60 | OXCT1,ACAT2,BDH2,HMGCL,ACAT1,BDH1,HMGCS1 |
| Bacterial invasion of epithelial cells | 0.00226 | 2.65 | CDH1,ARPC2,CDC42,ARPC5,CRK,VCL,CLTA,ARPC3,ARPC1A,CTTN,CTNNA1,CLTC,ACTG1,ARPC4 |
| Glyoxylate and dicarboxylate metabolism | 4.86E-09 | 8.31 | MDH2,MDH1,SHMT1,DLD,ACO2,ACAT2,MUT,HOGA1,GLUL,CAT,HAO2,ACO1,ACAT1,PGP |
| Selenocompound metabolism | 0.000129 | 3.89 | TXNRD3,INMT,TXNRD1,SCLY,SEPHS1,KYAT1,KYAT3,SEPHS2 |
| Valine, leucine and isoleucine degradation | 6.45E-13 | 12.19 | DLD,ALDH6A1,AUH,MCCC2,OXCT1,ACAT2,MUT,HSD17B10,ECHS1,HADHA,ALDH9A1,IVD,HADH,HMGCL,ALDH2,HIBADH,BCAT2,ACAT1,ACAA2,ALDH7A1,ABAT,HMGCS1 |
| Glutathione metabolism | 0.00244 | 2.61 | GPX3,GSTA3,GSS,GCLM,TXNDC12,IDH2,GSR,GCLC,LAP3,GSTM1,GPX1,GSTA1,GPX4 |
| Nitrogen metabolism | 0.0139 | 1.86 | GLUD1,GLUL,CA1,CA3,CA2,CA5B |
| Proteasome | 4.15E-08 | 7.38 | PSMD4,PSMB4,PSMB1,PSMA2,PSMB6,PSMA6,PSMB5,PSME1,PSMB7,PSMA7,PSMB2,PSMA1,PSMA4,PSMA3,PSMA5,PSMB3 |
| Drug metabolism - other enzymes | 0.016 | 1.80 | NME2,DPYS,HPRT1,GSTA3,CMPK1,CDA,CES1F,DPYD,NME1,CES1D,GSTM1,IMPDH2,NME3,GSTA1 |
| Proximal tubule bicarbonate reclamation | 0.00802 | 2.10 | MDH1,GLUD1,SLC25A10,ATP1B1,PCK1,CA2,ATP1A1 |
| Peroxisome | 0.0000366 | 4.44 | CROT,SOD2,PIPOX,EPHX2,AMACR,SOD1,HSD17B4,PRDX5,PECR,PHYH,CAT,SLC27A2,HAO2,SCP2,HMGCL,PRDX1,ACOX3,IDH2 |
| 2-Oxocarboxylic acid metabolism | 0.000212 | 3.67 | ACO2,ACY1,ACO1,IDH2,BCAT2,GOT2,IDH3A,AADAT |
| Propanoate metabolism | 0.00000167 | 5.78 | DLD,ALDH6A1,SUCLA2,ACAT2,MUT,ECHS1,HADHA,LDHB,ACAT1,SUCLG1,ABAT,SUCLG2 |
| Tight junction | 0.0336 | 1.47 | CDC42,ACTN1,ACTR2,PPP2CA,HSPA4,SLC9A3R1,PPP2R2A,DLG1,TUBA1B,TUBA4A,ACTR3,CTTN,MSN,RDX,NEDD4,EZR,ACTN4,ACTG1,AFDN,MYL6 |
| Pyruvate metabolism | 5.45E-12 | 11.26 | DLAT,MDH2,MDH1,DLD,PDHB,ACAT2,GLO1,HAGH,ALDH9A1,PCK1,ALDH2,LDHB,PDHA1,ACAT1,PKM,ME1,ALDH7A1,ACYP2 |
| Glycolysis / Gluconeogenesis | 0.000000497 | 6.30 | DLAT,ALDOC,DLD,FBP2,PDHB,TPI1,ALDH9A1,PCK1,AKR1A1,ALDH2,LDHB,PDHA1,PKM,GALM,ALDH7A1,GAPDH,FBP1,ADH1 |
| KEGG Pathway (AL) | p-val (adj) | -LOG10(p-val adj) | Involved Gene |
| No KEGG | | | |
